# Automatic identification of players in the flavonoid biosynthesis with application on the biomedicinal plant *Croton tiglium*

**DOI:** 10.1101/2020.06.27.175067

**Authors:** Boas Pucker, Franziska Reiher, Hanna Marie Schilbert

**Author notes:** Correspondence (H.M.S.).

## Abstract

The flavonoid biosynthesis is a well characterised model system for specialised metabolism and transcriptional regulation in plants. Flavonoids have numerous biological functions like UV protection and pollinator attraction, but also biotechnological potential. Here, we present Knowledge-based Identification of Pathway Enzymes (KIPEs) as an automatic approach for the identification of players in the flavonoid biosynthesis. KIPEs combines comprehensive sequence similarity analyses with the inspection of functionally relevant amino acid residues and domains in subjected peptide sequences. Comprehensive sequence sets of flavonoid biosynthesis enzymes and knowledge about functionally relevant amino acids were collected. As a proof of concept, KIPEs was applied to investigate the flavonoid biosynthesis of the medicinal plant *Croton tiglium* based on a transcriptome assembly. Enzyme candidates for all steps in the biosynthesis network were identified and matched to previous reports of corresponding metabolites in *Croton* species.

## 1. Introduction

Flavonoids are a group of specialised plant metabolites comprising more than 9,000 identified compounds [1] with numerous biological functions [2]. Flavonoids are derived from the aromatic amino acid phenylalanine in a branch of the phenylpropanoid pathway namely the flavonoid biosynthesis (Figure 1). Generally, flavonoids consist of two aromatic C6-rings and one heterocyclic pyran ring [3]. Products of the flavonoid biosynthesis can be assigned to different subgroups, including chalcones, flavones, flavonols, flavandiols, anthocyanins, proanthocyanidins (PA), and aurones [4]. These subclasses are characterised by different oxidation states [5]. In plants, these aglycons are often modified through the addition of various sugars leading to a huge diversity [6].

**Figure 1:**
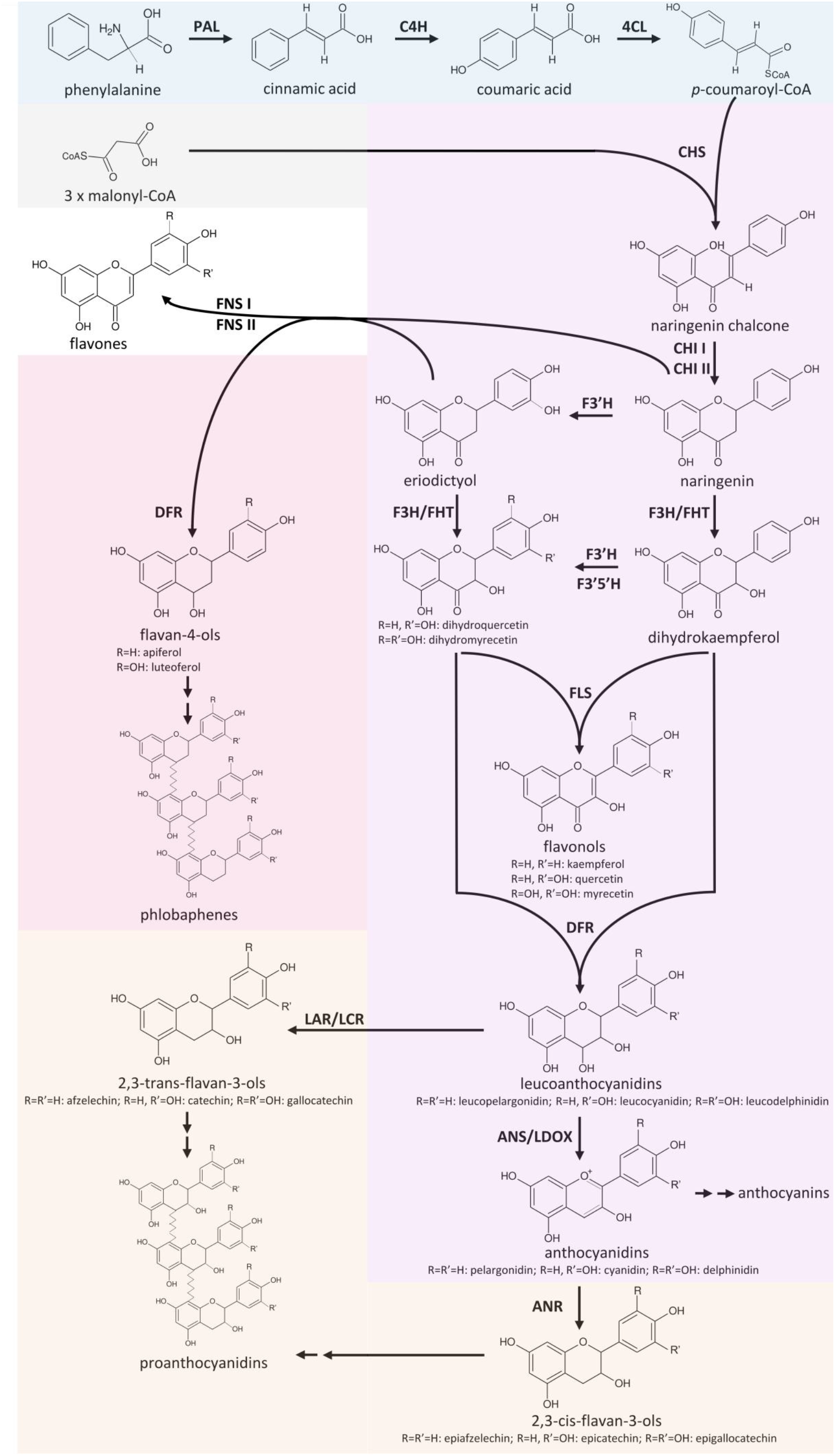
Simplified illustration of the general phenylpropanoid pathway and the core flavonoid aglycon biosynthesis network.

Flavonoids have important developmental and ecological roles in plants including the control of auxin transport [7], the attraction of pollinators [8], protection of plants against UV light [9], and defense against pathogens and herbivores [10]. Different types of flavonoids can take up these roles. Anthocyanins appear as violet, blue, orange, or red pigments in plants recruiting pollinators and seed dispersers [8]. PAs accumulate in the seed coat leading to the characteristic dark colour of seeds in many species [8]. Flavonols are stored in their glycosylated form in the vacuole of epidermal cells or on occasion in epicuticular waxes [4]. They possess several physiological functions including antimicrobial defense, scavenging of reactive oxygen species (ROS), UV protection, signaling, and colouration of flower pigmentation together with anthocyanins [9]. Consequently, the activity of different branches of the flavonoid biosynthesis needs to be adjusted in response to developmental stages and environmental conditions. While the biosynthesis of anthocyanins can be triggered by abiotic factors such as light, temperature, dryness or salts [11], PAs are formed independently of external stimuli in the course of seed development leading to a brown seed colour [11].

As the accumulation of flavonoids in fruits and vegetables [12] leads to colouration desired by customers, this pigment pathways is of biotechnological relevance. Therefore, the flavonoid biosynthesis was previously modified by genetic engineering in multiple species (as reviewed in [13]). Flavonoids are not just interesting colourants, but have been reported to have nutritional benefits [14] and even potential in medical applications [15]. Reported anti-oxidative, anti-inflammatory, anti-mutagenic, and anti-carcinogenic properties of flavonoids provide health benefits to humans [16]. For example, kaempferols are assumed to inhibit cancer cell growth and induce cancer cell apoptosis [17]. Heterologous production of flavonoids in plants is considered a promising option to meet customers’ demands. Studies already demonstrated that the production of anthocyanins in plant cell cultures is possible [18,19].

The flavonoid biosynthesis is one of the best-studied pathways in plants thus serving as a model system for the investigation of specialised metabolism [9]. Academic interest in the synthesis of flavonoids spans multiple fields including molecular genetics, chemical ecology, biochemistry, and health sciences [9,20]. Especially the three subgroups flavonols, anthocyanins, and PAs are well studied in the model organism *Arabidopsis thaliana* [21]. Since a partial lack of flavonoids is not lethal under most conditions, there are large numbers of mutants with visible phenotypes caused by the knockout of various genes in the pathway [22]. For example, seeds lacking PAs show a yellow phenotype due to the absence of brown pigments in the seed coat which inspired the name of mutants in this pathway: *transparent testa* [23]. While the early steps of the flavonoid aglycon biosynthesis are very well known, some later steps require further investigation. Especially the transfer of sugars to PAs and anthocyanidins offer potential for future discoveries [24].

The core pathway of the flavonoid aglycon biosynthesis comprises several key steps which allow effective channeling of substrates in specific branches (Figure 1). A type III polyketide synthase, the chalcone synthase (CHS), catalyses the initial step of the flavonoid biosynthesis which is the conversion of *p*-coumaroyl-CoA and three malonyl-CoA into naringenin chalcone [25]. Since a knock-out or down-regulation of this step influences all branches of the flavonoid biosynthesis, CHS is well studied in a broad range of species. Flower colour engineering with CHS resulted in the identification of mechanisms for the suppression of gene expression [26]. *A. thaliana* CHS can be distinguished from very similar stilbene synthases (STS) based on two diagnostic amino acid residues Q166 and Q167, while a STS would show Q166 H167 or H166 Q167 [27]. The chalcone isomerase (CHI) catalyses the conversion of bicyclic chalcones into tricyclic (S)-flavanones [28]. CHI I converts 6’-tetrahydroxychalcone to 5-hydroxyflavanone, while CHI II additionally converts 6’-deoxychalcone to 5-dexoyflavanone [29]. An investigation of CHI in early land plants revealed the presence of CHI II, which is in contrast to the initial assumption that CHI II activity would be restricted to legumes [30]. A detailed theory about the evolution of functional CHIs from non-enzymatic fatty acid binding proteins and the origin of CHI-like proteins was developed based on evolution experiments [31]. The CHI product naringenin can be processed by different enzymes broadening the flavonoid biosynthesis pathway to a metabolic network.

Flavanone 3β-hydroxylase (F3H/FHT) catalyses 3-hydroxylation of naringenin to dihydroflavonols [32]. As a member of the 2-oxoglutarate-dependent dioxygenase (2-ODD) family, F3H utilises the same cofactors and cosubstrate as the two other 2-ODD enzymes in the flavonoid biosynthesis: flavonol synthase (FLS) and leucoanthocyanidin dioxygenase (LDOX) / anthocyanidin synthase (ANS) [33]. The 2-ODD enzymes share overlapping substrate and product selectivities [34]. FLS was identified to be a bifunctional enzyme showing F3H activity in some species including *A. thaliana* [35], *Oryza sativa* [36], and *Ginkgo biloba* [37]. ANS, an enzyme of a late step in the flavonoid biosynthesis pathway, can have both, FLS and F3H activity [38–41]. Due to its FLS side-activity, ANS has to be considered as an additional candidate for the synthesis of flavonols. The flavonoid 3’-hydroxylase (F3’H) catalyses the conversion of naringenin to eriodictyol and the conversion of dihydrokaempferol to dihydroquercetin [42]. Expression and activity of flavonoid 3’5’-hydroxylase (F3’5’H) is essential for the formation of 5’-hydroxylated anthocyanins which cause the blue colour of flowers [13,43]. F3’5’H competes with FLS for dihydroflavonols thus it is possible that F3’5’H processes only the excess of these substrates that surpass the FLS capacity [44]. Functionality of enzymes like F3’5’H or F3’H is determined by only a few amino acids. A T487S mutation converted a *Gerbera hybrida* F3’H into a F3’5’H and the reverse mutation in an *Osteospermum hybrida* F3’5’H deleted the F3’5’H activity almost completely while F3’H activity remained [45]. The central enzyme in the flavonol biosynthesis is FLS, which converts a dihydroflavonol into the corresponding flavonol by introducing a double bond between C-2 and C-3 of the heterocylic pyran ring (Figure 1)[46,47]. FLS activity was first identified in irradiated parsley cells [48] and has then been characterised in several species including *Petunia hybrida* [46], *A. thaliana* [49], and *Zea mays* [24], revealing species-specific substrate specificities and affinities.

Another branching pathway channels naringenin into the flavone synthesis. Together with flavonols, flavones occur as primary pigments in white flowers and function as co-pigments with anthocyanins in blue flowers [50]. Flavanones can be oxidized to flavones by flavanone synthase I (FNS I) [51] and FNS II [52]. Hence, FNS I and FNS II compete with F3H for flavanones and present a branching reaction in the flavonoid biosynthesis [53]. Being a 2-ODD, FNS I shows only minor differences in its catalytic mechanism compared to F3H, which are determined by only seven amino acid residues [53]. The exchange of all seven residues in parsley F3H resulted in a complete change to FNS I activity [53].

Colourful pigments are generated in the anthocyanin and proanthocyanidin biosynthesis. The NADPH-dependent reduction of dihydroflavonols to leucoanthocyanidins by dihydroflavonol-4-reductase (DFR) is the first committed step of the anthocyanin and proanthocyanidin biosynthesis. There is a competition between FLS and DFR for dihydroflavonols [54]. DFR enzymes have different preferences for various dihydroflavonols (dihydrokaempferol, dihydroquercetin, and dihydromyricetin). The molecular basis of these preferences are probably due to differences in a 26-amino acid substrate binding domain of these enzymes [55]. N at position 3 of the substrate determining domain was associated with recognition of all three dihydroflavonols [55]. D at position 3 prevented the acceptance of dihydrokaempferols [55], while a L or A lead to a preference for dihydrokaempferol and substantially reduced the processing of dihydromyricetin [55,56]. Although this position is central for the substrate specificity, other positions contribute to the substrate specificity [57]. ANS catalyses the last step in the anthocyanin aglycon biosynthesis, the conversion of leucoanthocyanidins into anthocyanidins. The NADPH/NADH-dependent isoflavone-like reductases, leucoanthocyanidin reductase (LAR) / leucocyanidin reductase (LCR), and anthocyanidin reductase (ANR, encoded by *BANYULS* (*BAN*)) are members of the reductase epimerase dehydrogenase superfamily [58]. LAR channels leucoanthocyanidins into the proanthocyanidin biosynthesis which is in competition with the anthocyanidin formation catalysed by ANS. There is also a competition between 3-glucosyltransferases (3GT) and ANR for anthocyanidins [59]. While 3GT generates stable anthocyanins through the addition of a sugar group to anthocyanidins, ANR channels anthocyanidins into the proanthocyanidin biosynthesis. Anthocyanidins are instable in aqueous solution and fade rapidly unless the pH is extremely low [60]. Suppression of *ANR1* and *ANR2* in *Glycine max* caused the formation of red seeds through a reduction in proanthocyanidin biosynthesis and an increased anthocyanin biosynthesis [61]. Substrate preferences of ANR can differ between species as demonstrated for *A. thaliana* and *M. truncatula* [62].

As a complex metabolic network with many branches, the flavonoid biosynthesis requires sophisticated regulation. Activity of different branches is mainly regulated at the transcriptional level [63]. In *A. thaliana* as in many other plants, R2R3-MYBs [64,65] and basic helix-loop-helix proteins (bHLH) [66] are two main transcription factor families involved in the regulation of the flavonoid biosynthesis. The WD40 protein TTG1 facilitates the interaction of R2R3-MYBs and bHLHs in the regulation of the anthocyanin and proanthocyanidin biosynthesis in *A. thaliana* [67]. Due to its components, this trimeric complex is also referred to as MBW complex [67]. Examples of MBW complexes are MYB123 / bHLH42 / TTG1 and MYB75 / bHLH2 / TTG1, which are involved in anthocyanin biosynthesis regulation in a tissue-specific manner [68]. However, the bHLH-independent R2R3-MYBs like MYB12, MYB11, and MYB111 can activate as single transcriptional activators early genes of the flavonoid biosynthesis including *CHS*, *CHI*, *F3H*, and *FLS* [69].

Many previous studies performed a systematic investigation of the flavonoid biosynthesis in plant species including *Fragaria x ananassa* [70], *Musa acuminata* [71], Tricyrtis spp. [72], and multiple *Brassica* species [73]. In addition to these systematic investigations, genes of the flavonoid biosynthesis are often detected as differentially expressed in transcriptomic studies without particular focus on this pathway [74–76]. In depth investigation of the flavonoid biosynthesis starts with the identification of candidate genes for all steps. This identification of candidates is often relying on an existing annotation or requires tedious manual inspection of sequence alignments. As plant genome sequences and their structural annotations become available with an increasing pace [77], the timely addition of functional annotations is an ever increasing challenge. Therefore, we developed a pipeline for the automatic identification of flavonoid biosynthesis players in any given set of peptide, transcript, or genomic sequences. As a proof of concept, we validate the predictions made by Knowledge-based Identification of Pathway Enzymes (KIPEs) with a manual annotation of the flavonoid biosynthesis in the medicinal plant *Croton tiglium*. *C. tiglium* is a member of the family Euphorbiaceae [78] and was first mentioned over 2,200 years ago in China as a medicinal plant probably because of the huge variety of specialised metabolites [79]. Oil of *C. tiglium* was traditionally used to treat gastrointestinal disorders and may have abortifacient and counterirritant effects [80]. Additionally, *C. tiglium* produces phorbol esters and a ribonucleoside analog of guanosine with antitumor activity [81,82]. Characterization of the specialised metabolism of *C. tiglium* will facilitate the unlocking of its potential in agronomical, biotechnological, and medical applications. The flavonoid biosynthesis of *C. tiglium* is largely unexplored. To the best of our knowledge, previous studies only showed the presence of flavonoids through analysis of extracts [83–85]. However, transcriptomic resources are available [86] and provide the basis for a systematic investigation of the flavonoid biosynthesis in *C. tiglium*.

A huge number of publicly available genome and transcriptome assemblies of numerous plant species provide a valuable resource for comparative analysis of the flavonoid biosynthesis. Here, we present an automatic workflow for the identification of flavonoid biosynthesis genes applicable to any plant species and demonstrate the functionality by analyzing a *de novo* transcriptome assembly of *C. tiglium*.

## 2. Results

We developed a tool for the automatic identification of enzyme sequences in a set of peptide sequences, a transcriptome assembly, or a genome sequence. Knowledge-based Identification of Pathway Enzymes (KIPEs) identifies candidate sequences based on overall sequence similarity, functionally relevant amino acid residues, and functionally relevant domains (Figure 2). As a proof of concept, the transcriptome assembly of *Croton tiglium* was screened with KIPEs to identify the flavonoid aglycon biosynthesis network. Results of the automatic annotation are validated by a manually curated annotation.

**Figure 2.**
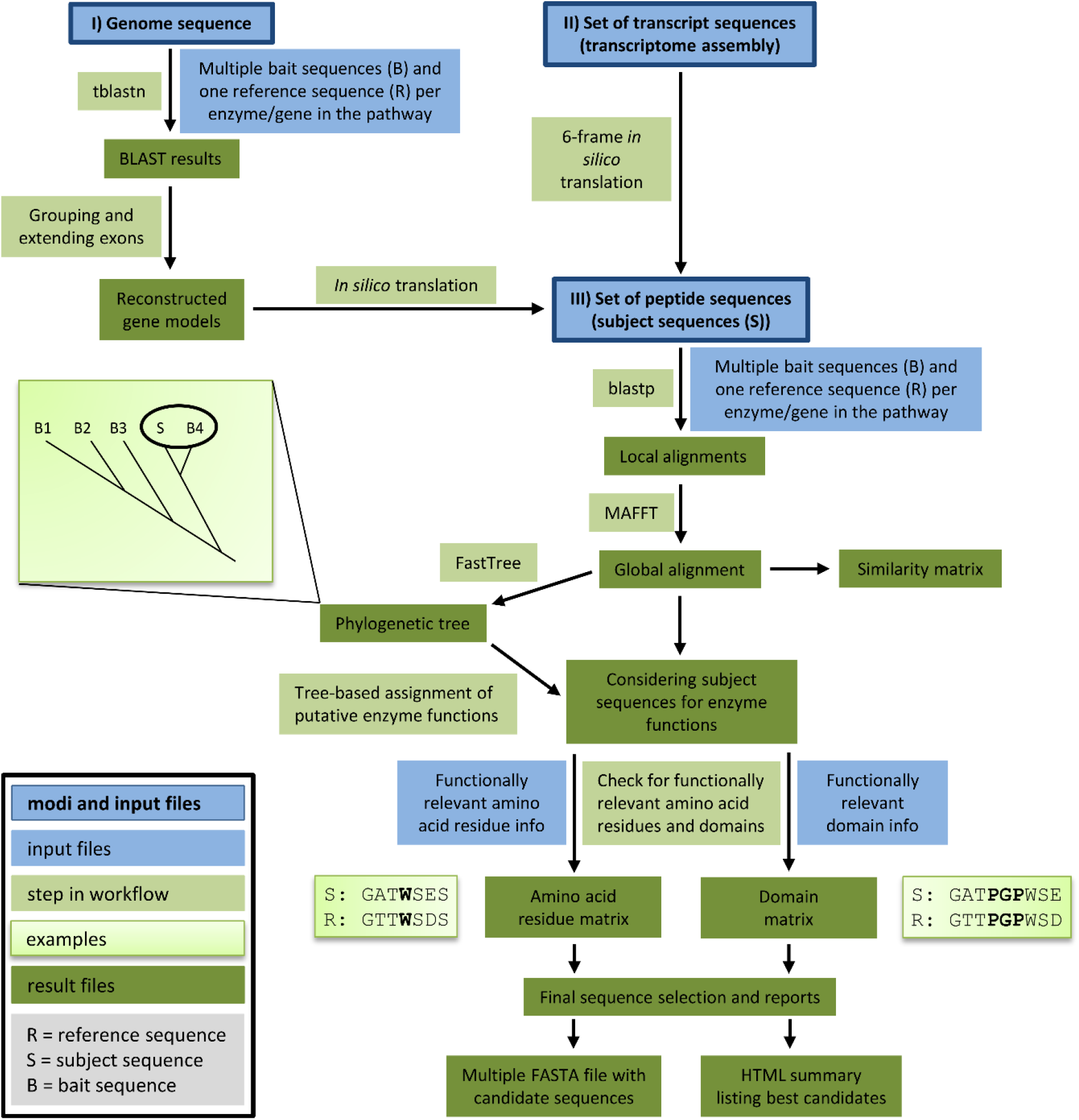
This overview illustrates the components and steps of Knowledge-based Identification of Pathway Enzymes (KIPEs). Three different modes allow the screening of peptide, transcript, or genome sequences for candidate sequences. Bait sequences and information about functionally relevant features (blue) are supplied by the user. Different modules of KIPEs (light green) are executed consecutively depending on the type of input data. Intermediate results and the final output (dark green) are stored to keep the process transparent.

### 2.1. Concept and components of Knowledge-based Identification of Pathway Enzymes (KIPEs)

#### 2.1.1. General concept

The automatic detection of sequences encoding enzymes of the flavonoid biosynthesis network requires (1) a set of bait sequences covering a broad taxonomic range and (2) information about functionally relevant amino acid residues and domains. Bait sequences were selected to encode enzymes with evidence of functionality i.e. mutant complementation studies or *in vitro* assays. Additional bait sequences were included which were previously studied in comparative analyses of the particular enzyme family. Positions of amino acids and domains with functional relevance need to refer to a reference sequence included in the bait sequence set. All bait sequences and one reference sequence related to one reaction in the network are supplied in one FASTA file. However, many FASTA files can be provided to cover all reactions of a complete metabolic network. Positions of functionally relevant residues and domains are specified in an additional text file based on the reference sequence (see manual for details, https://github.com/bpucker/KIPEs). Collections of bait sequences and detailed information about the relevant amino acid residues in flavonoid biosynthesis enzymes are provided along with KIPEs. However, these collections can be customized by users to reflect updated knowledge and specific research questions. KIPEs was developed to have a minimal amount of dependencies. Only the frequently used alignment tools BLAST and MAFFT are required. Both tools are freely available as precompiled binaries without the need for installation.

#### 2.1.2. Three modes

A user can choose between three different analysis modes depending on the available input sequences: peptide sequences, transcript sequences, or a genome sequence. If a reliable peptide sequence annotation is available, these peptide sequences should be subjected to the analysis. Costs in terms of time and computational efforts are substantially lower for the analysis of peptide sequences than for the analysis of genome sequences. The provided peptide sequences are screened via blastp for similarity to previously characterised bait sequences. If default criteria are applied, BLAST hits are considered if the sequence similarity is above 40% and if the score is above 30% of the score resulting from an alignment of the query sequence against itself. These lenient filter criteria are applied to collect a comprehensive set of candidate sequences which is subsequently refined through the construction of global alignments via MAFFT. Next, phylogenetic trees are generated to identify best candidates based on their position in a tree. Candidates are classified based on the closest distance to a bait sequence. Multiple closely related bait sequences can be considered if specified. When transcript sequences are supplied to KIPEs, *in silico* translation in all six possible frames generates a set of peptide sequences which are subsequently analysed as described above. Supplied DNA sequences are screened for similarity to the bait peptide sequences via tblastn. Hits reported by tblastn are considered exons or exon fragments and therefore assigned to groups which might represent candidate genes. The connection of these hits is attempted in a way that canonical GT-AG splice site combinations emerge. One isoform per locus is constructed and subsequently analysed as described above.

#### 2.1.3. Final filtering

After identification of initial candidates through overall sequence similarity, a detailed comparison against a well characterised reference sequence with described functionally relevant amino acid residues is performed. All candidates are screened for matching amino acid residues at functionally relevant positions. Sequences encoding functional enzymes are expected to display a matching amino acid residue at all checked positions. Additionally, the conservation of relevant domains is analysed. A prediction about the functionality/non-functionality of the encoded enzyme of all candidate sequences is performed at this step. Results of intermediate steps are stored to allow in depth inspection if necessary.

### 2.2. Technical validation of KIPEs

A first technical validation of KIPEs was performed based on sequence data sets of plant species with previously characterized flavonoid biosyntheses namely *A. lyrata*, *A. thaliana*, *Cicer arietinum*, *Fragaria vesca*, *Glycine max*, *Malus domestica*, *Medicago truncatula*, *Musa acuminata*, *Populus trichocarpa*, *Solanum lycopersicum*, *Solanum tuberosum*, *Theobroma cacao*, and *Vitis vinifera*. The flavonoid biosynthesis of these species was previously characterised thus providing an opportunity for validation. KIPEs identified candidate sequences with conservation of all functionally relevant amino acid residues for the expected enzymes in all species (File S1).

### 2.3. The flavonoid biosynthesis enzymes in Croton tiglium

Genes in the flavonoid biosynthesis of *C. tiglium* were identified based on bait sequences of over 200 plant species and well characterised reference sequences of *A. thaliana*, *G. max*, *M. sativa*, *Osteospermum* spec., *Petroselinum crispum*, *P. tomentosa*, and *V. vinifera*. The transcriptome assembly of *C. tiglium* revealed sequences encoding enzymes for all steps in the flavonoid biosynthesis (Table 1). Phylogenetic analyses placed the *C. tiglium* sequences of enzymes in the flavonoid biosynthesis close to the corresponding sequences of related *Malpighiales* species like *Populus tomentosa* (File S2). Conservation of functionally relevant amino acid residues was inspected in an alignment with sequences of characterised enzymes of the respective step (File S3).

**Table 1.**
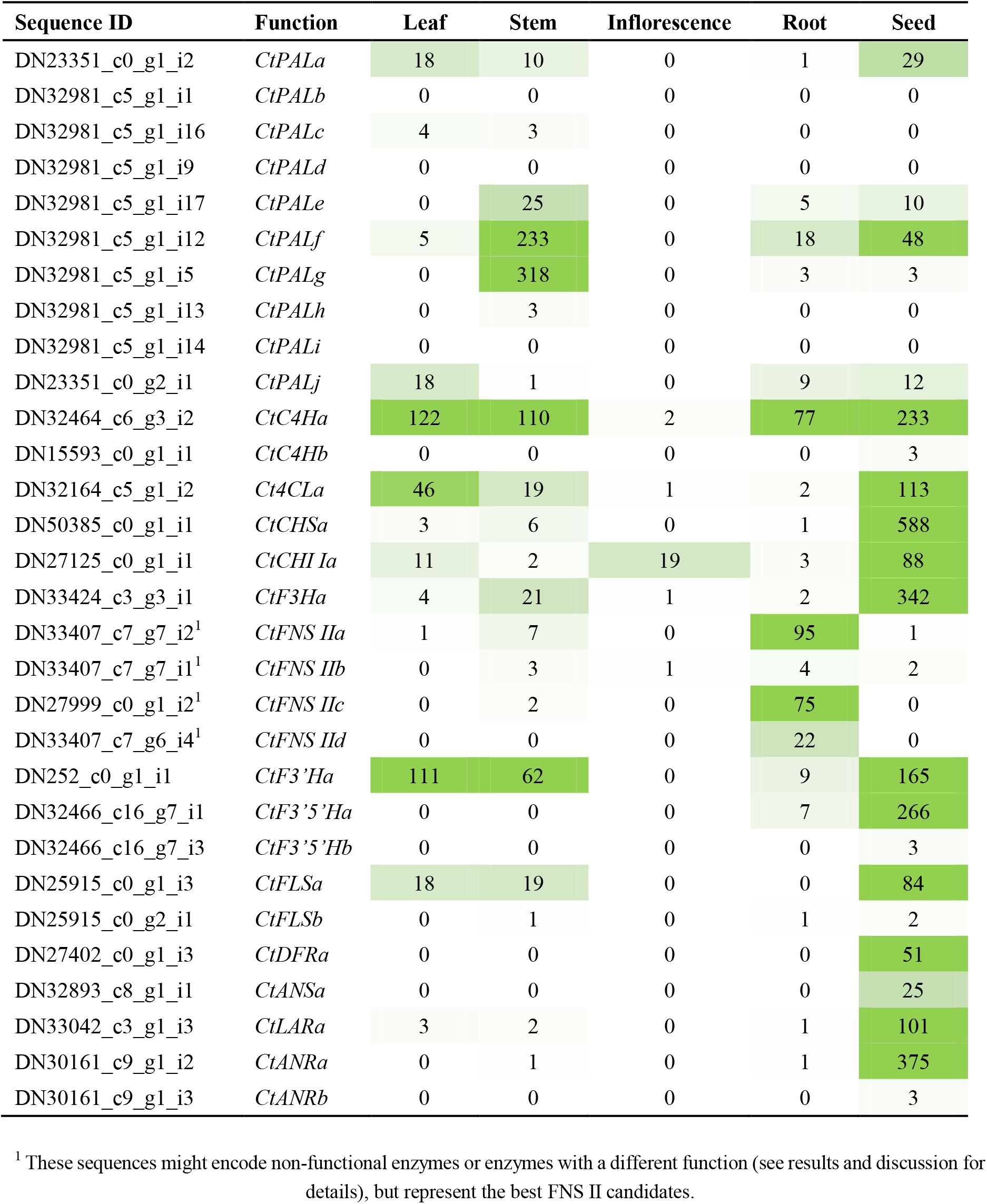
Candidates in the flavonoid biosynthesis of *Croton tiglium*. ‘TRINITY’ prefix of all sequence names was omitted for brevity. Candidates are sorted by their position in the respective pathway and decreasing similarity to bait sequences. Transcripts per million (TPM) values of the candidates in different tissues are shown: leaf (SRR6239848), stem (SRR6239849), inflorescence (SRR6239850), root (SRR6239851), and seed (SRR6239852). Displayed values are rounded to the closest integer thus extremely low abundances are listed as 0. A full table with all available RNA-Seq samples and transcript abundance values for all candidates is available in the supplements (File S6).

The general phenylpropanoid biosynthesis is represented by ten phenylalanine ammonia lyase (PAL) candidates, two cinnamate 4-hydroxylases (C4H) candidates, and one 4-coumarate-CoA ligase (4CL) candidate (Table 1, File S4, File S5). Many PAL sequences show a high overall sequence similarity indicating that multiple alleles or isoforms could contribute to the high number. A phylogenetic analysis supports the hypothesis that many PAL candidates might be alleles or alternative transcript variants of the same genes (File S2). Very low transcript abundances indicate that at least three of the PAL candidates can be neglected (Table 1).

Although multiple CHS candidates were identified based on overall sequence similarity to the *A. thaliana* CHS sequence, only CtCHSa showed all functionally relevant amino acid residues (File S3). Five other candidates were discarded due to the lack of Q166 and Q167, which differentiate CHS from other polyketide synthases like STS or LAP5. Additionally, a CHS signature sequence at the C-terminal end and the malonyl-CoA binding motif at position 313 to 329 in the *A. thaliana* sequence are conserved in CtCHSa. A phylogenetic analysis supported these findings by placing CtCHSa in a clade with *bona fide* chalcone synthases (File S2). There is only one CHI candidate, CtCHI Ia, which contains all functionally relevant amino acid residues (File S3). No CHI II candidate was detected. *C. tiglium* has one F3H candidate, one F3’H candidate, and two F3’5’H candidates. CtF3Ha, CtF3’Ha, CtF3’5’Ha, and CtF3’5’Hb show conservation of the respective functionally relevant amino acid residues (File S3). CtF3’Ha contains the N-terminal proline rich domain and a perfectly conserved oxygen binding pocket at position 302 to 307 in the *A. thaliana* reference sequence. Both, CtF3’5’Ha and CtF3’5’Hb, were also considered as F3’H candidates, but show overall a higher similarity to the F3’5’H bait sequences than to the F3’H bait sequences. The flavone biosynthesis capacities of *C. tiglium* remained elusive. No FNS I candidates with conservation of all functionally relevant amino acids were detected. However, there are four FNS II candidates which show only one substitution of an amino acid residue in the oxygen binding pocket (T313F). The committed step of the flavonol biosynthesis is represented by CtFLSa and CtFLSb which show all functionally relevant residues (File S3).

*C. tiglium* contains excellent candidates for all steps of the anthocyanidin and proanthocyanidin biosynthesis. CtDFR shows conservation of the functionally relevant amino acid residues (File S3). We investigated the substrate specificity domain to understand the enzymatic potential of the DFR in *C. tiglium*. Position 3 of this substrate specificity domain shows a D which is associated with low acceptance of dihydrokaempferols. CtLAR is the only LAR candidate with conservation of the functionally relevant amino acid residues (File S3). CtANS is the only ANS candidate with conservation of the functionally relevant amino acid residues (File S3). There are two ANR candidates in *C. tiglium*. CtANRa and CtANRb show conservation of all functionally relevant amino acid residues (File S3). CtANRa shows 74% identical amino acid residues when compared to the reference sequence, which exceeds the 49% of CtANRb substantially.

The identification of candidates in a transcriptome assembly already shows transcriptional activity of the respective gene. To resolve the transcriptional activity of genes in greater detail, we quantified the presence of candidate transcripts in different tissues of *C. tiglium* and compared it to *C. draco* through cross-species transcriptomics (File S6). High transcript abundance of almost all flavonoid biosynthesis candidates was observed in seeds, while only a few candidate transcripts were observed in other investigated tissues (Table 1). Transcripts involved in the proanthocyanidin biosynthesis show an exceptionally high abundance in seeds of *C. tiglium* and inflorescence of *C. draco*. Overall, the tissue specific abundance of many transcripts is similar between *C. tiglium* and *C. draco*. LAR and ANR show substantially higher transcript abundances in inflorescences of *C. draco* compared to *C. tiglium*. CHS and ANS show the highest transcript abundance in pink flowers of *C. draco* (File S6).

### 2.4. Transcriptional regulators of the flavonoid biosynthesis in Croton tiglium

To demonstrate the applicability of KIPEs for the investigation of non-enzyme sequences like transcription factor gene families, we screened the transcriptome assembly of *C. tiglium* for members of the MYB, bHLH, and WD40 family. This analysis revealed candidates for some key regulators of the flavonoid biosynthesis namely MYB11/MYB12/MYB111 (subgroup7), MYB123 (subgroup5), MYB75/MYB90/MYB113/MYB114 (subgroup6), bHLH2/bHLH42, and TTG1 according to the nomenclature in *A. thaliana* (Table 2, File S7). The MYB subgroups 6 and 7 have multiple members in *A. thaliana* and *C. tiglium*. Therefore, *C. tiglium* candidates are only assigned to an orthogroup (Table 2). The reliable identification of MYB orthologs between both species was not feasible (File S7). There are five homologous sequences of MYB123 in *C. tiglium* with one of them probably originating from the same gene. The R2R3 MYB domain was detected in the MYB candidates except for DN21046_c0_g1_i3, DN21046_c0_g1_i3, DN30455_c10_g1_i1, and DN33314_c5_g2_i4. With the exception of DN33314_c5_g2_i4 (truncated protein) all CtMYB candidates of subgroup6 have a conserved bHLH interaction domain, while the CtMYB candidates of the bHLH-independent subgroup7 do not show this conserved domain. There are seven *C. tiglium* sequences in a clade with the *A. thaliana* bHLH42 (File S7), but these might be alternative isoforms originating from the same gene. The same is true for the seven isoforms detected as homologous sequences of *A. thaliana* bHLH2 (File S7). Three TTG1 candidates exist in the *C. tiglium* transcriptome assembly, but two of them might be isoforms belonging to the same gene. The MYB, bHLH, and TTG1 transcription factor candidates show generally lower transcript abundances than the enzyme candidates (Table 1, Table 2). The highest transcript abundance of all three MBW complex components was observed in seeds.

**Table 2.** Transcriptional regulator candidates of the flavonoid biosynthesis. MYB11/MYB12/MYB111 candidates are summarised as subgroup7 MYBs. MYB75/MYB90/MYB113/MYB114 are summarised as subgroup6. Transcripts per million (TPM) values of the candidates in different tissues are shown: leaf (SRR6239848), stem (SRR6239849), inflorescence (SRR6239850), root (SRR6239851), and seed (SRR6239852). Displayed values are rounded to the closest integer thus extremely low abundances are listed as 0.

| Sequence ID | Group | Leaf | Stem | Inflorescence | Root | Seed |
| --- | --- | --- | --- | --- | --- | --- |
| DN30455_c10_g1_i1 | Subgroup7 | 0 | 1 | 0 | 0 | 4 |
| DN21046_c0_g1_i3 | Subgroup7 | 0 | 0 | 0 | 0 | 0 |
| DN21046_c0_g1_i2 | Subgroup7 | 0 | 1 | 0 | 0 | 9 |
| DN28041_c1_g1_i4 | Subgroup6 | 0 | 0 | 0 | 0 | 7 |
| DN28041_c1_g1_i2 | Subgroup6 | 0 | 0 | 0 | 0 | 0 |
| DN33356_c3_g1_i2 | Subgroup6 | 0 | 0 | 0 | 0 | 4 |
| DN31144_c5_g1_i2 | MYB123 | 0 | 0 | 0 | 10 | 29 |
| DN33314_c5_g2_i2 | MYB123 | 3 | 8 | 0 | 1 | 14 |
| DN33314_c5_g2_i3 | MYB123 | 0 | 0 | 0 | 0 | 1 |
| DN33314_c5_g2_i4 | MYB123 | 1 | 2 | 0 | 0 | 6 |
| DN31260_c4_g2_i2 | MYB123 | 0 | 0 | 0 | 0 | 5 |
| DN30681_c1_g1_i1 | bHLH2 | 0 | 0 | 0 | 0 | 0 |
| DN30681_c1_g1_i2 | bHLH2 | 0 | 2 | 0 | 2 | 4 |
| DN30681_c1_g1_i3 | bHLH2 | 0 | 0 | 0 | 0 | 0 |
| DN30681_c1_g1_i6 | bHLH2 | 0 | 0 | 0 | 0 | 0 |
| DN30681_c1_g1_i7 | bHLH2 | 2 | 13 | 0 | 16 | 16 |
| DN30681_c1_g1_i8 | bHLH2 | 2 | 9 | 0 | 20 | 18 |
| DN30681_c1_g1_i9 | bHLH2 | 0 | 0 | 0 | 0 | 0 |
| DN32219_c4_g2_i2 | bHLH42 | 0 | 0 | 0 | 0 | 0 |
| DN32219_c4_g2_i5 | bHLH42 | 0 | 0 | 0 | 0 | 2 |
| DN32219_c4_g2_i12 | bHLH42 | 0 | 0 | 0 | 0 | 2 |
| DN32219_c4_g2_i11 | bHLH42 | 0 | 0 | 0 | 0 | 3 |
| DN32219_c4_g2_i4 | bHLH42 | 0 | 1 | 0 | 0 | 25 |
| DN32219_c4_g2_i7 | bHLH42 | 0 | 0 | 0 | 0 | 4 |
| DN32219_c4_g2_i8 | bHLH42 | 0 | 0 | 0 | 0 | 5 |
| DN32272_c1_g1_i1 | TTG1 | 0 | 0 | 0 | 0 | 0 |
| DN32272_c1_g2_i2 | TTG1 | 12 | 8 | 4 | 10 | 9 |
| DN32604_c4_g1_i2 | TTG1 | 0 | 1 | 0 | 1 | 1 |

## 3. Discussion

As previous studies of extracts from *Croton tiglium* and various other *Croton* species revealed the presence of flavonoids [84,85,87–92], steps in the central flavonoid aglycon biosynthesis network should be represented by at least one functional enzyme each. However, this is the first identification of candidates involved in the biosynthesis. Previous reports [84,85,87–92] about flavonoids align well with our observation (Table 1) that at least one predicted peptide contains all previously described functionally relevant amino acid residues of the respective enzyme. The only exception is the flavone synthase step. While FNS I is frequently absent in flavonoid producing species outside the *Apiaceae*, FNS II is more broadly distributed across plants [53]. *C. tiglium* is not a member of the *Apiaceae* thus the absence of FNS I and the presence of FNS II candidates are expected.

All candidate sequences of presumably functional enzymes belong to actively transcribed genes as indicated by the presence of these sequences in a transcriptome assembly. Since the flavonoid biosynthesis is mainly regulated at the transcriptional level [63] and previously reported blocks in the pathway are expected to be due to transcriptional down-regulation [93,94], we expect most branches of the flavonoid biosynthesis in *C. tiglium* to be functional. No CHI II candidate was detected thus *C. tiglium* probably lacks a 6’-deoxychalcone to 5-dexoyflavanone catalytic activity like most non-leguminous plants [30,95].

A domination of proanthocyanidins has been reported for *Croton* species [88]. This high proanthocyanidin content correlates well with high transcript abundance of proanthocyanidin biosynthesis genes (*CtLAR*, *CtANR*). PAs have been reported to account for up to 90% of the dried weight of red sap of *Croton lechleri* [96]. Expression of *CtFLSa* in the leaves matches previous reports about flavonol extraction from leaves [90,97]. Interestingly, almost all analysed *Croton* species showed very high amounts of quercetin derivates compared to kaempferol derivates in their leaf extracts, which significantly correlated with antioxidant potential [97]. This high quercetin concentration might be due to a high expression level of *CtF3’Ha* in leaves. Since F3’H converts dihydrokaempferol (DHK) to dihydroquercetin (DHQ), a high gene expression might result in high amounts of DHQ which can be used from FLS to produce quercetin. At the same time, the production of kaempferols from DHK is reduced.

Flavonols have been extracted from several *Croton* species and several important functions have been attributed to these flavonols. Quercetin 3,7-dimethyl ether was extracted from *Croton schiedeanus* and elicits vasorelaxation in isolated aorta [91]. Casticin a methyoxylated flavonol from *Croton betulaster* modulates cerebral cortical progenitors in rats by directly decreasing neuronal death, and indirectly via astrocytes [98]. Besides the anticancer activity of flavonol rich extracts from *Croton celtidifolius* in mice [99], flavonols extracted from *Croton menyharthii* leaves possess antimicrobial activity [100]. Kaempferol 7-O-β-D-(6″-O-cumaroyl)-glucopyranoside isolated from *Croton piauhiensis* leaves enhanced the effect of antibiotics and showed antibacterial activity on its own [101]. Flavonols extracted from *Croton cajucara* showed anti-inflammatory activities [102].

The investigating of the CtDFR substrate specificity revealed aspartate at the third position of the substrate specificity domain which was previously reported to reduce the acceptance of dihydrokaempferol [55]. Although the substrate specificity of DFR is not completely resolved, a high DHQ affinity would fit to the high transcript abundance of *CtF3’Hs* which encode putative DHQ producing enzymes. Further investigations are needed to reveal how effectively *C. tiglium* produces anthocyanidins and proanthocyanidins based on different dihydroflavonols. As *C. tiglium* is known to produce various proanthocyanidins [83], a functional biosynthetic network must be present. Phlobatannine have been reported in leaves of *C. tiglium* [83] which aligns well with our identification of a probably functional CtDFRa.

Our automatic approach for the identification of flavonoid biosynthesis genes could be applied to identify target genes for an experimental validation in a species with a newly sequenced transcriptome or genome. Due to multiple refinement steps, the predictions of KIPEs have a substantially higher fidelity than frequently used BLAST results. Especially the distinction of different enzymes with very similar sequences (e.g. CHS, STS, LAP5) was substantially improved by KIPEs. Additionally, the automatic identification of flavonoid biosynthesis enzymes/genes across a large number of plant species facilitates comparative analyses which could be a valuable addition to functional studies or might even replace some studies. As functionally relevant amino acid residues are well described for many of the enzymes, an automatic classification of candidate sequences as functional or non-functional is feasible in many cases. It has not escaped our notice that ‘non-functionality’ only holds with respect to the initially expected enzyme function. Sub- and neofunctionalisation, especially following gene duplications, are likely. Results produced by KIPEs could be used to identify species-specific modifications of the general flavonoid biosynthesis. Bi- or even multifunctionality has been described for some members of the 2-ODDs (FLS [36,103,104], F3H, FNS I, and ANS [38–41]). Experimental characterization of these enzymes will still be required to determine the degree of the possible multifunctionalities in one enzyme. However, enzyme characterization experiments could be informed by the results produced by KIPEs. As KIPEs has a particular focus on high impact amino acid substitutions, it would also be possible to screen sequence data sets of phenotypically interesting plants to identify blocks in pathways. Another potential application is the assessment of the functional impact of amino acid substitutions e.g. in re-sequencing studies. There are established tools like SnpEff [105] for the annotation of sequence variants in re-sequencing studies. Additionally, KIPEs could operate on the set of modified peptide sequences to analyse the functional relevance of sequence variants. If functionally relevant amino acids are effected, KIPEs could predict that the variant might cause non-functionality.

Although KIPEs can be applied to screen a genome sequence, we recommend to supply peptide or transcript sequences as input whenever possible. Well established gene prediction tools like AUGUSTUS [106] and GeMoMa [107] generate gene models of superior quality in most cases. KIPEs is restricted to the identification of canonical GT-AG splice sites. The very low frequency of non-canonical splice sites in plant genomes [108] would cause extreme computational costs and could lead to a substantial numbers of mis-annotations. To the best of our knowledge, non-canonical splice sites have not been reported for genes in the flavonoid biosynthesis. Nevertheless, dedicated gene prediction tools can incorporate additional hints to predict non-canonical introns with high fidelity.

During the identification of amino acid residues which were previously reported to be relevant for the enzyme function, we observed additional patterns. Certain positions showed not perfect conservation, but multiple amino acids with similar biochemical properties occurred at the respective position. Low relevance of the amino acid at these positions for the enzymatic activity could be one explanation. However, these patterns could also point to lineage specific specializations of various enzymes. A previous study reported the evolution of different F3’H classes in monocots [109]. Subtle differences between isoforms might cause different enzyme properties e.g. altered substrate specificities which could explain the presence of multiple isoforms of the same enzyme in some species. For example, a single amino acid has substantial influence on the enzymatic functionality of F3’H and F3’5’H [45]. This report matches our observation of both F3’5’H candidates being initially also considered as F3’H candidates. A higher overall similarity to the F3’5’H bait sequences than to the F3’H bait sequences allowed an accurate classification. This example showcases the challenges when assigning enzyme functions to peptide sequences.

We developed KIPEs for the automatic identification and annotation of core flavonoid biosynthesis enzymes, because this pathway is well characterised in numerous plant species. Additionally, we demonstrate the applicability for the identification of gene families by screening the transcriptome assembly for MYB, bHLH, and WD40 candidates. Quality and fidelity of the KIPEs results depend on the quality of the bait sequence set and the knowledge about functionally relevant amino acid residues. Nevertheless, the implementation of KIPEs allows the analysis of additional steps of the flavonoid biosynthesis (e.g. the glycosylation of flavonoids) and even the analysis of other pathways. Here, we presented the identification of enzyme candidates based on single amino acid residues with functional relevance. Functionally characterized domains were subordinate in this enzyme detection process. However, KIPEs can also assess the conservation of domains. This function is not only relevant for the analysis of enzymes, but could be applied to the analysis of other proteins like transcription factors with specific binding domains.

## 4. Materials and Methods

### 4.1. Retrieval of bait and reference sequences

The NCBI protein database was screened for sequences of the respective enzyme for all steps in the core flavonoid biosynthesis by searching for the common names. Listed sequences were screened for associated publications about functionality of the respective sequence. Only peptide sequences with evidence for enzyme functionality were retrieved (File S8). To generate a comprehensive set of bait sequences, we also considered sequences with indirect evidence like clear differential expression associated with a phenotype and sequences which were previously included in analyses of the respective enzyme family. The set of bait and reference sequences used for the analyses described in this manuscript is designated FlavonoidBioSynBaits_v1.0.

### 4.2. Collection of information about important amino acid residues

All bait sequences and one reference sequence per step in the flavonoid biosynthesis were subjected to a global alignment via MAFFT v7 [111]. Highly conserved positions, which were also reported in the literature to be functionally relevant, are referred to as ‘functionally relevant amino acid residues’ in this manuscript (File S9). The amino acid residues and their positions in a designated reference sequence are provided in one table per reaction in the network (https://github.com/bpucker/KIPEs). A customized Python script was applied to identify contrasting residues between two sequence sets e.g. chalcone and stilbene synthases (https://github.com/bpucker/KIPEs).

### 4.3. Implementation and availability of KIPEs

KIPEs is implemented in Python 2.7. The script is freely available at github: https://github.com/bpucker/KIPEs. Details about the usage are described in the manual provided along with the Python script. Collections of bait and reference sequences as well as data tables about functionally relevant amino acid residues are included. In summary, these data sets allow the automatic identification of flavonoid biosynthesis genes in other plant species via KIPEs. Customization of all data sets is possible to enable the analysis of other pathways. Mandatory dependencies of KIPEs are blastp [112], tblastn [112], and MAFFT [111]. FastTree2 [113] is an optional dependency which substantially improves the fidelity of the candidate identification and classification. Positions of candidate sequences in a phylogenetic tree are used to identify the closest bait sequences. The function of the closest bait sequence is then transferred to the candidate. However, it is possible to consider a candidate sequence for multiple different functions. If the construction of phylogenetic trees is not possible, the highest similarity to a bait sequence in a global alignment is used instead to predict a function. An analysis of functionally relevant amino acid residues in the candidate sequences is finally used to assign a function.

### 4.4 Phylogenetic analysis

Alignments were generated with MAFFT v7 [111] and cleaned with pxclsq [114] to remove alignment columns with very low occupancy. Phylogenetic trees were constructed with FastTree v2.1.10 [113] using the WAG+CAT model. FigTree (http://tree.bio.ed.ac.uk/software/figtree/) was used to visualize the phylogenetic trees. Alignments were visualized online at http://espript.ibcp.fr/ESPript/ESPript/index.php v3.0 [115] using 3D structures of reference enzymes derived from the Protein Data Bank (PDB) [116] (File S10). If no PDB entry was available, the amino acid sequence of the respective reference enzyme was subjected to I-TASSER [117] for protein structure prediction and modelling (File S10, File S11). Functionally relevant amino acid residues in the *C. tiglium* sequences were subsequently highlighted in the generated PDFs (File S3).

### 4.5 Transcript abundance quantification

All available RNA-Seq data sets of *C. tiglium* [86,118] and *C. draco* [119] were retrieved from the Sequence Read Archive (https://www.ncbi.nlm.nih.gov/sra) via fastq-dump v2.9.6 (https://github.com/ncbi/sra-tools). Kallisto v0.44 [120] was applied with default parameters to quantify the abundance of transcripts based on the *C. tiglium* transcriptome assembly [86].

### 4.6 Application of KIPEs for the identification of transcription factors

KIPEs was run with sets of MYB, bHLH, and WD40 peptide sequences (MYB_bHLH_WD40_v1.0) to identify corresponding candidates in the *C. tiglium* transcriptome assembly. MYB sequences of *A. thaliana* [64], *Vitis vinifera* [121], *Beta vulgaris* [122], and *Musa acuminata* [123] were subject to KIPEs as baits. bHLH bait sequences were collected from *A. thaliana* [124], *V. vinifera* [125], *Nelumbo nucifera* [126], *Citrus grandis* [127], *M. acuminata* [128], and *Solanum melongena* [129]. WD40 sequences of *A. thaliana* [130], *Triticum aestivum* [131], and *Setaria italica* [132] were collected as bait sequences for the identification of the WD40 protein TTG1. Phylogenetic trees with the candidates reported by KIPEs, the sets of bait sequences derived from the genome-wide studies, and selected sequences retrieved from the NCBI were generated with FastTree v2.1.10 [113] based on alignments constructed with MAFFT v7 [111]. MYB domain and bHLH-interaction domain were identified with a Python script (https://github.com/bpucker/bananaMYB) based on previously defined patterns [123].

## 5. Conclusions

KIPEs enables the automatic identification of enzymes involved in flavonoid biosynthesis in uninvestigated sequence data sets of plants, thus paving the way for comparative studies and the identification of lineage specific differences. While we demonstrate the applicability of KIPEs for the identification and sequence-based characterization of players in the core flavonoid biosynthesis, we envision applications beyond this pathway. Various enzymes of entire metabolic networks can be identified if sufficient knowledge about functionally relevant amino acids is available.

## Supplementary Materials

The following are available online: File S1: KIPEs evaluation results, File S2: Phylogenetic trees of candidates, File S3: Multiple sequence alignments of candidates (yellow highlighting is used for functional relevant residues in the *C. tiglium* sequences, acc=relative accessibility, black background indicates perfect conservation across all sequences), File S4: Coding sequences of *C. tiglium* flavonoid biosynthesis genes, File S5: Peptide sequences of *C. tiglium* flavonoid biosynthesis genes, File S6: Gene expression heatmap of all candidate genes, File S7: Unrooted phylogenetic trees of MYB, bHLH, and WD40 candidates in *C. tiglium* and corresponding bait sequences; File S8: List of bait and reference sequences, File S9: Functionally relevant amino acid residues considered for analysis of flavonoid biosynthesis enzymes, File S10: Information about used crystal structures of previously characterized enzymes and protein models produced in this study, File S11: 3D models of flavonoid biosynthesis enzyme structures generated by I-TASSER.

## Author Contributions

B.P. and H.M.S. conceived the project. B.P., F.R., and H.M.S. conducted data analysis. B.P., F.R., and H.M.S. wrote the manuscript. B.P. supervised the project. All authors have read and agreed to the final version of this manuscript.

## Acknowledgments

We are extremely grateful to all researchers who characterised enzymes in the flavonoid biosynthesis, submitted the underlying sequences to the appropriate databases, and published their experimental findings.

