## Supplementary figures and images for "Automatic identification of players in the flavonoid biosynthesis with application on the biomedicinal plant *Croton tiglium*"

### File S2

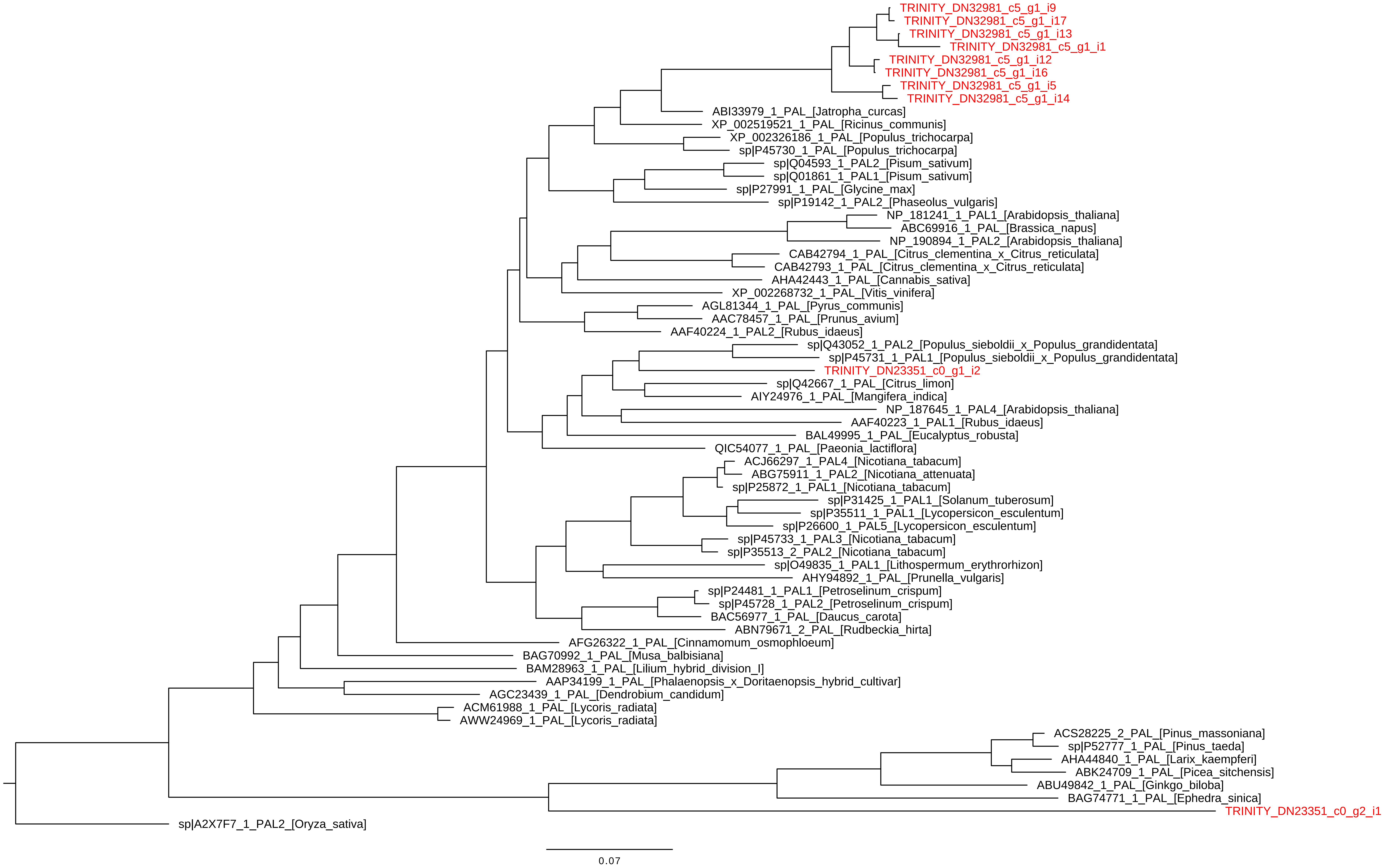

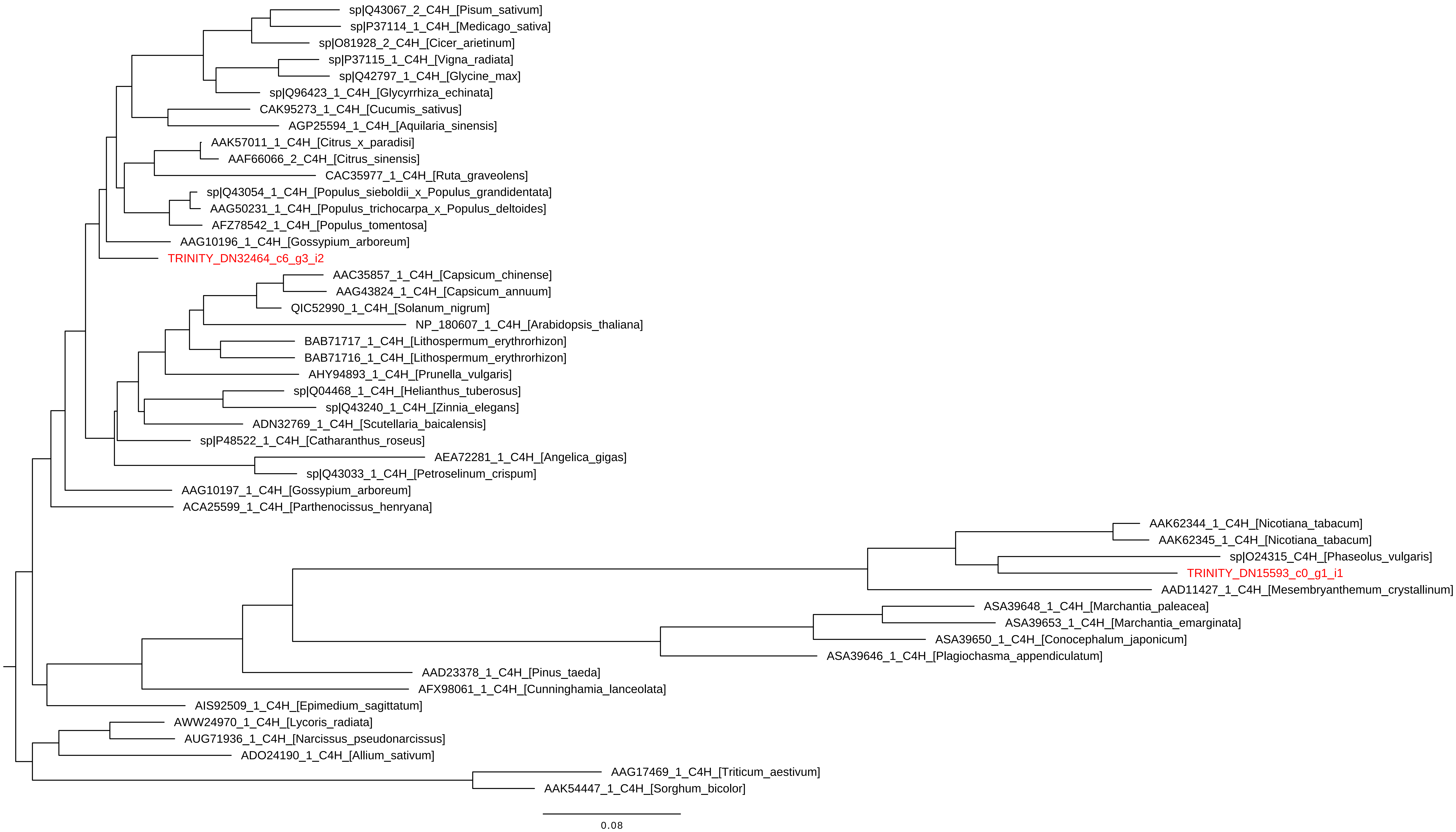

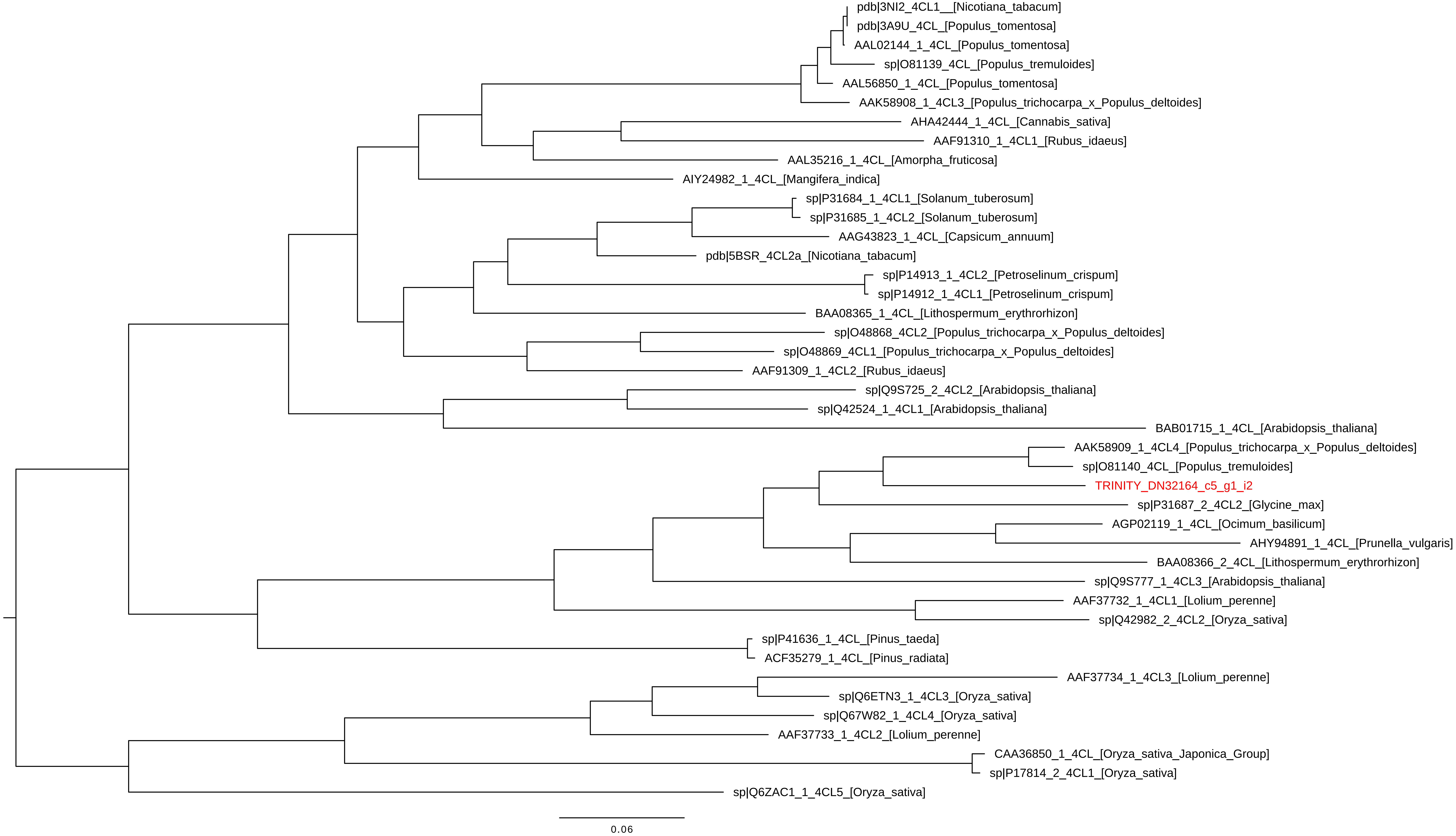

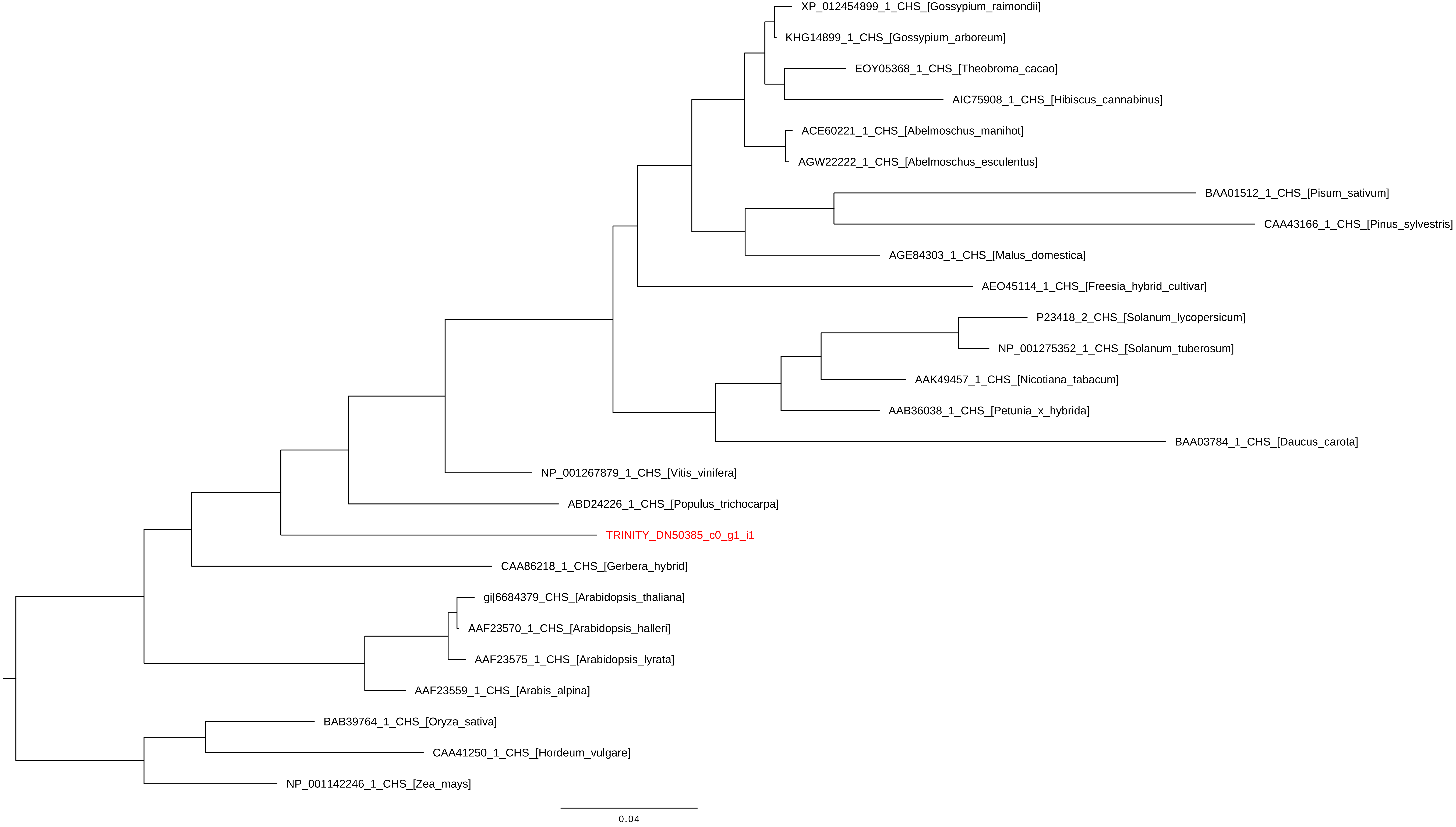

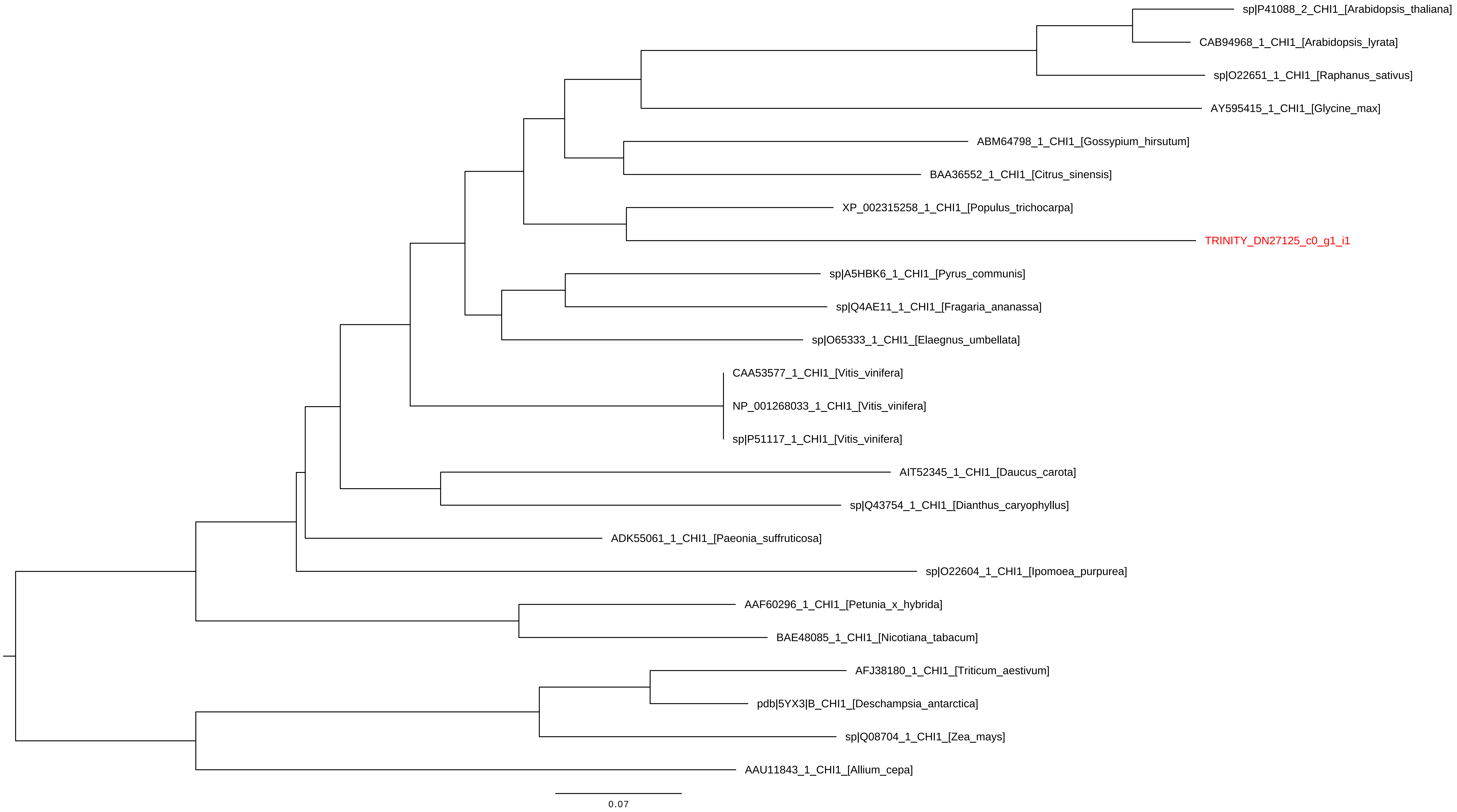

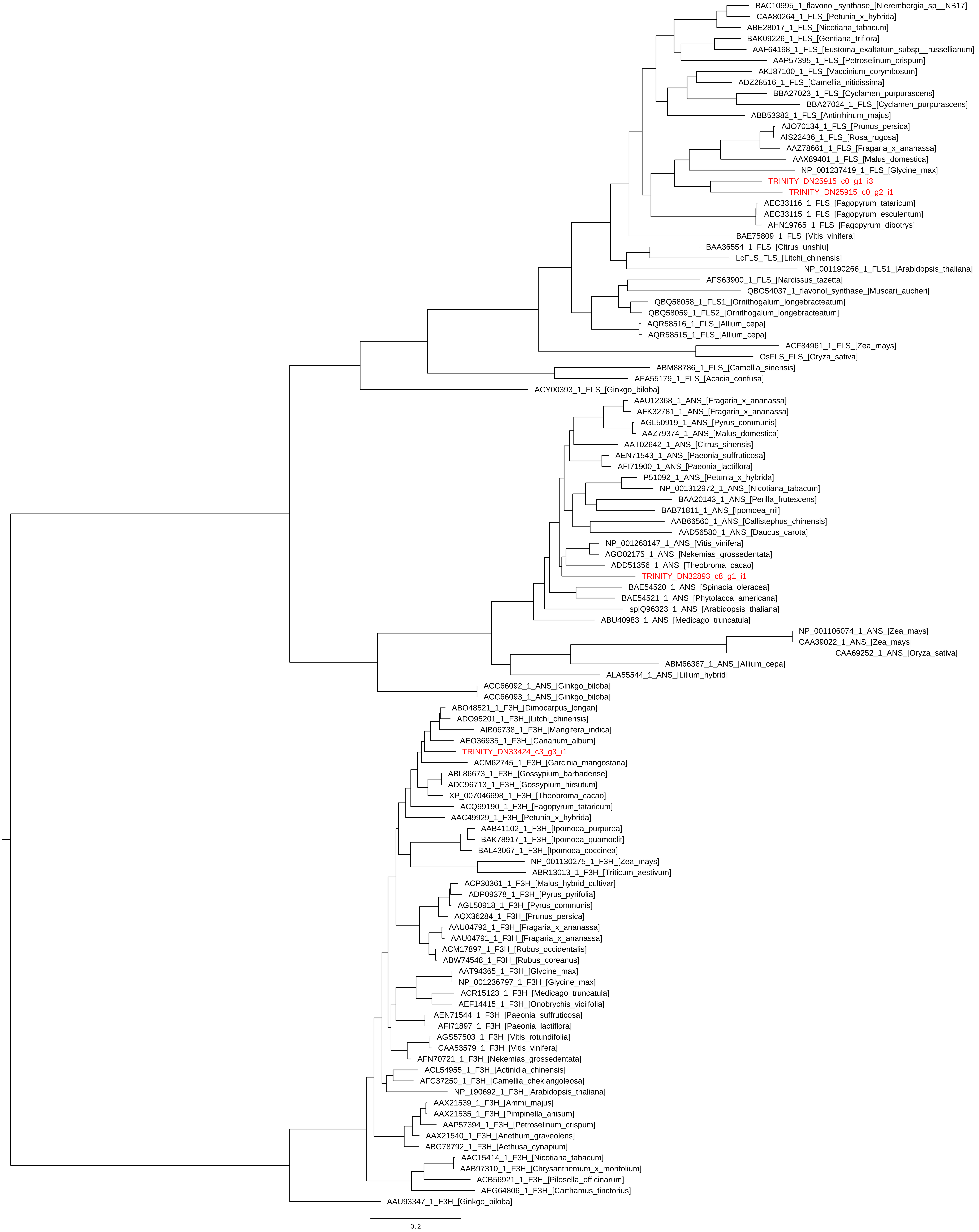

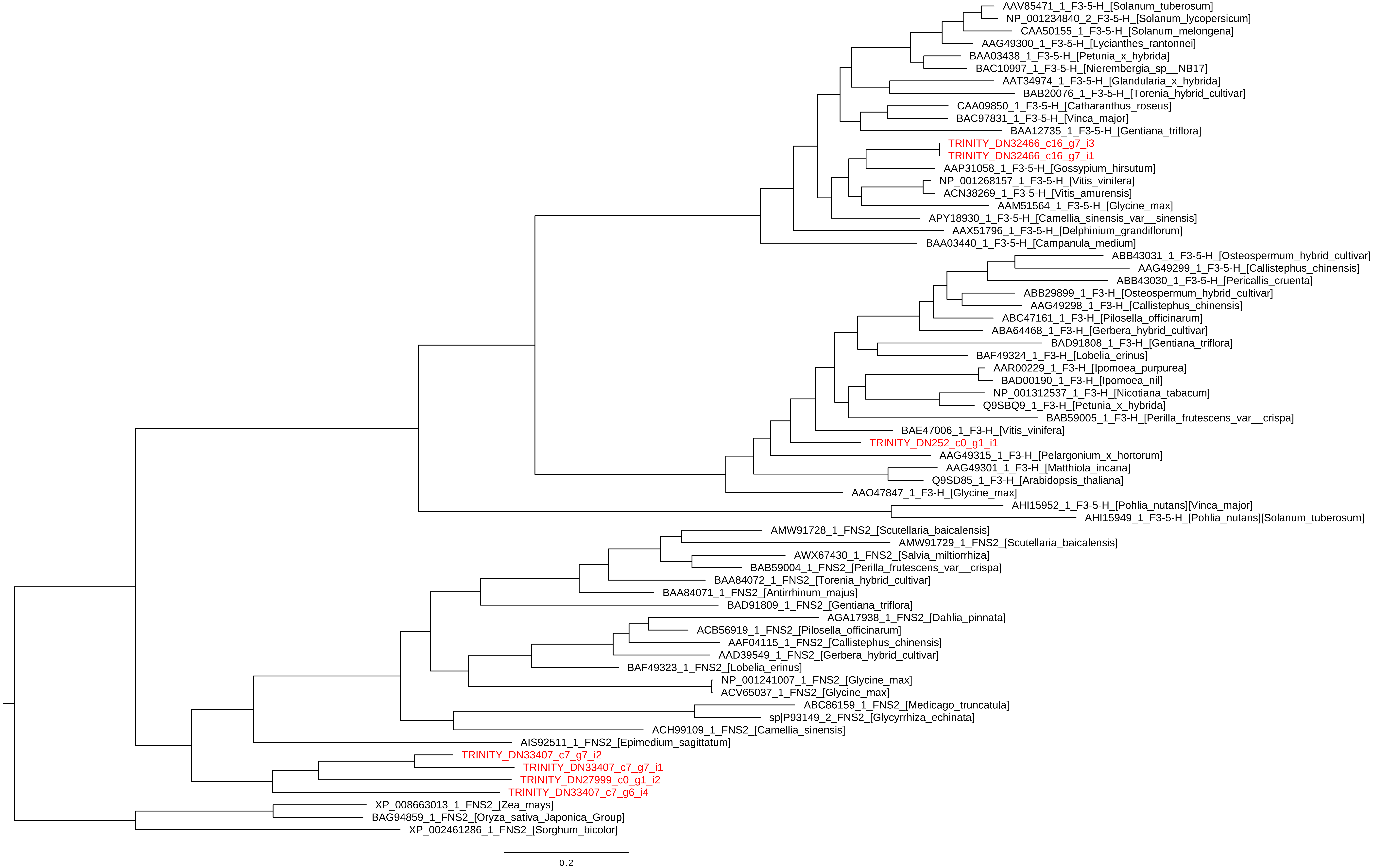

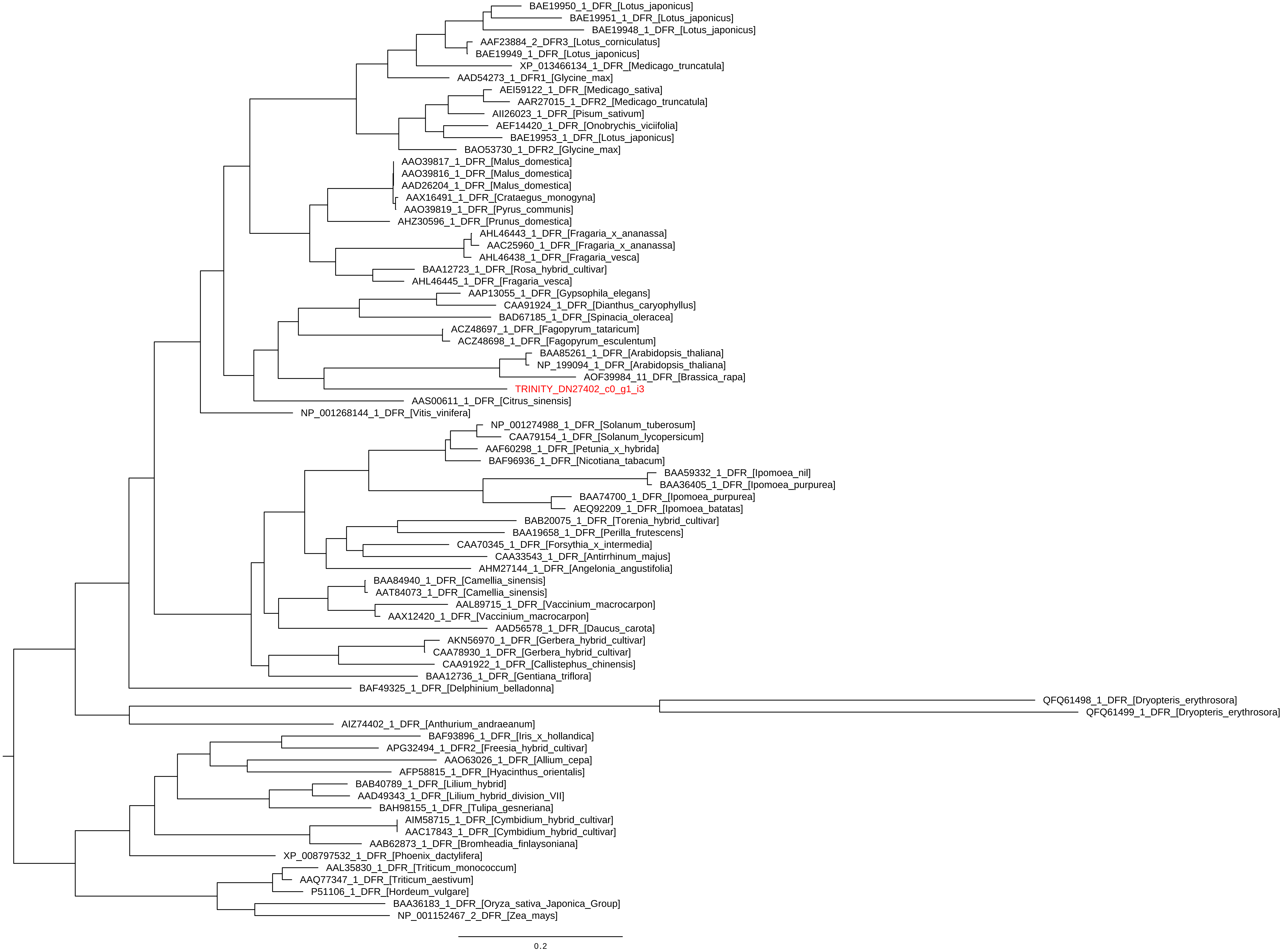

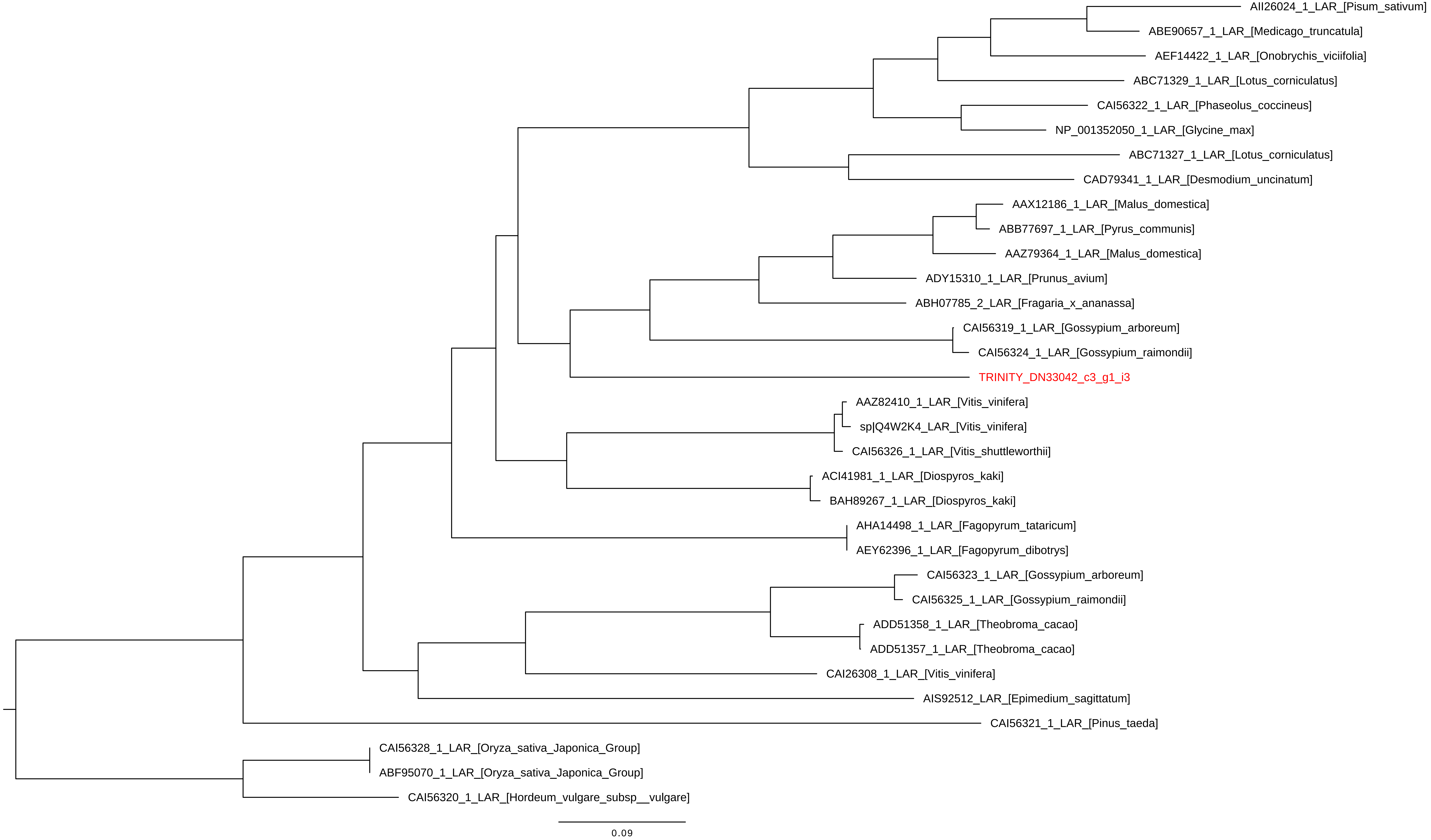

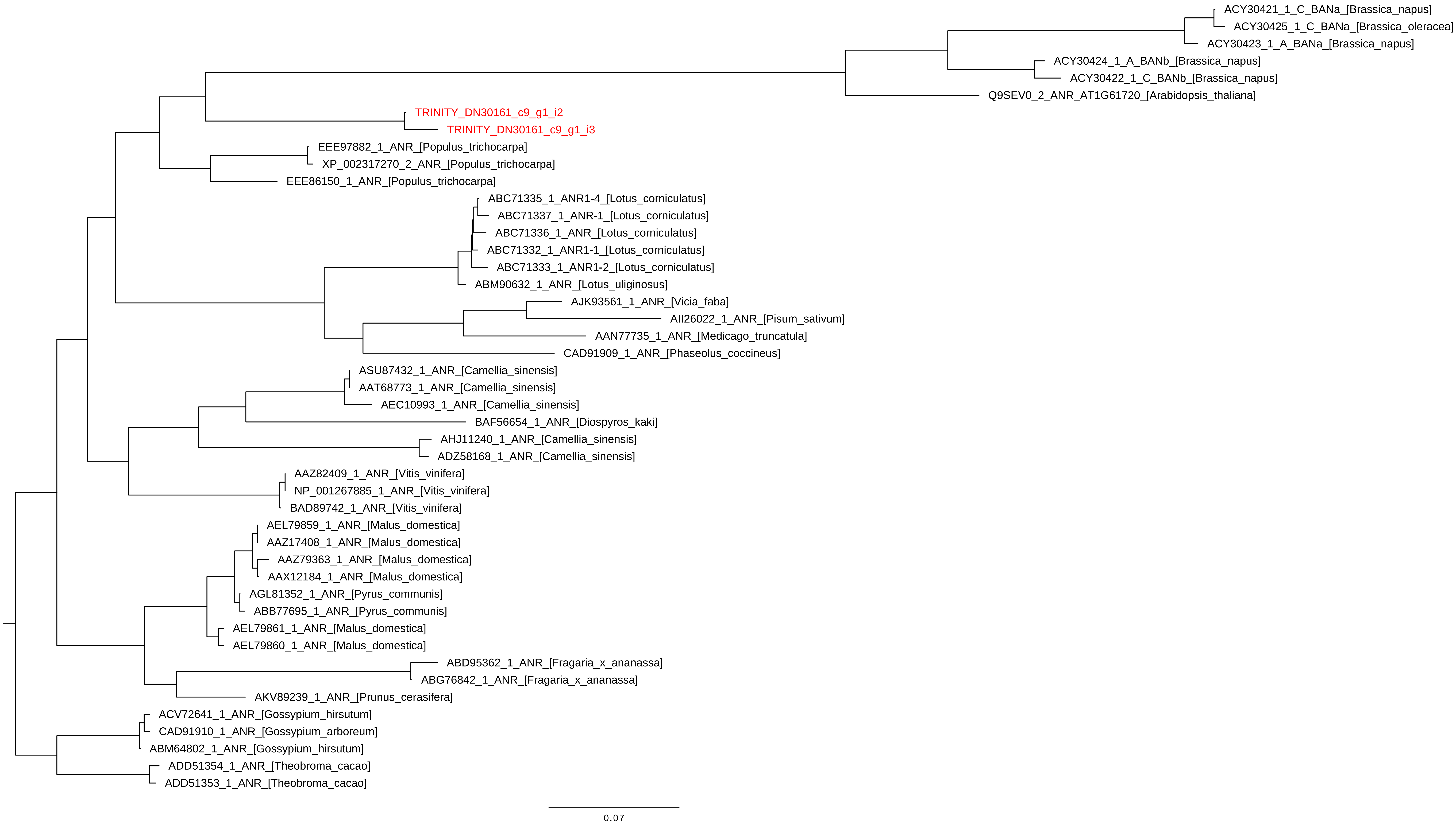

### File S7

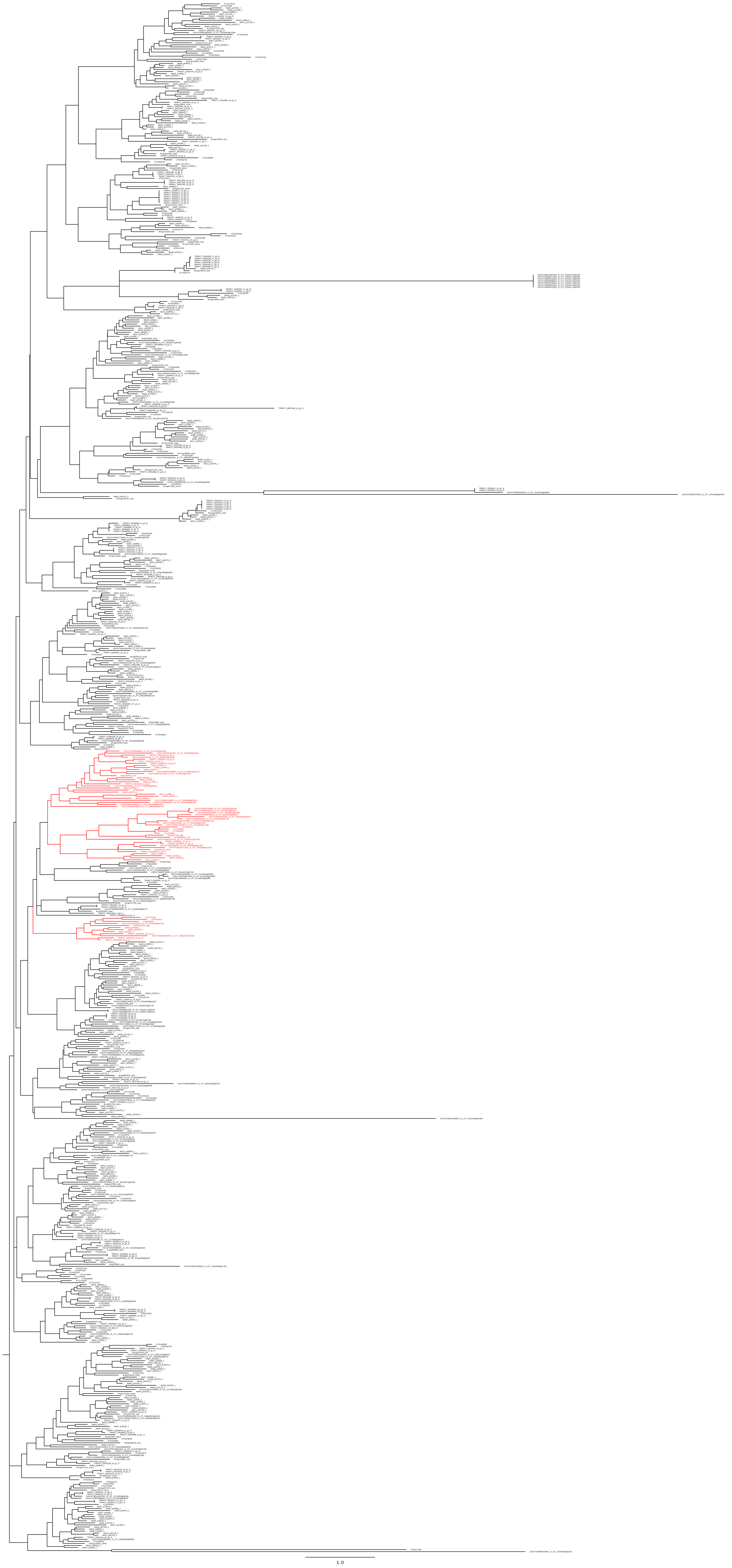

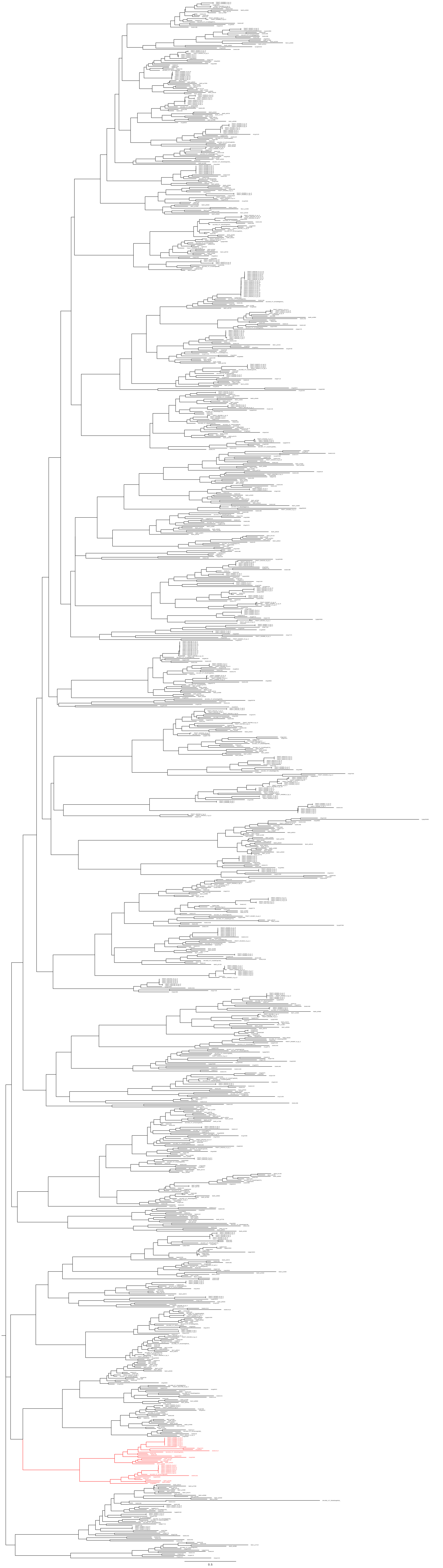

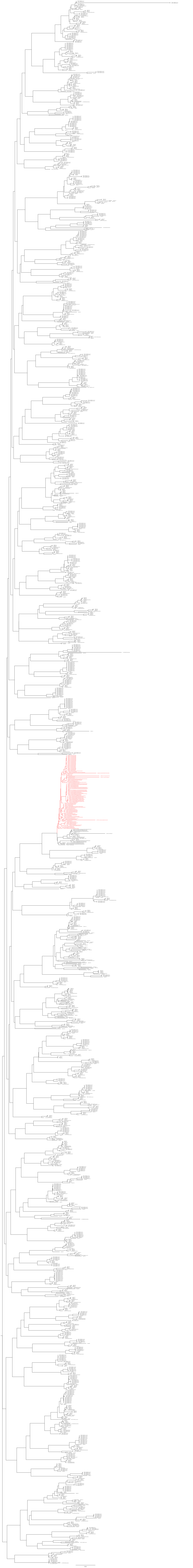
