## Supplementary material for "Automatic identification of players in the flavonoid biosynthesis with application on the biomedicinal plant *Croton tiglium*": File S3

```
                                                        η1
sp|P24481.1_PAL1_Petroselinum_crispum          ℓ....ℓℓ                             TT          T
                                             1          10                                   20
sp|P24481.1_PAL1_Petroselinum_crispum        ..ME.....N....GNGATTN...G....H....VN..GN.G..MD........FC.MKT
TRINITY_DN23351_c0_g1_i2                     ..ME.....F.CRKT....PN...G....N........AN.G...........WLG....
TRINITY_DN32981_c5_g1_i1                     ..MA.....K.I..........................GSFDD........LC.MT.
TRINITY_DN32981_c5_g1_i16                    ..MA.....K.I..........................GSFDD........LC.MT.
TRINITY_DN32981_c5_g1_i9                     ..MA.....K.I..........................GSFDD........LC.MT.
TRINITY_DN32981_c5_g1_i17                    ..MA.....K.I..........................GSFDD........LC.MT.
TRINITY_DN32981_c5_g1_i12                    ..MA.....K.I..........................GSFDD........LC.MT.
TRINITY_DN32981_c5_g1_i5                     ..MA.....K.I..........................GSFDD........LC.MT.
TRINITY_DN32981_c5_g1_i13                    ..MA.....K.I..........................GSFDD........LC.MT.
TRINITY_DN32981_c5_g1_i14                    ..MA.....K.I..........................GSFDD........LC.MT.
TRINITY_DN23351_c0_g2_i1                     ..MAFE............................................VC....
sp|P45728.1_PAL2_Petroselinum_crispum        ..ME.....N....GNGAITN...G....H....VN..GN.G..MD........FC.MKT
AWW24969.1_PAL_Lycoris_radiata               ..MA.....Y.A......NGN...G....NA..................NGFC.IL.
AIY24976.1_PAL_Mangifera_indica              ..ME.....F.CH.......D...N............RN.G...........GLC.T..
AHY94892.1_PAL_Prunella_vulgaris             ..MA.....A.........EN...G...HH......ASN.G............FC.VKQ
AHA42443.1_PAL_Cannabis_sativa               ..ME.....Q.........KN...D................VSLES........FC.VK.
AGL81344.1_PAL_Pyrus_communis                MEAE.....T.IT......QN...GKNGHH........QN.GAVES......P.LC.IK.
QIC54077.1_PAL_Paeonia_lactiflora            ..ME.....N.........GN...G....N...........GM..........SIC....
NP_181241.1_PAL1_Arabidopsis_thaliana        ..ME.....I..........N...GA...H.......KSNGGGVDA.......MLC.GGD
NP_190894.1_PAL2_Arabidopsis_thaliana        ..MD.....................................QIEA.......MLC.GGG
NP_187645.1_PAL4_Arabidopsis_thaliana        ..MELC............NQN.......NH.........................ITA
sp|Q42667.1_PAL_Citrus_limon                 ..ME.....L.SHETCNGIKN...D............RN.GGTSS......LGLC.TG.
CAB42793.1_PAL_Citrus_clementina_x_Citrus_r  ..MD.....RGAVI.....EN...G....H........QN.GCLEG........LC....
CAB42794.1_PAL_Citrus_clementina_x_Citrus_r  ..ME.....IGATT.....EN...G....H........QN.GGLEG........LC....
sp|P27991.1_PAL_Glycine_max                  ..ME.....A.........TN...G....H........QN.G...........SFC.LST
sp|O49835.1_PAL1_Lithospermum_erythrorhizon  ..ME.....T.IV......EN...G....N...........GKTME........FC....
sp|P35511.1_PAL1_Lycopersicon_esculentum     .........................................MD........LC.KKS
sp|P26600.1_PAL5_Lycopersicon_esculentum     ..MA.....SSIV......QN...G....H........VN.GEAMD........LC.KKS
sp|P25872.1_PAL1_Nicotiana_tabacum           ..MA.....S..........N...G....H........VNGGENFE........LC.KKS
sp|P35513.2_PAL2_Nicotiana_tabacum           ..MA.....G.VA......QN...G....H........QE....MD........FC.MK.
sp|P45733.1_PAL3_Nicotiana_tabacum           ..MA.....G.VA......QN...G....H........QE....MD........FC.VK.
sp|P19142.1_PAL2_Phaseolus_vulgaris          ..MD.....A.T.......PN...G....K........D.............AFV.VTA
sp|Q01861.1_PAL1_Pisum_sativum               ..ME.....T.VAAAITK.NN...G....Y........E.............SFC.VTN
sp|Q04593.1_PAL2_Pisum_sativum               ..ME.....A.IGAAITK.NNS..G....Y........D.............SFC.LTN
sp|P45731.1_PAL1_Populus_sieboldii_x_Populu  ............................................................
sp|P45730.1_PAL_Populus_trichocarpa          ..ME.....T.VT......KN...G....Y........QN.GSLES........LCVNQ.
sp|Q43052.1_PAL2_Populus_sieboldii_x_Populu  ..ME.....F.CQ.......D...S.............RN.GNGSP........GFN.T..
AAF40223.1_PAL1_Rubus_idaeus                 ..MEF.............YKN...G....N...........GAVES........FC....
AAF40224.1_PAL2_Rubus_idaeus                 ..ME.....S.IT......QN...G...HHH...QNGIQN.GSLDD......G.LC.IKT
AAC78457.1_PAL_Prunus_avium                  MATN.....S.IK......QN...G....H........KN.GSVEL......PELC.IK.
sp|P31425.1_PAL1_Solanum_tuberosum           ..MA.....PSIA......QN...G....H........VN.GEVEE......V.LW.KKS
sp|A2X7F7.1_PAL2_Oryza_sativa                ..ME..............CEN...G....RV......SAN.G.........MSGLC.VAA
sp|P52777.1_PAL_Pinus_taeda                  ..MV.....AAAEIT...QANEV......QV.......KSTG............LC.TDF
ABU49842.1_PAL_Ginkgo_biloba                 ..MV.....AGAERM...QSNPQNG..SQYV.......KSGGI.GD........LC.QSF
ACS28225.2_PAL_Pinus_massoniana              ..MV.....AAAEMT...QANEV......QV.......KSTG............LC.TDF
BAG74771.1_PAL_Ephedra_sinica                ..MV.....AGAEMA...QTAFV.....QHV.......KDGGIREF........LC.KGS
ABC69916.1_PAL_Brassica_napus                ..ME.....V..........N...GL............SHGGEVDA.......MLC.GGE
BAC56977.1_PAL_Daucus_carota                 ................MAYTN...G...HH....EN..GN.G..VD........LC.MKK
ABK24709.1_PAL_Picea_sitchensis              .MVA.....AAAEMA...QTNEV.....QQV.......KSTG............LC.TGL
AHA44840.1_PAL_Larix_kaempferi               ..MV.....AGAEIM...QTNEV.....QQV.......KSTG............LC.TSF
AGC23439.1_PAL_Dendrobium_candidum           ..ME.....H.V......KGN...G....YA........N.GAKAM......EGLC.LKG
ABI33979.1_PAL_Jatropha_curcas               ..MA.....T.II......GN...G....H........QN.GSLEG........LC.IT.
XP_002326186.1_PAL_Populus_trichocarpa       ..ME.....T.IT......KN...G....Y........QN.GSSES........LC.TQ.
XP_002519521.1_PAL_Ricinus_communis          ..MA.....A.MA......EN...G....S........KN.DSLES........FCNMG.
XP_002268732.1_PAL_Vitis_vinifera            ..MD.....A.T........N...C....H........GS.NKVKS........FC..V.
AFG26322.1_PAL_Cinnamomum_osmophloeum        ..ME.....T.T......QET...G....NV..................LQSFC.LK.
BAG70992.1_PAL_Musa_balbisiana               ..MEFAPKAQ.V......VEN...G.........................EAFC.LKA
AAP34199.1_PAL_Phalaenopsis_x_Doritaenopsis  ..ME.....V.S......KEN..............................GLC.LQG
ACM61988.1_PAL_Lycoris_radiata               ..MA.....Y.A......NGN...G....NA..................NGFC.IL.
ABN79671.2_PAL_Rudbeckia_hirta               ..ME.....N......GNHVN...G...NH....VN..GN.GAAAE........FC.IK.
ACJ66297.1_PAL4_Nicotiana_tabacum            ..MA.....S..........N...G....H........VNGGENFE........LC.KKS
ABG75911.1_PAL2_Nicotiana_attenuata          ..MA.....S..........N...G....H........VNGGEDFE........LC.KKS
BAL49995.1_PAL_Eucalyptus_robusta            MEME.....S.TTGT....GN...GL...H...........G...........LCATGS
ACM61988.1_PAL_Lycoris_radiata               ..MA.....Y.A......NGN...G....NA..................NGFC.IL.
BAM28963.1_PAL_Lilium_hybrid_division_I      ..MA.....H.IV...CENGN...G....FAATAADGNIN.GFIATAADGNSNGLC.LHA
                                       acc
```

```
                                                         α1            α2              β1        α3
sp|P24481.1_PAL1_Petroselinum_crispum         ...........T                         
                                                            30        40        50        60        70

sp|P24481.1_PAL1_Petroselinum_crispum         ............EDPLYWGIAAEAMTGSHLDEVKKMVAEY.RKPVVKLGGETLTISQVAA
TRINITY_DN23351_c0_g1_i2                      .......FDV..NDPLNWGAVAESLKGSHLDEVKRMVKEY.RKPVVKLGGETLTIAQVAS
TRINITY_DN32981_c5_g1_i1                      ............RDPLNWGVAAESMKGSHFDEVKQMVSDF.RSPLVKLGGETLTVAQVAA
TRINITY_DN32981_c5_g1_i16                     ............RDPLNWGVAAESMKGSHFDEVKQMVSDF.RSPLVKLGGETLTVAQVAA
TRINITY_DN32981_c5_g1_i9                      ............RDPLNWGVAAESMKGSHFDEVKQMVSDF.RSPLVKLGGETLTVAQVAA
TRINITY_DN32981_c5_g1_i17                     ............LDPLNWGVAAESMKGSHFDEVKQMVRDF.RSPLVKLGGETLTVAQVAA
TRINITY_DN32981_c5_g1_i12                     ............LDPLNWGVAAESMKGSHFDEVKQMVRDF.RSPLVKLGGETLTVAQVAA
TRINITY_DN32981_c5_g1_i5                      ............RDPLNWGVAAESMKGSHFDEVKQMVSDF.RSPLVKLGGETLTVAQVAA
TRINITY_DN32981_c5_g1_i13                     ............RDPLNWGVAAESMKGSHFDEVKQMVSDF.RSPLVKLGGETLTVAQVAA
TRINITY_DN32981_c5_g1_i14                     ............LDPLNWGVAAESMKGSHFDEVKQMVRDF.RSPLVKLGGETLTVAQVAA
TRINITY_DN23351_c0_g2_i1                      ............QTVSHWQKAAELLQSSHFDEVQEMVSQFANAITVDIKGTSLTVAEVTA
sp|P45728.1_PAL2_Petroselinum_crispum         ............EDPLYWGIAAEAMTGSHLDEVKKMVAEY.RKPVVKLGGETLTISQVAA
AWW24969.1_PAL_Lycoris_radiata                .............DPLNWGAAAEALTGSHLDEVKRMVKDY.REAYVRLEGATLKVAQIAA
AIY24976.1_PAL_Mangifera_indica               ............SDPLNWNFAAESLKGSHLDEVKRMVEEF.RKPVVKLGGETLTIGQVTA
AHY94892.1_PAL_Prunella_vulgaris              ............SDPLNWGAAAESMSGSHLQEVKRMVEEF.RKPVVRLGGETLTISQVAA
AHA42443.1_PAL_Cannabis_sativa                ............GDPLNWGLAAESMSGSHLDEVKRMVAEY.RKPVVGLGGETLTISQVAA
AGL81344.1_PAL_Pyrus_communis                 ............KDPLNWGLAADSLKGSHLDEVKRMVAEY.RKPVVKLGGESLTISQVAA
QIC54077.1_PAL_Paeonia_lactiflora             .........T..QDPLSWGVAAESLKGSHLDEVKRMVEEF.RKPVIRLAGKALTIAQVAG
NP_181241.1_PAL1_Arabidopsis_thaliana         IKTKNMVINA..EDPLNWGAAAEQMKGSHLDEVKRMVAEF.RKPVVNLGGETLTIGQVAA
NP_190894.1_PAL2_Arabidopsis_thaliana         EKTKVAVTTKTLADPLNWGLAADQMKGSHLDEVKKMVEEY.RRPVVNLGGETLTIGQVAA
NP_187645.1_PAL4_Arabidopsis_thaliana         VS..........GDPLNWNATAEALKGSHLDEVKRMVKEY.RKEAVKLGGETLTIGQVAA
sp|Q42667.1_PAL_Citrus_limon                  ............TDPLNWTVAADSLKGSHLDEVKRMIDEY.RRPVVKLGGESLTIGQVTA
CAB42793.1_PAL_Citrus_clementina_x_Citrus_r   .KDN..NYSS..GDALNWGVMAETLKGSHLEEVKRMVAEY.RKPVVNLGGETLTVAQVAA
CAB42794.1_PAL_Citrus_clementina_x_Citrus_r   .KNNNYNYSS..GDALNWGVMAETLKGSHLEEVKRMVAEY.RKPVVNLGGETLTVAQVAA
sp|P27991.1_PAL_Glycine_max                   AKGNN........DPLNWGAAAEAMKGSHLDEVKRMVAEY.RKPVVRLGGETLTIAQVAA
sp|O49835.1_PAL1_Lithospermum_erythrorhizon   .........M..KDPLNWEMASESMKGSHLDEVKNMVAEF.RKPVVQLAGKTLTIGQVAA
sp|P35511.1_PAL1_Lycopersicon_esculentum      I...........NDPLNWEMAADSLRGSHLDEVKKMVDEF.RKPIVKLGGETLSVAQVAS
sp|P26600.1_PAL5_Lycopersicon_esculentum      INV.........NDPLNWEMAAESLRGSHLDEVKKMVDEF.RKPIVKLGGETLTVAQVAS
sp|P25872.1_PAL1_Nicotiana_tabacum            A............DPLNWEMAAESLRGSHLDEVKKMVSEF.RKPMVKLGGESLTVAQVAA
sp|P35513.2_PAL2_Nicotiana_tabacum            ............VDPLNWEMAADSLKGSHLDEVKKMVAEF.RKPVVKLGGETLTVAQVAA
sp|P45733.1_PAL3_Nicotiana_tabacum            ............VDPLNWEMAADSLKGSHLDEVKKMVAEF.RKPVVKLGGETLTVAQVAA
sp|P19142.1_PAL2_Phaseolus_vulgaris           ANAAG........DPLNWAAAAEALSGSHLDEVKRMVAEY.RKPAVRLGGQTLTIAQVAA
sp|Q01861.1_PAL1_Pisum_sativum                AKNNNMKVNS..ADPLNWGVAAEAMKGSHLDEVKRMVEEY.RKPVVRLGGETLTISQVAA
sp|Q04593.1_PAL2_Pisum_sativum                AKNNNIKVSD..SDPLNWGVAAEAMKGSHLDEVKRMVDEY.RKPVVRLGGETLTISQVAA
sp|P45731.1_PAL1_Populus_sieboldii_x_Populu   ...................MAADSLKGSHLDEVKRMIEEY.RNPVVKLGGETLTIGQVTA
sp|P45730.1_PAL_Populus_trichocarpa           ............RDPLSWGVAAEAMKGSHLDEVKRMVADY.RKPVVKLGGETLTIAQVAS
sp|Q43052.1_PAL2_Populus_sieboldii_x_Populu   ............NDPLNWGMAAESLKGSHLDEVKRMIEEY.RNPVVKLGGETLTIGQVTA
AAF40223.1_PAL1_Rubus_idaeus                  ........EG..HDPLNWNMAAESLKGSHVDELKRMVSDY.RKPVVKLGGETLTIGQVAA
AAF40224.1_PAL2_Rubus_idaeus                  ESIKT.GYSV..SDPLNWGAAAESMTGSHLDEVRRMVAEY.RKPVVKLGGETLTISQVAA
AAC78457.1_PAL_Prunus_avium                   ............KDPLNWGVAAETLKGSHLDEVKRMVAEY.RKPVVKLGGESLTISQVAA
sp|P31425.1_PAL1_Solanum_tuberosum            I...........HDPLNWEMAVDSLRGSHLDEVKKMVDEF.RKPIVKLWGETLTVAQVAS
sp|A2X7F7.1_PAL2_Oryza_sativa                 PR..........ADPLNWGKATEEMTGSHLDEVKRMVAEY.RQPLVKIEGASLRIAQVAA
sp|P52777.1_PAL_Pinus_taeda                   GSSG........SDPLNWVRAAKAMEGSHFEEVKAMVDSYFGAKEISIEGKSLTISDVAA
ABU49842.1_PAL_Ginkgo_biloba                  .DST........TDPLNWARAAKALQGSHFEEVKQMVDSYFKSGEISIEGKTLTVADVTA
ACS28225.2_PAL_Pinus_massoniana               GSSG........SDPLNWVRAAKAMEGSHFEEVKAMVDSYLGVKEIFIEGKSLTISDVAA
BAG74771.1_PAL_Ephedra_sinica                 .DSS........NDPLNWVAAAKSMSGSHFDMVRDMVEVYLNAKEVSIEGKTLTVAQVTA
ABC69916.1_PAL_Brassica_napus                 IK.KNATVVG..ADPLNWGAAAEQMKGSHLDEVKRMVKEF.RRPVVNLGGETLTIGQVAA
BAC56977.1_PAL_Daucus_carota                  ............EDPLSWGVAAEALKGSHLEEVKRMVAEY.RKPVVKLGGETLTISQVAA
ABK24709.1_PAL_Picea_sitchensis               SSSS........SDPLNWIRAAKAMEGSHFEEVKTMVDSYFESKEIFIEGKTLTIADVTA
AHA44840.1_PAL_Larix_kaempferi                GSST........SDPLNWIQAAKAMEGSHFEEVKSMVNSYLGSKEISIEGKSLTISDVAA
AGC23439.1_PAL_Dendrobium_candidum            ............RDQLGWAAAAKALEGSHLDEVKRMVKEF.RSPVVRLEGAELKISQVAA
ABI33979.1_PAL_Jatropha_curcas                ............RDPLSWGVAAESMKGSHLDEVKKMVSEY.RKPLVKLGGETLTVAQVAA
XP_002326186.1_PAL_Populus_trichocarpa        ............RDPLSWGVAAEAMKGSHLDEVKRMVAEY.RKPVVNLAGQTLTIAQVAS
XP_002519521.1_PAL_Ricinus_communis           ............RDPLSWGLAAESMKGSHLDEVKKMVAEY.RKPFVKLGGETLTVAQVAA
XP_002268732.1_PAL_Vitis_vinifera             ............SDPLNWGMAAETLKGSHLDEVKRMVAEY.RKPVVRLGGETLTISQVAA
AFG26322.1_PAL_Cinnamomum_osmophloeum         .............DPLNWEMAAESLKGSHLEEVKRMVDEF.RKPLIQLEGATLTISQVAA
BAG70992.1_PAL_Musa_balbisiana                .............DPLNWIKAAESLTGSHLDEVKRMVEEF.RKPLVRLEGATLTISQVAA
AAP34199.1_PAL_Phalaenopsis_x_Doritaenopsis   ............HDPLNWGAAAAELQGSHLDEVKKMVEEF.RRPVVQLEGAKLKISQVAA
ACM61988.1_PAL_Lycoris_radiata                .............DPLNWGAAAEALTGSHLDEVKRMVKDY.REAYVKLEGATLKVAQIAA
ABN79671.2_PAL_Rudbeckia_hirta                .............DPLNWGAAAEALTGSHLDEVKKMVAEF.RKPVVKLGGETLTVSQVAG
ACJ66297.1_PAL4_Nicotiana_tabacum             SA..........TDPLNWEMAAESLRGSHLDEVKKMVSEF.RKPMVKLGGETLTVAQVAA
ABG75911.1_PAL2_Nicotiana_attenuata           SA..........TDPLNWEMASESLRGSHLDEVKKMVSEF.RKPMVKLGGETLTVAQVAA
BAL49995.1_PAL_Eucalyptus_robusta             RH..........ADPLNWGAAAAALTGSHLDEVKRMVEEY.RRPAVRLGGESLTIAQVAA
ACM61988.1_PAL_Lycoris_radiata                .............DPLNWGAAAEALTGSHLDEVKRMVKDY.REAYVKLEGATLKVAQIAA
BAM28963.1_PAL_Lilium_hybrid_division_I       .............DPLNWRAAAEAMSGSHLDEVRRMAAEF.RKPVVRLEGASLNISQVAA
                                        acc
```

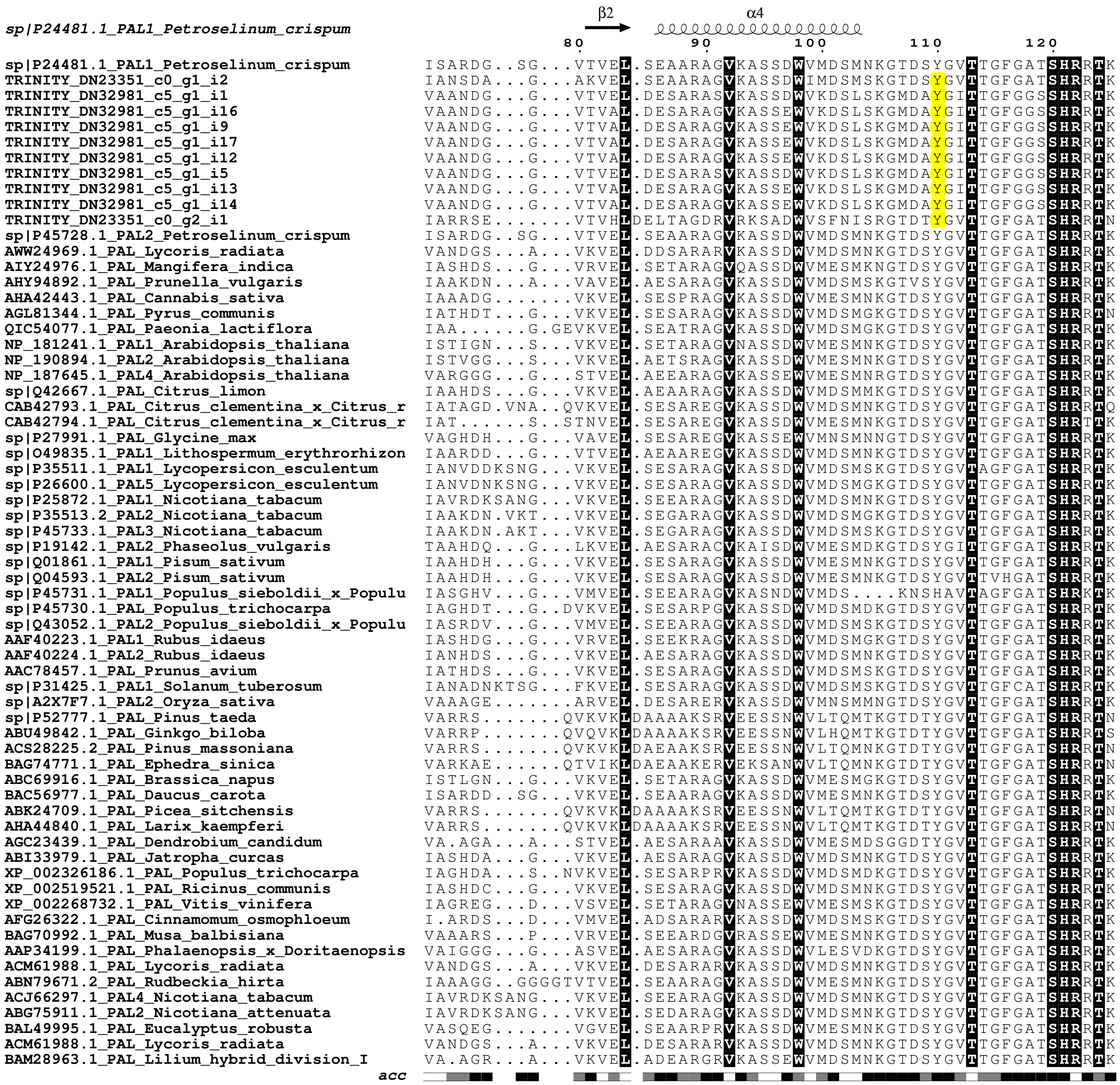
sp|P24481.1_PAL1_Petroselinum_crispum
β2
α4
80 90 100 110 120
sp|P24481.1_PAL1_Petroselinum_crispum ISARDG..SG...VTVEL.SEAARAGVKASSDWVMDSMNKGTDSYGVTTGFGATSHRRTK
TRINITY_DN23351_c0_g1_i2 IANSDA...G...AKVEL.SEAARAGVKASSDWIMDSMSKGTDSYGVTTGFGATSHRRTK
TRINITY_DN32981_c5_g1_i1 VAANDG...G...VTVEL.DESARASVKASSDWVKDSLSKGMDAYGITTGFGGSSHRRTK
TRINITY_DN32981_c5_g1_i16 VAANDG...G...VTVAL.DESARAGVKASSEWVKDSLSKGMDAYGITTGFGGSSHRRTK
TRINITY_DN32981_c5_g1_i9 VAANDG...G...VTVAL.DESARAGVKASSEWVKDSLSKGMDAYGITTGFGGSSHRRTK
TRINITY_DN32981_c5_g1_i17 VAANDG...G...VTVAL.DESARAGVKASSEWVKDSLSKGMDAYGITTGFGGSSHRRTK
TRINITY_DN32981_c5_g1_i12 VAANDG...G...VTVAL.DESARAGVKASSEWVKDSLSKGMDAYGITTGFGGSSHRRTK
TRINITY_DN32981_c5_g1_i5 VAANDG...G...VTVEL.DESARASVKASSDWVKDSLSKGMDAYGITTGFGGSSHRRTK
TRINITY_DN32981_c5_g1_i13 VAANDG...G...VTVAL.DESARAGVKASSEWVKDSLSKGMDAYGITTGFGGSSHRRTK
TRINITY_DN32981_c5_g1_i14 VAANDG...G...VTVAL.DESARAGVKASSEWVKDSLSKGMDAYGITTGFGGSSHRRTK
TRINITY_DN23351_c0_g2_i1 IARRSE......VTVHLDELTAGDRVRKSADWVSFNISRGTDTYGVTTGFGATSHRRTN
sp|P45728.1_PAL2_Petroselinum_crispum ISARDG..SG...VTVEL.SEAARAGVKASSDWVMDSMNKGTDSYGVTTGFGATSHRRTK
AWW24969.1_PAL_Lycoris_radiata VANDGS...A...VKVEL.DDSARARVKASSDWVMDSMNKGTDSYGVTTGFGATSHRRTK
AIY24976.1_PAL_Mangifera_indica IASHDS...G...VRVEL.SETARAGVQASSDWVMESMKNGTDSYGVTTGFGATSHRRTK
AHY94892.1_PAL_Prunella_vulgaris IAAKDN...A...VAVEL.AESARAGVKASSDWVMDSMSKGTVSYGVTTGFGATSHRRTK
AHA42443.1_PAL_Cannabis_sativa IAAADG......VKVEL.SESPRAGVKASSDWVMESMNKGTDSYGVTTGFGATSHRRTK
AGL81344.1_PAL_Pyrus_communis IATHDT...G...VKVEL.SESARAGVKASSDWVMDSMGKGTDSYGVTTGFGATSHRRTN
QIC54077.1_PAL_Paeonia_lactiflora IAA.....G.GEVKVEL.SEATRAGVKASSDWVMDSMGKGTDSYGVTTGFGATSHRRTK
NP_181241.1_PAL1_Arabidopsis_thaliana ISTIGN...S...VKVEL.SETARAGVNASSDWVMESMNKGTDSYGVTTGFGATSHRRTK
NP_190894.1_PAL2_Arabidopsis_thaliana ISTVGG...S...VKVEL.AETSRAGVKASSDWVMESMNKGTDSYGVTTGFGATSHRRTK
NP_187645.1_PAL4_Arabidopsis_thaliana VARGGG...G...STVEL.AEEARAGVKASSEWVMESMNRGTDSYGVTTGFGATSHRRTK
sp|Q42667.1_PAL_Citrus_limon IAAHDS...G...VKVEL.AEAARAGVKASSDWVMDSMMKGTDSYGVTTGFGATSHRRTK
CAB42793.1_PAL_Citrus_clementina_x_Citrus_r IATAGD.VNA..QVKVEL.SESAREGVKASSDWVMDSMNKGTDSYGVTTGFGATSHRRTQ
CAB42794.1_PAL_Citrus_clementina_x_Citrus_r IAT.....S..STNVEL.SESAREGVKASSDWVMESMNKGTDSYGVTTGFGATSHRTTK
sp|P27991.1_PAL_Glycine_max VAGHDH...G...VAVEL.SESAREGVKASSEWVMNSMNNGTDSYGVTTGFGATSHRRTK
sp|O49835.1_PAL1_Lithospermum_erythrorhizon IAARDD...G...VTVEL.AEAAREGVKASSDWVMDSMNKGTDSYGVTTGFGATSHRRTK
sp|P35511.1_PAL1_Lycopersicon_esculentum IANVDDKSNG...VKVEL.SESARAGVKASSDWVMDSMSKGTDSYGVTAGFGATSHRRTK
sp|P26600.1_PAL5_Lycopersicon_esculentum IANVDNKSNG...VKVEL.SESARAGVKASSDWVMDSMGKGTDSYGVTTGFGATSHRRTK
sp|P25872.1_PAL1_Nicotiana_tabacum IAVRDKSANG...VKVEL.SEEARAGVKASSDWVMDSMNKGTDSYGVTTGFGATSHRRTK
sp|P35513.2_PAL2_Nicotiana_tabacum IAAKDN.VKT...VKVEL.SEGARAGVKASSDWVMDSMGKGTDSYGVTTGFGATSHRRTK
sp|P45733.1_PAL3_Nicotiana_tabacum IAAKDN.AKT...VKVEL.SEGARAGVKASSDWVMDSMSKGTDSYGVTTGFGATSHRRTK
sp|P19142.1_PAL2_Phaseolus_vulgaris TAAHDQ...G...LKVEL.AESARACVKAISDWVMESMDKGTDSYGITTGFGATSHRRTK
sp|Q01861.1_PAL1_Pisum_sativum IAAHDH...G...VKVEL.SESARAGVKASSDWVMESMNKGTDSYGVTTGFGATSHRRTK
sp|Q04593.1_PAL2_Pisum_sativum IAAHDH...G...VKVEL.SESARAGVKASSDWVMESMNKGTDSYGVTTVHGATSHRRTK
sp|P45731.1_PAL1_Populus_sieboldii_x_Populu IASGHV...G...VMVEL.SEEARAGVKASNDWVMDS....KNSHAVTAGFGATSHRKTK
sp|P45730.1_PAL_Populus_trichocarpa IAGHDT...G..DVKVEL.SESARPGVKASSDWVMDSMDKGTDSYGVTTGFGATSHRRTK
sp|Q43052.1_PAL2_Populus_sieboldii_x_Populu IASRDV...G...VMVEL.SEEARAGVKASSDWVMDSMSKGTDSYGVTTGFGATSHRRTK
AAF40223.1_PAL1_Rubus_idaeus IASHDG...G...VRVEL.SEEKRAGVKASSDWVMDSMGKGTDSYGVTTGFGATSHRRTK
AAF40224.1_PAL2_Rubus_idaeus IANHDS...G...VKVEL.AESARAGVKASSDWVMDSMNKGTDSYGVTTGFGATSHRRTK
AAC78457.1_PAL_Prunus_avium IATHDS...G...VKVEL.SESARAGVKASSDWVMDSMSKGTDSYGVTTGFGATSHRRTK
sp|P31425.1_PAL1_Solanum_tuberosum IANADNKTSG...FKVEL.SESARAGVKASSDWVMDSMSKGTDSYGVTTGFCATSHRRTK
sp|A2X7F7.1_PAL2_Oryza_sativa VAAAGE......ARVEL.DESARERVKASSDWVMNSMMNGTDSYGVTTGFGATSHRRTK
sp|P52777.1_PAL_Pinus_taeda VARRS......QVKVKLDAAAAKSRVEESSNWVLTQMTKGTDTYGVTTGFGATSHRRTN
ABU49842.1_PAL_Ginkgo_biloba VARRP......QVQVKLDAAAAKSRVEESSNWVLHQMTKGTDTYGVTTGFGATSHRRTS
ACS28225.2_PAL_Pinus_massoniana VARRS......QVKVKLDAEAAKSRVEESSNWVLTQMNKGTDTYGVTTGFGATSHRRTN
BAG74771.1_PAL_Ephedra_sinica VARKAE.....QTVIKLDAEAAKERVEKSANWVLTQMNKGTDTYGVTTGFGATSHRRTN
ABC69916.1_PAL_Brassica_napus ISTLGN...G...VKVEL.SETARAGVKASSDWVMESMGKGTDSYGVTTGFGATSHRRTK
BAC56977.1_PAL_Daucus_carota ISARDD..SG...VKVEL.SEAARAGVKASSDWVMESMNKGTDSYGVTTGFGATSHRRTK
ABK24709.1_PAL_Picea_sitchensis VARRS......QVKVKLDAAAAKSRVEESSNWVLTQMTKGTDTYGVTTGFGATSHRRTN
AHA44840.1_PAL_Larix_kaempferi VARRS......QVKVKLDAAAAKSRVEESSNWVLTQMTKGTDTYGVTTGFGATSHRRTN
AGC23439.1_PAL_Dendrobium_candidum VA.AGA...A...STVEL.AESARAAVKASSDWVMESMDSGGDTYGVTTGFGATSHRRTK
ABI33979.1_PAL_Jatropha_curcas IASHDA...G...VKVEL.AESARAGVKASSDWVMDSMNKGTDSYGVTTGFGATSHRRTK
XP_002326186.1_PAL_Populus_trichocarpa IAGHDA...S..NVKVEL.SESARPRVKASSDWVMDSMDKGTDSYGVTTGFGATSHRRTK
XP_002519521.1_PAL_Ricinus_communis IASHDC...G...VKVEL.SESARAGVKASSDWVMDSMNKGTDSYGVTTGFGATSHRRTK
XP_002268732.1_PAL_Vitis_vinifera IAGREG...D...VSVEL.SETARAGVNASSEWVMESMSKGTDSYGVTTGFGATSHRRTK
AFG26322.1_PAL_Cinnamomum_osmophloeum I.ARDS...D...VMVEL.ADSARARVKASSDWVMDSMSKGTDSYGVTTGFGATSHRRTK
BAG70992.1_PAL_Musa_balbisiana VAAARS...P...VRVEL.SEEARDGVRASSEWVMESMNKGTDSYGVTTGFGATSHRRTK
AAP34199.1_PAL_Phalaenopsis_x_Doritaenopsis VAIGGG...G...ASVEL.AESARAGVKASSDWVLESVDKGTDSYGVTTGFGATSHRRTK
ACM61988.1_PAL_Lycoris_radiata VANDGS...A...VKVEL.DESARARVKASSDWVMDSMNKGTDSYGVTTGFGATSHRRTK
ABN79671.2_PAL_Rudbeckia_hirta IAAAGG..GGGGTVTVEL.SEAARAGVKASSDWVMESMNKGTDSYGVTTGFGATSHRRTK
ACJ66297.1_PAL4_Nicotiana_tabacum IAVRDKSANG...VKVEL.SEEARAGVKASSDWVMDSMNKGTDSYGVTTGFGATSHRRTK
ABG75911.1_PAL2_Nicotiana_attenuata IAVRDKSANG...VKVEL.SEDARAGVKASSDWVMDSMNKGTDSYGVTTGFGATSHRRTK
BAL49995.1_PAL_Eucalyptus_robusta VASQEG......VGVEL.SEAARPRVKASSDWVMESMNKGTDSYGVTTGFGATSHRRTK
ACM61988.1_PAL_Lycoris_radiata VANDGS...A...VKVEL.DESARARVKASSDWVMDSMNKGTDSYGVTTGFGATSHRRTK
BAM28963.1_PAL_Lilium_hybrid_division_I VA.AGR...A...VKVEL.ADEARGRVKASSEWVMDSMNKGTDSYGVTTGFGATSHRRTK
acc

```
                                            α5                          α6                 α7
sp|P24481.1_PAL1_Petroselinum_crispum                            TT                    TT
                                         130       140         150       160       170       180
sp|P24481.1_PAL1_Petroselinum_crispum    QGGALQKELIRFLNAGIFGNGS...DNTLPHSATRAAMLVRINTLLQGYSGIRFEILEAI
TRINITY_DN23351_c0_g1_i2                 QGGALQRELIRFFLNAGIFGNGAEN.SQILPHSATRAAMLVRINTLLQGYSGIRFEILEAI
TRINITY_DN32981_c5_g1_i1                 QGTALQNELLRHLNAGIFGNGIET.GHTLPVSATRAAMLVRINSNLQGYSGIRFEILEAI
TRINITY_DN32981_c5_g1_i16                QGTALQNELLRHLNAGIFGNGTET.GHTLPVSATRAAMLVRINSNLQGYSGIRFEILEAI
TRINITY_DN32981_c5_g1_i9                 QGTALQNELLRHLNAGIFGNGTET.GHTLPVSATRAAMLVRINSNLQGYSGIRFEILEAI
TRINITY_DN32981_c5_g1_i17                QGTALQNELLRHLNAGIFGNGTET.GHTLPVSATRAAMLVRINSNLQGYSGIRFEILEAI
TRINITY_DN32981_c5_g1_i12                QGTALQNELLRHLNAGIFGNGTET.GHTLPVSATRAAMLVRINSNLQGYSGIRFEILEAI
TRINITY_DN32981_c5_g1_i5                 QGTALQNELLRHLNAGIFGNGIET.GHTLPVSATRAAMLVRINSNLQGYSGIRFEILEAI
TRINITY_DN32981_c5_g1_i13                QGTALQNELLRHLNAGIFGNGTET.GHTLPVSATRAAMLVRINSNLQGYSGIRFEILEAI
TRINITY_DN32981_c5_g1_i14                QGTALQNELLRHLNAGIFGNGIET.GHTLPVSATRAAMLVRINSNLQGYSGIRFEILEAI
TRINITY_DN23351_c0_g2_i1                 KTADLQTELIRFLNAGVIG......KESLPLNYAKAAMLVRTNTLMQGYSGIRWEMLESI
sp|P45728.1_PAL2_Petroselinum_crispum    QGGALQKELIRFLNAGIFGNGS...DNTLPHSATRAAMLVRINTLLQGYSGIRFEILEAI
AWW24969.1_PAL_Lycoris_radiata           QGGALQKELIRFLNVGIFESGHNS.SNTLPASTTRAAMLVRINTLLQGYSGIRFEILEVI
AIY24976.1_PAL_Mangifera_indica          QGGALQKELIRFLNAGIFGSGTES.CHTLPHSATRAAMLVRINTLLQGYSGIRFDILEAI
AHY94892.1_PAL_Prunella_vulgaris         QGGALQKELIRFLNAGIFGNGTES.SHTLPHTATRAAMLVRINTLLQGYSGIRFEILEAI
AHA42443.1_PAL_Cannabis_sativa           NGGALQKELIRFLNAGIFGHGTES.CHTLPHSSTRAGLLVRINTLLQGYSGIRFEILEAM
AGL81344.1_PAL_Pyrus_communis            KGAALQKELIRFLNAGVFGSATES.GHTLPHQATRAAMLVRINTLLQGYSGIRFEILEAI
QIC54077.1_PAL_Paeonia_lactiflora        QGGALQNELIRFLNAGIFGNGRET.CHTLPHTTTRAAMLVRINTLLQGYSGIRFEILEAL
NP_181241.1_PAL1_Arabidopsis_thaliana    NGVALQKELIRFLNAGIFGSTKET.SHTLPHSATRAAMLVRINTLLQGFSGIRFEILEAI
NP_190894.1_PAL2_Arabidopsis_thaliana    NGTALQTELIRFLNAGIFGNTKET.CHTLPQSATRAAMLVRVNTLLQGYSGIRFEILEAI
NP_187645.1_PAL4_Arabidopsis_thaliana    QGGALQNELIRFLNAGIFGPGAGDTSHTLPKPTTRAAMLVRVNTLLQGYSGIRFEILEAI
sp|Q42667.1_PAL_Citrus_limon             QGGALQKELIRFLNSGIFGNGTES.SHTLPHSATRAAMLVRVNTLLQGYSGIRFEILETI
CAB42793.1_PAL_Citrus_clementina_x_Citrus_r NGGALQKELIKFLNAGIFGNGTKS.SHTLPHSATRAAMLVRVNTLLQGYSGIRFEILDAI
CAB42794.1_PAL_Citrus_clementina_x_Citrus_r NGGALQKELIKFLNAGIFGNGTKS.SHTLPHSATRAAMLVRVNTLLQGYSGIRFEILKAI
sp|P27991.1_PAL_Glycine_max              QGGALQKELIRFLNAGIFGNGTES.SHTLPHTATRAAMLVRINTLLQGYSGIRFEILEAI
sp|O49835.1_PAL1_Lithospermum_erythrorhizon QGGALQKELIRFLNAGIFGNGTET.SHTLPHSATRAAMLVRINTLLQGYSGIRFEILEAI
sp|P35511.1_PAL1_Lycopersicon_esculentum NGGALQKELIRFLNAGVFGNGIES.FHTLPHSATRAAMLVRINTLLQGYSGIRFEILEAI
sp|P26600.1_PAL5_Lycopersicon_esculentum NGGALQKELIRFLNAGVFGNGTES.SHTLPHSATRAAMLVRINTLLQGYSGIRFEILEAI
sp|P25872.1_PAL1_Nicotiana_tabacum       NGGALQKELIRFLNAGVFGNGTET.SHTLPHSATRAAMLVRINTLLQGYSGIRFEILEAI
sp|P35513.2_PAL2_Nicotiana_tabacum       NGGALQKELIRFLNAGVFGNGTES.CHTLPQSGTRAAMLVRINTLLQGYSGIRFEILEAI
sp|P45733.1_PAL3_Nicotiana_tabacum       NGGALQKELIRFLNAGVFGNGTES.CHTLPQSGTRAAMLVRINTLLQGYSGIRFEILEAI
sp|P19142.1_PAL2_Phaseolus_vulgaris      QGGALQKELIRFLNAGIFGNGTES.NCTLPHTATRAAMLVRVNTLLQGYSGIRFEILEAI
sp|Q01861.1_PAL1_Pisum_sativum           QGGALQKELIRFLNAGIFGNGTES.SHTLPHTATRAAMLVRINTLLQGYSGIRFEILEAI
sp|Q04593.1_PAL2_Pisum_sativum           QGGALQKELIRFLNAGIFGNGSES.THTLPHTATRAAMLVRINTLLQGYSGIRFEILEAI
sp|P45731.1_PAL1_Populus_sieboldii_x_Populu QGGELQKELIRFLNVGIFGNGTES.NHILPRSATRAAMLVRTNTLLQGYSGIRFEMLEAI
sp|P45730.1_PAL_Populus_trichocarpa      QGGALQKELIRFLNAGIFGNGTET.CHTLPHSATRAAMLVRINTLLQGYSGIRFEILEAI
sp|Q43052.1_PAL2_Populus_sieboldii_x_Populu QGGELQKELIRFLNAGIFGNGTES.SHTLPRSATRAAMLVRTNTLLQGYSGIRFEMLEAI
AAF40223.1_PAL1_Rubus_idaeus             NGGALQRELIRFLNAGIFGSSLDS.THKLPHTATRAAMLVRFNTLLQGYSGIRFEILEAI
AAF40224.1_PAL2_Rubus_idaeus             QGAALQKELIRFLNAGVLRNGTES.AHTLPHSATRAAMLVRINTLLQGYSGIRFEILEAI
AAC78457.1_PAL_Prunus_avium              QGAALQKELIRFLNAGVFGSTKES.GHTLPHQATRAAMLVRINTLLQGYSGIRFEILEVI
sp|P31425.1_PAL1_Solanum_tuberosum       NGGALQKELIKFLNAGVFGNGTES.THTLPHSATRAAMLVRINTLLQGYSGIRFEILEAI
sp|A2X7F7.1_PAL2_Oryza_sativa            EGGALQRELIRFLNAGAFGTGTD..GHVLPAEATRAAMLVRINTLLQGYSGIRFEILEAI
sp|P52777.1_PAL_Pinus_taeda              QGAELQKELIRFLNAGVLGKCPEN...VLSEDTTRAAMLVRTNTLLQGYSGVRWDILETV
ABU49842.1_PAL_Ginkgo_biloba             QGVELQKELIRFLNAGVFGSCEGN...VLPEATTRAAMLVRTNTLPQGYSGIRWALLETI
ACS28225.2_PAL_Pinus_massoniana          QGAELQKELIRFLNAGVLGKCPEN...VLSEDTTRAAMLVRTNTLLQGYSGIRWDILETV
BAG74771.1_PAL_Ephedra_sinica            QGAKLQKELIRFLNAGVLG.CDDN...VLPVETTRAAMLVRTNTLMQGYSGIRWEILAAV
ABC69916.1_PAL_Brassica_napus            NGVALQKELIRFLNAGIFGSTKET.SHTLPHSATRAAMLVRINTLLQGYSGIRFEILEAI
BAC56977.1_PAL_Daucus_carota             QGGALQKELIRFLNAGIFGSGAEAGNNTLPHSATRAAMLVRINTLLQGYSGIRFEILEAI
ABK24709.1_PAL_Picea_sitchensis          QGAELQKELIRFLNAGVLGKCQDN...VLSEDTTRAAMLVRTNTLLQGYSGIRWDILETV
AHA44840.1_PAL_Larix_kaempferi           QGAELQKELIRFLNAGVLGKCQDN...ILSEDTTRAAMLVRTNTLLQGYSGIRWDILETV
AGC23439.1_PAL_Dendrobium_candidum       QGGALQKELIRFLNAGIFGSGKD...NTLPPAATRAAMLVRINTLLQGYSGIRFEILEAI
ABI33979.1_PAL_Jatropha_curcas           QGAALQRELIRFLNAGIFGNGTET.CHTLPHSATRAAMLVRINTLLQGYSGIRFEILEAI
XP_002326186.1_PAL_Populus_trichocarpa   QGGALQKELIRFLNAGIFGNGTET.CHTLPHSATRAAMLVRINTLLQGYSGIRFEILEAI
XP_002519521.1_PAL_Ricinus_communis      QGGALQKELIRFLNAGIFGNGTES.CHTLPHSATRAAMLVRINTLLQGYSGIRFEILEAI
XP_002268732.1_PAL_Vitis_vinifera        QGGALQKELIRFLNAGIFGNGRES.CHTLPHSATRAAMLVRINTLLQGYSGIRFEILEAI
AFG26322.1_PAL_Cinnamomum_osmophloeum    QGGALQKELIRFLNAGVFGGAGES.GHTLPPSATRAAMLVRINTLMQGYSGIRFEILEAI
BAG70992.1_PAL_Musa_balbisiana           QGGALQKELIRFLNAGIFGSGPES.GNTLPSSAAKAAMLVRVNTLLQGYSGIRFEILEAI
AAP34199.1_PAL_Phalaenopsis_x_Doritaenopsis QGGALQKELIRFLNAGIFGSGNS...NTLPSSATRAAMLVRINTLLQGYSGIRFEILEAI
ACM61988.1_PAL_Lycoris_radiata           QGGALQKELIRFLNAGIFESGHNS.SNTLPASTTRAAMLVRINTLLQGYSGIRFEILEVI
ABN79671.2_PAL_Rudbeckia_hirta           NGGALQKELIRFLNAGIFGNGTES.SHTLPHSATRAAMLVRINTLLQGYSGIRFEILEAI
ACJ66297.1_PAL4_Nicotiana_tabacum        NGGALQKELIRFLNAGVFGNGTET.SHTLPHSATRAAMLVRINTLLQGYSGIRFEILEAI
ABG75911.1_PAL2_Nicotiana_attenuata      NGGALQKELIRFLNAGVFGNGTET.NHTLPHSATRAAMLVRINTLLQGYSGIRFEILEAI
BAL49995.1_PAL_Eucalyptus_robusta        QGGALQKELIRFLNAGIFGNGTES.CHTLPQSSTRAAMLVRVNTLLQGYSGIRFEILEAI
ACM61988.1_PAL_Lycoris_radiata           QGGALQKELIRFLNAGIFESGHNS.SNTLPASTTRAAMLVRINTLLQGYSGIRFEILEVI
BAM28963.1_PAL_Lilium_hybrid_division_I  QGGALQKELIRFLNAGIFGAGPEA.GQTLPPTTTRAAMLVRVNTLLQGYSGIRFEILEAV
                                    acc
```

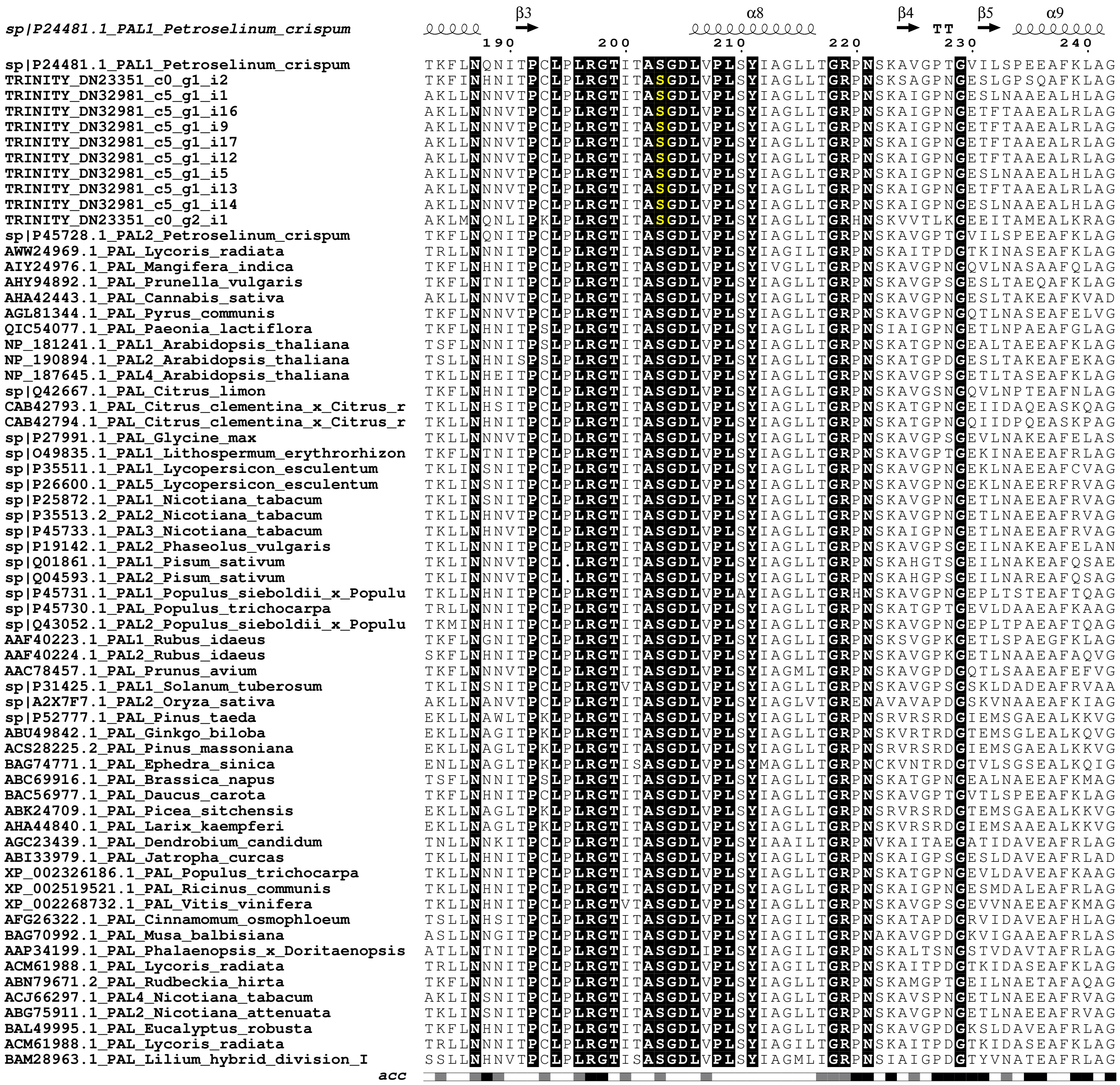
sp|P24481.1_PAL1_Petroselinum_crispum
β3 α8 β4 TT β5 α9
190 200 210 220 230 240
sp|P24481.1_PAL1_Petroselinum_crispum TKFLNQNITPCLPLRGTITASGDLVPLSYIAGLLTGRPNSKAVGPTGVILSPEEAFKLAG
TRINITY_DN23351_c0_g1_i2 TKFINHNITPCLPLRGTITASGDLVPLSYIAGLLTGRPNSKSAGPNGESLGPSQAFKLAG
TRINITY_DN32981_c5_g1_i1 AKLLNNNVTPCLPLRGTITASGDLVPLSYIAGLLTGRPNSKAIGPNGESLNAAEALHLAG
TRINITY_DN32981_c5_g1_i16 AKLLNNNVTPCLPLRGTITASGDLVPLSYIAGLLTGRPNSKAIGPNGETFTAAEALRLAG
TRINITY_DN32981_c5_g1_i9 AKLLNNNVTPCLPLRGTITASGDLVPLSYIAGLLTGRPNSKAIGPNGETFTAAEALRLAG
TRINITY_DN32981_c5_g1_i17 AKLLNNNVTPCLPLRGTITASGDLVPLSYIAGLLTGRPNSKAIGPNGETFTAAEALRLAG
TRINITY_DN32981_c5_g1_i12 AKLLNNNVTPCLPLRGTITASGDLVPLSYIAGLLTGRPNSKAIGPNGETFTAAEALRLAG
TRINITY_DN32981_c5_g1_i5 AKLLNNNVTPCLPLRGTITASGDLVPLSYIAGLLTGRPNSKAIGPNGESLNAAEALHLAG
TRINITY_DN32981_c5_g1_i13 AKLLNNNVTPCLPLRGTITASGDLVPLSYIAGLLTGRPNSKAIGPNGETFTAAEALRLAG
TRINITY_DN32981_c5_g1_i14 AKLLNNNVTPCLPLRGTITASGDLVPLSYIAGLLTGRPNSKAIGPNGESLNAAEALHLAG
TRINITY_DN23351_c0_g2_i1 AKLMNQNLIPKLPLRGTITASGDLVPLSYIAGLLTGRHNSKVVTLKGEEITAMEALKRAG
sp|P45728.1_PAL2_Petroselinum_crispum TKFLNQNITPCLPLRGTITASGDLVPLSYIAGLLTGRPNSKAVGPTGVILSPEEAFKLAG
AWW24969.1_PAL_Lycoris_radiata TRLLNNNITPCLPLRGTITASGDLVPLSYIAGILTGRPNSKAITPDGTKINASEAFKLAG
AIY24976.1_PAL_Mangifera_indica TKFLNHNITPCLPLRGTITASGDLVPLSYIVGLLTGRPNSKAVGPNGQVLNASAAFQLAG
AHY94892.1_PAL_Prunella_vulgaris TKFLNTNITPCLPLRGTITASGDLVPLSYIAGLLTGRPNSKAVGPSGESLTAEQAFKLAG
AHA42443.1_PAL_Cannabis_sativa AKLLNNNVTPCLPLRGTITASGDLVPLSYIAGLLTGRPNSKAVGPNGESLTAKEAFKVAD
AGL81344.1_PAL_Pyrus_communis TKFLNNNVTPCLPLRGTITASGDLVPLSYIAGLLTGRPNSKAVGPNGQTLNASEAFELVG
QIC54077.1_PAL_Paeonia_lactiflora TKFLNHNITPSLPLRGTITASGDLVPLSYIAGLLIGRPNSIAIGPNGETLNPAEAFGLAG
NP_181241.1_PAL1_Arabidopsis_thaliana TSFLNNNITPSLPLRGTITASGDLVPLSYIAGLLTGRPNSKATGPNGEALTAEEAFKLAG
NP_190894.1_PAL2_Arabidopsis_thaliana TSLLNHNISPSLPLRGTITASGDLVPLSYIAGLLTGRPNSKATGPDGESLTAKEAFEKAG
NP_187645.1_PAL4_Arabidopsis_thaliana TKLLNHEITPCLPLRGTITASGDLVPLSYIAGLLTGRPNSKAVGPSGETLTASEAFKLAG
sp|Q42667.1_PAL_Citrus_limon TKFLNHNITPCLPLRGTITASGDLVPLSYIAGLLTGRPNSKAVGSNGQVLNPTEAFNLAG
CAB42793.1_PAL_Citrus_clementina_x_Citrus_r TKLLNHSITPCLPLRGTITASGDLVPLSYIAGLLTGRPNSKATGPNGEIIDAQEASKQAG
CAB42794.1_PAL_Citrus_clementina_x_Citrus_r TKLLNHNITPCLPLRGTITASGDLVPLSYIAGLLTGRPNSKATGPNGQIIDPQEASKPAG
sp|P27991.1_PAL_Glycine_max TKLLNNNVTPCLDLRGTITASGDLVPLSYIAGLLTGRPNSKAVGPSGEVLNAKEAFELAS
sp|O49835.1_PAL1_Lithospermum_erythrorhizon TKFLNTNITPCLPLRGTITASGDLVPLSYIAGLLTGRPNSKAVGPTGEKINAEEAFRLAG
sp|P35511.1_PAL1_Lycopersicon_esculentum TKLINSNITPCLPLRGTITASGDLVPLSYIAGLLTGRPNSKAVGPNGEKLNAEEAFCVAG
sp|P26600.1_PAL5_Lycopersicon_esculentum TKLINSNITPCLPLRGTITASGDLVPLSYIAGLLTGRPNSKAVGPNGEKLNAEERFRVAG
sp|P25872.1_PAL1_Nicotiana_tabacum TKLINSNITPCLPLRGTITASGDLVPLSYIAGLLTGRPNSKAVGPNGETLNAEEAFRVAG
sp|P35513.2_PAL2_Nicotiana_tabacum TKLLNHNVTPCLPLRGTITASGDLVPLSYIAGLLTGRPNSKAVGPNGETLNAEEAFRVAG
sp|P45733.1_PAL3_Nicotiana_tabacum TKLLNHNVTPCLPLRGTITASGDLVPLSYIAGLLTGRPNSKAIGPNGETLNAEEAFRVAG
sp|P19142.1_PAL2_Phaseolus_vulgaris TKLLNNNITPCLPLRGTITASGDLVPLSYIAGLLTGRPNSKAVGPSGEILNAKEAFELAN
sp|Q01861.1_PAL1_Pisum_sativum TKLINNNVTPCL.LRGTITASGDLVPLSYIAGLLTGRPNSKAHGTSGEILNAKEAFQSAE
sp|Q04593.1_PAL2_Pisum_sativum TKLINNNVTPCL.LRGTITASGDLVPLSYIAGLLTGRPNSKAHGPSGEILNAREAFQSAG
sp|P45731.1_PAL1_Populus_sieboldii_x_Populu TKLLNHNITPCLPLRGTITASGDLVPLAYIAGLLTGRHNSKAVGPNGEPLTSTEAFTQAG
sp|P45730.1_PAL_Populus_trichocarpa TRLLNNNITPCLPLRGTITASGDLVPLSYIAGLLTGRPNSKATGPTGEVLDAAEAFKAAG
sp|Q43052.1_PAL2_Populus_sieboldii_x_Populu TKMINHNITPCLPLRGTITASGDLVPLSYIAGLLTGRPNSKAVGPNGEPLTPAEAFTQAG
AAF40223.1_PAL1_Rubus_idaeus TKFLNGNITPCLPLRGTITASGDLVPLSYIAGLLIGRPNSKSVGPKGETLSPAEGFKLAG
AAF40224.1_PAL2_Rubus_idaeus SKFLNHNITPCLPLRGTITASGDLVPLSYIAGLLTGRPNSKAVGPKGETLNAAEAFAQVG
AAC78457.1_PAL_Prunus_avium TKFLNNNVTPCLPLRGTITASGDLVPLSYIAGMLTGRPNSKAVGPDGQTLSAAEAFEFVG
sp|P31425.1_PAL1_Solanum_tuberosum TKLINSNITPCLPLRGTVTASGDLVPLSYIAGLLTGRPNSKAVGPSGSKLDADEAFRVAA
sp|A2X7F7.1_PAL2_Oryza_sativa AKLLNANVTPCLPLRGTITASGDLVPLSYIAGLVTGRENAVAVAPDGSKVNAAEAFKIAG
sp|P52777.1_PAL_Pinus_taeda EKLLNAWLTPKLPLRGTITASGDLVPLSYIAGLLTGRPNSRVRSRDGIEMSGAEALKKVG
ABU49842.1_PAL_Ginkgo_biloba EKLLNAGITPKLPLRGTITASGDLVPLSYIAGLLTGRPNSKVRTRDGTEMSGLEALKQVG
ACS28225.2_PAL_Pinus_massoniana EKLLNAGLTPKLPLRGTITASGDLVPLSYIAGLLTGRPNSRVRSRDGIEMSGAEALKKVG
BAG74771.1_PAL_Ephedra_sinica ENLLNAGLTPKLPLRGTISASGDLVPLSYMAGLLTGRPNCKVNTRDGTVLSGSEALKQIG
ABC69916.1_PAL_Brassica_napus TSFLNNNITPSLPLRGTITASGDLVPLSYIAGLLTGRPNSKATGPNGEALNAEEAFKMAG
BAC56977.1_PAL_Daucus_carota TKFLNHNITPCLPLRGTITASGDLVPLSYIAGLLTGRPNSKAVGPTGVTLSPEEAFKLAG
ABK24709.1_PAL_Picea_sitchensis EKLLNAGLTPKLPLRGTITASGDLVPLSYIAGLLTGRPNSRVRSRDGTEMSGAEALKKVG
AHA44840.1_PAL_Larix_kaempferi EKLLNAGLTPKLPLRGTITASGDLVPLSYIAGLLTGRPNSKVRSRDGIEMSAAEALKKVG
AGC23439.1_PAL_Dendrobium_candidum TNLLNNKITPCLPLRGTITASGDLVPLSYIAAILTGRPNVKAITAEGATIDAVEAFRLAG
ABI33979.1_PAL_Jatropha_curcas TKLLNHNITPCLPLRGTITASGDLVPLSYIAGLLTGRPNSKAIGPSGESLDAVEAFRLAD
XP_002326186.1_PAL_Populus_trichocarpa TKLLNNNITPCLPLRGTITASGDLVPLSYIAGLLTGRPNSKATGPNGEVLDAVEAFKAAG
XP_002519521.1_PAL_Ricinus_communis TKLLNHNITPCLPLRGTITASGDLVPLSYIAGLLTGRPNSKAIGPNGESMDALEAFRLAG
XP_002268732.1_PAL_Vitis_vinifera TKLLNHNITPCLPLRGTVTASGDLVPLSYIAGLLTGRPNSKAVGPSGEVVNAEEAFKMAG
AFG26322.1_PAL_Cinnamomum_osmophloeum TSLLNHSITPCLPLRGTITASGDLVPLSYIAGILTGRPNSKATAPDGRVIDAVEAFHLAG
BAG70992.1_PAL_Musa_balbisiana ASLLNNGITPCLPLRGTITASGDLVPLSYIAGILTGRPNAKAVGPDGKVIGAAEAFRLAS
AAP34199.1_PAL_Phalaenopsis_x_Doritaenopsis ATLLNTNITPCLPLRGTITASGDLIPLSYIAGILTGRPNSKALTSNGSTVDAVTAFRLAG
ACM61988.1_PAL_Lycoris_radiata TRLLNNNITPCLPLRGTITASGDLVPLSYIAGILTGRPNSKAITPDGTKIDASEAFKLAG
ABN79671.2_PAL_Rudbeckia_hirta TKFLNNNITPCLPLRGTITASGDLVPLSYIAGLLTGRPNSKAMGPTGEILNAETAFAQAG
ACJ66297.1_PAL4_Nicotiana_tabacum AKLINSNITPCLPLRGTITASGDLVPLSYIAGLLTGRPNSKAVSPNGETLNAEEAFRVAG
ABG75911.1_PAL2_Nicotiana_attenuata TKLINSNITPCLPLRGTITASGDLVPLSYIAGLLTGRPNSKAVGPNGETLNAEEAFRVAG
BAL49995.1_PAL_Eucalyptus_robusta TKFLNHNITPCLPLRGTITASGDLVPLSYIAGLLTGRPNSKAVGPDGKSLDAVEAFRLAG
ACM61988.1_PAL_Lycoris_radiata TRLLNNNITPCLPLRGTITASGDLVPLSYIAGILTGRPNSKAITPDGTKIDASEAFKLAG
BAM28963.1_PAL_Lilium_hybrid_division_I SSLLNHNVTPCLPLRGTISASGDLVPLSYIAGMLIGRPNSIAIGPDGTYVNATEAFRLAG
acc

```
                                                     α10              α11                    η2
sp|P24481.1_PAL1_Petroselinum_crispum
                                              250       260       270       280       290       300
sp|P24481.1_PAL1_Petroselinum_crispum        VEGGFFELQPKEGLALVNGTAVGSGMASMVLFEANILAVLAEVMSAIFAEVMQGKPEFTD
TRINITY_DN23351_c0_g1_i2                     IVTGFFELQPKEGLALVNGTAVGSGLASMVLFEANVLAILAEVLSGIFAEVMQGKPEFTD
TRINITY_DN32981_c5_g1_i1                     IDTGFFELQPKEGLAMVNGTAVGSGLASMVLFEANILAVLSEVFSAIFAEVMNGKPEFTD
TRINITY_DN32981_c5_g1_i16                    VDSGFFELQPKEGLAMVNGTAVGSGLASMVLFEANILAVLSEVFSAIFAEVMNGKPEFTD
TRINITY_DN32981_c5_g1_i9                     VDSGFFELQPKEGLAMVNGTAVGSGLASMVLFEANILAVLSEVFSAIFAEVMNGKPEFTD
TRINITY_DN32981_c5_g1_i17                    VDSGFFELQPKEGLAMVNGTAVGSGLASMVLFEANILAVLSEVFSAIFAEVMNGKPEFTD
TRINITY_DN32981_c5_g1_i12                    VDSGFFELQPKEGLAMVNGTAVGSGLASMVLFEANILAVLSEVFSAIFAEVMNGKPEFTD
TRINITY_DN32981_c5_g1_i5                     IDTGFFELQPKEGLAMVNGTAVGSGLASMVLFEANILAVLSEVFSAIFAEVMNGKPEFTD
TRINITY_DN32981_c5_g1_i13                    VDSGFFELQPKEGLAMVNGTAVGSGLASMVLFEANILAVLSEVFSAIFAEVMNGKPEFTD
TRINITY_DN32981_c5_g1_i14                    IDTGFFELQPKEGLAMVNGTAVGSGLASMVLFEANILAVLSEVFSAIFAEVMNGKPEFTD
TRINITY_DN23351_c0_g2_i1                     IKSP.FELQAKEGLALVNGTAVGSAVAATVCFDANILAFLAEIVSALFCEVMHGKPEYTD
sp|P45728.1_PAL2_Petroselinum_crispum        VEGGFFELQPKEGLALVNGTAVGSGMASMVLFEANILAVLAEVMSAIFAEVMQGKPEFTD
AWW24969.1_PAL_Lycoris_radiata               ITDGFFQLQPKEGLALVNGTAVGSGLASTVLYDTNILAVLAEVLSAIFCEVMQGKPEFTD
AIY24976.1_PAL_Mangifera_indica              IGSGFFELQPKEGLALVNGTAVGSGLASTVLFEANILAILSEVLSAIFAEVMNGKPEFTD
AHY94892.1_PAL_Prunella_vulgaris             VSGGFFELQPKEGLALVNGTAVGSGLATMALYNANLLAVLSVVMSAVFAEVMNGKPEFTD
AHA42443.1_PAL_Cannabis_sativa               INSSFFELQPKEGLALVNGTAVGSGVASTVLFEANILAVLAEVLSAIFAEVIQGKPEFTD
AGL81344.1_PAL_Pyrus_communis                INSGFFELQPKEGLALVNGTAVGSGLASTVLFETNILALLAEILSAIFAEVMHGKPEFTD
QIC54077.1_PAL_Paeonia_lactiflora            INTGFFELQPKEGLALVNGTAVGSGLASMVLFEANILAILSEVLSGLFAEVMQGKPEFTD
NP_181241.1_PAL1_Arabidopsis_thaliana        ISSGFFDLQPKEGLALVNGTAVGSGMASMVLFETNVLSVLAEILSAVFAEVMSGKPEFTD
NP_190894.1_PAL2_Arabidopsis_thaliana        ISTGFFDLQPKEGLALVNGTAVGSGMASMVLFEANVQAVLAEVLSAIFAEVMSGKPEFTD
NP_187645.1_PAL4_Arabidopsis_thaliana        VSS.FFELQPKEGLALVNGTAVGSGLASTVLFDANILAVLSEVMSAMFAEVMQGKPEFTD
sp|Q42667.1_PAL_Citrus_limon                 VTSGFFELQPKEGLALVNGTAVGSGLAATVLFEANILAIMSEVLSAIFAEVMNGKPEFTD
CAB42793.1_PAL_Citrus_clementina_x_Citrus_r  F..GFFELQPKEGLALVNGTAVGSGLASMVLFDANNLALLSEILSAIFAEVMQGKPEFTD
CAB42794.1_PAL_Citrus_clementina_x_Citrus_r  F..GFFELQPKEGLALVNGTAVGSGLASMVLFEANNLALLSEILSAIFAEVMQGKPEFTD
sp|P27991.1_PAL_Glycine_max                  INSEFFELQPKEGLALVNGTAVGSGLASMVLFEANILAVLSEVLSAIFAEVMQGKPEFTD
sp|O49835.1_PAL1_Lithospermum_erythrorhizon  ISTGFFELQPKEGLALVNGTAVGSGMASMVLYEANILAVLSEVISAIFAEVMNGKPEFTD
sp|P35511.1_PAL1_Lycopersicon_esculentum     ISGGFFELQPKEGLALVNGTAVGSAMASIVLFESNIFAVMSEVLSAIFTEVMNGKPEFTD
sp|P26600.1_PAL5_Lycopersicon_esculentum     VTSGFFELQPKEGLALVNGTAVGSGMASMVLFESNILAVMSEVLSAIFAEVMNGKPEFTD
sp|P25872.1_PAL1_Nicotiana_tabacum           VNGGFFELQPKEGLALVNGTAVGSGMASMVLFDSNILAVMSEVLSAIFAEVMNGKPEFTD
sp|P35513.2_PAL2_Nicotiana_tabacum           VNGGFFELQPKEGLALVNGTAVGSGLASMVLFDANVLAVFSEVLSAIFAEVMNGKPEFTD
sp|P45733.1_PAL3_Nicotiana_tabacum           VNSGFFELQPKEGLALVNGTAVGSGLASMVLFDANILAVFSEVLSAIFAEVMNGKPEFTD
sp|P19142.1_PAL2_Phaseolus_vulgaris          IGSEFFELQPKEGLALVNGTAVGSGLASIVLFEANILAVLSEVISAIFAEVMQGKPEFTD
sp|Q01861.1_PAL1_Pisum_sativum               INDGFFELQPKEGLALVNGTAVGSGLASIVLFEANILAVLSEVLSAIFAEVMQGKPEFTD
sp|Q04593.1_PAL2_Pisum_sativum               INDGFFELQPKEGLALVNGTAVGSGLASIVLFEANILAVLSEVLSAIFAEVMQGKPEFTD
sp|P45731.1_PAL1_Populus_sieboldii_x_Populu  INGGFFELQPKEGLALVNGTAVGSGLASMVLFEANVLAILSEVLSAIFAEVMQGKPEFTD
sp|P45730.1_PAL_Populus_trichocarpa          IESGFFELQPKEGLALVNGTAVGSGLASMVLFETNVLAVLSELLSAIFAEVMNGKPEFTD
sp|Q43052.1_PAL2_Populus_sieboldii_x_Populu  IDGGFFELQPKEGLALVNGTAVGSGLASMVLFEANVLAILSEVLSAIFAEVMQGKPEFTD
AAF40223.1_PAL1_Rubus_idaeus                 IDGGFFELQPKEGLALVNGTAVGSGMASMVLFDANTLAVLSEVMSAIFAEVMQGKPEFTD
AAF40224.1_PAL2_Rubus_idaeus                 ISSGFFELQPKEGLALVNGTAVGSGLASTVLFETNILALLSEILSAIFAEVMQGKPEFTD
AAC78457.1_PAL_Prunus_avium                  INSGFFELQPKEGLALVNGTAVGSGLASTVLFDTNILALLSEILSAIFAEVMQGKPEFTD
sp|P31425.1_PAL1_Solanum_tuberosum           VSGGFFELQPKEGLALVNGTAVGSGMASIVLYDSNILAVMFEVLSAIFAEVMNGKPEFTD
sp|A2X7F7.1_PAL2_Oryza_sativa                IQGGFFELQPKEGLAMVNGTAVGSGLASTVLFEANILAILAEVLSAVFCEVMNGKPEYTD
sp|P52777.1_PAL_Pinus_taeda                  LEKP.FELQPKEGLAIVNGTSVGAALASIVCFDANVLALLSEVISAMFCEVMNGKPEFTD
ABU49842.1_PAL_Ginkgo_biloba                 LEKP.FELQPKEGLAIVNGTSVGAALASIVCFDANVLAVLSEVMSAMFCEVMNGKPEFTD
ACS28225.2_PAL_Pinus_massoniana              LEKP.FELQPKEGLAIVNGTSVGAALASIVCFDANVLALLSEVISAMFCEVMNGKPEFTD
BAG74771.1_PAL_Ephedra_sinica                IEKP.FELQPKEGLAIVNGTAVCAALASLVCFDANVLALLSEVMAAMFCEVMNGKPEFAD
ABC69916.1_PAL_Brassica_napus                VTSGFFDLQPKESLALVNGTAVGSGMASMVLFEANVLSVLAEVLSAVFAEVMSGKPEFTD
BAC56977.1_PAL_Daucus_carota                 VEGGFFELQPKEGLALVNGTAVGSGMASMVLFEANILAVLAEVMSAIFAEVMQGKPEFTD
ABK24709.1_PAL_Picea_sitchensis              LEKP.FELQPKEGLAIVNGTSVGAALASIVCFDANVLALLSEVISALFCEVMNGKPEFTD
AHA44840.1_PAL_Larix_kaempferi               VEKP.FELAPKEGLAIVNGTSVGAALASIVCFDANVLALLSEVISAMFCEVMNGKPEFTD
AGC23439.1_PAL_Dendrobium_candidum           ISGGFFELQPKEGLALVNGTAVGSGLASTVLFEANILSVMAEVLSAVFCEVMQGKPEYTD
ABI33979.1_PAL_Jatropha_curcas               IDSGFFELQPKEGLALVNGTAVGSGLASMVLFEANVLAVLSEILSAIFAEVMNGKPEFTD
XP_002326186.1_PAL_Populus_trichocarpa       IDSGFFELQPKEGLALVNGTAVGSGLASMVLFETNVLAVLSELISAIFAEVMNGKPEFTD
XP_002519521.1_PAL_Ricinus_communis          IESGFFELQPKEGLALVNGTAVGSGLASMVLFEANILAVLSEILSAIFAEVMNGKPEFTD
XP_002268732.1_PAL_Vitis_vinifera            IESGFFELQPKEGLALVNGTAVGSGLASMVLFETNVLAVLSEVLSAVFAEVMQGKPEFTD
AFG26322.1_PAL_Cinnamomum_osmophloeum        ITDGFFELQPKEGLALVNGTAVGSGLASMVLFEANVLSVLSEVISAIFCEVMQGKPEYTD
BAG70992.1_PAL_Musa_balbisiana               IADGFFELQPKEGLALVNGTAVGSGLASMVLFEANVLAVLSEVLSAVFCEVMQGKPEFTD
AAP34199.1_PAL_Phalaenopsis_x_Doritaenopsis  ISTGFFELQPKEGLALVNGTAVGSGLASLVLFETNILAIMAEVLSALFCEVMQGKPEYTD
ACM61988.1_PAL_Lycoris_radiata               ITDGFFQLQPKEGLALVNGTAVGSGLASTVLYDTNILAVLAEVLSAIFCEVMQGKPEFTD
ABN79671.2_PAL_Rudbeckia_hirta               VKGGFFELQPKEGLALVNGTAVGSGMASMVLFEANVLALLSEVLSAIFAEVMQGKPEFTD
ACJ66297.1_PAL4_Nicotiana_tabacum            VNGGFFELQPKEGLALVNGTAVGSGMASMVLFDSNILAVMSEVLSAIFAEVMNGKPEFTD
ABG75911.1_PAL2_Nicotiana_attenuata          VNGGFFELQPKEGLALVNGTAVGSGMASMVLFDSNILAVMSEVLSAIFAEVMNGKPEFTD
BAL49995.1_PAL_Eucalyptus_robusta            IDTGFFELQPKEGLALVNGTAVGSGLASIVLFEANILAVLSEVLSAIFAEVMQGKPEFTD
ACM61988.1_PAL_Lycoris_radiata               ITDGFFQLQPKEGLALVNGTAVGSGLASTVLYDTNILAVLAEVLSAIFCEVMQGKPEFTD
BAM28963.1_PAL_Lilium_hybrid_division_I      ISGGFFELQPKEGLALVNGTAVGSGLASMVLFEANILAVLAEVLSAIFCEVMQGKPEFTD
                                        acc
```

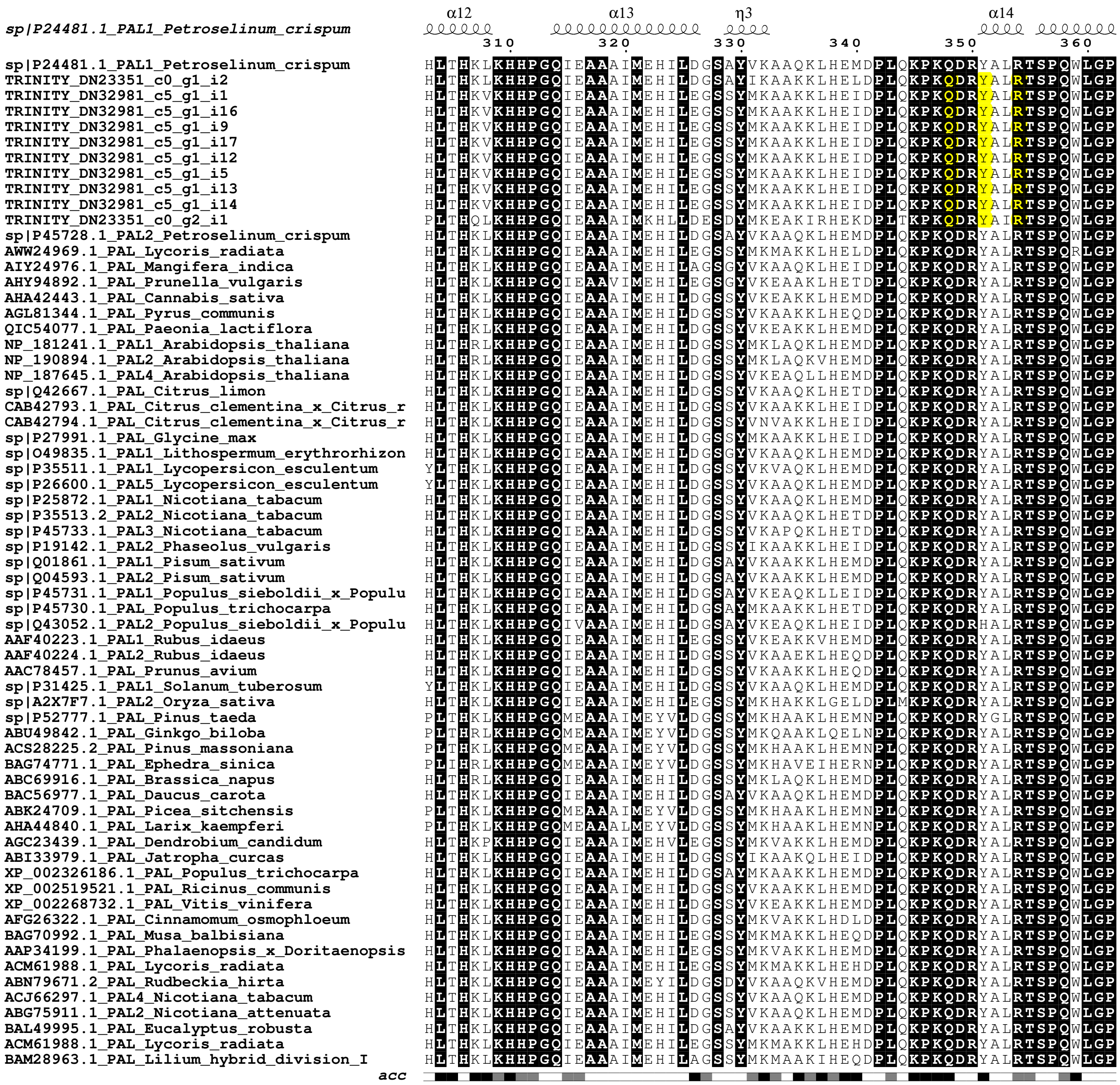
sp|P24481.1_PAL1_Petroselinum_crispum
α12
α13
η3
α14
310
320
330
340
350
360
sp|P24481.1_PAL1_Petroselinum_crispum
TRINITY_DN23351_c0_g1_i2
TRINITY_DN32981_c5_g1_i1
TRINITY_DN32981_c5_g1_i16
TRINITY_DN32981_c5_g1_i9
TRINITY_DN32981_c5_g1_i17
TRINITY_DN32981_c5_g1_i12
TRINITY_DN32981_c5_g1_i5
TRINITY_DN32981_c5_g1_i13
TRINITY_DN32981_c5_g1_i14
TRINITY_DN23351_c0_g2_i1
sp|P45728.1_PAL2_Petroselinum_crispum
AWW24969.1_PAL_Lycoris_radiata
AIY24976.1_PAL_Mangifera_indica
AHY94892.1_PAL_Prunella_vulgaris
AHA42443.1_PAL_Cannabis_sativa
AGL81344.1_PAL_Pyrus_communis
QIC54077.1_PAL_Paeonia_lactiflora
NP_181241.1_PAL1_Arabidopsis_thaliana
NP_190894.1_PAL2_Arabidopsis_thaliana
NP_187645.1_PAL4_Arabidopsis_thaliana
sp|Q42667.1_PAL_Citrus_limon
CAB42793.1_PAL_Citrus_clementina_x_Citrus_r
CAB42794.1_PAL_Citrus_clementina_x_Citrus_r
sp|P27991.1_PAL_Glycine_max
sp|O49835.1_PAL1_Lithospermum_erythrorhizon
sp|P35511.1_PAL1_Lycopersicon_esculentum
sp|P26600.1_PAL5_Lycopersicon_esculentum
sp|P25872.1_PAL1_Nicotiana_tabacum
sp|P35513.2_PAL2_Nicotiana_tabacum
sp|P45733.1_PAL3_Nicotiana_tabacum
sp|P19142.1_PAL2_Phaseolus_vulgaris
sp|Q01861.1_PAL1_Pisum_sativum
sp|Q04593.1_PAL2_Pisum_sativum
sp|P45731.1_PAL1_Populus_sieboldii_x_Populu
sp|P45730.1_PAL_Populus_trichocarpa
sp|Q43052.1_PAL2_Populus_sieboldii_x_Populu
AAF40223.1_PAL1_Rubus_idaeus
AAF40224.1_PAL2_Rubus_idaeus
AAC78457.1_PAL_Prunus_avium
sp|P31425.1_PAL1_Solanum_tuberosum
sp|A2X7F7.1_PAL2_Oryza_sativa
sp|P52777.1_PAL_Pinus_taeda
ABU49842.1_PAL_Ginkgo_biloba
ACS28225.2_PAL_Pinus_massoniana
BAG74771.1_PAL_Ephedra_sinica
ABC69916.1_PAL_Brassica_napus
BAC56977.1_PAL_Daucus_carota
ABK24709.1_PAL_Picea_sitchensis
AHA44840.1_PAL_Larix_kaempferi
AGC23439.1_PAL_Dendrobium_candidum
ABI33979.1_PAL_Jatropha_curcas
XP_002326186.1_PAL_Populus_trichocarpa
XP_002519521.1_PAL_Ricinus_communis
XP_002268732.1_PAL_Vitis_vinifera
AFG26322.1_PAL_Cinnamomum_osmophloeum
BAG70992.1_PAL_Musa_balbisiana
AAP34199.1_PAL_Phalaenopsis_x_Doritaenopsis
ACM61988.1_PAL_Lycoris_radiata
ABN79671.2_PAL_Rudbeckia_hirta
ACJ66297.1_PAL4_Nicotiana_tabacum
ABG75911.1_PAL2_Nicotiana_attenuata
BAL49995.1_PAL_Eucalyptus_robusta
ACM61988.1_PAL_Lycoris_radiata
BAM28963.1_PAL_Lilium_hybrid_division_I
acc

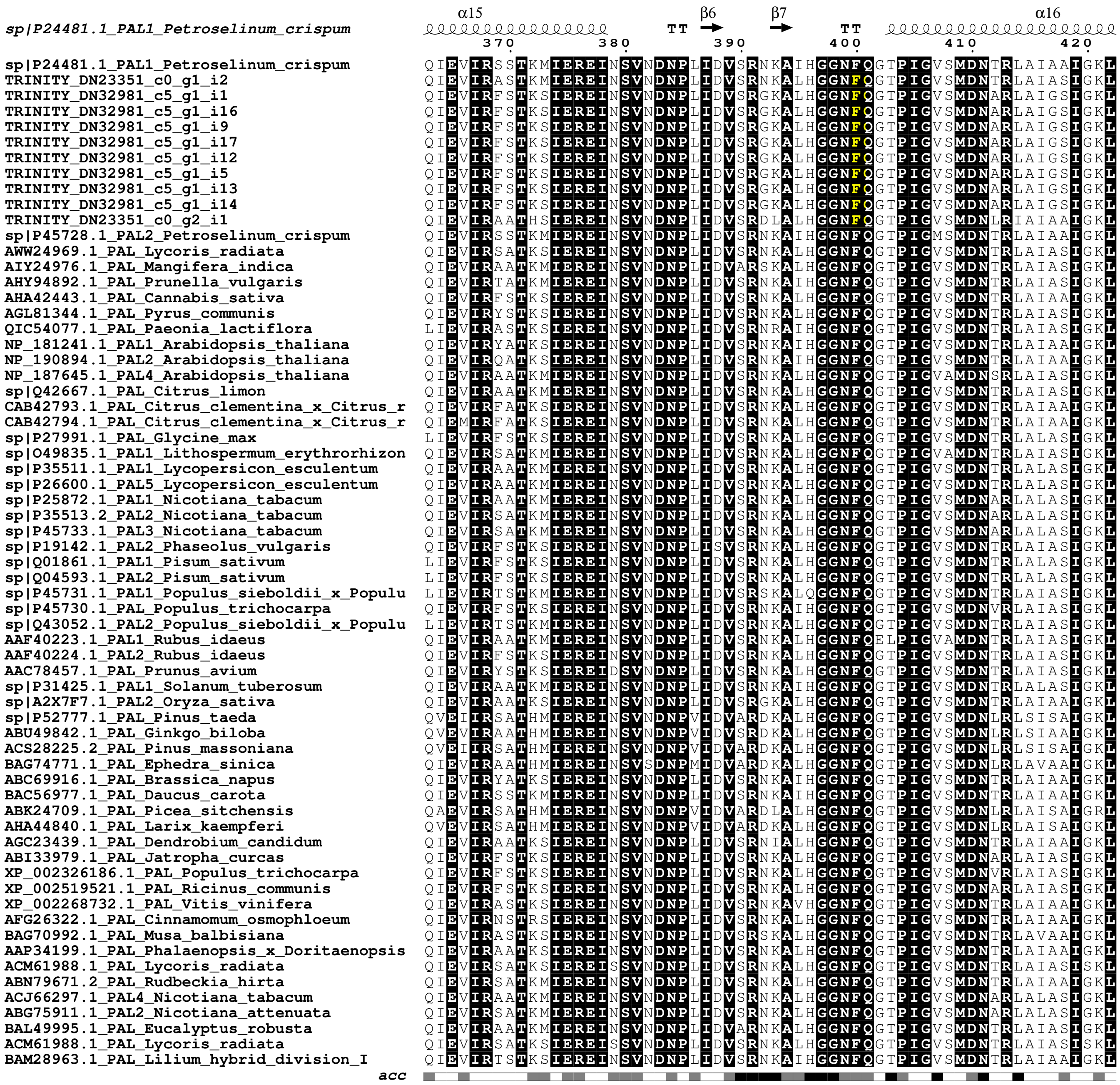
α15 β6 β7 α16
sp|P24481.1_PAL1_Petroselinum_crispum
TT TT
370 380 390 400 410 420
sp|P24481.1_PAL1_Petroselinum_crispum QIEVIRSSTKMIEREINSVNDNPLIDVSRNKAIHGGNFQGTPIGVSMDNTRLAIAAIGKL
TRINITY_DN23351_c0_g1_i2 QIEVIRASTKMIEREINSVNDNPLIDVSRNKALHGGNFQGTPIGVSMDNTRLAIASIGKL
TRINITY_DN32981_c5_g1_i1 QIEVIRFSTKSIEREINSVNDNPLIDVSRGKALHGGNFQGTPIGVSMDNARLAIGSIGKL
TRINITY_DN32981_c5_g1_i16 QIEVIRFSTKSIEREINSVNDNPLIDVSRGKALHGGNFQGTPIGVSMDNARLAIGSIGKL
TRINITY_DN32981_c5_g1_i9 QIEVIRFSTKSIEREINSVNDNPLIDVSRGKALHGGNFQGTPIGVSMDNARLAIGSIGKL
TRINITY_DN32981_c5_g1_i17 QIEVIRFSTKSIEREINSVNDNPLIDVSRGKALHGGNFQGTPIGVSMDNARLAIGSIGKL
TRINITY_DN32981_c5_g1_i12 QIEVIRFSTKSIEREINSVNDNPLIDVSRGKALHGGNFQGTPIGVSMDNARLAIGSIGKL
TRINITY_DN32981_c5_g1_i5 QIEVIRFSTKSIEREINSVNDNPLIDVSRGKALHGGNFQGTPIGVSMDNARLAIGSIGKL
TRINITY_DN32981_c5_g1_i13 QIEVIRFSTKSIEREINSVNDNPLIDVSRGKALHGGNFQGTPIGVSMDNARLAIGSIGKL
TRINITY_DN32981_c5_g1_i14 QIEVIRFSTKSIEREINSVNDNPLIDVSRGKALHGGNFQGTPIGVSMDNARLAIGSIGKL
TRINITY_DN23351_c0_g2_i1 QIEVIRAATHSIEREINSVNDNPIIDVSRDLALHGGNFQGTPIGVSMDNLRIAIAAIGKL
sp|P45728.1_PAL2_Petroselinum_crispum QIEVIRSSTKMIEREINSVNDNPLIDVSRNKAIHGGNFQGTPIGMSMDNTRLAIAAIGKL
AWW24969.1_PAL_Lycoris_radiata QIEVIRSATKSIEREINSVNDNPLIDVSRNKALHGGNFQGTPIGVSMDNTRLAIASIGKL
AIY24976.1_PAL_Mangifera_indica QIEVIRAATKMIEREINSVNDNPLIDVARSKALHGGNFQGTPIGVSMDNTRLAIASIGKL
AHY94892.1_PAL_Prunella_vulgaris QIEVIRTATKMIEREINSVNDNPLIDVSRNKAIHGGNFQGTPIGVSMDNTRLAIASIGKL
AHA42443.1_PAL_Cannabis_sativa QIEVIRFSTKSIEREINSVNDNPLIDVSRNKALHGGNFQGTPIGVSMDNTRLAIAAIGKL
AGL81344.1_PAL_Pyrus_communis QIEVIRYSTKSIEREINSVNDNPLIDVSRNKALHGGNFQGTPIGVSMDNTRLAIASIGKL
QIC54077.1_PAL_Paeonia_lactiflora LIEVIRASTKSIEREINSVNDNPLIDVSRNRAIHGGNFQGTPIGVSMDNTRLAIASIGKL
NP_181241.1_PAL1_Arabidopsis_thaliana QIEVIRYATKSIEREINSVNDNPLIDVSRNKAIHGGNFQGTPIGVSMDNTRLAIAAIGKL
NP_190894.1_PAL2_Arabidopsis_thaliana QIEVIRQATKSIEREINSVNDNPLIDVSRNKAIHGGNFQGTPIGVSMDNTRLAIAAIGKL
NP_187645.1_PAL4_Arabidopsis_thaliana QIEVIRAATKMIEREINSVNDNPLIDVSRNKALHGGNFQGTPIGVAMDNSRLAIASIGKL
sp|Q42667.1_PAL_Citrus_limon QIEVIRAATKMIEREINSVNDNPLIDVSRNKALHGGNFQGTPIGVSMDNTRLAIASIGKL
CAB42793.1_PAL_Citrus_clementina_x_Citrus_r QIEVIRFATKSIEREINSVNDNPLIDVSRNKALHGGNFQGTPIGVSMDNTRLAIAAIGKL
CAB42794.1_PAL_Citrus_clementina_x_Citrus_r QIEMIRFATKSIEREINSVNDNPLIDVSRNKALHGGNFQGTPIGVSMDNTRLAIAAIGKL
sp|P27991.1_PAL_Glycine_max LIEVIRFSTKSIEREINSVNDNPLIDVSRNKALHGGNFQGTPIGVSMDNTRLALASIGKL
sp|O49835.1_PAL1_Lithospermum_erythrorhizon QIEVIRSATKMIEREINSVNDNPLIDVSRNKALHGGNFQGTPIGVAMDNTRLAIASIGKL
sp|P35511.1_PAL1_Lycopersicon_esculentum QIEVIRAATKMIEREINSVNDNPLIDVSRNKALHGGNFQGTPIGVSMDNTRLALASIGKL
sp|P26600.1_PAL5_Lycopersicon_esculentum QIEVIRAATKMIEREINSVNDNPLIDVSRNKALHGGNFQGTPIGVSMDNTRLALASIGKL
sp|P25872.1_PAL1_Nicotiana_tabacum QIEVIRAATKMIEREINSVNDNPLIDVSRNKALHGGNFQGTPIGVSMDNARLALASIGKL
sp|P35513.2_PAL2_Nicotiana_tabacum QIEVIRSATKMIEREINSVNDNPLIDVSRNKALHGGNFQGTPIGVSMDNARLALASIGKL
sp|P45733.1_PAL3_Nicotiana_tabacum QIEVIRSATKMIEREINSVNDNPLIDVSRNKALHGGNFQGTPIGVSMDNARLALASIGKL
sp|P19142.1_PAL2_Phaseolus_vulgaris QIEVIRFSTKSIEREINSVNDNPLISVSRNKALHGGNFQGTPIGVSMDNTRLAIASIGKL
sp|Q01861.1_PAL1_Pisum_sativum LIEVIRFSTKSIEREINSVNDNPLIDVSRNKALHGGNFQGTPIGVSMDNTRLALASIGKL
sp|Q04593.1_PAL2_Pisum_sativum LIEVIRFSTKSIEREINSVNDNPLIDVSRNKALHGGNFQGTPIGVSMDNTRLALASIGKL
sp|P45731.1_PAL1_Populus_sieboldii_x_Populu LIEVIRTSTKMIEREINSVNDNPLIDVSRSKALQGGNFQGTPIGVSMDNTRLAIASIGKL
sp|P45730.1_PAL_Populus_trichocarpa QIEVIRFSTKSIEREINSVNDNPLIDVSRNKAIHGGNFQGTPIGVSMDNVRLAIASIGKL
sp|Q43052.1_PAL2_Populus_sieboldii_x_Populu LIEVIRTSTKMIEREINSVNDNPLIDVSRNKALHGGNFQGTPIGVSMDNTRLAIASIGKL
AAF40223.1_PAL1_Rubus_idaeus QIEVIRAATKMIEREINSVNDNPLIDVSRNKALHGGNFQELPIGVAMDNTRLAIASIGKL
AAF40224.1_PAL2_Rubus_idaeus QIEVIRFSTKSIEREINSVNDNPLIDVSRNKALHGGNFQGTPIGVSMDNTRLAIASIGKL
AAC78457.1_PAL_Prunus_avium QIEVIRYSTKSIEREIDSVNDNPLIDVSRNKALHGGNFQGTPIGVSMDNTRLAIASIGKL
sp|P31425.1_PAL1_Solanum_tuberosum QIEVIRAATKMIEREINSVNDNPLIDVSRNKAIHGGNFQGTPIGVSMDNTRLALASIGKL
sp|A2X7F7.1_PAL2_Oryza_sativa QIEVIRAATKSIEREINSVNDNPLIDVSRGKALHGGNFQGTPIGVSMDNTRLAIAAIGKL
sp|P52777.1_PAL_Pinus_taeda QVEIIRSATHMIEREINSVNDNPVIDVARDKALHGGNFQGTPIGVSMDNLRLSISAIGKL
ABU49842.1_PAL_Ginkgo_biloba QVEVIRAATHMIEREINSVNDNPVIDVSRDKALHGGNFQGTPIGVSMDNLRLSIAAIGKL
ACS28225.2_PAL_Pinus_massoniana QVEIIRSATHMIEREINSVNDNPVIDVARDKALHGGNFQGTPIGVSMDNLRLSISAIGKL
BAG74771.1_PAL_Ephedra_sinica QIEVIRAATHMIEREINSVSDNPMIDVARDKALHGGNFQGTPIGVSMDNLRLAVAAIGKL
ABC69916.1_PAL_Brassica_napus QIEVIRYATKSIEREINSVNDNPLIDVSRNKAIHGGNFQGTPIGVSMDNTRLAIAAIGKL
BAC56977.1_PAL_Daucus_carota QIEVIRSSTKMIEREINSVNDNPLIDVSRNKAIHGGNFQGTPIGVSMDNTRLAIAAIGKL
ABK24709.1_PAL_Picea_sitchensis QAEVIRSATHMIEREINSVNDNPVIDVARDLALHGGNFQGTPIGVSMDNLRLAISAIGRL
AHA44840.1_PAL_Larix_kaempferi QVEVIRSATHMIEREINSVNDNPVIDVARDKALHGGNFQGTPIGVSMDNLRLAISAIGKL
AGC23439.1_PAL_Dendrobium_candidum QIEVIRAATKSIEREINSVNDNPLIDVSRNIALHGGNFQGTPIGVSMDNTRLAIAAIGKL
ABI33979.1_PAL_Jatropha_curcas QIEVIRFSTKSIEREINSVNDNPLIDVSRNKALHGGNFQGTPIGVSMDNARLAIASIGKL
XP_002326186.1_PAL_Populus_trichocarpa QIEVIRFSTKSIEREINSVNDNPLIDVSRNKALHGGNFQGTPIGVSMDNVRLAIASIGKL
XP_002519521.1_PAL_Ricinus_communis QIEVIRFSTKSIEREINSVNDNPLIDVSRNKALHGGNFQGTPIGVSMDNARLAIASIGKL
XP_002268732.1_PAL_Vitis_vinifera QIEVIRASTKSIEREINSVNDNPLIDVSRNKAVHGGNFQGTPIGVSMDNTRLAIAAIGKL
AFG26322.1_PAL_Cinnamomum_osmophloeum QIEVIRNSTRSIEREINSVNDNPLIDVSRNKALHGGNFQGTPIGVSMDNTRLAIAAIGKL
BAG70992.1_PAL_Musa_balbisiana QIEVIRASTKSIEREINSVNDNPLIDVSRSKALHGGNFQGTPIGVSMDNTRLAVAAIGKL
AAP34199.1_PAL_Phalaenopsis_x_Doritaenopsis QIEVIRAATKSIEREINSVNDNPLIDVSRNKALHGGNFQGTPIGVSMDNTRLAIAAIGKL
ACM61988.1_PAL_Lycoris_radiata QIEVIRSATKSIEREISSVNDNPLIDVSRNKALHGGNFQGTPIGVSMDNTRLAIASISKL
ABN79671.2_PAL_Rudbeckia_hirta QIEVIRSATKMIEREINSVNDNPLIDVSRNKALHGGNFQGTPIGVSMDNTRLAIAAIGKL
ACJ66297.1_PAL4_Nicotiana_tabacum QIEVIRAATKMIEREINSVNDNPLIDVSRNKALHGGNFQGTPIGVSMDNARLALASIGKL
ABG75911.1_PAL2_Nicotiana_attenuata QIEVIRSATKMIEREINSVNDNPLIDVSRNKALHGGNFQGTPIGVSMDNARLALASIGKL
BAL49995.1_PAL_Eucalyptus_robusta QIEVIRAATKMIEREINSVNDNPLIDVARNKALHGGNFQGTPIGVSMDNTRLAIASIGKL
ACM61988.1_PAL_Lycoris_radiata QIEVIRSATKSIEREISSVNDNPLIDVSRNKALHGGNFQGTPIGVSMDNTRLAIASISKL
BAM28963.1_PAL_Lilium_hybrid_division_I QIEVIRTSTKSIEREINSVNDNPLIDVSRNKAIHGGNFQGTPIGVSMDNTRLAIAAIGKL
acc

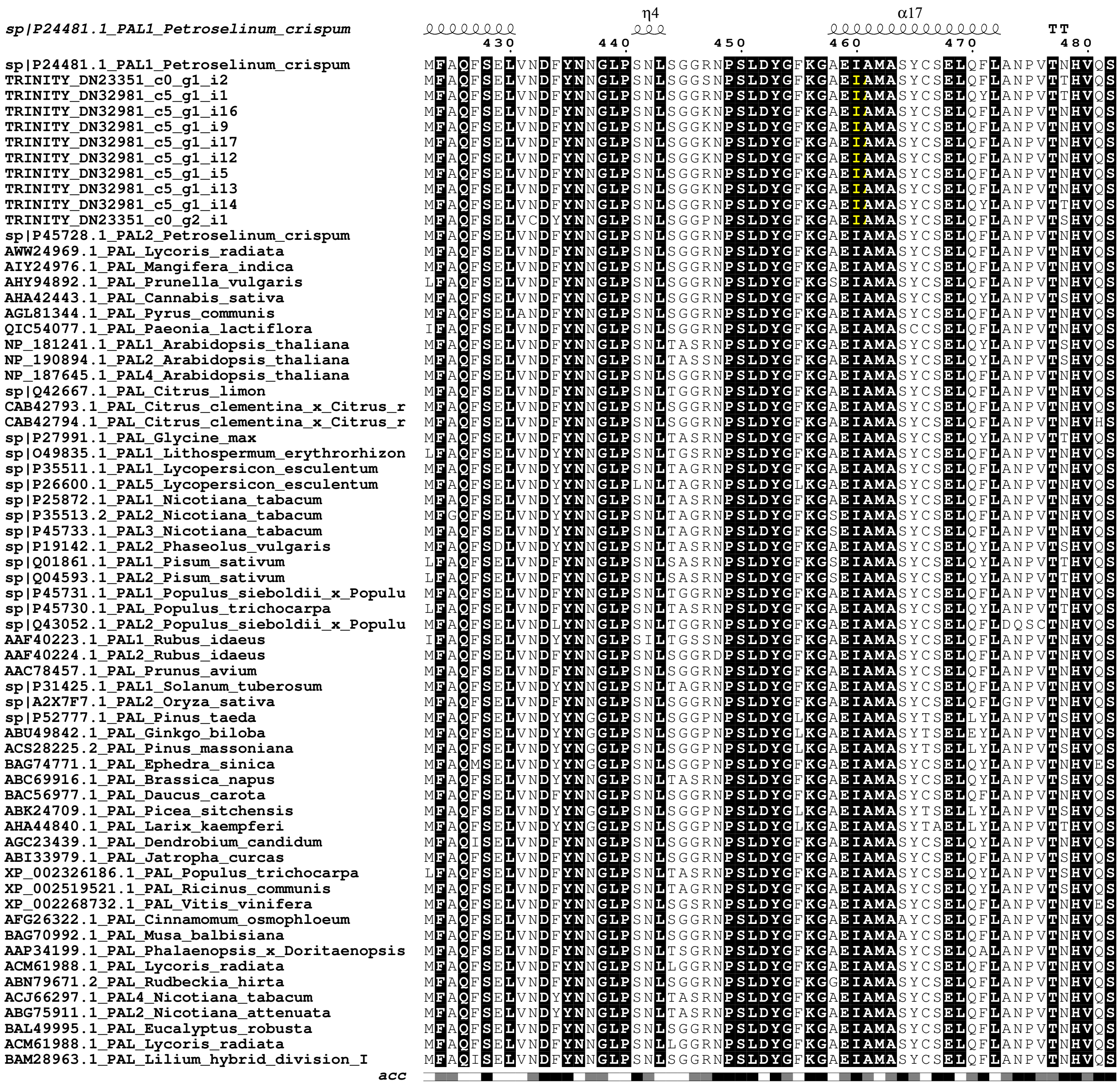

sp|P24481.1_PAL1_Petroselinum_crispum
η4
α17
TT
430
440
450
460
470
480
sp|P24481.1_PAL1_Petroselinum_crispum MFAQFSELVNDFYNNGLPSNLSGGRNPSLDYGFKGAEIAMASYCSELQFLANPVTNHVQS
TRINITY_DN23351_c0_g1_i2 MFAQFSELVNDFYNNGLPSNLSGGSNPSLDYGFKGAEIAMASYCSELQFLANPVTTHVQS
TRINITY_DN32981_c5_g1_i1 MFAQFSELVNDFYNNGLPSNLSGGRNPSLDYGFKGAEIAMASYCSELQYLANPVTTHVQS
TRINITY_DN32981_c5_g1_i16 MFAQFSELVNDFYNNGLPSNLSGGKNPSLDYGFKGAEIAMASYCSELQFLANPVTNHVQS
TRINITY_DN32981_c5_g1_i9 MFAQFSELVNDFYNNGLPSNLSGGKNPSLDYGFKGAEIAMASYCSELQFLANPVTNHVQS
TRINITY_DN32981_c5_g1_i17 MFAQFSELVNDFYNNGLPSNLSGGKNPSLDYGFKGAEIAMASYCSELQFLANPVTNHVQS
TRINITY_DN32981_c5_g1_i12 MFAQFSELVNDFYNNGLPSNLSGGKNPSLDYGFKGAEIAMASYCSELQFLANPVTNHVQS
TRINITY_DN32981_c5_g1_i5 MFAQFSELVNDFYNNGLPSNLSGGRNPSLDYGFKGAEIAMASYCSELQFLANPVTNHVQS
TRINITY_DN32981_c5_g1_i13 MFAQFSELVNDFYNNGLPSNLSGGKNPSLDYGFKGAEIAMASYCSELQFLANPVTNHVQS
TRINITY_DN32981_c5_g1_i14 MFAQFSELVNDFYNNGLPSNLSGGRNPSLDYGFKGAEIAMASYCSELQYLANPVTTHVQS
TRINITY_DN23351_c0_g2_i1 MFAQFSELVCDYYNNGLPSNLSGGPNPSLDYGFKGAEIAMASYCSELQFLANPVTSHVQS
sp|P45728.1_PAL2_Petroselinum_crispum MFAQFSELVNDFYNNGLPSNLSGGRNPSLDYGFKGAEIAMASYCSELQFLANPVTNHVQS
AWW24969.1_PAL_Lycoris_radiata MFAQFSELVNDFYNNGLPSNLSGGRNPSLDYGFKGAEIAMASYCSELQFLANPVTNHVQS
AIY24976.1_PAL_Mangifera_indica MFAQFSELVNDFYNNGLPSNLSGGRNPSLDYGFKGAEIAMASYCSELQFLANPVTNHVQS
AHY94892.1_PAL_Prunella_vulgaris LFAQFSELVNDFYNNGLPSNLSGGRNPSLDYGFKGSEIAMASYCSELQFLANPVTNHVQS
AHA42443.1_PAL_Cannabis_sativa MFAQFSELVNDFYNNGLPSNLSGGRNPSLDYGFKGAEIAMASYCSELQYLANPVTSHVQS
AGL81344.1_PAL_Pyrus_communis MFAQFSELANDFYNNGLPSNLSGGRNPSLDYGFKGAEIAMASYCSELQFLANPVTNHVQS
QIC54077.1_PAL_Paeonia_lactiflora IFAQFSELVNDFYNNGLPSNLSGGRNPSLDYGFKGAEIAMASCCSELQFLANPVTNHVQS
NP_181241.1_PAL1_Arabidopsis_thaliana MFAQFSELVNDFYNNGLPSNLTASRNPSLDYGFKGAEIAMASYCSELQYLANPVTSHVQS
NP_190894.1_PAL2_Arabidopsis_thaliana MFAQFSELVNDFYNNGLPSNLTASSNPSLDYGFKGAEIAMASYCSELQYLANPVTSHVQS
NP_187645.1_PAL4_Arabidopsis_thaliana MFAQFSELVNDFYNNGLPSNLSGGRNPSLDYGFKGAEIAMASYCSELQFLANPVTNHVQS
sp|Q42667.1_PAL_Citrus_limon MFAQFSELVNDFYNNGLPSNLTGGRNPSLDYGFKGAEIAMASYCSELQFLANPVTNHVQS
CAB42793.1_PAL_Citrus_clementina_x_Citrus_r MFAQFSELVNDFYNNGLPSNLSGGRNPSLDYGFKGAEIAMASYCSELQFLANPVTNHVQS
CAB42794.1_PAL_Citrus_clementina_x_Citrus_r MFAQFSELVNDFYNNGLPSNLSGGRNPSLDYGFKGAEIAMASYCSELQFLANPVTNHVHS
sp|P27991.1_PAL_Glycine_max MFAQFSELVNDFYNNGLPSNLTASRNPSLDYGFKGAEIAMASYCSELQYLANPVTTHVQS
sp|O49835.1_PAL1_Lithospermum_erythrorhizon LFAQFSELVNDYYNNGLPSNLTGSRNPSLDYGFKGAEIAMASYCSELQFLANPVTNHVQS
sp|P35511.1_PAL1_Lycopersicon_esculentum MFAQFSELVNDYYNNGLPSNLTAGRNPSLDYGFKGAEIAMASYCSELQFLANPVTNHVQS
sp|P26600.1_PAL5_Lycopersicon_esculentum MFAQFSELVNDYYNNGLPLNLTAGRNPSLDYGLKGAEIAMASYCSELQFLANPVTNHVQS
sp|P25872.1_PAL1_Nicotiana_tabacum MFAQFSELVNDYYNNGLPSNLTASRNPSLDYGFKGAEIAMASYCSELQFLANPVTNHVQS
sp|P35513.2_PAL2_Nicotiana_tabacum MFGQFSELVNDYYNNGLPSNLTAGRNPSLDYGFKGSEIAMASYCSELQFLANPVTNHVQS
sp|P45733.1_PAL3_Nicotiana_tabacum MFAQFSELVNDYYNNGLPSNLTAGRNPSLDYGFKGSEIAMASYCSELQFLANPVTNHVQS
sp|P19142.1_PAL2_Phaseolus_vulgaris MFAQFSDLVNDYYNNGLPSNLTASRNPSLDYGFKGAEIAMASYCSELQYLANPVTSHVQS
sp|Q01861.1_PAL1_Pisum_sativum LFAQFSELVNDFYNNGLPSNLSASRNPSLDYGFKGSEIAMASYCSELQYLANPVTTHVQS
sp|Q04593.1_PAL2_Pisum_sativum LFAQFSELVNDFYNNGLPSNLSASRNPSLDYGFKGSEIAMASYCSELQYLANPVTTHVQS
sp|P45731.1_PAL1_Populus_sieboldii_x_Populu MFAQFSELVNDFYNNGLPSNLTGGRNPSLDYGFKGAEIAMASYCSELQFLANPVTNHVQS
sp|P45730.1_PAL_Populus_trichocarpa LFAQFSELVNDFYNNGLPSNLTASRNPSLDYGFKGAEIAMASYCSELQYLANPVTTHVQS
sp|Q43052.1_PAL2_Populus_sieboldii_x_Populu MFAQFSELVNDLYNNGLPSNLTGGRNPSLDYGFKGAEIAMASYCSELQFLDQSCTNHVQS
AAF40223.1_PAL1_Rubus_idaeus IFAQFSELVNDYYNNGLPSILTGSSNPSLDYGFKGAEIAMASYCSELQFLANPVTNHVQS
AAF40224.1_PAL2_Rubus_idaeus MFAQFSELVNDFYNNGLPSNLSGGRDPSLDYGFKGAEIAMASYCSELQFLANPVTNHVQS
AAC78457.1_PAL_Prunus_avium MFAQFSELVNDFYNNGLPSNLSGGRNPSLDYGFKGAEIAMASYCSELQFLANPVTNHVQS
sp|P31425.1_PAL1_Solanum_tuberosum MFAQFSELVNDYYNNGLPSNLTAGRNPSLDYGFKGAEIAMASYCSELQFLANPVTNHVQS
sp|A2X7F7.1_PAL2_Oryza_sativa MFAQFSELVNDFYNNGLPSNLSGGRNPSLDYGFKGAEIAMASYCSELQFLGNPVTNHVQS
sp|P52777.1_PAL_Pinus_taeda MFAQFSELVNDYYNGGLPSNLSGGPNPSLDYGLKGAEIAMASYTSELLYLANPVTSHVQS
ABU49842.1_PAL_Ginkgo_biloba MFAQFSELVNDYYNGGLPSNLSGGPNPSLDYGLKGAEIAMASYTSELEYLANPVTNHVQS
ACS28225.2_PAL_Pinus_massoniana MFAQFSELVNDYYNGGLPSNLSGGPNPSLDYGLKGAEIAMASYTSELLYLANPVTSHVQS
BAG74771.1_PAL_Ephedra_sinica MFAQMSELVNDYYNGGLPSNLSGGPNPSLDYGFKGAEIAMASYCSELQYLANPVTNHVES
ABC69916.1_PAL_Brassica_napus MFAQFSELVNDYYNNGLPSNLTASRNPSLDYGFKGAEIAMASYCSELQYLANPVTSHVQS
BAC56977.1_PAL_Daucus_carota MFAQFSELVNDFYNNGLPSNLSGGRNPSLDYGFKGAEIAMASYCSELQFLANPVTNHVQS
ABK24709.1_PAL_Picea_sitchensis MFAQFSELVNDYYNGGLPSNLSGGPNPSLDYGLKGAEIAMASYTSELLYLANPVTSHVQS
AHA44840.1_PAL_Larix_kaempferi MFAQFSELVNDYYNGGLPSNLSGGPNPSLDYGLKGAEIAMASYTAELLYLANPVTTHVQS
AGC23439.1_PAL_Dendrobium_candidum MFAQISELVNDFYNNGLPSNLSGGRNPSLDYGFKGAEIAMASYCSELQYLANPVTNHVQS
ABI33979.1_PAL_Jatropha_curcas MFAQFSELVNDFYNNGLPSNLSGGRNPSLDYGFKGAEIAMASYCSELQFLANPVTSHVQS
XP_002326186.1_PAL_Populus_trichocarpa LFAQFSELVNDFYNNGLPSNLTASRNPSLDYGFKGAEIAMASYCSELQYLANPVTSHVQS
XP_002519521.1_PAL_Ricinus_communis MFAQFSELVNDFYNNGLPSNLTAGRNPSLDYGFKGAEIAMASYCSELQYLANPVTSHVQS
XP_002268732.1_PAL_Vitis_vinifera MFAQFSELVNDFYNNGLPSNLSGSRNPSLDYGFKGAEIAMASYCSELQFLANPVTNHVES
AFG26322.1_PAL_Cinnamomum_osmophloeum MFAQFSELVNDFYNNGLPSNLSGGRNPSLDYGFKGAEIAMAAYCSELQFLANPVTNHVQS
BAG70992.1_PAL_Musa_balbisiana MFAQFSELVNDFYNNGLPSNLSGGRNPSLDYGFKGAEIAMAAYCSELQFLANPVTNHVQS
AAP34199.1_PAL_Phalaenopsis_x_Doritaenopsis MFAQFSELVNDFYNNGLPSNLTSGRNPSLDYGFKGAEIAMASYCSELQALANPVTNHVQS
ACM61988.1_PAL_Lycoris_radiata MFAQFSELVNDFYNNGLPSNLLGGRNPSLDYGFKGAEIAMASYCSELQFLANPVTNHVQS
ABN79671.2_PAL_Rudbeckia_hirta MFAQFSELVNDFYNNGLPSNLSGGRNPSLDYGFKGGEIAMASYCSELQFLANPVTNHVQS
ACJ66297.1_PAL4_Nicotiana_tabacum MFAQFSELVNDYYNNGLPSNLTASRNPSLDYGFKGAEIAMASYCSELQFLANPVTNHVQS
ABG75911.1_PAL2_Nicotiana_attenuata MFAQFSELVNDYYNNGLPSNLTASRNPSLDYGFKGAEIAMASYCSELQFLANPVTNHVQS
BAL49995.1_PAL_Eucalyptus_robusta MFAQFSELVNDFYNNGLPSNLSGGRNPSLDYGFKGAEIAMASYCSELQFLANPVTNHVQS
ACM61988.1_PAL_Lycoris_radiata MFAQFSELVNDFYNNGLPSNLLGGRNPSLDYGFKGAEIAMASYCSELQFLANPVTNHVQS
BAM28963.1_PAL_Lilium_hybrid_division_I MFAQISELVNDFYNNGLPSNLSGGRNPSLDYGFKGAEIAMASYCSELQFLANPVTNHVQS
acc

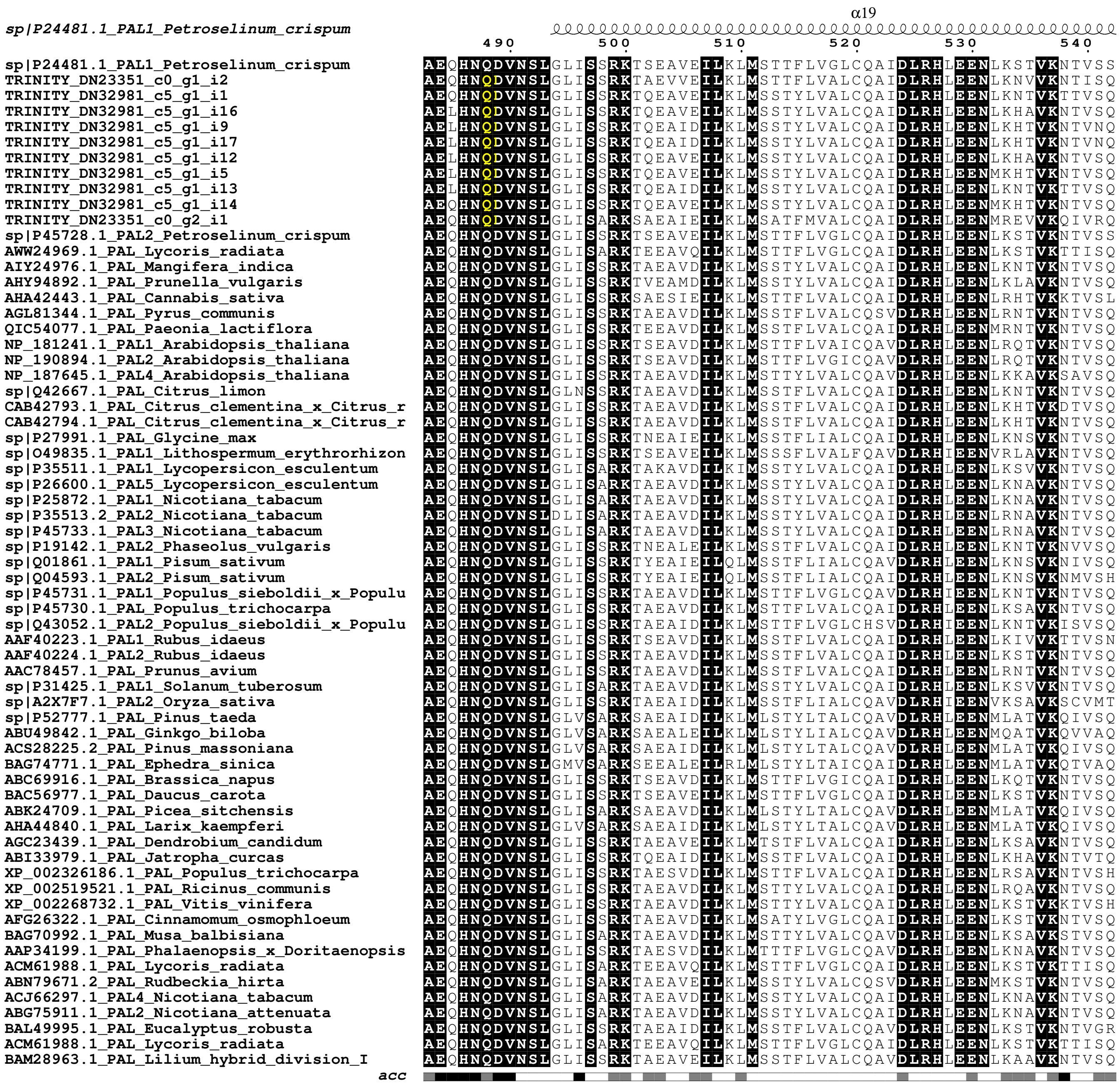
α19
sp|P24481.1_PAL1_Petroselinum_crispum
490 500 510 520 530 540
sp|P24481.1_PAL1_Petroselinum_crispum AEQHNQDVNSLGLISSRKTSEAVEILKLMSTTFLVGLCQAIDLRHLEENLKSTVKNTVSS
TRINITY_DN23351_c0_g1_i2 AEQHNQDVNSLGLISSRKTAEVVEILKLMSTTFLVALCQAIDLRHLEENLKNVVKNTISQ
TRINITY_DN32981_c5_g1_i1 AEQHNQDVNSLGLISSRKTQEAVEILKLMSSTYLVALCQAIDLRHLEENLKNTVKTTVSQ
TRINITY_DN32981_c5_g1_i16 AELHNQDVNSLGLISSRKTQEAVEILKLMSSTYLVALCQAIDLRHLEENLKHAVKNTVSQ
TRINITY_DN32981_c5_g1_i9 AELHNQDVNSLGLISSRKTQEAIDILKLMSSTYLVALCQAIDLRHLEENLKHTVKNTVNQ
TRINITY_DN32981_c5_g1_i17 AELHNQDVNSLGLISSRKTQEAIDILKLMSSTYLVALCQAIDLRHLEENLKHTVKNTVNQ
TRINITY_DN32981_c5_g1_i12 AELHNQDVNSLGLISSRKTQEAVEILKLMSSTYLVALCQAIDLRHLEENLKHAVKNTVSQ
TRINITY_DN32981_c5_g1_i5 AELHNQDVNSLGLISSRKTQEAVEILKLMSSTYLVALCQAIDLRHLEENMKHTVKNTVSQ
TRINITY_DN32981_c5_g1_i13 AELHNQDVNSLGLISSRKTQEAIDILKLMSSTYLVALCQAIDLRHLEENLKNTVKTTVSQ
TRINITY_DN32981_c5_g1_i14 AEQHNQDVNSLGLISSRKTQEAVEILKLMSSTYLVALCQAIDLRHLEENMKHTVKNTVSQ
TRINITY_DN23351_c0_g2_i1 AEQHNQDVNSLGLISARKSAEAIEILKLMSATFMVALCQAIDLRHLEENMREVVKQIVRQ
sp|P45728.1_PAL2_Petroselinum_crispum AEQHNQDVNSLGLISSRKTSEAVEILKLMSTTFLVGLCQAIDLRHLEENLKSTVKNTVSS
AWW24969.1_PAL_Lycoris_radiata AEQHNQDVNSLGLISARKTEEAVQILKLMSTTFLVGLCQAIDLRHLEENLKSTVKTTISQ
AIY24976.1_PAL_Mangifera_indica AEQHNQDVNSLGLISSRKTAEAVDILKLMSSTYLVALCQAIDLRHLEENLKNTVKNTVSQ
AHY94892.1_PAL_Prunella_vulgaris AEQHNQDVNSLGLISSRKTVEAMDILKLMSSTYLVALCQAIDLRHLEENLKLAVKNTVSQ
AHA42443.1_PAL_Cannabis_sativa AEQHNQDVNSLGLISSRKSAESIEILKLMSTTFLVALCQAIDLRHLEENLRHTVKKTVSL
AGL81344.1_PAL_Pyrus_communis AEQHNQDVNSLGLISSRKTAEAVDILKLMSSTFLVALCQSVDLRHLEENLRNTVKNTVSQ
QIC54077.1_PAL_Paeonia_lactiflora AEQHNQDVNSLGLISSRKTEEAVDILKLMSSTFLVALCQAIDLRHLEENMRNTVKNTVSQ
NP_181241.1_PAL1_Arabidopsis_thaliana AEQHNQDVNSLGLISSRKTSEAVDILKLMSTTFLVAICQAVDLRHLEENLRQTVKNTVSQ
NP_190894.1_PAL2_Arabidopsis_thaliana AEQHNQDVNSLGLISSRKTSEAVDILKLMSTTFLVGICQAVDLRHLEENLRQTVKNTVSQ
NP_187645.1_PAL4_Arabidopsis_thaliana AEQHNQDVNSLGLISSRKTAEAVDILKLMSTTYLVALCQAVDLRHLEENLKKAVKSAVSQ
sp|Q42667.1_PAL_Citrus_limon AEQHNQDVNSLGLNSSRKTAEAVDILKLMSSTFLVALCQAIDLRHLEENLKNTVKNTVSQ
CAB42793.1_PAL_Citrus_clementina_x_Citrus_r AEQHNQDVNSLGLISSRKTAEAVDILKLMSSTFLVALCQAIDLRHLEENLKHTVKDTVSQ
CAB42794.1_PAL_Citrus_clementina_x_Citrus_r AEQHNQDVNSLGLISSRKTAEAVDILKLMSSTFLVALCQAIDLRHLEENLKHTVKNTVSQ
sp|P27991.1_PAL_Glycine_max AEQHNQDVNSLGLISSRKTNEAIEILKLMSSTFLIALCQAIDLRHLEENLKNSVKNTVSQ
sp|O49835.1_PAL1_Lithospermum_erythrorhizon AEQHNQDVNSLGLISSRKTSEAVEILKLMSSSFLVALFQAVDLRHIEENVRLAVKNTVSQ
sp|P35511.1_PAL1_Lycopersicon_esculentum AEQHNQDVNSLGLISARKTAKAVDILKIMSSTYLVALCQAIDLRHLEENLKSVVKNTVSQ
sp|P26600.1_PAL5_Lycopersicon_esculentum AEQHNQDVNSLGLISARKTAEAVDILKLMSSTYLVALCQAIDLRHLEENLKNAVKNTVSQ
sp|P25872.1_PAL1_Nicotiana_tabacum AEQHNQDVNSLGLISARKTAEAVDILKLMSSTYLVALCQAIDLRHLEENLKNAVKNTVSQ
sp|P35513.2_PAL2_Nicotiana_tabacum AEQHNQDVNSLDLISARKTAEAVDILKLMSSTYLVALCQAIDLRHLEENLRNAVKNTVSQ
sp|P45733.1_PAL3_Nicotiana_tabacum AEQHNQDVNSLGLISARKTAEAVDILKLMSSTYLVALCQAIDLRHLEENLRNAVKNTVSQ
sp|P19142.1_PAL2_Phaseolus_vulgaris AEQHNQDVNSLGLISSRKTNEALEILKLMSSTFLVALCQAIDLRHLEENLKNTVKNVVSQ
sp|Q01861.1_PAL1_Pisum_sativum AEQHNQDVNSLGLISSRKTYEAIEILQLMSSTFLIALCQAVDLRHLEENLKNSVKNIVSQ
sp|Q04593.1_PAL2_Pisum_sativum AEQHNQDVNSLGLISSRKTYEAIEILQLMSSTFLIALCQAVDLRHLEENLKNSVKNMVSH
sp|P45731.1_PAL1_Populus_sieboldii_x_Populu AEQHNQDVNSLGLISSRKTAEAVDILKLMSTTFLVGLCQAVDLRHIEENLKNTVKNTVSQ
sp|P45730.1_PAL_Populus_trichocarpa AEQHNQDVNSLGLISSRKTAEAVDILKLMSTTFLVALCQAIDLRHLEENLKSAVKNTVSQ
sp|Q43052.1_PAL2_Populus_sieboldii_x_Populu AEQHNQDVNSLGLISSRKTAEAIDILKLMSTTFLVGLCHSVDLRHIEENLKNTVKISVSQ
AAF40223.1_PAL1_Rubus_idaeus AEQHNQDVNSLGLISSRKTSEAVDILKLMSSTFLVALCQAIDLRHLEENLKIVVKTTVSN
AAF40224.1_PAL2_Rubus_idaeus AEQHNQDVNSLGLISSRKTAEAVDILKLMSSTFLVALCQAIDLRHLEENLKSTVKNTVSQ
AAC78457.1_PAL_Prunus_avium AEQHNQDVNSLGLISSRKTAEAVDILKLMSSTFLVALCQAIDLRHLEENLRNTVKNTVSQ
sp|P31425.1_PAL1_Solanum_tuberosum AEQHNQDVNSLGLISARKTAEAVDILKLMSSTYLVALCQAIDLRHLEENLKSVVKNTVSQ
sp|A2X7F7.1_PAL2_Oryza_sativa AEQHNQDVNSLGLISSRKTAEAIDILKLMSSTFLIALCQAVDLRHIEENVKSAVKSCVMT
sp|P52777.1_PAL_Pinus_taeda AEQHNQDVNSLGLVSARKSAEAIDILKLMLSTYLTALCQAVDLRHLEENMLATVKQIVSQ
ABU49842.1_PAL_Ginkgo_biloba AEQHNQDVNSLGLVSARKSAEALEILKLMLSTYLIALCQAVDLRHLEENMQATVKQVVAQ
ACS28225.2_PAL_Pinus_massoniana AEQHNQDVNSLGLVSARKSAEAIDILKLMLSTYLTALCQAVDLRHLEENMLATVKQIVSQ
BAG74771.1_PAL_Ephedra_sinica AEQHNQDVNSLGMVSARKSEEALEILRLMLSTYLTAICQAIDLRHLEENMLATVKQTVAQ
ABC69916.1_PAL_Brassica_napus AEQHNQDVNSLGLISSRKTSEAVDILKLMSTTFLVGICQAIDLRHLEENLKQTVKNTVSQ
BAC56977.1_PAL_Daucus_carota AEQHNQDVNSLGLISSRKTSEAVEILKLMSTTFLVGLCQAIDLRHLEENLKSTVKNTVSQ
ABK24709.1_PAL_Picea_sitchensis AEQHNQDVNSLGLISARKSAEAVDILKLMLSTYLTALCQAVDLRHLEENMLATVKQIVSQ
AHA44840.1_PAL_Larix_kaempferi AEQHNQDVNSLGLVSARKSAEAIDILKLMLSTYLTALCQAVDLRHLEENMLATVKQIVSQ
AGC23439.1_PAL_Dendrobium_candidum AEQHNQDVNSLGLISSRKTAEAVEILKLMTSTFLVALCQAIDLRHLEENLKSAVKNTVSQ
ABI33979.1_PAL_Jatropha_curcas AEQHNQDVNSLGLISSRKTQEAIDILKLMSSTFLVALCQAIDLRHLEENLKHAVKNTVTQ
XP_002326186.1_PAL_Populus_trichocarpa AEQHNQDVNSLGLISSRKTAESVDILKLMSTTFLVALCQAIDLRHLEENLRSAVKNTVSH
XP_002519521.1_PAL_Ricinus_communis AEQHNQDVNSLGLISSRKTAEAVDILKLMSTTYLVALCQAIDLRHLEENLRQAVKNTVSQ
XP_002268732.1_PAL_Vitis_vinifera AEQHNQDVNSLGLISSRKTAEAVDILKLMSTTYLVALCQAIDLRHLEENLKSTVKKTVSH
AFG26322.1_PAL_Cinnamomum_osmophloeum AEQHNQDVNSLGLISSRKTAEAVDILKLMSATYLVGLCQAIDLRHLEENLKSTVKNTVSQ
BAG70992.1_PAL_Musa_balbisiana AEQHNQDVNSLGLISARKTAEAVDILKLMTTTYLVALCQAVDLRHLEENLKSAVKSTVSQ
AAP34199.1_PAL_Phalaenopsis_x_Doritaenopsis AEQHNQDVNSLGLISSRKTAESVDILKLMTTTFLVALCQAVDLRHLEENLKNAVKNTVSQ
ACM61988.1_PAL_Lycoris_radiata AEQHNQDVNSLGLISARKTEEAVQILKLMSTTFLVGLCQAIDLRHLEENLKSTVKTTISQ
ABN79671.2_PAL_Rudbeckia_hirta AEQHNQDVNSLGLISARKTAEAVEILKLMSSTYLVALCQSVDLRHLEENMKSTVKNTVSQ
ACJ66297.1_PAL4_Nicotiana_tabacum AEQHNQDVNSLGLISARKTAEAVDILKLMSSTYLVALCQAIDLRHLEENLKNAVKNTVSQ
ABG75911.1_PAL2_Nicotiana_attenuata AEQHNQDVNSLGLISARKTAEAVDILKLMSSTYLVALCQAIDLRHLEENLKNAVKNTVSQ
BAL49995.1_PAL_Eucalyptus_robusta AEQHNQDVNSLGLISSRKTAEAIDILKLMSSTFLVALCQAVDLRHLEENLKSTVKNTVGR
ACM61988.1_PAL_Lycoris_radiata AEQHNQDVNSLGLISARKTEEAVQILKLMSTTFLVGLCQAIDLRHLEENLKSTVKTTISQ
BAM28963.1_PAL_Lilium_hybrid_division_I AEQHNQDVNSLGLISSRKTAEAVEILKLMSSTFLVALCQAVDLRHLEENLKAAVKNTVSQ
acc

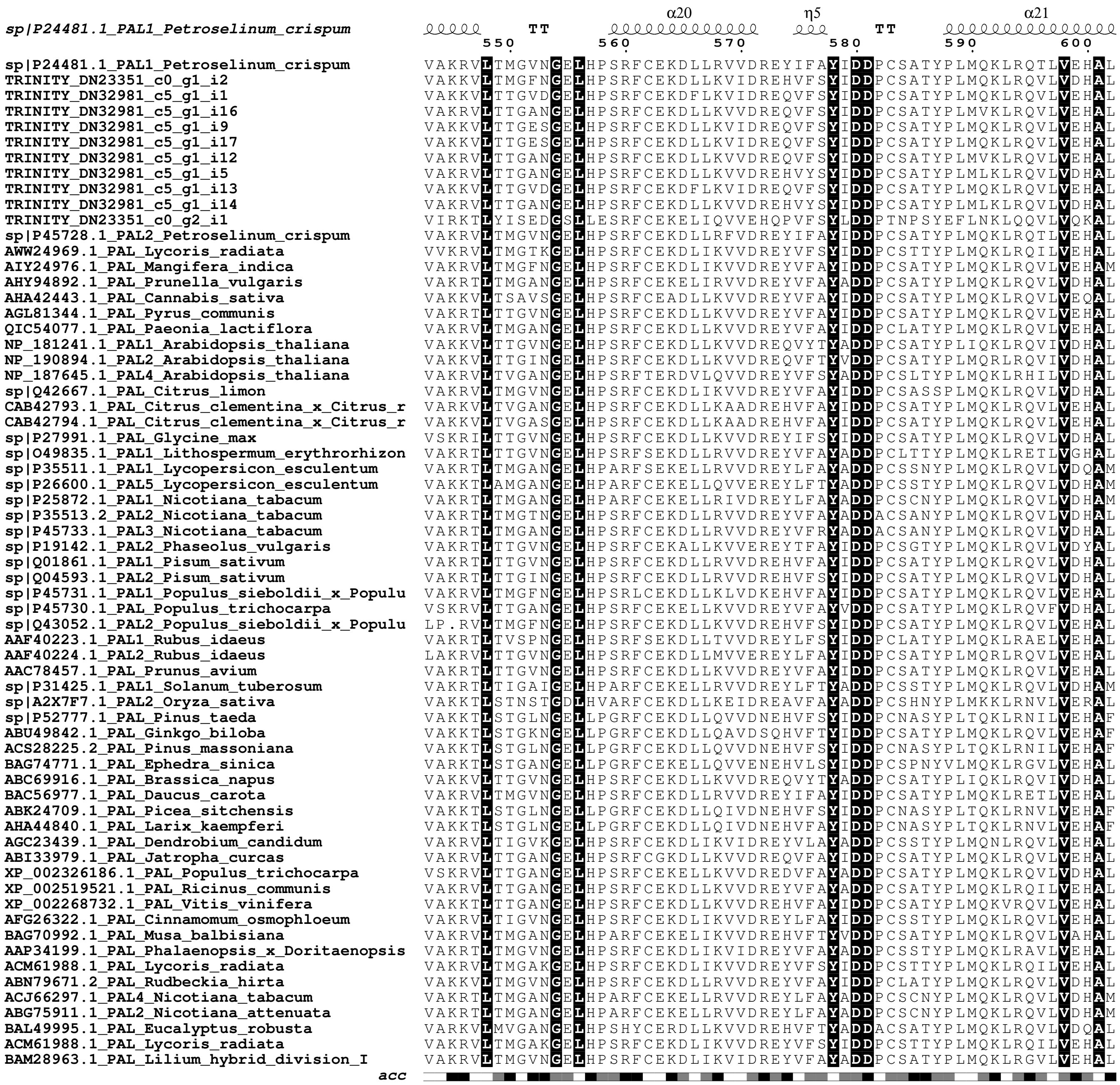
sp|P24481.1_PAL1_Petroselinum_crispum
α20
η5
α21
TT
TT
550
560
570
580
590
600
sp|P24481.1_PAL1_Petroselinum_crispum
TRINITY_DN23351_c0_g1_i2
TRINITY_DN32981_c5_g1_i1
TRINITY_DN32981_c5_g1_i16
TRINITY_DN32981_c5_g1_i9
TRINITY_DN32981_c5_g1_i17
TRINITY_DN32981_c5_g1_i12
TRINITY_DN32981_c5_g1_i5
TRINITY_DN32981_c5_g1_i13
TRINITY_DN32981_c5_g1_i14
TRINITY_DN23351_c0_g2_i1
sp|P45728.1_PAL2_Petroselinum_crispum
AWW24969.1_PAL_Lycoris_radiata
AIY24976.1_PAL_Mangifera_indica
AHY94892.1_PAL_Prunella_vulgaris
AHA42443.1_PAL_Cannabis_sativa
AGL81344.1_PAL_Pyrus_communis
QIC54077.1_PAL_Paeonia_lactiflora
NP_181241.1_PAL1_Arabidopsis_thaliana
NP_190894.1_PAL2_Arabidopsis_thaliana
NP_187645.1_PAL4_Arabidopsis_thaliana
sp|Q42667.1_PAL_Citrus_limon
CAB42793.1_PAL_Citrus_clementina_x_Citrus_r
CAB42794.1_PAL_Citrus_clementina_x_Citrus_r
sp|P27991.1_PAL_Glycine_max
sp|O49835.1_PAL1_Lithospermum_erythrorhizon
sp|P35511.1_PAL1_Lycopersicon_esculentum
sp|P26600.1_PAL5_Lycopersicon_esculentum
sp|P25872.1_PAL1_Nicotiana_tabacum
sp|P35513.2_PAL2_Nicotiana_tabacum
sp|P45733.1_PAL3_Nicotiana_tabacum
sp|P19142.1_PAL2_Phaseolus_vulgaris
sp|Q01861.1_PAL1_Pisum_sativum
sp|Q04593.1_PAL2_Pisum_sativum
sp|P45731.1_PAL1_Populus_sieboldii_x_Populu
sp|P45730.1_PAL_Populus_trichocarpa
sp|Q43052.1_PAL2_Populus_sieboldii_x_Populu
AAF40223.1_PAL1_Rubus_idaeus
AAF40224.1_PAL2_Rubus_idaeus
AAC78457.1_PAL_Prunus_avium
sp|P31425.1_PAL1_Solanum_tuberosum
sp|A2X7F7.1_PAL2_Oryza_sativa
sp|P52777.1_PAL_Pinus_taeda
ABU49842.1_PAL_Ginkgo_biloba
ACS28225.2_PAL_Pinus_massoniana
BAG74771.1_PAL_Ephedra_sinica
ABC69916.1_PAL_Brassica_napus
BAC56977.1_PAL_Daucus_carota
ABK24709.1_PAL_Picea_sitchensis
AHA44840.1_PAL_Larix_kaempferi
AGC23439.1_PAL_Dendrobium_candidum
ABI33979.1_PAL_Jatropha_curcas
XP_002326186.1_PAL_Populus_trichocarpa
XP_002519521.1_PAL_Ricinus_communis
XP_002268732.1_PAL_Vitis_vinifera
AFG26322.1_PAL_Cinnamomum_osmophloeum
BAG70992.1_PAL_Musa_balbisiana
AAP34199.1_PAL_Phalaenopsis_x_Doritaenopsis
ACM61988.1_PAL_Lycoris_radiata
ABN79671.2_PAL_Rudbeckia_hirta
ACJ66297.1_PAL4_Nicotiana_tabacum
ABG75911.1_PAL2_Nicotiana_attenuata
BAL49995.1_PAL_Eucalyptus_robusta
ACM61988.1_PAL_Lycoris_radiata
BAM28963.1_PAL_Lilium_hybrid_division_I
acc

```
                                              α22  η6          α23                        η7
sp|P24481.1_PAL1_Petroselinum_crispum
                                                  610       620       630       640       650
sp|P24481.1_PAL1_Petroselinum_crispum        KN...GDNERNLSTSIFQKIATFEDELKALLPKEVESARAALESGNPA.IPNRIEECRSY
TRINITY_DN23351_c0_g1_i2                     MN...GEKEKNSATSIFQKIGIFEDEIKTLLPKDVESARVNFENGNAA.IPNKIKECRSY
TRINITY_DN32981_c5_g1_i1                     GN...GENEKNISTSVYQKIGAFEEELKALLPKEVESAREAYETGNVA.VGNKIKECRSY
TRINITY_DN32981_c5_g1_i16                    GN...GENEKNVSTSVYQKIGAFEEELKALLPKEVESAREAFESGNTT.VDNKIKECRSY
TRINITY_DN32981_c5_g1_i9                     EN...GENEKNISTSVYQKIGAFEEELKALLPKEVETAREAYESGNVA.VGNKIKECRSY
TRINITY_DN32981_c5_g1_i17                    EN...GENEKNISTSVYQKIGAFEEELKALLPKEVETAREAYESGNVA.VGNKIKECRSY
TRINITY_DN32981_c5_g1_i12                    GN...GENEKNVSTSVYQKIGAFEEELKALLPKEVESAREAFESGNTT.VDNKIKECRSY
TRINITY_DN32981_c5_g1_i5                     AN...GENEKNVSNSVFQKIGAFEEELKALLPKEAEGVRSAFASGNVA.VANKIKECRSY
TRINITY_DN32981_c5_g1_i13                    GN...GENEKNISTSVYQKIGAFEEELKALLPKEVESAREAYETGNVA.VGNKIKECRSY
TRINITY_DN32981_c5_g1_i14                    AN...GENEKNVSNSVFQKIGAFEEELKALLPKEAEGVRSAFASGNVA.VANKIKECRSY
TRINITY_DN23351_c0_g2_i1                     KDPKSGNSDENGYLT.FKRIPVFMMELKARLEEDVPKARERFDNGEFG.VENRIMKCRTS
sp|P45728.1_PAL2_Petroselinum_crispum        KN...GDNERNMNTSIFQKIATFEDELKALLPKEVESARAALESGNPA.IPNRIEECRSY
AWW24969.1_PAL_Lycoris_radiata               NN...GEKEKDANTSIFQKISAFEEELNVVLPKEVENAWVAYENGTSA.IKNRIEDCRSY
AIY24976.1_PAL_Mangifera_indica              AN...GEREKKSSTSIFQKIGAFEEELKTLLPKEVESTRIEIENGNAA.VPNKIKECRSY
AHY94892.1_PAL_Prunella_vulgaris             KN...GDGEKNVSTSIFHKIEAFEEELKALLPKEVESARIALESGAPA.VANRIAECRSY
AHA42443.1_PAL_Cannabis_sativa               VN...GETEKNPNTSIFQKIGAFEEELKNILPKEVESSRVALESGNPT.LTNKIKDCRSY
AGL81344.1_PAL_Pyrus_communis                TN...GESEKNASTSIFQKIGAFEEELKTLLPKEVESARSAIESGNAA.VPNRIAECRSY
QIC54077.1_PAL_Paeonia_lactiflora            VN...GEREKNSSTSIFQKIAAFEEELMTLLPKEVDSARSEYESQSASLIPNRIKECRSY
NP_181241.1_PAL1_Arabidopsis_thaliana        IN...GESEKNAVTSIFHKIGAFEEELKAVLPKEVEAARAAYDNGTSA.IPNRIKECRSY
NP_190894.1_PAL2_Arabidopsis_thaliana        SN...GETEKNAVTSIFQKIGAFEEELKAVLPKEVEAARAAYGNGTAP.IPNRIKECRSY
NP_187645.1_PAL4_Arabidopsis_thaliana        AD...PEREANSATSVFHKIGAFEAELKLLLPKEVERVRVEYEEGTSA.IANRIKECRSY
sp|Q42667.1_PAL_Citrus_limon                 DN...GDREKNSTTSIFQKIGAFEDELKTLLPKEVEIARTELESGNAA.IPNRIKECRSY
CAB42793.1_PAL_Citrus_clementina_x_Citrus_r  NN...GENEKNANSSIFQKIAAFEEELKAVLPKEVENARQTVENGNPT.IPNRIKECRSY
CAB42794.1_PAL_Citrus_clementina_x_Citrus_r  NN...GENEKTANSSIFQKIAAFEEELKTVLPKEVENARQTVENGSPT.IPNRIKECRSY
sp|P27991.1_PAL_Glycine_max                  VN...AECEKDVNSSIFQKIAIFEEELKNLLPKEVEGARAAYESGKAA.IPNKIQECRSY
sp|O49835.1_PAL1_Lithospermum_erythrorhizon  DN...GENEKDVNTSIFHKIAIFEEELKAILPKEVENARASVENGIPA.ISNRIEECRSY
sp|P35511.1_PAL1_Lycopersicon_esculentum     KN...GESEKNVNSSIFQKIGAFEDELIAVLPKEVESVRAVFESGNPL.IRNRITECRSY
sp|P26600.1_PAL5_Lycopersicon_esculentum     KN...GESEKNLNSSIFQKIVAFEDELKAVLPKEVESARAVVESGNPA.IPNRITECRSY
sp|P25872.1_PAL1_Nicotiana_tabacum           NN...GESEKNVNSSIFQKIGAFEDELKAVLPKEVESARAALESGNPA.IPNRITECRSY
sp|P35513.2_PAL2_Nicotiana_tabacum           QN...GENEKNANSSIFQKILAFEDELKAVLPKEVESARAALESGNPA.IANRIKECRSY
sp|P45733.1_PAL3_Nicotiana_tabacum           EN...GENEKNANSSIFQKILAFEGELKAVLPKEVESARISLENGNPA.IANRIKECRSY
sp|P19142.1_PAL2_Phaseolus_vulgaris          AN...GENEKNLNTSIFQKIASFEEELKTLLPKEVEGARLAYENDQCA.IPNKIKDCRSY
sp|Q01861.1_PAL1_Pisum_sativum               VN...GESEKNLNTSIFQKIATFEDELKTLLPKEVESTRAAYESGNPT.VPNKINGCRSY
sp|Q04593.1_PAL2_Pisum_sativum               VN...GESEKNLNTSIFQKIATFEDELKTLLPKEVESARGAYENGNTT.ISNKIKECRSY
sp|P45731.1_PAL1_Populus_sieboldii_x_Populu  VN...GERETNSTTSIFQKIRSFEEELKTLLPKEVESARLEVENGNPV.VPNRIKECRSY
sp|P45730.1_PAL_Populus_trichocarpa          EN...GENEKNFSTSVFQKIEAFEEELKALLPKEVESARAAYDSGNSA.IDNKIKECRSY
sp|Q43052.1_PAL2_Populus_sieboldii_x_Populu  VN...GEKVRNSTTSIFQKIGSFEEELKTLLPKEVESARLEVENGNPA.IPNRIKECRSY
AAF40223.1_PAL1_Rubus_idaeus                 KN...GERERSANTSIFHKIAAFEEELKTILPKEVDNARIEIENGKSE.IPNRIKECRSY
AAF40224.1_PAL2_Rubus_idaeus                 TN...GENEKNASTSIFQKITAFEEELKTILPKEVESARAAYESGNAA.IPNRIVECRSY
AAC78457.1_PAL_Prunus_avium                  TN...GENEKNASTSIFQKIVAFEEELKVLLPKEVDSARAALDSGSAG.VPNRITECRSY
sp|P31425.1_PAL1_Solanum_tuberosum           KN...GESEKNINSSIFQKIGAFEDELNAVLPKEVESARALLESGNPS.IPNRITECRSY
sp|A2X7F7.1_PAL2_Oryza_sativa                AN...GAAEFNADTSVFAKVAQFEEELRATLPGAIEAARAAVENGTAA.IPSRITECRSY
sp|P52777.1_PAL_Pinus_taeda                  KN...AEGEKDPNTSIFNKIPVFEAELKAQLEPQVSLARESYDKGTSP.LPDRIQECRSY
ABU49842.1_PAL_Ginkgo_biloba                 KN...TESEKDPNTSIFNKINLFEQELKSQLESEVDLTRANYDKGNSA.VANRIHDCRSY
ACS28225.2_PAL_Pinus_massoniana              KN...AEGEKDPNTSIFNKIPLFEAELKAQLELQVSLARESYDKGTSP.LPNRIQECRSY
BAG74771.1_PAL_Ephedra_sinica                KN...REAEKDLRTSIFHRIEEFETELKVQLESQATIVRANFDNGVTA.LPNRIKECRSY
ABC69916.1_PAL_Brassica_napus                VN...GESEKNAMTSIFHKIGAFEEELKAVLPKEVEAARAAYDNGTAA.IPNRIKECRSY
BAC56977.1_PAL_Daucus_carota                 NN...GDKERNLSTSIFQKIAAFEDELKALLPKEVESARAAVESGNPA.IPNRIKECRSY
ABK24709.1_PAL_Picea_sitchensis              KN...TDGEKDPNTSIFNKITLFEAELKTQLELQVNLARESYDKGISP.LPNRIQECRSY
AHA44840.1_PAL_Larix_kaempferi               KN...TDGEKDPNTSIFNKITLFEAELKTQLESQVNLARDSYDKGISP.LPNRIQECRSY
AGC23439.1_PAL_Dendrobium_candidum           NN...GEKEKEADSSIFQKITAFEEELKAVLPKEVEATRLAFENGTST.IGNRIKDCRSY
ABI33979.1_PAL_Jatropha_curcas               AN...GENEKNASTSVFQKIGAFEEELKTLLPKEVESAREAYESGSAA.IGNKIKECRSY
XP_002326186.1_PAL_Populus_trichocarpa       AN...GENEKNTSTSVFQKITAFEEELKALLPKEVESARAAYDSGNSA.IENKIKECRSY
XP_002519521.1_PAL_Ricinus_communis          AN...GENEKNAGTSVFQKISAFEEELKTLLPKEVESVRIAYESGNPA.TANRIKECRSY
XP_002268732.1_PAL_Vitis_vinifera            NN...GENEKNGSTSIFQKIGAFEEELKAVLPKEVESARDGVESGNPS.IPNRIKECRSY
AFG26322.1_PAL_Cinnamomum_osmophloeum        NN...EEKEKDLNTSIFQKIAAFEEELKTHLPKEVESARAEWESGNPA.IANRIKECRSY
BAG70992.1_PAL_Musa_balbisiana               EN...GDKEKDAGSSIFQKIATFEEELTAQLPKEVEAARAAVEGGKAA.IPNRIEECRSY
AAP34199.1_PAL_Phalaenopsis_x_Doritaenopsis  NN...GDKEKTSSTSIFQKISSFEAELKAALPKEVEAARAAFENGSPA.IENRIKDCRSY
ACM61988.1_PAL_Lycoris_radiata               NN...GEKEKDANTSIFQKISAFEEELNVVLPKEVENAWVAYENGTSA.IKNRIEDCRSY
ABN79671.2_PAL_Rudbeckia_hirta               NN...GETEKNTNTSIFQKIATFEDELKAILPKEVESVRVAFENGTMS.IPNRIKACRSY
ACJ66297.1_PAL4_Nicotiana_tabacum            NN...GESEKNVNSSIFQKIGAFEDELKAVLPKEVESARAALECGNPA.IANRITECRSY
ABG75911.1_PAL2_Nicotiana_attenuata          KN...GESEKNVNSSIFQKIGAFEDELKAVLPKEVENARAALESGNPA.IANRITECRSY
BAL49995.1_PAL_Eucalyptus_robusta            VN...GESELNPSTSIFQKIVAFEEELKAQLPKDVEGIRVQYETGSLA.IPNQIKECRSY
ACM61988.1_PAL_Lycoris_radiata               NN...GEKEKDANTSIFQKISAFEEELNVVLPKEVENAWVAYENGTSA.IKNRIEDCRSY
BAM28963.1_PAL_Lilium_hybrid_division_I      VN...GEKEKEVGTSIFQKITAFEEELKATLPKEVEGVRMAFDNGTNV.IANRISECRSY
                                        acc
```

```
                                              α24                  α25             α26
sp|P24481.1_PAL1_Petroselinum_crispum      ℓℓℓℓℓℓℓℓℓ     TT  ℓℓℓℓℓℓℓℓℓℓℓℓℓ  ℓℓℓℓℓℓℓℓ
                                          660       670       680       690       700       710
sp|P24481.1_PAL1_Petroselinum_crispum     PLYKFVRKELGTEYLTGEKVTSPGEEFEKVFIAMSKGEIIDPLLECLESWNGA.......
TRINITY_DN23351_c0_g1_i2                  PLYKFVREDLSTSLLTGENLRSPGEDFDKVFEAICAGKLIDPLFECLKEWNGA.......
TRINITY_DN32981_c5_g1_i1                  PLYKFVREELGTRLLTGEKGRSPGQDFDKVFTAMCEGKIIDPMMECLKEWNGT.......
TRINITY_DN32981_c5_g1_i16                 PLYKFVRQELSTGLLTGEKGRSPGEDFDKVFTAMCQGKIIDPMMDCLKEWNGA.......
TRINITY_DN32981_c5_g1_i9                  PLYKFVREELGTGLLTGEKGRSPGQDFDKVFTAMCEGKIIDPMMECLKEWNGS.......
TRINITY_DN32981_c5_g1_i17                 PLYKFVREELGTGLLTGEKGRSPGQDFDKVFTAMCEGKIIDPMMECLKEWNGS.......
TRINITY_DN32981_c5_g1_i12                 PLYKFVRQELSTGLLTGEKGRSPGEDFDKVFTAMCQGKIIDPMMDCLKEWNGA.......
TRINITY_DN32981_c5_g1_i5                  PLYKFLREELGTGLLTGEKGRSPGEDFDKVFTAMCQGKLIDPMMDCLKEWNGA.......
TRINITY_DN32981_c5_g1_i13                 PLYKFVREELGTRLLTGEKGRSPGQDFDKVFTAMCEGKIIDPMMECLKEWNGT.......
TRINITY_DN32981_c5_g1_i14                 PLYKFLREELGTGLLTGEKGR...........................................
TRINITY_DN23351_c0_g2_i1                  PIYEFVRSEVGTSLLSGAKKVSPGEDIEKVHEAINEGKLGQVLIRCLKDWKQA.......
sp|P45728.1_PAL2_Petroselinum_crispum     PLYKFVRKELGIEYLTGEKVTSPGEEFDKVFIAMSKGEIIDPLLECLESWNGA.......
AWW24969.1_PAL_Lycoris_radiata            PLYRFVREEIGTSLLTGEKVRSPGEEFDKVFNAICKGKLVDPLLECLEDWNGA.......
AIY24976.1_PAL_Mangifera_indica           PLYKFVREELGASFLTGEKVKSPGEEFDKVFSAICAGKLIDPMMECLKEWNGA.......
AHY94892.1_PAL_Prunella_vulgaris          PLYKFIREGLGTGFLTGEKAVSPGEECDKVFTALCDGLIVDPLLECLQGWNGE.......
AHA42443.1_PAL_Cannabis_sativa            PLYKFVRDNLETNLLSGENVRSPGEEFDKVFTAMCQGKIIDPLLECLSNWNGT.......
AGL81344.1_PAL_Pyrus_communis             PLYKFVREELGGEYLTGEKVRSPGEECDRVFQAICQGKIIDPILGCLEGWNGA.......
QIC54077.1_PAL_Paeonia_lactiflora         PLYKFVREELGSGLLTGDKTRSPGEDFDKVFSAMCEGKLIDPMLDCLKEWNGA.......
NP_181241.1_PAL1_Arabidopsis_thaliana     PLYRFVREELGTELLTGEKVTSPGEEFDKVFTAICEGKIIDPMMECLNEWNGA.......
NP_190894.1_PAL2_Arabidopsis_thaliana     PLYRFVREELGTKLLTGEKVVSPGEEFDKVFTAMCEGKLIDPLMDCLKEWNGA.......
NP_187645.1_PAL4_Arabidopsis_thaliana     PLYRFVRDELNTELLTGENVRSPGEEFDKVFLAISDGKLIDPLLECLKEWNGA.......
sp|Q42667.1_PAL_Citrus_limon              PLYKIVREDIGTSLLTGEKVRSPGEEFDKVFTAMCEGKLIDPMLECLKEWNGA.......
CAB42793.1_PAL_Citrus_clementina_x_Citrus_r PLYRLVREELGTNFLTGEKVTSPGEKFDKVFTAMCQGKIIDPMLECLREWNGA.......
CAB42794.1_PAL_Citrus_clementina_x_Citrus_r PLYRLVREGLGSNFLTGEKVTSPGEEFDKVFTAMCQGKIIDPMLECLREWNGA.......
sp|P27991.1_PAL_Glycine_max               PLYKFVREELGTGLLTGEKVRSPGEEFDKLFTAMCQGKIIDPLMECLGEWNGA.......
sp|O49835.1_PAL1_Lithospermum_erythrorhizon PLYKFVREELGTELLTGEKVRSPGEELDKVFTAMCEGKLVDPLLACLEAWNGA.......
sp|P35511.1_PAL1_Lycopersicon_esculentum  PLYRLVREELGTELLTGEKVRSPGEEIDKVFTAICNGQIIDPLLECLKSWNGA.......
sp|P26600.1_PAL5_Lycopersicon_esculentum  PLYRLVRQEVGTELLTGEKVRSPGEEIDKVFTAFCNGQIIDPLLECLKSWNGA.......
sp|P25872.1_PAL1_Nicotiana_tabacum        PLYRFVRKELGTELLTGEKVRSPGEECDKVFTAMCNGQIIDPMLECLKSWNGA.......
sp|P35513.2_PAL2_Nicotiana_tabacum        PLYRFVRGELGAELLTGEKVRSPGEECDKVFTAMCNGQIIDSLLECLKEWNGA.......
sp|P45733.1_PAL3_Nicotiana_tabacum        PLYRFVREELGAELLTGEKVRSPGEECDKVFTAMCNGQIIDSLLECLKEWNGA.......
sp|P19142.1_PAL2_Phaseolus_vulgaris       PLYKFVREELGTSLLTGEKVISPGEECDKVFSAMCQGKIIDPLLECLGEWNGA.......
sp|Q01861.1_PAL1_Pisum_sativum            PLYRFVRQELGTGLLTGEKVISPGEECDKLFTAICQGKIIDPLLQCLGDWNGA.......
sp|Q04593.1_PAL2_Pisum_sativum            PLYKFVREELGTSLLTGEKVISPGEECDKLFTAICQGKIIDPLLECLGDWNGA.......
sp|P45731.1_PAL1_Populus_sieboldii_x_Populu PLYKFVREELGTSLLTGEKVKSPGEDFDKVFTAICAGKLMDPLLECLKEWNGA.......
sp|P45730.1_PAL_Populus_trichocarpa       PLYKFVREELGTVLLTGEKVQSPGEEFDKVFTAMCQGKIIDPMLECLGEWNGS.......
sp|Q43052.1_PAL2_Populus_sieboldii_x_Populu PLYKFVREELGTSLLTGEKVKSPGEEFDKVFTAICAGKLIDPLLECLKEWDGA.......
AAF40223.1_PAL1_Rubus_idaeus              PLYRFVREELGTSLLTGEKIKSPGEECYKVFNAICAGKLVDPLLECLKEWNGA.......
AAF40224.1_PAL2_Rubus_idaeus              PLYKFVREELGGEFLTGEKVRSPGEECDKVFTAMCQGNIIDPILDCLSGWNGE.......
AAC78457.1_PAL_Prunus_avium               PLYKFVREELGAEYLTGEKVRSPGEECDKVFTAICEGKIIDPILDCLEGWNGA.......
sp|P31425.1_PAL1_Solanum_tuberosum        PLYRLVRQELGTELLTGEKVRSPGEEIEKVFTAMCNGQINDPLLECLKSWNGE.......
sp|A2X7F7.1_PAL2_Oryza_sativa             PLYRFVREELGTKYLTGEKTRSPGEELNKVLVAINEGKHIDPLLECLKEWNGE.......
sp|P52777.1_PAL_Pinus_taeda               PLYEFVRNQLGTKLLSGTRTISPGEVIEVVYDAISEDKVIVPLFKCLDGWKGTLAHSEIN
ABU49842.1_PAL_Ginkgo_biloba              PLYEFVRTQLGTKLLSGTRATSPGEYIEKVFDAICEDKIIEPIIKCLDGWKGT.......
ACS28225.2_PAL_Pinus_massoniana           PLYEFVRNQLGTKLLSGTRTTSPGEVIEVVYDAISEDKVIVPLFKCLDGWKGT.......
BAG74771.1_PAL_Ephedra_sinica             PLYSFVRETLGTQLLSGDRDISPGEDIEVVYKAVKANDIIVPLFKCLHGWKGT.......
ABC69916.1_PAL_Brassica_napus             PLYRFVREELGTELLTGEKATSPGEEFDKVFTAICEGKIIDPLMECLDEWNGA.......
BAC56977.1_PAL_Daucus_carota              PLYKFVREELGTEYLTGEKVTSPGEEFDKVFTAMTKGEIIDPLLECLQSWNGA.......
ABK24709.1_PAL_Picea_sitchensis           PLYEFVRTQLGTKLLSGTRTTSPGEVIELVYDAISEDKIIGPLLKCVEGWKAT.......
AHA44840.1_PAL_Larix_kaempferi            PLYEFVRIQLGTKLLSGARTTSPGEVIELVYDAISEDKIIGPLFKCLDGWKAT.......
AGC23439.1_PAL_Dendrobium_candidum        PIYRFVREELGTSFLTGEEVRSPGEEFDKVFNAICEGKAIDPLLECLQEWDGA.......
ABI33979.1_PAL_Jatropha_curcas            PLYKFVREELGSGLLTGEKVRSPGEEFDKVFTAMCEGKIIDPMMECLKEWNGA.......
XP_002326186.1_PAL_Populus_trichocarpa    PLYKFVREELGTGLLTGEKVRSPGEEFDKVFTAMCQGKIIDPMLECLGEWNGA.......
XP_002519521.1_PAL_Ricinus_communis       PLYKFVREELGTGLLTGDKVMSPGEEFDKVFTAMCQGKIIDPMMDCLKEWNGA.......
XP_002268732.1_PAL_Vitis_vinifera         PLYKFVREELGTGLLTGETVRSPGEDFDKVFTAMCEGKIIDPLLDCLSAWNGA.......
AFG26322.1_PAL_Cinnamomum_osmophloeum     PLYKFVREELKTSLLTGERVRSPGEEFDKVFAAICEGKVIDPFLECLREWNGA.......
BAG70992.1_PAL_Musa_balbisiana            PLYRLVREELKTGFLTGEKVRSPGEEFDKVFDAICQGKVIDPLLECLKEWNGA.......
AAP34199.1_PAL_Phalaenopsis_x_Doritaenopsis PLYRFVK.QVGAGFLTGEKIVSPGEELDKVFNAICEGKAIDPMLDCLKEWNGA.......
ACM61988.1_PAL_Lycoris_radiata            PLYRFVREEIGTSLLTGEKVRSPGEEFDKVFNAICKGKLVDPLLECLEDWNGA.......
ABN79671.2_PAL_Rudbeckia_hirta            PLYRFVREELSGAYLTGEKVTSPGEEFDKVFTAMSKGQIIDPLLECVEGWNGA.......
ACJ66297.1_PAL4_Nicotiana_tabacum         PLYRFVRKELGTELLTGERVRSPGEECEKVFTAMCNGQIIDPMLECLKSWNGA.......
ABG75911.1_PAL2_Nicotiana_attenuata       PLYRFVRKELGTELLTGERVRSPGEECDKVFTAMCNGQIIDPMLECLKSWNGA.......
BAL49995.1_PAL_Eucalyptus_robusta         PLYKLVREELGTALLTGEGVISPGEDFDKVFTAICAGKLIDPLLECLSGWNGA.......
ACM61988.1_PAL_Lycoris_radiata            PLYRFVREEIGTSLLTGEKVRSPGEEFDKVFNAICKGKLVDPLLECLEDWNGA.......
BAM28963.1_PAL_Lilium_hybrid_division_I   PLYKFVREELGAGYLTGEKVRSPGEEFNKVFIAMSQWKVIDPLLECLSDWNGA.......
acc
```

*sp|P24481.1_PAL1_Petroselinum_crispum*

```
sp|P24481.1_PAL1_Petroselinum_crispum         ...PLPI...C......................
TRINITY_DN23351_c0_g1_i2                      ...PLPI...C......................
TRINITY_DN32981_c5_g1_i1                      ...PIPI...C......................
TRINITY_DN32981_c5_g1_i16                     ...PLPL...S......................
TRINITY_DN32981_c5_g1_i9                      ...PLPI...C......................
TRINITY_DN32981_c5_g1_i17                     ...PLPI...C......................
TRINITY_DN32981_c5_g1_i12                     ...PLPL...S......................
TRINITY_DN32981_c5_g1_i5                      ...PVPI...C......................
TRINITY_DN32981_c5_g1_i13                     ...PIPI...C......................
TRINITY_DN32981_c5_g1_i14                     .................................
TRINITY_DN23351_c0_g2_i1                      .................................
sp|P45728.1_PAL2_Petroselinum_crispum         ...PLPI...C......................
AWW24969.1_PAL_Lycoris_radiata                ...PLPI...C......................
AIY24976.1_PAL_Mangifera_indica               ...PLPL...C......................
AHY94892.1_PAL_Prunella_vulgaris              ...PLPI...C......................
AHA42443.1_PAL_Cannabis_sativa                ...PLPL...C......................
AGL81344.1_PAL_Pyrus_communis                 ...PLPI...C......................
QIC54077.1_PAL_Paeonia_lactiflora             ...PLPI...C......................
NP_181241.1_PAL1_Arabidopsis_thaliana         ...PIPI...C......................
NP_190894.1_PAL2_Arabidopsis_thaliana         ...PIPI...C......................
NP_187645.1_PAL4_Arabidopsis_thaliana         ...PVSI...C......................
sp|Q42667.1_PAL_Citrus_limon                  ...PLPI...CQN....................
CAB42793.1_PAL_Citrus_clementina_x_Citrus_r   ...PLPI...C......................
CAB42794.1_PAL_Citrus_clementina_x_Citrus_r   ...PLPI...C......................
sp|P27991.1_PAL_Glycine_max                   ...PLPI...S......................
sp|O49835.1_PAL1_Lithospermum_erythrorhizon   ...PLPI...C......................
sp|P35511.1_PAL1_Lycopersicon_esculentum      ...PLPI...C......................
sp|P26600.1_PAL5_Lycopersicon_esculentum      ...PIPI...C......................
sp|P25872.1_PAL1_Nicotiana_tabacum            ...PLPI...C......................
sp|P35513.2_PAL2_Nicotiana_tabacum            ...PLPI...C......................
sp|P45733.1_PAL3_Nicotiana_tabacum            ...PLPI...C......................
sp|P19142.1_PAL2_Phaseolus_vulgaris           ...PLPI...C......................
sp|Q01861.1_PAL1_Pisum_sativum                ...PLPI...S......................
sp|Q04593.1_PAL2_Pisum_sativum                ...PLPI...S......................
sp|P45731.1_PAL1_Populus_sieboldii_x_Populu   ...PLPI...C......................
sp|P45730.1_PAL_Populus_trichocarpa           ...PLPI...C......................
sp|Q43052.1_PAL2_Populus_sieboldii_x_Populu   ...PLPI...C......................
AAF40223.1_PAL1_Rubus_idaeus                  ...PLPI...S......................
AAF40224.1_PAL2_Rubus_idaeus                  ...PLPI...C......................
AAC78457.1_PAL_Prunus_avium                   ...PLPI...C......................
sp|P31425.1_PAL1_Solanum_tuberosum            ...PLPI...C......................
sp|A2X7F7.1_PAL2_Oryza_sativa                 ...PLPI...C......................
sp|P52777.1_PAL_Pinus_taeda                   NLPRSPLYNDCYDLSPRMLLLMLLFSDPEFDWS
ABU49842.1_PAL_Ginkgo_biloba                  ...PGPF..........................
ACS28225.2_PAL_Pinus_massoniana               ...PGPF..........................
BAG74771.1_PAL_Ephedra_sinica                 ...PGPF..........................
ABC69916.1_PAL_Brassica_napus                 ...PIPI...C......................
BAC56977.1_PAL_Daucus_carota                  ...PLPI...C......................
ABK24709.1_PAL_Picea_sitchensis               ...PGPF..........................
AHA44840.1_PAL_Larix_kaempferi                ...PGPF..........................
AGC23439.1_PAL_Dendrobium_candidum            ...PLPI...C......................
ABI33979.1_PAL_Jatropha_curcas                ...PLPI...C......................
XP_002326186.1_PAL_Populus_trichocarpa        ...PLPI...C......................
XP_002519521.1_PAL_Ricinus_communis           ...PLPI...C......................
XP_002268732.1_PAL_Vitis_vinifera             ...PLPI...C......................
AFG26322.1_PAL_Cinnamomum_osmophloeum         ...PLPI...C......................
BAG70992.1_PAL_Musa_balbisiana                ...PLPI...C......................
AAP34199.1_PAL_Phalaenopsis_x_Doritaenopsis   ...PLPI...C......................
ACM61988.1_PAL_Lycoris_radiata                ...PLPI...C......................
ABN79671.2_PAL_Rudbeckia_hirta                ...PLPI...C......................
ACJ66297.1_PAL4_Nicotiana_tabacum             ...PLPI...C......................
ABG75911.1_PAL2_Nicotiana_attenuata           ...PLPI...C......................
BAL49995.1_PAL_Eucalyptus_robusta             ...PLPI...S......................
ACM61988.1_PAL_Lycoris_radiata                ...PLPI...C......................
BAM28963.1_PAL_Lilium_hybrid_division_I       ...PLPI...C......................
                                        acc
```

***AAK54447.1_C4H_Sorghum_bicolor***

```
                                                               1                                 10
                                                               .                                  .
AAK54447.1_C4H_Sorghum_bicolor                  ....................MDL.........................VLLEKALLGLFA
TRINITY_DN32464_c6_g3_i2                        ....................MDL.........................ILLEKTLLGLFA
TRINITY_DN15593_c0_g1_i1                        ...................................MP...........LSIAKSI..IFG
NP_180607.1_C4H_Arabidopsis_thaliana            ....................MDL.........................LLLEKSLIAVFV
ACA25599.1_C4H_Parthenocissus_henryana          ....................MDL.........................ILIEKALLAVFC
CAK95273.1_C4H_Cucumis_sativus                  ....................MDL.........................LLLEKTLLGLFL
QIC52990.1_C4H_Solanum_nigrum                   ....................MDL.........................LLLEKTLIGLFF
AHY94893.1_C4H_Prunella_vulgaris                ....................MDL.........................LLVEKTLIGLFF
AWW24970.1_C4H_Lycoris_radiata                  ....................MDL.........................ILLEKSLLAVFF
ASA39653.1_C4H_Marchantia_emarginata            ...................MSEL........LEKVL...PGSSAA...FTVQNFLYALLA
ASA39650.1_C4H_Conocephalum_japonicum           ...................MTEL........MEKVL...PGSKAN...FTFQNVLLALLA
ASA39648.1_C4H_Marchantia_paleacea              ...................MVDQ........LERLL...PGSSAA...LTMQNVLYGLLA
ASA39646.1_C4H_Plagiochasma_appendiculatum      MGTGFSTMAVCLIAAGMSKVTAE........AEEVVAAPSGSNSA...WTLQNVLYGLLA
AFX98061.1_C4H_Cunninghamia_lanceolata          ...................MADV........QQGSL............AVVEKSLVALFV
AFZ78542.1_C4H_Populus_tomentosa                ....................MDL.........................LLLEKTLLGSFV
AUG71936.1_C4H_Narcissus_pseudonarcissus        ....................MDL.........................ILLEKSLLAVFF
AEA72281.1_C4H_Angelica_gigas                   ....................MDF.........................LLLEKALLGLFI
ADO24190.1_C4H_Allium_sativum                   ....................MEL.........................LLLEKTLLSLFF
ADN32769.1_C4H_Scutellaria_baicalensis          ....................MDL.........................LLLEKSLLGLFL
AIS92509.1_C4H_Epimedium_sagittatum             ....................MDL.........................LLLEKSLIASFI
AGP25594.1_C4H_Aquilaria_sinensis               ....................MDL.........................LLLEKTLLALFA
AAG50231.1_C4H_Populus_trichocarpa_x_Populus    ....................MDL.........................LLLEKTLLGSFV
sp|Q43054.1_C4H_Populus_sieboldii_x_Populus_    ....................MDL.........................LLLEKTLLGSFV
AAG10197.1_C4H_Gossypium_arboreum               ....................MDL.........................LFLEKVLISLFF
AAG10196.1_C4H_Gossypium_arboreum               ....................MDL.........................LFLEKALLGLFV
sp|Q43033.1_C4H_Petroselinum_crispum            ...................MMDF.........................VLLEKALLGLFI
CAC35977.1_C4H_Ruta_graveolens                  ....................MDL.........................LLLEKALLGLFA
AAF66066.2_C4H_Citrus_sinensis                  ....................MDLNGWCNSGNQNMCCCQ.SYVKRGYDRVLSFNGL.....
AAK57011.1_C4H_Citrus_x_paradisi                ....................MDL.........................LLLEKTLLALFA
sp|P48522.1_C4H_Catharanthus_roseus             ....................MDL.........................LLLEKTLLGLFA
BAB71717.1_C4H_Lithospermum_erythrorhizon       ....................MDL.........................LLLEKVLIGLFI
BAB71716.1_C4H_Lithospermum_erythrorhizon       ....................MDL.........................LLLEKALIGLFF
AAC35857.1_C4H_Capsicum_chinense                ............................................................
AAG43824.1_C4H_Capsicum_annuum                  ....................MDL.........................LLLEKTLVGLFF
sp|Q43240.1_C4H_Zinnia_elegans                  ....................MDL.........................LLVEKTLLALFA
sp|Q04468.1_C4H_Helianthus_tuberosus            ....................MDL.........................LLIEKTLVALFA
sp|Q42797.1_C4H_Glycine_max                     ....................MDL.........................LLLEKTLIGLFL
sp|P37115.1_C4H_Vigna_radiata                   ....................MDL.........................LLLEKTLLGLFL
sp|Q96423.1_C4H_Glycyrrhiza_echinata            ....................MDL.........................LLLEKTLLGLFI
sp|O81928.2_C4H_Cicer_arietinum                 ....................MDL.........................LLLEKTLLALFI
sp|P37114.1_C4H_Medicago_sativa                 ....................MDL.........................LLLEKTLLALFI
sp|Q43067.2_C4H_Pisum_sativum                   ....................MDL.........................LLLEKTLLALFL
AAD11427.1_C4H_Mesembryanthemum_crystallinum    ......MA...........KMET........HSKPMS...........KKLARNL..ILL
sp|O24315_C4H_Phaseolus_vulgaris                ............................................................
AAK62344.1_C4H_Nicotiana_tabacum                ................................MKNMA...........KLLNKTIFCILF
AAK62345.1_C4H_Nicotiana_tabacum                ...................................MA...........KLLNNTIFCILF
AAG17469.1_C4H_Triticum_aestivum                ....................MDV.........................LLLEKALLGLFA
AAD23378.1_C4H_Pinus_taeda                      ....................MEI..........MTV............ASLEKGLLAIFA
                                             acc
```

***AAK54447.1_C4H_Sorghum_bicolor***

```
                                                   20                                                 30
                                                    .                                                  .
AAK54447.1_C4H_Sorghum_bicolor                  AAVLAVAVAK.....LT...................................GKRYRLPP
TRINITY_DN32464_c6_g3_i2                        AIIVAIVVSK.....LR...................................GKRFKLPP
TRINITY_DN15593_c0_g1_i1                        SLIIAIFTKF.....L..YLNI.FN..NVSFPHVWKIVLFLLPFAYLFTSICKSSKNLPP
NP_180607.1_C4H_Arabidopsis_thaliana            AVILATVISK.....LR...................................GKKLKLPP
ACA25599.1_C4H_Parthenocissus_henryana          AIILAITISK.....LL...................................GKKFKLPP
CAK95273.1_C4H_Cucumis_sativus                  SVVLAIAISK.....LR...................................GKRFKPPP
QIC52990.1_C4H_Solanum_nigrum                   AILIAIIVSK.....LR...................................SKRFKLPP
AHY94893.1_C4H_Prunella_vulgaris                AIVVAAVVSE.....LR...................................GKKFKLPP
AWW24970.1_C4H_Lycoris_radiata                  AIILSILISK.....LR...................................GKKLRLPP
ASA39653.1_C4H_Marchantia_emarginata            ATLLVILIIE.....LR...................................KEKLNLPP
ASA39650.1_C4H_Conocephalum_japonicum           ATLLVIVIVE.....LT...................................RKRLKLPP
ASA39648.1_C4H_Marchantia_paleacea              ATLLVILILE.....LR...................................KERLKLPP
ASA39646.1_C4H_Plagiochasma_appendiculatum      ATLGVILLMK.....LK...................................EKKLKLPP
AFX98061.1_C4H_Cunninghamia_lanceolata          VVVGAVLVNK.....FR...................................GKKLKLPP
AFZ78542.1_C4H_Populus_tomentosa                AVLVAILVSK.....LR...................................GKRFKLPP
AUG71936.1_C4H_Narcissus_pseudonarcissus        AVIFSIIVSK.....LR...................................GKKLKLPP
AEA72281.1_C4H_Angelica_gigas                   ATIVAITVSK.....LR...................................GKKLKLPP
ADO24190.1_C4H_Allium_sativum                   AITLAIVISK.....LR...................................GRKFKLPP
ADN32769.1_C4H_Scutellaria_baicalensis          AIIVATMVSK.....LR...................................GKKFKLPP
AIS92509.1_C4H_Epimedium_sagittatum             AIIVAIAISK.....LR...................................GKRFKLPP
AGP25594.1_C4H_Aquilaria_sinensis               TVILAIAISK.....LC...................................GKKYRLPP
AAG50231.1_C4H_Populus_trichocarpa_x_Populus    AILVAILVSK.....LR...................................GKRFKLPP
sp|Q43054.1_C4H_Populus_sieboldii_x_Populus_    AVLVAILVSK.....LR...................................GKRFKLPP
AAG10197.1_C4H_Gossypium_arboreum               TIIFAILVSK.....LR...................................GKRFKLPP
AAG10196.1_C4H_Gossypium_arboreum               AVVLAITISK.....LR...................................GKRFKLPP
sp|Q43033.1_C4H_Petroselinum_crispum            ATIVAITISK.....LR...................................GKKLKLPP
CAC35977.1_C4H_Ruta_graveolens                  AAVVAIAVSK.....LR...................................GKRFKLPP
AAF66066.2_C4H_Citrus_sinensis                  .....ITVSK.....LR...................................GKRFKLPP
AAK57011.1_C4H_Citrus_x_paradisi                AVVVAITVSK.....LR...................................GKRFKLPP
sp|P48522.1_C4H_Catharanthus_roseus             AIIVASIVSK.....LR...................................GKKFKLPP
BAB71717.1_C4H_Lithospermum_erythrorhizon       AIILSIIISK.....LG...................................GKKFKLPP
BAB71716.1_C4H_Lithospermum_erythrorhizon       SFIIAIVISK.....LR...................................GKKFKLPP
AAC35857.1_C4H_Capsicum_chinense                ............................................................
AAG43824.1_C4H_Capsicum_annuum                  AIVVAIIVSK.....LR...................................SKRFKLPP
sp|Q43240.1_C4H_Zinnia_elegans                  AIIASIFISK.....LR...................................GKRFKLPP
sp|Q04468.1_C4H_Helianthus_tuberosus            AIIGAILISK.....LR...................................GKKFKLPP
sp|Q42797.1_C4H_Glycine_max                     AAVVAIAVST.....LR...................................GRKFKLPP
sp|P37115.1_C4H_Vigna_radiata                   AAVVAIVVSK.....LR...................................GKRFKLPP
sp|Q96423.1_C4H_Glycyrrhiza_echinata            AAITAIAISK.....LR...................................GRRFKLPP
sp|O81928.2_C4H_Cicer_arietinum                 AATIAITISK.....LR...................................GKRFKLPP
sp|P37114.1_C4H_Medicago_sativa                 AATIAVTISK.....LR...................................GKRFKLPP
sp|Q43067.2_C4H_Pisum_sativum                   AAITAITISK.....LR...................................GKPFKLPP
AAD11427.1_C4H_Mesembryanthemum_crystallinum    AISIIVLTTSSSNPNFSYYLAIFLP............IIVYLVHSICFHRA.QNSGTTPP
sp|O24315_C4H_Phaseolus_vulgaris                ......MTKL.....LHSYFSIPFSPFYVSIP..IATVLFVL.IIYNFFLASKNHSSTPP
AAK62344.1_C4H_Nicotiana_tabacum                TIAFLSFAKL.....LSSYLSMPFPLKYMSL......IVPLLPLIINFLYV.KPQNNLPP
AAK62345.1_C4H_Nicotiana_tabacum                SIVFLSFAKL.....LSSYLSIPFPLEYISL......IVLLLPLIINFLCV.KPQNNLPP
AAG17469.1_C4H_Triticum_aestivum                AAVLAIAVAK.....LT...................................GKRFRLPP
AAD23378.1_C4H_Pinus_taeda                      VIVGAIFISK.....LK...................................SKKLKLPP
                                             acc
```

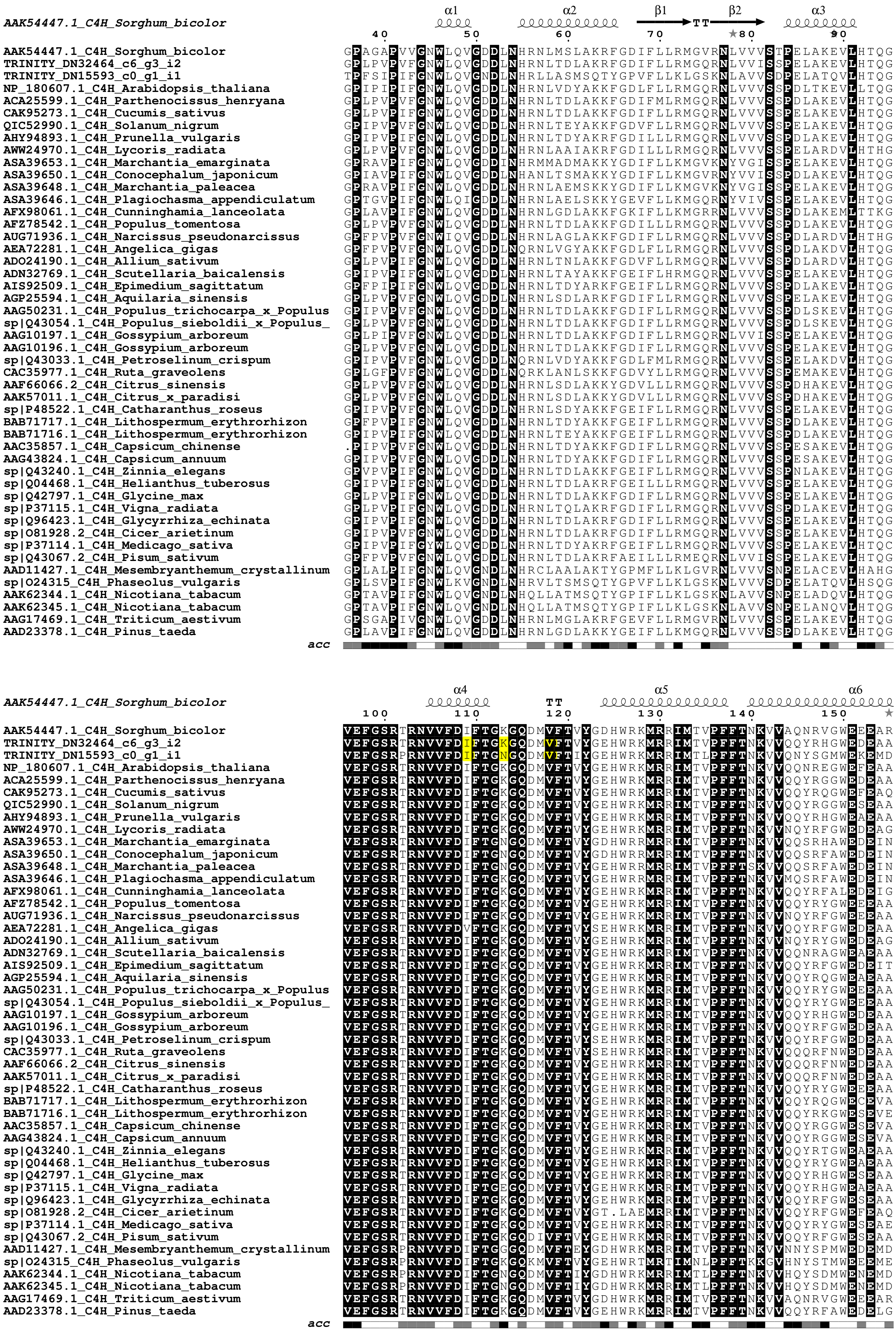

AAK54447.1_C4H_Sorghum_bicolor
α1
α2
β1
β2
α3
α4
α5
α6
TT
acc
AAK54447.1_C4H_Sorghum_bicolor
TRINITY_DN32464_c6_g3_i2
TRINITY_DN15593_c0_g1_i1
NP_180607.1_C4H_Arabidopsis_thaliana
ACA25599.1_C4H_Parthenocissus_henryana
CAK95273.1_C4H_Cucumis_sativus
QIC52990.1_C4H_Solanum_nigrum
AHY94893.1_C4H_Prunella_vulgaris
AWW24970.1_C4H_Lycoris_radiata
ASA39653.1_C4H_Marchantia_emarginata
ASA39650.1_C4H_Conocephalum_japonicum
ASA39648.1_C4H_Marchantia_paleacea
ASA39646.1_C4H_Plagiochasma_appendiculatum
AFX98061.1_C4H_Cunninghamia_lanceolata
AFZ78542.1_C4H_Populus_tomentosa
AUG71936.1_C4H_Narcissus_pseudonarcissus
AEA72281.1_C4H_Angelica_gigas
ADO24190.1_C4H_Allium_sativum
ADN32769.1_C4H_Scutellaria_baicalensis
AIS92509.1_C4H_Epimedium_sagittatum
AGP25594.1_C4H_Aquilaria_sinensis
AAG50231.1_C4H_Populus_trichocarpa_x_Populus
sp|Q43054.1_C4H_Populus_sieboldii_x_Populus_
AAG10197.1_C4H_Gossypium_arboreum
AAG10196.1_C4H_Gossypium_arboreum
sp|Q43033.1_C4H_Petroselinum_crispum
CAC35977.1_C4H_Ruta_graveolens
AAF66066.2_C4H_Citrus_sinensis
AAK57011.1_C4H_Citrus_x_paradisi
sp|P48522.1_C4H_Catharanthus_roseus
BAB71717.1_C4H_Lithospermum_erythrorhizon
BAB71716.1_C4H_Lithospermum_erythrorhizon
AAC35857.1_C4H_Capsicum_chinense
AAG43824.1_C4H_Capsicum_annuum
sp|Q43240.1_C4H_Zinnia_elegans
sp|Q04468.1_C4H_Helianthus_tuberosus
sp|Q42797.1_C4H_Glycine_max
sp|P37115.1_C4H_Vigna_radiata
sp|Q96423.1_C4H_Glycyrrhiza_echinata
sp|O81928.2_C4H_Cicer_arietinum
sp|P37114.1_C4H_Medicago_sativa
sp|Q43067.2_C4H_Pisum_sativum
AAD11427.1_C4H_Mesembryanthemum_crystallinum
sp|O24315_C4H_Phaseolus_vulgaris
AAK62344.1_C4H_Nicotiana_tabacum
AAK62345.1_C4H_Nicotiana_tabacum
AAG17469.1_C4H_Triticum_aestivum
AAD23378.1_C4H_Pinus_taeda

```
AAK54447.1_C4H_Sorghum_bicolor          η1        β3      α7                    α8
                                        160         170       180       190       200       210
AAK54447.1_C4H_Sorghum_bicolor                  LVVEDVRKDPRA....AAEGVVIRRRLQLMMYNDMFRIMFDTRFESEQDPLFNKLKALNA
TRINITY_DN32464_c6_g3_i2                        KVVEDVKKNPEA....ATRGVVLRRRLQLLMYNNMYRIMFDRRFESEEDPLFVKLKALNG
TRINITY_DN15593_c0_g1_i1                        FVVSDLSDDLKA....RNEGIVIRKRLQLMLYNIMYGMMFDARFESQEDPLFIEATMFNS
NP_180607.1_C4H_Arabidopsis_thaliana            SVVEDVKKNPDS....ATKGIVLRKRLQLMMYNNMFRIMFDRRFESEDDPLFLRLKALNG
ACA25599.1_C4H_Parthenocissus_henryana          RVVEDVKKNPEA....STNGIVLRRRLQLMMYNNMYRIMFDRRFESEEDPLFVKLKALNG
CAK95273.1_C4H_Cucumis_sativus                  SVVDDVKKNPES....ATTGIVLRRRLQLMMYNNMYRIMFDRRFESEDDPLFHKLRALNG
QIC52990.1_C4H_Solanum_nigrum                   SVVEDVKKNPES....ATNGIVLRKRLQLMMYNNMFRIMFDRRFESEDDPLFVKLRALNG
AHY94893.1_C4H_Prunella_vulgaris                AVIDDVKKNPES....ATNGIVLRRRLQLTMYNNMYRIMFDRRFESEEDPLFVKLKALNG
AWW24970.1_C4H_Lycoris_radiata                  RVVEDVKKDPMA....ATVGIVLRRRLQLMMYNNMYRIMFDRRFESEEDPLFLRLRALNG
ASA39653.1_C4H_Marchantia_emarginata            HVIKDLKADPEA....STTGCIIRRRLQLMMYNVMYRMMFDRRFESTKDELFIKLSTVNR
ASA39650.1_C4H_Conocephalum_japonicum           HVIKDLKADPAA....STSGCIIRRRLQLMMYNIMYRLMFDRRFESMSDELFVKLSTVNR
ASA39648.1_C4H_Marchantia_paleacea              HVIKDLKADPEA....STTGCIIRRRLQLMMYNVMYRMMFDRRFESVKDELFIRLSTVNR
ASA39646.1_C4H_Plagiochasma_appendiculatum      HVIKDLTASKEA....STTGVIIRRRLQLMMYNVMYRLMFDRRFETETDPLFVKLKAVNG
AFX98061.1_C4H_Cunninghamia_lanceolata          RVVEDVKSRPEA....STTGIVIRRRLQLMMYNIMYRIMFDRRFESEKDPLFLQLKALND
AFZ78542.1_C4H_Populus_tomentosa                QVVEDVKKNPEA....ATHGIVLRRRLQLMMYNNMYRIMFDRRFESEEDPLFNKLKALNG
AUG71936.1_C4H_Narcissus_pseudonarcissus        RVVEDVKKDPMA....AAEGIVLRRRLQLLMYNNMYRIMFDRRFESVDDPLFLKLTALNG
AEA72281.1_C4H_Angelica_gigas                   RVVEDVKANTEA....ATNGIVLRNRLQLLMYNNMYRIMFDRRFESVDDPLFLKLKALNG
ADO24190.1_C4H_Allium_sativum                   RVVEDVKKDTKA....ATEGIVLRKRLQLMMYNNMYRIMFDRRFESGNDPLFLKLKALNN
ADN32769.1_C4H_Scutellaria_baicalensis          AVVDDVKKNPEA....ATSGIVLRRRLQLMMYNNMYRIMFDRRFESEDDPLFVKLKALNG
AIS92509.1_C4H_Epimedium_sagittatum             KVVEDVRKMPEA....ATNGIVLRKRLQLMMYNNMYRIMFDRRFESEEDPLFLKLKALNG
AGP25594.1_C4H_Aquilaria_sinensis               SVVEDVKKRPES....ATTGIVLRRRLQLMMYNNMYRIMFDRRFESEEDPLFVKLRALNG
AAG50231.1_C4H_Populus_trichocarpa_x_Populus    QVVEDVKKNPEA....ATNGIVLRRRLQLMMYNNMYRIMFDRRFESEDDPLFNKLKALNG
sp|Q43054.1_C4H_Populus_sieboldii_x_Populus_    QVVEDVKKNPEA....ATNGIVLRRRLQLMMYNNMYRIMFDRRFESEDDPLFNKLKALNG
AAG10197.1_C4H_Gossypium_arboreum               SVVEDVKKNPEA....ATNGIVLRKRLQLMMYNNMYRIMFDRRFESEDDPLFVKLKALNG
AAG10196.1_C4H_Gossypium_arboreum               RVVEDVRKNPEA....ATNGIVLRRRLQLMMYNNMYRIMFDTRFESEDDPLFVRLKALNG
sp|Q43033.1_C4H_Petroselinum_crispum            RVVEDVKANPEA....ATNGIVLRNRLQLLMYNNMYRIMFDRRFESVDDPLFLKLKALNG
CAC35977.1_C4H_Ruta_graveolens                  RVVEDVKKDPQA....ATTGIVLRRRLQLLMYNNMYRIMFDRRFESVDDPLFNKLKALNG
AAF66066.2_C4H_Citrus_sinensis                  RVVEDVKKDPEA....ATNGIVLRRRLQLMMYNNMYRIMFDRRFESQDDPLFNRLKALNG
AAK57011.1_C4H_Citrus_x_paradisi                RVVEDVKKDPEA....ATNGIVLRRRLQLMMYNNMYRIMFDRRFESQDDPLFNRLKALNG
sp|P48522.1_C4H_Catharanthus_roseus             RVVEDVKKNPES....ATNGIVLRRRLQLMMYNNMYRIMFDRRFESEDDPLFVKLKALNG
BAB71717.1_C4H_Lithospermum_erythrorhizon       SVVEDVKKNPES....RTNGIVLRKRLQLMMYNNMFRIMFDRRFESEDDPLFIKLKALNG
BAB71716.1_C4H_Lithospermum_erythrorhizon       SVIEDVKKIPES....ETVGIVLRKRLQLMMYNNMFRIMFDRRFESENDPLFMKLRALNG
AAC35857.1_C4H_Capsicum_chinense                SVVEDVKKNPES....ATNGIFLRKRLQLMMYNNMFRIMFDRRFESEDDPLFVKLRALNG
AAG43824.1_C4H_Capsicum_annuum                  SVVEDVKKNPES....ATNGIVLRKRLQLMMYNNMFRIMFDRRFESEDDPPFVKLRALNA
sp|Q43240.1_C4H_Zinnia_elegans                  AVVDDVKKNPKA....ATEGVVIRKRLQLMMYNNMFRIMFDRRFESEDDPLFVKLKMLNG
sp|Q04468.1_C4H_Helianthus_tuberosus            AVVDDVKKNPAA....ATEGIVIRKRLQLMMYNNMFRIMFDRRFESEDDPLFLKLKALNG
sp|Q42797.1_C4H_Glycine_max                     AVVEDVKKNPDA....AVSGTVIRRRLQLMMYNNMYRIMFDRRFESEEDPIFQRLRALNG
sp|P37115.1_C4H_Vigna_radiata                   AVVDDVKKNPDA....AVSGLVIRRRLQLMMYNNMYRIMFDRRFESEEDPLFQRLKALNG
sp|Q96423.1_C4H_Glycyrrhiza_echinata            SVVDDVRRNPDA....AAGGIVLRRRLQLMMYNNMYRIMFDRRFESEEDPLFVKLKALNG
sp|O81928.2_C4H_Cicer_arietinum                 SVVDDVKKNPEA....CSSGIVLRRRLQLMMYNIMYRIMFDRRFESEEDPLFVKLKALNG
sp|P37114.1_C4H_Medicago_sativa                 SVVNDVKNNAEA....SVGGIVIRKRLQLMMYNIMYRIMFDRRFESEEDPLFVKLKALNG
sp|Q43067.2_C4H_Pisum_sativum                   SVVDDVKKNSKA....SVGGIVIRRRLQLMMYNIMYRIMFDRRFESEEDPLFVKLKALNG
AAD11427.1_C4H_Mesembryanthemum_crystallinum    KVVNDLNHNEKISIKAKHEGFVIRKRLQLMLYNIMYRMMFDEGFESMEDPMFIDATKFNS
sp|O24315_C4H_Phaseolus_vulgaris                LVVRDLKVNEHV....RSEGIVIRKRLQLMLYNIMYRMMFDAKFESQEDPLFIQATRFNS
AAK62344.1_C4H_Nicotiana_tabacum                LVVNDLKKNEKV....KYEGIVIRKRLQLMLYNIMYRMMFDAKFESQNDPLFIEATKFNS
AAK62345.1_C4H_Nicotiana_tabacum                LVVDDLKKNEKV....KYDGIVIRKRLQLMLYNIMYRMMFDAKFESQDDPLFIEATKFNS
AAG17469.1_C4H_Triticum_aestivum                LVVEDLKADPAA....ATAGVVVRRRLQLMMYNDMFRIMFDRRFESVADPLFNQLKALNA
AAD23378.1_C4H_Pinus_taeda                      RAVDDIKKRPEA....STTGIVIRRRLQLVMYNIMYRMMFDRRFESEEDPLFLRLKALNG
                                           acc
```

```
AAK54447.1_C4H_Sorghum_bicolor                          η2  α9      η3         α10                 α11
                                                     220         230       240       250       260
AAK54447.1_C4H_Sorghum_bicolor                  ERSRLSQSFEYNYGDFIPVLRPFLRGYLNRCHDLKTRRMKVFEDNFVQERKKVMA.....
TRINITY_DN32464_c6_g3_i2                        ERSRLAQSFEYNYGDFIPILRPFLRGYLKICKEVKERLQLFKDYFVEERKKLASTKSMN
TRINITY_DN15593_c0_g1_i1                        ERSRLAQSFEYNYGDFIPILRPFLRGYLNKCKDLQQRLAFFNNHFVQRRKIMGANGEK
NP_180607.1_C4H_Arabidopsis_thaliana            ERSRLAQSFEYNYGDFIPILRPFLRGYLKICQDVKDRRIALFKKYFVDERKQIASSKPTG
ACA25599.1_C4H_Parthenocissus_henryana          ERSRLAQSFEYNYGDFIPILRPFLRGYLKICKEVKERRLQLFKDHFLEERKKLASTKSTD
CAK95273.1_C4H_Cucumis_sativus                  ERSRLAQSFEYNYGDFIPILRPFLRGYLKICKEVKERRLKLFKDYFVEERKKLANTKSTT
QIC52990.1_C4H_Solanum_nigrum                   ERSRLAQSFEYNYGDFIPILRPFLRGYLKICKEVKEKRLKLFKDYFVDERKKLANTKSMD
AHY94893.1_C4H_Prunella_vulgaris                ERSRLAQSFEYNYGDFIPILRPFLRGYLKMCQQVKERRLKLFKDYFVDERKKLVSTKPAD
AWW24970.1_C4H_Lycoris_radiata                  ERSRLAQSFEYNYGDFIPILRPFLRGYLKICKQVKERRLQLFKDYFLDERKKLASTKPTD
ASA39653.1_C4H_Marchantia_emarginata            ERSQLAQSFDYNYGDFIPILRPFLRGYLKRCQALKDKRLGIFRDYFIEERKKVLEARAKK
ASA39650.1_C4H_Conocephalum_japonicum           ERSRLAQSFEYNYGDFIPILRPFLKGYLNQCQVLKDKRLAIFKEYFLDERKKVLAARQAK
ASA39648.1_C4H_Marchantia_paleacea              ERSRLAQSFDYNYGDFIPILRPFLKKYLARCQALKDKRLAIFKEYFIDERKKVLAAREAR
ASA39646.1_C4H_Plagiochasma_appendiculatum      ERSRLAQSFEYNYGDFIPILRPFLRKYLAYCQELKDKRLALFKEYFIEERKKLLNTRGAN
AFX98061.1_C4H_Cunninghamia_lanceolata          ERSRLAQSFDYNYGDFIPILRPFLRGYLNICKEIKEARLNLFKEYFVEERKKLASVKTVE
AFZ78542.1_C4H_Populus_tomentosa                ERSRLAQSFDYNYGDFIPILRPFLRGYLKICKEVKERRLQLFKDYFVDERKKLGSTKSMS
AUG71936.1_C4H_Narcissus_pseudonarcissus        ERSRLAQSFEYNYGDFIPILRPFLRGYLKICKQVKERRFKLFKDYFLEERKKLASTKPTD
AEA72281.1_C4H_Angelica_gigas                   ERSRLAQSFEYNYGDFIPVLRPLLKGYLKLCQEIKDKRLKLFKDYFVDERKKLESIKSVD
ADO24190.1_C4H_Allium_sativum                   ERSRLAQSFEYNYGDFIPILRPFLRGYLKICKEVKERRIQLFKDYFLEERKKLSSTTTTD
ADN32769.1_C4H_Scutellaria_baicalensis          ERSRLAQSFEYNYGDFIPILRPFLRGYLQICKDVKTKRLQLFKDYFVDERKKVASTKSSS
AIS92509.1_C4H_Epimedium_sagittatum             ERSRLAQSFDYNYGDFIPILRPLLRGYLKICKEVKERRLQLFKDYFLDERKRLTSTTTPS
AGP25594.1_C4H_Aquilaria_sinensis               ERSRLAQSFEYNYGDFIPILRPFLRGYLRICKEVKERRLQLFNDYFVQER.KLASTTSVS
AAG50231.1_C4H_Populus_trichocarpa_x_Populus    ERSRLAQSFEYNYGDFIPILRPFLRGYLKICQEVKERRLQLFKDYFVDERKKLASTKNMS
sp|Q43054.1_C4H_Populus_sieboldii_x_Populus_    ERSRLAQSFDYNYGDFIPILRPFLRGYLKICQEVKERRLQLFKDYFVDERKKLASTKNMS
AAG10197.1_C4H_Gossypium_arboreum               ERSRLAQSFEYNYGDFIPILRPFLRGYLKLCKEVKERRIQLFRDQFLEERKKLATTKRID
AAG10196.1_C4H_Gossypium_arboreum               ERSRLAQSFEYNYGDFIPILRPFLRGYLKICKEVKDRRLQLFKDHFVEERKKLGSTKSMN
sp|Q43033.1_C4H_Petroselinum_crispum            ERSRLAQSFEYHFGDFIPILRPFLRGYLKLCQEIKDKRLKLFKDYFVDERKKLESIKSVD
CAC35977.1_C4H_Ruta_graveolens                  ERSRLAQSFEYNYGDFIPILRPFLRGYLKLVKEVKERRLKLFKDYFVEERKKLTSTKSMT
AAF66066.2_C4H_Citrus_sinensis                  ERSRLAQSFEYNYGDFIPILRPFLRGYLKICKEVKERRLQLFKDYFVEERKKLASTKSMS
AAK57011.1_C4H_Citrus_x_paradisi                ERSRLAQSFEYNYGDFIPILRPFLRGYLKICKEVKERRLQLFKDYFVEERKKLASTKSMS
sp|P48522.1_C4H_Catharanthus_roseus             ERSRLAQGFEYNYGDFIPILRPFLRGYLRICKEVKERRLQLFKDYFVDERKKFGSTKSMD
BAB71717.1_C4H_Lithospermum_erythrorhizon       ERSRLAQSFDYNYGDFIPILRPFLRGYLKMCKEVKQTRLKLFKDYFVDERKKLASSKRMD
BAB71716.1_C4H_Lithospermum_erythrorhizon       ERSRLAQSFDYNYGDFIPILRPFLRGYLKICKEVKETRLKLFKDYFVEERKKIASTKSTT
AAC35857.1_C4H_Capsicum_chinense                ERSRLAQSFEYNYGDFIPIFRPFLRGYLKICKEVKEKRLQLFKDYFVDERKKLANTKSMD
AAG43824.1_C4H_Capsicum_annuum                  ERSRLAQSFEYNYGDFIPILRPFLRGYLKICKEVKEKRLQLFKDYFVDERKKLSNTKSMD
sp|Q43240.1_C4H_Zinnia_elegans                  ERSRLAQSFEYNYGDFIPILRPFLKGYLKLCKEVKEKRFQLFKDYFVDERKKLGSTKSMD
sp|Q04468.1_C4H_Helianthus_tuberosus            ERSRLAQSFEYNYGDFIPILRPFLRNYLKLCKEVKDKRIQLFKDYFVDERKKIGSTKKMD
sp|Q42797.1_C4H_Glycine_max                     ERSRLAQSFEYNYGDFIPILRPFLKGYLKICKEVKETRLKLFKDYFVDERKKLGSTKSTN
sp|P37115.1_C4H_Vigna_radiata                   ERSRLAQSFEYNYGDFIPILRPFLKGYLKICKEVKETRLKLFKDYFVDERKNIGSTKSTN
sp|Q96423.1_C4H_Glycyrrhiza_echinata            ERSRLAQSFEYNYGDFIPILRPFLKGYLKICKEVKERRLKLFKDYFVDERMKLESTKSTS
sp|O81928.2_C4H_Cicer_arietinum                 ERSRLAQSFEYNYGDFIPILRPFLKGYLKLCKEVKERRLQLFKDYFVDERKKLGSTKSTT
sp|P37114.1_C4H_Medicago_sativa                 ERSRLAQSFEYNYGDFIPILRPFLKGYLKVCKEVKDRRLQLFKDYFVDERKKLESTKSTT
sp|Q43067.2_C4H_Pisum_sativum                   ERSRLAQSFEYNYGDFIPILRPFLKGYLKVCKEVKDRRLQLFKDYFVDERKKLGSTKSTY
AAD11427.1_C4H_Mesembryanthemum_crystallinum    ERSRLAQSFEYNYGDFIPFLRPFLRSYLSKCRDLQKSRLAFFNNYFVEKRRKIMAANGEH
sp|O24315_C4H_Phaseolus_vulgaris                ERSRLAQSFEYNYGDFIPILRPFLRGYLNKCKDLQSRRLAFFNTHYVQKRRQIMAANGEK
AAK62344.1_C4H_Nicotiana_tabacum                ERSRLAQSFDYNYGDFIPLLRPFLRGYLNKCKDLQTRRLAFFNNYFVEKRRKIMDENGEK
AAK62345.1_C4H_Nicotiana_tabacum                ERSRLAQSFDYNYGDFIPLLRPFLKGYLNKCKDLQTRRLAFFNNYFVGKRRKIMGENGEK
AAG17469.1_C4H_Triticum_aestivum                ERSILSQSFDYNYGDFIPVLRPFLRRYLNRCTNLKTKRMKVFEDHFVQQRKEALE....
AAD23378.1_C4H_Pinus_taeda                      ERSRLAQSFEYNYGDFIPVLRPFLRGYLKICKEVKESRLLLFKDYFIDERKKLASTTGSK
                                           acc
```

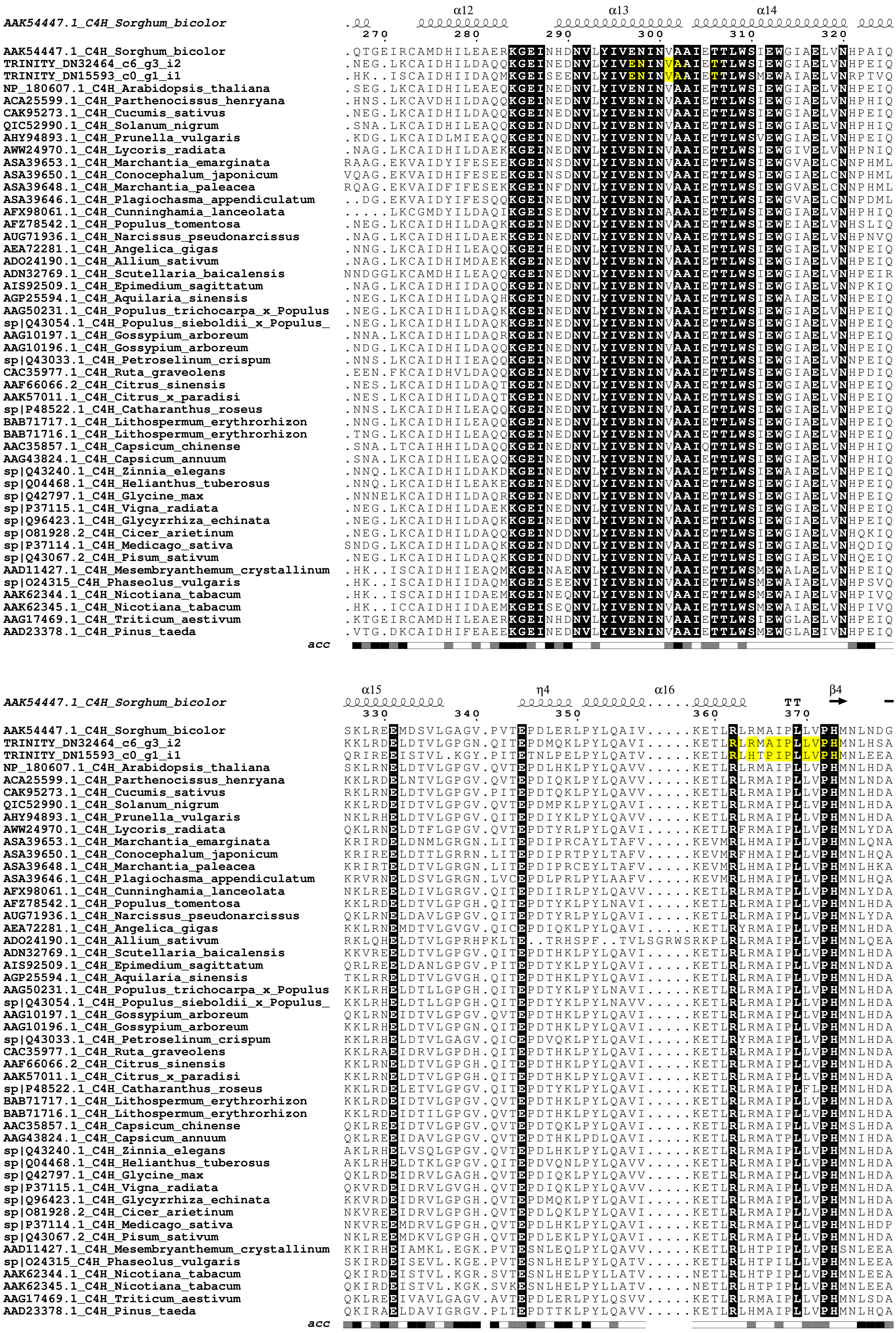
AAK54447.1_C4H_Sorghum_bicolor
α12
α13
α14
270
280
290
300
310
320
AAK54447.1_C4H_Sorghum_bicolor
TRINITY_DN32464_c6_g3_i2
TRINITY_DN15593_c0_g1_i1
NP_180607.1_C4H_Arabidopsis_thaliana
ACA25599.1_C4H_Parthenocissus_henryana
CAK95273.1_C4H_Cucumis_sativus
QIC52990.1_C4H_Solanum_nigrum
AHY94893.1_C4H_Prunella_vulgaris
AWW24970.1_C4H_Lycoris_radiata
ASA39653.1_C4H_Marchantia_emarginata
ASA39650.1_C4H_Conocephalum_japonicum
ASA39648.1_C4H_Marchantia_paleacea
ASA39646.1_C4H_Plagiochasma_appendiculatum
AFX98061.1_C4H_Cunninghamia_lanceolata
AFZ78542.1_C4H_Populus_tomentosa
AUG71936.1_C4H_Narcissus_pseudonarcissus
AEA72281.1_C4H_Angelica_gigas
ADO24190.1_C4H_Allium_sativum
ADN32769.1_C4H_Scutellaria_baicalensis
AIS92509.1_C4H_Epimedium_sagittatum
AGP25594.1_C4H_Aquilaria_sinensis
AAG50231.1_C4H_Populus_trichocarpa_x_Populus
sp|Q43054.1_C4H_Populus_sieboldii_x_Populus_
AAG10197.1_C4H_Gossypium_arboreum
AAG10196.1_C4H_Gossypium_arboreum
sp|Q43033.1_C4H_Petroselinum_crispum
CAC35977.1_C4H_Ruta_graveolens
AAF66066.2_C4H_Citrus_sinensis
AAK57011.1_C4H_Citrus_x_paradisi
sp|P48522.1_C4H_Catharanthus_roseus
BAB71717.1_C4H_Lithospermum_erythrorhizon
BAB71716.1_C4H_Lithospermum_erythrorhizon
AAC35857.1_C4H_Capsicum_chinense
AAG43824.1_C4H_Capsicum_annuum
sp|Q43240.1_C4H_Zinnia_elegans
sp|Q04468.1_C4H_Helianthus_tuberosus
sp|Q42797.1_C4H_Glycine_max
sp|P37115.1_C4H_Vigna_radiata
sp|Q96423.1_C4H_Glycyrrhiza_echinata
sp|O81928.2_C4H_Cicer_arietinum
sp|P37114.1_C4H_Medicago_sativa
sp|Q43067.2_C4H_Pisum_sativum
AAD11427.1_C4H_Mesembryanthemum_crystallinum
sp|O24315_C4H_Phaseolus_vulgaris
AAK62344.1_C4H_Nicotiana_tabacum
AAK62345.1_C4H_Nicotiana_tabacum
AAG17469.1_C4H_Triticum_aestivum
AAD23378.1_C4H_Pinus_taeda
acc
α15
η4
α16
β4
TT
330
340
350
360
370
acc

```
                                               β5   β6
AAK54447.1_C4H_Sorghum_bicolor                 →TT→  TT ——..............................................
                                             380         390
                                               .           .
AAK54447.1_C4H_Sorghum_bicolor                 KLAGYDIPAESKIL..............................................
TRINITY_DN32464_c6_g3_i2                       KLAGYDIPAESKIL..............................................
TRINITY_DN15593_c0_g1_i1                       KLGGFTIPKESKVV..............................................
NP_180607.1_C4H_Arabidopsis_thaliana           KLAGYDIPAESKIL..............................................
ACA25599.1_C4H_Parthenocissus_henryana         KLGSYDIPAESKIL..............................................
CAK95273.1_C4H_Cucumis_sativus                 KLAGYDIPAESKIL..............................................
QIC52990.1_C4H_Solanum_nigrum                  KLAGYDIPAESKIL..............................................
AHY94893.1_C4H_Prunella_vulgaris               KLGGFDIPAESKVL..............................................
AWW24970.1_C4H_Lycoris_radiata                 KLGGYDIPAESKIL..............................................
ASA39653.1_C4H_Marchantia_emarginata           KLGGYDIPAESKIL..............................................
ASA39650.1_C4H_Conocephalum_japonicum          KLGGYDIPAESKIL..............................................
ASA39648.1_C4H_Marchantia_paleacea             KLAGYDIPAESKIL..............................................
ASA39646.1_C4H_Plagiochasma_appendiculatum     KLAGYDIPAESKIL..............................................
AFX98061.1_C4H_Cunninghamia_lanceolata         KLGGYDIPAESKIL..............................................
AFZ78542.1_C4H_Populus_tomentosa               KLGGFDIPAESKIL..............................................
AUG71936.1_C4H_Narcissus_pseudonarcissus       KLGSYDIPAESKIL..............................................
AEA72281.1_C4H_Angelica_gigas                  KLAGYDIPAESKILVNAWWLANNPAHWKNPNEFRPERFLEEESKVEANGNDFKYIPFGVG
ADO24190.1_C4H_Allium_sativum                  KLSGYDIPAESKIL..............................................
ADN32769.1_C4H_Scutellaria_baicalensis         KLNGYDIPAESKIL..............................................
AIS92509.1_C4H_Epimedium_sagittatum            KLNGFDIPAESKIL..............................................
AGP25594.1_C4H_Aquilaria_sinensis              KLGGYDIPAESKIL..............................................
AAG50231.1_C4H_Populus_trichocarpa_x_Populus   KLGGFDIPAESKIL..............................................
sp|Q43054.1_C4H_Populus_sieboldii_x_Populus_   KLGGFDIPAESKIL..............................................
AAG10197.1_C4H_Gossypium_arboreum              KLGGYDIPAESKIL..............................................
AAG10196.1_C4H_Gossypium_arboreum              KLGGYDIPAESKIL..............................................
sp|Q43033.1_C4H_Petroselinum_crispum           KLAGYDIPAESKIL..............................................
CAC35977.1_C4H_Ruta_graveolens                 KLAGYDIPAESKIL..............................................
AAF66066.2_C4H_Citrus_sinensis                 KLGGYDVPAESKIL..............................................
AAK57011.1_C4H_Citrus_x_paradisi               KLGGYDVPAESKIL..............................................
sp|P48522.1_C4H_Catharanthus_roseus            KLGGYDIPAESKIL..............................................
BAB71717.1_C4H_Lithospermum_erythrorhizon      KLGEYDIPAESKSL..............................................
BAB71716.1_C4H_Lithospermum_erythrorhizon      KLNGYDIPAESKIL..............................................
AAC35857.1_C4H_Capsicum_chinense               KLAVYDIPAESKIL..............................................
AAG43824.1_C4H_Capsicum_annuum                 KLAGYDIPAESKIL..............................................
sp|Q43240.1_C4H_Zinnia_elegans                 KLGGYDIPAESKIL..............................................
sp|Q04468.1_C4H_Helianthus_tuberosus           KLGGFDIPAESKIL..............................................
sp|Q42797.1_C4H_Glycine_max                    KLGGYDIPAESKIL..............................................
sp|P37115.1_C4H_Vigna_radiata                  KLGGYDIPAESKIL..............................................
sp|Q96423.1_C4H_Glycyrrhiza_echinata           KLGGYDIPAESKIL..............................................
sp|O81928.2_C4H_Cicer_arietinum                KLSGFDIPAESKIL..............................................
sp|P37114.1_C4H_Medicago_sativa                KLNGFDIPAESKIL..............................................
sp|Q43067.2_C4H_Pisum_sativum                  KLGGFDIPAESKIL..............................................
AAD11427.1_C4H_Mesembryanthemum_crystallinum   KLGGYTIPKNSKVV..............................................
sp|O24315_C4H_Phaseolus_vulgaris               KLGGYTVPKESKVV..............................................
AAK62344.1_C4H_Nicotiana_tabacum               KLGGYTIPKETKVV..............................................
AAK62345.1_C4H_Nicotiana_tabacum               KLGGYTIPKETKVV..............................................
AAG17469.1_C4H_Triticum_aestivum               KLAGYDIPAESKIL..............................................
AAD23378.1_C4H_Pinus_taeda                     KLGGYDIPAESKIL..............................................
                                          acc

                                                                      β7
AAK54447.1_C4H_Sorghum_bicolor                 ............................................................

AAK54447.1_C4H_Sorghum_bicolor                 ............................................................
TRINITY_DN32464_c6_g3_i2                       ............................................................
TRINITY_DN15593_c0_g1_i1                       ............................................................
NP_180607.1_C4H_Arabidopsis_thaliana           ............................................................
ACA25599.1_C4H_Parthenocissus_henryana         ............................................................
CAK95273.1_C4H_Cucumis_sativus                 ............................................................
QIC52990.1_C4H_Solanum_nigrum                  ............................................................
AHY94893.1_C4H_Prunella_vulgaris               ............................................................
AWW24970.1_C4H_Lycoris_radiata                 ............................................................
ASA39653.1_C4H_Marchantia_emarginata           ............................................................
ASA39650.1_C4H_Conocephalum_japonicum          ............................................................
ASA39648.1_C4H_Marchantia_paleacea             ............................................................
ASA39646.1_C4H_Plagiochasma_appendiculatum     ............................................................
AFX98061.1_C4H_Cunninghamia_lanceolata         ............................................................
AFZ78542.1_C4H_Populus_tomentosa               ............................................................
AUG71936.1_C4H_Narcissus_pseudonarcissus       ............................................................
AEA72281.1_C4H_Angelica_gigas                  RRSCPGIILALPILGITIGRMVQNFELLPPPGMSKIDTAEKAGQFSLQISKHYTIVCKPR
ADO24190.1_C4H_Allium_sativum                  ............................................................
ADN32769.1_C4H_Scutellaria_baicalensis         ............................................................
AIS92509.1_C4H_Epimedium_sagittatum            ............................................................
AGP25594.1_C4H_Aquilaria_sinensis              ............................................................
AAG50231.1_C4H_Populus_trichocarpa_x_Populus   ............................................................
sp|Q43054.1_C4H_Populus_sieboldii_x_Populus_   ............................................................
AAG10197.1_C4H_Gossypium_arboreum              ............................................................
AAG10196.1_C4H_Gossypium_arboreum              ............................................................
sp|Q43033.1_C4H_Petroselinum_crispum           ............................................................
CAC35977.1_C4H_Ruta_graveolens                 ............................................................
AAF66066.2_C4H_Citrus_sinensis                 ............................................................
AAK57011.1_C4H_Citrus_x_paradisi               ............................................................
sp|P48522.1_C4H_Catharanthus_roseus            ............................................................
BAB71717.1_C4H_Lithospermum_erythrorhizon      ............................................................
BAB71716.1_C4H_Lithospermum_erythrorhizon      ............................................................
AAC35857.1_C4H_Capsicum_chinense               ............................................................
AAG43824.1_C4H_Capsicum_annuum                 ............................................................
sp|Q43240.1_C4H_Zinnia_elegans                 ............................................................
sp|Q04468.1_C4H_Helianthus_tuberosus           ............................................................
sp|Q42797.1_C4H_Glycine_max                    ............................................................
sp|P37115.1_C4H_Vigna_radiata                  ............................................................
sp|Q96423.1_C4H_Glycyrrhiza_echinata           ............................................................
sp|O81928.2_C4H_Cicer_arietinum                ............................................................
sp|P37114.1_C4H_Medicago_sativa                ............................................................
sp|Q43067.2_C4H_Pisum_sativum                  ............................................................
AAD11427.1_C4H_Mesembryanthemum_crystallinum   ............................................................
sp|O24315_C4H_Phaseolus_vulgaris               ............................................................
AAK62344.1_C4H_Nicotiana_tabacum               ............................................................
AAK62345.1_C4H_Nicotiana_tabacum               ............................................................
AAG17469.1_C4H_Triticum_aestivum               ............................................................
AAD23378.1_C4H_Pinus_taeda                     ............................................................
                                          acc
```

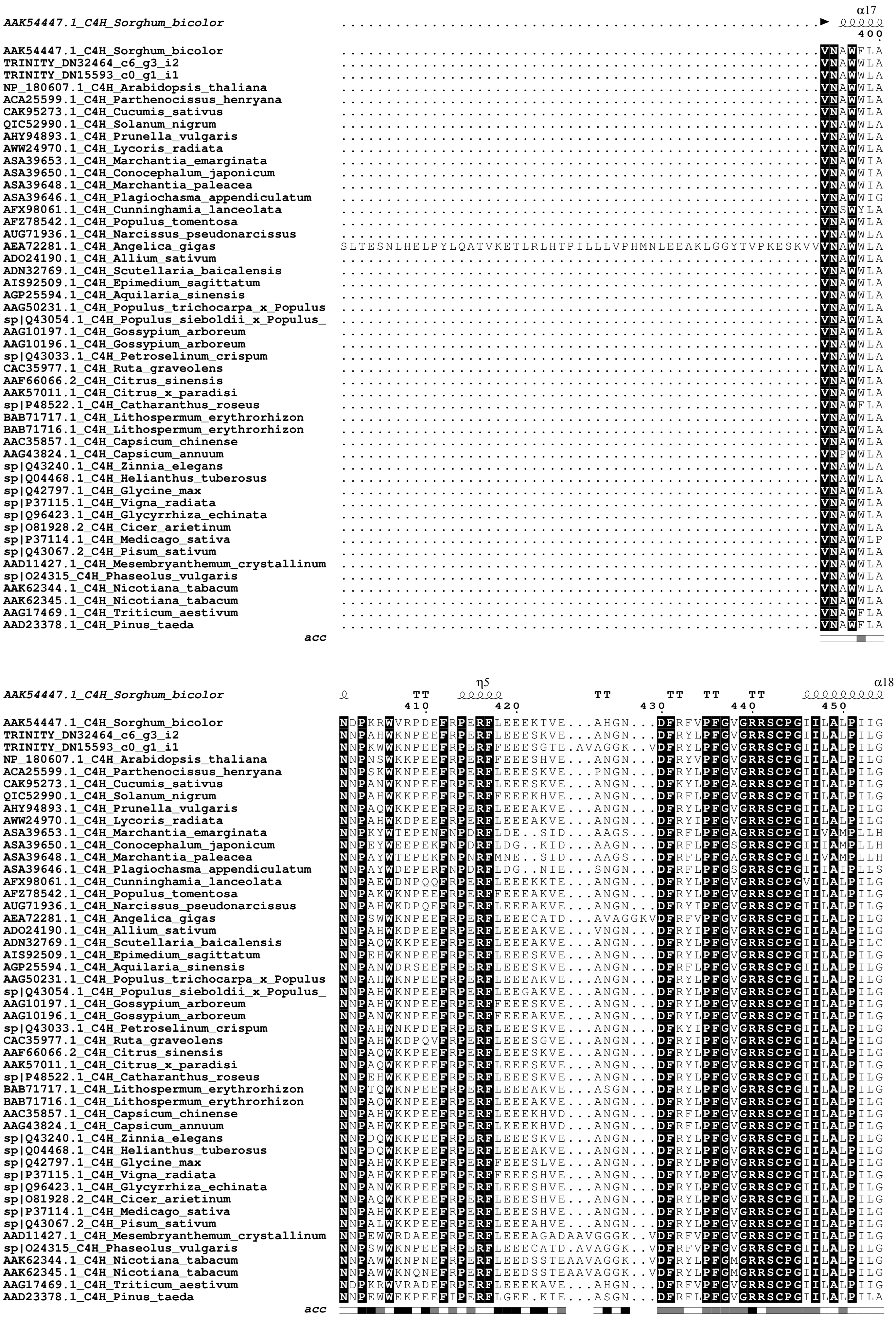
AAK54447.1_C4H_Sorghum_bicolor
α17
400
AAK54447.1_C4H_Sorghum_bicolor VNAWFLA
TRINITY_DN32464_c6_g3_i2 VNAWWLA
TRINITY_DN15593_c0_g1_i1 VNAWWLA
NP_180607.1_C4H_Arabidopsis_thaliana VNAWWLA
ACA25599.1_C4H_Parthenocissus_henryana VNAWWLA
CAK95273.1_C4H_Cucumis_sativus VNAWWLA
QIC52990.1_C4H_Solanum_nigrum VNAWWLA
AHY94893.1_C4H_Prunella_vulgaris VNAWWLA
AWW24970.1_C4H_Lycoris_radiata VNAWWLA
ASA39653.1_C4H_Marchantia_emarginata VNAWWIA
ASA39650.1_C4H_Conocephalum_japonicum VNAWWIA
ASA39648.1_C4H_Marchantia_paleacea VNAWWIA
ASA39646.1_C4H_Plagiochasma_appendiculatum VNAWWIG
AFX98061.1_C4H_Cunninghamia_lanceolata VNSWYLA
AFZ78542.1_C4H_Populus_tomentosa VNAWWLA
AUG71936.1_C4H_Narcissus_pseudonarcissus VNAWWLA
AEA72281.1_C4H_Angelica_gigas SLTESNLHELPYLQATVKETLRLHTPILLLVPHMNLEEAKLGGYTVPKESKVVVNAWWLA
ADO24190.1_C4H_Allium_sativum VNAWWLA
ADN32769.1_C4H_Scutellaria_baicalensis VNAWWLA
AIS92509.1_C4H_Epimedium_sagittatum VNAWWLA
AGP25594.1_C4H_Aquilaria_sinensis VNAWWLA
AAG50231.1_C4H_Populus_trichocarpa_x_Populus VNAWWLA
sp|Q43054.1_C4H_Populus_sieboldii_x_Populus_ VNAWWLA
AAG10197.1_C4H_Gossypium_arboreum VNAWWLA
AAG10196.1_C4H_Gossypium_arboreum VNAWWLA
sp|Q43033.1_C4H_Petroselinum_crispum VNAWWLA
CAC35977.1_C4H_Ruta_graveolens VNAWWLA
AAF66066.2_C4H_Citrus_sinensis VNAWWLA
AAK57011.1_C4H_Citrus_x_paradisi VNAWWLA
sp|P48522.1_C4H_Catharanthus_roseus VNAWFLA
BAB71717.1_C4H_Lithospermum_erythrorhizon VNAWWLA
BAB71716.1_C4H_Lithospermum_erythrorhizon VNAWWLA
AAC35857.1_C4H_Capsicum_chinense VNAWWLA
AAG43824.1_C4H_Capsicum_annuum VNPWWLA
sp|Q43240.1_C4H_Zinnia_elegans VNAWWLA
sp|Q04468.1_C4H_Helianthus_tuberosus VNAWWLA
sp|Q42797.1_C4H_Glycine_max VNAWWLA
sp|P37115.1_C4H_Vigna_radiata VNAWWLA
sp|Q96423.1_C4H_Glycyrrhiza_echinata VNAWWLA
sp|O81928.2_C4H_Cicer_arietinum VNAWWLA
sp|P37114.1_C4H_Medicago_sativa VNAWWLP
sp|Q43067.2_C4H_Pisum_sativum VNAWWLA
AAD11427.1_C4H_Mesembryanthemum_crystallinum VNAWWLA
sp|O24315_C4H_Phaseolus_vulgaris VNAWWLA
AAK62344.1_C4H_Nicotiana_tabacum VNAWWLA
AAK62345.1_C4H_Nicotiana_tabacum VNAWWLA
AAG17469.1_C4H_Triticum_aestivum VNAWFLA
AAD23378.1_C4H_Pinus_taeda VNAWFLA
acc
AAK54447.1_C4H_Sorghum_bicolor
TT η5 TT TT TT TT α18
410 420 430 440 450
AAK54447.1_C4H_Sorghum_bicolor NDPKRWVRPDEFRPERFLEEEKTVE...AHGN...DFRFVPFGVGRRSCPGIILALPIIG
TRINITY_DN32464_c6_g3_i2 NNPAHWKNPEEFRPERFLEEESKVE...ANGN...DFRYLPFGVGRRSCPGIILALPILG
TRINITY_DN15593_c0_g1_i1 NNPKWWKNPEEFRPERFFEEESGTE.AVAGGK..VDFRYLPFGVGRRSCPGIILALPILG
NP_180607.1_C4H_Arabidopsis_thaliana NNPNSWKKPEEFRPERFFEEESHVE...ANGN...DFRYVPFGVGRRSCPGIILALPILG
ACA25599.1_C4H_Parthenocissus_henryana NNPSKWKNPEEFRPERFLEEESKVE...PNGN...DFRYLPFGVGRRSCPGIILALPILG
CAK95273.1_C4H_Cucumis_sativus NNPANWKNPEEFRPERFLEEESKVE...ANGN...DFKYLPFGAGRRSCPGIILALPILG
QIC52990.1_C4H_Solanum_nigrum NNPAHWKKPEEFRPERFFEEEKHVE...ANGN...DFRFLPFGVGRRSCPGIILALPILG
AHY94893.1_C4H_Prunella_vulgaris NNPAQWKKPEEFRPERFLEEEAKVE...ANGN...DFRYLPFGVGRRSCPGIILALPILG
AWW24970.1_C4H_Lycoris_radiata NNPAHWKNPEEFRPERFLEEEAKVE...ANGN...DFRYIPFGVGRRSCPGIILALPILG
ASA39653.1_C4H_Marchantia_emarginata NNPKYWTEPENFNPDRFLDE..SID...AAGS...DFRFLPFGAGRRSCPGIIVAMPLLH
ASA39650.1_C4H_Conocephalum_japonicum NNPEYWEEPEKFNPDRFLDG..KID...AAGN...DFRFLPFGSGRRSCPGIIIAMPLLH
ASA39648.1_C4H_Marchantia_paleacea NNPAYWTEPEKFNPNRFMNE..SID...AAGS...DFRFLPFGAGRRSCPGIIVAMPLLH
ASA39646.1_C4H_Plagiochasma_appendiculatum NNPAYWDEPERFNPDRFLDG..NIE...SNGN...DFRFLPFGSGRRSCPGIIIAIPLLS
AFX98061.1_C4H_Cunninghamia_lanceolata NNPAEWDNPQQFRPERFLEEEKKTE...ANGN...DFRYLPFGVGRRSCPGVILALPILG
AFZ78542.1_C4H_Populus_tomentosa NNPAHWKNPEEFRPERFFEEEAKVE...ANGN...DFRYLPFGVGRRSCPGIILALPILG
AUG71936.1_C4H_Narcissus_pseudonarcissus NNPAHWKDPQEFRPERFLEEEAKVE...ANGN...DFRYIPFGVGRRSCPGIILALPILG
AEA72281.1_C4H_Angelica_gigas NNPSWWKNPEEFRPERFLEEECATD...AVAGGKVDFRFVPFGVGRRSCPGIILALPILG
ADO24190.1_C4H_Allium_sativum NNPAHWKDPEEFRPERFLEEEAKVE...VNGN...DFRYIPFGVGRRSCPGIILALPILG
ADN32769.1_C4H_Scutellaria_baicalensis NNPAQWKKPEEFRPERFLEEEAKVE...ANGN...DFRYLPFGVGRRSCPGIILALPILC
AIS92509.1_C4H_Epimedium_sagittatum NNPEHWKNPEEFRPERFLEEESKVE...ANGN...DFRYLPFGVGRRSCPGIILALPILG
AGP25594.1_C4H_Aquilaria_sinensis NNPANWDRSEEFRPERFLEEESKVE...ANGN...DFRYLPFGVGRRSCPGIILALPILG
AAG50231.1_C4H_Populus_trichocarpa_x_Populus NNPAHWKNPEEFRPERFLEEEAKVE...ANGN...DFRYLPFGVGRRSCPGIILALPILG
sp|Q43054.1_C4H_Populus_sieboldii_x_Populus_ NNPAHWKNPEEFRPERFLEEGAKVE...ANGN...DFRYLPFGVGRRSCPGIILALPILG
AAG10197.1_C4H_Gossypium_arboreum NNPAHWKNPEEFRPERFFEEESKVE...ANGN...DFRYLPFGVGRRSCPGIILALPILG
AAG10196.1_C4H_Gossypium_arboreum NNPANWKNPEEFRPERFFEEEAKVE...ANGN...DFRYLPFGVGRRSCPGIILALPILG
sp|Q43033.1_C4H_Petroselinum_crispum NNPAHWNKPDEFRPERFLEEESKVE...ANGN...DFKYIPFGVGRRSCPGIILALPILG
CAC35977.1_C4H_Ruta_graveolens NNPAHWKDPQVFRPERFLEEESGVE...ANGN...DFRYIPFGVGRRSCPGIILALPILG
AAF66066.2_C4H_Citrus_sinensis NNPAQWKKPEEFRPERFLEEESKVE...ANGN...DFRYLPFGVGRRSCPGIILALPILG
AAK57011.1_C4H_Citrus_x_paradisi NNPAQWKKPEEFRPERFLEEESKVE...ANGN...DFRYLPFGVGRRSCPGIILALPILG
sp|P48522.1_C4H_Catharanthus_roseus NNPEHWKKPEEFRPERFLEEESKVE...ANGN...DFRYLPFGVGRRSCPGIILALPILG
BAB71717.1_C4H_Lithospermum_erythrorhizon NNPTQWKNPEEFRPERFLEEEAKVE...ASGN...DFRYLPFGVGRRSCPGIILALPILG
BAB71716.1_C4H_Lithospermum_erythrorhizon NNPAQWKNPEEFRPERFLEEEAKVE...ANGN...DFRYLPFGVGRRSCPGIILALPILG
AAC35857.1_C4H_Capsicum_chinense NNPAHWKKPEEFRPERFLEEEKHVD...ANGN...DFRFLPFGVGRRSCPGIILALPILG
AAG43824.1_C4H_Capsicum_annuum NNPAHWKKPEEFRPERFLKEEKHVD...ANGN...DFRFLPFGVGRRSCPGIILALPILG
sp|Q43240.1_C4H_Zinnia_elegans NNPDQWKKPEEFRPERFLEEESKVE...ANGN...DFRYLPFGVGRRSCPGIILALPILG
sp|Q04468.1_C4H_Helianthus_tuberosus NNPDQWKKPEEFRPERFLEEEAKVE...ANGN...DFRYLPFGVGRRSCPGIILALPILG
sp|Q42797.1_C4H_Glycine_max NNPAHWKKPEEFRPERFFEEESLVE...ANGN...DFRYLPFGVGRRSCPGIILALPILG
sp|P37115.1_C4H_Vigna_radiata NNPAHWKKPEEFRPERFFEEESHVE...ANGN...DFRYLPFGVGRRSCPGIILALPILG
sp|Q96423.1_C4H_Glycyrrhiza_echinata NNPANWKKPEEFRPERFLEEESHVE...ANGN...DFRYLPFGVGRRSCPGIILALPILG
sp|O81928.2_C4H_Cicer_arietinum NNPAQWKKPEEFRPERFLEEESHVE...ANGN...DFRYLPFGVGRRSCPGIILALPILG
sp|P37114.1_C4H_Medicago_sativa NNPAHWKKPEEFRPERFLEEESHVE...ANGN...DFRYLPFGVGRRSCPGIILALPILG
sp|Q43067.2_C4H_Pisum_sativum NNPALWKKPEEFRPERFLEEEAHVE...ANGN...DFRYLPFGVGRRSCPGIILALPILG
AAD11427.1_C4H_Mesembryanthemum_crystallinum NNPEWWRDAEEFRPERFLEEEAGADAVAGGGK..VDFRFVPFGVGRRSCPGIILALPILG
sp|O24315_C4H_Phaseolus_vulgaris NNPSWWKNPEEFRPERFLEEECATD.AVAGGK..VDFRFVPFGVGRRSCPGIILALPILG
AAK62344.1_C4H_Nicotiana_tabacum NNPAWWKNPNEFRPERFLEEDSSTEAAVAGGK..VDFRYLPFGMGRRSCPGIILALPILG
AAK62345.1_C4H_Nicotiana_tabacum NNPAWWKNQNEFRPERFLEEDSSTEAAVAGGK..VDFRYLPFGMGRRSCPGIILALPILG
AAG17469.1_C4H_Triticum_aestivum NNPKRWVRADEFRPERFLEEEKAVE...AHGN...DFRFVPFGVGRRSCPGIILALPIIG
AAD23378.1_C4H_Pinus_taeda NNPEWWEKPEEFIPERFLGEE.KIE...ASGN...DFRFLPFGVGRRSCPGIILALPILA
acc

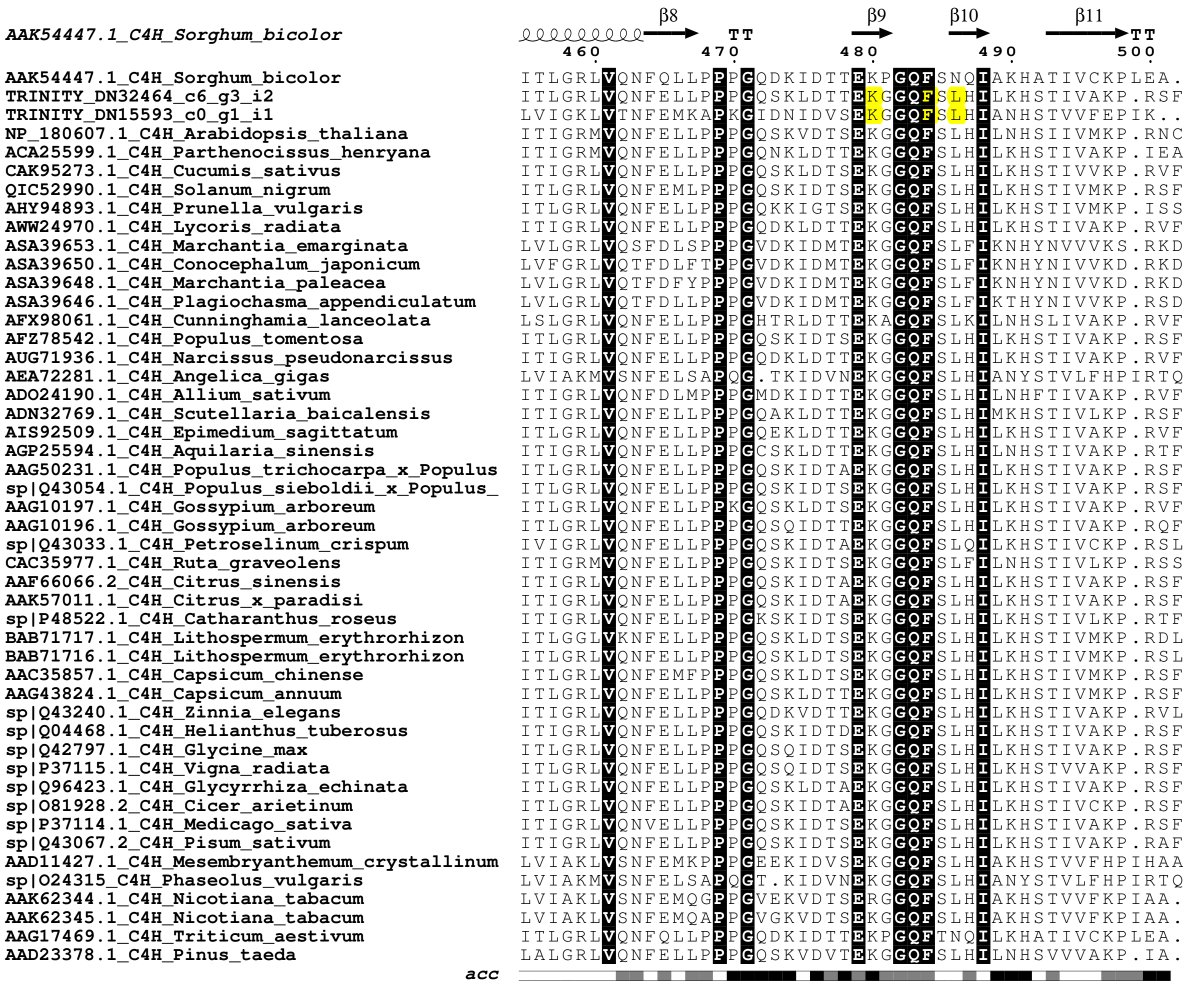
AAK54447.1_C4H_Sorghum_bicolor
β8
β9
β10
β11
TT
TT
460
470
480
490
500
AAK54447.1_C4H_Sorghum_bicolor ITLGRLVQNFQLLPPPGQDKIDTTEKPGQFSNQIAKHATIVCKPLEA..
TRINITY_DN32464_c6_g3_i2 ITLGRLVQNFELLPPPGQSKLDTTEKGGQFSLHILKHSTIVAKP.RSF.
TRINITY_DN15593_c0_g1_i1 LVIGKLVTNFEMKAPKGIDNIDVSEKGGQFSLHIANHSTVVFEPIK...
NP_180607.1_C4H_Arabidopsis_thaliana ITIGRMVQNFELLPPPGQSKVDTSEKGGQFSLHILNHSIIVMKP.RNC.
ACA25599.1_C4H_Parthenocissus_henryana ITIGRMVQNFELLPPPGQNKLDTTEKGGQFSLHILKHSTIVAKP.IEA.
CAK95273.1_C4H_Cucumis_sativus ITIGRLVQNFELLPPPGQSKLDTSEKGGQFSLHILKHSTIVVKP.RVF.
QIC52990.1_C4H_Solanum_nigrum ITLGRLVQNFEMLPPPGQSKIDTSEKGGQFSLHILKHSTIVMKP.RSF.
AHY94893.1_C4H_Prunella_vulgaris ITLGRLVQNFELLPPPGQKKIGTSEKGGQFSLHILKHSTIVMKP.ISS.
AWW24970.1_C4H_Lycoris_radiata ITIGRLVQNFELLPPPGQDKLDTTEKGGQFSLHILKHSTIVAKP.RVF.
ASA39653.1_C4H_Marchantia_emarginata LVLGRLVQSFDLSPPPGVDKIDMTEKGGQFSLFIKNHYNVVVKS.RKD.
ASA39650.1_C4H_Conocephalum_japonicum LVFGRLVQTFDLFTPPGVDKIDMTEKGGQFSLFIKNHYNIVVKD.RKD.
ASA39648.1_C4H_Marchantia_paleacea LVLGRLVQTFDFYPPPGVDKIDMTEKGGQFSLFIKNHYNIVVKD.RKD.
ASA39646.1_C4H_Plagiochasma_appendiculatum LVLGRLVQTFDLLPPPGVDKIDMTEKGGQFSLFIKTHYNIVVKP.RSD.
AFX98061.1_C4H_Cunninghamia_lanceolata LSLGRLVQNFELLPPPGHTRLDTTEKAGQFSLKILNHSLIVAKP.RVF.
AFZ78542.1_C4H_Populus_tomentosa ITLGRLVQNFELLPPPGQSKIDTSEKGGQFSLHILKHSTIVAKP.RSF.
AUG71936.1_C4H_Narcissus_pseudonarcissus ITIGRLVQNFELLPPPGQDKLDTTEKGGQFSLHILKHSTIVAKP.RVF.
AEA72281.1_C4H_Angelica_gigas LVIAKMVSNFELSAPQG.TKIDVNEKGGQFSLHIANYSTVLFHPIRTQ.
ADO24190.1_C4H_Allium_sativum ITIGRLVQNFDLMPPPGMDKIDTTEKGGQFSLHILNHFTIVAKP.RVF.
ADN32769.1_C4H_Scutellaria_baicalensis ITIGRLVQNFELLPPPGQAKLDTTEKGGQFSLHIMKHSTIVLKP.RSF.
AIS92509.1_C4H_Epimedium_sagittatum ITLGRLVQNFELLPPPGQEKLDTTEKGGQFSLHILKHSTIVAKP.RVF.
AGP25594.1_C4H_Aquilaria_sinensis ITIGRLVQNFELLPPPGCSKLDTTEKGGQFSLHILNHSTIVAKP.RTF.
AAG50231.1_C4H_Populus_trichocarpa_x_Populus ITLGRLVQNFELLPPPGQSKIDTAEKGGQFSLHILKHSTIVAKP.RSF.
sp|Q43054.1_C4H_Populus_sieboldii_x_Populus_ ITLGRLVQNFELLPPPGQSKIDTSEKGGQFSLHILKHSTIVAKP.RSF.
AAG10197.1_C4H_Gossypium_arboreum ITLGRLVQNFELLPPKGQSKLDTSEKGGQFSLHILKHSTIVAKP.RVF.
AAG10196.1_C4H_Gossypium_arboreum ITLGRLVQNFELLPPPGQSQIDTTEKGGQFSLHILKHSTIVAKP.RQF.
sp|Q43033.1_C4H_Petroselinum_crispum IVIGRLVQNFELLPPPGQSKIDTAEKGGQFSLQILKHSTIVCKP.RSL.
CAC35977.1_C4H_Ruta_graveolens ITIGRMVQNFELLPPPGQSKIDTSEKGGQFSLFILNHSTIVLKP.RSSV
AAF66066.2_C4H_Citrus_sinensis ITIGRLVQNFELLPPPGQSKIDTAEKGGQFSLHILKHSTIVAKP.RSF.
AAK57011.1_C4H_Citrus_x_paradisi ITIGRLVQNFELLPPPGQSKIDTAEKGGQFSLHILKHSTIVAKP.RSF.
sp|P48522.1_C4H_Catharanthus_roseus ITIGRLVQNFELLPPPGKSKIDTSEKGGQFSLHILKHSTIVLKP.RTF.
BAB71717.1_C4H_Lithospermum_erythrorhizon ITLGGLVKNFELLPPPGQSKLDTSEKGGQFSLHILKHSTIVMKP.RDL.
BAB71716.1_C4H_Lithospermum_erythrorhizon ITLGRLVQNFELLPPPGQSKLDTSEKGGQFSLHILKHSTIVMKP.RSL.
AAC35857.1_C4H_Capsicum_chinense ITLGRLVQNFEMFPPPGQSKLDTSEKGGQFSLHILKHSTIVMKP.RSF.
AAG43824.1_C4H_Capsicum_annuum ITLGRLVQNFELLPPPGQSKLDTTEKGGQFSLHILKHSTIVMKP.RSF.
sp|Q43240.1_C4H_Zinnia_elegans ITIGRLVQNFELLPPPGQDKVDTTEKGGQFSLHILKHSTIVAKP.RVL.
sp|Q04468.1_C4H_Helianthus_tuberosus ITIGRLVQNFELLPPPGQSKIDTDEKGGQFSLHILKHSTIVAKP.RSF.
sp|Q42797.1_C4H_Glycine_max ITLGRLVQNFELLPPPGQSQIDTSEKGGQFSLHILKHSTIVAKP.RSF.
sp|P37115.1_C4H_Vigna_radiata ITLGRLVQNFELLPPPGQSQIDTSEKGGQFSLHILKHSTVVAKP.RSF.
sp|Q96423.1_C4H_Glycyrrhiza_echinata ITLGRLVQNFELLPPPGQSKLDTAEKGGQFSLHILKHSTIVAKP.RSF.
sp|O81928.2_C4H_Cicer_arietinum ITLGRLVQNFELLPPPGQSKIDTAEKGGQFSLHILKHSTIVCKP.RSFN
sp|P37114.1_C4H_Medicago_sativa ITIGRLVQNVELLPPPGQSKIDTSEKGGQFSLHILKHSTIVAKP.RSF.
sp|Q43067.2_C4H_Pisum_sativum ITIGRLVQNFELLPPPGQSKIDTSEKGGQFSLHILKHSTIVAKP.RAF.
AAD11427.1_C4H_Mesembryanthemum_crystallinum LVIAKLVSNFEMKPPPGEEKIDVSEKGGQFSLHIAKHSTVVFHPIHAA.
sp|O24315_C4H_Phaseolus_vulgaris LVIAKMVSNFELSAPQGT.KIDVNEKGGQFSLHIANYSTVLFHPIRTQ.
AAK62344.1_C4H_Nicotiana_tabacum LVIAKLVSNFEMQGPPGVEKVDTSERGGQFSLHIAKHSTVVFKPIAA..
AAK62345.1_C4H_Nicotiana_tabacum LVIAKLVSNFEMQAPPGVGKVDTSEKGGQFSLHIAKHSTVVFKPIAA..
AAG17469.1_C4H_Triticum_aestivum ITLGRLVQNFQLLPPPGQDKIDTTEKPGQFTNQILKHATIVCKPLEA..
AAD23378.1_C4H_Pinus_taeda LALGRLVQNFELLPPPGQSKVDVTEKGGQFSLHILNHSVVVAKP.IA..
acc

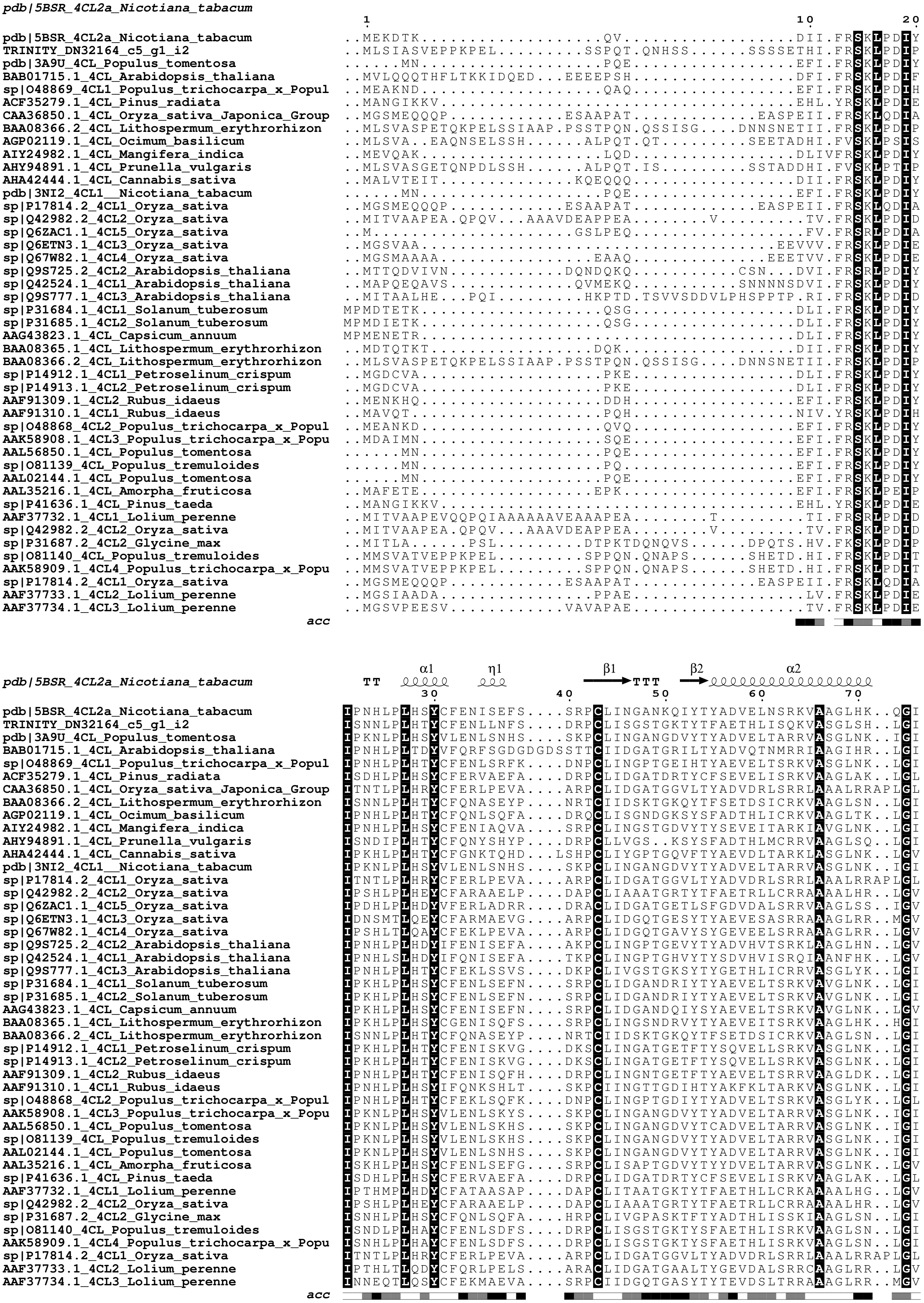
pdb|5BSR_4CL2a_Nicotiana_tabacum
1 10 20
pdb|5BSR_4CL2a_Nicotiana_tabacum ..MEKDTK.................QV...............DII.FRSKLPDIY
TRINITY_DN32164_c5_g1_i2 ..MLSIASVEPPKPEL........SSPQT.QNHSS......SSSSETHI.FRSKLPDIP
pdb|3A9U_4CL_Populus_tomentosa ......MN.................PQE...............EFI.FRSKLPDIY
BAB01715.1_4CL_Arabidopsis_thaliana ..MVLQQQTHFLTKKIDQED...EEEEPSH.............DFI.FRSKLPDIF
sp|O48869_4CL1_Populus_trichocarpa_x_Popul ..MEAKND.................QAQ...............EFI.FRSKLPDIH
ACF35279.1_4CL_Pinus_radiata ..MANGIKKV...............................EHL.YRSKLPDIE
CAA36850.1_4CL_Oryza_sativa_Japonica_Group ..MGSMEQQQP...........ESAAPAT............EASPEII.FRSKLQDIA
BAA08366.2_4CL_Lithospermum_erythrorhizon ..MLSVASPETQKPELSSIAAP.PSSTPQN.QSSISG....DNNSNETII.FRSKLPDIP
AGP02119.1_4CL_Ocimum_basilicum ..MLSVA..EAQNSELSSH......ALPQT.QT........SEETADHI.FVSKLPSIS
AIY24982.1_4CL_Mangifera_indica ..MEVQAK.................LQD...............DLIVFRSKLPDIY
AHY94891.1_4CL_Prunella_vulgaris ..MLSVASGETQNPDLSSH......ALPQT.IS........SSTADDHI.FVSKLPTIP
AHA42444.1_4CL_Cannabis_sativa ..MALVTEIT............KQEQQQ...............DII.FRSKLPDIY
pdb|3NI2_4CL1__Nicotiana_tabacum ......MN.................PQE...............EFI.FRSKLPDIY
sp|P17814.2_4CL1_Oryza_sativa ..MGSMEQQQP...........ESAAPAT............EASPEII.FRSKLQDIA
sp|Q42982.2_4CL2_Oryza_sativa ..MITVAAPEA.QPQV...AAAVDEAPPEA.......V.........TV.FRSKLPDID
sp|Q6ZAC1.1_4CL5_Oryza_sativa ..M.....................GSLPEQ...............FV.FRSRLPDIA
sp|Q6ETN3.1_4CL3_Oryza_sativa ..MGSVAA..............................EEVVV.FRSKLPDIE
sp|Q67W82.1_4CL4_Oryza_sativa ..MGSMAAAA.............EAAQ............EEETVV.FRSKLPDIE
sp|Q9S725.2_4CL2_Arabidopsis_thaliana ..MTTQDVIVN...........DQNDQKQ..........CSN...DVI.FRSRLPDIY
sp|Q42524.1_4CL1_Arabidopsis_thaliana ..MAPQEQAVS...........QVMEKQ..........SNNNNSDVI.FRSKLPDIY
sp|Q9S777.1_4CL3_Arabidopsis_thaliana ..MITAALHE...PQI.........HKPTD.TSVVSDDVLPHSPPTP.RI.FRSKLPDID
sp|P31684.1_4CL1_Solanum_tuberosum MPMDTETK.................QSG...............DLI.FRSKLPDIY
sp|P31685.1_4CL2_Solanum_tuberosum MPMDIETK.................QSG...............DLI.FRSKLPDIY
AAG43823.1_4CL_Capsicum_annuum MPMENETR.................QSG...............DLI.FRSKLPDIY
BAA08365.1_4CL_Lithospermum_erythrorhizon ..MDTQTKT...............DQK...............DII.FRSKLPDIY
BAA08366.2_4CL_Lithospermum_erythrorhizon ..MLSVASPETQKPELSSIAAP.PSSTPQN.QSSISG....DNNSNETII.FRSKLPDIP
sp|P14912.1_4CL1_Petroselinum_crispum ..MGDCVA.................PKE...............DLI.FRSKLPDIY
sp|P14913.1_4CL2_Petroselinum_crispum ..MGDCVA.................PKE...............DLI.FRSKLPDIY
AAF91309.1_4CL2_Rubus_idaeus ..MENKHQ.................DDH...............EFI.FRSKLPDIY
AAF91310.1_4CL1_Rubus_idaeus ..MAVQT..................PQH...............NIV.YRSKLPDIH
sp|O48868_4CL2_Populus_trichocarpa_x_Popul ..MEANKD.................QVQ...............EFI.FRSKLPDIY
AAK58908.1_4CL3_Populus_trichocarpa_x_Popu ..MDAIMN.................SQE...............EFI.FRSKLPDIY
AAL56850.1_4CL_Populus_tomentosa ......MN.................PQE...............EFI.FRSKLPDIY
sp|O81139_4CL_Populus_tremuloides ......MN.................PQ................EFI.FRSKLPDIY
AAL02144.1_4CL_Populus_tomentosa ......MN.................PQE...............EFI.FRSKLPDIY
AAL35216.1_4CL_Amorpha_fruticosa ..MAFETE.................EPK...............EFI.FRSKLPEIP
sp|P41636.1_4CL_Pinus_taeda ..MANGIKKV...............................EHL.YRSKLPDIE
AAF37732.1_4CL1_Lolium_perenne ..MITVAAPEVQQPQIAAAAAAVEAAAPEA.......T.........TI.FRSRLPDID
sp|Q42982.2_4CL2_Oryza_sativa ..MITVAAPEA.QPQV...AAAVDEAPPEA.......V.........TV.FRSKLPDID
sp|P31687.2_4CL2_Glycine_max ..MITLA......PSL.........DTPKTDQNQVS......DPQTS.HV.FKSKLPDIP
sp|O81140_4CL_Populus_tremuloides ..MMSVATVEPPKPEL........SPPQN.QNAPS......SHETD.HI.FRSKLPDIT
AAK58909.1_4CL4_Populus_trichocarpa_x_Popu ..MMSVATVEPPKPEL........SPPQN.QNAPS......SHETD.HI.FRSKLPDIT
sp|P17814.2_4CL1_Oryza_sativa ..MGSMEQQQP...........ESAAPAT............EASPEII.FRSKLQDIA
AAF37733.1_4CL2_Lolium_perenne ..MGSIAADA...............PPAE................LV.FRSKLPDIE
AAF37734.1_4CL3_Lolium_perenne ..MGSVPEESV.............VAVAPAE................TV.FRSKLPDIE
acc
pdb|5BSR_4CL2a_Nicotiana_tabacum
TT α1 η1 β1 TTT β2 α2
30 40 50 60 70
pdb|5BSR_4CL2a_Nicotiana_tabacum IPNHLPLHSYCFENISEFS....SRPCLINGANKQIYTYADVELNSRKVAAGLHK..QGI
TRINITY_DN32164_c5_g1_i2 ISNNLPLHTYCFENLLNFS....DRPCLISGSTGKTYTFAETHLISQKVASGLSH..LGI
pdb|3A9U_4CL_Populus_tomentosa IPKNLPLHSYVLENLSNHS....SKPCLINGANGDVYTYADVELTARRVASGLNK..IGI
BAB01715.1_4CL_Arabidopsis_thaliana IPNHLPLTDYVFQRFSGDGDGDSSTTCIIDGATGRILTYADVQTNMRRIAAGIHR..LGI
sp|O48869_4CL1_Populus_trichocarpa_x_Popul IPNHLPLHTYCFENLSRFK....DNPCLINGPTGEIHTYAEVELTSRKVASGLNK..LGI
ACF35279.1_4CL_Pinus_radiata ISDHLPLHSYCFERVAEFA....DRPCLIDGATDRTYCFSEVELISRKVAAGLAK..LGL
CAA36850.1_4CL_Oryza_sativa_Japonica_Group ITNTLPLHRYCFERLPEVA....ARPCLIDGATGGVLTYADVDRLSRRLAAALRRAPLGL
BAA08366.2_4CL_Lithospermum_erythrorhizon ISNNLPLHTYCFQNASEYP....NRTCIIDSKTGKQYTFSETDSICRKVAAGLSN..LGI
AGP02119.1_4CL_Ocimum_basilicum IPNHLPLHTYCFQNLSQFA....DRQCLISGNDGKSYSFADTHLICRKVAAGLTK..LGI
AIY24982.1_4CL_Mangifera_indica IPNHLPLHSYCFENIAQVA....SRPCLINGSTGDVYTYSEVEITARKIAVGLNK..LGI
AHY94891.1_4CL_Prunella_vulgaris ISNDIPLHTYCFQNYSHYP....DRPCLLVGS..KSYSFADTHLMCRRVAAGLSQ..LGI
AHA42444.1_4CL_Cannabis_sativa IPKHLPLHTYCFGNKTQHD...LSHPCLIYGPTGQVFTYAEVDLTARKLASGLNK..LGV
pdb|3NI2_4CL1__Nicotiana_tabacum IPKNLPLHSYVLENLSNHS....SKPCLINGANGDVYTYADVELTARRVASGLNK..IGI
sp|P17814.2_4CL1_Oryza_sativa ITNTLPLHRYCFERLPEVA....ARPCLIDGATGGVLTYADVDRLSRRLAAALRRAPLGL
sp|Q42982.2_4CL2_Oryza_sativa IPSHLPLHEYCFARAAELP....DAPCLIAAATGRTYTFAETRLLCRRAAAALHR..LGV
sp|Q6ZAC1.1_4CL5_Oryza_sativa IPDHLPLHDYVFERLADRR....DRACLIDGATGETLSFGDVDALSRRVAAGLSS..IGV
sp|Q6ETN3.1_4CL3_Oryza_sativa IDNSMTLQEYCFARMAEVG....ARPCLIDGQTGESYTYAEVESASRRAAAGLRR..MGV
sp|Q67W82.1_4CL4_Oryza_sativa IPSHLTLQAYCFEKLPEVA....ARPCLIDGQTGAVYSYGEVEELSRRAAAGLRR..LGV
sp|Q9S725.2_4CL2_Arabidopsis_thaliana IPNHLPLHDYIFENISEFA....AKPCLINGPTGEVYTYADVHVTSRKLAAGLHN..LGV
sp|Q42524.1_4CL1_Arabidopsis_thaliana IPNHLSLHDYIFQNISEFA....TKPCLINGPTGHVYTYSDVHVISRQIAANFHK..LGV
sp|Q9S777.1_4CL3_Arabidopsis_thaliana IPNHLPLHTYCFEKLSSVS....DKPCLIVGSTGKSYTYGETHLICRRVASGLYK..LGI
sp|P31684.1_4CL1_Solanum_tuberosum IPKHLPLHSYCFENLSEFN....SRPCLIDGANDRIYTYAEVELTSRKVAVGLNK..LGI
sp|P31685.1_4CL2_Solanum_tuberosum IPKHLPLHSYCFENLSEFN....SRPCLIDGANDRIYTYAEVELTSRKVAVGLNK..LGI
AAG43823.1_4CL_Capsicum_annuum IPKHLPLHSYCFENISEFN....SRPCLIDGANDQIYSYAEVELTSRKVAVGLNK..LGV
BAA08365.1_4CL_Lithospermum_erythrorhizon IPKHLPLHSYCGENISQFS....SRPCLINGSNDRVYTYAEVEITSRKVAAGLHK..HGI
BAA08366.2_4CL_Lithospermum_erythrorhizon ISNNLPLHTYCFQNASEYP....NRTCIIDSKTGKQYTFSETDSICRKVAAGLSN..LGI
sp|P14912.1_4CL1_Petroselinum_crispum IPKHLPLHTYCFENISKVG....DKSCLINGATGETFTYSQVELLSRKVASGLNK..LGI
sp|P14913.1_4CL2_Petroselinum_crispum IPKHLPLHTYCFENISKVG....DKSCLINGATGETFTYSQVELLSRKVASGLNK..LGI
AAF91309.1_4CL2_Rubus_idaeus IPNHLPLHTYCFENISQFH....DRPCLINGNTGETFTYAEVELTSRRVAAGLDK..LGI
AAF91310.1_4CL1_Rubus_idaeus IPNHLPLHSYIFQNKSHLT....SKPCIINGTTGDIHTYAKFKLTARKVASGLNK..LGI
sp|O48868_4CL2_Populus_trichocarpa_x_Popul IPNHLPLHTYCFEKLSQFK....DNPCLINGPTGDIYTYADVELTSRKVASGLYK..LGL
AAK58908.1_4CL3_Populus_trichocarpa_x_Popu IPKNLPLHSYVLENLSKYS....SKPCLINGANGDVYTYADVELTARRVASGLNK..IGI
AAL56850.1_4CL_Populus_tomentosa IPKNLPLHSYVLENLSKHS....SKPCLINGANGDVYTCADVELTARRVASGLNK..IGI
sp|O81139_4CL_Populus_tremuloides IPKNLPLHSYVLENLSKHS....SKPCLINGANGDVYTYADVELTARRVASGLNK..IGI
AAL02144.1_4CL_Populus_tomentosa IPKNLPLHSYVLENLSNHS....SKPCLINGANGDVYTYADVELTARRVASGLNK..IGI
AAL35216.1_4CL_Amorpha_fruticosa ISKHLPLHSYCFENLSEFG....SRPCLISAPTGDVYTYYDVELTARRVASGLNK..LGV
sp|P41636.1_4CL_Pinus_taeda ISDHLPLHSYCFERVAEFA....DRPCLIDGATDRTYCFSEVELISRKVAAGLAK..LGL
AAF37732.1_4CL1_Lolium_perenne IPTHMPLHDYCFATAASAP....DAPCLITAATGKTYTFAETHLLCRKAAAALHG..LGV
sp|Q42982.2_4CL2_Oryza_sativa IPSHLPLHEYCFARAAELP....DAPCLIAAATGRTYTFAETRLLCRRAAAALHR..LGV
sp|P31687.2_4CL2_Glycine_max ISNHLPLHSYCFQNLSQFA....HRPCLIVGPASKTFTYADTHLISSKIAAGLSN..LGI
sp|O81140_4CL_Populus_tremuloides ISNDLPLHAYCFENLSDFS....DRPCLISGSTGKTYSFAETHLISRKVAAGLSN..LGI
AAK58909.1_4CL4_Populus_trichocarpa_x_Popu ISNHLPLHAYCFENLSDFS....DRPCLISGSTGKTYSFAETHLISRKVAAGLSN..LGI
sp|P17814.2_4CL1_Oryza_sativa ITNTLPLHRYCFERLPEVA....ARPCLIDGATGGVLTYADVDRLSRRLAAALRRAPLGL
AAF37733.1_4CL2_Lolium_perenne IPTHLTLQDYCFQRLPELS....ARACLIDGATGAALTYGEVDALSRRCAAGLRR..LGV
AAF37734.1_4CL3_Lolium_perenne INNEQTLQSYCFEKMAEVA....SRPCIIDGQTGASYTYTEVDSLTRRAAAGLRR..MGV
acc

```
pdb|5BSR_4CL2a_Nicotiana_tabacum              TT   β3          α3              β4     TT       α4            β5     η2
                                                      80         90        100        110        120        130
pdb|5BSR_4CL2a_Nicotiana_tabacum              QPKDTIMILLPNSPEFVFAFIGASYLGAISTMANPLFTPAEVVKQAKASSAKIIVTQACH
TRINITY_DN32164_c5_g1_i2                      KKGDVIMILLQNCPEFVFSFMGASMIGAITTTANPFYTKNEIFKQLTSSRAKLIITQSLY
pdb|3A9U_4CL_Populus_tomentosa                QQGDVIMLFLPSSPEFVLAFLGASHRGAIITAANPFSTPAELAKHAKASRAKLLITQACY
BAB01715.1_4CL_Arabidopsis_thaliana           RHGDVVMLLLPNSPEFALSFLAVAYLGAVSTTANPFYTQPEIAKQAKASAAKMIITKKCL
sp|O48869_4CL1_Populus_trichocarpa_x_Popul    KQGDVILLLLQNSPEFVFAFLGASIIGAISTTANPFYTPAEVAKQATASKAKLIITQAVY
ACF35279.1_4CL_Pinus_radiata                  QQGQVVMLLLPNCIEFAFVFMGASVRGAIVTTANPFYKPGEIAKQAKAAGARIIVTLAAY
CAA36850.1_4CL_Oryza_sativa_Japonica_Group    RRGGVVMSLLRNSPEFVLSFFAASRVGAAVTTANPMSTPHEIESQLAAAGATVVITESMA
BAA08366.2_4CL_Lithospermum_erythrorhizon     QKGDVIMVLLQNCAEFVFTFMGASIIGAVITTGNPFYTTAEIFKQVNVSNTKLIITQSNY
AGP02119.1_4CL_Ocimum_basilicum               KKGDVIMVLLQNCAEFVFSFIGASMIGAVTTTANPFCTSKEIFKQFNASRSKLIVTQSQY
AIY24982.1_4CL_Mangifera_indica               QQYDVILLLLPNFPEFILSFLGASYRGAITAANPFFTPAEIAKQADASNAKLIITQACY
AHY94891.1_4CL_Prunella_vulgaris              KKGDVVMALLQNSAEFVFTFMGASMIGAVTTTANPFCTNKEIYKQFHASKSKLIVTQSIY
AHA42444.1_4CL_Cannabis_sativa                KQGDVVMVLLPNSPEYVFTFLGASYRGAMTTAANPFFTAAEIQKQAKASKTKLIVTQASY
pdb|3NI2_4CL1__Nicotiana_tabacum              QQGDVIMLFLPSSPEFVLAFLGASHRGAIITAANPFSTPAELAKHAKASRAKLLITQACY
sp|P17814.2_4CL1_Oryza_sativa                 RRGGVVMSLLRNSPEFVLSFFAASRVGAAVTTANPMSTPHEIESQLAAAGATVVITESMA
sp|Q42982.2_4CL2_Oryza_sativa                 GHGDRVMVLLQNCVEFAVAFFAASFLGAVTTAANPFCTPQEIHKQFKASGVKLILTQSVY
sp|Q6ZAC1.1_4CL5_Oryza_sativa                 CHGSTVMLLLPNSVEFAVAFLASSRLGAVTTTANPLHTPPEIAKQVAASGATVVVTEPAF
sp|Q6ETN3.1_4CL3_Oryza_sativa                 GKGDVVMSLLRNCPEFAFSFLGAARLGAATTTANPFYTPHEVHRQAEAAGARVIVTEACA
sp|Q67W82.1_4CL4_Oryza_sativa                 GKGDVVMSLLRNCPEFAFTFLGAARLGAATTTANPFYTPHEIHRQASAAGARVIVTEACA
sp|Q9S725.2_4CL2_Arabidopsis_thaliana         KQHDVVMILLPNSPEVVLTFLAASFIGAITTSANPFFTPAEISKQAKASAAKLIVTQSRY
sp|Q42524.1_4CL1_Arabidopsis_thaliana         NQNDVVMLLLPNCPEFVLSFLAASFRGATATAANPFFTPAEIAKQAKASNTKLIITEARY
sp|Q9S777.1_4CL3_Arabidopsis_thaliana         RKGDVIMILLQNSAEFVFSFMGASMIGAVSTTANPFYTSQELYKQLKSSGAKLIITHSQY
sp|P31684.1_4CL1_Solanum_tuberosum            QQKDTIMILLPNCPEFVFAFIGASYLGAISTMANPLFTPAEVVKQAKASSAKIVITQACF
sp|P31685.1_4CL2_Solanum_tuberosum            QQKDTIMILLPNCPEFVFAFIGASYLGAISTMANPLFTPAEVVKQAKASSAKIVITQACF
AAG43823.1_4CL_Capsicum_annuum                QQKDTIMILLPNSPEFVFAFMGASYLGAISTMANPLFTPAEVLKQAKASSAKIIITLACY
BAA08365.1_4CL_Lithospermum_erythrorhizon     KQTETIMLLLPNCPEFVFAFLGASYIGAVSTTANPFFTSSEIIKQAKASKTKLIITVSTT
BAA08366.2_4CL_Lithospermum_erythrorhizon     QKGDVIMVLLQNCAEFVFTFMGASIIGAVITTGNPFYTTAEIFKQVNVSNTKLIITQSNY
sp|P14912.1_4CL1_Petroselinum_crispum         QQGDTIMLLLPNSPEYFFAFLGASYRGAISTMANPFFTSAEVIKQLKASQAKLIITQACY
sp|P14913.1_4CL2_Petroselinum_crispum         QQGDTIMLLLPNSPEYFFAFLGASYRGAISTMANPFFTSAEVIKQLKASLAKLIITQACY
AAF91309.1_4CL2_Rubus_idaeus                  QQNDVVMLLLQNCAEFAFAFLGASYIGAMSTTANPFYTPAEVAKQAKASNAKLIITQSAY
AAF91310.1_4CL1_Rubus_idaeus                  EKGDVFMLLLPNTSEFVFAFLGASFCGAMMTAANPFFTPAEIAKQAKASKAKLIITFACY
sp|O48868_4CL2_Populus_trichocarpa_x_Popul    QQGDVILLLLQNSPEFVFAFLGASFIGAISSTANPFYTSAEIAKQATASKAKLIITHAAY
AAK58908.1_4CL3_Populus_trichocarpa_x_Popu    QQGDVIMLFLPSSPEFVLAFLGASHRGAIVTAANPFSTPAELAKHAKPPRTKLLITQACY
AAL56850.1_4CL_Populus_tomentosa              QQGDVIMLFLPSSPEFVLAFLGASHRGAIITAANPFSTPAELAKHAKASRAKLLITQACY
sp|O81139_4CL_Populus_tremuloides             QQGDVIMLFLPSSPEFVLAFLGASHRGAMITAANPFSTPAELAKHAKASRAKLLITQACY
AAL02144.1_4CL_Populus_tomentosa              QQGDVIMLFLPSSPEFVLAFLGASHRGAIITAANPFSTPAELAKHAKASRAKLLITQACY
AAL35216.1_4CL_Amorpha_fruticosa              QQGDVIMLLLPNSPEFVFSFLGASYRGAMITAANPFFTSAEIAKQAKASNTKLLITQASY
sp|P41636.1_4CL_Pinus_taeda                   QQGQVVMLLLPNCIEFAFVFMGASVRGAIVTTANPFYKPGEIAKQAKAAGARIIVTLAAY
AAF37732.1_4CL1_Lolium_perenne                RHGDRIMLLLQNSVEFALAFLGASMLGAVSTAANPFCTPQEIHKQLVASGAKLVVTQSAY
sp|Q42982.2_4CL2_Oryza_sativa                 GHGDRVMVLLQNCVEFAVAFFAASFLGAVTTAANPFCTPQEIHKQFKASGVKLILTQSVY
sp|P31687.2_4CL2_Glycine_max                  LKGDVVMILLQNSADFVFSFLAISMIGAVATTANPFYTAPEIFKQFTVSKAKLIITQAMY
sp|O81140_4CL_Populus_tremuloides             KKGDVIMTLLQNCPEFVFSFIGASMIGAVITTANPFYTQSEIFKQFSASRAKLIITQSQY
AAK58909.1_4CL4_Populus_trichocarpa_x_Popu    KKGDVIMTLLQNCPEFVFSFMGASMIGAVTTTVNPFYTPGEIFKQFSASRAKLIITQSQH
sp|P17814.2_4CL1_Oryza_sativa                 RRGGVVMSLLRNSPEFVLSFFAASRVGAAVTTANPMSTPHEIESQLAAAGATVVITESMA
AAF37733.1_4CL2_Lolium_perenne                GKGDVVMALLRNCPEFAFVFLGAARLGAATTTANPFYTPHEIHRQATAAGARVIVTEACA
AAF37734.1_4CL3_Lolium_perenne                GKGDVVMNLLRNCPEFAFSFLGAARLGAATTTANPFYTPHEIHRQAEAAGAKLIVTEACA
                                         acc
```

```
pdb|5BSR_4CL2a_Nicotiana_tabacum                        α5                   β6                  T.T β7  α6        η3
                                                 140                    150                    160       170
pdb|5BSR_4CL2a_Nicotiana_tabacum              VNKVKDYA..........FEND.VKIICID.......SAPE.GCLHF.SVL.TQ.ANEH
TRINITY_DN32164_c5_g1_i2                      VEKLKDS...........EEN.LILITIDDP.......PE.NCLHF.SVL.SE.SNET
pdb|3A9U_4CL_Populus_tomentosa                YEKVKDFA..........RESD.VKVMCVD.......SAPD.GCLHF.SEL.TQ.ADEN
BAB01715.1_4CL_Arabidopsis_thaliana           VDKLTNLK............NDGVLIVCLDDDGDNGVVSSSDDGCVSF.TEL.TQ.ADET
sp|O48869_4CL1_Populus_trichocarpa_x_Popul    AEKVQEFV..........KENVHVKIVTVD.......SPPE.NYLHF.SEL.TN.SDED
ACF35279.1_4CL_Pinus_radiata                  VEKLADLQ..........SHD..VLVITID.......DAPKEGCQHI.SVL.TE.ADET
CAA36850.1_4CL_Oryza_sativa_Japonica_Group    ADKLPSHSHGA............LTVVLID.......ERRD.GCLHFWDDL.MS.EDEA
BAA08366.2_4CL_Lithospermum_erythrorhizon     VDKLRNTTINESDNKYPKLGED.FKVITIDTP........PE.NCLPF.SLL.IENTQEN
AGP02119.1_4CL_Ocimum_basilicum               VDKLRDTG.....DDSVVFGED.FSVVTIDAP........PE.GCLDF.SVL.CE.ADEA
AIY24982.1_4CL_Mangifera_indica               VDKVKALA..........KKRD.VKIMCID.......SAPD.GCLHF.SEL.TQ.ADEN
AHY94891.1_4CL_Prunella_vulgaris              VDKLRDTG.....DDSLVLGVD.FPVVTVDSP........PQ.GCLPF.SAL.SD.ADPS
AHA42444.1_4CL_Cannabis_sativa                YDKVKDLD..........QEGL.VKFVCVDS.......PVPE.GWFHF.SELCNN.SDEN
pdb|3NI2_4CL1__Nicotiana_tabacum              YEKVKDFA..........RESD.VKVMCVD.......SAPD.GCLHF.SEL.TQ.ADEN
sp|P17814.2_4CL1_Oryza_sativa                 ADKLPSHSHGA............LTVVLID.......ERRD.GCLHFWDDL.MS.EDEA
sp|Q42982.2_4CL2_Oryza_sativa                 VDKLRQHEAFPRIDACTVGDDT.LTVITIDDDE.....ATPE.GCLPF.WDL.IADADEG
sp|Q6ZAC1.1_4CL5_Oryza_sativa                 VAKVSGLAG............VTVVATG.......GGAE.RCASF.AGL.AA.ADGS
sp|Q6ETN3.1_4CL3_Oryza_sativa                 VEKVREFA..........AERG.VPVVTVD.......GAFD.GCVEFREVL...AAEEL
sp|Q67W82.1_4CL4_Oryza_sativa                 VEKVRGFA..........ADRG.IPVVAVD.......GDFD.GCVGFGEAMLDASIEPL
sp|Q9S725.2_4CL2_Arabidopsis_thaliana         VDKIKNLQ..........N.DG.VLIVTTDSD......AIPE.NCLRF.SEL.TQ.SEEP
sp|Q42524.1_4CL1_Arabidopsis_thaliana         VDKIKPLQ..........NDDG.VVIVCIDDNES....VPIPE.GCLRF.TEL.TQ.STTE
sp|Q9S777.1_4CL3_Arabidopsis_thaliana         VDKLKN............LGEN.LTLITTDEP.......TPE.NCLPF.STL.IT.DDET
sp|P31684.1_4CL1_Solanum_tuberosum            AGKVKDYA..........IEND.LKVICVD.......SVPE.GCVHF.SEL.IQ.SDEH
sp|P31685.1_4CL2_Solanum_tuberosum            AGKVKDYA..........IEND.LKVICVD.......SAPE.GCVHF.SEL.IQ.SDEH
AAG43823.1_4CL_Capsicum_annuum                IGKVKDYA..........TEND.VKLICID.......SAPE.GCIHF.SEL.TQ.SDEH
BAA08365.1_4CL_Lithospermum_erythrorhizon     VPKLKDFS..........QENH.VKIMCID.......DKID.GCLHFSSDL.EN.SDET
BAA08366.2_4CL_Lithospermum_erythrorhizon     VDKLRNTTINESDNKYPKLGED.FKVITIDTP........PE.NCLPF.SLL.IENTQEN
sp|P14912.1_4CL1_Petroselinum_crispum         VDKVKDYA..........AEKN.IQIICID.......DAPQ.DCLHF.SKL.ME.ADES
sp|P14913.1_4CL2_Petroselinum_crispum         VDKVKDYA..........AEKN.IQIICID.......DAPQ.DCLHF.SKL.ME.ADES
AAF91309.1_4CL2_Rubus_idaeus                  VDKVKDFA..........KLND.VKVMCVDE......TSSE.DVLHF.SEL.MS.ADES
AAF91310.1_4CL1_Rubus_idaeus                  YDKVKDLS..........CDE..VKLMCIDS......PPPDSSCLHF.SEL.TQ.SDEN
sp|O48868_4CL2_Populus_trichocarpa_x_Popul    AEKVQQFA..........QENDHVKIMTID.......SLTE.NCLHF.SEL.TS.SDEN
AAK58908.1_4CL3_Populus_trichocarpa_x_Popu    YDKVKDFA..........RESD.VKVMCVD.......SAPD.GCLHF.SEL.TQ.ADEN
AAL56850.1_4CL_Populus_tomentosa              YEKVKDFA..........RESD.VKVMCVD.......SAPD.GCLHF.SEL.TQ.ADEN
sp|O81139_4CL_Populus_tremuloides             YEKVKDFA..........RESD.VKVMCVD.......SAPD.GASLF.RAH.TQ.ADEN
AAL02144.1_4CL_Populus_tomentosa              YEKVKDFA..........RESD.VKVMCVD.......SAPD.GCLHF.SEL.TQ.ADEN
AAL35216.1_4CL_Amorpha_fruticosa              YDKVKDL.............D.VKLVFVD.......SPPD.GHMHY.SEL.RE.ADES
sp|P41636.1_4CL_Pinus_taeda                   VEKLADLQ..........SHD..VLVITID.......DAPKEGCQHI.SVL.TE.ADET
AAF37732.1_4CL1_Lolium_perenne                VDKLR.HEAFPRI......GEA.LTVITIDEDD.....GTPD.GCQPF.WAL.VSAADEN
sp|Q42982.2_4CL2_Oryza_sativa                 VDKLRQHEAFPRIDACTVGDDT.LTVITIDDDE.....ATPE.GCLPF.WDL.IADADEG
sp|P31687.2_4CL2_Glycine_max                  VDKLRNHD.......GAKLGED.FKVVTVDDP........PE.NCLHF.SVL.SE.ANES
sp|O81140_4CL_Populus_tremuloides             VNKLGDSDCHE...NNQKPGED.FIVITIDDP........PE.NCLHF.NVL.VE.ASES
AAK58909.1_4CL4_Populus_trichocarpa_x_Popu    VNKLRDSDYHE...NNQKPEED.FIVITIDDP........PE.NCLHF.NVL.VE.ANES
sp|P17814.2_4CL1_Oryza_sativa                 ADKLPSHSHGA............LTVVLID.......ERRD.GCLHFWDDL.MS.EDEA
AAF37733.1_4CL2_Lolium_perenne                VEKVRAFA..........AERG.IPVVSVD.......EGVDGGCLPFAETLLGEESGER
AAF37734.1_4CL3_Lolium_perenne                VEKVLEFA..........AGRG.VPVVTVD.......GRRD.GCVDFAELIAGEELPEA
                                         acc
```

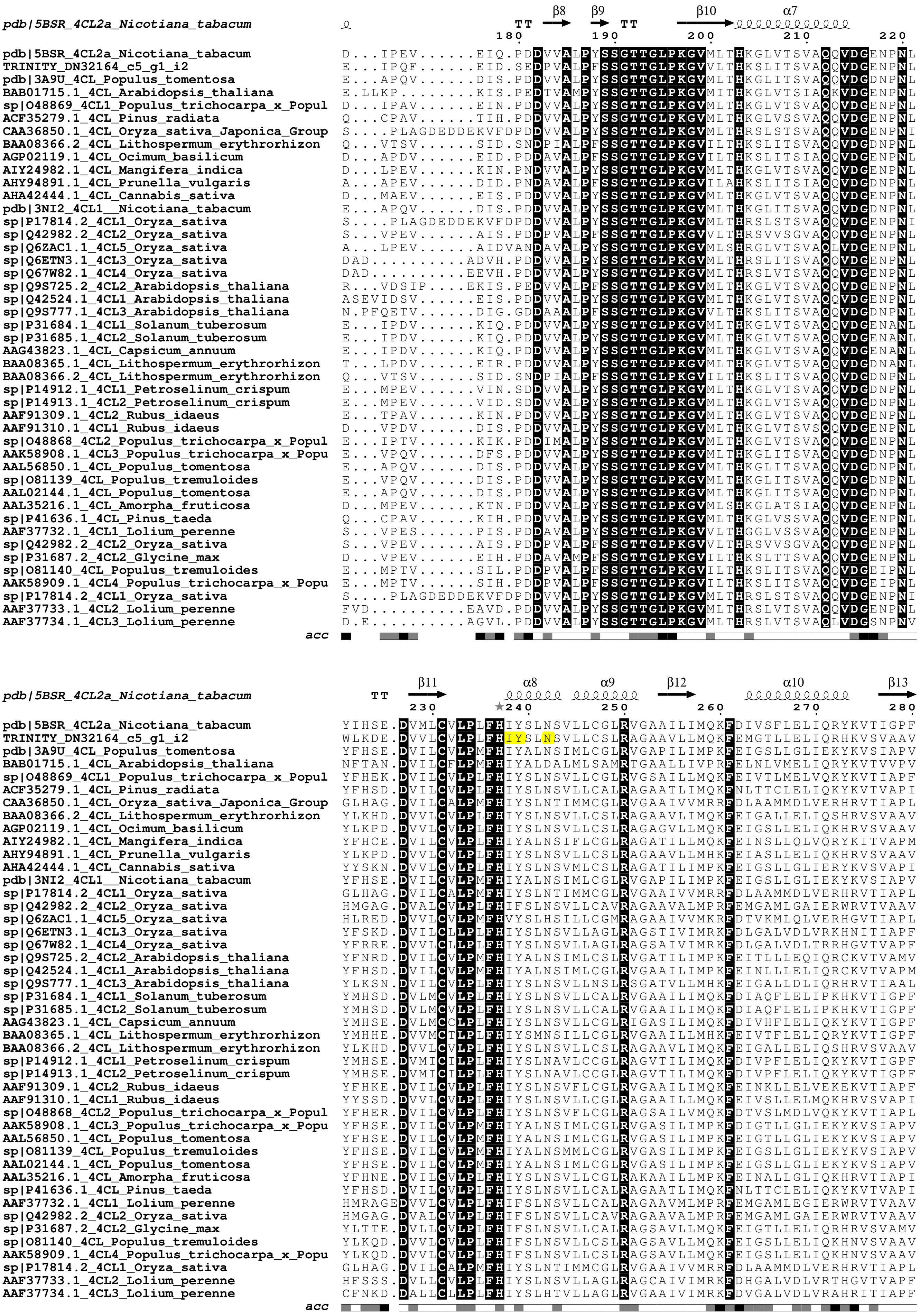

pdb|5BSR_4CL2a_Nicotiana_tabacum
β8
β9
β10
α7
180
190
200
210
220
acc
β11
α8
α9
β12
α10
β13
230
240
250
260
270
280

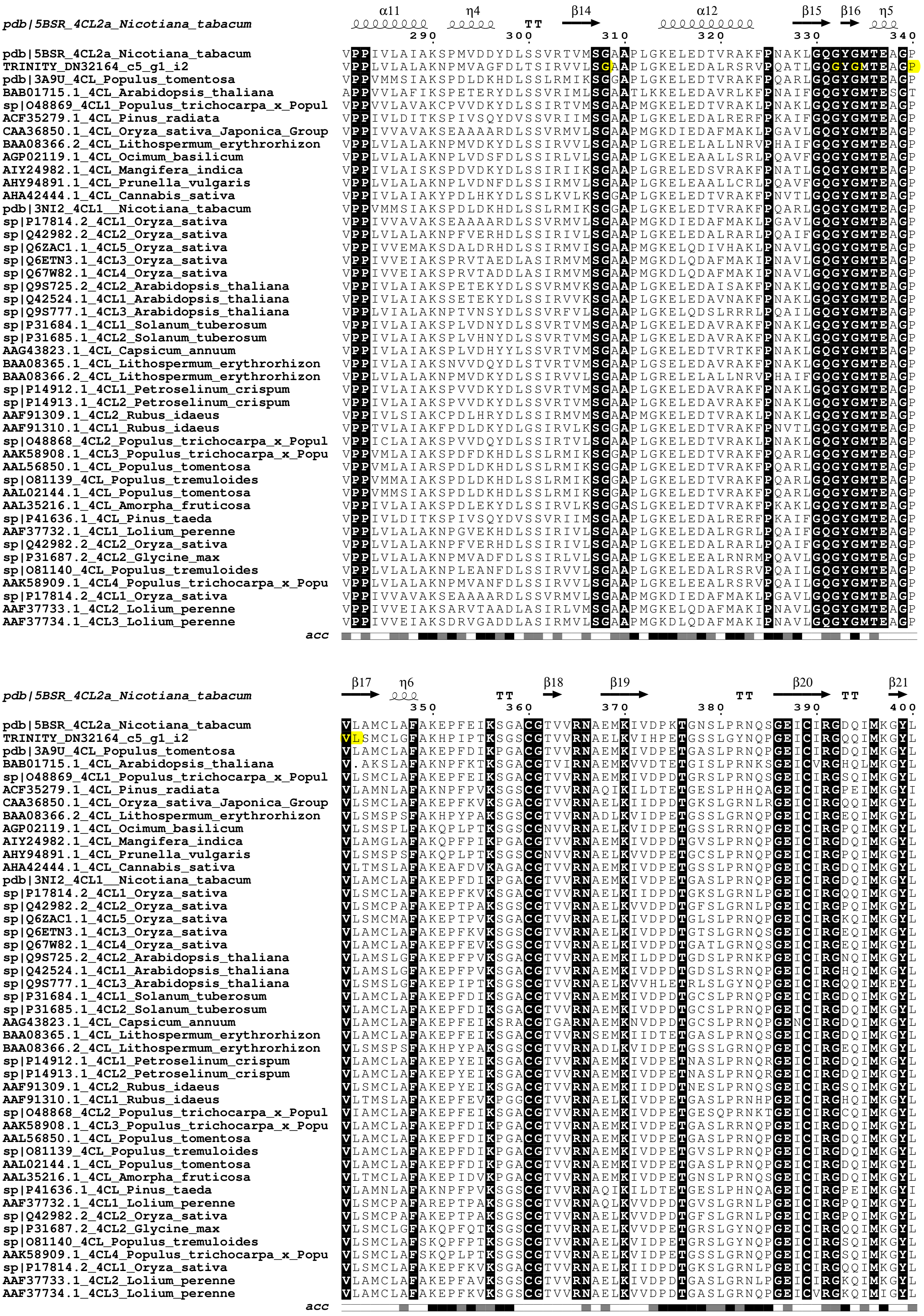
pdb|5BSR_4CL2a_Nicotiana_tabacum
α11 η4 TT β14 α12 β15 β16 η5
290 300 310 320 330 340
pdb|5BSR_4CL2a_Nicotiana_tabacum VPPIVLAIAKSPMVDDYDLSSVRTVMSGAAPLGKELEDTVRAKFPNAKLGQGYGMTEAGP
TRINITY_DN32164_c5_g1_i2 VPPLVLALAKNPMVAGFDLTSIRVVLSGAAPLGKELEDALRSRVPQATLGQGYGMTEAGP
pdb|3A9U_4CL_Populus_tomentosa VPPVMMSIAKSPDLDKHDLSSLRMIKSGGAPLGKELEDTVRAKFPQARLGQGYGMTEAGP
BAB01715.1_4CL_Arabidopsis_thaliana APPVVLAFIKSPETERYDLSSVRIMLSGAATLKKELEDAVRLKFPNAIFGQGYGMTESGT
sp|O48869_4CL1_Populus_trichocarpa_x_Popul VPPVVLAVAKCPVVDKYDLSSIRTVMSGAAPMGKELEDTVRAKLPNAKLGQGYGMTEAGP
ACF35279.1_4CL_Pinus_radiata VPPIVLDITKSPIVSQYDVSSVRIIMSGAAPLGKELEDALRERFPKAIFGQGYGMTEAGP
CAA36850.1_4CL_Oryza_sativa_Japonica_Group VPPIVVAVAKSEAAAARDLSSVRMVLSGAAPMGKDIEDAFMAKLPGAVLGQGYGMTEAGP
BAA08366.2_4CL_Lithospermum_erythrorhizon VPPLVLALAKNPMVDKYDLSSIRVVLSGAAPLGRELELALLNRVPHAIFGQGYGMTEAGP
AGP02119.1_4CL_Ocimum_basilicum VPPLVLALAKNPLVDSFDLSSIRLVLSGAAPLGKELEAALLSRLPQAVFGQGYGMTEAGP
AIY24982.1_4CL_Mangifera_indica VPPIVLAISKSPDVDKYDLSSIRIVLSGAAPLGKELEDAVRAKFPNATLGQGYGMTEAGP
AHY94891.1_4CL_Prunella_vulgaris VPPLVLALAKNPLVDNFDLSSIRMVLSGAAPLGKELEAALLCRLPQAVFGQGYGMTEAGP
AHA42444.1_4CL_Cannabis_sativa VPPIVLAIAKYPDLHKYDLSSLKVLKSGGAPLGKELEDTVRAKFPNVTLGQGYGMTEAGP
pdb|3NI2_4CL1__Nicotiana_tabacum VPPVMMSIAKSPDLDKHDLSSLRMIKSGGAPLGKELEDTVRAKFPQARLGQGYGMTEAGP
sp|P17814.2_4CL1_Oryza_sativa VPPIVVAVAKSEAAAARDLSSVRMVLSGAAPMGKDIEDAFMAKLPGAVLGQGYGMTEAGP
sp|Q42982.2_4CL2_Oryza_sativa VPPLVLALAKNPFVERHDLSSIRIVLSGAAPLGKELEDALRARLPQAIFGQGYGMTEAGP
sp|Q6ZAC1.1_4CL5_Oryza_sativa VPPIVVEMAKSDALDRHDLSSIRMVISGAAPMGKELQDIVHAKLPNAVLGQGYGMTEAGP
sp|Q6ETN3.1_4CL3_Oryza_sativa VPPIVVEIAKSPRVTAEDLASIRMVMSGAAPMGKDLQDAFMAKIPNAVLGQGYGMTEAGP
sp|Q67W82.1_4CL4_Oryza_sativa VPPIVVEIAKSPRVTADDLASIRMVMSGAAPMGKDLQDAFMAKIPNAVLGQGYGMTEAGP
sp|Q9S725.2_4CL2_Arabidopsis_thaliana VPPIVLAIAKSPETEKYDLSSVRMVKSGAAPLGKELEDAISAKFPNAKLGQGYGMTEAGP
sp|Q42524.1_4CL1_Arabidopsis_thaliana VPPIVLAIAKSSETEKYDLSSIRVVKSGAAPLGKELEDAVNAKFPNAKLGQGYGMTEAGP
sp|Q9S777.1_4CL3_Arabidopsis_thaliana VPPLVIALAKNPTVNSYDLSSVRFVLSGAAPLGKELQDSLRRRLPQAILGQGYGMTEAGP
sp|P31684.1_4CL1_Solanum_tuberosum VPPIVLAIAKSPLVDNYDLSSVRTVMSGAAPLGKELEDAVRAKFPNAKLGQGYGMTEAGP
sp|P31685.1_4CL2_Solanum_tuberosum VPPIVLAIAKSPLVHNYDLSSVRTVMSGAAPLGKELEDAVRAKFPNAKLGQGYGMTEAGP
AAG43823.1_4CL_Capsicum_annuum VPPIVLAIAKSPLVDHYYLSSVRTVMSGAAPLGKELEDTVRTKFPNAKLGQGYGMTEAGP
BAA08365.1_4CL_Lithospermum_erythrorhizon VPPIVLAIAKSNVVDQYDLSTVRTVMSGAAPLGSELEDAVRAKFPNAKLGQGYGMTEAGP
BAA08366.2_4CL_Lithospermum_erythrorhizon VPPLVLALAKNPMVDKYDLSSIRVVLSGAAPLGRELELALLNRVPHAIFGQGYGMTEAGP
sp|P14912.1_4CL1_Petroselinum_crispum VPPIVLAIAKSPVVDKYDLSSVRTVMSGAAPLGKELEDAVRAKFPNAKLGQGYGMTEAGP
sp|P14913.1_4CL2_Petroselinum_crispum VPPIVLAIAKSPVVDKYDLSSVRTVMSGAAPLGKELEDAVRAKFPNAKLGQGYGMTEAGP
AAF91309.1_4CL2_Rubus_idaeus VPPIVLSIAKCPDLHRYDLSSIRMVMSGAAPMGKELEDTVRAKLPNAKLGQGYGMTEAGP
AAF91310.1_4CL1_Rubus_idaeus VPPTVLAIAKFPDLDKYDLGSIRVLKSGGAPLGKELEDTVRAKFPNVTLGQGYGMTEAGP
sp|O48868_4CL2_Populus_trichocarpa_x_Popul VPPICLAIAKSPVVDQYDLSSIRTVLSGAAPLGKELEDTVRAKLPNAKLGQGYGMTEAGP
AAK58908.1_4CL3_Populus_trichocarpa_x_Popu VPPVMLAIAKSPDFDKHDLSSLRMIKSGGAPLGKELEDTVRAKFPQARLGQGYGMTEAGP
AAL56850.1_4CL_Populus_tomentosa VPPVMLAIARSPDLDKHDLSSLRMIKSGGAPLGKELEDTVRAKFPQARLGQGYGMTEAGP
sp|O81139_4CL_Populus_tremuloides VPPVMMAIAKSPDLDKHDLSSLRMIKSGGAPLGKELEDTVRAKFPQARLGQGYGMTEAGP
AAL02144.1_4CL_Populus_tomentosa VPPVMMSIAKSPDLDKHDLSSLRMIKSGGAPLGKELEDTVRAKFPQARLGQGYGMTEAGP
AAL35216.1_4CL_Amorpha_fruticosa VPPIVLAIAKSPDLEKYDLSSIRVLKSGGASLGKELEDTVRAKFPKAKLGQGYGMTEAGP
sp|P41636.1_4CL_Pinus_taeda VPPIVLDITKSPIVSQYDVSSVRIIMSGAAPLGKELEDALRERFPKAIFGQGYGMTEAGP
AAF37732.1_4CL1_Lolium_perenne VPPLVLALAKNPGVEKHDLSSIRIVLSGAAPLGKELEDALRGRLPQAIFGQGYGMTEAGP
sp|Q42982.2_4CL2_Oryza_sativa VPPLVLALAKNPFVERHDLSSIRIVLSGAAPLGKELEDALRARLPQAIFGQGYGMTEAGP
sp|P31687.2_4CL2_Glycine_max VPPLVLALAKNPMVADFDLSSIRLVLSGAAPLGKELEEALRNRMPQAVLGQGYGMTEAGP
sp|O81140_4CL_Populus_tremuloides VPPLVLALAKNPLEANFDLSSIRVVLSGAAPLGKELEDALRSRVPQAILGQGYGMTEAGP
AAK58909.1_4CL4_Populus_trichocarpa_x_Popu VPPLVLALAKNPMVANFDLSSIRVVLSGAAPLGKELEEALRSRVPQAILGQGYGMTEAGP
sp|P17814.2_4CL1_Oryza_sativa VPPIVVAVAKSEAAAARDLSSVRMVLSGAAPMGKDIEDAFMAKLPGAVLGQGYGMTEAGP
AAF37733.1_4CL2_Lolium_perenne VPPIVVEIAKSARVTAADLASIRLVMSGAAPMGKELQDAFMAKIPNAVLGQGYGMTEAGP
AAF37734.1_4CL3_Lolium_perenne VPPIVVEIAKSDRVGADDLASIRMVLSGAAPMGKDLQDAFMAKIPNAVLGQGYGMTEAGP
acc
pdb|5BSR_4CL2a_Nicotiana_tabacum
β17 η6 TT β18 β19 TT β20 TT β21
350 360 370 380 390 400
pdb|5BSR_4CL2a_Nicotiana_tabacum VLAMCLAFAKEPFEIKSGACGTVVRNAEMKIVDPKTGNSLPRNQSGEICIRGDQIMKGYL
TRINITY_DN32164_c5_g1_i2 VLSMCLGFAKHPIPTKSGSCGTVVRNAELKVIHPETGSSLGYNQPGEICIRGQQIMKGYL
pdb|3A9U_4CL_Populus_tomentosa VLAMCLAFAKEPFDIKPGACGTVVRNAEMKIVDPETGASLPRNQPGEICIRGDQIMKGYL
BAB01715.1_4CL_Arabidopsis_thaliana V.AKSLAFAKNPFKTKSGACGTVIRNAEMKVVDTETGISLPRNKSGEICVRGHQLMKGYL
sp|O48869_4CL1_Populus_trichocarpa_x_Popul VLSMCLAFAKEPFEIKSGACGTVVRNAEMKIVDPETGRSLPRNQSGEICIRGSQIMKGYL
ACF35279.1_4CL_Pinus_radiata VLAMNLAFAKNPFPVKSGSCGTVVRNAQIKILDTETGESLPHHQAGEICIRGPEIMKGYI
CAA36850.1_4CL_Oryza_sativa_Japonica_Group VLSMCLAFAKEPFKVKSGACGTVVRNAELKIIDPDTGKSLGRNLRGEICIRGQQIMKGYL
BAA08366.2_4CL_Lithospermum_erythrorhizon VLSMSPSFAKHPYPAKSGSCGTVVRNADLKVIDPETGSSLGRNQPGEICIRGEQIMKGYL
AGP02119.1_4CL_Ocimum_basilicum VLSMSPLFAKQPLPTKSGSCGNVVRNAELKVIDPETGSSLRRNQPGEICIRGPQIMKGYL
AIY24982.1_4CL_Mangifera_indica VLAMGLAFAKQPFPIKPGACGTVVRNAEMKIVDPDTGVSLPRNQPGEICIRGDQIMKGYL
AHY94891.1_4CL_Prunella_vulgaris VLSMSPSFAKQPLPTKSGSCGNVVRNAELKVVDPETGCSLPRNQPGEICIRGPQIMKGYL
AHA42444.1_4CL_Cannabis_sativa VLTMSLAFAKEAFDVKAGACGTVVRNAEMKIVDPETGSSLPRNQPGEICIRGDQIMKGYL
pdb|3NI2_4CL1__Nicotiana_tabacum VLAMCLAFAKEPFDIKPGACGTVVRNAEMKIVDPETGASLPRNQPGEICIRGDQIMKGYL
sp|P17814.2_4CL1_Oryza_sativa VLSMCLAFAKEPFKVKSGACGTVVRNAELKIIDPDTGKSLGRNLPGEICIRGQQIMKGYL
sp|Q42982.2_4CL2_Oryza_sativa VLSMCPAFAKEPTPAKSGSCGTVVRNAELKVVDPDTGFSLGRNLPGEICIRGPQIMKGYL
sp|Q6ZAC1.1_4CL5_Oryza_sativa VLSMCMAFAKEPTPVKSGACGTVVRNAELKIVDPDTGLSLPRNQPGEICIRGKQIMKGYL
sp|Q6ETN3.1_4CL3_Oryza_sativa VLAMCLAFAKEPFKVKSGSCGTVVRNAELKIVDPDTGTSLGRNQSGEICIRGEQIMKGYL
sp|Q67W82.1_4CL4_Oryza_sativa VLAMCLAFAKEPFEVKSGSCGTVVRNAELKIVDPDTGATLGRNQSGEICIRGEQIMKGYL
sp|Q9S725.2_4CL2_Arabidopsis_thaliana VLAMSLGFAKEPFPVKSGACGTVVRNAEMKILDPDTGDSLPRNKPGEICIRGNQIMKGYL
sp|Q42524.1_4CL1_Arabidopsis_thaliana VLAMSLGFAKEPFPVKSGACGTVVRNAEMKIVDPDTGDSLSRNQPGEICIRGHQIMKGYL
sp|Q9S777.1_4CL3_Arabidopsis_thaliana VLSMSLGFAKEPIPTKSGSCGTVVRNAELKVVHLETRLSLGYNQPGEICIRGQQIMKEYL
sp|P31684.1_4CL1_Solanum_tuberosum VLAMCLAFAKEPFDIKSGACGTVVRNAEMKIVDPDTGCSLPRNQPGEICIRGDQIMKGYL
sp|P31685.1_4CL2_Solanum_tuberosum VLAMCLAFAKEPFDIKSGACGTVVRNAEMKIVDPDTGCSLPRNQPGEICIRGDQIMKGYL
AAG43823.1_4CL_Capsicum_annuum VLAMCLAFAKEPFEIKSRACGTGARNAEMKNVDPDTGCSLPRNQPGENCIRGDQIMKGYL
BAA08365.1_4CL_Lithospermum_erythrorhizon VLAMCLAFAKEPFEIKSGACGTVVRNSEMKIIDTETGASLPRNQSGEICIRGDQIMKGYL
BAA08366.2_4CL_Lithospermum_erythrorhizon VLSMSPSFAKHPYPAKSGSCGTVVRNADLKVIDPETGSSLGRNQPGEICIRGEQIMKGYL
sp|P14912.1_4CL1_Petroselinum_crispum VLAMCLAFAKEPYEIKSGACGTVVRNAEMKIVDPETNASLPRNQRGEICIRGDQIMKGYL
sp|P14913.1_4CL2_Petroselinum_crispum VLAMCLAFAKEPYEIKSGACGTVVRNAEMKIVDPETNASLPRNQRGEICIRGDQIMKGYL
AAF91309.1_4CL2_Rubus_idaeus VLSMCLAFAKEPYEIKSGACGTVVRNAEMKIIDPDTNESLPRNQSGEICIRGSQIMKGYL
AAF91310.1_4CL1_Rubus_idaeus VLTMSLAFAKEPFEVKPGGCGTVVRNAELKIVDPETGASLPRNHPGEICIRGHQIMKGYL
sp|O48868_4CL2_Populus_trichocarpa_x_Popul VIAMCLAFAKEPFEIKSGACGTVVRNAEMKIVDPETGESQPRNKTGEICIRGCQIMKGYL
AAK58908.1_4CL3_Populus_trichocarpa_x_Popu VLAMCLAFAKEPFDIKPGACGTVVRNAEMKIVDPETGASLRRNQPGEICIRGDQIMKGYL
AAL56850.1_4CL_Populus_tomentosa VLAMCLAFAKEPFDIKPGACGTVVRNAEMKIVDPETGASLPRNQPGEICIRGDQIMKGYL
sp|O81139_4CL_Populus_tremuloides VLAMCLAFAKEPFDIKPGACGTVVRNAEMKIVDPETGVSLPRNQPGEICIRGDQIMKGYL
AAL02144.1_4CL_Populus_tomentosa VLAMCLAFAKEPFDIKPGACGTVVRNAEMKIVDPETGASLPRNQPGEICIRGDQIMKGYL
AAL35216.1_4CL_Amorpha_fruticosa VLTMCLAFAKEPIDVKPGACGTVVRNAEMKIVDPETGNSLPRNQSGEICIRGDQIMKGYL
sp|P41636.1_4CL_Pinus_taeda VLAMNLAFAKNPFPVKSGSCGTVVRNAQIKILDTETGESLPHNQAGEICIRGPEIMKGYI
AAF37732.1_4CL1_Lolium_perenne VLSMCPAFAREPTPAKSGSCGTVVRNAQLKVVDPDTGVSLGRNLPGEICIRGPQIMKGYL
sp|Q42982.2_4CL2_Oryza_sativa VLSMCPAFAKEPTPAKSGSCGTVVRNAELKVVDPDTGFSLGRNLPGEICIRGPQIMKGYL
sp|P31687.2_4CL2_Glycine_max VLSMCLGFAKQPFQTKSGSCGTVVRNAELKVVDPETGRSLGYNQPGEICIRGQQIMKGYL
sp|O81140_4CL_Populus_tremuloides VLSMCLAFSKQPFPTKSGSCGTVVRNAELKVIDPETGRSLGYNQPGEICIRGSQIMKGYL
AAK58909.1_4CL4_Populus_trichocarpa_x_Popu VLSMCLAFSKQPLPTKSGSCGTVVRNAELKVIDPETGSSLGRNQPGEICIRGSQIMKGYL
sp|P17814.2_4CL1_Oryza_sativa VLSMCLAFAKEPFKVKSGSCGTVVRNAELKIIDPDTGKSLGRNLPGEICIRGQQIMKGYL
AAF37733.1_4CL2_Lolium_perenne VLAMCLAFAKEPFAVKSGSCGTVVRNAELKIVDPDTGASLGRNLPGEICIRGKQIMKGYL
AAF37734.1_4CL3_Lolium_perenne VLAMCLAFAKEPFKVKSGSCGTVVRNAELKVVDPDTGASLGRNQPGEICVRGKQIMIGYL
acc

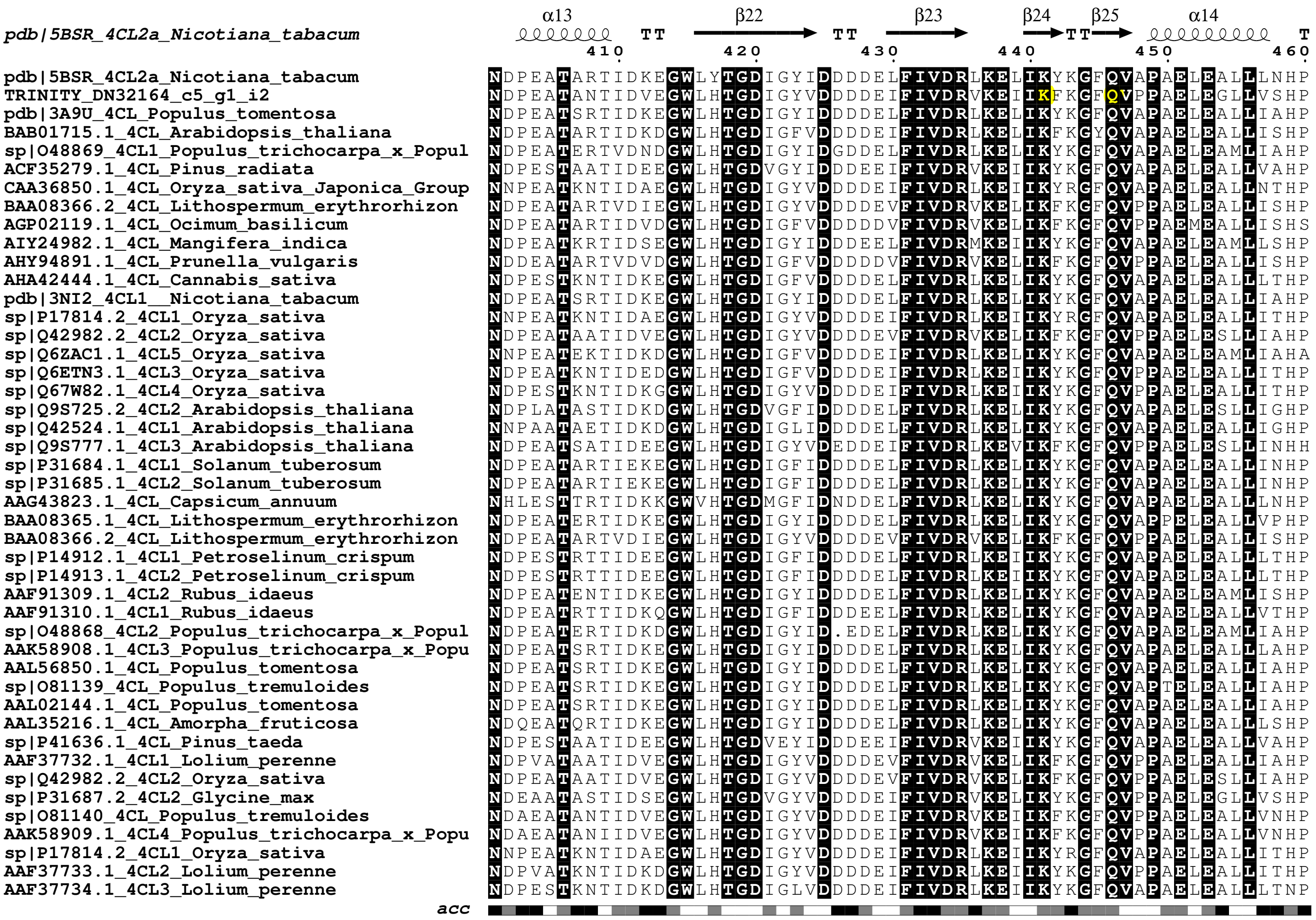
pdb|5BSR_4CL2a_Nicotiana_tabacum
α13 TT β22 TT β23 β24 TT β25 α14 T
410 420 430 440 450 460
pdb|5BSR_4CL2a_Nicotiana_tabacum
TRINITY_DN32164_c5_g1_i2
pdb|3A9U_4CL_Populus_tomentosa
BAB01715.1_4CL_Arabidopsis_thaliana
sp|O48869_4CL1_Populus_trichocarpa_x_Popul
ACF35279.1_4CL_Pinus_radiata
CAA36850.1_4CL_Oryza_sativa_Japonica_Group
BAA08366.2_4CL_Lithospermum_erythrorhizon
AGP02119.1_4CL_Ocimum_basilicum
AIY24982.1_4CL_Mangifera_indica
AHY94891.1_4CL_Prunella_vulgaris
AHA42444.1_4CL_Cannabis_sativa
pdb|3NI2_4CL1__Nicotiana_tabacum
sp|P17814.2_4CL1_Oryza_sativa
sp|Q42982.2_4CL2_Oryza_sativa
sp|Q6ZAC1.1_4CL5_Oryza_sativa
sp|Q6ETN3.1_4CL3_Oryza_sativa
sp|Q67W82.1_4CL4_Oryza_sativa
sp|Q9S725.2_4CL2_Arabidopsis_thaliana
sp|Q42524.1_4CL1_Arabidopsis_thaliana
sp|Q9S777.1_4CL3_Arabidopsis_thaliana
sp|P31684.1_4CL1_Solanum_tuberosum
sp|P31685.1_4CL2_Solanum_tuberosum
AAG43823.1_4CL_Capsicum_annuum
BAA08365.1_4CL_Lithospermum_erythrorhizon
BAA08366.2_4CL_Lithospermum_erythrorhizon
sp|P14912.1_4CL1_Petroselinum_crispum
sp|P14913.1_4CL2_Petroselinum_crispum
AAF91309.1_4CL2_Rubus_idaeus
AAF91310.1_4CL1_Rubus_idaeus
sp|O48868_4CL2_Populus_trichocarpa_x_Popul
AAK58908.1_4CL3_Populus_trichocarpa_x_Popu
AAL56850.1_4CL_Populus_tomentosa
sp|O81139_4CL_Populus_tremuloides
AAL02144.1_4CL_Populus_tomentosa
AAL35216.1_4CL_Amorpha_fruticosa
sp|P41636.1_4CL_Pinus_taeda
AAF37732.1_4CL1_Lolium_perenne
sp|Q42982.2_4CL2_Oryza_sativa
sp|P31687.2_4CL2_Glycine_max
sp|O81140_4CL_Populus_tremuloides
AAK58909.1_4CL4_Populus_trichocarpa_x_Popu
sp|P17814.2_4CL1_Oryza_sativa
AAF37733.1_4CL2_Lolium_perenne
AAF37734.1_4CL3_Lolium_perenne
acc

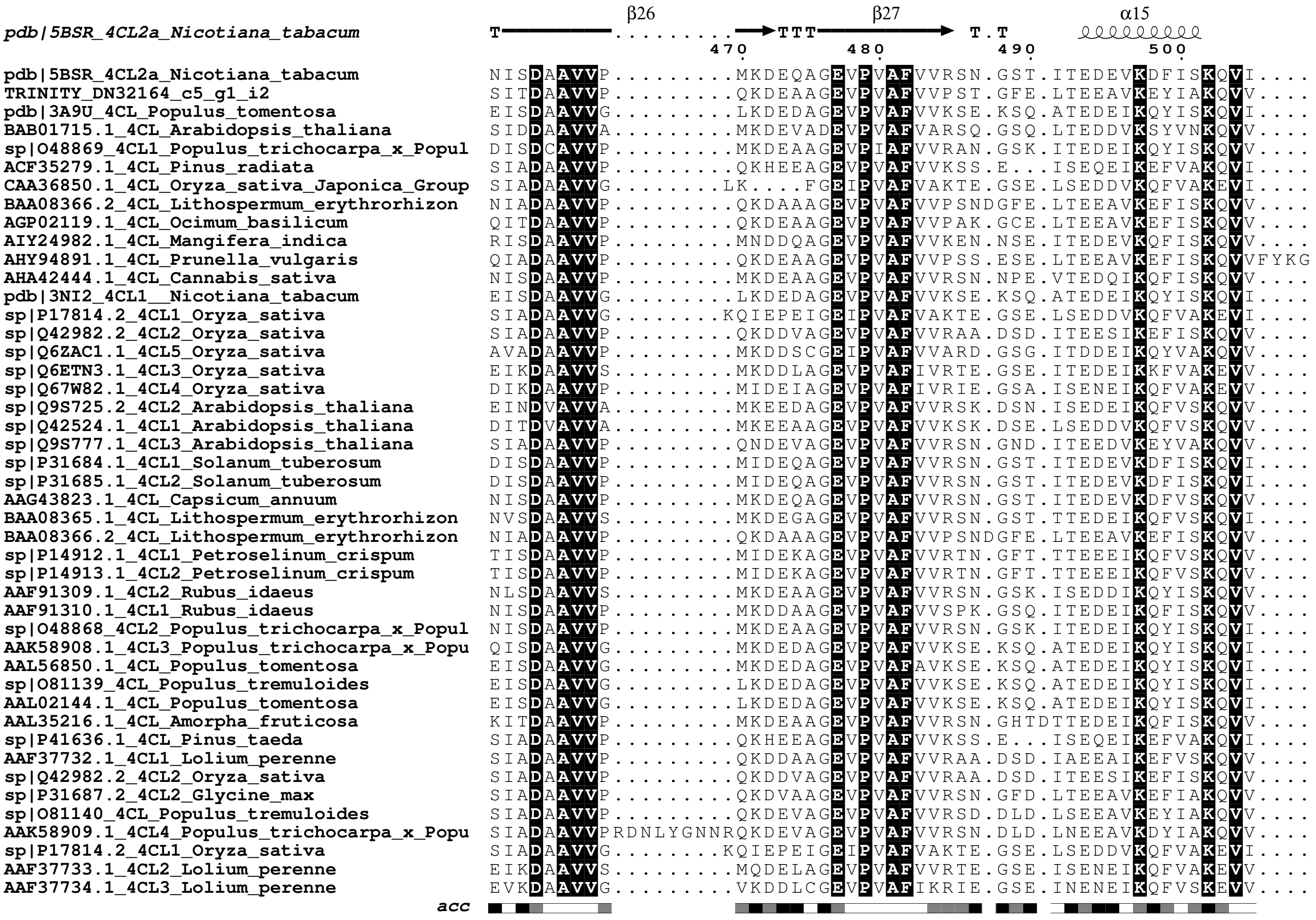
pdb|5BSR_4CL2a_Nicotiana_tabacum
T β26 TTT β27 T.T α15
470 480 490 500
acc

```
pdb|5BSR_4CL2a_Nicotiana_tabacum

pdb|5BSR_4CL2a_Nicotiana_tabacum                ............................................................
TRINITY_DN32164_c5_g1_i2                        ............................................................
pdb|3A9U_4CL_Populus_tomentosa                  ............................................................
BAB01715.1_4CL_Arabidopsis_thaliana             ............................................................
sp|O48869_4CL1_Populus_trichocarpa_x_Popul      ............................................................
ACF35279.1_4CL_Pinus_radiata                    ............................................................
CAA36850.1_4CL_Oryza_sativa_Japonica_Group      ............................................................
BAA08366.2_4CL_Lithospermum_erythrorhizon       ............................................................
AGP02119.1_4CL_Ocimum_basilicum                 ............................................................
AIY24982.1_4CL_Mangifera_indica                 ............................................................
AHY94891.1_4CL_Prunella_vulgaris                MPKGFFVQAIPKSPFGKILGKNFRAQIGSPTFLFIVDRLKELIKYKGYQVAPAELEALLL
AHA42444.1_4CL_Cannabis_sativa                  ............................................................
pdb|3NI2_4CL1__Nicotiana_tabacum                ............................................................
sp|P17814.2_4CL1_Oryza_sativa                   ............................................................
sp|Q42982.2_4CL2_Oryza_sativa                   ............................................................
sp|Q6ZAC1.1_4CL5_Oryza_sativa                   ............................................................
sp|Q6ETN3.1_4CL3_Oryza_sativa                   ............................................................
sp|Q67W82.1_4CL4_Oryza_sativa                   ............................................................
sp|Q9S725.2_4CL2_Arabidopsis_thaliana           ............................................................
sp|Q42524.1_4CL1_Arabidopsis_thaliana           ............................................................
sp|Q9S777.1_4CL3_Arabidopsis_thaliana           ............................................................
sp|P31684.1_4CL1_Solanum_tuberosum              ............................................................
sp|P31685.1_4CL2_Solanum_tuberosum              ............................................................
AAG43823.1_4CL_Capsicum_annuum                  ............................................................
BAA08365.1_4CL_Lithospermum_erythrorhizon       ............................................................
BAA08366.2_4CL_Lithospermum_erythrorhizon       ............................................................
sp|P14912.1_4CL1_Petroselinum_crispum           ............................................................
sp|P14913.1_4CL2_Petroselinum_crispum           ............................................................
AAF91309.1_4CL2_Rubus_idaeus                    ............................................................
AAF91310.1_4CL1_Rubus_idaeus                    ............................................................
sp|O48868_4CL2_Populus_trichocarpa_x_Popul      ............................................................
AAK58908.1_4CL3_Populus_trichocarpa_x_Popu      ............................................................
AAL56850.1_4CL_Populus_tomentosa                ............................................................
sp|O81139_4CL_Populus_tremuloides               ............................................................
AAL02144.1_4CL_Populus_tomentosa                ............................................................
AAL35216.1_4CL_Amorpha_fruticosa                ............................................................
sp|P41636.1_4CL_Pinus_taeda                     ............................................................
AAF37732.1_4CL1_Lolium_perenne                  ............................................................
sp|Q42982.2_4CL2_Oryza_sativa                   ............................................................
sp|P31687.2_4CL2_Glycine_max                    ............................................................
sp|O81140_4CL_Populus_tremuloides               ............................................................
AAK58909.1_4CL4_Populus_trichocarpa_x_Popu      ............................................................
sp|P17814.2_4CL1_Oryza_sativa                   ............................................................
AAF37733.1_4CL2_Lolium_perenne                  ............................................................
AAF37734.1_4CL3_Lolium_perenne                  ............................................................
                                           acc
```

```
                                                                                                η7     β28
pdb|5BSR_4CL2a_Nicotiana_tabacum                                                                  510
                                                                                                    .
pdb|5BSR_4CL2a_Nicotiana_tabacum                ................................................FYKRIKRVFFVD
TRINITY_DN32164_c5_g1_i2                        ................................................FYKKLHKVYFVH
pdb|3A9U_4CL_Populus_tomentosa                  ................................................FYKRIKRVFFIE
BAB01715.1_4CL_Arabidopsis_thaliana             ................................................HYKRIKMVFFIE
sp|O48869_4CL1_Populus_trichocarpa_x_Popul      ................................................FYKRISRVFFTE
ACF35279.1_4CL_Pinus_radiata                    ................................................FYKKIHRVYFVD
CAA36850.1_4CL_Oryza_sativa_Japonica_Group      ................................................YYKKIREVFFVD
BAA08366.2_4CL_Lithospermum_erythrorhizon       ................................................FYKRLHKVYFVH
AGP02119.1_4CL_Ocimum_basilicum                 ................................................FYKRLHKVYFVH
AIY24982.1_4CL_Mangifera_indica                 ................................................FYKRINRVFFID
AHY94891.1_4CL_Prunella_vulgaris                NHPNISDAAVVPMKDEQAGEVPVAFVVKSKGSVITEDDIKAFVSKQVIFYKRIKRVFFVD
AHA42444.1_4CL_Cannabis_sativa                  ................................................FYKRISRVFFID
pdb|3NI2_4CL1__Nicotiana_tabacum                ................................................FYKRIKRVFFIE
sp|P17814.2_4CL1_Oryza_sativa                   ................................................YYKKIREVFFVD
sp|Q42982.2_4CL2_Oryza_sativa                   ................................................FYKRLHKVHFIH
sp|Q6ZAC1.1_4CL5_Oryza_sativa                   ................................................FYKRLHKIFFVD
sp|Q6ETN3.1_4CL3_Oryza_sativa                   ................................................FYKRINKVFFTD
sp|Q67W82.1_4CL4_Oryza_sativa                   ................................................FYKRLNKVFFAD
sp|Q9S725.2_4CL2_Arabidopsis_thaliana           ................................................FYKRINKVFFTD
sp|Q42524.1_4CL1_Arabidopsis_thaliana           ................................................FYKRINKVFFTE
sp|Q9S777.1_4CL3_Arabidopsis_thaliana           ................................................FYKRLHKVFFVA
sp|P31684.1_4CL1_Solanum_tuberosum              ................................................FYKRIKRVFFVE
sp|P31685.1_4CL2_Solanum_tuberosum              ................................................FYKRIKRVFFVE
AAG43823.1_4CL_Capsicum_annuum                  ................................................FYKRIKRVFFVE
BAA08365.1_4CL_Lithospermum_erythrorhizon       ................................................FYKRINRVFGVD
BAA08366.2_4CL_Lithospermum_erythrorhizon       ................................................FYKRLHKVYFVH
sp|P14912.1_4CL1_Petroselinum_crispum           ................................................FYKRIFRVFFVD
sp|P14913.1_4CL2_Petroselinum_crispum           ................................................FYKRIFRVFFVD
AAF91309.1_4CL2_Rubus_idaeus                    ................................................FYKRISKVFFTD
AAF91310.1_4CL1_Rubus_idaeus                    ................................................FYKRIKRVFFIE
sp|O48868_4CL2_Populus_trichocarpa_x_Popul      ................................................FYKRIGRVFFTE
AAK58908.1_4CL3_Populus_trichocarpa_x_Popu      ................................................FYKRIKRVFFIE
AAL56850.1_4CL_Populus_tomentosa                ................................................FYKRIKRVFFIE
sp|O81139_4CL_Populus_tremuloides               ................................................FYKRIKRVFFIE
AAL02144.1_4CL_Populus_tomentosa                ................................................FYKRIKRVFFIE
AAL35216.1_4CL_Amorpha_fruticosa                ................................................FYKRISRVFFID
sp|P41636.1_4CL_Pinus_taeda                     ................................................FYKKIHRVYFVD
AAF37732.1_4CL1_Lolium_perenne                  ................................................FYKRLHKVYFTH
sp|Q42982.2_4CL2_Oryza_sativa                   ................................................FYKRLHKVHFIH
sp|P31687.2_4CL2_Glycine_max                    ................................................FYKRLHKVYFVH
sp|O81140_4CL_Populus_tremuloides               ................................................FYKKLHKVFFVH
AAK58909.1_4CL4_Populus_trichocarpa_x_Popu      ................................................FYKKLHKVFFVH
sp|P17814.2_4CL1_Oryza_sativa                   ................................................YYKKIREVFFVD
AAF37733.1_4CL2_Lolium_perenne                  ................................................FYKRICKVFFAD
AAF37734.1_4CL3_Lolium_perenne                  ................................................FYKRINKVYFTD
                                           acc
```

```
                                                                        α16
pdb|5BSR_4CL2a_Nicotiana_tabacum                  TT   
                                       520          530
                                         .            .
pdb|5BSR_4CL2a_Nicotiana_tabacum         AIPKSPSGKILRKDLRAKLA.....A..................................
TRINITY_DN32164_c5_g1_i2                 AIPKSPSGKILRKDLKAKLA.....S..................................
pdb|3A9U_4CL_Populus_tomentosa           AIPKAPSGKILRKNLKEKL......A..................................
BAB01715.1_4CL_Arabidopsis_thaliana      VIPKAVSGKILRKDLRAKLE.....T..................................
sp|O48869_4CL1_Populus_trichocarpa_x_Popul AIPKAPSGKILRKDLRARLA.....T..................................
ACF35279.1_4CL_Pinus_radiata             AIPKSPSGKILRKDLRSRLA.....AK.................................
CAA36850.1_4CL_Oryza_sativa_Japonica_Group KIPKAPSGKILRKELRKQLQHLQQEA..................................
BAA08366.2_4CL_Lithospermum_erythrorhizon SIPKSPSGKILRKDLRAKLA.....A..................................
AGP02119.1_4CL_Ocimum_basilicum          AIPKSPSGKILRKDLRAKLA.....A..................................
AIY24982.1_4CL_Mangifera_indica          AIPKAPSGKILRKDLRAKLA.....A..................................
AHY94891.1_4CL_Prunella_vulgaris         AVPKSPSGKILRKELRARLAD...GV..................................
AHA42444.1_4CL_Cannabis_sativa           AIPKSPSGKILRKDLRAKLA.....L..................................
pdb|3NI2_4CL1__Nicotiana_tabacum         AIPKAPSGKILRKNLKEKL......A..................................
sp|P17814.2_4CL1_Oryza_sativa            KIPKAPSGKILRKELRKQLQHLQQEA..................................
sp|Q42982.2_4CL2_Oryza_sativa            AIPKSASGKILRRELRAKLA.....A..................................
sp|Q6ZAC1.1_4CL5_Oryza_sativa            AIPKAPSGKILRKDLRAKLA.....A..................................
sp|Q6ETN3.1_4CL3_Oryza_sativa            SIPKNPSGKILRKDLRARLA.....A..................................
sp|Q67W82.1_4CL4_Oryza_sativa            SIPKSPSGKILRKDLRAKLA.....A..................................
sp|Q9S725.2_4CL2_Arabidopsis_thaliana    SIPKAPSGKILRKDLRARLA.....N..................................
sp|Q42524.1_4CL1_Arabidopsis_thaliana    SIPKAPSGKILRKDLRAKLA.....N..................................
sp|Q9S777.1_4CL3_Arabidopsis_thaliana    SIPKSPSGKILRKDLKAKLC........................................
sp|P31684.1_4CL1_Solanum_tuberosum       TVPKSPSGKILRKDLRARLA.....A..................................
sp|P31685.1_4CL2_Solanum_tuberosum       TVPKSPSGKILRKDLRARLA.....A..................................
AAG43823.1_4CL_Capsicum_annuum           TVPKSPSGKILRKDLRARLA.....A..................................
BAA08365.1_4CL_Lithospermum_erythrorhizon SIPKSPSGKIVRKDLRAKLA.....ARFLNGPTTNVVPNGGNVAKDNVPNGVSNGVSKAN
BAA08366.2_4CL_Lithospermum_erythrorhizon SIPKSPSGKILRKDLRAKLA.....A..................................
sp|P14912.1_4CL1_Petroselinum_crispum    AIPKSPSGKILRKDLRARIA.....S..................................
sp|P14913.1_4CL2_Petroselinum_crispum    AIPKSPSGKILRKDLRAKIA.....S..................................
AAF91309.1_4CL2_Rubus_idaeus             KIPKSPSGKILRKDLRAKLA.....A..................................
AAF91310.1_4CL1_Rubus_idaeus             AIPKSPSGKILRKELRAKLA.....A..................................
sp|O48868_4CL2_Populus_trichocarpa_x_Popul AIPKAPSGKILRKDLRARVS.....A..................................
AAK58908.1_4CL3_Populus_trichocarpa_x_Popu AIPKAPSGKILRKNLRETL......P..................................
AAL56850.1_4CL_Populus_tomentosa         AIPKAPSGKILRKNLKEKL......P..................................
sp|O81139_4CL_Populus_tremuloides        AIPKAPSGKILRKNLKEKL......P..................................
AAL02144.1_4CL_Populus_tomentosa         AIPKAPSGKILRKNLKEKL......A..................................
AAL35216.1_4CL_Amorpha_fruticosa         AIPKSPSGKILRKDLRAKLA.....A..................................
sp|P41636.1_4CL_Pinus_taeda              AIPKSPSGKILRKDLRSRLA.....AK.................................
AAF37732.1_4CL1_Lolium_perenne           AIPKSASGKILRKELRAKLA.....A..................................
sp|Q42982.2_4CL2_Oryza_sativa            AIPKSASGKILRRELRAKLA.....A..................................
sp|P31687.2_4CL2_Glycine_max             AIPKSPSGKILRKDLRAKLE.....T..................................
sp|O81140_4CL_Populus_tremuloides        SIPKSASGKILRKDLRAKLA.....T..................................
AAK58909.1_4CL4_Populus_trichocarpa_x_Popu SIPKSASGKILRKDLRAKLA.....T..................................
sp|P17814.2_4CL1_Oryza_sativa            KIPKAPSGKILRKELRKQLQHLQQEA..................................
AAF37733.1_4CL2_Lolium_perenne           SIPKSPSGKILRKDLRAKLA.....A..................................
AAF37734.1_4CL3_Lolium_perenne           SIPKNPSGKILRKDLRARLA.....A..................................
                                     acc

pdb|5BSR_4CL2a_Nicotiana_tabacum
                                                                     540
                                                                       .
pdb|5BSR_4CL2a_Nicotiana_tabacum         ...........................G...LPN..........................
TRINITY_DN32164_c5_g1_i2                 ..........................ASQ..LS...........................
pdb|3A9U_4CL_Populus_tomentosa           ...........................G...I............................
BAB01715.1_4CL_Arabidopsis_thaliana      ...............................MCSK.........................
sp|O48869_4CL1_Populus_trichocarpa_x_Popul ...........................GDFLIKF..........................
ACF35279.1_4CL_Pinus_radiata             ............................................................
CAA36850.1_4CL_Oryza_sativa_Japonica_Group ...............................LTN..........................
BAA08366.2_4CL_Lithospermum_erythrorhizon ..........................AAS..S............................
AGP02119.1_4CL_Ocimum_basilicum          ..........................PSS..TS...........................
AIY24982.1_4CL_Mangifera_indica          ...........................G...LPN..........................
AHY94891.1_4CL_Prunella_vulgaris         ..........................PA................................
AHA42444.1_4CL_Cannabis_sativa           ...........................GN..L............................
pdb|3NI2_4CL1__Nicotiana_tabacum         ...........................G...I............................
sp|P17814.2_4CL1_Oryza_sativa            ............................................................
sp|Q42982.2_4CL2_Oryza_sativa            ..........................C.................................
sp|Q6ZAC1.1_4CL5_Oryza_sativa            ...........................G...IPAC.........................
sp|Q6ETN3.1_4CL3_Oryza_sativa            ...........................G...IP..DAVAAAAA................D
sp|Q67W82.1_4CL4_Oryza_sativa            ...........................G...IPTNDNTQLKS..................
sp|Q9S725.2_4CL2_Arabidopsis_thaliana    ...........................G...LMN..........................
sp|Q42524.1_4CL1_Arabidopsis_thaliana    ...........................G...L............................
sp|Q9S777.1_4CL3_Arabidopsis_thaliana    ............................................................
sp|P31684.1_4CL1_Solanum_tuberosum       ...........................G...ISN..........................
sp|P31685.1_4CL2_Solanum_tuberosum       ...........................G...ISN..........................
AAG43823.1_4CL_Capsicum_annuum           ...........................G...VTN..........................
BAA08365.1_4CL_Lithospermum_erythrorhizon GGVAKEGVANGVPTDGDYGVATKGVANG...ISNGVYKQVSNGVVSNGVANGIVSNGIAN
BAA08366.2_4CL_Lithospermum_erythrorhizon ..........................AAS..S............................
sp|P14912.1_4CL1_Petroselinum_crispum    ..........................GD...LPK..........................
sp|P14913.1_4CL2_Petroselinum_crispum    ..........................GD...LPK..........................
AAF91309.1_4CL2_Rubus_idaeus             ...........................G...LPN..........................
AAF91310.1_4CL1_Rubus_idaeus             ...........................G...FAN..........................
sp|O48868_4CL2_Populus_trichocarpa_x_Popul ...........................GD..LPC..........................
AAK58908.1_4CL3_Populus_trichocarpa_x_Popu ...........................G...I............................
AAL56850.1_4CL_Populus_tomentosa         ...........................G...I............................
sp|O81139_4CL_Populus_tremuloides        ...........................G...I............................
AAL02144.1_4CL_Populus_tomentosa         ...........................G...I............................
AAL35216.1_4CL_Amorpha_fruticosa         ...........................G...VPN..........................
sp|P41636.1_4CL_Pinus_taeda              ............................................................
AAF37732.1_4CL1_Lolium_perenne           ..........................PAT..A............................
sp|Q42982.2_4CL2_Oryza_sativa            ..........................C.................................
sp|P31687.2_4CL2_Glycine_max             ..........................AAT..QTP..........................
sp|O81140_4CL_Populus_tremuloides        ..........................ATT..MS...........................
AAK58909.1_4CL4_Populus_trichocarpa_x_Popu ..........................ATT..MS...........................
sp|P17814.2_4CL1_Oryza_sativa            ............................................................
AAF37733.1_4CL2_Lolium_perenne           ...........................G...IPSSNTTQSKS..................
AAF37734.1_4CL3_Lolium_perenne           ...........................G...IP....TEVAA..................
                                     acc
```

*pdb|5BSR_4CL2a_Nicotiana_tabacum*

```
pdb|5BSR_4CL2a_Nicotiana_tabacum           ...........
TRINITY_DN32164_c5_g1_i2                   ...........
pdb|3A9U_4CL_Populus_tomentosa             ...........
BAB01715.1_4CL_Arabidopsis_thaliana        ...........
sp|O48869_4CL1_Populus_trichocarpa_x_Popul .QHDTYMQKQQ
ACF35279.1_4CL_Pinus_radiata               ...........
CAA36850.1_4CL_Oryza_sativa_Japonica_Group ...........
BAA08366.2_4CL_Lithospermum_erythrorhizon  ...........
AGP02119.1_4CL_Ocimum_basilicum            ...........
AIY24982.1_4CL_Mangifera_indica            ...........
AHY94891.1_4CL_Prunella_vulgaris           ...........
AHA42444.1_4CL_Cannabis_sativa             ...........
pdb|3NI2_4CL1__Nicotiana_tabacum           ...........
sp|P17814.2_4CL1_Oryza_sativa              ...........
sp|Q42982.2_4CL2_Oryza_sativa              ...........
sp|Q6ZAC1.1_4CL5_Oryza_sativa              ...........
sp|Q6ETN3.1_4CL3_Oryza_sativa              APKSS......
sp|Q67W82.1_4CL4_Oryza_sativa              ...........
sp|Q9S725.2_4CL2_Arabidopsis_thaliana      ...........
sp|Q42524.1_4CL1_Arabidopsis_thaliana      ...........
sp|Q9S777.1_4CL3_Arabidopsis_thaliana      ...........
sp|P31684.1_4CL1_Solanum_tuberosum         ...........
sp|P31685.1_4CL2_Solanum_tuberosum         ...........
AAG43823.1_4CL_Capsicum_annuum             ...........
BAA08365.1_4CL_Lithospermum_erythrorhizon  GVHN.......
BAA08366.2_4CL_Lithospermum_erythrorhizon  ...........
sp|P14912.1_4CL1_Petroselinum_crispum      ...........
sp|P14913.1_4CL2_Petroselinum_crispum      ...........
AAF91309.1_4CL2_Rubus_idaeus               ...........
AAF91310.1_4CL1_Rubus_idaeus               ...........
sp|O48868_4CL2_Populus_trichocarpa_x_Popul .TSDS......
AAK58908.1_4CL3_Populus_trichocarpa_x_Popu ...........
AAL56850.1_4CL_Populus_tomentosa           ...........
sp|O81139_4CL_Populus_tremuloides          ...........
AAL02144.1_4CL_Populus_tomentosa           ...........
AAL35216.1_4CL_Amorpha_fruticosa           ...........
sp|P41636.1_4CL_Pinus_taeda                ...........
AAF37732.1_4CL1_Lolium_perenne             ...........
sp|Q42982.2_4CL2_Oryza_sativa              ...........
sp|P31687.2_4CL2_Glycine_max               ...........
sp|O81140_4CL_Populus_tremuloides          ...........
AAK58909.1_4CL4_Populus_trichocarpa_x_Popu ...........
sp|P17814.2_4CL1_Oryza_sativa              ...........
AAF37733.1_4CL2_Lolium_perenne             ...........
AAF37734.1_4CL3_Lolium_perenne             .PRS.......
                                       acc
```

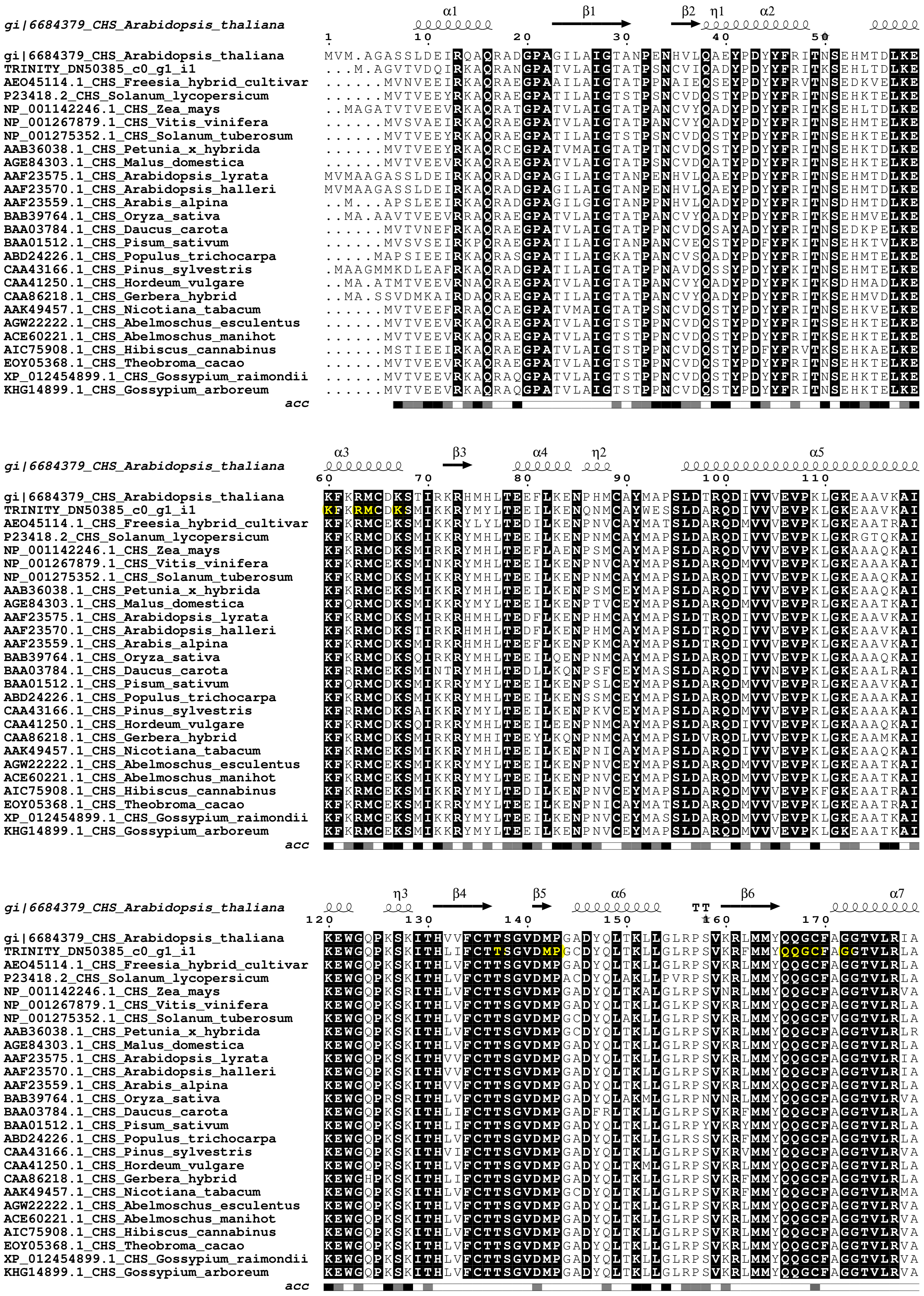
gi|6684379_CHS_Arabidopsis_thaliana
α1 β1 β2 η1 α2
1 10 20 30 40 50
gi|6684379_CHS_Arabidopsis_thaliana MVM.AGASSLDEIRQAQRADGPAGILAIGTANPENHVLQAEYPDYYFRITNSEHMTDLKE
TRINITY_DN50385_c0_g1_i1 ...M.AGVTVDQIRKAQRANGPATVLAIGTATPSNCVIQADYPDYYFRITKSEHLTDLKE
AEO45114.1_CHS_Freesia_hybrid_cultivar ......MVNVEEIRKAQRAEGPAAILAIGTATPPNAIEQSEYPDYYFRVTNSEDKVELKE
P23418.2_CHS_Solanum_lycopersicum ......MVTVEEYRKAQRAEGPATILAIGTSTPSNCVDQSTYPDYYFRITNSEHKTELKE
NP_001142246.1_CHS_Zea_mays ..MAGATVTVEEVRKAQRATGPATVLAIGTATPANCVYQADYPDYYFRITKSEHLTDLKE
NP_001267879.1_CHS_Vitis_vinifera ......MVSVAEIRKAQRAEGPATVLAIGTATPANCVYQADYPDYYFRITNSEHMTELKE
NP_001275352.1_CHS_Solanum_tuberosum ......MVTVEEYRKAQRAEGPATILAIGTSTPSNCVDQSTYPDYYFRITNSEHKTELKE
AAB36038.1_CHS_Petunia_x_hybrida ......MVTVEEYRKAQRCEGPATVMAIGTATPTNCVDQSTYPDYYFRITNSEHKTDLKE
AGE84303.1_CHS_Malus_domestica ......MVTVEEVRKAQRAEGPATVLAIGTATPSNCVDQATYPDYYFRITNSEHKTELKE
AAF23575.1_CHS_Arabidopsis_lyrata MVMAAGASSLDEIRKAQRADGPAGILAIGTANPENHVLQAEYPDYYFRITNSEHMTDLKE
AAF23570.1_CHS_Arabidopsis_halleri MVMAAGASSLDEIRKAQRADGPAGILAIGTANPENHVLQAEYPDYYFRITNSEHMTDLKE
AAF23559.1_CHS_Arabis_alpina ..M...APSLEEIRKAQRADGPAGILGIGTANPPNHVLQADYPDYYFRITNSDHMTDLKE
BAB39764.1_CHS_Oryza_sativa ..MA.AAVTVEEVRRAQRAEGPATVLAIGTATPANCVYQADYPDYYFRITKSEHMVELKE
BAA03784.1_CHS_Daucus_carota ......MVTVNEFRKAQRAEGPATVLAIGTATPPNCVDQSAYADYYFRITNSEDKPELKE
BAA01512.1_CHS_Pisum_sativum ......MVSVSEIRKPQRAEGPATILAIGTANPANCVEQSTYPDFYFKITNSEHKTVLKE
ABD24226.1_CHS_Populus_trichocarpa .....MAPSIEEIRKAQRASGPATILAIGKATPANCVSQADYPDYYFRITNSEHMTELKE
CAA43166.1_CHS_Pinus_sylvestris .MAAGMMKDLEAFRKAQRADGPATILAIGTATPPNAVDQSSYPDYYFKITNSEHMTELKE
CAA41250.1_CHS_Hordeum_vulgare ..MA.ATMTVEEVRNAQRAEGPATVLAIGTATPANCVYQADYPDYYFKITKSDHMADLKE
CAA86218.1_CHS_Gerbera_hybrid ..MA.SSVDMKAIRDAQRAEGPATILAIGTATPANCVYQADYPDYYFRITKSEHMVDLKE
AAK49457.1_CHS_Nicotiana_tabacum ......MVTVEEFRRAQCAEGPATVMAIGTATPSNCVDQSTYPDYYFRITNSEHKVELKE
AGW22222.1_CHS_Abelmoschus_esculentus ......MVTVEEVRKAQRAEGPATVLAIGTSTPPNCVDQSTYPDYYFRITSSEHKTELKE
ACE60221.1_CHS_Abelmoschus_manihot ......MVTVEEVRKAQRAEGPATVLAIGTSTPPNCVDQSTYPDYYFRITKSEHKTELKE
AIC75908.1_CHS_Hibiscus_cannabinus ......MSTIEEIRKAQRAEGPATVLAIGTSTPPNCVDQSTYPDFYFRVTKSEHKAELKE
EOY05368.1_CHS_Theobroma_cacao ......MVTVEEVRKAQRAEGPATVLAIGTSTPPNCVDQSTYPDYYFRITNSEHKTELKE
XP_012454899.1_CHS_Gossypium_raimondii ......MVTVEEVRKAQRAQGPATVLAIGTSTPPNCVDQSTYPDYYFRITNSEHKTELKE
KHG14899.1_CHS_Gossypium_arboreum ......MVTVEEVRKAQRAQGPATVLAIGTSTPPNCVDQSTYPDYYFRITNSEHKTELKE
acc
gi|6684379_CHS_Arabidopsis_thaliana
α3 β3 α4 η2 α5
60 70 80 90 100 110
gi|6684379_CHS_Arabidopsis_thaliana KFKRMCDKSTIRKRHMHLTEEFLKENPHMCAYMAPSLDTRQDIVVVEVPKLGKEAAVKAI
TRINITY_DN50385_c0_g1_i1 KFKRMCDKSMIKKRYLYLTEEILKENQNMCAYWESSLDARQDIVVVEVPKLGKEAAVKAI
AEO45114.1_CHS_Freesia_hybrid_cultivar KFKRMCEKSMIKKRYLYLTEDILKENPNVCAYMATSLDARQDMVVVEVPKLGKEAATRAI
P23418.2_CHS_Solanum_lycopersicum KFKRMCEKSMIKKRYMHLTEEILKENPNMCAYMAPSLDARQDIVVVEVPKLGKRGTQKAI
NP_001142246.1_CHS_Zea_mays KFKRMCDKSMIRKRYMHLTEEFLAENPSMCAYMAPSLDARQDVVVVEVPKLGKAAAQKAI
NP_001267879.1_CHS_Vitis_vinifera KFKRMCEKSMINKRYMHLTEEILKENPNVCAYMAPSLDARQDMVVVEVPKLGKEAAAKAI
NP_001275352.1_CHS_Solanum_tuberosum KFKRMCDKSMIKKRYMHLTEEILKENPNMCAYMAPSLDARQDIVVVEVPKLGKEAAQKAI
AAB36038.1_CHS_Petunia_x_hybrida KFKRMCEKSMIKKRYMHLTEEILKENPSMCEYMAPSLDARQDIVVVEVPKLGKEAAQKAI
AGE84303.1_CHS_Malus_domestica KFQRMCDKSMIKKRYMYLTEEILKENPTVCEYMAPSLDARQDMVVVEVPRLGKEAATKAI
AAF23575.1_CHS_Arabidopsis_lyrata KFKRMCDKSMIRKRHMHLTEDFLKENPHMCAYMAPSLDTRQDIVVVEVPKLGKEAAVKAI
AAF23570.1_CHS_Arabidopsis_halleri KFKRMCDKSTIRKRHMHLTEDFLKENPHMCAYMAPSLDTRQDIVVVEVPKLGKEAAVKAI
AAF23559.1_CHS_Arabis_alpina KFKRMCDKSMIRKRHMHLTEEFLKENPKMCAYMAPSLDTRQDIVVVEVPKLGKEAAVKAI
BAB39764.1_CHS_Oryza_sativa KFKRMCDKSQIRKRYMHLTEEILQENPNMCAYMAPSLDARQDIVVVEVPKLGKAAAQKAI
BAA03784.1_CHS_Daucus_carota KFRRMCEKSMINTRYMHLTEDLLKQNPSFCEYMASSLDARQDIVVNEVPKLGKEAALRAI
BAA01512.1_CHS_Pisum_sativum KFQRMCDKSMIKKRYMYLTEEILKENPSLCEYMAPSLDARQDMVVVEVPRLGKEAAVKAI
ABD24226.1_CHS_Populus_trichocarpa KFKRMCDKSMIKKRYMHLTEEILKENSSMCEYMAPSLDARQDMVVVEVPKLGKEAAAKAI
CAA43166.1_CHS_Pinus_sylvestris KFRRMCDKSAIKKRYMYLTEEILKENPKVCEYMAPSLDARQDMVVVEVPRLGKEAAAKAI
CAA41250.1_CHS_Hordeum_vulgare KFKRMCDKSQIRKRYMHLTEEILEENPNMCAYMAPSLDARQDIVVVEVPKLGKAAAQKAI
CAA86218.1_CHS_Gerbera_hybrid KFKRMCDKSMIRKRYMHITEEYLKQNPNMCAYMAPSLDVRQDIVVVEVPKLGKEAAMKAI
AAK49457.1_CHS_Nicotiana_tabacum KFKRMCEKSMIKKRYMHLTEEILKENPNICAYMAPSLDARQDIVVVEVPKLGKEAAQKAI
AGW22222.1_CHS_Abelmoschus_esculentus KFKRMCEKSMIKKRYMYLTEEILKENPNVCEYMAPSLDARQDMVVVEVPKLGKEAATKAI
ACE60221.1_CHS_Abelmoschus_manihot KFKRMCEKSMIKKRYMYLTEEILKENPNVCEYMAPSLDARQDMVVVEVPKLGKEAATKAI
AIC75908.1_CHS_Hibiscus_cannabinus KFKRMCEKSMIKKRYMYLTEDILKENPNVCEYMAPSLDARQDMVVVEVPKFGKEAATRAI
EOY05368.1_CHS_Theobroma_cacao KFKRMCEKSMIKRRYMYLTEEILKENPNICAYMATSLDARQDMVVVEVPKLGKEAATRAI
XP_012454899.1_CHS_Gossypium_raimondii KFKRMCEKSMIKKRYMYLTEEILKENPNVCEYMASSLDARQDMVVVEVPKLGKEAATKAI
KHG14899.1_CHS_Gossypium_arboreum KFKRMCEKSMIKKRYMYLTEEILKENPNVCEYMAPSLDARQDMVVVEVPKLGKEAATKAI
acc
gi|6684379_CHS_Arabidopsis_thaliana
η3 β4 β5 α6 TT β6 α7
120 130 140 150 160 170
gi|6684379_CHS_Arabidopsis_thaliana KEWGQPKSKITHVVFCTTSGVDMPGADYQLTKLLGLRPSVKRLMMYQQGCFAGGTVLRIA
TRINITY_DN50385_c0_g1_i1 KEWGQPKSKITHLIFCTTSGVDMPGCDYQLTKLLGLRPSVKRFMMYQQGCFAGGTVLRLA
AEO45114.1_CHS_Freesia_hybrid_cultivar KEWGQPKSKITHLVFCTTSGVDMPGADYQLTKLLGLRPSVKRLMMYQQGCFAGGTVLRLA
P23418.2_CHS_Solanum_lycopersicum KEWGQPKSKITHLVFCTTSGVDMPACDYQLAKLLPVRPSVKRLMMYQQGCFAGGTVLRLA
NP_001142246.1_CHS_Zea_mays KEWGQPKSRITHLVFCTTSGVDMPGADYQLTKALGLRPSVNRLMMYQQGCFAGGTVLRVA
NP_001267879.1_CHS_Vitis_vinifera KEWGQPKSKITHLVFCTTSGVDMPGADYQLTKLLGLKPSVKRLMMYQQGCFAGGTVLRLA
NP_001275352.1_CHS_Solanum_tuberosum KEWGQPKSKITHLVFCTTSGVDMPGCDYQLAKLLGLRPSVKRLMMYQQGCFVGGTVLRLA
AAB36038.1_CHS_Petunia_x_hybrida KEWGQPKSKITHLVFCTTSGVDMPGCDYQLTKLLGLRPSVKRLMMYQQGCFAGGTVLRLR
AGE84303.1_CHS_Malus_domestica KEWGQPKSKITHLVFCTTSGVDMPGADYQLTKLLGLRPSVKRLMMYQQGCFAGGTVLRLA
AAF23575.1_CHS_Arabidopsis_lyrata KEWGQPKSKITHVVFCTTSGVDMPGADYQLTKLLGLRPSVKRLMMYQQGCFAGGTVLRIA
AAF23570.1_CHS_Arabidopsis_halleri KEWGQPKSKITHVVFCTTSGVDMPGADYQLTKLLGLRPSVKRLMMYQQGCFAGGTVLRIA
AAF23559.1_CHS_Arabis_alpina KEWGQPKSKITHVVFCTTSGVDMPGADYQLTKLLGLRPSVKRLMMXQQGCFAGGTVLRLA
BAB39764.1_CHS_Oryza_sativa KEWGQPRSRITHLVFCTTSGVDMPGADYQLAKMLGLRPNVNRLMMYQQGCFAGGTVLRVA
BAA03784.1_CHS_Daucus_carota KEWGQPKSKITHLIFCTTSGVDMPGADFRLTKLLGLRPSVKRFMMYQQGCFAGGTVLRLA
BAA01512.1_CHS_Pisum_sativum KEWGQPKSKITHLIFCTTSGVDMPGADYQLTKLLGLRPYVKRYMMYQQGCFAGGTVLRLA
ABD24226.1_CHS_Populus_trichocarpa KEWGQPKSKITHLVFCTTSGVDMPGADYQLTKLLGLRSSVKRFMMYQQGCFAGGTVLRLA
CAA43166.1_CHS_Pinus_sylvestris KEWGQPKSKITHVIFCTTSGVDMPGADYQLTKLLGLRPSVKRVMMYQQGCFAGGTVLRVA
CAA41250.1_CHS_Hordeum_vulgare KEWGQPRSKITHLVFCTTSGVDMPGADYQLTKMLGLRPSVKRLMMYQQGCFAGGTVLRLA
CAA86218.1_CHS_Gerbera_hybrid KEWGHPKSKITHLIFCTTSGVDMPGADYQLTKLLGLRPSVKRFMMYQQGCFAGGTVLRLA
AAK49457.1_CHS_Nicotiana_tabacum KEWGQPKSKITHLVFCTTSGVDMPGADYQLTKLLGLRPSVKRFMMYQQGCFAGGTVLRMA
AGW22222.1_CHS_Abelmoschus_esculentus KEWGQPKSKITHLVFCTTSGVDMPGADYQLTKLLGLRPSVKRLMMYQQGCFAGGTVLRVA
ACE60221.1_CHS_Abelmoschus_manihot KEWGQPKSKITHLVFCTTSGVDMPGADYQLTKLLGLRPSVKRLMMYQQGCFAGGTVLRVA
AIC75908.1_CHS_Hibiscus_cannabinus KEWGQPKSKITHLVFCTTSGVDMPGADYQLTKLLGLRPSVKRLMMYQQGCFAGGTVLRVA
EOY05368.1_CHS_Theobroma_cacao KEWGQPKSKITHLVFCTTSGVDMPGADYQLTKLLGLRPSVKRLMMYQQGCFAGGTVLRVA
XP_012454899.1_CHS_Gossypium_raimondii KEWGQPKSKITHLVFCTTSGVDMPGADYQLTKLLGLRPSVKRLMMYQQGCFAGGTVLRVA
KHG14899.1_CHS_Gossypium_arboreum KEWGQPKSKITHLVFCTTSGVDMPGADYQLTKLLGLRPSVKRLMMYQQGCFAGGTVLRVA
acc

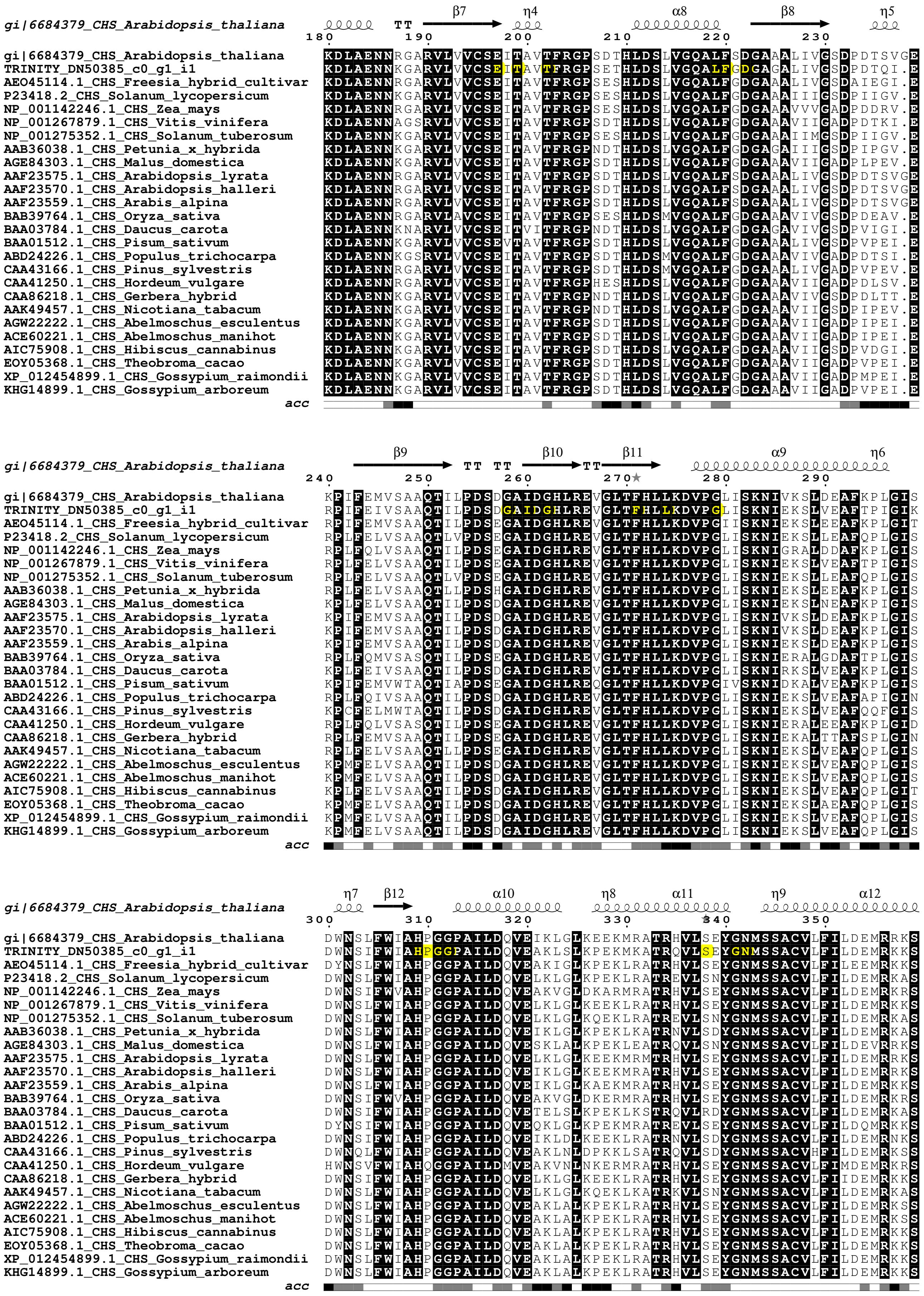
gi|6684379_CHS_Arabidopsis_thaliana
β7 η4 α8 β8 η5
β9 β10 β11 α9 η6
η7 β12 α10 η8 α11 η9 α12
gi|6684379_CHS_Arabidopsis_thaliana
TRINITY_DN50385_c0_g1_i1
AEO45114.1_CHS_Freesia_hybrid_cultivar
P23418.2_CHS_Solanum_lycopersicum
NP_001142246.1_CHS_Zea_mays
NP_001267879.1_CHS_Vitis_vinifera
NP_001275352.1_CHS_Solanum_tuberosum
AAB36038.1_CHS_Petunia_x_hybrida
AGE84303.1_CHS_Malus_domestica
AAF23575.1_CHS_Arabidopsis_lyrata
AAF23570.1_CHS_Arabidopsis_halleri
AAF23559.1_CHS_Arabis_alpina
BAB39764.1_CHS_Oryza_sativa
BAA03784.1_CHS_Daucus_carota
BAA01512.1_CHS_Pisum_sativum
ABD24226.1_CHS_Populus_trichocarpa
CAA43166.1_CHS_Pinus_sylvestris
CAA41250.1_CHS_Hordeum_vulgare
CAA86218.1_CHS_Gerbera_hybrid
AAK49457.1_CHS_Nicotiana_tabacum
AGW22222.1_CHS_Abelmoschus_esculentus
ACE60221.1_CHS_Abelmoschus_manihot
AIC75908.1_CHS_Hibiscus_cannabinus
EOY05368.1_CHS_Theobroma_cacao
XP_012454899.1_CHS_Gossypium_raimondii
KHG14899.1_CHS_Gossypium_arboreum
acc

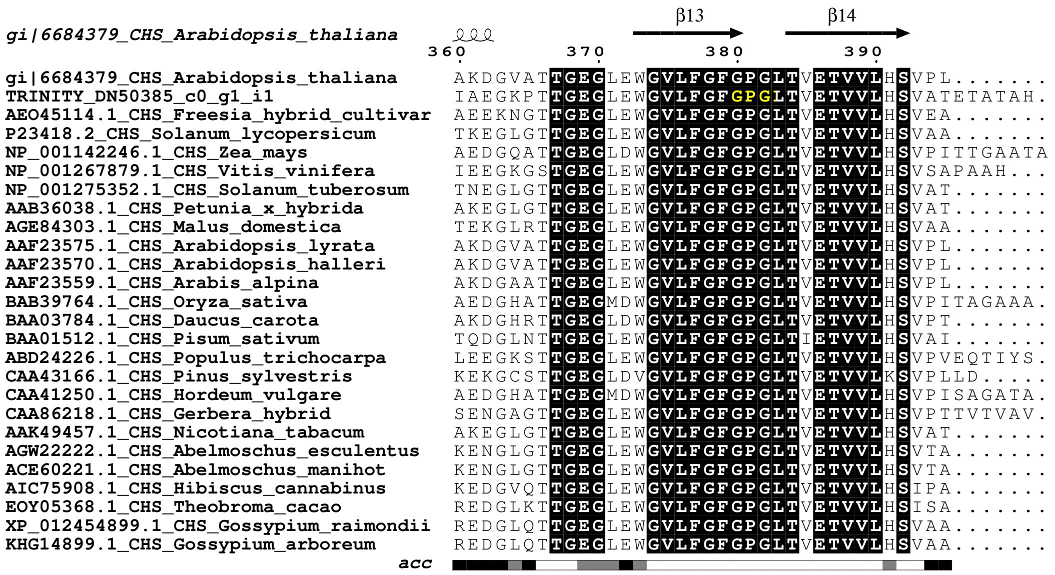
gi|6684379_CHS_Arabidopsis_thaliana
β13
β14
360
370
380
390
gi|6684379_CHS_Arabidopsis_thaliana AKDGVATTGEGLEWGVLFGFGPGLTVETVVLHSVPL.......
TRINITY_DN50385_c0_g1_i1 IAEGKPTTGEGLEWGVLFGFGPGLTVETVVLHSVATETATAH.
AEO45114.1_CHS_Freesia_hybrid_cultivar AEEKNGTTGEGLEWGVLFGFGPGLTVETVVLHSVEA.......
P23418.2_CHS_Solanum_lycopersicum TKEGLGTTGEGLEWGVLFGFGPGLTVETVVLHSVAA.......
NP_001142246.1_CHS_Zea_mays AEDGQATTGEGLDWGVLFGFGPGLTVETVVLHSVPITTGAATA
NP_001267879.1_CHS_Vitis_vinifera IEEGKGSTGEGLEWGVLFGFGPGLTVETVVLHSVSAPAAH...
NP_001275352.1_CHS_Solanum_tuberosum TNEGLGTTGEGLEWGVLFGFGPGLTVETVVLHSVAT.......
AAB36038.1_CHS_Petunia_x_hybrida AKEGLGTTGEGLEWGVLFGFGPGLTVETVVLHSVAT.......
AGE84303.1_CHS_Malus_domestica TEKGLRTTGEGLEWGVLFGFGPGLTVETVVLHSVAA.......
AAF23575.1_CHS_Arabidopsis_lyrata AKDGVATTGEGLEWGVLFGFGPGLTVETVVLHSVPL.......
AAF23570.1_CHS_Arabidopsis_halleri AKDGVATTGEGLEWGVLFGFGPGLTVETVVLHSVPL.......
AAF23559.1_CHS_Arabis_alpina AKDGAATTGEGLEWGVLFGFGPGLTVETVVLHSVPL.......
BAB39764.1_CHS_Oryza_sativa AEDGHATTGEGMDWGVLFGFGPGLTVETVVLHSVPITAGAAA.
BAA03784.1_CHS_Daucus_carota AKDGHRTTGEGLDWGVLFGFGPGLTVETVVLHSVPT.......
BAA01512.1_CHS_Pisum_sativum TQDGLNTTGEGLEWGVLFGFGPGLTIETVVLHSVAI.......
ABD24226.1_CHS_Populus_trichocarpa LEEGKSTTGEGLEWGVLFGFGPGLTVETVVLHSVPVEQTIYS.
CAA43166.1_CHS_Pinus_sylvestris KEKGCSTTGEGLDVGVLFGFGPGLTVETVVLKSVPLLD.....
CAA41250.1_CHS_Hordeum_vulgare AEDGHATTGEGMDWGVLFGFGPGLTVETVVLHSVPISAGATA.
CAA86218.1_CHS_Gerbera_hybrid SENGAGTTGEGLEWGVLFGFGPGLTVETVVLHSVPTTVTVAV.
AAK49457.1_CHS_Nicotiana_tabacum AKEGLGTTGEGLEWGVLFGFGPGLTVETVVLHSVAT.......
AGW22222.1_CHS_Abelmoschus_esculentus KENGLGTTGEGLEWGVLFGFGPGLTVETVVLHSVTA.......
ACE60221.1_CHS_Abelmoschus_manihot KENGLGTTGEGLEWGVLFGFGPGLTVETVVLHSVTA.......
AIC75908.1_CHS_Hibiscus_cannabinus KEDGVQTTGEGLEWGVLFGFGPGLTVETVVLHSIPA.......
EOY05368.1_CHS_Theobroma_cacao REDGLKTTGEGLEWGVLFGFGPGLTVETVVLHSISA.......
XP_012454899.1_CHS_Gossypium_raimondii REDGLQTTGEGLEWGVLFGFGPGLTVETVVLHSVAA.......
KHG14899.1_CHS_Gossypium_arboreum REDGLQTTGEGLEWGVLFGFGPGLTVETVVLHSVAA.......
acc

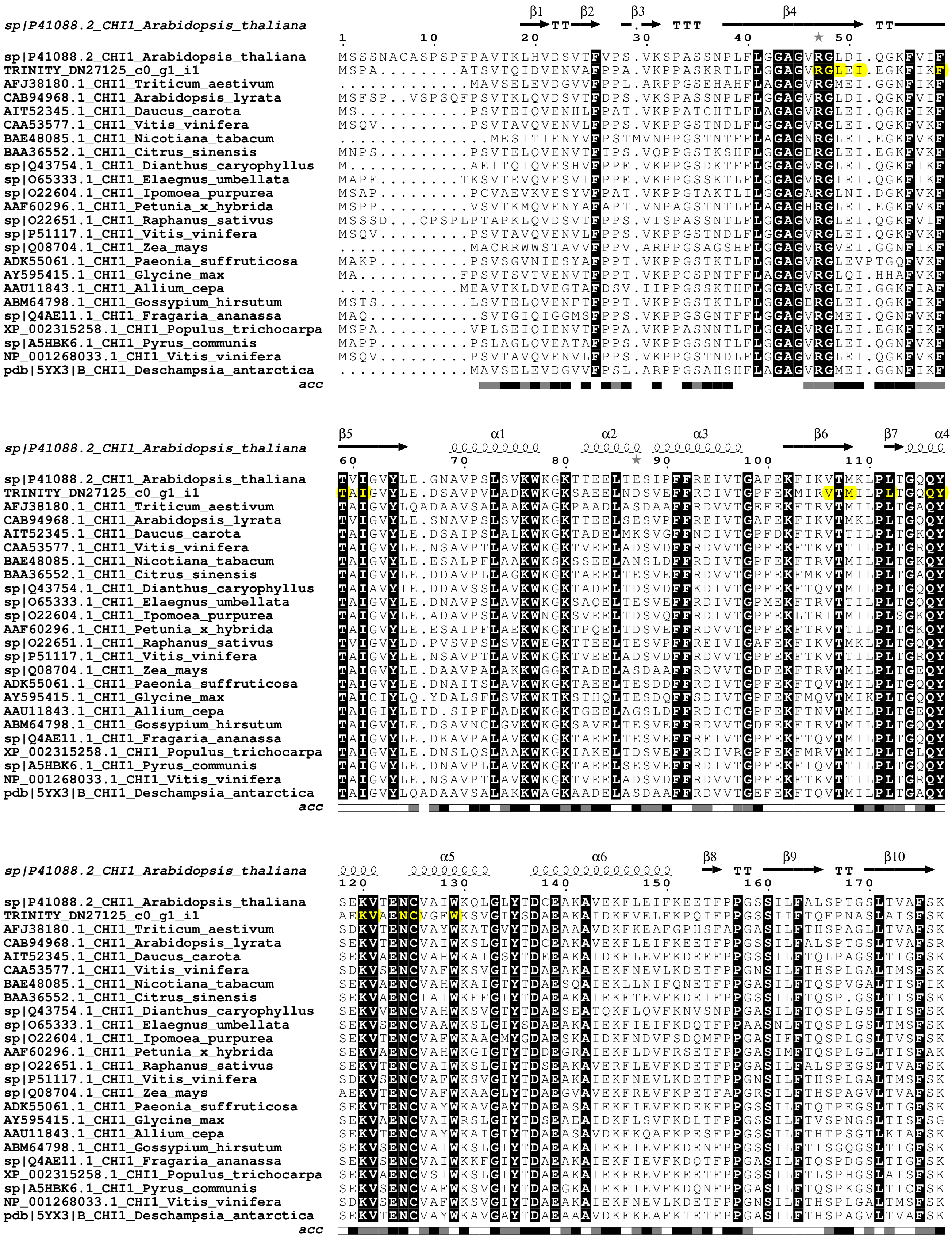
sp|P41088.2_CHI1_Arabidopsis_thaliana
β1 β2 β3 β4 β5 β6 β7 β8 β9 β10
α1 α2 α3 α4 α5 α6
sp|P41088.2_CHI1_Arabidopsis_thaliana
TRINITY_DN27125_c0_g1_i1
AFJ38180.1_CHI1_Triticum_aestivum
CAB94968.1_CHI1_Arabidopsis_lyrata
AIT52345.1_CHI1_Daucus_carota
CAA53577.1_CHI1_Vitis_vinifera
BAE48085.1_CHI1_Nicotiana_tabacum
BAA36552.1_CHI1_Citrus_sinensis
sp|Q43754.1_CHI1_Dianthus_caryophyllus
sp|O65333.1_CHI1_Elaegnus_umbellata
sp|O22604.1_CHI1_Ipomoea_purpurea
AAF60296.1_CHI1_Petunia_x_hybrida
sp|O22651.1_CHI1_Raphanus_sativus
sp|P51117.1_CHI1_Vitis_vinifera
sp|Q08704.1_CHI1_Zea_mays
ADK55061.1_CHI1_Paeonia_suffruticosa
AY595415.1_CHI1_Glycine_max
AAU11843.1_CHI1_Allium_cepa
ABM64798.1_CHI1_Gossypium_hirsutum
sp|Q4AE11.1_CHI1_Fragaria_ananassa
XP_002315258.1_CHI1_Populus_trichocarpa
sp|A5HBK6.1_CHI1_Pyrus_communis
NP_001268033.1_CHI1_Vitis_vinifera
pdb|5YX3|B_CHI1_Deschampsia_antarctica
acc

```
                                             β11      α7                       α8                   α9
sp|P41088.2_CHI1_Arabidopsis_thaliana
                                          180       190       200       210       220
sp|P41088.2_CHI1_Arabidopsis_thaliana     DDSIPETGIAVIENKLLAEAVLESIIGKNGVSPGTRLSVAERLSQLMMKNKD........
TRINITY_DN27125_c0_g1_i1                  DGCIPDAENCLIENKLLSEAVLESIIGKHGVSPEARKCMAVRLVELFEKDAQ.......K
AFJ38180.1_CHI1_Triticum_aestivum         DSSVPESGGVAIDNARLCEAVLESIIGEHGVSPAAKLSLATRVAELL.EGAA........
CAB94968.1_CHI1_Arabidopsis_lyrata        DDSIPETGIAVIENKLLAEAVLESIIGKNGVSPGTRLSIAERLAKLMTKNKV........
AIT52345.1_CHI1_Daucus_carota             ETSIPKDGKAVIHNKQLAEAVLDSIIGEHGVSPAAKHSLAVRISDLLKHHDH........
CAA53577.1_CHI1_Vitis_vinifera            DGSLPEVGNAVIENKLLTEAVLESIIGKHGVSPEAKKSLAARLSELF..CKE........
BAE48085.1_CHI1_Nicotiana_tabacum         DDSITGTGNAVIENKQLSEAVLESIIGKHGVSPAAKCSIAERVSGLFKKSYA........
BAA36552.1_CHI1_Citrus_sinensis           DGSIPKDGVAVIESNLLSEAVLESMIGKNGVSPAAKKSLAERLSALLNVTSD........
sp|Q43754.1_CHI1_Dianthus_caryophyllus    DGSLPENGKEVIVNKQLSEAVLQSIIGKHGVSPEAKKSLSARLSDLINQHEN........
sp|O65333.1_CHI1_Elaegnus_umbellata       DGSIPEVGNAVLENKLLSEAVLESIIGKHGVSPEAKQNLATRLVQLLNENSTTDLNESEN
sp|O22604.1_CHI1_Ipomoea_purpurea         DGSMPEIASAVIENKPLSEAVLESIIGSKGVSPEAKQSLAVRLSEMFINGGD........
AAF60296.1_CHI1_Petunia_x_hybrida         DDSVTGTANAVIENKQLSEAVLESIIGKHGVSPAAKCNVAERVAELLKKSYA........
sp|O22651.1_CHI1_Raphanus_sativus         DDSIPETGKAVIENKLLAEAVLESIIGKNRVSPGARLRVAERLAQLMKENKV........
sp|P51117.1_CHI1_Vitis_vinifera           DGSLPEVGNAVIENKLLTEAVLESIIGKHGVSPEAKKSLAARLSELF..CKE........
sp|Q08704.1_CHI1_Zea_mays                 DSSVPAAGGVAIENKRLCEAVLESIIGERGVSPAAKLSLAARVSELLAKETA........
ADK55061.1_CHI1_Paeonia_suffruticosa      DGVLPEVGNAVVENKQLSEAVLESIIGKHGVSPEAKQSLAARISELLK............
AY595415.1_CHI1_Glycine_max               DETIPEVTSAVIENKLLSEAVLESMIGKNGVSPAAKQSLASRLSHLFKEPGVC.......
AAU11843.1_CHI1_Allium_cepa               DGLVPKDEGVLIENKALTQAVLESIIGEHGVSPAAKLSIASRLSEIMNKVGN........
ABM64798.1_CHI1_Gossypium_hirsutum        DSSVPEGGKVVIENKLLANSVLESVIGKNGVSPAAKESLASRLSPLFNDCG.........
sp|Q4AE11.1_CHI1_Fragaria_ananassa        DGCIPEVGNAVIENKLLSESVLESIIGKPGVSPEARKSVATRLSELLKESDHC.......
XP_002315258.1_CHI1_Populus_trichocarpa   DGSVPEIENAVIENKLLSEAVLESMIGKHGVSPAAKQSLATTLSELLKESNE........
sp|A5HBK6.1_CHI1_Pyrus_communis           DASVPEAANAVIENKLLSEAVLESILGKHGVSPAAKRSLAARLSELLNGCK.........
NP_001268033.1_CHI1_Vitis_vinifera        DGSLPEVGNAVIENKLLTEAVLESIIGKHGVSPEAKKSLAARLSELF..CKE........
pdb|5YX3|B_CHI1_Deschampsia_antarctica    NSSVPESGGAAIENRPLCEAVLESIIGEHGVSPAAKLSLATRVAELL.KEAA........
                                    acc
```

```
sp|P41088.2_CHI1_Arabidopsis_thaliana     ....
                                            230       240
sp|P41088.2_CHI1_Arabidopsis_thaliana     ....EKEVSDHSVEE....KLAKEN...................
TRINITY_DN27125_c0_g1_i1                  EELIDKSPQNVGMVE....KPKETADFIQI..T..PQKNVN...
AFJ38180.1_CHI1_Triticum_aestivum         ......LAGGEPAAE.......................AVSVSV
CAB94968.1_CHI1_Arabidopsis_lyrata        ....EEDASNHSIDE....KLAKEN...................
AIT52345.1_CHI1_Daucus_carota             .........EVTGNG....KA.EVGKMASKDTE..TETPVQV..
CAA53577.1_CHI1_Vitis_vinifera            ...........AGDE....KI.EAEKVA..........PVAC..
BAE48085.1_CHI1_Nicotiana_tabacum         .........DASVFE....KPGIEKSSDPVIEEKPTIPEIGV..
BAA36552.1_CHI1_Citrus_sinensis           ......................................KMK...
sp|Q43754.1_CHI1_Dianthus_caryophyllus    ...........PKKE.............................
sp|O65333.1_CHI1_Elaegnus_umbellata       EKL...NSNEVSKEE....KPLQVEKSAF........KEVEV..
sp|O22604.1_CHI1_Ipomoea_purpurea         ...........AISG....KVGVENDVIP........QTVVVSK
AAF60296.1_CHI1_Petunia_x_hybrida         ........EEASVFG....KPETEKS.........TIPVIGV..
sp|O22651.1_CHI1_Raphanus_sativus         ....EEDATKTDQEEANDLSLAKEN...................
sp|P51117.1_CHI1_Vitis_vinifera           ...........AGDE....KI.EAEKVA..........PVAC..
sp|Q08704.1_CHI1_Zea_mays                 ......AAADAPQAE.......................PVSITA
ADK55061.1_CHI1_Paeonia_suffruticosa      ............................................
AY595415.1_CHI1_Glycine_max               ...................DPQSHK...................
AAU11843.1_CHI1_Allium_cepa               ......VEEKLPVLS.............................
ABM64798.1_CHI1_Gossypium_hirsutum        ...........ADSE....KPQS.....................
sp|Q4AE11.1_CHI1_Fragaria_ananassa        ..........VAGNG....KVDECTKEA..........EVKA..
XP_002315258.1_CHI1_Populus_trichocarpa   ...........NGN..............................
sp|A5HBK6.1_CHI1_Pyrus_communis           ............................................
NP_001268033.1_CHI1_Vitis_vinifera        ...........AGDE....KI.EAEKVA..........PVAC..
pdb|5YX3|B_CHI1_Deschampsia_antarctica    ......SVDEPAAAE.......................PVSVSA
                                    acc
```

```
                                                                      η1        η2          β1
NP_190692.1_F3H_Arabidopsis_thaliana
                                     1        10        20        30        40
NP_190692.1_F3H_Arabidopsis_thaliana   MA..........PGTL....TELAGESKLNSKFVRDEDERPKVAYNVFSDEIPVISL..A
TRINITY_DN33424_c3_g3_i1               MAP.........PSTL....TALSEEKTLQASFVRDEDERPKVAYNEFSNEIPVISL..A
AAU04792.1_F3H_Fragaria_x_ananassa     MAP........TPTTL....TAIAGEKTLQQSFVRDEDERPKVAYNQFSNDIPIISL..S
NP_001236797.1_F3H_Glycine_max         MAP........TAKTL....TYLAQEKTLESSFVRDEEERPKVAYNEFSDEIPVISL..A
ACR15123.1_F3H_Medicago_truncatula     MAP.........AQTL....TYLAQEKTLESSFVREEDERPKVAYNNFSNEIPIISL..D
AQX36284.1_F3H_Prunus_persica          MA.........PATTL....TSIAGEKTLQQKFVRDEDERPKVAYNNFSNEIPIISL..A
ABR13013.1_F3H_Triticum_aestivum       MAPV.......SNET..FLPTAAWGEATLRPSFVRDEDERPKVAHDRLSDAVPVISL..D
NP_001130275.1_F3H_Zea_mays            MAPV.......SISAVPFLPTAAEGETKLRASFVREEDERPKVPHDRFSDEVPVVSL..E
ACL54955.1_F3H_Actinidia_chinensis     MAP.........TTTL....TALAEEKTLQSKFVRDEDERPKVAYNVFSSEIPVISL..A
AFN70721.1_F3H_Nekemias_grossedentata  MA..........PTTL....TALAGEKTLQSSFVRDEDERPKVAYNEFSNEIPIISL..E
ACM62745.1_F3H_Garcinia_mangostana     MAP........TPTTL....TALEGETILRASFVRDEDERPKVAYNQFSNDVPVISL..E
AEO36935.1_F3H_Canarium_album          MAP.........PATL....TTLAQQKTLEASFVRDEDERPKVAYNQFSNEIPVISL..A
AEG64806.1_F3H_Carthamus_tinctorius    MAP.........ISSL.....KWDQGSAVENKYVRDEDERPKVPYDQFSNDIPVISL..E
AFC37250.1_F3H_Camellia_chekiangoleosa MAP.........TTTL....TALAQEKSLQQKFVRDEDERPKAAYNVFSNEIPVISL..A
ACB56921.1_F3H_Pilosella_officinarum   MAP..........ASM.....KWDENSLLEKGFLRDEDERPKVPYNQFSNEIPVISL..E
AAB97310.1_F3H_Chrysanthemum_x_morifolium ......................MDDNSLHENRFVRDEDERPKVPYNKFSSEIPVISL..K
AAC15414.1_F3H_Nicotiana_tabacum       ......................MDDNSLHENRFVRDEDERPKVPYNKFSSEIPVISL..K
ABL86673.1_F3H_Gossypium_barbadense    MA..........PSTL....TALAEEKTLQASFVRDEDERPKVAYNQFSNDIPVISL..A
ADC96713.1_F3H_Gossypium_hirsutum      MA..........PSTL....TALAEEKTLQASFVRDEDERPKVAYNQFSNDIPVISL..A
XP_007046698.1_F3H_Theobroma_cacao     MA..........PSTL....TALAEEKTLQASFVRDEDERPKVAYNQFSNEIPVISL..A
ABO48521.1_F3H_Dimocarpus_longan       MA..........PATL....TALAQEKTLNASFVRDEDERPKVAYNEFSKEIPVISL..A
ACQ99190.1_F3H_Fagopyrum_tataricum     MAPV......AKPSTL....TALAEEKTLNLSFVRDEDERPKVAYNHFSNEIPVISL..A
BAL43067.1_F3H_Ipomoea_coccinea        MT.........TLSTL....TALAGETALRSSFVRDEDERPKVGYNEFSDEIPVISL..K
AAB41102.1_F3H_Ipomoea_purpurea        MT.........TVSTL....TALAGETALRSSFVRDEDERPKVGYNEFSDEIPVISL..K
BAK78917.1_F3H_Ipomoea_quamoclit       MT.........TLSTL....TALAGETALRSSFVRDEDERPKVGYNEFSDEIPVISL..K
ADO95201.1_F3H_Litchi_chinensis        MA..........LATL....TALAQEKTLNASFVRDEDERPKVAYNEFSNEIPVISL..A
AIB06738.1_F3H_Mangifera_indica        MA..........PATL....TALAQEKTLNSKFVRDEDERPKVAYNEFSNEIPVISL..A
AEF14415.1_F3H_Onobrychis_viciifolia   MAP........AARTL....THLAKEKTLESSFVRDEDERPKVAYNNFSTEIPVISL..T
AFI71897.1_F3H_Paeonia_lactiflora      MAP........TATTL....TALAGEKTLQSSFVRDEDERPKVAYNQFSNEIPIISL..A
AEN71544.1_F3H_Paeonia_suffruticosa    MAP........TATTL....TALAGEKTLQSSFVRDEDERPKVAYNQFSNEIPIISL..A
CAA53579.1_F3H_Vitis_vinifera          MA..........PTTL....TALAGEKTLQSSFVRDEDERPKVAYNDFSNEIPVISLTKE
AGS57503.1_F3H_Vitis_rotundifolia      MA..........PTTL....TALAGEKTLQSSFVRDEDERPKVAYNDFSNEIPVISLTKE
ACM17897.1_F3H_Rubus_occidentalis      MAP........TPTTL....TAIAGEKTLQQSFVRDEDERPKVAYNQFSNEIPIISL..S
ABW74548.1_F3H_Rubus_coreanus          MAP........TPTTL....TAIAGEKTLQQSFVRDEDERPKVAYNQFSNEIPIISL..S
ADP09378.1_F3H_Pyrus_pyrifolia         MAP........PAATL....TSIAHEKTLQQKFVRDEDERPKVAYNEFSNEIPIISL..A
AGL50918.1_F3H_Pyrus_communis          MA.........PATTL....TSIAHEKTLQQKFVRDEDERPKVAYNEFSNEIPIISL..A
ACP30361.1_F3H_Malus_hybrid_cultivar   MA.........PATTL....TSIAHEKTLQQKFVRDEDERPKVAYNDFSNEIPIISL..A
ABG78792.1_F3H_Aethusa_cynapium        MA..........PTTL....TALAQEKTLNSKFVRDEDERPKVAYNQFSNEIPVISL..A
AAX21540.1_F3H_Anethum_graveolens      MA..........PSTL....TALAQEKTLNSKFVRDEDERPKIAYNQFSDEIPVISL..A
AAX21539.1_F3H_Ammi_majus              MA..........PTTL....TALAQEKTLNSKFVRDEDERPKIAYNQFSDEIPVISL..A
AAU04791.1_F3H_Fragaria_x_ananassa     MAP........TPTTL....TAIAGENTLQQSFVRDEDERPKVAYNQFSIDIPIISL..S
AAU93347.1_F3H_Ginkgo_biloba           MAPVQSVSAAPVPKFL....PTENEAKVLQSIFVRDEDERPKVAYNEFSKEIPIISL..A
AAT94365.1_F3H_Glycine_max             MAP........TAKTL....TYLAQEKTLESSFVRDEEERPKVAYNEFSDEIPVISL..A
AAX21535.1_F3H_Pimpinella_anisum       MA..........PSTL....TALAQEKTLNSKFVRDEDERPKIAYNQFSDEIPVISL..A
AAP57394.1_F3H_Petroselinum_crispum    MA..........PSTL....TALAQEKTLNSKFVRDEDERPKIAYNKFSDEIPVISL..A
AAC49929.1_F3H_Petunia_x_hybrida       MA..........PSTL....TALAEEKTLQTSFIRDEDERPKVAYNQFSNEIPIISL..E
                                 acc

                                                    α1              β2             α2
NP_190692.1_F3H_Arabidopsis_thaliana   TT
                                            50          60        70        80        90
NP_190692.1_F3H_Arabidopsis_thaliana   GID.DVD.......GKRGEICRQIVEACENWGIFQVVDHGVDTNLVADMTRLARDFFALP
TRINITY_DN33424_c3_g3_i1               GIE...G.......ERRAEICKKIVEACEDWGIFQVIDHGVDSKLISEMTRLAKEFFALP
AAU04792.1_F3H_Fragaria_x_ananassa     GIE.EVE.......GRRAEICKKIVEACEDWGVFQIVDHGIDPKLISEMTRLAREFFALP
NP_001236797.1_F3H_Glycine_max         GID.EVD.......GRRREICEKIVEACENWGIFQVVDHGVDQQLVAEMTRLAKEFFALP
ACR15123.1_F3H_Medicago_truncatula     GID.DAG.......GRRAEICNKIVEACENWGIFQVVDHGVDSKLISEMTRFAKGFFDLP
AQX36284.1_F3H_Prunus_persica          GID.EVE.......GRRADICKKIVEACEDWGIFQIVDHGVDTKLISEMTGLAREFFALP
ABR13013.1_F3H_Triticum_aestivum       GID........G..ARRAEIRDRVAAACEGWGIFQVVDHGVDADLIADMTRLSREFFALP
NP_001130275.1_F3H_Zea_mays            GID........G..ARRAEIRARVAAACEDWGIFQVVDHGVDAALVADMARLARDFFALP
ACL54955.1_F3H_Actinidia_chinensis     GID.EVD.......GRRSEICRKIVEACEDWGIFQVVDHGVDAKLVGEMTRLARDFFALP
AFN70721.1_F3H_Nekemias_grossedentata  GID.EVG.......GRRDEICRKIVEACEDWGIFQVVNHGVDSNLISEMTRLAREFFALP
ACM62745.1_F3H_Garcinia_mangostana     GID.EGG.......QKRAEICKKIVEACEEWGIFQVVDHGVDTKLVSEMTRLARAFFALP
AEO36935.1_F3H_Canarium_album          GID.EVD.......GRRAEICKKIVEACEDWGIFQVIDHGVDANLISDMTRLAREFFALP
AEG64806.1_F3H_Carthamus_tinctorius    GID.DADGAG....SRRGEICAKIVKACEDWGIFQAVDHGVDTKLMSEMTRLAREFFQLP
AFC37250.1_F3H_Camellia_chekiangoleosa GID.EIE.......GRRSEICRKIVEACEDWGIFQVVDHGVDANLVAEMTRLAREFFALP
ACB56921.1_F3H_Pilosella_officinarum   GID.DVEGSGGG.RSRRAEICEKIVKAFEDWGIFQVVDHGVDAKLLSEMKRLAREFFLMP
AAB97310.1_F3H_Chrysanthemum_x_morifolium GID.DVEESSGGIKSRRAEICEKIIKACEDWGIFQVVDHGVDTKMLSEMTRLAREFFEMP
AAC15414.1_F3H_Nicotiana_tabacum       GID.DVEESSGGIKSRRAEICEKIIKACEDWGIFQVVDHGVDTKMLSEMTRLAREFFEMP
ABL86673.1_F3H_Gossypium_barbadense    GID.DVD.......GKRGEICKKIVEACEDWGVFQVVDHGVDTKLVSEMTRFAREFFALP
ADC96713.1_F3H_Gossypium_hirsutum      GID.DVD.......GKRGEICKKIVEACEDWGVFQVVDHGVDTKLVSEMTRFAREFFALP
XP_007046698.1_F3H_Theobroma_cacao     GID.DVD.......GNRAEICKKIVEACEDWGVFQVIDHGVDTKLISEMTRLAREFFALP
ABO48521.1_F3H_Dimocarpus_longan       GID.EVD.......GRRAEICKKIVEACEDWGVFQVIDHGVDTKLISDMTRLAREFFALP
ACQ99190.1_F3H_Fagopyrum_tataricum     GID.EVD.......GKRAEICNKIVKACEEWGIFQVIDHGVDTSLVSEMTRLATEFFALP
BAL43067.1_F3H_Ipomoea_coccinea        GID.DFL.......GRRVQICNDIVKACEDWGIFQVVDHGVDAELIGEMTRLSKDFFSLP
AAB41102.1_F3H_Ipomoea_purpurea        GID.DVL.......GRRVQIRKEIVKACEDWGIFQVVDHGVDAGLIGEMTRLSKDFFSLP
BAK78917.1_F3H_Ipomoea_quamoclit       GID.DVL.......GRRVQIRDEVVKACEDWGIFQVVDHGVDAGLIAEMTRLSKEFFSLP
ADO95201.1_F3H_Litchi_chinensis        GID.EVD.......GRRAEICQKIVEACEDWGIFQVIDHGVDTKLISDMTRLAREFFALP
AIB06738.1_F3H_Mangifera_indica        GID.EVD.......GRRAEICKKIVEACEEWGIFQVIDHGVDIKLISDMTALAREFFSLP
AEF14415.1_F3H_Onobrychis_viciifolia   GID.EVD.......GRRAEICKKIVEACENWGIFQVIDHGIDTNLISDMTRLAKLFFDLP
AFI71897.1_F3H_Paeonia_lactiflora      GID.D.D.......GQRSEICRKIVDACEDWGIFQVIDHGVDANLISDMNRLAREFFALP
AEN71544.1_F3H_Paeonia_suffruticosa    GID.D.D.......GQRSEICRKIVGACEDWGIFQVVDHGVDANLISDMNRLAREFFALP
CAA53579.1_F3H_Vitis_vinifera          SM..KLA.......AVVDEICRKIVEACEDWGIFQVVNHGVDSNLISEMTRLAREFFALP
AGS57503.1_F3H_Vitis_rotundifolia      SM..KLA.......AVVDEICRKIVEACEDWGIFQVVNHGVDSNLISEMTRLAREFFALP
ACM17897.1_F3H_Rubus_occidentalis      GID.EVE.......GRRAEICNKIVEACEDWGVFQIVDHGVDAKLISEMTRLARDFFALP
ABW74548.1_F3H_Rubus_coreanus          GID.EVE.......GRRAEICNKIVEACEDWGVFQIVDHGVDAKLISEMTRLARDFFALP
ADP09378.1_F3H_Pyrus_pyrifolia         GID.EVE.......GRRAEICKKIVEACEDWGIFQIVDHGVDAELVSEMTGLAREFFALP
AGL50918.1_F3H_Pyrus_communis          GID.EVE.......GRRAEICKKIVEACEDWGIFQIVDHGVDAELVSEMTGLAREFFALP
ACP30361.1_F3H_Malus_hybrid_cultivar   GID.EVE.......GRRGEICKKIVAACEDWGIFQIVDHGVDAELISEMTGLAREFFALP
ABG78792.1_F3H_Aethusa_cynapium        GIDDDSG.......DKRSQICRKIVEACEDWGIFQVVDHGIDSHLISEMTRLAREFFALP
AAX21540.1_F3H_Anethum_graveolens      GID..SD.......DKRSEICRKIVEACEDWGIFQVVDHGVDSDLISEMTRLAREFFALP
AAX21539.1_F3H_Ammi_majus              GIDDDSD.......DKRSQICRKIVEACEGWGIFQVVDHGIDTDLISEMTRLAREFFALP
AAU04791.1_F3H_Fragaria_x_ananassa     GIE.EVE.......GRRAEICKKIVEACEDWGVFQIVDHGIDPKLISEMTRLAREFFALP
AAU93347.1_F3H_Ginkgo_biloba           GME........G..EERGRVREEVRAACEEWGIFQVLHHGVPSDLVHRMSQLSRSFFALP
AAT94365.1_F3H_Glycine_max             GID.EVD.......GRRREICEKIVEACENWGIFQVVDHGVDQQLVAEMTRLAKEFFALP
AAX21535.1_F3H_Pimpinella_anisum       GID...D.......DKRSQICRKIVEACEGWGIFQVVDHGIDTDLISEMTRLAREFFALP
AAP57394.1_F3H_Petroselinum_crispum    GIDDDSV.......DKRSQICRKIVEACEDWGIFQVVDHGIDIDLISEMTRLARQFFALP
AAC49929.1_F3H_Petunia_x_hybrida       GID.DET.......GKRAEICDKIVKACEDWGVFQVVDHGVDAELISQMTTFAKEFFALP
                                 acc
```

```
                                              α3                                          β3     β4    η3      α4
NP_190692.1_F3H_Arabidopsis_thaliana                      TT                 TT
                                              100       110       120       130       140       150
NP_190692.1_F3H_Arabidopsis_thaliana          PEDKLRFDMSGGKKGGFIVSSHLQGEAVQDWREIVTYFSYPVRNRDYSRWPDKPEGWVKV
TRINITY_DN33424_c3_g3_i1                      PEEKLRFDMSGGKKGGFIVSSHLQGEAVQDWREIVTYFSYPLRTRDYSRWPDNPKEWRAV
AAU04792.1_F3H_Fragaria_x_ananassa            PEEKLRFDMSGGKKGGFIVSSHLQGEAVQDWREIVTYFSYPVRHRDYSRWPDKPEGWRDV
NP_001236797.1_F3H_Glycine_max                PDEKLRFDMSGAKKGGFIVSSHLQGESVQDWREIVTYFSYPKRERDYSRWPDTPEGWRSV
ACR15123.1_F3H_Medicago_truncatula            PEEKLRFDMSGGKKGGFIVSSHLQGEAVKDWRELVTYFSYPIRQRDYSRWPDKPEGWKEV
AQX36284.1_F3H_Prunus_persica                 SEEKLRFDMSGGKKGGFIVSSHLQGEAVQDWREIVTYFSYPIRHRDYSRWPDKPEGWREV
ABR13013.1_F3H_Triticum_aestivum              AEDKLRYDMSGGKKGGFIVSSHLQGEAVQDWREIVTYFSYPVKARDYGRWPEKPAGWRAV
NP_001130275.1_F3H_Zea_mays                   PEDKLRFDMSGGKKGGFIVSSHLQGEAVQDWREIVTYFSYPVKARDYSRWPDKPAAWRAV
ACL54955.1_F3H_Actinidia_chinensis            PEEKLRFDMSGGKKGGFIVSSHLQGEAVQDWREIVTYFSYPIRARDYSRWPDKPDGWRAV
AFN70721.1_F3H_Nekemias_grossedentata         PEEKLRFDMSGGKKGGFIVSSHLQGEAVQDWREIVTYFSYPLRTRDYSRWPDKPEGWKSV
ACM62745.1_F3H_Garcinia_mangostana            PEEKLRFDMSGGKKGGFIVSSHLQGEAVQDWREIVTYFSYPTRTRDYSRWPDKPDGWVDV
AEO36935.1_F3H_Canarium_album                 PEEKLRFDMTGGKKGGFIVSSHLQGEAVQDWREIVTYFSYPLRSRDYSRWPDKPEGWIEV
AEG64806.1_F3H_Carthamus_tinctorius           REEKLRFDMTGGKKGGYIVSSHLEGEPVEDWRELAIFFSYPINSRDYSRWPDKPKEWRAV
AFC37250.1_F3H_Camellia_chekiangoleosa        PEEKLRFDMSGGKKGGFIVSSHLQGEAVQDWREIVTYFSYPIRARDYSRWPDKPEGWRAV
ACB56921.1_F3H_Pilosella_officinarum          EEEKLKFDMAGGKKGGFLVSSHLKGETVQNWREFVTYFSYPIKARDYSRWPNKPNEWRSV
AAB97310.1_F3H_Chrysanthemum_x_morifolium     AEEKLRFDMTGGKKGGFIVSSHLQGEAVQDWREIVTYFSYPIKARDYSRWPDKPKEWRAV
AAC15414.1_F3H_Nicotiana_tabacum              AEEKLRFDMTGGKKGGFIVSSHLQGEAVQDWREIVTYFSYPIKARDYSRWPDKPKEWRAV
ABL86673.1_F3H_Gossypium_barbadense           AEEKLRFDMSGGKKGGFIVSSHLQGEAVQDWREIVTYFSYPLKSRDYSRWPDKPEGWIEV
ADC96713.1_F3H_Gossypium_hirsutum             AEEKLRFDMSGGKKGGFIVSSHLQGEAVQDWREIVTYFSYPLKSRDYSRWPDKPEGWIEV
XP_007046698.1_F3H_Theobroma_cacao            AEEKLRFDMSGGKKGGFIVSSHLQGEAVQDWREIVTYFSYPLKARDYSRWPDKPEGWVEV
ABO48521.1_F3H_Dimocarpus_longan              PEEKLRFDMSGGKKGGFIVSSHLQGEAVQDWREIVTYFSYPMRTRDYSRWPDKPEGWIDV
ACQ99190.1_F3H_Fagopyrum_tataricum            PEEKLKFDMSGGKKGGFIVSSHLQGEAVQDWREIVTYFSYPLKSRDYSRWPNNPEGWIKV
BAL43067.1_F3H_Ipomoea_coccinea               PEEKLRFDMSGGKKGGFIVSSHLQGEAVKDWREIVTYFSYPIKARDYSRWPDKPEGWRAV
AAB41102.1_F3H_Ipomoea_purpurea               PEEKLRFDMSGGKKGGFIVSSHLQGEAVKDWREIVTYFSYPVRARDYSRWPDKPEGLRAV
BAK78917.1_F3H_Ipomoea_quamoclit              PEEKLRFDMSGGKKGGFIVSSHLQGEAVKDWREIVTYFSYPVRARDYSRWPDKPEGWRAV
ADO95201.1_F3H_Litchi_chinensis               PEEKLRFDMSGGKKGGFIVSSHLQGEAVQDWREIVTYFSYPMRTRDYSRWPDKPQGWIDV
AIB06738.1_F3H_Mangifera_indica               PVEKLRFDMSGGKKGGFIVSSHLQGETVQDWREIVTYFSYPIKNRDYSRWPDKPEGWRDV
AEF14415.1_F3H_Onobrychis_viciifolia          PEEKLRFDMSGGKKGGFIVSSHLQGEAVKDWREIVTYFSYPIRARDYSRWPDKPEGWKEV
AFI71897.1_F3H_Paeonia_lactiflora             PEDKLRFDMSGGKKGGFIVSSHLQGEAVRDWREIVTYFSYPIRNRDYSRWPDKPGGWKRV
AEN71544.1_F3H_Paeonia_suffruticosa           PDDKLRFDMSGGKKGGFIVSSHLQGEAVRDWREIVTYFSYPIRNRDYSRWPDKPEGWKRV
CAA53579.1_F3H_Vitis_vinifera                 PEENVRFDMSGGKKGGFIVSSHLQGEAVQDWREIVTYFSYPLRTRDYSRWPDKPEGWRSV
AGS57503.1_F3H_Vitis_rotundifolia             PEEKLRFDMSGGKKGGFIVSSHLQGEAVQDWREIVTYFSYPLRTRDYSRWPDKPEGWRSV
ACM17897.1_F3H_Rubus_occidentalis             PEEKLRFDMSGGKKGGFIVSSHLQGEAVQDWREIVTYFSYPVRHRDYSRWPDKPEGWRAV
ABW74548.1_F3H_Rubus_coreanus                 PEEKLRFDMSGGKKGGFIVSSHLQGEAVQDWREIVTYFSYPVRHRDYSRWPDKPEGWRAV
ADP09378.1_F3H_Pyrus_pyrifolia                SEEKLRFDMSGGKKGGFIVSSHLQGEAVQDWREIVTYFSYPIRHRDYSRWPDKPEAWREV
AGL50918.1_F3H_Pyrus_communis                 SEEKLRFDMSGGKKGGFIVSSHLQGEAVQDWREIVTYFSYPIRHRDYSRWPDKPEAWREV
ACP30361.1_F3H_Malus_hybrid_cultivar          SEEKLRFDMSGGKKGGFIVSSHLQGEAVQDWREIVTYFSYPIRHRDYSRWPDKPEAWREV
ABG78792.1_F3H_Aethusa_cynapium               AEEKLRFDMTGGKKGGFIVSSHLQGEAVQDWREIVTFFSYPVPARDYSRWPDKPEGWRSV
AAX21540.1_F3H_Anethum_graveolens             AEEKLRFDMSGGKKGGFIVSSHLQGEAVQDWREIVTYFSYPIQARDYSRWPDKPEGWRSI
AAX21539.1_F3H_Ammi_majus                     AEEKLRFDMTGGKKGGFIVSSHLQGEAVQDWREIVTYFSYPVPARDYSRWPDKPEGWRSV
AAU04791.1_F3H_Fragaria_x_ananassa            PEEKLRFDMSGGKKGGFIVSSHLQGEAVQDWREIVTYFSYPVRHRDYSRWPDKPEGWRDV
AAU93347.1_F3H_Ginkgo_biloba                  SHEKLKFDMTGGKRGGFVVSSHLQGESVLDWREIFTYFSYPLRSRDYSRWPQNPDGWREV
AAT94365.1_F3H_Glycine_max                    PDEKLRFDMSGAKKGGFIVSSHLQGESVQDWREIVTYFSYPKRERDYSRWPDTPEGWRSV
AAX21535.1_F3H_Pimpinella_anisum              AEEKLRFDMTGGKKGGFIVSSHLQGEAVQDWREIVTYFSYPVPARDYSRWPDKPEGWRSV
AAP57394.1_F3H_Petroselinum_crispum           AEEKLRFDMTGGKKGGFIVSSHLQGEAVQDWREIVTYFSYPIQARDYSRWPDKPEGWRSI
AAC49929.1_F3H_Petunia_x_hybrida              PEEKLRFDMSGGKKGGFIVSSHLQGEVVQDWREIVTYFSYPTRARDYSRWPDKPEGWIAV
                                        acc
```

```
                                                      α5                   α6  η4           β5             β6
NP_190692.1_F3H_Arabidopsis_thaliana
                                              160       170       180       190       200       210
NP_190692.1_F3H_Arabidopsis_thaliana          TEEYSERLMSLACKLLEVLSEAMGLEKESLTNACVDMDQKIVVNYYPKCPQPDLTLGLKR
TRINITY_DN33424_c3_g3_i1                      TEEYSEKLMGLACKLLEVLSEAMGLEKEALTKACVDMDQKVVVNYYPKCPQPDLTLGLKR
AAU04792.1_F3H_Fragaria_x_ananassa            TTQYSDELMGLACKLLEVLSEAMGLEKEALTKACVDMDQKVVVNFYPKCPQPDLTLGLKR
NP_001236797.1_F3H_Glycine_max                TEEYSDKVMGLACKLMEVLSEAMGLEKEGLSKACVDMDQKVVVNYYPKCPQPDLTLGLKR
ACR15123.1_F3H_Medicago_truncatula            TEQYSEKLMNLACKLLEVLSEAMGLEKDALTKACVDMDQKVVINYYPKCPQPDLTLGLKR
AQX36284.1_F3H_Prunus_persica                 TQKYSDELMGLACKLLGVLSEAMGLDTEALTKACVDMDQKVVVNFYPKCPQPDLTLGLKR
ABR13013.1_F3H_Triticum_aestivum              VERYSERLMGLSCKLLGVLSEAMGLESEALAKACVDMDQKVVVNFYPRCPQPDLTLGLKR
NP_001130275.1_F3H_Zea_mays                   VERYSEQLMALACRLLGVLSEAMGLDTEALARACVDMDQKVVVNFYPRCPQPDLTLGLKR
ACL54955.1_F3H_Actinidia_chinensis            TQAYSENLMGLACKLLEVLSEAMGLEKEALTKACIDMDQKVVVNFYPKCPQPDLTLGLKR
AFN70721.1_F3H_Nekemias_grossedentata         TQEYSEKLMGLACKLLEVLSEAMDLEKEALTKACVDMDQKVVVNFYPQCPQPGLTLGLKR
ACM62745.1_F3H_Garcinia_mangostana            TKDYGDQLMGLACKLLEVLSEAMGLEKEALTKACVDMDQKIVVNYYPKCPQPDLTLGLKR
AEO36935.1_F3H_Canarium_album                 TKEYSDKLMGLACKLLEVLSEAMGLEKEALTKACVDMDQKVVVNYYPKCPQPDLTLGLKR
AEG64806.1_F3H_Carthamus_tinctorius           TAEYSETLMGLACKLLEVLSEAMGLEKEALTKACVNMDQKMVVNYYPKCPHPDLTLGLKR
AFC37250.1_F3H_Camellia_chekiangoleosa        TETYSEKLMDLACKLLEVLSEAMGLEKEALTKACVDMDQKVVVNFYPKCPQPDLTLGLKR
ACB56921.1_F3H_Pilosella_officinarum          TEEYSEKSMGLACKLLEVLSEAMGLEKEALTKACVDMDQKVVVNYYPKCPQPDLTLGLKR
AAB97310.1_F3H_Chrysanthemum_x_morifolium     TEKYSEDLMGLGCKLLEVLSEAMGLEKEALKDACVDMDQKVVLNYYPKCPQPDLTLGLKR
AAC15414.1_F3H_Nicotiana_tabacum              TEKYSEDLMGLGCKLLEVLSEAMGLEKEALKDACVDMDQKVVLNSYPKCPQPDLTLGLKR
ABL86673.1_F3H_Gossypium_barbadense           TKEYSEKLMGLACKLLEVLSEAMGLEKEALTKACVDMDQKVVVNFYPKCPQPDLTLGLKR
ADC96713.1_F3H_Gossypium_hirsutum             TKEYSEKLMGLACKLLEVLSEAMGLEKEALTKACVDMDQKVVVNFYPKCPQPDLTLGLKR
XP_007046698.1_F3H_Theobroma_cacao            TKEYSDKLMGLACKLLEVLSEAMDLDKEALTKACVDMDQKVVVNFYPKCPQPDLTLGLKR
ABO48521.1_F3H_Dimocarpus_longan              TKEYSDKLMGLACKLLEVLSEAMGLEKEALTKACVDMDQKVVVNYYPKCPQPDLTLGLKR
ACQ99190.1_F3H_Fagopyrum_tataricum            TEKYSENLMALACKLLGVLSEAMGLETEALTKACVDMDQKVVVNCYPKCPQPDLTLGLKR
BAL43067.1_F3H_Ipomoea_coccinea               TEKYSEKLMDLACKLLEVLSEAMGLENEALTKACVEMDQKLVVNFYPKCPQPDLTLGLKR
AAB41102.1_F3H_Ipomoea_purpurea               TEKYSEKLMDLACKLLEVLSEAMGLEKEALTKACVEMDQKLVVNFYPKCPQPDLTLGLKR
BAK78917.1_F3H_Ipomoea_quamoclit              TERYSEKLMDLACKLLEVLSEAMGLEKEALTKACVEMDQKLVVNFYPKCPQPDLTLGLKR
ADO95201.1_F3H_Litchi_chinensis               TKEYSDKLMGLACKLLEVLSEAMGLEKEALTNACVDMDQKVVVNYYPKCPQSDLTLGLKR
AIB06738.1_F3H_Mangifera_indica               TKEYSDKLMGLACKLLEVLSEAMGLDKEALTKACVDMDQKVVVNFYPTCPEPELTLGLKR
AEF14415.1_F3H_Onobrychis_viciifolia          TEEYSEKLMGLACKLLEVLSEAMGLEKEALTKACVDMDQKVVINYYPKCPEPDLTLGLKR
AFI71897.1_F3H_Paeonia_lactiflora             TEEYSEKLMGLACKLLEVLSEAMGLEKEALTNACVDMDQKVVVNYYPKCPQPDLTLGLKR
AEN71544.1_F3H_Paeonia_suffruticosa           TEEYSEKLMGLACKLLEVLSEAMGLEKEALTKACVDMDQKVVVNYYPKCPQPDLTLGLKR
CAA53579.1_F3H_Vitis_vinifera                 TQEYSEKLMGLACKLLEVLSEAMDLDKDALTNACVDMDQKVVVNFYPQCPQPDLTLGLKR
AGS57503.1_F3H_Vitis_rotundifolia             TQEYSEKLMGLACKLLEVLSEAMDLDKDALTNACVDMDQKVVVNFYPQCPQPDLTLGLKR
ACM17897.1_F3H_Rubus_occidentalis             TQQYSDELMGLACKLLEVLSEAMGLEKEALTKACVDMDQKVVVNFYPKCPQPDLTLGLKR
ABW74548.1_F3H_Rubus_coreanus                 TQQYSDELMGLACKLLEVLSEAMGLEKEALTKACVDMDQKVVVNFYPKCPQPDLTLGLKR
ADP09378.1_F3H_Pyrus_pyrifolia                TKKYSDELMGLACKLLGVLSEAMGLDTEALTKACVDMDQKVVVNFYPKCPQPDLTLGLKR
AGL50918.1_F3H_Pyrus_communis                 TKKYSDELMGLACKLLGVLSEAMGLDTEALTKACVDMDQKVVVNFYPKCPQPDLTLGLKR
ACP30361.1_F3H_Malus_hybrid_cultivar          TKKYSDELMGLACKLLGVLSEAMGLDTEALTKACVDMDQKVVVNFYPKCPQPDLTLGLKR
ABG78792.1_F3H_Aethusa_cynapium               TEVYSEKLMALGCKLLEVLSEAMGLEKETITKACVDMDQKVIVNYYPKCPQPDLTLGLKR
AAX21540.1_F3H_Anethum_graveolens             TEVYSEKLMALACKLLEVLSEAMGLEKETITKACVDMDQKVIVNYYPKCPQPDLTLGLKR
AAX21539.1_F3H_Ammi_majus                     TEVYSDKLMGLACKLLEVLSEAMGLEKEALTKACVDMDQKVIVNYYPKCPQPDLTLGLKR
AAU04791.1_F3H_Fragaria_x_ananassa            TTQYSDELMGLACKLLEVLSEAMGLEKEALTKACVDMDQKVVVNFYPKCPQPDLTLGLKR
AAU93347.1_F3H_Ginkgo_biloba                  VEEYSQELMKLACKLLEIISESLGLEPQAVTKACLDMDQKVVINFYPKCPQPDMTLGLKR
AAT94365.1_F3H_Glycine_max                    TEEYSDKVMGLACKLMEVLSEAMGLEKEGLSKACVDMDQKVVVNYYPKCPQPDLTLGLKR
AAX21535.1_F3H_Pimpinella_anisum              TEVYSDKLMGLACKLLEVLSEAMGLEKEALTKACVDMDQKVIVNYYPKCPQPDLTLGLKR
AAP57394.1_F3H_Petroselinum_crispum           TEMYSDELMALACKLLEVLSEAMGLEKEGLTKACVDMDQKVIVNYYPKCPQPNLTLGLKR
AAC49929.1_F3H_Petunia_x_hybrida              TQKYSEKLMELACKLLDVLSEAMGLEKEALTKACVDMDQKVVVNFYPKCPEPDLTLGLKR
                                        acc
```

```
NP_190692.1_F3H_Arabidopsis_thaliana        β7  β8  β9  TT  β10  α7  β11
                                            220       230       240       250       260       270
NP_190692.1_F3H_Arabidopsis_thaliana        HTDPGTITLLLQDQVGGLQATRDNGKTWITVQPVEGAFVVNLGDHGHFLSNGRFKNADHQ
TRINITY_DN33424_c3_g3_i1                    HTDPGTITLLLQDQVGGLQATRDNGKTWITVQPVEGAFVVNLGDHGHFLSNGRFRNADHQ
AAU04792.1_F3H_Fragaria_x_ananassa          HTDPGTITLLLQDQVGGLQATRDGGKTWITVQPVEGAFVVNLGDHGHFLSNGRFKNADHQ
NP_001236797.1_F3H_Glycine_max              HTDPGTITLLLQDQVGGLQATRDNGKTWITVQPVEAAFVVNLGDHAHYLSNGRFKNADHQ
ACR15123.1_F3H_Medicago_truncatula          HTDPGTITLLLQDQVGGLQATKDNGKTWITVQPVEGAFVVNLGDHGHYLSNGRFKNADHQ
AQX36284.1_F3H_Prunus_persica               HTDPGTITLLLQDQVGGLQATRDDGKTWITVQPVEGAFVVNLGDHGHFLSNGRFKNADHQ
ABR13013.1_F3H_Triticum_aestivum            HTDPGTITLLLQDLVGGLQATRDGGKTWITVQPISGAFVVNLGDHGHFMSNGRFKNADHQ
NP_001130275.1_F3H_Zea_mays                 HTDPGTITLLLQDLVGGLQATRDGGRTWITVQPVEGAFVVNLGDHGHLLSNGRFKNADHQ
ACL54955.1_F3H_Actinidia_chinensis          HTDPGTITLLLQDQVGGLQATRDGGKTWITVQPVDGAFVVNLGDHGHYLSNGRFKNADHQ
AFN70721.1_F3H_Nekemias_grossedentata       HTDPGTITLLLQDQVGGLQATRDGGKTWITVQPVEGAFVVNLGDHGHYLSNGRFKNADHQ
ACM62745.1_F3H_Garcinia_mangostana          HTDPGTITLLLQDQVGGLQATRDNGETWITVQPIEGAFVVNLGDHGHFLSNGRFRNADHQ
AEO36935.1_F3H_Canarium_album               HTDPGTITLLLQDQVGGLQATRDDGKTWITVQPVEEAFVVNLGDHGHHLSNGRFRNADHQ
AEG64806.1_F3H_Carthamus_tinctorius         HTDAGTITLLLQDQVGGLQATRDGGESWITVQPIEGAFVVNLGDHGHYLSNGRFKNADHQ
AFC37250.1_F3H_Camellia_chekiangoleosa      HTDPGTITLLLQDQVGGLQATRDGGKTWITVQPVEGAFVVNLGDHGHYLSNGRFKNADHQ
ACB56921.1_F3H_Pilosella_officinarum        HTDPGTITLVLQDLVGGLQATRDGGKSWITVEPIEGAFVVNLGDHGHYLSNGRFKNADHQ
AAB97310.1_F3H_Chrysanthemum_x_morifolium   HTDPGTITLLLQDQVGGLQATRDGGKTWITVEPIEGAFVVNLGDHGHYLSNGRFKNADHQ
AAC15414.1_F3H_Nicotiana_tabacum            HTDPGTITLLLQDQVGGLQATRDGGKTWITVEPIEGAFVVNLGDHGHYLSNGRFKNADHQ
ABL86673.1_F3H_Gossypium_barbadense         HTDPGTITLLLQDQVGGLQATRDNGKTWITVQPVEGAFVVNLGDHGHYLSNGRFKNADHQ
ADC96713.1_F3H_Gossypium_hirsutum           HTDPGTITLLLQDQVGGLQATRDNGKTWITVQPVEGAFVVNLGDHGHYLSNGRFKNADHQ
XP_007046698.1_F3H_Theobroma_cacao          HTDPGTITLLLQDQVGGLQATRDNGKTWITVQPVEGAFVVNLGDHGHFLSNGRFKNADHQ
ABO48521.1_F3H_Dimocarpus_longan            HTDPGTITLLLQDQVGGLQATRDNGKTWITVQPVDGAFVVNLGDHGHYLSNGRFKNADHQ
ACQ99190.1_F3H_Fagopyrum_tataricum          HTDPGTITLLLQDQVGGLQATRDDGKTWITVQPIEGAFVVNLGDHGHYLSNGRFRNADHQ
BAL43067.1_F3H_Ipomoea_coccinea             HTDPGTITLLLQDQVGGLQATKDGGKTWITVQPVDGAFVVNLGDHGHFLSNGRFKNADHQ
AAB41102.1_F3H_Ipomoea_purpurea             HTDPGTITLLLQDQVGGLQATKDGGKTWITVQPVDGAFVVNLGDHGHFLSNGRFKNADHQ
BAK78917.1_F3H_Ipomoea_quamoclit            HTDPGTITLLLQDQVGGLQATKDGGKTWITVQPVDGAFVVNLGDHGHFLSNGRFKNADHQ
ADO95201.1_F3H_Litchi_chinensis             HTDPGTITLLLQDQVGGLQATRDNGKTWITVQPVDGAFVVNLGDHGHYLSNGRFKNADHQ
AIB06738.1_F3H_Mangifera_indica             HTDPGTITLLLQDQVGGLQATRDNGETWITVQPVDGAFVVNLGDHGHYLSNGRFKNADHQ
AEF14415.1_F3H_Onobrychis_viciifolia        HTDPGTITLLLQDQVGGLQATKDNGKTWITVQPIEGAFVVNLGDHGHFLSNGRFKNADHQ
AFI71897.1_F3H_Paeonia_lactiflora           HTDPGTITLLLQDQVGGLQATRDGGKTWITVQPVEGAFVVNLGDHGHYLSNGRFKNADHQ
AEN71544.1_F3H_Paeonia_suffruticosa         HTDPGTITLLLQDQVGGLQATRDGGKTWITVQPVEGAFVVNLGDHGHYLSNGRFKNADHQ
CAA53579.1_F3H_Vitis_vinifera               HTDPGTITLLLQDQVGGLQATRDGGKTWITVQPVEGAFVVNLGDHGHYLSNGRFKNADHQ
AGS57503.1_F3H_Vitis_rotundifolia           HTDPGTITLLLQDQVGGLQATRDGGKTWITVQPVEGAFVVNLGDHGHYLSNGRFKNADHQ
ACM17897.1_F3H_Rubus_occidentalis           HTDPGTITLLLQDQVGGLQATRDGGKTWITVQPVEGAFVVNLGDHGHFLSNGRFKNADHQ
ABW74548.1_F3H_Rubus_coreanus               HTDPGTITLLLQDQVGGLQATRDGGKTWITGQPVEGAFVVNLGDHGHFLSNGRFKNADHQ
ADP09378.1_F3H_Pyrus_pyrifolia              HTDPGTITLLLQDQVGGLQATRDDGKTWITVQPVEGAFVVNLGDHGHFLSNGRFKNADHQ
AGL50918.1_F3H_Pyrus_communis               HTDPGTITLLLQDQVGGLQATRDGGKTWITVQPVEGAFVVNLGDHGHFLSNGRFKNADHQ
ACP30361.1_F3H_Malus_hybrid_cultivar        HTDPGTITLLLQDQVGGLQATRDDGKTWITVQPVEGAFVVNLGDHGHLLSNGRFKNADHQ
ABG78792.1_F3H_Aethusa_cynapium             HTDPGTITLLLQDQVGGLQATRDGGKTWITVQPVEGAFVVNLGDHGHYLSNGRFKNADHQ
AAX21540.1_F3H_Anethum_graveolens           HTDPGTITLLLQDQVGGLQATRDGGKTWITVQPVEGAFVVNLGDHGHYLSNGRFKNADHQ
AAX21539.1_F3H_Ammi_majus                   HTDPGTITLLLQDQVGGLQATRDGGKTWITVQPVEGAFVVNLGDHGHYLSNGRFKNADHQ
AAU04791.1_F3H_Fragaria_x_ananassa          HTDPGTITLLLQDQVGGLQATRDGGKTWITVQPVEGAFVVNLGDHGHFLSNGRFKNADHQ
AAU93347.1_F3H_Ginkgo_biloba                HTDPGTITLLLQDQVGGLQATKD.GLNWITVEPVEGAFVVNLGDHMHYLSNGKYKNADHQ
AAT94365.1_F3H_Glycine_max                  HTDPGTITLLLQDQVGGLQATRDNGKTWITVQPVEAAFVVNLGDHAHYLSNGRFKNADHQ
AAX21535.1_F3H_Pimpinella_anisum            HTDPGTITLLLQDQVGGLQATRDGGKTWITVQPVEGAFVVNLGDHGHYLSNGRFKNADHQ
AAP57394.1_F3H_Petroselinum_crispum         HTDPGTITLLLQDQVGGLQATRDGGKTWITVQPVEGAFVVNLGDHGHYLSNGRFKNADHQ
AAC49929.1_F3H_Petunia_x_hybrida            HTDPGTITLLLQDQVGGLQATKDNGKTWITVQPVEGAFVVNLGDHGHFLSNGRFKNADHQ
acc
```

```
NP_190692.1_F3H_Arabidopsis_thaliana        β12  TT  α8
                                            280       290       300       310       320       330
NP_190692.1_F3H_Arabidopsis_thaliana        AVVNSNSSRLSIATFQNPAPDATVYPLK..VREGEKAILEEPITFAEMYKRKMGRDLELA
TRINITY_DN33424_c3_g3_i1                    AVVNSNCSRLSIATFQNPAPEATVYPLK..IEEGVKPVLDEPMTFAEMYRRKMSKDLELA
AAU04792.1_F3H_Fragaria_x_ananassa          AVVNSNHSRLSIATFQNPAQEAIVYPLK..VREGEKPILEEPMTYTEMYKKKMSKDLELA
NP_001236797.1_F3H_Glycine_max              AVVNSNHSRLSIATFQNPAPNATVYPLK..IREGEKPVMEEPITFAEMYRRKMSKDIEIA
ACR15123.1_F3H_Medicago_truncatula          AVVNSNYSRLSIATFQNPAPDATVYPLK..IRDGEKSVMEEPITFAEMYRRKMSKDIEIA
AQX36284.1_F3H_Prunus_persica               AVVNSNSSRLSIATFQNPAQEATVYPLS..IREGEKPILEGPITYTEMYKKKMTKDLELA
ABR13013.1_F3H_Triticum_aestivum            AVVNGESSRLSIATFQNPAPDARVWPLA..VREGEEPILEEPITFAEMYRRKMERDLDLA
NP_001130275.1_F3H_Zea_mays                 AVVNSECSRLSIATFQNPAPDATVYPLA..VRDGEAPILDHPITFAEMYRRKMARDIELA
ACL54955.1_F3H_Actinidia_chinensis          AVVNSNCSRLSIATFQNPAPEATVYPLA..IQEGEKPVLEEPITFAEMYRRKMSKDLELA
AFN70721.1_F3H_Nekemias_grossedentata       AVVNSNYSRLSIATFQNPAPEATVYPLK..IREGEKAVLEEPITFAEMYRRKMSKDLELA
ACM62745.1_F3H_Garcinia_mangostana          AVVNSNCSRLSIATFQNPAPDAIVYPLK..IREGEKSILEEPITFSEMYRRKMAKDLELA
AEO36935.1_F3H_Canarium_album               AVVNSDCSRLSIATFQNPAPNATVYPLK..IREGEKPVLEEPITFAEMYRRKMSRDLELA
AEG64806.1_F3H_Carthamus_tinctorius         ARVNSNTNRLSIATFQNPAPDAIVYPLK..VNEGDRSIMEKPITSTEMYERKMSKDLELA
AFC37250.1_F3H_Camellia_chekiangoleosa      AVVNSNCSRLSIATFQNPAPEATVYPLK..IREGEKPVLEEPITFADMYKRKMSKDIELA
ACB56921.1_F3H_Pilosella_officinarum        AVVNSSTSRLSIATFQNPAPEAIVYPLK..INEGEKSIMEEAMTFMDMYKKKMSGDLELA
AAB97310.1_F3H_Chrysanthemum_x_morifolium   AVVNSNTSRLSIATFQNPAPNAIVYPLI..VNEGEKSIMEEAITFMDMYKKKMSTDLELA
AAC15414.1_F3H_Nicotiana_tabacum            AVVNSNTSRLSIATFQNPAPNAIVYPLI..VNEGEKSIMEEAITFMDMYKKKMSTDLELA
ABL86673.1_F3H_Gossypium_barbadense         AVVNSNCSRLSIATFQNPAPDATVYPLK..IREGEKPILEEPITFAEMYRRKMSKDLELA
ADC96713.1_F3H_Gossypium_hirsutum           AVVNSNCSRLSIATFQNPAPDATVYPLK..IREGEKPILEEPITFAEMYRRKMSKDLELA
XP_007046698.1_F3H_Theobroma_cacao          AVVNSDCSRLSIATFQNPAPDATVYPLR..IREGEQPILEEPITFAEMYRRKMSKDLELA
ABO48521.1_F3H_Dimocarpus_longan            AVVDSNCSRLSIATFQNPAPEATVYPLK..IREGEKPILEEPITFAEMYRRKMSKDLELA
ACQ99190.1_F3H_Fagopyrum_tataricum          AVVNSNSSRLSIATFQNPAPEATVYPLA..IREGEKPILEEPITFAEMYRRKMSKDLELA
BAL43067.1_F3H_Ipomoea_coccinea             AVVNSERSRVSIATFQNPAPEAKVYPLK..VREGEKPILEEPITFAEMYRRKMSKDLELA
AAB41102.1_F3H_Ipomoea_purpurea             AVVNSEHSRMSIATFQNPAPEAKVYPLK..VREGEKAILEEPITFAEMYRRKMSKDLELA
BAK78917.1_F3H_Ipomoea_quamoclit            AVVNSERSRVSIATFQNPAPEAKVYPLT..VREGEKPILEEPITFAEMYRRKMSKDLELA
ADO95201.1_F3H_Litchi_chinensis             AVVNSNCSRLSIATFQNPAPEATVYPLK..IREGEKPILEQPITFAEMYRRKMSKDLELA
AIB06738.1_F3H_Mangifera_indica             AVVNSNCSRLSIATFQNPAPEATVYPLK..IREGEEAILEEPITFAEMYRRKMSKDLELA
AEF14415.1_F3H_Onobrychis_viciifolia        AVVNSNYSRLSIATFQNPAPDATVYPLK..VREGEKSVMEEPITFAEMYRRKMSKDLEIA
AFI71897.1_F3H_Paeonia_lactiflora           AVVNSNYSRLSIATFQNPAPEAIVYPLK..IRDGEKSVLEKPITFTEMYRTKMSKDIELA
AEN71544.1_F3H_Paeonia_suffruticosa         AVVNSNYSRLSIATFQNPAPEAIVYPLK..IRDGEKPVLEKPITFTEMYRTKMSKDIELA
CAA53579.1_F3H_Vitis_vinifera               AVVNSNHSRLSIATFQNPAPEATVYPLK..IREGEKAVLEGPITFAEMYRRKMSKDLELA
AGS57503.1_F3H_Vitis_rotundifolia           AVVNSNHSRLSIATFQNPAPEATVYPLK..IREGEKAVLEGPITFAEMYRRKMSKDLELA
ACM17897.1_F3H_Rubus_occidentalis           AVVNSNHSRLSIATFQNPAPEATVYPLK..VREGEKPILEEPITYTEMYKKKMSKDLELA
ABW74548.1_F3H_Rubus_coreanus               AVVNSNHSRLSIATFQNPAQEAIVYPLK..VREGEKPILEEPITYTEMYKKKMSKDLELA
ADP09378.1_F3H_Pyrus_pyrifolia              AVVNSNSSRLSIATFQNPAQDAIVYPLS..VREGEKPILEAPITYTEMYKKKMSKDLEVA
AGL50918.1_F3H_Pyrus_communis               AVVNSNSSRLSIATFQNPAQEAIVYPLS..VREGEKPILEAPITYTEMYKKKMSKDLELA
ACP30361.1_F3H_Malus_hybrid_cultivar        AVVNSNSSRLSIATFQNPAQEATVYPLS..VREGEKPILEAPITYTEMYKKKMSKDLELA
ABG78792.1_F3H_Aethusa_cynapium             AVVNSDSSRMSIATFQNPAPNATVYPLK..IREGEKSVMEEAITFAEMYRRKMSKDIEVA
AAX21540.1_F3H_Anethum_graveolens           AVVNSNSSRMSIATFQNPAPNATVYPLK..IREGEKAVMEEPITFAEMYKRKMSKDIELA
AAX21539.1_F3H_Ammi_majus                   AVVNSNSSRMSIATFQNPAPNATVYPLK..IREGEKAVMEEPITFAEMYKRKMSKDIEMA
AAU04791.1_F3H_Fragaria_x_ananassa          AVVNSNHSRLSIATFQNPAQEAIVYPLK..VREGEKPILEEPMTYTEMYKKKMSKDLELA
AAU93347.1_F3H_Ginkgo_biloba                AVVNADTSRLSIATFQNPAPEATVYPLEGVVKAEDKCVMEEPITFAQMYSHKMNRDLELA
AAT94365.1_F3H_Glycine_max                  AVVNSNHSRLSIATFQNPAPNATVYPLK..IREGEKPVMEEPITFAEMYRRKMSKDIEIA
AAX21535.1_F3H_Pimpinella_anisum            AVVNSNSSRMSIATFQNPAPNATVYPLT..IREGEKAVMEEPITFAEMYKRKMSKDIEMA
AAP57394.1_F3H_Petroselinum_crispum         AVVNSNSSRMSIATFQNPAPNATVYPLK..IREGEKAVMEEPITFAEMYKRKMSRDIEMA
AAC49929.1_F3H_Petunia_x_hybrida            AVVNSNSSRMSIATFQNPAPNATVYPLK..IREGEKSIMDEPITFAEMYRRKMSKDLELA
acc
```

```
                                                   α9
NP_190692.1_F3H_Arabidopsis_thaliana     ℓℓℓℓℓℓ.ℓℓ....................ℓℓ
                                           340                 350
NP_190692.1_F3H_Arabidopsis_thaliana     RLKKLAK.EE...................RDHK........EVDKP..VDQIFA......
TRINITY_DN33424_c3_g3_i1                 RLKKLAK.EQ....QAKDL..........EKSK........LEGKP..IEEIFA......
AAU04792.1_F3H_Fragaria_x_ananassa       RLKKHAK.EQ....L.QDS..........EKTK........LEAKP..VDDIFA......
NP_001236797.1_F3H_Glycine_max           RMKKLAK.EK....HLQDLENEKHLQELDQKAK........LEAKP..LKEILA......
ACR15123.1_F3H_Medicago_truncatula       RMKKLAK.EEK...ELRDL..........EKAK........IEAKP..LNEILA......
AQX36284.1_F3H_Prunus_persica            RLKKLAK.EQ....QLQDS..........EK...........EGKP..KDDIFA......
ABR13013.1_F3H_Triticum_aestivum         KRKKQAK.DQL...MQQQLQLQ.......QQQQAVAAAPMPTATKS..LNEILA......
NP_001130275.1_F3H_Zea_mays              RLKKQVKADKS...ANKEF         .............ADSKP..LDAILA......
ACL54955.1_F3H_Actinidia_chinensis       RLKKLAK.ETQL..QEQEL..........EKAK........LGAKG..VEEIFA......
AFN70721.1_F3H_Nekemias_grossedentata    RLKKLAK.EQ....QLQDL..........EKAK........LESKP..IDEIFA......
ACM62745.1_F3H_Garcinia_mangostana       RLKKLAK.EQ....QLTDV..........EKAK........LEAKP..IEKILA......
AEO36935.1_F3H_Canarium_album            RLKKLAK.EK....QLQDL..........EKAK........LEGKP..LEQILA......
AEG64806.1_F3H_Carthamus_tinctorius      RLKKLAK.EKQ.....QNL..........EKE............KS..IEKISA......
AFC37250.1_F3H_Camellia_chekiangoleosa   KLKKLAK.EKKLLQDQQDI..........EKAK........LEIKS..TDEIFA......
ACB56921.1_F3H_Pilosella_officinarum     RLKKLAK.DKK.....DNL..........EKV............KP..TKNIFV......
AAB97310.1_F3H_Chrysanthemum_x_morifolium RLKKLAK.AKQ.....QDL..........EKA............KP..IESILA......
AAC15414.1_F3H_Nicotiana_tabacum         RLKKLAK.AKQ.....QDL..........EKA............KP..IESILAKRLADF
ABL86673.1_F3H_Gossypium_barbadense      RLKKVAK.EQQ...QLKEKEAE......NEKPK........LEAKP..LEEILA......
ADC96713.1_F3H_Gossypium_hirsutum        RLKKVAK.EQQ...QLKEKEAE......NEKPK........LEAKP..LEEILA......
XP_007046698.1_F3H_Theobroma_cacao       RLKKLAK.EQQ...QVQEI..........EKTK........LEAKP..LEKILA......
ABO48521.1_F3H_Dimocarpus_longan         RLKKLAK.EQKL..QLQDN..........EKSK........LEAKP..IEQILA......
ACQ99190.1_F3H_Fagopyrum_tataricum       RLKKLAK.EQ....QSSDL..........QKAK........LDSKP..IQDIFA......
BAL43067.1_F3H_Ipomoea_coccinea          RMKKFSK.EQK...QQMIIKAA.......ADTN........LETKP..IDQILA......
AAB41102.1_F3H_Ipomoea_purpurea          RIKKFAK.EQ....QMIKAAAA.......ADTN........LETKP..IDQILA......
BAK78917.1_F3H_Ipomoea_quamoclit         RIKKFAK.EQK...QMIKAAAA.......ADTN........LETKP..IDQILA......
ADO95201.1_F3H_Litchi_chinensis          RLKKLAK.EQ....QQQDT..........EKSK........LEAKP..IEEILA......
AIB06738.1_F3H_Mangifera_indica          RLKKLAK.EQ....QLQES..........ETAK........LDAQP..IEKILA......
AEF14415.1_F3H_Onobrychis_viciifolia     RMKKIAK..EK...ELRDL..........EKAK........LEAKP..LNQILA......
AFI71897.1_F3H_Paeonia_lactiflora        RLKKLAK.E.....QMQDH.....L....EKTQ........LESKP..LEQILA......
AEN71544.1_F3H_Paeonia_suffruticosa      RLK...K.Q.....QMQDH.....L....EKTQ........LESKP..LDQILA......
CAA53579.1_F3H_Vitis_vinifera            RLKKLAK.EQ....QLQDV..........EKAK........LESKP..IDQILA......
AGS57503.1_F3H_Vitis_rotundifolia        RLKKLAK.EL....QLQDV..........EKAK........LESKP..IDQILA......
ACM17897.1_F3H_Rubus_occidentalis        RLKKLAK.EQ....QPEDS..........EKAK........LEVKQ..VDDIFA......
ABW74548.1_F3H_Rubus_coreanus            RLKKLAK.EQ....QPEDS..........EKAK........LEVKQ..VDDIFA......
ADP09378.1_F3H_Pyrus_pyrifolia           RLKKLAK.EQ....ELQDL..........EKDK........VEIKP..ANDIFA......
AGL50918.1_F3H_Pyrus_communis            RLKKLAK.EQ....QLQDL..........ETAK........VDTKP..VDDIFA......
ACP30361.1_F3H_Malus_hybrid_cultivar     RLKKLAK.EQ....QSQDL..........EKAK........VDTKP..VDDIFA......
ABG78792.1_F3H_Aethusa_cynapium          KLKKLAK.EKLL..QDQDT..........EKAK........LQTKPKSADEIFA......
AAX21540.1_F3H_Anethum_graveolens        KLKKLAK.DKLL..QEQDA..........DKAK........LQTTPKTADEIFA......
AAX21539.1_F3H_Ammi_majus                KLKKLAK.EKLL..QEQEA..........EKAK........LQMTPKSADEIFA......
AAU04791.1_F3H_Fragaria_x_ananassa       RLKKHAK.EQ....L.QDS..........EKTK........LEAKP..VDDIFA......
AAU93347.1_F3H_Ginkgo_biloba             RQKKLAKLQQ...............................DESK...............
AAT94365.1_F3H_Glycine_max               RMKKLAK.EK....HLQDLENEKHLQELDQKAK........LEAKP..LKEILA......
AAX21535.1_F3H_Pimpinella_anisum         KLKKLAK.EKVL..QEQEA..........EKAK........LQMTPKSADEIFA......
AAP57394.1_F3H_Petroselinum_crispum      TLKKLAK.EKVL..QDQEV..........EKAK........LQMTPKSADEIFA......
AAC49929.1_F3H_Petunia_x_hybrida         RLKKQAK.EQ....QLQAEVAA.......EKAK........LESKP..IEEILA......
                                     acc

NP_190692.1_F3H_Arabidopsis_thaliana

NP_190692.1_F3H_Arabidopsis_thaliana     ............................................................
TRINITY_DN33424_c3_g3_i1                 ............................................................
AAU04792.1_F3H_Fragaria_x_ananassa       ............................................................
NP_001236797.1_F3H_Glycine_max           ............................................................
ACR15123.1_F3H_Medicago_truncatula       ............................................................
AQX36284.1_F3H_Prunus_persica            ............................................................
ABR13013.1_F3H_Triticum_aestivum         ............................................................
NP_001130275.1_F3H_Zea_mays              ............................................................
ACL54955.1_F3H_Actinidia_chinensis       ............................................................
AFN70721.1_F3H_Nekemias_grossedentata    ............................................................
ACM62745.1_F3H_Garcinia_mangostana       ............................................................
AEO36935.1_F3H_Canarium_album            ............................................................
AEG64806.1_F3H_Carthamus_tinctorius      ............................................................
AFC37250.1_F3H_Camellia_chekiangoleosa   ............................................................
ACB56921.1_F3H_Pilosella_officinarum     ............................................................
AAB97310.1_F3H_Chrysanthemum_x_morifolium ............................................................
AAC15414.1_F3H_Nicotiana_tabacum         SLCHPVSVCSCQILLVEINLAIVCRNIVAYNPDFEKRILEAAANSNGTYILEMDDNSLHE
ABL86673.1_F3H_Gossypium_barbadense      ............................................................
ADC96713.1_F3H_Gossypium_hirsutum        ............................................................
XP_007046698.1_F3H_Theobroma_cacao       ............................................................
ABO48521.1_F3H_Dimocarpus_longan         ............................................................
ACQ99190.1_F3H_Fagopyrum_tataricum       ............................................................
BAL43067.1_F3H_Ipomoea_coccinea          ............................................................
AAB41102.1_F3H_Ipomoea_purpurea          ............................................................
BAK78917.1_F3H_Ipomoea_quamoclit         ............................................................
ADO95201.1_F3H_Litchi_chinensis          ............................................................
AIB06738.1_F3H_Mangifera_indica          ............................................................
AEF14415.1_F3H_Onobrychis_viciifolia     ............................................................
AFI71897.1_F3H_Paeonia_lactiflora        ............................................................
AEN71544.1_F3H_Paeonia_suffruticosa      ............................................................
CAA53579.1_F3H_Vitis_vinifera            ............................................................
AGS57503.1_F3H_Vitis_rotundifolia        ............................................................
ACM17897.1_F3H_Rubus_occidentalis        ............................................................
ABW74548.1_F3H_Rubus_coreanus            ............................................................
ADP09378.1_F3H_Pyrus_pyrifolia           ............................................................
AGL50918.1_F3H_Pyrus_communis            ............................................................
ACP30361.1_F3H_Malus_hybrid_cultivar     ............................................................
ABG78792.1_F3H_Aethusa_cynapium          ............................................................
AAX21540.1_F3H_Anethum_graveolens        ............................................................
AAX21539.1_F3H_Ammi_majus                ............................................................
AAU04791.1_F3H_Fragaria_x_ananassa       ............................................................
AAU93347.1_F3H_Ginkgo_biloba             ............................................................
AAT94365.1_F3H_Glycine_max               ............................................................
AAX21535.1_F3H_Pimpinella_anisum         ............................................................
AAP57394.1_F3H_Petroselinum_crispum      ............................................................
AAC49929.1_F3H_Petunia_x_hybrida         ............................................................
                                     acc
```

*NP_190692.1_F3H_Arabidopsis_thaliana*

```
NP_190692.1_F3H_Arabidopsis_thaliana              ............................................................
TRINITY_DN33424_c3_g3_i1                          ............................................................
AAU04792.1_F3H_Fragaria_x_ananassa                ............................................................
NP_001236797.1_F3H_Glycine_max                    ............................................................
ACR15123.1_F3H_Medicago_truncatula                ............................................................
AQX36284.1_F3H_Prunus_persica                     ............................................................
ABR13013.1_F3H_Triticum_aestivum                  ............................................................
NP_001130275.1_F3H_Zea_mays                       ............................................................
ACL54955.1_F3H_Actinidia_chinensis                ............................................................
AFN70721.1_F3H_Nekemias_grossedentata             ............................................................
ACM62745.1_F3H_Garcinia_mangostana                ............................................................
AEO36935.1_F3H_Canarium_album                     ............................................................
AEG64806.1_F3H_Carthamus_tinctorius               ............................................................
AFC37250.1_F3H_Camellia_chekiangoleosa            ............................................................
ACB56921.1_F3H_Pilosella_officinarum              ............................................................
AAB97310.1_F3H_Chrysanthemum_x_morifolium         ............................................................
AAC15414.1_F3H_Nicotiana_tabacum                  NRFVRDEDERPKVPYNKFSSEIPVISLKGIDDVEESSGGIKSRRAEICEKIIACEDWGIF
ABL86673.1_F3H_Gossypium_barbadense               ............................................................
ADC96713.1_F3H_Gossypium_hirsutum                 ............................................................
XP_007046698.1_F3H_Theobroma_cacao                ............................................................
ABO48521.1_F3H_Dimocarpus_longan                  ............................................................
ACQ99190.1_F3H_Fagopyrum_tataricum                ............................................................
BAL43067.1_F3H_Ipomoea_coccinea                   ............................................................
AAB41102.1_F3H_Ipomoea_purpurea                   ............................................................
BAK78917.1_F3H_Ipomoea_quamoclit                  ............................................................
ADO95201.1_F3H_Litchi_chinensis                   ............................................................
AIB06738.1_F3H_Mangifera_indica                   ............................................................
AEF14415.1_F3H_Onobrychis_viciifolia              ............................................................
AFI71897.1_F3H_Paeonia_lactiflora                 ............................................................
AEN71544.1_F3H_Paeonia_suffruticosa               ............................................................
CAA53579.1_F3H_Vitis_vinifera                     ............................................................
AGS57503.1_F3H_Vitis_rotundifolia                 ............................................................
ACM17897.1_F3H_Rubus_occidentalis                 ............................................................
ABW74548.1_F3H_Rubus_coreanus                     ............................................................
ADP09378.1_F3H_Pyrus_pyrifolia                    ............................................................
AGL50918.1_F3H_Pyrus_communis                     ............................................................
ACP30361.1_F3H_Malus_hybrid_cultivar              ............................................................
ABG78792.1_F3H_Aethusa_cynapium                   ............................................................
AAX21540.1_F3H_Anethum_graveolens                 ............................................................
AAX21539.1_F3H_Ammi_majus                         ............................................................
AAU04791.1_F3H_Fragaria_x_ananassa                ............................................................
AAU93347.1_F3H_Ginkgo_biloba                      ............................................................
AAT94365.1_F3H_Glycine_max                        ............................................................
AAX21535.1_F3H_Pimpinella_anisum                  ............................................................
AAP57394.1_F3H_Petroselinum_crispum               ............................................................
AAC49929.1_F3H_Petunia_x_hybrida                  ............................................................
                                           acc
```

*NP_190692.1_F3H_Arabidopsis_thaliana*

```
NP_190692.1_F3H_Arabidopsis_thaliana              ............................................................
TRINITY_DN33424_c3_g3_i1                          ............................................................
AAU04792.1_F3H_Fragaria_x_ananassa                ............................................................
NP_001236797.1_F3H_Glycine_max                    ............................................................
ACR15123.1_F3H_Medicago_truncatula                ............................................................
AQX36284.1_F3H_Prunus_persica                     ............................................................
ABR13013.1_F3H_Triticum_aestivum                  ............................................................
NP_001130275.1_F3H_Zea_mays                       ............................................................
ACL54955.1_F3H_Actinidia_chinensis                ............................................................
AFN70721.1_F3H_Nekemias_grossedentata             ............................................................
ACM62745.1_F3H_Garcinia_mangostana                ............................................................
AEO36935.1_F3H_Canarium_album                     ............................................................
AEG64806.1_F3H_Carthamus_tinctorius               ............................................................
AFC37250.1_F3H_Camellia_chekiangoleosa            ............................................................
ACB56921.1_F3H_Pilosella_officinarum              ............................................................
AAB97310.1_F3H_Chrysanthemum_x_morifolium         ............................................................
AAC15414.1_F3H_Nicotiana_tabacum                  QVVDHGVDTKMLSEMTRLAREFFEMSAEEKLRFDMTGGKKGGFIVSSHLQGEAVQDWREI
ABL86673.1_F3H_Gossypium_barbadense               ............................................................
ADC96713.1_F3H_Gossypium_hirsutum                 ............................................................
XP_007046698.1_F3H_Theobroma_cacao                ............................................................
ABO48521.1_F3H_Dimocarpus_longan                  ............................................................
ACQ99190.1_F3H_Fagopyrum_tataricum                ............................................................
BAL43067.1_F3H_Ipomoea_coccinea                   ............................................................
AAB41102.1_F3H_Ipomoea_purpurea                   ............................................................
BAK78917.1_F3H_Ipomoea_quamoclit                  ............................................................
ADO95201.1_F3H_Litchi_chinensis                   ............................................................
AIB06738.1_F3H_Mangifera_indica                   ............................................................
AEF14415.1_F3H_Onobrychis_viciifolia              ............................................................
AFI71897.1_F3H_Paeonia_lactiflora                 ............................................................
AEN71544.1_F3H_Paeonia_suffruticosa               ............................................................
CAA53579.1_F3H_Vitis_vinifera                     ............................................................
AGS57503.1_F3H_Vitis_rotundifolia                 ............................................................
ACM17897.1_F3H_Rubus_occidentalis                 ............................................................
ABW74548.1_F3H_Rubus_coreanus                     ............................................................
ADP09378.1_F3H_Pyrus_pyrifolia                    ............................................................
AGL50918.1_F3H_Pyrus_communis                     ............................................................
ACP30361.1_F3H_Malus_hybrid_cultivar              ............................................................
ABG78792.1_F3H_Aethusa_cynapium                   ............................................................
AAX21540.1_F3H_Anethum_graveolens                 ............................................................
AAX21539.1_F3H_Ammi_majus                         ............................................................
AAU04791.1_F3H_Fragaria_x_ananassa                ............................................................
AAU93347.1_F3H_Ginkgo_biloba                      ............................................................
AAT94365.1_F3H_Glycine_max                        ............................................................
AAX21535.1_F3H_Pimpinella_anisum                  ............................................................
AAP57394.1_F3H_Petroselinum_crispum               ............................................................
AAC49929.1_F3H_Petunia_x_hybrida                  ............................................................
                                           acc
```

*NP_190692.1_F3H_Arabidopsis_thaliana*

```
NP_190692.1_F3H_Arabidopsis_thaliana        ............................................................
TRINITY_DN33424_c3_g3_i1                    ............................................................
AAU04792.1_F3H_Fragaria_x_ananassa          ............................................................
NP_001236797.1_F3H_Glycine_max              ............................................................
ACR15123.1_F3H_Medicago_truncatula          ............................................................
AQX36284.1_F3H_Prunus_persica               ............................................................
ABR13013.1_F3H_Triticum_aestivum            ............................................................
NP_001130275.1_F3H_Zea_mays                 ............................................................
ACL54955.1_F3H_Actinidia_chinensis          ............................................................
AFN70721.1_F3H_Nekemias_grossedentata       ............................................................
ACM62745.1_F3H_Garcinia_mangostana          ............................................................
AEO36935.1_F3H_Canarium_album               ............................................................
AEG64806.1_F3H_Carthamus_tinctorius         ............................................................
AFC37250.1_F3H_Camellia_chekiangoleosa      ............................................................
ACB56921.1_F3H_Pilosella_officinarum        ............................................................
AAB97310.1_F3H_Chrysanthemum_x_morifolium   ............................................................
AAC15414.1_F3H_Nicotiana_tabacum            VTYFSYPIKARDYSRWPDKPKEWRAVTEKYSEDLMGLGCKLLEMLSEAMGLEKEALKDAC
ABL86673.1_F3H_Gossypium_barbadense         ............................................................
ADC96713.1_F3H_Gossypium_hirsutum           ............................................................
XP_007046698.1_F3H_Theobroma_cacao          ............................................................
ABO48521.1_F3H_Dimocarpus_longan            ............................................................
ACQ99190.1_F3H_Fagopyrum_tataricum          ............................................................
BAL43067.1_F3H_Ipomoea_coccinea             ............................................................
AAB41102.1_F3H_Ipomoea_purpurea             ............................................................
BAK78917.1_F3H_Ipomoea_quamoclit            ............................................................
ADO95201.1_F3H_Litchi_chinensis             ............................................................
AIB06738.1_F3H_Mangifera_indica             ............................................................
AEF14415.1_F3H_Onobrychis_viciifolia        ............................................................
AFI71897.1_F3H_Paeonia_lactiflora           ............................................................
AEN71544.1_F3H_Paeonia_suffruticosa         ............................................................
CAA53579.1_F3H_Vitis_vinifera               ............................................................
AGS57503.1_F3H_Vitis_rotundifolia           ............................................................
ACM17897.1_F3H_Rubus_occidentalis           ............................................................
ABW74548.1_F3H_Rubus_coreanus               ............................................................
ADP09378.1_F3H_Pyrus_pyrifolia              ............................................................
AGL50918.1_F3H_Pyrus_communis               ............................................................
ACP30361.1_F3H_Malus_hybrid_cultivar        ............................................................
ABG78792.1_F3H_Aethusa_cynapium             ............................................................
AAX21540.1_F3H_Anethum_graveolens           ............................................................
AAX21539.1_F3H_Ammi_majus                   ............................................................
AAU04791.1_F3H_Fragaria_x_ananassa          ............................................................
AAU93347.1_F3H_Ginkgo_biloba                ............................................................
AAT94365.1_F3H_Glycine_max                  ............................................................
AAX21535.1_F3H_Pimpinella_anisum            ............................................................
AAP57394.1_F3H_Petroselinum_crispum         ............................................................
AAC49929.1_F3H_Petunia_x_hybrida            ............................................................
                                      acc
```

*NP_190692.1_F3H_Arabidopsis_thaliana*

```
NP_190692.1_F3H_Arabidopsis_thaliana        ..................
TRINITY_DN33424_c3_g3_i1                    ..................
AAU04792.1_F3H_Fragaria_x_ananassa          ..................
NP_001236797.1_F3H_Glycine_max              ..................
ACR15123.1_F3H_Medicago_truncatula          ..................
AQX36284.1_F3H_Prunus_persica               ..................
ABR13013.1_F3H_Triticum_aestivum            ..................
NP_001130275.1_F3H_Zea_mays                 ..................
ACL54955.1_F3H_Actinidia_chinensis          ..................
AFN70721.1_F3H_Nekemias_grossedentata       ..................
ACM62745.1_F3H_Garcinia_mangostana          ..................
AEO36935.1_F3H_Canarium_album               ..................
AEG64806.1_F3H_Carthamus_tinctorius         ..................
AFC37250.1_F3H_Camellia_chekiangoleosa      ..................
ACB56921.1_F3H_Pilosella_officinarum        ..................
AAB97310.1_F3H_Chrysanthemum_x_morifolium   ..................
AAC15414.1_F3H_Nicotiana_tabacum            VDMDQKVVLNSYPKCPQP
ABL86673.1_F3H_Gossypium_barbadense         ..................
ADC96713.1_F3H_Gossypium_hirsutum           ..................
XP_007046698.1_F3H_Theobroma_cacao          ..................
ABO48521.1_F3H_Dimocarpus_longan            ..................
ACQ99190.1_F3H_Fagopyrum_tataricum          ..................
BAL43067.1_F3H_Ipomoea_coccinea             ..................
AAB41102.1_F3H_Ipomoea_purpurea             ..................
BAK78917.1_F3H_Ipomoea_quamoclit            ..................
ADO95201.1_F3H_Litchi_chinensis             ..................
AIB06738.1_F3H_Mangifera_indica             ..................
AEF14415.1_F3H_Onobrychis_viciifolia        ..................
AFI71897.1_F3H_Paeonia_lactiflora           ..................
AEN71544.1_F3H_Paeonia_suffruticosa         ..................
CAA53579.1_F3H_Vitis_vinifera               ..................
AGS57503.1_F3H_Vitis_rotundifolia           ..................
ACM17897.1_F3H_Rubus_occidentalis           ..................
ABW74548.1_F3H_Rubus_coreanus               ..................
ADP09378.1_F3H_Pyrus_pyrifolia              ..................
AGL50918.1_F3H_Pyrus_communis               ..................
ACP30361.1_F3H_Malus_hybrid_cultivar        ..................
ABG78792.1_F3H_Aethusa_cynapium             ..................
AAX21540.1_F3H_Anethum_graveolens           ..................
AAX21539.1_F3H_Ammi_majus                   ..................
AAU04791.1_F3H_Fragaria_x_ananassa          ..................
AAU93347.1_F3H_Ginkgo_biloba                ..................
AAT94365.1_F3H_Glycine_max                  ..................
AAX21535.1_F3H_Pimpinella_anisum            ..................
AAP57394.1_F3H_Petroselinum_crispum         ..................
AAC49929.1_F3H_Petunia_x_hybrida            ..................
                                      acc
```

```
Q9SD85.1_F3-H_Arabidopsis_thaliana                    TT    TT      TT                    TT                η1
                                                    1         10        20        30        40

Q9SD85.1_F3-H_Arabidopsis_thaliana          ......MATLFLTILLATVLFLIL...RIFSHRRNR..SHNNRLPPGPNPWPIIGNLPHM
TRINITY_DN252_c0_g1_i1                      ...MSSFLLYF..ILFFISIYFL....SSLRSFLR.....RRLPPGPKPWPIVGNLPHL
NP_001312537.1_F3-H_Nicotiana_tabacum       ...MAIFSLILYTVIFSFLLHSIF...SLFFRKRYP.....LTLPPGPKPWPIIGNLVHM
Q9SBQ9.1_F3-H_Petunia_x_hybrida             ...MEILSLILYTVIFSFLLQFIL...RSFFRKRYP.....LPLPPGPKPWPIIGNLVHL
AAG49298.1_F3-H_Callistephus_chinensis      ...MTILPFIFYTCITALVLYVL....LNLLTRNP.....NRLPPGPTPWPIVGNLPHL
BAD91808.1_F3-H_Gentiana_triflora           .MEMNPSPFIVHTAVFGLFVVFIF..IFLKLVSRKKGHGRSLPLPPGPRPWPILGNIPHL
ABA64468.1_F3-H_Gerbera_hybrid_cultivar     ...MTPLTLLIGTCVTGLFLYVL....LNRCTRNP.....NRLPPGPTPWPVVGNLPHL
AAO47847.1_F3-H_Glycine_max                 ...MSPLIVALATIAAAILIYRI....IKFITRPS.....LPLPPGPKPWPIVGNLPHM
ABC47161.1_F3-H_Pilosella_officinarum       ...MTLLSLIIYLCITGVTAYVL....VNLRTRRA.....NRLPPGPTPWPIVGNLPHL
BAD00190.1_F3-H_Ipomoea_nil                 ...MATLTLIFCTLVFAIFLYFLI...LRVKQRYP.....LPLPPGPKPWPVLGNLPHL
BAF49324.1_F3-H_Lobelia_erinus              ......MSLIFYTCIIAFLLYVLLSTLRTFLLNRNPA..SRKPLPPGPKPWPIVGNLPHL
AAG49301.1_F3-H_Matthiola_incana            ......MTTLILTILLATFLSLFI...FFLLRRNR..NRNHRLPPGPNPWPIVGNLPHM
ABB29899.1_F3-H_Osteospermum_hybrid_cultivar ...MTILPLVLYSCITGLVIYVL....LNLRTRHS.....NRLPPGPTPWPIVGNLPHL
AAG49315.1_F3-H_Pelargonium_x_hortorum      MYNMS.LYLLLGSSALAFAAYLV....LFSFSKSR.....RRLPPGPKAWPIVGNLPHM
BAB59005.1_F3-H_Perilla_frutescens_var._crispa .MISAAVSLIICTSILGVLVYFLF.......LRRGGG.SNGRPLPPGPRPWPIVGNLPQL
BAE47006.1_F3-H_Vitis_vinifera              ...MNPLALFFCTALFCVLLYHF....LTRRSV........RLPPGPKPWPIVGNLPHL
AAR00229.1_F3-H_Ipomoea_purpurea            ...MATLTLIFCTLVFAIFLYFLI...LRVKQRYP.....LPLPPGPKPWPVLGNLPHL
                                       acc
```

```
Q9SD85.1_F3-H_Arabidopsis_thaliana            α1          β1   TT β2        α2        α3        η2
                                            50        60        70        80        90        100

Q9SD85.1_F3-H_Arabidopsis_thaliana          GTKPHRTLSAMVTTYGPILHLRLGFVDVVVAASKSVAEQFLKIHDANFASRPPNSGAKHM
TRINITY_DN252_c0_g1_i1                      GPMPHHSIAALARTYGPLMHLRMGFVDVVVAASAAVASQFLKFHDANFSSRPPNSGAKYV
NP_001312537.1_F3-H_Nicotiana_tabacum       GPKPHQSTAAMARTYGPLMHLKMGFVDVVVASASVAAQFLKTHDANFSSRPPNSGAKHL
Q9SBQ9.1_F3-H_Petunia_x_hybrida             GPKPHQSTAAMAQTYGPLMYLKMGFVDVVVAASASVAAQFLKTHDANFSSRPPNSGAEHM
AAG49298.1_F3-H_Callistephus_chinensis      GMIPHHSLAALAQKYGPLMHLRLGFVDVVVAASASVAAQFLKTHDANFASRPPNSGAKHI
BAD91808.1_F3-H_Gentiana_triflora           GSKPHQTLAEMAKTYGPLMHLKFGLKDAVVASSASVAEQFLKKHDVNFSNRPPNSGAKHI
ABA64468.1_F3-H_Gerbera_hybrid_cultivar     GTIPHHSLAAMAKKYGPLMHLRLGFVDVVVAASASVAAQFLKTHDANFADRPPNSGAKHI
AAO47847.1_F3-H_Glycine_max                 GPVPHHSLAALARIHGPLMHLRLGFVDVVVAASASVAEQFLKIHDSNFSSRPPNAGAKYI
ABC47161.1_F3-H_Pilosella_officinarum       GTIPHHSLADLATRYGPLMHLRLGFVDVVVAASASVAAQFLKTHDANFASRPPNSGAKHM
BAD00190.1_F3-H_Ipomoea_nil                 GKKPHQSIAAMAERYGPLMHLRLGFVDVVVAASAAVAAQFLKVHDSNFSNRPPNSGAEHI
BAF49324.1_F3-H_Lobelia_erinus              GTMPHQTMAALARTYGPLMHLRLGSVDVVVAASGAVAEQFLKVHDANFLNRPPNSGAVHI
AAG49301.1_F3-H_Matthiola_incana            GPKPHQTLAAMVTTYGPILHLRLGFVNVVVAASKSVAEQFLKIHDANFASRPPNSGAKHI
ABB29899.1_F3-H_Osteospermum_hybrid_cultivar GVVPHHSLAAMAEKYGPLMHLRLGFVDVVVAASAAVAAQFLKVHDANFASRPPNSGAKHI
AAG49315.1_F3-H_Pelargonium_x_hortorum      GSMPHQNLAAMARTYGPLVYLRLGFVDVVVALSASMASQFLKTHDSNFSSRPPNAGAKHI
BAB59005.1_F3-H_Perilla_frutescens_var._crispa GPKPHQSMAALARVHGPLMHLKMGFVHVVVAASATVAEKFLKVHDTNFLSRPPNSGAEHI
BAE47006.1_F3-H_Vitis_vinifera              GPVPHHSIAALAKTYGPLMHLRMGFVDVVVAASASVAAQFLKTHDANFSNRPPNSGAKHI
AAR00229.1_F3-H_Ipomoea_purpurea            GEKPHQSIAAMAERHGPLMHLRMGFVDVVVAASAAVAAQFLKVHDANFSNRPPNSGAEHI
                                       acc
```

```
Q9SD85.1_F3-H_Arabidopsis_thaliana                            α4                 α5                β3
                                            110       120       130       140       150       160

Q9SD85.1_F3-H_Arabidopsis_thaliana          AYNYQDLVFAP.YGHRWRLLRKISSVHLFSAKALEDFKHVRQEEVGTLTRELVRVGTKPV
TRINITY_DN252_c0_g1_i1                      AYNYQDLVFAP.YGPRWRMLRKISSVNLFSGKALDDFRHIRQHEVSVLTRALASAGSTPV
NP_001312537.1_F3-H_Nicotiana_tabacum       AYNYQDLVFAP.YGPRWRMLRKICSVHLFSAKALDDFSHVRQDEVRTLTRALASAAQKPV
Q9SBQ9.1_F3-H_Petunia_x_hybrida             AYNYQDLVFAP.YGPRWRMLRKICSVHLFSTKALDDFRHVRQDEVKTLTRALASAGQKPV
AAG49298.1_F3-H_Callistephus_chinensis      AYNYQDLVFAP.YGPRWRMLRKICSVHLFSTKALDDFRHVREEEVAILTRVLVHAGESAV
BAD91808.1_F3-H_Gentiana_triflora           AYNYQDLVFAP.YGPRWRLLRKICSVHLFSSKALDDFQHVRHEEICILIRAIASGGHAPV
ABA64468.1_F3-H_Gerbera_hybrid_cultivar     AYNYQDLVFAP.YGPRWRMLRKICSVHLFSTKALDDFRHVRQEEVAILARALVGAGKSPV
AAO47847.1_F3-H_Glycine_max                 AYNYQDLVFAP.YGPRWRLLRKLTSVHLFSGKAMNEFRHLRQEEVARLTCNLASSDTKAV
ABC47161.1_F3-H_Pilosella_officinarum       AYNYQDLVFAP.YGPRWRMLRKICSVHLFSAKSLDDFRHVRQEEVAILTRALVGAGKSTV
BAD00190.1_F3-H_Ipomoea_nil                 AYNYQDLVFAP.YGPRWRMLRKITSVHLFSAKALDDFCHVRQEEVATLTRSLASAGKTPV
BAF49324.1_F3-H_Lobelia_erinus              AYNYQDLVFAP.YGPRWRLLRKVSAVHLFSAKALDDFRHVRQEEVAVLTRDLASAGNAPV
AAG49301.1_F3-H_Matthiola_incana            AYNYQDLVFAP.YGQRWRMLRKISSVHLFSAKALEDFKHVRQEEIGRLTREVARADTKPV
ABB29899.1_F3-H_Osteospermum_hybrid_cultivar AYNYQDLVFAPYYGPRWRMLRKICSVHLFSSKALDDFRHVRQEEVAILTRALIGAGDSPV
AAG49315.1_F3-H_Pelargonium_x_hortorum      AYNYHDLVFAP.YGPRWRLFRKITSIHLFSGKALDDYRHVRQEEVGVLASNLARAVSTIV
BAB59005.1_F3-H_Perilla_frutescens_var._crispa AYNYNDLVFAP.HGPRWRLLRKICALHLFSSKALDDFRHVREEEVGILIRNLASVGEMPA
BAE47006.1_F3-H_Vitis_vinifera              AYNYQDLVFAP.YGPRWRMLRKICSVHLFSGKALDDFRHIRQEEVAVLTRALARAGQTPV
AAR00229.1_F3-H_Ipomoea_purpurea            AYNYQDLVFAP.YGPRWRMLRKITSVHLFSAKALDDFCHVRQEEVATLTRSLASAGKTPV
                                       acc
```

```
Q9SD85.1_F3-H_Arabidopsis_thaliana                α6                                α7              η3
                                            170       180       190       200       210       220

Q9SD85.1_F3-H_Arabidopsis_thaliana          NLGQLVNMCVVNALGREMIGRRLFGADA.......DHKADEFRSMVTEMMALAGVFNIGD
TRINITY_DN252_c0_g1_i1                      NLGQLLNVCTANALGRVMIGRRVFGDGSGGG....DRKADEFKTMVVELMVLAGVFNIGD
NP_001312537.1_F3-H_Nicotiana_tabacum       KLGQLLNVCTTNALAQVMLGRRVFADANGGV....DPQAEEFKSMVVEAMVLAGVFNVGD
Q9SBQ9.1_F3-H_Petunia_x_hybrida             KLGQLLNVCTTNALARVMLGKRVFADGSGDV....DPQAAEFKSMVVEMMVVAGVFNIGD
AAG49298.1_F3-H_Callistephus_chinensis      KLGQLLNVCTTNALARVMLGRRVFADGSEGRG..VDPKADEFKDMVVELMELAGEFNIGD
BAD91808.1_F3-H_Gentiana_triflora           NLGKLLGVCTTNALARVMLGRRVF.EGDGGE....NPHADEFKSMVVEIMVLAGAFNLGD
ABA64468.1_F3-H_Gerbera_hybrid_cultivar     KLGQLLNVCTTNALARVMLGRRVFDSG........DAQADEFKDMVVELMVLAGEFNIGD
AAO47847.1_F3-H_Glycine_max                 NLGQLLNVCTTNALARAMIGRRVFNDGNGGC....DPRADEFKAMVMEVMVLAGVFNIGD
ABC47161.1_F3-H_Pilosella_officinarum       KLGQLLHVCTTNALVRVMLGRRVFGDGSGGG....DPKADEFKNMVIEMMVLAGEFNLGD
BAD00190.1_F3-H_Ipomoea_nil                 KLGQLLNVCTTNALARVMLGRKVFNDGGSKS....DPKAEEFKSMVEEMMVLAGSFNIGD
BAF49324.1_F3-H_Lobelia_erinus              HLNQLLSVCTTNALSRVMFGKRVFGGGG.......DSKADEFKQMVVEAMVLAGVFNIGD
AAG49301.1_F3-H_Matthiola_incana            NLGQLVNMCVVNALGREMIGRRLFGDGA.......DHKAEEFRSMVTEMMALAGVFNVGD
ABB29899.1_F3-H_Osteospermum_hybrid_cultivar KLGQLLNVCTTNALARVMLGKRVFGDRSGGG....DPKADEFKDMVVEVMELAGEFNIGD
AAG49315.1_F3-H_Pelargonium_x_hortorum      NLGQLLNICATNALGRAVIGKKVFKDGTDDV....DPKADEFKSMVVELMVLAGVFNIGD
BAB59005.1_F3-H_Perilla_frutescens_var._crispa SIGQMMYVCATNAISRVMLGRHVLGDEHRGAAGGGDTTAEEFKAMVVELMALAGVFNVGD
BAE47006.1_F3-H_Vitis_vinifera              NLGQLLNVCTTNALGRVMLGRRVFGDGSGGE....DPKADEFKEMVVELMVLAGVFNIGD
AAR00229.1_F3-H_Ipomoea_purpurea            KLGQLLNVCTTNALARVMLGRKVFNDGGSKS....DPKAEEFKSMVEEMMVLAGSFNIGD
                                       acc
```

```
                                                          α8                    α9                                 α10
Q9SD85.1_F3-H_Arabidopsis_thaliana                                    230       240       250       260       270
Q9SD85.1_F3-H_Arabidopsis_thaliana                  FVPSLDWLDLQGVAGKMKRLHKRFDAFLSSILKEHE....M.NGQDQKHTDMLSTLISLK
TRINITY_DN252_c0_g1_i1                              FVPALEWLDLQGVAAKMKKLHKRFDAFLSEIVEDHK.MNLKGSDGSEKHTDMLSTLIALE
NP_001312537.1_F3-H_Nicotiana_tabacum               FIPALDWLDIQGVAAKMKKLHARFDAFLTSILEEHK..S.NQFGETKEHEDLLSTLISLK
Q9SBQ9.1_F3-H_Petunia_x_hybrida                     FIPQLNWLDIQGVAAKMKKLHARFDAFLTDILEEHK..G.KIFGEMK...DLLSTLISLK
AAG49298.1_F3-H_Callistephus_chinensis              FIPPLDCLDLQGITKKMKKLHARFDKFLNIILDDHK.IE.KG.AAGRRHSDLLTTLISLK
BAD91808.1_F3-H_Gentiana_triflora                   FIPVLDWFDLQGIAGKMKKLHARFDKFLNGILEDRK..S.NGSNGAEQYVDLLSVLISLQ
ABA64468.1_F3-H_Gerbera_hybrid_cultivar             FIPVLDWLDLQGVTKKMKKLHAKFDSFLNTILEEHK.TG.AGDGVASGKVDLLSTLISLK
AAO47847.1_F3-H_Glycine_max                         FIPSLEWLDLQGVQAKMKKLHKRFDAFLTSIIEEHN....NSSSKNENHKNFLSILLSLK
ABC47161.1_F3-H_Pilosella_officinarum               FIPVLDLLDLQGVTKKMKKLHTRFDSFLNSILEEHR.TS.SG.GASG.HVDLLSTLISLK
BAD00190.1_F3-H_Ipomoea_nil                         FIPVLGWFDVQGIVGKMKKLHARFDAFLNTILEEHKCVN.NQHTTLSKDVDFLSTLIRLK
BAF49324.1_F3-H_Lobelia_erinus                      FIPALEWFDLQGVVGKMKKLHVRFDSFLTAILEEHK.TT..GSRGGGKHVDLLSTLISLK
AAG49301.1_F3-H_Matthiola_incana                    FVPALDWLDLQGVAGKMKRLHKRFDAFLSSILKEHE....INNGGDQKHTDMLTTLISLK
ABB29899.1_F3-H_Osteospermum_hybrid_cultivar        FIPVLDSLDLQGIAKKMKELHVRFDSFLGKILEEHK.TG.NG.GASSQHTDLLTTLISLK
AAG49315.1_F3-H_Pelargonium_x_hortorum              FIPPLDCLDLQGVASKMKNLHKRFDAFLSAILQEHN.IN....SAASATPSMLTTLISLK
BAB59005.1_F3-H_Perilla_frutescens_var._crispa      FIPPLKGLDLQGVVAKMKKLHQRFDAFFSGILHDHK.IN..GSNAAEGHVDLLTTLISLK
BAE47006.1_F3-H_Vitis_vinifera                      FVPALEWLDLQGVAAKMKKLHARFDAFLGAIVEEHK.IS..GSAGSERHVDLLSTLISVR
AAR00229.1_F3-H_Ipomoea_purpurea                    FIPILGWFDLQGIVGKMKKLHARFDAFLNSILEEHKFVN.NQHTTLSKDVDFLTTLIRLK
acc

                                                                            α11                            α12
Q9SD85.1_F3-H_Arabidopsis_thaliana                      280       290       300       310       320       330
Q9SD85.1_F3-H_Arabidopsis_thaliana                  GT..DLDGDGGSLTDTEIKALLLNMFTAGTDTSASTVDWAIAELIRHPDIMVKAQEELDI
TRINITY_DN252_c0_g1_i1                              E...DADGEGGKLTDTEIKALLLNMFTAGTDTSSSTVEWAIAELIRHPKILSKVKQELDS
NP_001312537.1_F3-H_Nicotiana_tabacum               KE..EGDNEGGKLTDSEIKALLLNLFTAGTDTSSSTVEWAVAELIRNPRILAQAQHEIDK
Q9SBQ9.1_F3-H_Petunia_x_hybrida                     ND..DADNDGGKLTDTEIKALLLNLFVAGTDTSSSTVEWAIAELIRNPKILAQAQQEIDK
AAG49298.1_F3-H_Callistephus_chinensis              DVDAADDDEEGKLSDIEIKALLLNLFAAGTDTSSSTVEWAVAELIRHPELLKQAREEMDI
BAD91808.1_F3-H_Gentiana_triflora                   DSNIDGGDEGTKLTDTEIKALLLNLFIAGTDTSSSTVEWAMAELIRNPKLLVQAQEELDR
ABA64468.1_F3-H_Gerbera_hybrid_cultivar             D...DADGEGGKLSDIEIKALLLNLFTAGTDTSSSTIEWAIAELIRNPQLLNQARKEMDT
AAO47847.1_F3-H_Glycine_max                         D...VRDDHGNHLTDTEIKALLLNMFTAGTDTSSSTTEWAVAELIKNPQILAKLQQELDT
ABC47161.1_F3-H_Pilosella_officinarum               D...EADGEGGKLTDTEIKALLLNLFVAGTDTSSSTVEWAIAELIRNPQLLKQAQQELDT
BAD00190.1_F3-H_Ipomoea_nil                         DNGADMDCEEGKLTDTEIKALLLNLFTAGTDTSSSTVEWAIAELLRNPKILNQAQQELDL
BAF49324.1_F3-H_Lobelia_erinus                      DN..DLAGEGEKLSDTEIKALLLNMFIAGTDTSSSTVEWALAELIRHPKILAQAQQELDE
AAG49301.1_F3-H_Matthiola_incana                    GT..DFDGDGASITDTEIKALLLNMFTAGTDTSASTVDWAIAELIRHPHIMKRTQEELDA
ABB29899.1_F3-H_Osteospermum_hybrid_cultivar        D...DTDEEGGKLSDIEIKALLLNLFTAGTDTSSSTVEWAIAELIRHPQLLKQAQEEIDN
AAG49315.1_F3-H_Pelargonium_x_hortorum              DS..VEDSEGGKLTDTEIKALLLNMFTAGTDTTSSTVEWAIAELIRQPEILIRAQKEIDS
BAB59005.1_F3-H_Perilla_frutescens_var._crispa      DV..DNNGEGGKLTDTEIKALLLNLFTAGTDTTSSTVEWAITELIRNPNILARVRKELDL
BAE47006.1_F3-H_Vitis_vinifera                      D...NADGEGGKLTDVEIKALLLNLFTAGTDTSSSTVEWAIAELIRHPEMMAQAQQELDA
AAR00229.1_F3-H_Ipomoea_purpurea                    DNGADTDSEEGKLTDTEIKALLLNLFTAGTDTSSSTVEWAFAELLRNPKILNQAQQELDS
acc

                                                            η4        α13          TT    β4     β5  TT  β6   TT   β7
Q9SD85.1_F3-H_Arabidopsis_thaliana                     340       350       360       370       380       390
Q9SD85.1_F3-H_Arabidopsis_thaliana                  VVGRDRPVNESDIAQLPYLQAVIKENFRLHPPTPLSLPHIASESCEINGYHIPKGSTLLT
TRINITY_DN252_c0_g1_i1                              VVGRDRLVTELDLAQLTYFQAVVKEIFRLHPSTPLSLPRLAAESCEINGYHIPKGSTLLV
NP_001312537.1_F3-H_Nicotiana_tabacum               VVGKNRLVMESDLAQLTYLEAIVKETLRLHPSTPLSLPRIASESCEINGYFIPKGSTLLV
Q9SBQ9.1_F3-H_Petunia_x_hybrida                     VVGRDRLVGELDLAQLTYLEAIVKETFRLHPSTPLSLPRIASESCEINGYFIPKGSTLLL
AAG49298.1_F3-H_Callistephus_chinensis              VVGRDRLVTELDLSRLTFLQAIVKETFRLHPSTPLSLPRMASESCEVDGYYIPKGSTLLV
BAD91808.1_F3-H_Gentiana_triflora                   VVGPNRFVTESDLPQLTFLQAVIKETFRLHPSTPLSLPRMAAEDCEINGYYVSEGSTLLV
ABA64468.1_F3-H_Gerbera_hybrid_cultivar             IVGQDRLVTESDLGQLTFLQAIIKETFRLHPSTPLSLPRMALESCEVGGYYIPKGSTLLV
AAO47847.1_F3-H_Glycine_max                         VVGRDRSVKEEDLAHLPYLQAVIKETFRLHPSTPLSVPRAAAESCEIFGYHIPKGATLLV
ABC47161.1_F3-H_Pilosella_officinarum               VVGQGRLVNESDLSQLTFLQAIVKETFRLHPSTPLSLPRIASESCEINGYNIPKGSTLLV
BAD00190.1_F3-H_Ipomoea_nil                         VVGQNQLVTESDLTDLPFLQAIVKETFRLHPSTPLSLPRMGAQGCEINGYFIPKGATLLV
BAF49324.1_F3-H_Lobelia_erinus                      VVGRDRLVSESDLSQLTFFQAIIKETFRLHPSTPLSLPRMAAESCEINGYFIPKGSTVLV
AAG49301.1_F3-H_Matthiola_incana                    VVGRNRPINESDLSRLPYLQAVIKENFRLHPPTPLSLPHIAAESCEINGYHIPKGSTLLT
ABB29899.1_F3-H_Osteospermum_hybrid_cultivar        VVGRDHLVTELDLTQLPFLQAIVKETFRLHPSTPLSLPRIASESCEVNGYHIPKGSTLLV
AAG49315.1_F3-H_Pelargonium_x_hortorum              VVGRDRLVTELDLSKLPYLQAIVKETFRLHSSTPLSLPRIATQSCEINGYHIPKGATLLV
BAB59005.1_F3-H_Perilla_frutescens_var._crispa      IVGKDKLVKESDLGQLTYLQAVIKENFRLHPSTPLSLPRVAQESCEINGYYIPKDSTLLV
BAE47006.1_F3-H_Vitis_vinifera                      VVGRGRLVTDLDLPQLTYLQAIVKETFRLHPSTPLSLPRMAAESCEINGYHIPKNATLLV
AAR00229.1_F3-H_Ipomoea_purpurea                    VVGQNRLVTESDLTHLPFLQAIVKETFRLHPSTPLSLPRMGAQGCEINGYFIPKGATLLV
acc

                                                     α14    α15    TT     η5                          TT     TT
Q9SD85.1_F3-H_Arabidopsis_thaliana                      400       410       420       430       440       450
Q9SD85.1_F3-H_Arabidopsis_thaliana                  NIWAIARDPDQWSDPLAFKPERFLPGGEKSGVDVKGSDFELIPFGAGRRICAGLSLGLRT
TRINITY_DN252_c0_g1_i1                              NVWAIARDPDVWSDPLEFKPERFLPGGEKANVDVRGNDFELIPFGAGRRICAGMSLGLRM
NP_001312537.1_F3-H_Nicotiana_tabacum               NVWAIARDPNEWADPLEFRPERFLPGGEKPKVDVRGNDFEVIPFGAGRRICAGMNSGIRM
Q9SBQ9.1_F3-H_Petunia_x_hybrida                     NVWAIARDPNAWADPLEFRPERFLPGGEKPKVDVRGNDFEVIPFGAGRRICAGMNLGIRM
AAG49298.1_F3-H_Callistephus_chinensis              NVWAIARDPKMWTNPLEFRPSRFLPGGEKPDADIKGNDFEVIPFGAGRRICAGMSLGMRM
BAD91808.1_F3-H_Gentiana_triflora                   NVWAIARDPNAWANPLDFNPTRFLAGGEKPNVDVKGNDFEVIPFGAGRRICAGMSLGIRM
ABA64468.1_F3-H_Gerbera_hybrid_cultivar             NVWAISRDPKIWADPLEFQPTRFLPGGEKPNTDIKGNDFEVIPFGAGRRICVGMSLGLRM
AAO47847.1_F3-H_Glycine_max                         NIWAIARDPKEWNDPLEFRPERFLLGGEKADVDVRGNDFEVIPFGAGRRICAGLSLGLQM
ABC47161.1_F3-H_Pilosella_officinarum               NVWAIARDPKMWTEPLEFRPSRFLPDGEKPNADVKGNDFEVIPFGAGRRICAGMSLGLRM
BAD00190.1_F3-H_Ipomoea_nil                         NVWAIARDPNVWTNPLEFNPHRFLPGGEKPNVDIKGNDFEVIPFGAGRRICSGMSLGIRM
BAF49324.1_F3-H_Lobelia_erinus                      NVWAIARDPDIWAEPLEFRPQRFLPGGEKANVDVKGNDFEVIPFGAGRRVCAGLSLGLRM
AAG49301.1_F3-H_Matthiola_incana                    NIWAIARDPEQWSDPLAFRPERFLPGGEKFGVDVKGSDFELIPFGAGRRICAGLSLGLRT
ABB29899.1_F3-H_Osteospermum_hybrid_cultivar        NVWAIARDPKMWSEPLEFRPARFLPGGEKPDADVKGNDFEVIPFGAGRRSCAGMSLGLRM
AAG49315.1_F3-H_Pelargonium_x_hortorum              NVWAIARDPDVWADPLSFRPERFLPGSEKENVDVKGNDFELIPFGAGRRICAGMSLGLRM
BAB59005.1_F3-H_Perilla_frutescens_var._crispa      NVWAIGRDPNVWPDPLEFRPERFLMGGEKPNVDVRGNDFELIPFGSGRRICAGMNLGIRM
BAE47006.1_F3-H_Vitis_vinifera                      NVWAIARDPEVWEKPLEFRPSRFLPGGERPNADVRGNDFEVIPFGAGRRICAGMSLGLRM
AAR00229.1_F3-H_Ipomoea_purpurea                    NVWAIARDPNVWTNPLEFKPHRFLPGGEKPNVDIKGNDFEVIPFGAGRRICSGMSLGIRM
acc
```

```
Q9SD85.1_F3-H_Arabidopsis_thaliana           α16          β8        α17     β9     β10      β11
                                                   460       470       480       490       500       510
Q9SD85.1_F3-H_Arabidopsis_thaliana           IQFLTATLVQGFDWELAGGVTPEKLNMEESYGLTLQRAVPLVVHPKPRLAPNVYGLGSG
TRINITY_DN252_c0_g1_i1                       VQLLIATLVHGFDWELADGLMPDKLNMDEAFGLTLQRAVPLMVHPRPRLPLHAYGASN.
NP_001312537.1_F3-H_Nicotiana_tabacum        VQLMTATLIHAFNWDLSIGQSPEKLNMEEAFGLTLQRADPLVVHPCLRLEAQAYIG...
Q9SBQ9.1_F3-H_Petunia_x_hybrida              VQLMIATLIHAFNWDLVSGQLPEMLNMEEAYGLTLQRADPLVVHPRPRLEAQAYIG...
AAG49298.1_F3-H_Callistephus_chinensis       VQLLIATLVQTFDWELANGLDPEKLNMEEAYGLTLQRAEPLMVHPRPRLSPHVYESR..
BAD91808.1_F3-H_Gentiana_triflora            VQLVTASLVHSFDWALLDGLKPEKLDMEEGYGLTLQRASPLIVHPKPRLSAQVYCM...
ABA64468.1_F3-H_Gerbera_hybrid_cultivar      VQLLTATLIHAFDWELADGLNPKKLNMEEAYGLTLQRAAPLVVHPRPRLAPHVYETTKV
AAO47847.1_F3-H_Glycine_max                  VQLLTAALAHSFDWELEDCMNPEKLNMDEAYGLTLQRAVPLSVHPRPRLAPHVYSMSS.
ABC47161.1_F3-H_Pilosella_officinarum        VQLLTATLIQAFDWELANGLEPRNLNMEEAYGLTLQRAQPLMVHPRPRLAPHVYGTG..
BAD00190.1_F3-H_Ipomoea_nil                  VHLLVATLVHAFDWDLVNGQSVETLNMEEAYGLTLQRAVPLMLHPKPRLQPHLYTLN..
BAF49324.1_F3-H_Lobelia_erinus               VQLVTATLVHSFDWELADGQKPEELNMEEGYGLTLQRAKPLKVHPRPRLSEHVYQA...
AAG49301.1_F3-H_Matthiola_incana             IQLLTATLVHGFEWELAGGVTPEKLNMEETYGITVQRAVPLIVHPKPRLALNVYGVGSG
ABB29899.1_F3-H_Osteospermum_hybrid_cultivar VQLLVATLVQTFDWELANGLKPEKLNMEEAYGLTLQRAAPLLVHPKPRLAPHVYGSN..
AAG49315.1_F3-H_Pelargonium_x_hortorum       VQLLTATLLHAFNWDLPQGQIPQELNMDEAYGLTLQRASPLHVRPRPRLPSHLY.....
BAB59005.1_F3-H_Perilla_frutescens_var._crispa VQLLIATMVHAFDFELANGQLAKDLNMEEAYGITLQRADPLVVHPRPRLARHVYQAQV.
BAE47006.1_F3-H_Vitis_vinifera               VHLLTATLVHAFNWELPEGQVAEKLNMDEAYGLTLQRAPPLMVHPRPRLSPQVFGK...
AAR00229.1_F3-H_Ipomoea_purpurea             VHLLVATLVHAFDWDLMNGQSVETLNMEEAYGLTLQRAVPLMLHPKPRLQPHLYTLN..
                                         acc
```

```
NP_001234840.2_F3-5-H_Solanum_lycopersicum                 π1
                                                           1        10

NP_001234840.2_F3-5-H_Solanum_lycopersicum            .................................................MALRINELF.V
TRINITY_DN32466_c16_g7_i1                             .............................................MAFHALFFLQELS.V
TRINITY_DN32466_c16_g7_i3                             .............................................MAFHALFFLQELS.V
ACN38269.1_F3-5-H_Vitis_amurensis                     .............................................MAI.DTSLLLELA.A
NP_001268157.1_F3-5-H_Vitis_vinifera                  .............................................MAI.DTSLLLEFA.A
APY18930.1_F3-5-H_Camellia_sinensis_var._sinensis     .............................................MALDTVFLLRELS.F
BAA03438.1_F3-5-H_Petunia_x_hybrida                   ..................................................MMLLTELG.A
BAA03440.1_F3-5-H_Campanula_medium                    .............................................MSIDISTLFYELV.A
CAA50155.1_F3-5-H_Solanum_melongena                   .................................................MVILPSELI.G
BAC97831.1_F3-5-H_Vinca_major                         ...................................................MAIVDFL.A
CAA09850.1_F3-5-H_Catharanthus_roseus                 .............................................MAL.EKLVLFDFL.A
AAX51796.1_F3-5-H_Delphinium_grandiflorum             ..................................................MSISLFLA.G
BAA12735.1_F3-5-H_Gentiana_triflora                   .............................................MSPIYTTLTLHLA.T
AAM51564.1_F3-5-H_Glycine_max                         ...............................................MDSLLLLKEIA.T
AAP31058.1_F3-5-H_Gossypium_hirsutum                  .............................................MPSFDTILLRDLV.A
BAC10997.1_F3-5-H_Nierembergia_sp._NB17               .................................................MMQLSTELA.I
AAG49300.1_F3-5-H_Lycianthes_rantonnei                ...............................................M.VLLLISELC.A
AAV85471.1_F3-5-H_Solanum_tuberosum                   .................................................MTLRISELF.A
BAB20076.1_F3-5-H_Torenia_hybrid_cultivar             .............................................MTV.ATTLVVVLS.R
AAT34974.1_F3-5-H_Glandularia_x_hybrida               ...........................................MTFSELINILFWDLT.A
AAG49299.1_F3-5-H_Callistephus_chinensis              .............................................MS..ILSLLVYFC.I
ABB43030.1_F3-5-H_Pericallis_cruenta                  .............................................MS..ILTLICTF..I
ABB43031.1_F3-5-H_Osteospermum_hybrid_cultivar        .............................................MAI.IMTLLMDTC.I
AHI15949.1_F3-5-H_Pohlia_nutansSolanum_tuberosum      .............................................MGA.VEDAVAHVC.G
AHI15952.1_F3-5-H_Pohlia_nutansVinca_major            MKRLLLLVVSISTSVIRALVVAVGLCRIHIFAKENFPAFACFCKNMGASIVAAVAHVTRH
                                                 acc
```

```
NP_001234840.2_F3-5-H_Solanum_lycopersicum            π2                η1                          α1
                                                          20        30        40        50        60

NP_001234840.2_F3-5-H_Solanum_lycopersicum            AAIIY...IIVHIIISKLITTVRE.R.GRRLPLPPGPTGWPVIGALPLLGSMPHVALAKM
TRINITY_DN32466_c16_g7_i1                             SAILY...FVTKIIIEQLLPKPYRPR......LPPGPKGWPVIGALPLLGTMPHVSLAKL
TRINITY_DN32466_c16_g7_i3                             SAILY...FVTKIIIEQLLPKPYRPR......LPPGPKGWPVIGALPLLGTMPHVSLAKL
ACN38269.1_F3-5-H_Vitis_amurensis                     ATLLF...FITRFFIRSLLPKSSR.K......VPPGPKGWPLVGALPLLGNMPHVALAKM
NP_001268157.1_F3-5-H_Vitis_vinifera                  ATLLF...FITRFFIRSLLPKPSR.K......LPPGPKGWPLLGALPLLGNMPHVALAKM
APY18930.1_F3-5-H_Camellia_sinensis_var._sinensis     ATLVI...LITHIFMRSILSKPLR.M......LPPGPTGLPLIGAIPHLGSMPHVALAKM
BAA03438.1_F3-5-H_Petunia_x_hybrida                   ATSIF...LIAHIIISTLISKTTG.R.H......LPPGPRGWPVIGALPLLGAMPHVSLAKM
BAA03440.1_F3-5-H_Campanula_medium                    AISLY...LATYSFIRFLFKPSHHHH......LPPGPTGWPIIGALPLLGTMPHVSLADM
CAA50155.1_F3-5-H_Solanum_melongena                   ATIIY...IIVYIIIQKLIATGSW.R.RRR..LPPGPEGWPVIGALPLLGGMPHVALAKM
BAC97831.1_F3-5-H_Vinca_major                         AFLIF...ILTQKLIQTLFRRTYR.R.K....LPPGPKGWPVIGALPYLGTMPHTSLAYM
CAA09850.1_F3-5-H_Catharanthus_roseus                 AISIL...ILVQKFIQIVFLRSSS.RIR....LPPGPKGWPIIGALPYLGTMPHSILANM
AAX51796.1_F3-5-H_Delphinium_grandiflorum             AAILF...FVTHLLLSPT.......RTRK...LPPGPKGWPVVGALPMLGNMPHVALANL
BAA12735.1_F3-5-H_Gentiana_triflora                   ALFLF...FHVQKLVHYLHGKATGHRCRR....LPPGPTGWPILGALPLLGNMPHVTFANM
AAM51564.1_F3-5-H_Glycine_max                         SILIF...LITRLSIQTFLKSYRQ.K......LPPGPKGWPVVGALPLMGSMPHVTLAKM
AAP31058.1_F3-5-H_Gossypium_hirsutum                  AACLF...FITRYFIRRLLSNPKR.T......LPPGPKGWPIVGALPLLGSMPHVELAKL
BAC10997.1_F3-5-H_Nierembergia_sp._NB17               AAIIF...LLAHI....LISKTSG.R.R....LPPGPRGWPVIGALPLLGDMPHVSLAKM
AAG49300.1_F3-5-H_Lycianthes_rantonnei                SAIIF...IIVHIIISKLIATGWG.R.RQR..LPPGPMGWPVIGALPLLGTMPHVALAKM
AAV85471.1_F3-5-H_Solanum_tuberosum                   AAIIY...IIVHIIISKLIATVRE.R.GRR..LPPGPTGWPVIGALSLLGSMPHVALAKM
BAB20076.1_F3-5-H_Torenia_hybrid_cultivar             ELLYV..LVVYILVSKSLSTIIVVS.RKR..LPPGPTGFPVVGALPLLGSMPHVALAKM
AAT34974.1_F3-5-H_Glandularia_x_hybrida               AILLY...VALILVCSLYSKPSTVS.RN...LPPGPKGFPVVGALPLLGTMPHIALAKM
AAG49299.1_F3-5-H_Callistephus_chinensis              SLLVI......IALVNMFITRHTN.R......LPPGPAPWPVVGNLPHLGAIPHHTLAAL
ABB43030.1_F3-5-H_Pericallis_cruenta                  TGLMF......YGLVNL.LSRRAS.R......LPPGPTPWPIIGNLMHLGKLPHHSLADL
ABB43031.1_F3-5-H_Osteospermum_hybrid_cultivar        TVLVF......YVLLNL.RTRHPN.R......LPPGPTPWPIVGNLPHLGPLPHHTLAAF
AHI15949.1_F3-5-H_Pohlia_nutansSolanum_tuberosum      SFFEYGVAVLAIFLLIVLIQRP...K.....NLPPGPTGLPIIGALHLLGQRPHETLARM
AHI15952.1_F3-5-H_Pohlia_nutansVinca_major            SLLEYGVAVLAILGLVMVLRRPLK.K.....KLPPGPVGLPIMGSLHLLGPRPHQRLAQM
                                                 acc
```

```
NP_001234840.2_F3-5-H_Solanum_lycopersicum                    β1     β2        α2       η2        η3
                                                          70   TT  80        90       100       110       120   TT

NP_001234840.2_F3-5-H_Solanum_lycopersicum            AKKYGPIMYLKVGTCGMVVASTPNAAKAFLKTLDINFSNRPPNAGATHLAYNAQDMVFAP
TRINITY_DN32466_c16_g7_i1                             SKKYGEVMYLKMGTCNMVVASSPESAKAFLKTLDLNFSNRPPNAGATHLAYNAQDMVFAD
TRINITY_DN32466_c16_g7_i3                             SKKYGEVMYLKMGTCNMVVASSPESAKAFLKTLDLNFSNRPPNAGATHLAYNAQDMVFAD
ACN38269.1_F3-5-H_Vitis_amurensis                     AKRYGPVMFLKMGTNGMVVASTPGAARAFLKTLDINFSNRPPNAGATLLAYHAQDMVFAD
NP_001268157.1_F3-5-H_Vitis_vinifera                  AKRYGPVMFLKMGTNSMVVASTPEAARAFLKTLDINFSNRPPNAGATLLAYHAQDMVFAD
APY18930.1_F3-5-H_Camellia_sinensis_var._sinensis     AKIYGPIVYLKMGTCGMVVASTPDSARAFLKTLDLNFSNRPPNAGATHLAYGAQDMVFAD
BAA03438.1_F3-5-H_Petunia_x_hybrida                   AKKYGAIMYLKVGTCGMAVASTPDAAKAFLKTLDINFSNRPPNAGATHLAYNAQDMVFAH
BAA03440.1_F3-5-H_Campanula_medium                    AVKYGPIMYLKLGSKGTVVASNPKAARAFLKTHDANFSNRPIDGGPTYLAYNAQDMVFAE
CAA50155.1_F3-5-H_Solanum_melongena                   AKKYGPIMYLKVGTCGMVVASTPNAAKAFLKTLDINFSNRPPNAGATHMAYNAQDMVFAP
BAC97831.1_F3-5-H_Vinca_major                         AKKYGPVMYLKVGTNDMVVVSTPDAARAFLKTLDINFSNRPPNAGATHLAYGAQDMVFAH
CAA09850.1_F3-5-H_Catharanthus_roseus                 AKKYGPIMYLKLGTNGMVVASTPDAVKAFLRTLDMNFSNRPIDAGATHLAYNAQDMVFAH
AAX51796.1_F3-5-H_Delphinium_grandiflorum             SRRYGPIVYLKLGSRGMVVASTPDSARAFLKTQDLNFSNRPTDAGATHIAYNSQDMVFAD
BAA12735.1_F3-5-H_Gentiana_triflora                   AKKYGSVMYLKVGSHGLAIASTPDAAKAFLKTLDLNFSNRPPNAGATHLAYNAQDMVFAH
AAM51564.1_F3-5-H_Glycine_max                         AKKYGPIMYLKMGTNNMVVASTPAAARAFLKTLDQNFSNRPSNAGATHLAYDARDMVFAH
AAP31058.1_F3-5-H_Gossypium_hirsutum                  AKKYGPIMYLKMGTCNMVVASTPNAAKAFLKTLDLNFSNRPSNAGATHIAYNSQDMVFAE
BAC10997.1_F3-5-H_Nierembergia_sp._NB17               AKKYGPIMYLKVGTCGMAVASTPHAAKAFLKTLDINFSNRPPNAGATHFAYNAQDMVFAP
AAG49300.1_F3-5-H_Lycianthes_rantonnei                AKKYGPIMYLKVGTCGMVVASTPNAAKAFLKTLDINFSNRPPNAGATHLAYDAQDMVFAH
AAV85471.1_F3-5-H_Solanum_tuberosum                   AKNYGPIMYLKVGTCGMVVASTPNAAKAFLKTLDSNFSNRPPNAGATHLAYNAQDMVFAP
BAB20076.1_F3-5-H_Torenia_hybrid_cultivar             AKKYGPVMYLKTGTLGMVVASTPSSARAFLKTLDSNFSNRPANAGATHLAYGAQDMVFAP
AAT34974.1_F3-5-H_Glandularia_x_hybrida               AKTYGPVMYLKVGTWGMAVASTPDAARAFLKTLDTTFSDRPPNAGATLLAYNAQDMVFAH
AAG49299.1_F3-5-H_Callistephus_chinensis              ATKYGPLVYLRLGFVHVVVASSPSVAAQFLKVHDLKFASRPPNSGAKHIAYNYQDMVFAP
ABB43030.1_F3-5-H_Pericallis_cruenta                  AKKYGPLIHVRLGSVDVVVVASSASVAGQFLKVHDANFANRPPNSGAKHMAYNYHDMVFAP
ABB43031.1_F3-5-H_Osteospermum_hybrid_cultivar        AKKYGPLIHLRFGFVDVVVASSPTVASQFLRDNDLNFASRPPNSGAKHMAYNYQDLVFAP
AHI15949.1_F3-5-H_Pohlia_nutansSolanum_tuberosum      ADKYGPLMSLYMGQQLCIVASSRDTAMEFLKRQDAVFCSRPPQRGFEVIF..PKDITFSD
AHI15952.1_F3-5-H_Pohlia_nutansVinca_major            AEKYGPVMSLYMGQKLCVVATSADTAMEFLKAQDAVFCSRPPLRAAQVIF..PTDITFSD
                                                 acc
```

```
                                                          α3                         α4                          β3
NP_001234840.2_F3-5-H_Solanum_lycopersicum               130       140       150       160       170        180

NP_001234840.2_F3-5-H_Solanum_lycopersicum               YGPRWKLLRKLSNLHMLGGKALENWANVRANELGHMLKSMFDASQDG...EC.VVIADVL
TRINITY_DN32466_c16_g7_i1                                YGPRWKLLRKLSNLHMLGGKALDDWSNVRSQELGHMVRAMCEASQRN...EA.VVVPEML
TRINITY_DN32466_c16_g7_i3                                YGPRWKLLRKLSNLHMLGGKALDDWSNVRSQELGHMVRAMCEASQRN...EA.VVVPEML
ACN38269.1_F3-5-H_Vitis_amurensis                        YGARWKLLRKLSNLHMLGGKALEDWSQVRAVELGHMLRAMLELSQRT...EP.VVAPEML
NP_001268157.1_F3-5-H_Vitis_vinifera                     YGARWKLLRKLSNLHMLGGKALEDWSQVRAVELGHMLRAMLELCQRA...EP.VVVPEML
APY18930.1_F3-5-H_Camellia_sinensis_var._sinensis        YGPRWKLLRKLSNLHMLGGKALENWAQVRSSELGHMLKAMYDLSCRR...EP.VVVAEML
BAA03438.1_F3-5-H_Petunia_x_hybrida                      YGPRWKLLRKLSNLHMLGGKALENWANVRANELGHMLKSMSDMSREG...QR.VVVAEML
BAA03440.1_F3-5-H_Campanula_medium                       YGPKWKLLRKLCSLHMLGPKALEDWAHVKVSEVGHMLKEMYEQSSKSV.PVP.VVVPEML
CAA50155.1_F3-5-H_Solanum_melongena                      YGPRWKLLRKLSNLHMLGGKALENWANVRANELGHMLKSMFDASHVG...ER.IVVADML
BAC97831.1_F3-5-H_Vinca_major                            YGPKWKLLRKLSNLHMRGGKALEDWSNVRISELGHMLETMYESSKKG...EA.VVVAEML
CAA09850.1_F3-5-H_Catharanthus_roseus                    YGPKWKLLRKLSNLHMLGGKALENWSNVRATELGYMLQAMYESSRKG...ET.VVVPEML
AAX51796.1_F3-5-H_Delphinium_grandiflorum                YGPRWKLLRKLSSLHMLGGKAIEDWAVVRRDEVGYMVKAIYESSCAG...EA.VHVPDML
BAA12735.1_F3-5-H_Gentiana_triflora                      YGPKWKLLRKLSNLHMLGGKALENWADVRKTELGYMLKAMFESSQNN...EP.VMISEML
AAM51564.1_F3-5-H_Glycine_max                            YGSRWKLLRKLSNLHMLGGKALDDWAQIRDEEMGHMLGAMYDCNKRD...EA.VVVAEML
AAP31058.1_F3-5-H_Gossypium_hirsutum                     YGPRWKLLRKLSNLHMLGGKALEDWSQVRAVELGHMLRAMCESSRKG...EP.VVVPEML
BAC10997.1_F3-5-H_Nierembergia_sp._NB17                  YGPRWKLLRKLSNLHMLGGKALEDWSNVRANELGHMLKSMLDLSREG...QR.VVIADML
AAG49300.1_F3-5-H_Lycianthes_rantonnei                   YGPRWKLLRKLSNLHMLGGKALEDWANVRANELGHMLKSMFDASRVG...GR.VVVADML
AAV85471.1_F3-5-H_Solanum_tuberosum                      YGPRWKLLRKLSNLQMLGGKALENWANVRANELGHMIKSMFDASHVG...EC.VVIADML
BAB20076.1_F3-5-H_Torenia_hybrid_cultivar                YGPRWRLLRRLSNLHMLGGKALDGWANVRASELGHMLEAMNRASREG...EV.VVVPEML
AAT34974.1_F3-5-H_Glandularia_x_hybrida                  YGPRWRLLRKLSNLHMLGAKALEDWADVRSSEVGHMLQAMLTSSLRH...EA.VALPEML
AAG49299.1_F3-5-H_Callistephus_chinensis                 YGPQWTMFRKICKDHLFSSKALDDFRHVRQEEVAILARGLAGAGR.....SK.VNLGQQL
ABB43030.1_F3-5-H_Pericallis_cruenta                     YGPRWRMLRKMCSMHLFSAKALTDFRQVRQEEVTILTRVLARTGQ.....SA.VKLDQQL
ABB43031.1_F3-5-H_Osteospermum_hybrid_cultivar           YGPRWTMLRKICKDHLFSSKALDNFRHVREEEVAILTRILASAGE.....ST.VKLSQLL
AHI15949.1_F3-5-H_Pohlia_nutansSolanum_tuberosum         ITPASRHLRKILHSQLTSGRRIEEAEHIRADEIAQTVRSIA........PGL.VDLKVRL
AHI15952.1_F3-5-H_Pohlia_nutansVinca_major               ITPASRQMRKMLHLQLTSGKRIEESEQIRSEEMAHMVRTIAASGRGGEEPELTVNVKASL
acc
```

```
                                                         α5                                      α6                α7   α8
NP_001234840.2_F3-5-H_Solanum_lycopersicum                     190       200                 210       220        230

NP_001234840.2_F3-5-H_Solanum_lycopersicum               TFAMANMIGQVMLSKRVFV.........EKGVEVNEFKNMVVELMTVAGYFNIGDFIPKL
TRINITY_DN32466_c16_g7_i1                                TYAMANMIGQIILSRRVFV.........TKGSESNEFKDMVVELMTSAGYFNIGDFIPSI
TRINITY_DN32466_c16_g7_i3                                TYAMANMIGQIILSRRVFV.........TKGSESNEFKDMVVELMTSAGYFNIGDFIPSI
ACN38269.1_F3-5-H_Vitis_amurensis                        TFSMANMIGQVILSRRVFE.........TKGSESNEFKDMVVELMTTAGYFNIGDFIPSI
NP_001268157.1_F3-5-H_Vitis_vinifera                     TFSMANMIGQVILSRRVFE.........TKGSESNEFKDMVVELMTSAGYFNIGDFIPSI
APY18930.1_F3-5-H_Camellia_sinensis_var._sinensis        CYAMANMIGQVILSRRVFE.........TKGSESNEFKDMVVELMTTAGYFNIGDFIPSI
BAA03438.1_F3-5-H_Petunia_x_hybrida                      TFAMANMIGQVMLSKRVFV.........DKGVEVNEFKDMVVELMTIAGYFNIGDFIPCL
BAA03440.1_F3-5-H_Campanula_medium                       TYAMANMIGRIILSRRPFVITSKLDSSASASASVSEFQYMVMELMRMAGLFNIGDFIPYI
CAA50155.1_F3-5-H_Solanum_melongena                      TFAMANMIGQVMLSKRVFV.........EKGKEVNEFKNMVVELMTVAGYFNIGDFIPQI
BAC97831.1_F3-5-H_Vinca_major                            TYAMANMIGQVILSRRVFV.........TKGSDSNEFKDMVVELMTSAGLFNIGDFIPSI
CAA09850.1_F3-5-H_Catharanthus_roseus                    TYAMANMIGQVILSRRVFV.........TKSLESNEFKDMVVELMTTAGYFNIGDFIPSI
AAX51796.1_F3-5-H_Delphinium_grandiflorum                VFAMANMLGQVILSRRVFV.........TKGVESNEFKEMVIELMTSAGLFNVGDFIPSI
BAA12735.1_F3-5-H_Gentiana_triflora                      TYAMANMLSQVILSRRVFN.........KKGAKSNEFKDMVVELMTSAGYFNIGDFIPSI
AAM51564.1_F3-5-H_Glycine_max                            IF.MANMIGQVILSRRVFE.........TKGSESNEFKDMVVELMTVAGYFNIGDFIPFL
AAP31058.1_F3-5-H_Gossypium_hirsutum                     TYAMANMIGQVILSRRVFV.........TKGSESNEFKDMVVELMTSAGLFNIGDFIPSI
BAC10997.1_F3-5-H_Nierembergia_sp._NB17                  TFAMANMIGQVILSKRVFV.........DKGVEVNEFKNMVVELMTVAGYFNIGDYIPCL
AAG49300.1_F3-5-H_Lycianthes_rantonnei                   TFAMANMIGQVILSKRVFV.........EKGAEVNKFKDMVVELMTVAGYFNIGDFIPQL
AAV85471.1_F3-5-H_Solanum_tuberosum                      TFAMANMIGQVMLSKRVFV.........EKGAEVNEFKNMVVELMTVAGYFNIGDFIPKL
BAB20076.1_F3-5-H_Torenia_hybrid_cultivar                VYAMANMIGQVILSRRVFV.........TKGEEVNEFKDMVVELMTSAGYFNIGDFVPIL
AAT34974.1_F3-5-H_Glandularia_x_hybrida                  IYSMANMIGQVILSRRVFE.........TKGKEVNEFKDMVVELMTSAGYFNIGDFIPAL
AAG49299.1_F3-5-H_Callistephus_chinensis                 NMCTANTLARMMLDKRVFG....NESGGDDPKANEFKEMATELMFLAGQFNIGDYIPVL
ABB43030.1_F3-5-H_Pericallis_cruenta                     NVCFANTLSRMMLDRRVFG.....D....GDPKADDYKDMVVELMTLAGQFNIGDYIPWL
ABB43031.1_F3-5-H_Osteospermum_hybrid_cultivar           NVCTANTLARMMLDMRLFG.....DGNGSGDPKADEFKDMVVELMTLAGKFNIGDFIPML
AHI15949.1_F3-5-H_Pohlia_nutansSolanum_tuberosum         AVMTANILTRLIISKRFMGRSG.MNS..TEEKELQDFVDITEEIGDCLGTANPRDVIPAF
AHI15952.1_F3-5-H_Pohlia_nutansVinca_major               DAMTANIFTRLIISKRFFSPGGILRG..MQEAELREFIAITDEIDLCLGLPNPRDLIPAF
acc
```

```
                                                                       α9                                 α10
NP_001234840.2_F3-5-H_Solanum_lycopersicum                     240       250       260       270       280

NP_001234840.2_F3-5-H_Solanum_lycopersicum               AWMD.I.QGIEKGMKNLHKKFDDLLTKMFDEHEATSNERKENPDFLDVVMANR.DN..SE
TRINITY_DN32466_c16_g7_i1                                AWMD.L.QGIEKGMKKLHKKFDVLLTKMMEEHMATTHLRKGKPDFLDVIMANR.QN..SD
TRINITY_DN32466_c16_g7_i3                                AWMD.L.QGIEKGMKKLHKKFDVLLTKMMEEHMATTHLRKGKPDFLDVIMANR.QN..SD
ACN38269.1_F3-5-H_Vitis_amurensis                        AWLD.I.QGIQHGMKHLHRKFDRLLTKMMEEHTASAHERKGNPDFLDVIMANQ.EN..ST
NP_001268157.1_F3-5-H_Vitis_vinifera                     AWLD.I.QGIQRGMKHLHRKFDWLLTKMMEEHTASAHERKGNPDFLDVIMANQ.EN..ST
APY18930.1_F3-5-H_Camellia_sinensis_var._sinensis        AWMD.L.QGIERGMKRLHKKFDVLITKMIEEHKASAYQRKETPDFLDVFMAQQ.EN..PG
BAA03438.1_F3-5-H_Petunia_x_hybrida                      AWMD.L.QGIEKRMKRLHKKFDALLTKMFDEHKATTYERKGKPDFLDVVMENG.DN..SE
BAA03440.1_F3-5-H_Campanula_medium                       AWMD.L.QGIQRDMKVIQKKFDVLLNKMIKEHTESAHDRKDNPDFLDILMAATQEN..TE
CAA50155.1_F3-5-H_Solanum_melongena                      AWMD.L.QGIEKGMKKLHKKFDDLLTKMFEEHEATSNERKGKPDFLDFIMANR.DN..SE
BAC97831.1_F3-5-H_Vinca_major                            AWMD.L.QGIEGGMKRLHKKFDVLLTKLLEEHKESSSKRKGKPDLLDYVLANR.DN..SE
CAA09850.1_F3-5-H_Catharanthus_roseus                    AWMD.L.QGIERGMKRLHKKFDALLTKMLEEHKSSSHKRKEKPDFLDYVLANR.DN..SE
AAX51796.1_F3-5-H_Delphinium_grandiflorum                AWMD.L.QGIVRGMKRLHNKFDALLDKILREHTATRSERKGKPDLVDVLMDNRDNK..SE
BAA12735.1_F3-5-H_Gentiana_triflora                      GWMD.L.QGIEGGMKRLHKKFDVLLTRLLDDHKRTSQERKQKPDFLDFVIANG.DN..SD
AAM51564.1_F3-5-H_Glycine_max                            AKLD.L.QGIERGMKKLHKKFDALLTSMIEEHVASSHKRKGKPDFLDMVMAHHSEN..SD
AAP31058.1_F3-5-H_Gossypium_hirsutum                     AWMD.L.QGIEGEMKKLHNRWDVLLTKMMKEHEETAYERKGKPDFLDIIMDNR.EN..SA
BAC10997.1_F3-5-H_Nierembergia_sp._NB17                  AWLD.L.QGIEKGMKILHKKFDVLLTKMFDEHKTTSHEREGKSDFLDMVMANR.DN..SE
AAG49300.1_F3-5-H_Lycianthes_rantonnei                   AWMD.L.QGIEKGMKCLHKKFDDLLTKMFDEHKATTNERKGKPDFLDVVMANR.DN..SE
AAV85471.1_F3-5-H_Solanum_tuberosum                      AWMD.L.QGIEKGMKSLHKKFDDLLTKMFDEHEATTYERKENPDFLDVVMANK.DN..SE
BAB20076.1_F3-5-H_Torenia_hybrid_cultivar                AWMD.L.QGIERGMKALHKKFDKLITRMLEQHEASAH....KPDFLDAIIAASLDDYSSE
AAT34974.1_F3-5-H_Glandularia_x_hybrida                  AWMD.L.QGIEKGMKSLHKKFDDLIGRMLDKHLESAHKRNAKPDFLDVILANR.HN..SE
AAG49299.1_F3-5-H_Callistephus_chinensis                 DWLD.L.QGIVKKMKKLHTRFDKFLDVILDEHKVIAS...GHIDMLSTLISLK.DDTSVD
ABB43030.1_F3-5-H_Pericallis_cruenta                     DLLD.L.QGIVKRMKKVHSQFDSFLDTIIDEHTIGTG...RHVDMLSTMISLK.DNADGE
ABB43031.1_F3-5-H_Osteospermum_hybrid_cultivar           DWMD.L.QGVTTKMKKVHDRFDSFLDNILDEHKIRKG...AHDDMLSTLISLK.EDVNGE
AHI15949.1_F3-5-H_Pohlia_nutansSolanum_tuberosum         KCIDILGLGLDRRFTNLRRHMESFLSKIIAERRQNSK..LPEKDLLDVLLEQL......D
AHI15952.1_F3-5-H_Pohlia_nutansVinca_major               RWLD.LASGLDKRFHSLRRRMESFLSTIIAEHRQHHS..FEKLDLLHVLLQQMEKD..DD
acc
```

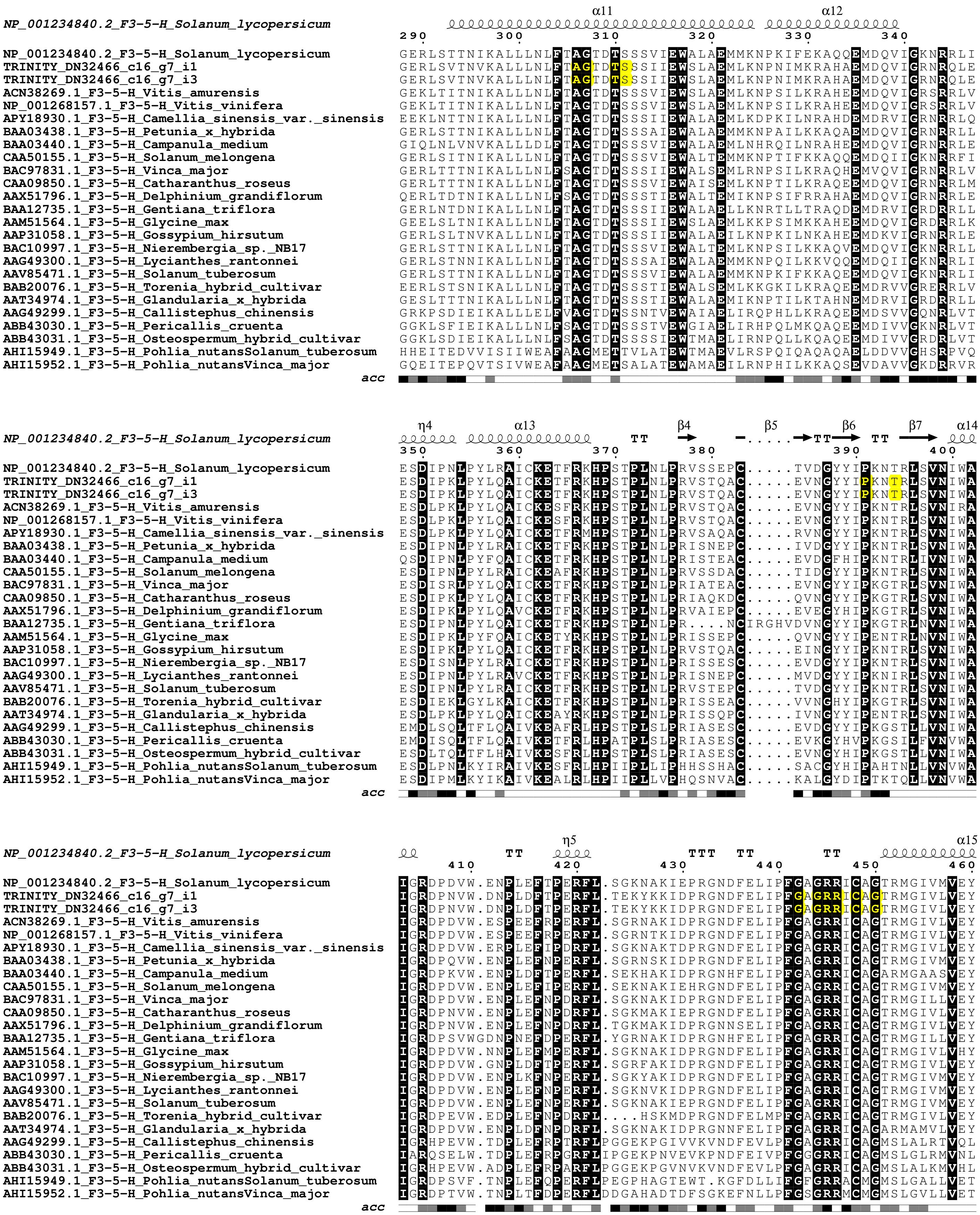

NP_001234840.2_F3-5-H_Solanum_lycopersicum
α11
α12
290
300
310
320
330
340
NP_001234840.2_F3-5-H_Solanum_lycopersicum
TRINITY_DN32466_c16_g7_i1
TRINITY_DN32466_c16_g7_i3
ACN38269.1_F3-5-H_Vitis_amurensis
NP_001268157.1_F3-5-H_Vitis_vinifera
APY18930.1_F3-5-H_Camellia_sinensis_var._sinensis
BAA03438.1_F3-5-H_Petunia_x_hybrida
BAA03440.1_F3-5-H_Campanula_medium
CAA50155.1_F3-5-H_Solanum_melongena
BAC97831.1_F3-5-H_Vinca_major
CAA09850.1_F3-5-H_Catharanthus_roseus
AAX51796.1_F3-5-H_Delphinium_grandiflorum
BAA12735.1_F3-5-H_Gentiana_triflora
AAM51564.1_F3-5-H_Glycine_max
AAP31058.1_F3-5-H_Gossypium_hirsutum
BAC10997.1_F3-5-H_Nierembergia_sp._NB17
AAG49300.1_F3-5-H_Lycianthes_rantonnei
AAV85471.1_F3-5-H_Solanum_tuberosum
BAB20076.1_F3-5-H_Torenia_hybrid_cultivar
AAT34974.1_F3-5-H_Glandularia_x_hybrida
AAG49299.1_F3-5-H_Callistephus_chinensis
ABB43030.1_F3-5-H_Pericallis_cruenta
ABB43031.1_F3-5-H_Osteospermum_hybrid_cultivar
AHI15949.1_F3-5-H_Pohlia_nutansSolanum_tuberosum
AHI15952.1_F3-5-H_Pohlia_nutansVinca_major
acc
η4
α13
TT
β4
β5
TT
β6
TT
β7
α14
350
360
370
380
390
400
acc
η5
TT
TTT
TT
TT
α15
410
420
430
440
450
460
acc

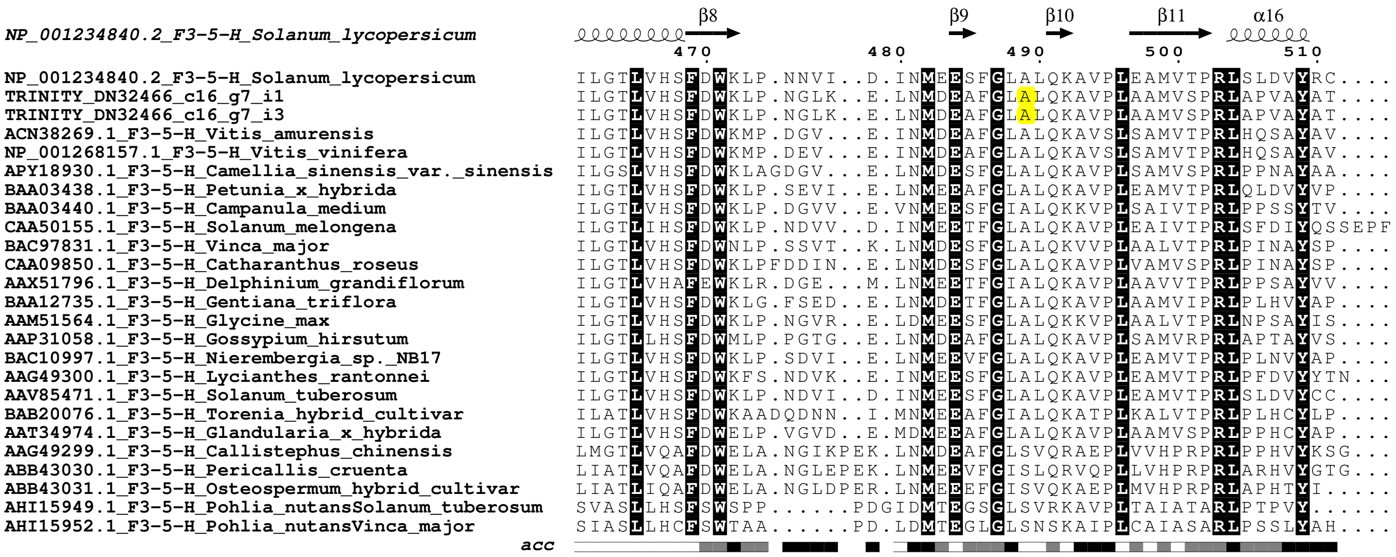

NP_001234840.2_F3-5-H_Solanum_lycopersicum
β8
β9
β10
β11
α16
470
480
490
500
510
NP_001234840.2_F3-5-H_Solanum_lycopersicum ILGTLVHSFDWKLP.NNVI..D.INMEESFGLALQKAVPLEAMVTPRLSLDVYRC....
TRINITY_DN32466_c16_g7_i1 ILGTLVHSFDWKLP.NGLK..E.LNMDEAFGLALQKAVPLAAMVSPRLAPVAYAT....
TRINITY_DN32466_c16_g7_i3 ILGTLVHSFDWKLP.NGLK..E.LNMDEAFGLALQKAVPLAAMVSPRLAPVAYAT....
ACN38269.1_F3-5-H_Vitis_amurensis ILGTLVHSFDWKMP.DGV...E.INMDEAFGLALQKAVSLSAMVTPRLHQSAYAV....
NP_001268157.1_F3-5-H_Vitis_vinifera ILGTLVHSFDWKMP.DEV...E.INMDEAFGLALQKAVSLSAMVTPRLHQSAYAV....
APY18930.1_F3-5-H_Camellia_sinensis_var._sinensis ILGSLVHSFDWKLAGDGV...E.LNMDESFGLALQKAVPLSAMVSPRLPPNAYAA....
BAA03438.1_F3-5-H_Petunia_x_hybrida ILGTLVHSFDWKLP.SEVI..E.LNMEEAFGLALQKAVPLEAMVTPRLQLDVYVP....
BAA03440.1_F3-5-H_Campanula_medium ILGTLVHSFDWKLP.DGVV..E.VNMEESFGIALQKKVPLSAIVTPRLPPSSYTV....
CAA50155.1_F3-5-H_Solanum_melongena ILGTLIHSFDWKLP.NDVV..D.INMEETFGLALQKAVPLEAIVTPRLSFDIYQSSEPF
BAC97831.1_F3-5-H_Vinca_major ILGTLVHSFDWNLP.SSVT..K.LNMDESFGLALQKVVPLAALVTPRLPINAYSP....
CAA09850.1_F3-5-H_Catharanthus_roseus ILGTLVHSFDWKLPFDDIN..E.LNMDESFGLALQKAVPLVAMVSPRLPINAYSP....
AAX51796.1_F3-5-H_Delphinium_grandiflorum ILGTLVHAFEWKLR.DGE...M.LNMEETFGIALQKAVPLAAVVTPRLPPSAYVV....
BAA12735.1_F3-5-H_Gentiana_triflora ILGTLVHSFDWKLG.FSED..E.LNMDETFGLALQKAVPLAAMVIPRLPLHVYAP....
AAM51564.1_F3-5-H_Glycine_max ILGTLVHSFDWKLP.NGVR..E.LDMEESFGLALQKKVPLAALVTPRLNPSAYIS....
AAP31058.1_F3-5-H_Gossypium_hirsutum ILGTLLHSFDWMLP.PGTG..E.LNMDEAFGLALQKAVPLSAMVRPRLAPTAYVS....
BAC10997.1_F3-5-H_Nierembergia_sp._NB17 ILGTLVHSFDWKLP.SDVI..E.LNMEEVFGLALQKAVPLEAMVTPRLPLNVYAP....
AAG49300.1_F3-5-H_Lycianthes_rantonnei ILGTLVHSFDWKFS.NDVK..E.INMEESFGLALQKAVPLEAMVTPRLPFDVYYTN...
AAV85471.1_F3-5-H_Solanum_tuberosum ILGTLVHSFDWKLP.NDVI..D.INMEESFGLALQKAVPLEAMVTPRLSLDVYCC....
BAB20076.1_F3-5-H_Torenia_hybrid_cultivar ILATLVHSFDWKAADQDNN..I.MNMEEAFGIALQKATPLKALVTPRLPLHCYLP....
AAT34974.1_F3-5-H_Glandularia_x_hybrida ILGTLVHSFDWELP.VGVD..E.MDMEEAFGLALQKAVPLAAMVSPRLPPHCYAP....
AAG49299.1_F3-5-H_Callistephus_chinensis LMGTLVQAFDWELA.NGIKPEK.LNMDEAFGLSVQRAEPLVVHPRPRLPPHVYKSG...
ABB43030.1_F3-5-H_Pericallis_cruenta LIATLVQAFDWELA.NGLEPEK.LNMEEVFGISLQRVQPLLVHPRPRLARHVYGTG...
ABB43031.1_F3-5-H_Osteospermum_hybrid_cultivar LIATLIQAFDWELA.NGLDPER.LNMEEEFGISVQKAEPLMVHPRPRLAPHTYI.....
AHI15949.1_F3-5-H_Pohlia_nutansSolanum_tuberosum SVASLLHSFSWSPP......PDGIDMTEGSGLSVRKAVPLTAIATARLPTPVY......
AHI15952.1_F3-5-H_Pohlia_nutansVinca_major SIASLLHCFSWTAA......PD.LDMTEGLGLSNSKAIPLCAIASARLPSSLYAH....
acc

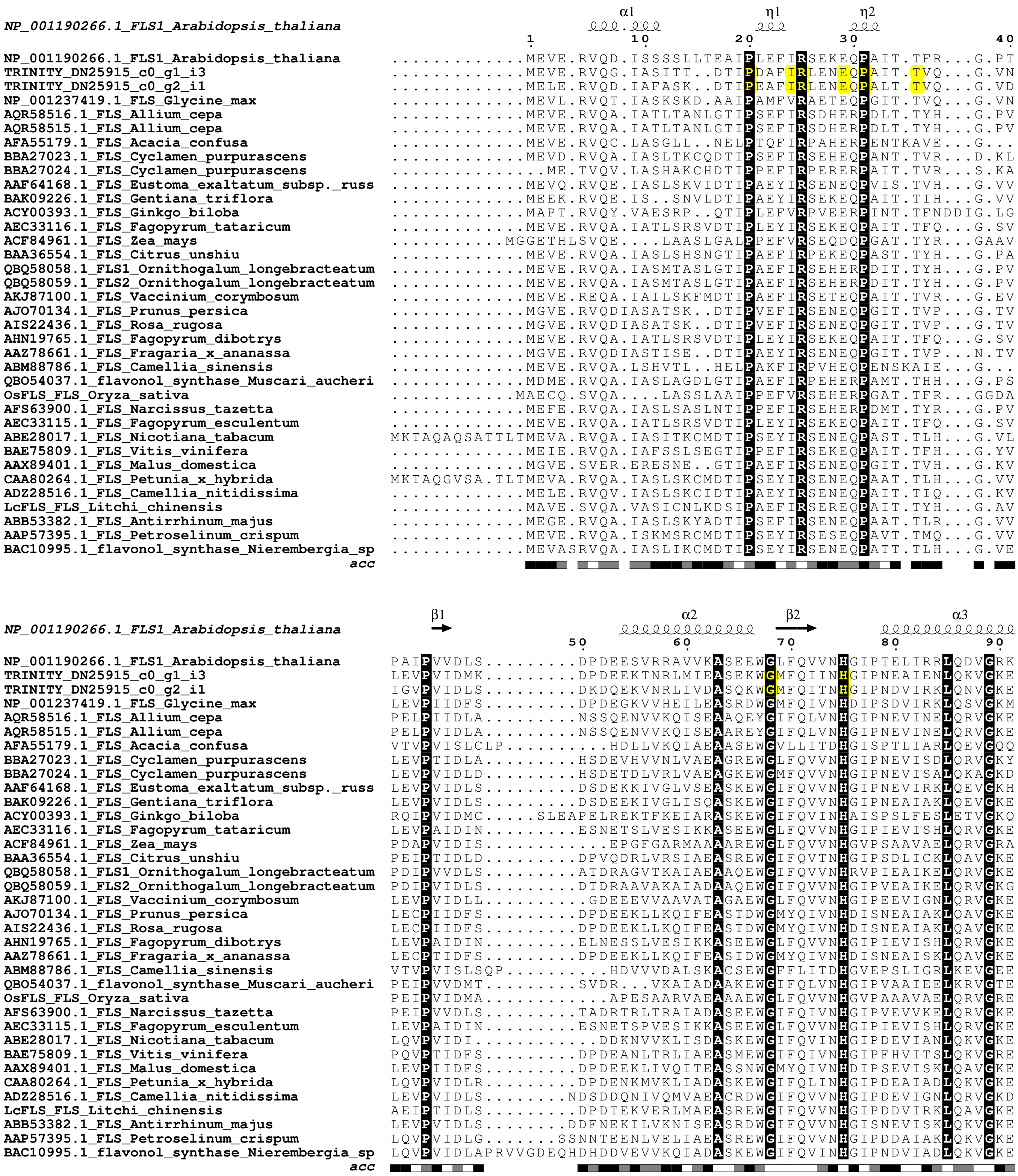
NP_001190266.1_FLS1_Arabidopsis_thaliana
α1 η1 η2
1 10 20 30 40
NP_001190266.1_FLS1_Arabidopsis_thaliana ............MEVE.RVQD.ISSSSLLTEAIPLEFIRSEKEQPAIT.TFR...G.PT
TRINITY_DN25915_c0_g1_i3 ............MEVE.RVQG.IASITT..DTIPDAFIRLENEQPAIT.TVQ...G.VN
TRINITY_DN25915_c0_g2_i1 ............MELE.RVQD.IAFASK..DTIPEAFIRLENEQPALT.TVQ...G.VD
NP_001237419.1_FLS_Glycine_max ............MEVL.RVQT.IASKSKD.AAIPAMFVRAETEQPGIT.TVQ...G.VN
AQR58516.1_FLS_Allium_cepa ............MEVE.RVQA.IATLTANLGTIPSEFIRSDHERPDLT.TYH...G.PV
AQR58515.1_FLS_Allium_cepa ............MEVE.RVQA.IATLTANLGTIPSEFIRSDHERPDLT.TYH...G.PV
AFA55179.1_FLS_Acacia_confusa ............MEVE.RVQC.LASGLL..NELPTQFIRPAHERPENTKAVE...G...
BBA27023.1_FLS_Cyclamen_purpurascens ............MEVD.RVQA.IASLTKCQDTIPSEFIRSEHEQPANT.TVR...D.KL
BBA27024.1_FLS_Cyclamen_purpurascens ..............ME.TVQV.LASHAKCHDTIPPEFIRPERERPAIT.TVR...S.KA
AAF64168.1_FLS_Eustoma_exaltatum_subsp._russ ............MEVQ.RVQE.IASLSKVIDTIPAEYIRSENEQPVIS.TVH...G.VV
BAK09226.1_FLS_Gentiana_triflora ............MEEK.RVQE.IS..SNVLDTIPAEYIRSEKEQPAIT.TIH...G.VV
ACY00393.1_FLS_Ginkgo_biloba ............MAPT.RVQY.VAESRP..QTIPLEFVRPVEERPINT.TFNDDIG.LG
AEC33116.1_FLS_Fagopyrum_tataricum ............MEVE.RVQA.IATLSRSVDTIPLEYIRSEKEQPAIT.TFQ...G.SV
ACF84961.1_FLS_Zea_mays ..........MGGETHLSVQE....LAASLGALPPEFVRSEQDQPGAT.TYR...GAAV
BAA36554.1_FLS_Citrus_unshiu ............MEVE.RVQA.IASLSHSNGTIPAEFIRPEKEQPAST.TYH...G.PA
QBQ58058.1_FLS1_Ornithogalum_longebracteatum ............MEVE.RVQA.IASMTASLGTIPAEFIRSEHERPDIT.TYH...G.PV
QBQ58059.1_FLS2_Ornithogalum_longebracteatum ............MEVE.RVQA.IASMTASLGTIPAEFIRSEHERPDIT.TYH...G.PV
AKJ87100.1_FLS_Vaccinium_corymbosum ............MEVE.REQA.IAILSKFMDTIPAEFIRSETEQPAIT.TVR...G.EV
AJO70134.1_FLS_Prunus_persica ............MGVE.RVQDIASATSK..DTIPVEFIRSENEQPGIT.TVP...G.TV
AIS22436.1_FLS_Rosa_rugosa ............MGVE.RVQDIASATSK..DTIPVEFIRSENEQPGIT.TVP...G.TV
AHN19765.1_FLS_Fagopyrum_dibotrys ............MEVE.RVQA.IATLSRSVDTIPLEYIRSEKEQPAIT.TFQ...G.SV
AAZ78661.1_FLS_Fragaria_x_ananassa ............MGVE.RVQDIASTISE..DTIPAEYIRSENEQPGIT.TVP...N.TV
ABM88786.1_FLS_Camellia_sinensis ............MEVE.RVQA.LSHVTL..HELPAKFIRPVHEQPENSKAIE...G...
QBO54037.1_flavonol_synthase_Muscari_aucheri ............MDME.RVQA.IASLAGDLGTIPAEFIRPEHERPAMT.THH...G.PS
OsFLS_FLS_Oryza_sativa ...........MAECQ.SVQA....LASSLAAIPPEFVRSEHERPGAT.TFR...GGDA
AFS63900.1_FLS_Narcissus_tazetta ............MEFE.RVQA.IASLSASLNTIPPEFIRSEHERPDMT.TYR...G.PV
AEC33115.1_FLS_Fagopyrum_esculentum ............MEVE.RVQA.IATLSRSVDTIPLEYIRSEKEQPAIT.TFQ...G.SV
ABE28017.1_FLS_Nicotiana_tabacum MKTAQAQSATTLTMEVA.RVQA.IASITKCMDTIPSEYIRSENEQPAST.TLH...G.VL
BAE75809.1_FLS_Vitis_vinifera ............MEIE.RVQA.IAFSSLSEGTIPPEFIRSEKEQPAIT.TFH...G.YV
AAX89401.1_FLS_Malus_domestica ............MGVE.SVER.ERESNE..GTIPAEFIRSENEQPGIT.TVH...G.KV
CAA80264.1_FLS_Petunia_x_hybrida MKTAQGVSA.TLTMEVA.RVQA.IASLSKCMDTIPSEYIRSENEQPAAT.TLH...G.VV
ADZ28516.1_FLS_Camellia_nitidissima ............MELE.RVQV.IASLSKCIDTIPAEYIRSENEQPAIT.TIQ...G.KV
LcFLS_FLS_Litchi_chinensis ............MAVE.SVQA.VASICNLKDSIPAEFIRSEREQPAIT.TYH...G.AV
ABB53382.1_FLS_Antirrhinum_majus ............MEGE.RVQA.IASLSKYADTIPSEFIRSENEQPAAT.TLR...G.VV
AAP57395.1_FLS_Petroselinum_crispum ............MEVE.RVQA.ISKMSRCMDTIPSEYIRSESEQPAVT.TMQ...G.VV
BAC10995.1_flavonol_synthase_Nierembergia_sp ............MEVASRVQA.IASLIKCMDTIPSEYIRSENEQPATT.TLH...G.VE
acc
NP_001190266.1_FLS1_Arabidopsis_thaliana
β1 α2 β2 α3
50 60 70 80 90
NP_001190266.1_FLS1_Arabidopsis_thaliana PAIPVVDLS.........DPDEESVRRAVVKASEEWGLFQVVNHGIPTELIRRLQDVGRK
TRINITY_DN25915_c0_g1_i3 LEVPVIDMK.........DPDEEKTNRLMIEASEKWGMFQIINHGIPNEAIENLQKVGKE
TRINITY_DN25915_c0_g2_i1 IGVPVIDLS.........DKDQEKVNRLIVDASQKWGMFQITNHGIPNDVIRKLQNVGKE
NP_001237419.1_FLS_Glycine_max LEVPIIDFS.........DPDEGKVVHEILEASRDWGMFQIVNHDIPSDVIRKLQSVGKM
AQR58516.1_FLS_Allium_cepa PELPIIDLA.........NSSQENVVKQISEAAQEYGIFQLVNHGIPNEVINELQRVGKE
AQR58515.1_FLS_Allium_cepa PELPVIDLA.........NSSQENVVKQISEAAREYGIFQLVNHGIPNEVINELQRVGKE
AFA55179.1_FLS_Acacia_confusa VTVPVISLCLP..........HDLLVKQIAEAASEWGVLLITDHGISPTLIARLQEVGQQ
BBA27023.1_FLS_Cyclamen_purpurascens LEVPTIDLA.........HSDEVHVVNLVAEAGREWGMFQVVNHGIPNEVISDLQRVGKY
BBA27024.1_FLS_Cyclamen_purpurascens LEVPVIDLD.........HSDETDLVRLVAEAGKEWGMFQVVNHGIPNEVISALQKAGKD
AAF64168.1_FLS_Eustoma_exaltatum_subsp._russ LEVPVIDLS.........DSDEKKIVGLVSEASKEWGIFQVVNHGIPNEVIRKLQEVGKH
BAK09226.1_FLS_Gentiana_triflora LEVPVIDLS.........DSDEEKIVGLISQASKEWGIFQVVNHGIPNEAIAKLQEVGKE
ACY00393.1_FLS_Ginkgo_biloba RQIPVIDMC.....SLEAPELREKTFKEIARASKEWGIFQVINHAISPSLFESLETVGKQ
AEC33116.1_FLS_Fagopyrum_tataricum LEVPAIDIN.........ESNETSLVESIKKASEEWGLFQVVNHGIPIEVISHLQRVGKE
ACF84961.1_FLS_Zea_mays PDAPVIDIS............EPGFGARMAAAAREWGLFQVVNHGVPSAAVAELQRVGRA
BAA36554.1_FLS_Citrus_unshiu PEIPTIDLD.........DPVQDRLVRSIAEASREWGIFQVTNHGIPSDLICKLQAVGKE
QBQ58058.1_FLS1_Ornithogalum_longebracteatum PDIPVVDLS.........ATDRAGVTKAIAEAAQEWGIFQVVNHRVPIEAIKELQRVGKE
QBQ58059.1_FLS2_Ornithogalum_longebracteatum PDIPVIDLS.........DTDRAAVAKAIADAAQEWGIFQVVNHGIPVEAIKELQRVGKG
AKJ87100.1_FLS_Vaccinium_corymbosum LEVPVIDLL..........GDEEEVVAAVATAGAEWGIFQVVNHGIPEEVIGNLQRVGKE
AJO70134.1_FLS_Prunus_persica LECPIIDFS.........DPDEEKLLKQIFEASTDWGMYQIVNHDISNEAIAKLQAVGKE
AIS22436.1_FLS_Rosa_rugosa LECPIIDFS.........DPDEEKLLKQIFEASTDWGMYQIVNHDISNEAIAKLQAVGKE
AHN19765.1_FLS_Fagopyrum_dibotrys LEVPAIDIN.........ELNESSLVESIKKASEEWGLFQVVNHGIPIEVISHLQRVGKE
AAZ78661.1_FLS_Fragaria_x_ananassa LECPTIDFS.........DPDEEKLLKQIFEASIDWGMYQIVNHDISNEAISKLQAVGKE
ABM88786.1_FLS_Camellia_sinensis VTVPVISLSQP..........HDVVVDALSKACSEWGFFLITDHGVEPSLIGRLKEVGEE
QBO54037.1_flavonol_synthase_Muscari_aucheri PEIPVVDMT.........SVDR...VKAIADAAKEWGIFQLVNHGIPVAAIEELKRVGTE
OsFLS_FLS_Oryza_sativa PEIPVIDMA............APESAARVAEAAAEWGLFQVVNHGVPAAAVAELQRVGRE
AFS63900.1_FLS_Narcissus_tazetta PEIPVVDLS.........TADRTRLTRAIADASEEWGIFQIVNHGIPVEVVKELQRVGKE
AEC33115.1_FLS_Fagopyrum_esculentum LEVPAIDIN.........ESNETSPVESIKKASEEWGLFQVVNHGIPIEVISHLQRVGKE
ABE28017.1_FLS_Nicotiana_tabacum LQVPVIDI...........DDKNVVKLISDASKEWGIFQVINHGIPDEVIANLQKVGKE
BAE75809.1_FLS_Vitis_vinifera PQVPTIDFS.........DPDEANLTRLIAEASMEWGIFQIVNHGIPFHVITSLQKVGRE
AAX89401.1_FLS_Malus_domestica LEVPIIDFS.........DPDEEKLIVQITEASSNWGMYQIVNHDIPSEVISKLQAVGKE
CAA80264.1_FLS_Petunia_x_hybrida LQVPVIDLR.........DPDENKMVKLIADASKEWGIFQLINHGIPDEAIADLQKVGKE
ADZ28516.1_FLS_Camellia_nitidissima LEVPVIDLS........NDSDDQNIVQMVAEACRDWGIFQVVNHGIPDEVIGNLQRVGKD
LcFLS_FLS_Litchi_chinensis AEIPTIDLS.........DPDTEKVERLMAEASREWGIFQVVNHGIPDDVIRKLQAVGKE
ABB53382.1_FLS_Antirrhinum_majus LEVPVIDLS........DDFNEKILVKNISEASRDWGIFQVVNHGISNEVISKLQKVGKE
AAP57395.1_FLS_Petroselinum_crispum LQVPVIDLG.......SSNNTEENLVELIAEASREWGIFQVVNHGIPDDAIAKLQKVGKE
BAC10995.1_flavonol_synthase_Nierembergia_sp LQVPVIDLAPRVVGDEQHDHDDVEVVKQIADASKEWGIFQVINHGIPNDVIADLQKVGKE
acc

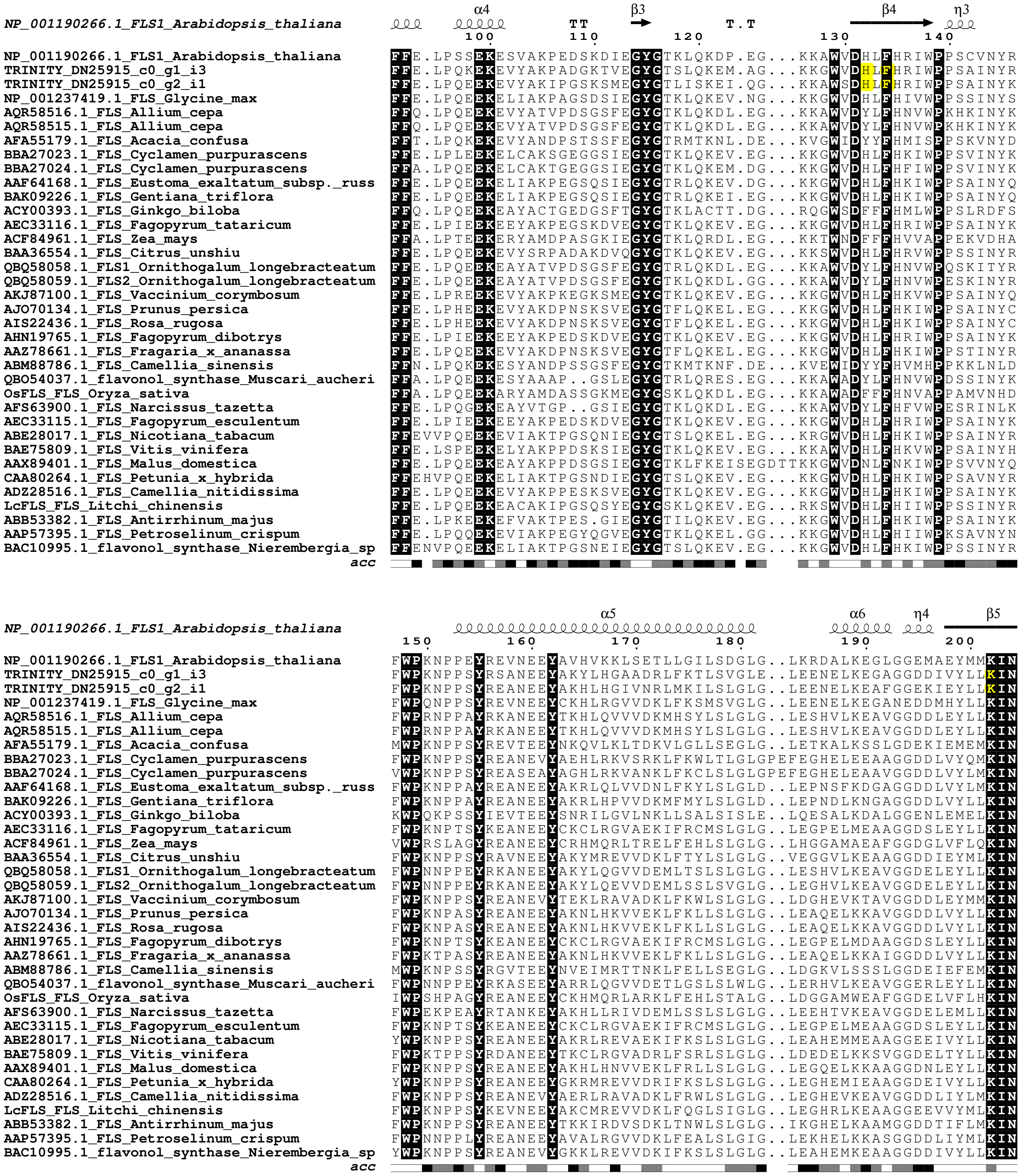

NP_001190266.1_FLS1_Arabidopsis_thaliana
α4 TT β3 T.T β4 η3
100 110 120 130 140
NP_001190266.1_FLS1_Arabidopsis_thaliana FFE.LPSSEKESVAKPEDSKDIEGYGTKLQKDP.EG...KKAWVDHLFHRIWPPSCVNYR
TRINITY_DN25915_c0_g1_i3 FFE.LPQKEKEVYAKPADGKTVEGYGTSLQKEM.AG...KKGWVDHLFHRIWPPSAINYR
TRINITY_DN25915_c0_g2_i1 FFE.LPQEEKEVYAKIPGSKSMEGYGTLISKEI.QG...KKAWSDHLFHRIWPPSAINYK
NP_001237419.1_FLS_Glycine_max FFE.LPQEEKELIAKPAGSDSIEGYGTKLQKEV.NG...KKGWVDHLFHIVWPPSSINYS
AQR58516.1_FLS_Allium_cepa FFQ.LPQEEKEVYATVPDSGSFEGYGTKLQKDL.EG...KKAWVDYLFHNVWPKHKINYK
AQR58515.1_FLS_Allium_cepa FFQ.LPQEEKEVYATVPDSGSFEGYGTKLQKDL.EG...KKAWVDYLFHNVWPKHKINYK
AFA55179.1_FLS_Acacia_confusa FFT.LPQKEKEVYANDPSTSSFEGYGTRMTKNL.DE...KVGWIDYYFHMISPPSKVNYD
BBA27023.1_FLS_Cyclamen_purpurascens FFE.LPLEEKELCAKSGEGGSIEGYGTKLQKEV.EG...KKGWVDHLFHKIWPPSVINYK
BBA27024.1_FLS_Cyclamen_purpurascens FFA.LPQEEKELCAKTGEGGSIEGYGTKLQKEM.DG...KKGWVDHLFHFIWPPSVINYK
AAF64168.1_FLS_Eustoma_exaltatum_subsp._russ FFE.LPQEEKELIAKPEGSQSIEGYGTKLQKEV.DG...KKGWVDHLFHKIWPPSAINYQ
BAK09226.1_FLS_Gentiana_triflora FFE.LPQEEKELIAKPEGSQSIEGYGTRLQKEI.EG...KKGWVDHLFHKIWPPSAINYQ
ACY00393.1_FLS_Ginkgo_biloba FFQ.LPQEEKEAYACTGEDGSFTGYGTKLACTT.DG...RQGWSDFFFHMLWPPSLRDFS
AEC33116.1_FLS_Fagopyrum_tataricum FFE.LPIEEKEEYAKPEDSKDVEGYGTKLQKEV.EG...KKGWVDHLFHRIWPPSAINYK
ACF84961.1_FLS_Zea_mays FFA.LPTEEKERYAMDPASGKIEGYGTKLQRDL.EG...KKTWNDFFFHVVAPPEKVDHA
BAA36554.1_FLS_Citrus_unshiu FFE.LPQEEKEVYSRPADAKDVQGYGTKLQKEV.EG...KKSWVDHLFHRVWPPSSINYR
QBQ58058.1_FLS1_Ornithogalum_longebracteatum FFE.LPQEEKEAYATVPDSGSFEGYGTRLQKDL.EG...KKAWVDYLFHNVWPQSKITYR
QBQ58059.1_FLS2_Ornithogalum_longebracteatum FFE.LPQEEKEAYATVPDSGSFEGYGTKLQKDL.GG...KKAWVDYLFHNVWPESKINYR
AKJ87100.1_FLS_Vaccinium_corymbosum FFE.LPREEKEVYAKPKEGKSMEGYGTKLQKEV.EG...KKGWVDHLFHKVWPPSAINYQ
AJO70134.1_FLS_Prunus_persica FFE.LPHEEKEVYAKDPNSKSVEGYGTFLQKEL.EG...KKGWVDHLFHKIWPPSAINYC
AIS22436.1_FLS_Rosa_rugosa FFE.LPHEEKEVYAKDPNSKSVEGYGTFLQKEL.EG...KKGWVDHLFHKIWPPSAINYC
AHN19765.1_FLS_Fagopyrum_dibotrys FFE.LPIEEKEEYAKPEDSKDVEGYGTKLQKEV.EG...KKGWVDHLFHRIWPPSAINYK
AAZ78661.1_FLS_Fragaria_x_ananassa FFE.LPQEEKEVYAKDPNSKSVEGYGTFLQKEL.EG...KKGWVDHLFHKIWPPSTINYR
ABM88786.1_FLS_Camellia_sinensis FFN.LPQKEKESYANDPSSGSFEGYGTKMTKNF.DE...KVEWIDYYFHVMHPPKKLNLD
QBO54037.1_flavonol_synthase_Muscari_aucheri FFA.LPQEEKESYAAAP..GSLEGYGTRLQRES.EG...KKAWADYLFHNVWPDSSINYK
OsFLS_FLS_Oryza_sativa FFA.LPQEEKARYAMDASSGKMEGYGSKLQKDL.EG...KKAWADFFFHNVAPPAMVNHD
AFS63900.1_FLS_Narcissus_tazetta FFE.LPQGEKEAYVTGP..GSIEGYGTKLQKDL.EG...KKAWVDYLFHFVWPESRINLK
AEC33115.1_FLS_Fagopyrum_esculentum FFE.LPIEEKEEYAKPEDSKDVEGYGTKLQKEV.EG...KKGWVDHLFHRIWPPSAINYK
ABE28017.1_FLS_Nicotiana_tabacum FFEVVPQEEKEVIAKTPGSQNIEGYGTSLQKEL.EG...KRGWVDHLFHKIWPPSAINYR
BAE75809.1_FLS_Vitis_vinifera FFE.LSPEEKELYAKPPDSKSIEGYGTKLQKEV.EG...KKAWVDHLFHKVWPPSAINYH
AAX89401.1_FLS_Malus_domestica FFE.LPQEEKEAYAKPPDSGSIEGYGTKLFKEISEGDTTKKGWVDNLFNKIWPPSVVNYQ
CAA80264.1_FLS_Petunia_x_hybrida FFEHVPQEEKELIAKTPGSNDIEGYGTSLQKEV.EG...KKGWVDHLFHKIWPPSAVNYR
ADZ28516.1_FLS_Camellia_nitidissima FFE.LPQEEKEVYAKPPESKSVEGYGTKLQKEV.EG...KKGWVDHLFHKIWPPSAINYK
LcFLS_FLS_Litchi_chinensis FFE.LPQEEKEACAKIPGSQSYEGYGSKLQKEV.EG...RKSWVDHLFHIIWPPSSINYR
ABB53382.1_FLS_Antirrhinum_majus FFE.LPKEEKEFVAKTPES.GIEGYGTILQKEV.EG...KKGWVDHLFHKIWPPSAINYK
AAP57395.1_FLS_Petroselinum_crispum FFE.LPQQEKEVIAKPEGYQGVEGYGTKLQKEL.GG...KKGWVDHLFHIIWPKSAVNYN
BAC10995.1_flavonol_synthase_Nierembergia_sp FFENVPQEEKELIAKTPGSNEIEGYGTSLQKEV.EG...KKGWVDHLFHKIWPPSSINYR
acc

NP_001190266.1_FLS1_Arabidopsis_thaliana
α5 α6 η4 β5
150 160 170 180 190 200
NP_001190266.1_FLS1_Arabidopsis_thaliana FWPKNPPEYREVNEEYAVHVKKLSETLLGILSDGLG..LKRDALKEGLGGEMAEYMMKIN
TRINITY_DN25915_c0_g1_i3 FWPKNPPSYRSANEEYAKYLHGAADRLFKTLSVGLE..LEENELKEGAGGDDIVYLLKIN
TRINITY_DN25915_c0_g2_i1 FWPKNPPSYREANEEYAKHLHGIVNRLMKILSLGLG..LEENELKEAFGGEKIEYLLKIN
NP_001237419.1_FLS_Glycine_max FWPQNPPSYREVNEEYCKHLRGVVDKLFKSMSVGLG..LEENELKEGANEDDMHYLLKIN
AQR58516.1_FLS_Allium_cepa FWPRNPPAYRKANEEYTKHLQVVVDKMHSYLSLGLG..LESHVLKEAVGGDDLEYLLKIN
AQR58515.1_FLS_Allium_cepa FWPRNPPAYRKANEEYTKHLQVVVDKMHSYLSLGLG..LESHVLKEAVGGDDLEYLLKIN
AFA55179.1_FLS_Acacia_confusa MWPKNPPSYREVTEEYNKQVLKLTDKVLGLLSEGLG..LETKALKSSLGDEKIEMEMKIN
BBA27023.1_FLS_Cyclamen_purpurascens FWPKNPPSYREANEVYAEHLRKVSRKLFKWLTLGLGPEFEGHELEEAAGGDDLVYQMKIN
BBA27024.1_FLS_Cyclamen_purpurascens VWPKNPPSYREASEAYAGHLRKVANKLFKCLSLGLGPEFEGHELEAAVGGDDLVYLMKIN
AAF64168.1_FLS_Eustoma_exaltatum_subsp._russ FWPKNPPAYREANEEYAKRLQLVVDNLFKYLSLGLD..LEPNSFKDGAGGDDLVYLMKIN
BAK09226.1_FLS_Gentiana_triflora FWPKNPPAYREANEEYAKRLHPVVDKMFMYLSLGLD..LEPNDLKNGAGGDDLVYLLKIN
ACY00393.1_FLS_Ginkgo_biloba KWPQKPSSYIEVTEEYSNRILGVLNKLLSALSISLE..LQESALKDALGGENLEMELKIN
AEC33116.1_FLS_Fagopyrum_tataricum FWPKNPTSYKEANEEYCKCLRGVAEKIFRCMSLGLG..LEGPELMEAAGGDSLEYLLKIN
ACF84961.1_FLS_Zea_mays VWPRSLAGYREANEEYCRHMQRLTRELFEHLSLGLG..LHGGAMAEAFGGDGLVFLQKIN
BAA36554.1_FLS_Citrus_unshiu FWPKNPPSYRAVNEEYAKYMREVVDKLFTYLSLGLG..VEGGVLKEAAGGDDIEYMLKIN
QBQ58058.1_FLS1_Ornithogalum_longebracteatum FWPNNPPEYRKANEEYAKYLQGVVDEMLTSLSLGLG..LESHVLKEAVGGDELEYLLKIN
QBQ58059.1_FLS2_Ornithogalum_longebracteatum FWPNNPPEYRKANEEYAKYLQEVVDEMLSSLSVGLG..LEGHVLKEAVGGDELEYLLKIN
AKJ87100.1_FLS_Vaccinium_corymbosum FWPKNPPSYREANEVYTEKLRAVADKLFKWLSLGLG..LDGHEVKTAVGGDDLEYMMKIN
AJO70134.1_FLS_Prunus_persica FWPKNPASYREANEEYAKNLHKVVEKLFKLLSLGLG..LEAQELKKAVGGDDLVYLLKIN
AIS22436.1_FLS_Rosa_rugosa FWPKNPASYREANEEYAKNLHKVVEKLFKLLSLGLG..LEAQELKKAVGGDDLVYLLKIN
AHN19765.1_FLS_Fagopyrum_dibotrys FWPKNPTSYKEANEEYCKCLRGVAEKIFRCMSLGLG..LEGPELMDAAGGDSLEYLLKIN
AAZ78661.1_FLS_Fragaria_x_ananassa FWPKTPASYREANEEYAKNLHKVVEKLFKLLSLGLG..LEAQELKKAIGGDDLVYLLKIN
ABM88786.1_FLS_Camellia_sinensis MWPKNPSSYRGVTEEYNVEIMRTTNKLFELLSEGLG..LDGKVLSSSLGGDEIEFEMKIN
QBO54037.1_flavonol_synthase_Muscari_aucheri FWPNNPPEYRKASEEYARRLQGVVDETLGSLSLWLG..LERHVLKEAVGGERLEYLLKIN
OsFLS_FLS_Oryza_sativa IWPSHPAGYREANEEYCKHMQRLARKLFEHLSTALG..LDGGAMWEAFGGDELVFLHKIN
AFS63900.1_FLS_Narcissus_tazetta FWPEKPEAYRTANKEYAKHLLRIVDEMLSSLSLGLG..LEEHTVKEAVGGDELEMLLKIN
AEC33115.1_FLS_Fagopyrum_esculentum FWPKNPTSYKEANEEYCKCLRGVAEKIFRCMSLGLG..LEGPELMEAAGGDSLEYLLKIN
ABE28017.1_FLS_Nicotiana_tabacum YWPKNPPSYREANEEYAKRLREVAEKIFKSLSLGLG..LEAHEMMEAAGGEEIVYLLKIN
BAE75809.1_FLS_Vitis_vinifera FWPKTPPSYRDANEEYTKCLRGVADRLFSRLSLGLG..LDEDELKKSVGGDELTYLLKIN
AAX89401.1_FLS_Malus_domestica FWPKNPPSYREANEEYAKHLHNVVEKLFRLLSLGLG..LEGQELKKAAGGDNLEYLLKIN
CAA80264.1_FLS_Petunia_x_hybrida YWPKNPPSYREANEEYGKRMREVVDRIFKSLSLGLG..LEGHEMIEAAGGDEIVYLLKIN
ADZ28516.1_FLS_Camellia_nitidissima FWPKNPPSYREANEVYAERLRAVADKLFRWLSLGLG..LEGHEVKEAVGGDDLIYLMKIN
LcFLS_FLS_Litchi_chinensis FWPKNPPSYKEVNEEYAKCMREVVDKLFQGLSIGLG..LEGHRLKEAAGGEEVVYMLKIN
ABB53382.1_FLS_Antirrhinum_majus FWPKNPPSYREANEEYTKKIRDVSDKLTNWLSLGLG..IKGHELKAAMGGDDTIFLMKIN
AAP57395.1_FLS_Petroselinum_crispum FWPNNPPLYREANEEYAVALRGVVDKLFEALSLGIG..LEKHELKKASGGDDLIYMLKIN
BAC10995.1_flavonol_synthase_Nierembergia_sp YWPKNPPSYREANEVYGKKLREVVDKIFKSLSLGLG..LEAHEMKEAAGGDDIVYLLKIN
acc

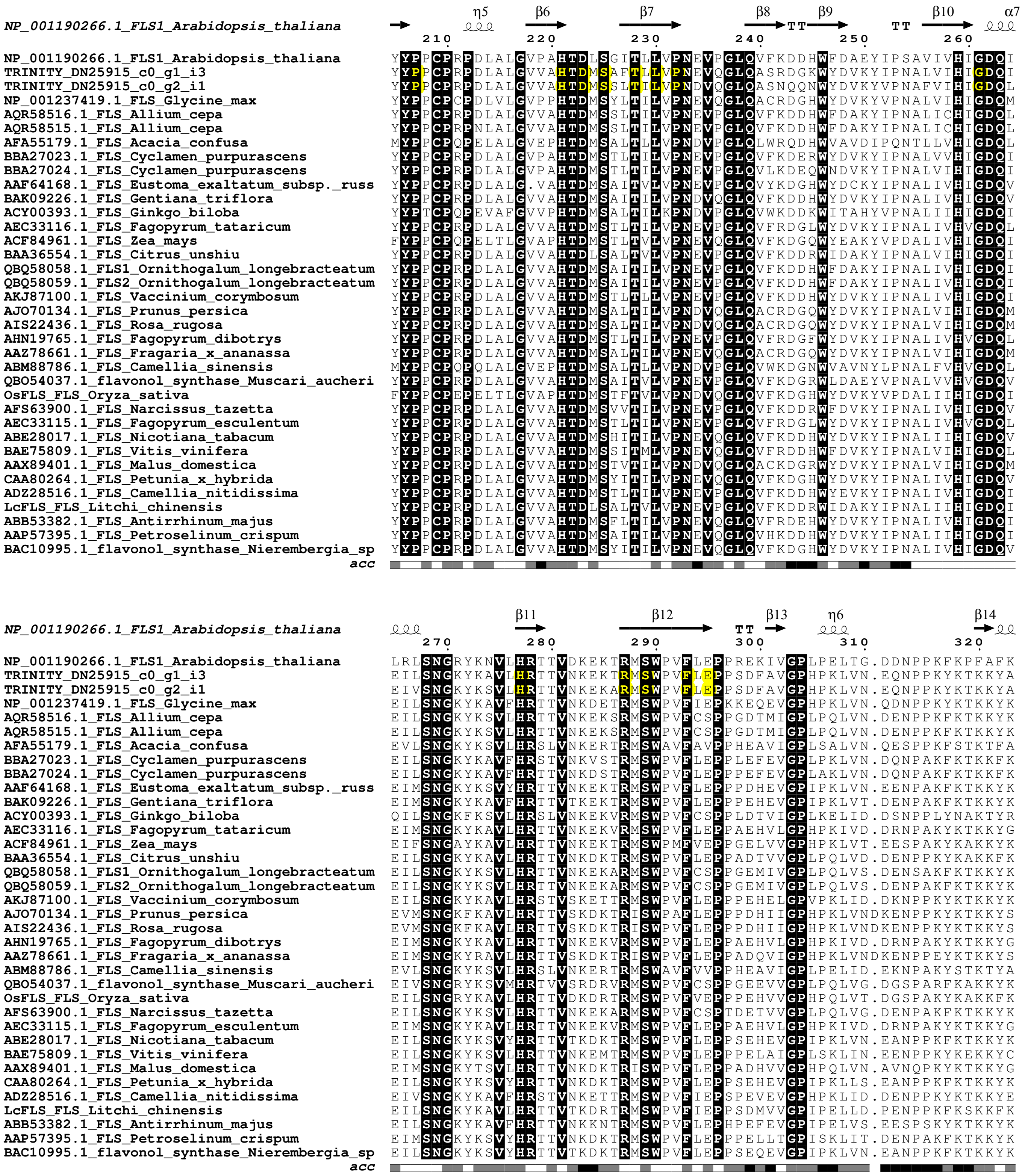
NP_001190266.1_FLS1_Arabidopsis_thaliana
η5 β6 β7 β8 TT β9 TT β10 α7
210 220 230 240 250 260
acc
β11 β12 TT β13 η6 β14
270 280 290 300 310 320
acc

| | |
|---|---|
| *NP_001190266.1_FLS1_Arabidopsis_thaliana* | α8 TT |
| | 330 |
| NP_001190266.1_FLS1_Arabidopsis_thaliana | DYSYRKLNKLPLD |
| TRINITY_DN25915_c0_g1_i3 | DYCYCKLNKIPQ. |
| TRINITY_DN25915_c0_g2_i1 | DYCYCKLNKIPQ. |
| NP_001237419.1_FLS_Glycine_max | DYAYCKLNKIPQ. |
| AQR58516.1_FLS_Allium_cepa | DYAYCKINKLPQ. |
| AQR58515.1_FLS_Allium_cepa | DYAYCKINKLPQ. |
| AFA55179.1_FLS_Acacia_confusa | EFRHRKFNKLPQ. |
| BBA27023.1_FLS_Cyclamen_purpurascens | DYLYCKLNKLPQ. |
| BBA27024.1_FLS_Cyclamen_purpurascens | DYFYCKLNKLPQ. |
| AAF64168.1_FLS_Eustoma_exaltatum_subsp._russ | DYAYCKLNKLPQ. |
| BAK09226.1_FLS_Gentiana_triflora | DYAYCKLNKLPQ. |
| ACY00393.1_FLS_Ginkgo_biloba | EYKHRKINKLGQ. |
| AEC33116.1_FLS_Fagopyrum_tataricum | DYMYCKLNKLPQ. |
| ACF84961.1_FLS_Zea_mays | DYQHCKINKLPM. |
| BAA36554.1_FLS_Citrus_unshiu | DYSYCKLNKLPQ. |
| QBQ58058.1_FLS1_Ornithogalum_longebracteatum | DYSYCKINKLPQ. |
| QBQ58059.1_FLS2_Ornithogalum_longebracteatum | EYSYCKINKLPQ. |
| AKJ87100.1_FLS_Vaccinium_corymbosum | DYVYCKLNKLPQ. |
| AJO70134.1_FLS_Prunus_persica | EYVYNKLNKIPQ. |
| AIS22436.1_FLS_Rosa_rugosa | EYVYNKLNKIPQ. |
| AHN19765.1_FLS_Fagopyrum_dibotrys | DYMYCKLNKLPQ. |
| AAZ78661.1_FLS_Fragaria_x_ananassa | EYVYNKLNKMPQ. |
| ABM88786.1_FLS_Camellia_sinensis | EYRHRKFNKIPQ. |
| QBO54037.1_flavonol_synthase_Muscari_aucheri | DYSYCKINKLPQ. |
| OsFLS_FLS_Oryza_sativa | DYRHCKINKLPM. |
| AFS63900.1_FLS_Narcissus_tazetta | DYCYCKLNKLPQ. |
| AEC33115.1_FLS_Fagopyrum_esculentum | DYMYCKLNKLPQ. |
| ABE28017.1_FLS_Nicotiana_tabacum | DYVYCKLNKLPQ. |
| BAE75809.1_FLS_Vitis_vinifera | DYVYCKLNKIPQ. |
| AAX89401.1_FLS_Malus_domestica | DYVYCKINKLPQ. |
| CAA80264.1_FLS_Petunia_x_hybrida | DYVYCKLNKLPQ. |
| ADZ28516.1_FLS_Camellia_nitidissima | DYMYCKLNKLPQ. |
| LcFLS_FLS_Litchi_chinensis | DYSYCKLSKIP.. |
| ABB53382.1_FLS_Antirrhinum_majus | DYVYCKLNKIPQ. |
| AAP57395.1_FLS_Petroselinum_crispum | DYVYCKLNKLPQ. |
| BAC10995.1_flavonol_synthase_Nierembergia_sp | DYVYCKLNKLPQ. |
| *acc* | |

```
NP_001241007.1_FNS2_Glycine_max
                                                     1            10        20
NP_001241007.1_FNS2_Glycine_max                      .MISESL..............................LLVFLIVFISASLLKLL...FVR
TRINITY_DN252_c0_g1_i1                               .MSSFLL.........YFI..................LFFISIYFLSSLRSFLR......
AAD39549.1_FNS2_Gerbera_hybrid_cultivar              .MNTLQL................................IFLLFFFPTLLFLY...CLPY
AGA17938.1_FNS2_Dahlia_pinnata                       .MNTLLVLQMV.........................IPAIIAFVIFHL.LFL........
AWX67430.1_FNS2_Salvia_miltiorrhiza                  .MELVEM..............................GAYAALFLVSAALSR.....SIL
ACB56919.1_FNS2_Pilosella_officinarum                .MNIFVLFQSL.........................SPAVIAAVVLPS.LFLY....L.L
AIS92511.1_FNS2_Epimedium_sagittatum                 ..MVLDL..............................VLYTIIFFLSALLVR.....IIS
BAB59004.1_FNS2_Perilla_frutescens_var._crispa       ......M..............................ALYAALFLLSAAVVR.....SVL
AMW91728.1_FNS2_Scutellaria_baicalensis              .MDLVEV..............................TLYAALFLLSAAFLL.....LIF
AMW91729.1_FNS2_Scutellaria_baicalensis              ....MEV..............................TLNVALLLLSAAVCL.....MVF
BAA84072.1_FNS2_Torenia_hybrid_cultivar              .MDTVLI..............................TLYTALFVITTTFL......LLL
ACV65037.1_FNS2_Glycine_max                          .MISESL..............................LVVFLIVFISASLLKLL...FVR
BAA84071.1_FNS2_Antirrhinum_majus                    .MSTL.................................VYSTLFILSTLLLT.....LLT
BAF49323.1_FNS2_Lobelia_erinus                       .....................................VAIPTIFLFFTSLFF.....LQR
BAD91809.1_FNS2_Gentiana_triflora                    .MMLLDF..............................FYSASIFVLSILLFR.....AIY
AAF04115.1_FNS2_Callistephus_chinensis               .MNIFEVFQSV.........................SPAIIAIFFISS.LFIY....LVL
ACH99109.1_FNS2_Camellia_sinensis                    ...MFDL...I.........................SIATLFFVIISTTILLLSINHFKK
XP_002461286.1_FNS2_Sorghum_bicolor                  ....MEAAEAV..........TVGVGGSIGAPSAPGGVLLLFLLALSTLIIIIR...WRW
sp|P93149.2_FNS2_Glycyrrhiza_echinata                ......M.............................EPQLVAVSVLVSALICY....FFF
BAG94859.1_FNS2_Oryza_sativa_Japonica_Group          MASLMEVQVPL...........LGMGTTMGA.........LAL..ALVVVVVVH...VAV
ABC86159.1_FNS2_Medicago_truncatula                  ......M.............................EPLLLAFTLFLSSLICY....IIF
XP_008663013.1_FNS2_Zea_mays                         ....MEEQQPRPRPSIMFVLSSLAKNNPESV.........LALIAVLTVVALRH...LIS
                                                acc

NP_001241007.1_FNS2_Glycine_max
                                                                        η1       α1          β1  T.T β2
                                                       30        40        50        60        70        80
NP_001241007.1_FNS2_Glycine_max                      ENKPKAH.LKNPPSP.PAIPIIGHLHLLKPLIHHSFRDLSLRYGPLLSLRIG.SVKFIVA
TRINITY_DN252_c0_g1_i1                               ........RRLPPGP.KPWPIVGNLPHLGPMPHHSIAALARTYGPLMHLRMG.FVDVVVA
AAD39549.1_FNS2_Gerbera_hybrid_cultivar              KRNQNHR..RLPPSP.PSFPIIGHLHHLGPLIHQSFHALSTRYGSLIHLRLG.SVPCVVV
AGA17938.1_FNS2_Dahlia_pinnata                       .KSKPNR..RLPPSP.PSLPIIGHLHHLGPLIHQSFNRLSARYGPLIHLRLG.SVSCVVA
AWX67430.1_FNS2_Salvia_miltiorrhiza                  RSKLRR....HAPGP.FPLPIIGHLHLLGPRLHQSFHELSQRYGPLMQLRLG.SIKCVVA
ACB56919.1_FNS2_Pilosella_officinarum                SKSKQNH..RLPPSP.PSLPIIGHLHHLGPLIHQSFHNLSTRYGPLIHLRLG.SVPCVVA
AIS92511.1_FNS2_Epimedium_sagittatum                 GAIKPIR.SRLPPSP.ISLPIIGHLHLLDIVPYKSFHKLATKYGPFFHLRLG.SVPCIVI
BAB59004.1_FNS2_Perilla_frutescens_var._crispa       DRKRGRP..PYPPGP.FPLPIIGHLHLLGPRLHQTFHDLSQRYGPLMQLRLG.SIRCVIA
AMW91728.1_FNS2_Scutellaria_baicalensis              AGDRSS.....PPGP.FPLPIIGHLHLLGPKLHQSFHGLSQRHGPLMQIRLG.SINCVVA
AMW91729.1_FNS2_Scutellaria_baicalensis              TGKRRRR.LPNPPGP.FPLPLIGNLNLVSPRLHHTFHMLAQRYGPIMKFRLG.SIPCLVV
BAA84072.1_FNS2_Torenia_hybrid_cultivar              RRRGPP....SPPGP.LSLPIIGHLHLLGPRLHHTFHEFSLKYGPLIQLKLG.SIPCVVA
ACV65037.1_FNS2_Glycine_max                          ENKPKAH.LKNPPSP.PAIPIIGHLHLLKPLIHHSFRDLSLRYGPLLSLRIG.SVKFIVA
BAA84071.1_FNS2_Antirrhinum_majus                    RTRRKT....RPPGP.LALPLIGHLHLLGPKLHHTFHQFSQRYGPLIQLYLG.SVPCVVA
BAF49323.1_FNS2_Lobelia_erinus                       TKSKTNH.LPLPPSP.WALPIIGHLHHLGPLIHHSFHDLSSRYGPLIHLRLG.SVPCVVA
BAD91809.1_FNS2_Gentiana_triflora                    TTKNRRL..RLPPSP.FGLPIIGHLHLLGPKIHHSFHNLYKRYGPIFHLRLG.SNRCIVV
AAF04115.1_FNS2_Callistephus_chinensis               IRNQKSL..SLPPSP.PALPIIGHLHHLGPLIHHSFHDLSTRYGPLIHLRLG.SVPCVVA
ACH99109.1_FNS2_Camellia_sinensis                    PPHLRRR.LSLPPTP.FALPIIGHLHLLGPIIHRSFHDLSSRYGPLFHLRLG.SVPCFVV
XP_002461286.1_FNS2_Sorghum_bicolor                  NNNRNSR...LPPSP.MALPLVGHLHLIRSPPHRSLDRIVKRYGPLVYLRLGPSTHCVVA
sp|P93149.2_FNS2_Glycyrrhiza_echinata                RPYFHRYGKNLPPSPFFRLPIIGHMHMLGPLLHQSFHNLSHRYGPLFSLNFG.SVLCVVA
BAG94859.1_FNS2_Oryza_sativa_Japonica_Group          NAFGRRR...LPPSP.ASLPVIGHLHLLRPPVHRTFHELAARLGPLMHVRLG.STHCVVA
ABC86159.1_FNS2_Medicago_truncatula                  QPILNRH.KNLPPSPLFKLPIIGHMHMLGPLLHHSFDRLSQKYGPIFSLNFG.SVLCVVA
XP_008663013.1_FNS2_Zea_mays                         SWRQQAR...LPPSP.TSLPVIGHLHLLRPPVHRTFQELASRIGPLMHIRLG.STHCVVA
                                                acc

NP_001241007.1_FNS2_Glycine_max
                                                       α2         α3         α4      α5                 α6
                                                         90        100       110       120       130       140
NP_001241007.1_FNS2_Glycine_max                      STPSLAQEFLKTNELTYSSRKMNMAINMVTYHNATFAFAPYDTYWKFMKKLSTTELLGNK
TRINITY_DN252_c0_g1_i1                               ASAAVASQFLKFHDANFSSRPPNSGAKYVAYNYQDLVFAPYGPRWRMLRKISSVNLFSGK
AAD39549.1_FNS2_Gerbera_hybrid_cultivar              STPDLAKDFLKTNELAFSSRKHSLAIDHITY.GVAFAFAPYGTYWKFIKKLFTVELLGTQ
AGA17938.1_FNS2_Dahlia_pinnata                       DAPDLAQELLQKNDLAFADRKHTLAIDHVTY.GVAFAFAPYGPYWRFVKKMSTVELLGIQ
AWX67430.1_FNS2_Salvia_miltiorrhiza                  SSPELAKEFLKTHDLVFSSRKHSTAVDIVTY.DSSFAFSPLGPYWKFIKKLCTYELLGAR
ACB56919.1_FNS2_Pilosella_officinarum                STPDLARDFLKTNELAFSSRKHSLAIDHITY.GVAFAFAPYGPYWKFIKKLSTVELLGNQ
AIS92511.1_FNS2_Epimedium_sagittatum                 SSPELTKELMATNELTFAARPVTMAIDHLTYNSSAFAFAPYGAQWKFMKKICMTELLSGR
BAB59004.1_FNS2_Perilla_frutescens_var._crispa       ASPELAKECLKTHELVFSSRKHSTAIDIVTY.DSSFAFSPYGPYWKFIKKLCTYELLGAR
AMW91728.1_FNS2_Scutellaria_baicalensis              STPELAKEFLKTNELVFSSRKHSTAIDIVTY.NSSFAFSPYGPYWKYIKKLCTYELLGAR
AMW91729.1_FNS2_Scutellaria_baicalensis              STPELAKDILKTHELIFSSRVKSTAIDIVTY.GVSFAFSPYGPYWKYIKKLCTYELLGSR
BAA84072.1_FNS2_Torenia_hybrid_cultivar              STPELAREFLKTNELAFSSRKHSTAIDIVTY.DSSFAFSPYGPYWKYIKKLCTYELLGAR
ACV65037.1_FNS2_Glycine_max                          STPSLAQEFLKTNELTYSSRKMNMAINMVTYHNATFAFAPYDTYWKFMKKLSTTELLGNK
BAA84071.1_FNS2_Antirrhinum_majus                    STPELAREFLKTHELDFSSRKHSTAIDIVTY.DSSFAFAPYGPYWKFIKKLCTYELLGAR
BAF49323.1_FNS2_Lobelia_erinus                       STPELARDFLKTNELTFSSRKHSAAIKRLSY.DVAFAFAPYGPYWKFIKKMSTFELLGVR
BAD91809.1_FNS2_Gentiana_triflora                    STPELAKEFLKTHELDFAYRKNSSAISLLTY.HVSFAFAPYGPYWKYIKKITTYQLLGNR
AAF04115.1_FNS2_Callistephus_chinensis               STPDLARDFLKTNELAFSSRKHSLAIDHVTY.GVSFAFAPYGPYWKFIKKTSIVELLGNQ
ACH99109.1_FNS2_Camellia_sinensis                    STPELAKEFLLTHELKFSSRRDSIAIQRLTY.DSAFAFAPYGPYWKFLKKLCTCDLLGAR
XP_002461286.1_FNS2_Sorghum_bicolor                  GTADAARDLLK.HEASIPERPITVVAHHLAYGDAGFAFAPYGAHWRFMKRLCMSELLGPR
sp|P93149.2_FNS2_Glycyrrhiza_echinata                STPHFAKQLLQTNELAFNCRIESTAVKKLTY.ESSLAFAPYGDYWRFIKKLSMNELLGSR
BAG94859.1_FNS2_Oryza_sativa_Japonica_Group          SSAEVAAELIRSHEAKISERPLTAVARQFAYESAGFAFAPYSPHWRFMKRLCMSELLGPR
ABC86159.1_FNS2_Medicago_truncatula                  STPHYAKQILQINEHAFNCRNESTAIKRLTY.EASLAFAPYGEYWRFIKKLSMNELLGSR
XP_008663013.1_FNS2_Zea_mays                         STPEVASELIRGHEGSISERPLTAVARQFAYDSAGFAFAPYNTHWRFMKRLCMSELLGPR
                                                acc
```

```
                                                   α7                        β3        α8
NP_001241007.1_FNS2_Glycine_max
                                                    150       160       170       180       190
NP_001241007.1_FNS2_Glycine_max                  TLGHFLPIRTREVHDIIQFLFHKSKAQESVNLTEALLSLSNNVISQMMLSIK.....SSG
TRINITY_DN252_c0_g1_i1                           ALDDFRHIRQHEVSVLTRAL..ASAGSTPVNLGQLLNVCTANALGRVMIGRRVFGDGSGG
AAD39549.1_FNS2_Gerbera_hybrid_cultivar          NLSHFLPIRTHEIRELLRTLMVKSRAKERVNLTEELLKLTNNVISQMMMSIR.....CSG
AGA17938.1_FNS2_Dahlia_pinnata                   NLGHFLPIRTQEIHGLLLTLTEKSKQNESVNMTNELLKLSNNIICQMMMGIR.....CSG
AWX67430.1_FNS2_Salvia_miltiorrhiza              NLLHFQPVRTLEVNTFLQILLRKGKSGESFNLTEELVKLTSNVISHMMLSIR.....CSG
ACB56919.1_FNS2_Pilosella_officinarum            NLGHFLPIRTQEIHELLHTLMEKSKRKESVNLTEELLKLTNNVICQMMMSIR.....CSG
AIS92511.1_FNS2_Epimedium_sagittatum             ALNQLYPIRKEELLLLVQVLVHKSSEGETVNLSEALVNISNNIISRMMWGAT.....YSG
BAB59004.1_FNS2_Perilla_frutescens_var._crispa   NLAHFQPIRTLEVKSFLQILMRKGESGESFNVTEELVKLTSNVISHMMLSIR.....CSE
AMW91728.1_FNS2_Scutellaria_baicalensis          NLHHFQPIRTFEVHTFLRLLMEKSESGESFNVTEELIKLTSNVMSNMMLGTR.....CSA
AMW91729.1_FNS2_Scutellaria_baicalensis          MLNHFEPLRALEVREFLKDVMAMGKAGKSFNVTEELMKLTSNVMSNMMLSIR.....AAE
BAA84072.1_FNS2_Torenia_hybrid_cultivar          NLGHFQPIRNLEVRSFLQLLMHKSFKGESVNVTDELVRLTSNVISHMMLSIR.....CSE
ACV65037.1_FNS2_Glycine_max                      TLGHFLPIRTREVHDIIQFLFHKSKAQESVNLTEALLSLSNNVISQMMLSIK.....SSG
BAA84071.1_FNS2_Antirrhinum_majus                NLSHFQPIRALEVNSFLRILYEKTEQKQSVNVTEELVKLTSNVISNMMLGIR.....CSG
BAF49323.1_FNS2_Lobelia_erinus                   NLSHFLPIRTREIHGLIHLLMKKSKGKERVNVTEELVRLTNNVISQMMMSIR.....CSG
BAD91809.1_FNS2_Gentiana_triflora                NLTHFEPIRRLETNRVLYDLMVSSKHGKSVNLTEEMIKLTSNIISQMMLSIR.....CSD
AAF04115.1_FNS2_Callistephus_chinensis           NLSNFLPIRTQEVHELLQTLMVKSKKNESVNLSEELLKLTNNVICQMMMSIR.....CSG
ACH99109.1_FNS2_Camellia_sinensis                SINHFLPTRTRELHCFVRLLIDKAVACEPVNITKELSTLANNIISQMMIGVR.....CSG
XP_002461286.1_FNS2_Sorghum_bicolor              TVDQLRPVREAELAAVLGAARQASASGERIDVSRYLISMSNNAIMRMVASAL.....PG.
sp|P93149.2_FNS2_Glycyrrhiza_echinata            SINNFQHLRAQETHQLLRLLSNRARAFEAVNITEELLKLTNNVISIMMV...........
BAG94859.1_FNS2_Oryza_sativa_Japonica_Group      TVEQLRPVRRAGLVSLLRHVL.SQPEAEAVDLTRELIRMSNTSIIRMAASTV.....PS.
ABC86159.1_FNS2_Medicago_truncatula              SISSFQHLRLQETHNLLKLFADKAKNYEAVNVTQELLKLSNNVISKMML...........
XP_008663013.1_FNS2_Zea_mays                     TVEQLRPVRRAGTVSLLADLLASSARGETVDLTRHLIRLSNTSIIRMVASTV.....PG.
                                             acc
```

```
                                                        α9             α10    α11   η2            α12
NP_001241007.1_FNS2_Glycine_max
                                                 200       210       220       230       240       250
NP_001241007.1_FNS2_Glycine_max                  TDSQAEQARTLVREVTQIFGEFNVSDFLGFCKNLDLQGFRKRALDIHKRYDALLEKIISD
TRINITY_DN252_c0_g1_i1                           GDRKADEFKTMVVELMVLAGVFNIGDFVPALEWLDLQGVAAKMKKLHKRFDAFLSEIVED
AAD39549.1_FNS2_Gerbera_hybrid_cultivar          TNSEADEAKNLVREVTKIFGQFNVSDFIWFCKNIDLQGFKKRYEGTHRRYDALLERIIMG
AGA17938.1_FNS2_Dahlia_pinnata                   NKTEAEEAKNLVREVTTIFGEFNVSDFIWFCKKLDLQGFKKRYEDIRTRYDALLERIIFA
AWX67430.1_FNS2_Salvia_miltiorrhiza              TEGEAEAARTVVREVTQIFGEFDVADIICFCKNFDFQGIRKRSQDIRRRYDALLENIITL
ACB56919.1_FNS2_Pilosella_officinarum            TNSEADEAKNLVREVTTIFGQFNVSDFIWFCKNLDLQGFKKRYEDTHRRYDALLEKIISE
AIS92511.1_FNS2_Epimedium_sagittatum             KDDQAKEARVLIREVTQIFGTFNLSDFIWLLRKFDIQGLLKKSKDARKRYNVMIERIIKE
BAB59004.1_FNS2_Perilla_frutescens_var._crispa   TESEAEAARTVIREVTQIFGEFDVSDIIWLCKNFDFQGIRKRSEDIQRRYDALLEKIITD
AMW91728.1_FNS2_Scutellaria_baicalensis          TDGEAEAARTVIREVTEIFGEFDAADIIWFCKNFDLQGIRKRSEDIQRRYDALLEKIITD
AMW91729.1_FNS2_Scutellaria_baicalensis          SEEQAEVARTLIREVSQLFGEFDFGDMLWFCKSFDFQGIKKRSKDIKVRYDALLEKILTD
BAA84072.1_FNS2_Torenia_hybrid_cultivar          DEGDAEAARTVIREVTQIFGEFDVTDIIWFCKKFDLQGIKKRSEDIQRRYDALLEKIISD
ACV65037.1_FNS2_Glycine_max                      TDSQAEQARTLVREVTQIFGEFNVSDFLGFCKNLDLQGFRKRALDIHKRYDALLEKIISD
BAA84071.1_FNS2_Antirrhinum_majus                TEGEAEVARTVIREVTQIFGEFDVSEIVWFCKNLDLQGIRKRSEDIRRRYDALLEKIISD
BAF49323.1_FNS2_Lobelia_erinus                   TDGEADEAKNLVREVTQIFGQFNLSDFVWFCRNVDLQGIKKRYEDTHRRYDALLEKIISE
BAD91809.1_FNS2_Gentiana_triflora                TESGATNVRNVIRDVTELFGEFDVSDIIWFCKNTDLQGIKKRANGIHERYDALLEKIISD
AAF04115.1_FNS2_Callistephus_chinensis           TNNEADEAKNLVREVTKIFGEFNISDFICLFKNIDLQGFKKRYVDTHTRYNALLEKMIFE
ACH99109.1_FNS2_Camellia_sinensis                TTGEAEEATTLAREVTKIFGEFNVSDFMWVIRNFDLQGFRKRVEDIYTRYDALLERIITN
XP_002461286.1_FNS2_Sorghum_bicolor              ..HMTEAARDCAKHVAELVGAFNIEDYVGICRGWDLQGITRRTREVRDKFDALMEIMITA
sp|P93149.2_FNS2_Glycyrrhiza_echinata            ..GEAEEARDVVRDVTEIFGEFNVSDFIWLFKKMDLQGFGKRIEDLFQRFDTLVERIISK
BAG94859.1_FNS2_Oryza_sativa_Japonica_Group      ..SVTEEAQELVKVVAELVGAFNADDYIALCRGWDLQGLGRRAADVHKRFDALLEEMIRH
ABC86159.1_FNS2_Medicago_truncatula              ..GEAEEARDVVRDVTEIFGEFNVSDFIWLFKKLDLQGFGKRIEDLFMRFDTLVERIISK
XP_008663013.1_FNS2_Zea_mays                     ..SVTDEAQEVVKDVAELVGAFNVDDYISLVRGWDLQGLRRRAAGVHRRFDALLEDILRH
                                             acc
```

```
                                                                                 α13
NP_001241007.1_FNS2_Glycine_max                                    TT
                                                 260       270       280       290       300
NP_001241007.1_FNS2_Glycine_max                  REELR...RKSKVDGCED..GDDEK.........VKDFLDILLDVAEQKECE.VQLTRNH
TRINITY_DN252_c0_g1_i1                           HKMNL...KGS.........DGSEKH.........TDMLSTLIALEEDADGEGGKLTDTE
AAD39549.1_FNS2_Gerbera_hybrid_cultivar          REENR...RR.....GKI..KDGEG..........KDFLDMLLDVLEDGKAE.IKITRDH
AGA17938.1_FNS2_Dahlia_pinnata                   REEMR...KE....GKGM..EDGKG..........KDFLDMLLDVLEDDKAE.IKITRNH
AWX67430.1_FNS2_Salvia_miltiorrhiza              RENHR...RS.......A..AAAVP..........RDFLDMFLDIMESGKAE.VTFTRDH
ACB56919.1_FNS2_Pilosella_officinarum            REEKR...RK....GGKR..DDVKG..........TDFLDMLLDVLEDGKAE.IKITRDH
AIS92511.1_FNS2_Epimedium_sagittatum             REEFR.TNKKL.........GSTEGKAD....GGVKDFLDILLDISEDDNAE.MKLNRDH
BAB59004.1_FNS2_Perilla_frutescens_var._crispa   REKQR...RTH...GGGG..GGGEA..........KDFLDMFLDIMESGKAE.VKFTREH
AMW91728.1_FNS2_Scutellaria_baicalensis          REKLR...RSH...R......GGEA..........KDFLDIFLDIMDSGNSE.VKFSREH
AMW91729.1_FNS2_Scutellaria_baicalensis          RENVR...RQN.........GVVEP..........KDMLDMFLDIMEGGKTD.VEFTREH
BAA84072.1_FNS2_Torenia_hybrid_cultivar          RERSR...RQNRDKHGGG..NNEEA..........KDFLDMLLDVMESGDTE.VKFTREH
ACV65037.1_FNS2_Glycine_max                      REELR...RKSKVDGCED..GDDEK.........VKDFLDILLDVAEQKECE.VQLTRNH
BAA84071.1_FNS2_Antirrhinum_majus                RERLR...LRG...GGGG..GGGEV..........KDFLDMLLDVMESEKSE.VEFTREH
BAF49323.1_FNS2_Lobelia_erinus                   REEKR...RGK.MGGGKIGEGEGEG..........KDFLDMLLDVLEDKKAE.VKITRDH
BAD91809.1_FNS2_Gentiana_triflora                RERTRIVEKKNSGAGGGS..GDGER..........NDFLDILMDAMEDDTSE.VKLSRNH
AAF04115.1_FNS2_Callistephus_chinensis           REEKR...KQ.....KKS..EDGKG..........KDFLDILLDVLEDENAE.IKITRDH
ACH99109.1_FNS2_Camellia_sinensis                REEVR...EKN.VQERKL..GVGEGH.......HVKDFLDLLLDVLEEDHSE.INFSRDN
XP_002461286.1_FNS2_Sorghum_bicolor              KEEKR...RSQGDDGAE..............TPTKDLLDILMDAAADENAE.VRLTREN
sp|P93149.2_FNS2_Glycyrrhiza_echinata            REQTR...KDRRRNGKKGEQGSGDG.........IRDFLDILLDCTEDENSE.IKIQRVH
BAG94859.1_FNS2_Oryza_sativa_Japonica_Group      KEEAR...MRKKTDTD...............VGSKDLLDILLDKAEDGAAE.VKLTRDN
ABC86159.1_FNS2_Medicago_truncatula              REELR...KNKGRKENKGEQGA..E.........FRDFLDILLDCAEDQNSE.IKVQRVH
XP_008663013.1_FNS2_Zea_mays                     KEEAR...AARRLDQDDD..GRGSSKQDKKQATHSKDLLDILMDKADDPAAE.IKLTREN
                                             acc
```

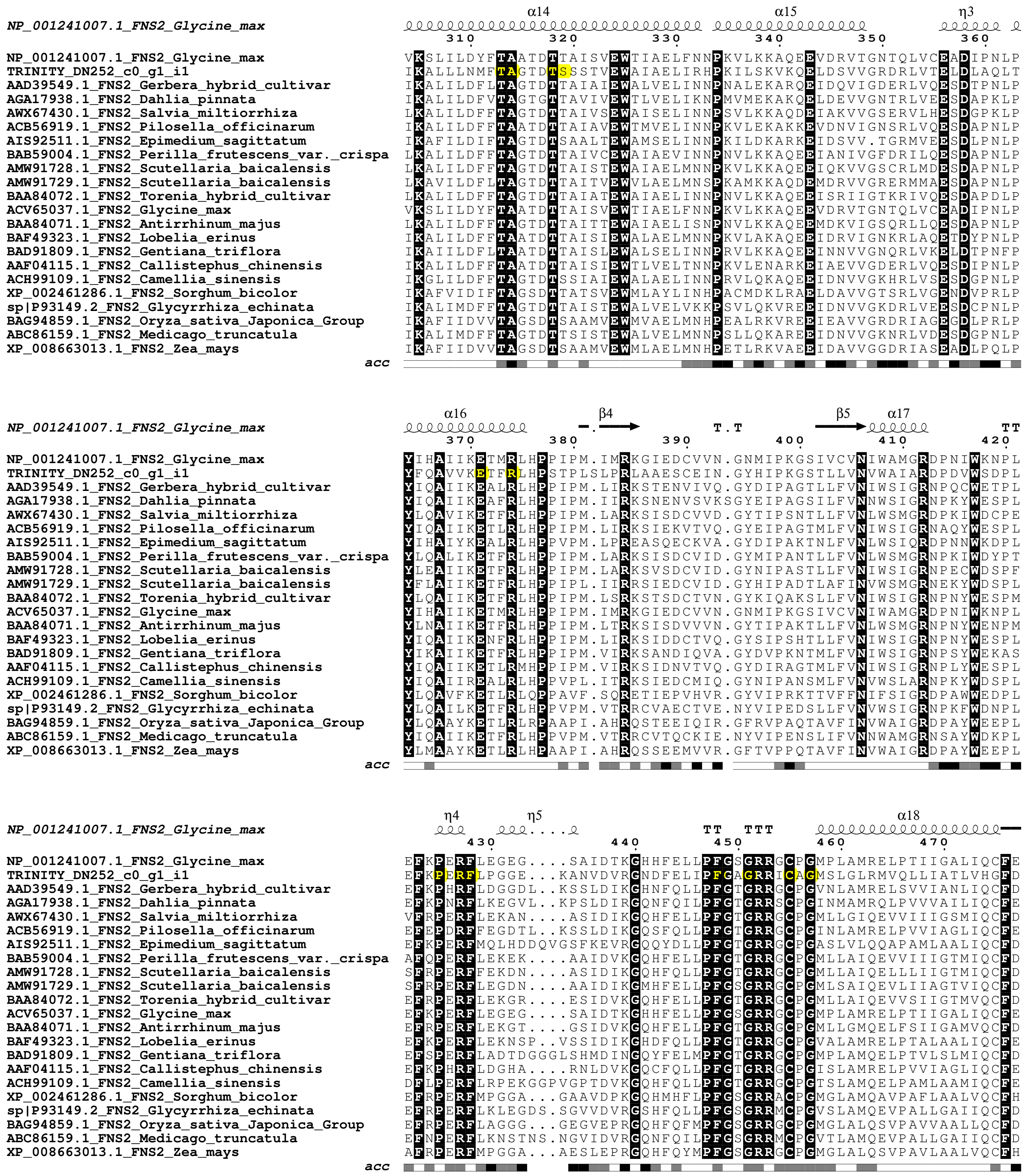
NP_001241007.1_FNS2_Glycine_max
α14 α15 η3
310 320 330 340 350 360
NP_001241007.1_FNS2_Glycine_max VKSLILDYFTAATDTTAISVEWTIAELFNNPKVLKKAQEEVDRVTGNTQLVCEADIPNLP
TRINITY_DN252_c0_g1_i1 IKALLLNMFTAGTDTTSSSTVEWAIAELIRHPKILSKVKQELDSVVGRDRLVTELDLAQLT
AAD39549.1_FNS2_Gerbera_hybrid_cultivar IKALILDFLTAGTDTTAIAIEWALVELINNPNALEKARQEIDQVIGDERLVQESDTPNLP
AGA17938.1_FNS2_Dahlia_pinnata IKALILDFVTAGTGTTAVIVEWTLVELIKNPMVMEKAKQELDEVVGNTRLVEESDAPKLP
AWX67430.1_FNS2_Salvia_miltiorrhiza LKALILDFFTAGTDTTAIVSEWAISELINNPSVLKKAQDEIAKVVGSERVLHESDGPKLP
ACB56919.1_FNS2_Pilosella_officinarum IKALILDFFTAATDTTAIAVEWTMVELINNPKVLEKAKKEVDNVIGNSRLVQESDAPNLP
AIS92511.1_FNS2_Epimedium_sagittatum IKAFILDIFTAGTDTSAALTEWAMSELINNPKILDKARKEIDSVV.TGRMVEESDLPNLP
BAB59004.1_FNS2_Perilla_frutescens_var._crispa LKALILDFFTAGTDTTAIVCEWAIAEVINNPNVLKKAQEEIANIVGFDRILQESDAPNLP
AMW91728.1_FNS2_Scutellaria_baicalensis LKALILDFFTAGTDTTAISTEWAIAELMNNPKVLKKAQEEIQKVVGSCRLMDESDAPNLP
AMW91729.1_FNS2_Scutellaria_baicalensis LKAVILDFLTAGTDTTAITVEWVLAELMNSPKAMKKAQDEMDRVVGRERMMAESDAPNLP
BAA84072.1_FNS2_Torenia_hybrid_cultivar LKALILDFFTAGTDTTAIATEWAIAELINNPNVLKKAQEEISRIIGTKRIVQESDAPDLP
ACV65037.1_FNS2_Glycine_max VKSLILDYFTAATDTTAISVEWTIAELFNNPKVLKKAQEEVDRVTGNTQLVCEADIPNLP
BAA84071.1_FNS2_Antirrhinum_majus LKALILDFFTAGTDTTAITTEWAIAELISNPNVLKKAQEEMDKVIGSQRLLQESDAPNLP
BAF49323.1_FNS2_Lobelia_erinus IKALILDFFTAATDTTAISIEWALAELMNNPKVLAKAQEEIDRVIGNKRLAQETDYPNLP
BAD91809.1_FNS2_Gentiana_triflora IKAIILDFLTAATDTTAISLEWALSELINNPRVLKKAQEEINNVVGNQRLVKELDTPNFP
AAF04115.1_FNS2_Callistephus_chinensis IKALILDFFTAATDTTAISIEWTLVELTNNPKVLENARKEIAEVVGDERLVQESDIPNLP
ACH99109.1_FNS2_Camellia_sinensis IKGLILDFFTAGTDTSSIAIEWALAELINNPRVLQKAQEEIDNVVGKHRLVSESDGPNLP
XP_002461286.1_FNS2_Sorghum_bicolor IKAFVIDIFTAGSDTTATSVEWMLAYLINHPACMDKLRAELDAVVGTSRLVGENDVPRLP
sp|P93149.2_FNS2_Glycyrrhiza_echinata IKALIMDFFTAGTDTTAISTEWALVELVKKPSVLQKVREEIDNVVGKDRLVEESDCPNLP
BAG94859.1_FNS2_Oryza_sativa_Japonica_Group IKAFIIDVVTAGSDTSAAMVEWMVAELMNHPEALRKVREEIEAVVGRDRIAGEGDLPRLP
ABC86159.1_FNS2_Medicago_truncatula IKALIMDFFTAGTDTTSISTEWALVELMNNPSLLQKAREEIDNVVGKNRLVDESDGPNLP
XP_008663013.1_FNS2_Zea_mays IKAFIIDVVTAGSDTSAAMVEWMLAELMNHPETLRKVAEEIDAVVGGDRIASEADLPQLP
acc
NP_001241007.1_FNS2_Glycine_max
α16 β4 T.T β5 α17 TT
370 380 390 400 410 420
NP_001241007.1_FNS2_Glycine_max YIHAIIKETMRLHPPIPM.IMRKGIEDCVVN.GNMIPKGSIVCVNIWAMGRDPNIWKNPL
TRINITY_DN252_c0_g1_i1 YFQAVVKETFRLHPSTPLSLPRLAAESCEIN.GYHIPKGSTLLVNVWAIARDPDVWSDPL
AAD39549.1_FNS2_Gerbera_hybrid_cultivar YIQAIIKEALRLHPPIPM.LIRKSTENVIVQ.GYDIPAGTLLFVNIWSIGRNPQCWETPL
AGA17938.1_FNS2_Dahlia_pinnata YIQAIIKEAFRLHPPIPM.IIRKSNENVSVKSGYEIPAGSILFVNNWSIGRNPKYWESPL
AWX67430.1_FNS2_Salvia_miltiorrhiza YLQAVIKETFRLHPPIPM.LARKSISDCVVD.GYTIPSNTLLFVNLWSMGRDPKIWDCPE
ACB56919.1_FNS2_Pilosella_officinarum YIQAIIKETLRLHPPIPM.LIRKSIEKVTVQ.GYEIPAGTMLFVNIWSIGRNAQYWESPL
AIS92511.1_FNS2_Epimedium_sagittatum YIHAIYKEALRLHPPVPM.LPREASQECKVA.GYDIPAKTNLFVNLWSIQRDPNNWKDPL
BAB59004.1_FNS2_Perilla_frutescens_var._crispa YLQALIKETFRLHPPIPM.LARKSISDCVID.GYMIPANTLLFVNLWSMGRNPKIWDYPT
AMW91728.1_FNS2_Scutellaria_baicalensis YLEAIIKETFRLHPPIPM.LARKSVSDCVID.GYNIPASTLLFVNIWSIGRNPECWDSPF
AMW91729.1_FNS2_Scutellaria_baicalensis YFLAIIKETFRLHPPIPL.IIRRSIEDCVID.GYHIPADTLAFINVWSMGRNEKYWDSPL
BAA84072.1_FNS2_Torenia_hybrid_cultivar YLQAIIKETFRLHPPIPM.LSRKSTSDCTVN.GYKIQAKSLLFVNIWSIGRNPNYWESPM
ACV65037.1_FNS2_Glycine_max YIHAIIKETMRLHPPIPM.IMRKGIEDCVVN.GNMIPKGSIVCVNIWAMGRDPNIWKNPL
BAA84071.1_FNS2_Antirrhinum_majus YLNAIIKETFRLHPPIPM.LTRKSISDVVVN.GYTIPAKTLLFVNLWSMGRNPNYWENPM
BAF49323.1_FNS2_Lobelia_erinus YIQAIIKENFRLHPPIPM.LIRKSIDDCTVQ.GYSIPSHTLLFVNIWSIGRNPNYWESPL
BAD91809.1_FNS2_Gentiana_triflora YIKAIIKETFRLHPPIPM.VIRKSANDIQVA.GYDVPKNTMLFVNIWSIGRNPSYWEKAS
AAF04115.1_FNS2_Callistephus_chinensis YIQAIIKETLRMHPPIPM.VIRKSIDNVTVQ.GYDIRAGTMLFVNIWSIGRNPLYWESPL
ACH99109.1_FNS2_Camellia_sinensis YIQAIIREALRLHPPVPL.ITRKSIEDCMIQ.GYNIPANSMLFVNVWSLARNPKYWDSPL
XP_002461286.1_FNS2_Sorghum_bicolor YLQAVFKETLRLQPPAVF.SQRETIEPVHVR.GYVIPRKTTVFFNIFSIGRDPAWWEDPL
sp|P93149.2_FNS2_Glycyrrhiza_echinata YLQAILKETLRLHPPVPM.VTRRCVAECTVE.NYVIPEDSLLFVNVWSIGRNPKFWDNPL
BAG94859.1_FNS2_Oryza_sativa_Japonica_Group YLQAAYKETLRLRPAAPI.AHRQSTEEIQIR.GFRVPAQTAVFINVWAIGRDPAYWEEPL
ABC86159.1_FNS2_Medicago_truncatula YIQAIIKETFRLHPPVPM.VTRRCVTQCKIE.NYVIPENSLIFVNNWAMGRNSAYWDKPL
XP_008663013.1_FNS2_Zea_mays YLMAAYKETLRLHPAAPI.AHRQSSEEMVVR.GFTVPPQTAVFINVWAIGRDPAYWEEPL
acc
NP_001241007.1_FNS2_Glycine_max
η4 η5 TT TTT α18
430 440 450 460 470
NP_001241007.1_FNS2_Glycine_max EFKPERFLEGEG....SAIDTKGHHFELLPFGSGRRGCPGMPLAMRELPTIIGALIQCFE
TRINITY_DN252_c0_g1_i1 EFKPERFLPGGE...KANVDVRGNDFELIPFGAGRRICAGMSLGLRMVQLLIATLVHGFD
AAD39549.1_FNS2_Gerbera_hybrid_cultivar EFKPHRFLDGGDL..KSSLDIKGHNFQLLPFGTGRRGCPGVNLAMRELSVVIANLIQCFD
AGA17938.1_FNS2_Dahlia_pinnata EFKPNRFLKEGVL..KPSLDIRGQNFQILPFGTGRRSCPGINMAMRQLPVVVAILIQCFE
AWX67430.1_FNS2_Salvia_miltiorrhiza VFRPERFLEKAN....ASIDIKGQHFELLPFGTGRRGCPGMLLGIQEVVIIIGSMIQCFD
ACB56919.1_FNS2_Pilosella_officinarum EFEPDRFFEGDTL..KSSLDIKGQSFQLLPFGTGRRGCPGINLAMRELPVVIAGLIQCFE
AIS92511.1_FNS2_Epimedium_sagittatum EFKPERFMQLHDDQVGSFKEVRGQQYDLLPFGTGRRGCPGASLVLQQAPAMLAALIQCFD
BAB59004.1_FNS2_Perilla_frutescens_var._crispa AFQPERFLEKEK....AAIDVKGQHFELLPFGTGRRGCPGMLLAIQEVVIIIGTMIQCFD
AMW91728.1_FNS2_Scutellaria_baicalensis SFRPERFFEKDN....ASIDIKGQHFQLLPFGTGRRGCPGMLLAIQELLLIIGTMIQCFD
AMW91729.1_FNS2_Scutellaria_baicalensis SFRPERFLEGDN....AAIDIKGMHFELLPFGSGRRGCPGMLSAIQEVLIIAGTVIQCFD
BAA84072.1_FNS2_Torenia_hybrid_cultivar EFRPERFLEKGR....ESIDVKGQHFELLPFGTGRRGCPGMLLAIQEVVSIIGTMVQCFD
ACV65037.1_FNS2_Glycine_max EFKPERFLEGEG....SAIDTKGHHFELLPFGSGRRGCPGMPLAMRELPTIIGALIQCFE
BAA84071.1_FNS2_Antirrhinum_majus EFRPERFLEKGT....GSIDVKGQHFELLPFGTGRRGCPGMLLGMQELFSIIGAMVQCFD
BAF49323.1_FNS2_Lobelia_erinus EFKPERFLEKNSP..VSSIDIKGHDFQLLPFGTGRRGCPGVALAMRELPTALAALIQCFE
BAD91809.1_FNS2_Gentiana_triflora EFSPERFLADTDGGGLSHMDINGQYFELMPFGTGRRGCPGMPLAMQELPTVLSALIQCFD
AAF04115.1_FNS2_Callistephus_chinensis EFKPHRFLDGHA....RNLDVKGQCFQLLPFGTGRRGCPGISLAMRELPVVIAGLIQCFD
ACH99109.1_FNS2_Camellia_sinensis DFLPERFLRPEKGGPVGPTDVKGQHFQLLPFGTGRRGCPGTSLAMQELPAMLAAMIQCFE
XP_002461286.1_FNS2_Sorghum_bicolor EFRPERFMPGGA...AAVDPKGQHMHFLPFGSGRRACPGMGLAMQAVPAFLAALVQCFH
sp|P93149.2_FNS2_Glycyrrhiza_echinata EFRPERFLKLEGDS.SGVVDVRGSHFQLLPFGSGRRMCPGVSLAMQEVPALLGAIIQCFD
BAG94859.1_FNS2_Oryza_sativa_Japonica_Group EFRPERFLAGGG...GEGVEPRGQHFQFMPFGSGRRGCPGMGLALQSVPAVVAALLQCFD
ABC86159.1_FNS2_Medicago_truncatula EFNPERFLKNSTNS.NGVIDVRGQNFQILPFGSGRRMCPGVTLAMQEVPALLGAIIQCFD
XP_008663013.1_FNS2_Zea_mays AFRPERFMPGGA...AESLEPRGQHFQYMPFGSGRRGCPGMGLALQSVPAVLAALVQCFH
acc

```
NP_001241007.1_FNS2_Glycine_max              β6 ⟶                                   β7 ⟶
                                                     TT.T
                                             480       490       500       510       520
NP_001241007.1_FNS2_Glycine_max              WKMLGSQGEI.LDHGRSLISMDERPGLTAPRANDLIGIPVARLNPTPFRQM........
TRINITY_DN252_c0_g1_i1                       WELADGL....M...PDKLNMDEAFGLTLQRAVPLMVHPRPRL.PLHAYGASN......
AAD39549.1_FNS2_Gerbera_hybrid_cultivar      WDVVGER..........LLNTDERAGLTAPRAVDFVCVPLERGNTLKILGS...N....
AGA17938.1_FNS2_Dahlia_pinnata               WTVNDKQ..........VLNMDEKGGLTTPRATDLVCFPLLRKNSPHSMFT...SV...
AWX67430.1_FNS2_Salvia_miltiorrhiza          WKLPQGT.........RAIDMAERSGLTAPRAYDLICRVVPRIDPTLVFGALPDSV...
ACB56919.1_FNS2_Pilosella_officinarum        WDVNNKE..........ALITDERAGLTAPRAVDFVCVPSMRENCPKVF..........
AIS92511.1_FNS2_Epimedium_sagittatum         WEVAGHDDGIKL....ATVDMIERPGLTVPRANALLLVPTTRFNPFTDTVVHP......
BAB59004.1_FNS2_Perilla_frutescens_var._crispa WKLPDGS.........GHVDMAERPGLTAPRETDLFCRVVPRVDPLVVSTQ........
AMW91728.1_FNS2_Scutellaria_baicalensis      WELPEGS.........GPVDMTERAGLTAPRAEDLICRVSCRVDPKIVF..........
AMW91729.1_FNS2_Scutellaria_baicalensis      WEQADGS.........GRVDMSERPGLTTPREIDLVCRVVPRVDERVISGH........
BAA84072.1_FNS2_Torenia_hybrid_cultivar      WKLADGSG........NNVDMTERSGLTAPRAFDLVCRLYPRVDPATISGA........
ACV65037.1_FNS2_Glycine_max                  WKMLGSQGEI.LDHGRSLISMDERPGLTAPRANDLIGIPVARLNPTPFRQM........
BAA84071.1_FNS2_Antirrhinum_majus            WKLPDGV.........KSVDMTERPGLTAPRANDLVCQLVPRIDPVVVSGP........
BAF49323.1_FNS2_Lobelia_erinus               WKAEGGE..........ALDMSERAGLTAPRAHDLVCVPVARINSPDIFDC........
BAD91809.1_FNS2_Gentiana_triflora            YIPLDFKGE....KAERVMDMSERPGLTAPRANELMCLLKPRIDLPNLLGNVKGE....
AAF04115.1_FNS2_Callistephus_chinensis       WNANDKE..........VLSMDERAGLTAPRAVDLEFVPLMRQNCPNIFVS...A....
ACH99109.1_FNS2_Camellia_sinensis            WKVVNQSGDV.M.NGDGALDMTEQPGMTAPRAHDLVCMPIPRIDQLYALLDP.......
XP_002461286.1_FNS2_Sorghum_bicolor          WAVPIPQGQ....STAPPLDMEEEAGLVTARKHHLVLIPTPRLNPLPVPVPVPATGKAT
sp|P93149.2_FNS2_Glycyrrhiza_echinata        FHVVGPKGEI.LKGDDIVINVDERPGLTAPRAHNLVCVPVDRTSGGGPLKIIEC.....
BAG94859.1_FNS2_Oryza_sativa_Japonica_Group  WQCMD..........NKLIDMEEADGLVCARKHRLLLHAHPRLHPFPPLL.........
ABC86159.1_FNS2_Medicago_truncatula          FNFVGPKGEI.LKGGDIVIDVNERPGLTAPRVHDLVCVPVERFACGGPLQSLGC.....
XP_008663013.1_FNS2_Zea_mays                 WATVDGDG.....GVNK.IDMSESDGLVCARKKPLLLRPTPRLTPFPAVV.........
                                       acc
```

*NP_001268144.1_DFR_Vitis_vinifera*

β1 (positions ~9–12), α1 (positions ~15–26), β2 (position ~30 onward)

| Sequence | 1 . . . 10 . . . 20 . . . 30 |
|---|---|
| **NP_001268144.1_DFR_Vitis_vinifera** | .........MG..................SQSETVCVTGASGFIGSWLVMRLLERGYTVR |
| **TRINITY_DN27402_c0_g1_i3** | .........MET..................QTKTLCVTGASGFIGSWLVMRLLECGYMVR |
| **NP_199094.1_DFR_Arabidopsis_thaliana** | .........MV..................SQKETVCVTGASGFIGSWLVMRLLERGYFVR |
| **BAA85261.1_DFR_Arabidopsis_thaliana** | .........MV..................SQKETVCVTGASGFIGSWLVMRLLERGYFVR |
| **AEI59122.1_DFR_Medicago_sativa** | .........MG..................SVSETVCVTGASGFIGSWLVMRLIEHGYTVR |
| **BAD67185.1_DFR_Spinacia_oleracea** | .........MVV..................QGEIVCVTGAAGFIGSWLVKRLLEHGYIVR |
| **BAB40789.1_DFR_Lilium_hybrid** | .........MEN..................VKGPVVVTGASGYVGSWLVMKLLQYGYTVR |
| **CAA79154.1_DFR_Solanum_lycopersicum** | .........MASEAHAVV......DAHSPPKTTTVWVTGGAGFIGSWLVMRLLERGYNVH |
| **AHZ30596.1_DFR_Prunus_domestica** | .................................SVCVXGASGFXGSWLVMRLLERGYTVR |
| **AAD26204.1_DFR_Malus_domestica** | .........MG..................SESESVCVTGASGFIGSWLVMRLLEHGYTVR |
| **AAO39819.1_DFR_Pyrus_communis** | .........MG..................SESESVCVTGASGFIGSWLVMRLLEHGYTVR |
| **AAD56578.1_DFR_Daucus_carota** | .........MV..................KELHTVCVTGASGFIGSWLVLRLLQRGYTVR |
| **AKN56970.1_DFR_Gerbera_hybrid_cultivar** | .........MEE.................DSPATVCVTGAAGFIGSWLVMRLLERGYVVH |
| **AOF39984.11_DFR_Brassica_rapa** | .........MV..................AHKETVCVTGASGFIGSWLVMRLLERGYFVR |
| **P51106.1_DFR_Hordeum_vulgare** | .........MDG..................NKGPVVVTGASGFVGSWLVMKLLQAGYTVR |
| **BAE19953.1_DFR_Lotus_japonicus** | .........MS..................SESETVCVTGAAGFIGSWLVMRLIERGYTVR |
| **AII26023.1_DFR_Pisum_sativum** | .........MG..................SVSETVCVTGASGFIGSWLVMRLIEHGYTVR |
| **NP_001274988.1_DFR_Solanum_tuberosum** | .........MASEVHSVV......DAHSPPKTPTVCVTGAAGFIGSWLVMRLLERGYNVH |
| **NP_001152467.2_DFR_Zea_mays** | .MEG...GAGAS..................EKGKVLVTGASGFVGSWLVMKLLQAGYTVR |
| **AIZ74402.1_DFR_Anthurium_andraeanum** | .........MM...................HKGTVCVTGAAGFVGSWLIMRLLEQGYSVK |
| **AAB62873.1_DFR_Bromheadia_finlaysoniana** | .........MEN.................EKKGPVVVTGASGYVGSWLVMKLLQKGYDVR |
| **CAA91922.1_DFR_Callistephus_chinensis** | .........MKE.................DSPPTVCVTGAAGFIGSWLVMRLLERGYIVR |
| **AIM58715.1_DFR_Cymbidium_hybrid_cultivar** | .........MET.................ERKGPVVVTGASGYVGSWLVMKLLQKGYEVR |
| **ACZ48698.1_DFR_Fagopyrum_esculentum** | .........MV..................AEGEIVCVTGASGFVGSWLVMRLLEHGYVVR |
| **ACZ48697.1_DFR_Fagopyrum_tataricum** | .........MV..................AEGEIVCVTGASGFVGSWLVMRLLEHGYVVR |
| **BAO53730.1_DFR2_Glycine_max** | .........MGS................SSASESVCVTGASGFIGSWLVMRLIERGYTVR |
| **AAP13055.1_DFR_Gypsophila_elegans** | .........MVSSNNTTETLDGKHDTKPV.QGETVCVTGASGFIGSWLIMRLLERGYTVR |
| **XP_013466134.1_DFR_Medicago_truncatula** | .........MG..................SMAETVCVTGASGFIGSWLVMRLMERGYMVR |
| **BAF96936.1_DFR_Nicotiana_tabacum** | .........MASEAHAAV......HA.PPPVAPTVCVTGAAGFIGSWLVMRLLERGYNVH |
| **BAA36183.1_DFR_Oryza_sativa_Japonica_Group** | ........MGEA..................VKGPVVVTGASGFVGSWLVMKLLQAGYTVR |
| **BAA12723.1_DFR_Rosa_hybrid_cultivar** | .........MA..................SESESVCVTGASGFIGSWLVMRLLDRGYTVR |
| **AHL46438.1_DFR_Fragaria_vesca** | .M......GLG..................AESGSVCVTGASGFVGSWLVMRLLERGYTVR |
| **AHL46443.1_DFR_Fragaria_x_ananassa** | .M......GLG..................AESGSVCVTGASGFVGSWLVMRLLEHGYTVR |
| **AHL46445.1_DFR_Fragaria_vesca** | .........MG..................SESESVCVHGASGFVGSWLVMRLLERGYTVR |
| **AHM27144.1_DFR_Angelonia_angustifolia** | .........METTATQPPP....PSS.AAAVPATVCVTGAAGFIGSWLVMKLLQRGYNVR |
| **AAT84073.1_DFR_Camellia_sinensis** | .........MKDSVA.........SA.TASAPGTVCVTGAAGFIGSWLVMRLLERGYIVR |
| **AAS00611.1_DFR_Citrus_sinensis** | .........MG..................SIAETVCVTGASGFIGSWLIMRLLERGYAVR |
| **AAX16491.1_DFR_Crataegus_monogyna** | .........MG..................SESESVCVTGASGFIGSWLVMRLLEHGYTVR |
| **BAF49325.1_DFR_Delphinium_belladonna** | .........MTV....................ETVCVTGAAGFIGSWLVMRLLERGYLVR |
| **CAA78930.1_DFR_Gerbera_hybrid_cultivar** | .........MEE.................DSPATVCVTGAAGFIGSWLVMRLLERGYVVH |
| **AAO39816.1_DFR_Malus_domestica** | .........MG..................SESESVCVTGASGFIGSWLVMRLLEHGYTVR |
| **AAO39817.1_DFR_Malus_domestica** | .........MG..................SESESVCVTGASGFIGSWLVMRLLEHGYTVR |
| **AAR27015.1_DFR2_Medicago_truncatula** | .........MG..................SVSETVCVTGASGFIGSWLVMRLMERGYTVR |
| **AEF14420.1_DFR_Onobrychis_viciifolia** | .........MG..................STSETVCVTGAAGFVGSWLVMRLIERGYTVR |
| **AAF60298.1_DFR_Petunia_x_hybrida** | ...........MPLHLRC......SA.......TVCVTGAAGFIGSWLVMRLLERGYNVH |
| **AAQ77347.1_DFR_Triticum_aestivum** | .........MDG..................NKGPVVVTGASGFVGSWLVMKLLQVGYTVR |
| **AAL89715.1_DFR_Vaccinium_macrocarpon** | .........MKD.VN.........SG.LGT...TVCVTGAAGFIGSWLIMRLLERGYVVR |
| **AAX12420.1_DFR_Vaccinium_macrocarpon** | .........MKD.VN.........SG.LGT...TVCVTGAAGFIGSWLIMRLLERGYVVR |
| **CAA33543.1_DFR_Antirrhinum_majus** | .........MSPTSLNTSS.....ET.APPSSTTVCVTGAAGFIGSWLVMRLLERGYTVR |
| **AAB62873.1_DFR_Bromheadia_finlaysoniana** | .........MEN.................EKKGPVVVTGASGYVGSWLVMKLLQKGYDVR |
| **BAA84940.1_DFR_Camellia_sinensis** | .........MKDSVA.........SA.TASAPGTVCVTGAAGFIGSWLVMRLLERGYIVR |
| **AAC17843.1_DFR_Cymbidium_hybrid_cultivar** | .........MET.................ERKGPVVVTGASGYVGSWLVMKLLQKGYEVR |
| **CAA91924.1_DFR_Dianthus_caryophyllus** | .........MVSS.TINETLDGKHDINKVGQGETVCVTGASGFIGSWLIMRLLERGYTVR |
| **CAA70345.1_DFR_Forsythia_x_intermedia** | .........MET............DA.LPLPATTVCVTGASGFIGSWLVMRLLERGYFVR |
| **AAC25960.1_DFR_Fragaria_x_ananassa** | .M......GLG..................AESGSVCVTGASGFVGSWLVMRLLEHGYTVR |
| **BAA12736.1_DFR_Gentiana_triflora** | .........ME.............GG.ILSNATTVCVTGASGYIGSWLAMRLLERGYTVR |
| **AAD54273.1_DFR1_Glycine_max** | .........MG..................SASESVCVTGASGFIGSWLVMRLIERGYTVR |
| **BAA59332.1_DFR_Ipomoea_nil** | .........MVGGNHTPA...........SPAPTVCVTGAAGFVGSWLVMKLLQRGYIVH |
| **BAA36405.1_DFR_Ipomoea_purpurea** | .........MVGGNHTPA...........SPAPTVCVTGAAGFVGSWLVMKLLQRGYIVH |
| **AAD49343.1_DFR_Lilium_hybrid_division_VII** | .........MEN..................AKGPVVVTGASGYVGSWLVMKLLQYGYTIR |
| **AAF23884.2_DFR3_Lotus_corniculatus** | .........MG..................SVPETVCVTGAAGFIGSWLVMRLMERGYMVR |
| **BAE19948.1_DFR_Lotus_japonicus** | .........MG..................SAAKTVCVTGSTGFIGSWLVMRLMERGYMVR |
| **BAE19949.1_DFR_Lotus_japonicus** | .........MG..................SVPETVCVTGAAGFIGSWLVMRLMERGYMVR |
| **BAE19950.1_DFR_Lotus_japonicus** | .........MG..................SAAKTVCVTGSTGFIGSWLVMRLMEGGYTVR |
| **BAE19951.1_DFR_Lotus_japonicus** | .........MG..................SAAKTVCVTGSTGFIGSWLVMRLMERGYMVR |
| **BAE19953.1_DFR_Lotus_japonicus** | .........MS..................SESETVCVTGAAGFIGSWLVMRLIERGYTVR |
| **BAA19658.1_DFR_Perilla_frutescens** | .........MSLETVAAP...........PPATTVCVTGASGFIGSWIIMRLLERGYVVR |
| **BAB20075.1_DFR_Torenia_hybrid_cultivar** | .........MSMEVVVPK...........AQPITVCVTGASGFIGSWLVMKLLNRGYTVH |
| **AAL35830.1_DFR_Triticum_monococcum** | .........MDG..................SKGPVVVTGASGFVGSWLVMKLLQAGYTVR |
| **QFQ61498.1_DFR_Dryopteris_erythrosora** | .........MDK.................PLHSTVLVTGCTGHIGSWLVLRLLEKGYSVR |
| **QFQ61499.1_DFR_Dryopteris_erythrosora** | .MAPNAVAVMAL.................P..ELVCVTGASGHLGTWLVKRLLERGYRVR |
| **BAA74700.1_DFR_Ipomoea_purpurea** | .........MVDGNHPL.............PAPKVCVTGAAGFIGSWLVKTLLQRGYHVH |
| **BAF93896.1_DFR_Iris_x_hollandica** | ..............................MMSPVVVTGASGYVGSWLVMKLLRDGYAVR |
| **XP_008797532.1_DFR_Phoenix_dactylifera** | .........ME...................TKGPVVVTGASGFIGTWLVMRLLEEGYIVR |
| **AEQ92209.1_DFR_Ipomoea_batatas** | .........MVDGNHP................KVVCVTGAAGFIGSWLVMTLLQRGYHVH |
| **BAH98155.1_DFR_Tulipa_gesneriana** | .........MKV..................VKGPVVATGASGYVGSWLVMKLLENGYTVR |
| **APG32494.1_DFR2_Freesia_hybrid_cultivar** | ...............................MGTVVVTGASGYVGSWLLMKLLQNGYAVR |
| **AAO63026.1_DFR_Allium_cepa** | MMK.....EIGA..................AGGAVVVTGAGGYVGSWLVMKLLHYGYTVR |
| **AFP58815.1_DFR_Hyacinthus_orientalis** | .........MEM..................EKGPVVATGAGGYIGSWLVMKLLRAGYTVR |
| *acc* | |

```
                                                                                  α2
NP_001268144.1_DFR_Vitis_vinifera        →                                       40        50
NP_001268144.1_DFR_Vitis_vinifera        ATV.RDP....................................TNVKKVKHLLDL.PKA.
TRINITY_DN27402_c0_g1_i3                 ATV.RDP....................................DNAKKVQHLLDL.PNA.
NP_199094.1_DFR_Arabidopsis_thaliana     ATV.RDP....................................GNLKKVQHLLDL.PNA.
BAA85261.1_DFR_Arabidopsis_thaliana      ATV.RDP....................................GNLKKVQHLLDL.PNA.
AEI59122.1_DFR_Medicago_sativa           ATV.RDP....................................DNIKKVKHLLEL.PGA.
BAD67185.1_DFR_Spinacia_oleracea         ATV.RDP....................................GNVKKVQHLLDL.PNA.
BAB40789.1_DFR_Lilium_hybrid             ATV.RDP....................................RDLRKTKPLLDL.PGA.
CAA79154.1_DFR_Solanum_lycopersicum      ATV.RDP....................................ENQKKVKHLLEL.PKA.
AHZ30596.1_DFR_Prunus_domestica          ATV.RDP....................................TNQKKVKHLLEL.PKA.
AAD26204.1_DFR_Malus_domestica           ATV.RDP....................................TNQKKVKHLLDL.PKA.
AAO39819.1_DFR_Pyrus_communis            ATV.RDP....................................TNQKKVKHLLDL.PKA.
AAD56578.1_DFR_Daucus_carota             ATV.RDP....................................GNPQKVKHLLQL.PKA.
AKN56970.1_DFR_Gerbera_hybrid_cultivar   ATV.RDP....................................GDLKKVKHLLEL.PKA.
AOF39984.11_DFR_Brassica_rapa            ATV.RDP....................................GNLKKVQHLLDL.PNA.
P51106.1_DFR_Hordeum_vulgare             ATV.RDP....................................ANVEKTKPLLEL.PGA.
BAE19953.1_DFR_Lotus_japonicus           ATI.RDP....................................ANMKKVKHLLEL.PDA.
AII26023.1_DFR_Pisum_sativum             ATV.RDP....................................DNVKKVKHLLEL.PDA.
NP_001274988.1_DFR_Solanum_tuberosum     ATV.RDP....................................ENQKKVKHLLEL.PKA.
NP_001152467.2_DFR_Zea_mays              ATV.RDP....................................ANVGKTKPLMDL.PGA.
AIZ74402.1_DFR_Anthurium_andraeanum      ATV.RDP....................................SNMKKVKHLLDL.PGA.
AAB62873.1_DFR_Bromheadia_finlaysoniana  ATI.RDP....................................TNLEKVKPLLDL.PRS.
CAA91922.1_DFR_Callistephus_chinensis    ATV.RNP....................................GDMKKVKHLLEL.PKA.
AIM58715.1_DFR_Cymbidium_hybrid_cultivar AAV.RDS....................................TNFEKVKPLLDL.PGS.
ACZ48698.1_DFR_Fagopyrum_esculentum      ATV.RDP....................................TNMKKVKHLLDL.PKS.
ACZ48697.1_DFR_Fagopyrum_tataricum       ATV.RDP....................................SNMKKVKHLLDL.PKS.
BAO53730.1_DFR2_Glycine_max              ATV.HDP....................................ANMKKVKHLVEL.PGA.
AAP13055.1_DFR_Gypsophila_elegans        GTV.RDP....................................DNTKKVQHLLDL.PQA.
XP_013466134.1_DFR_Medicago_truncatula   ATV.RDP....................................ENLKKVSHLLEL.PGA.
BAF96936.1_DFR_Nicotiana_tabacum         ATV.RDP....................................ENKKKVKHLFEL.PKA.
BAA36183.1_DFR_Oryza_sativa_Japonica_Group ATV.RDP..................................SNVGKTKPLLEL.AGS.
BAA12723.1_DFR_Rosa_hybrid_cultivar      ATV.RDP....................................ANKKKVNHLLDL.PKA.
AHL46438.1_DFR_Fragaria_vesca            ATV.RDP....................................ANLKKVRHLLEL.PQA.
AHL46443.1_DFR_Fragaria_x_ananassa       ATV.RDP....................................ANLKKVRHLLEL.PQA.
AHL46445.1_DFR_Fragaria_vesca            ATV.RDP....................................TNAKKVKHLLDL.PKA.
AHM27144.1_DFR_Angelonia_angustifolia    ATV.RDP....................................GNLKKIKHLTQL.PKA.
AAT84073.1_DFR_Camellia_sinensis         ATV.RDP....................................ANLKKVKHLLDL.PKA.
AAS00611.1_DFR_Citrus_sinensis           ATV.RDP....................................DNKKKVKHLLEL.PKA.
AAX16491.1_DFR_Crataegus_monogyna        ATV.RDP....................................TNQKKVKHLLDL.PKA.
BAF49325.1_DFR_Delphinium_belladonna     ATV.RNP....................................DNLKKLRHLLEL.PNA.
CAA78930.1_DFR_Gerbera_hybrid_cultivar   ATV.RDP....................................GDLKKVKHLLEL.PKA.
AAO39816.1_DFR_Malus_domestica           ATV.RDP....................................TNQKKVKHLLDL.PKA.
AAO39817.1_DFR_Malus_domestica           ATV.RDP....................................TNQKKVKHLLDL.PKA.
AAR27015.1_DFR2_Medicago_truncatula      ATV.RDP....................................DNMKKVKHLLEL.PGA.
AEF14420.1_DFR_Onobrychis_viciifolia     ATV.RDP....................................ANMKKVKHLLDL.PDA.
AAF60298.1_DFR_Petunia_x_hybrida         ATV.RDP....................................ENKKKVKHLLEL.PKA.
AAQ77347.1_DFR_Triticum_aestivum         ATV.RDPGERSLRLRLPHFLSPTHTRLTRYLTAFVICFCVFVSANVEKNKPLLEL.PGA.
AAL89715.1_DFR_Vaccinium_macrocarpon     ATV.RDP....................................GNLKKVKHLLEL.PKA.
AAX12420.1_DFR_Vaccinium_macrocarpon     ATV.RDP....................................GNLKKVKHLLEL.PKA.
CAA33543.1_DFR_Antirrhinum_majus         ATV.RDP....................................GNMKKVKHLIEL.PKA.
AAB62873.1_DFR_Bromheadia_finlaysoniana  ATI.RDP....................................TNLEKVKPLLDL.PRS.
BAA84940.1_DFR_Camellia_sinensis         ATV.RDP....................................ANLKKVKHLLDL.PKA.
AAC17843.1_DFR_Cymbidium_hybrid_cultivar AAV.RDS....................................TNFEKVKPLLDL.PGS.
CAA91924.1_DFR_Dianthus_caryophyllus     ATV.RDP....................................DNTKKVQHLLDL.PNA.
CAA70345.1_DFR_Forsythia_x_intermedia    ATA.RDP....................................ENKQKVKHLLEL.PRA.
AAC25960.1_DFR_Fragaria_x_ananassa       ATV.RDP....................................ANLKKVRHLLEL.PQA.
BAA12736.1_DFR_Gentiana_triflora         ATV.RDP....................................GNLKKVQHLLEL.PKA.
AAD54273.1_DFR1_Glycine_max              ATV.RDP....................................VNMKKVKHLVEL.PGA.
BAA59332.1_DFR_Ipomoea_nil               ATV.RDP....................................GNAQKVKHLLEL.PKG.
BAA36405.1_DFR_Ipomoea_purpurea          ATV.RDP....................................GNTQKVKHLLEL.PKA.
AAD49343.1_DFR_Lilium_hybrid_division_VII ATV.RDP...................................RDLRKTKPLLDI.PGA.
AAF23884.2_DFR3_Lotus_corniculatus       ATV.RDP....................................ANMKKVKHLLEL.PEA.
BAE19948.1_DFR_Lotus_japonicus           ATVQRDP....................................DNMKKVKHLLEL.PGA.
BAE19949.1_DFR_Lotus_japonicus           ATV.RDP....................................ANMKKVKHLLEL.PEA.
BAE19950.1_DFR_Lotus_japonicus           ATVQRDP....................................DNMKKVKHLLEL.PGA.
BAE19951.1_DFR_Lotus_japonicus           ATVQRDP....................................ENMKKVKHLLEL.PGA.
BAE19953.1_DFR_Lotus_japonicus           ATI.RDP....................................ANMKKVKHLLEL.PDA.
BAA19658.1_DFR_Perilla_frutescens        ATV.RDP....................................GDSKKVKHLIEL.PGA.
BAB20075.1_DFR_Torenia_hybrid_cultivar   ATV.RDP....................................ENMKKVKHLLEL.PRAD
AAL35830.1_DFR_Triticum_monococcum       ATV.RDP....................................ANVEKNKPLLEL.PGA.
QFQ61498.1_DFR_Dryopteris_erythrosora    AAV.LDP....................................EDKQDVQPLLGLAPSL.
QFQ61499.1_DFR_Dryopteris_erythrosora    ATV.RNP....................................GDPTKTAILRNL.PGA.
BAA74700.1_DFR_Ipomoea_purpurea          ATV.RDP....................................GNTKKVKHLVEL.PKA.
BAF93896.1_DFR_Iris_x_hollandica         ATV.RDP....................................TNVEKTKPLLDL.PGA.
XP_008797532.1_DFR_Phoenix_dactylifera   ATV.RDP....................................TNLKKTQPLLDL.PGA.
AEQ92209.1_DFR_Ipomoea_batatas           ATV.RDP....................................GNTKKVKHLLEL.PKA.
BAH98155.1_DFR_Tulipa_gesneriana         ATV.RDP....................................KDQGKTKPLLDL.RGA.
APG32494.1_DFR2_Freesia_hybrid_cultivar  ATV.RDP....................................TNLRKTKPLLDL.SGA.
AAO63026.1_DFR_Allium_cepa               ATL.RDS....................................SDEAKTKPLLEL.PGA.
AFP58815.1_DFR_Hyacinthus_orientalis     ATV.RDP....................................ANTKKLKPLLDI.PGA.
                                   acc
```

```
                                          α3   β3        α4      β4            α5
NP_001268144.1_DFR_Vitis_vinifera                   TT
                                               60        70        80        90

NP_001268144.1_DFR_Vitis_vinifera          ETHLTLWKADLAD.EGSFDEAIKGCTGVFHVATPMDFESKDPE.................
TRINITY_DN27402_c0_g1_i3                   KTHMSLWKADLSV.EGSFDEPIQGCNGVFHVATPMDFESKDPE.................
NP_199094.1_DFR_Arabidopsis_thaliana       KTLLTLWKADLSE.EGSYDDAINGCDGVFHVATPMDFESKDPE.................
BAA85261.1_DFR_Arabidopsis_thaliana        KTQLTLWKADLSE.EGSYDDAINGCDGVFHVATPMDFESKDPE.................
AEI59122.1_DFR_Medicago_sativa             NSKLSLWKADLGE.EGSFDEAIKGCTGVFHVATPMDFESKDPE.................
BAD67185.1_DFR_Spinacia_oleracea           NTHLTLWKADLNE.QGSFDEAISGCAGVFHVATPMDFDSNDPE.................
BAB40789.1_DFR_Lilium_hybrid               DERLTIWKADLSE.DGSFDEAINGCTGVFHVATPMDFDSEDPE.................
CAA79154.1_DFR_Solanum_lycopersicum        DTNLTLWKADLAV.EGSFDEAIQGCQGVFHVATPMDFESKDPE.................
AHZ30596.1_DFR_Prunus_domestica            ETHLTLWKADLAD.EGSFDEAIQGCTGVFHVATPMDFESKDPE.................
AAD26204.1_DFR_Malus_domestica             ETHLTLWKADLAD.EGSFDEAIQGCSGVFHVATPMDFESKDPE.................
AAO39819.1_DFR_Pyrus_communis              ETHLTLWKADLAD.EGSFDEAIQGCSGVFHVATPMDFESKDPE.................
AAD56578.1_DFR_Daucus_carota               ETNLILCRADLNE.EGSFDDAVKGCHAVFHMATPMDIESQDPE.................
AKN56970.1_DFR_Gerbera_hybrid_cultivar     QTNLKLWKADLTQ.EGSFDEAVQGCHGVFHLATPMDFESKDPE.................
AOF39984.11_DFR_Brassica_rapa              KTQLTLWKADLSD.EGSYDDAINGCDGVFHIATPMDFESKDPE.................
P51106.1_DFR_Hordeum_vulgare               KERLSIWKADLSE.DGSFNEAIAGCTGVFHVATPMDFDSQDPE.................
BAE19953.1_DFR_Lotus_japonicus             KTKLSLWKADLAE.EGSFDEAIRGCTGVFHVATPMDFESKDPE.................
AII26023.1_DFR_Pisum_sativum               KSKLSLWKADLAE.EGSFDEAIKGCVGVFHVATPMDFESKDPE.................
NP_001274988.1_DFR_Solanum_tuberosum       DTNLTLWKADLAV.EGSFDEAIQGCQGVFHVATPMDFESKDPE.................
NP_001152467.2_DFR_Zea_mays                TERLSIWKADLAE.EGSFHDAIRGCTGVFHVATPMDFLSKDPE.................
AIZ74402.1_DFR_Anthurium_andraeanum        ANRLTLWKADLVD.EDSFDEPIQGCTGVFHVATPMDFESKDPE.................
AAB62873.1_DFR_Bromheadia_finlaysoniana    NELLSIWKADLNDIEGSFDEVIRGCVGVFHVATPMNFQSKDPE.................
CAA91922.1_DFR_Callistephus_chinensis      ETNLTLWKADLTQ.EGSFDEAIEGCHGVFHVATPMDFESKDPE.................
AIM58715.1_DFR_Cymbidium_hybrid_cultivar   NELLSIWKADLNDIDETFDEVTRGSVGLFHVATPMNFQSEDPE.................
ACZ48698.1_DFR_Fagopyrum_esculentum        KTNLSLWKADLSE.EGSFDEAIQGCAGVFHVATPMDFESKDPE.................
ACZ48697.1_DFR_Fagopyrum_tataricum         KTNLSLWKADLSE.EGSFDEAIQGCAGVFHVATPMDFESKDPE.................
BAO53730.1_DFR2_Glycine_max                KTKLSLWKADLAQ.EGSFDEAIKGCTGVFHVATPMDFDSKDPE.................
AAP13055.1_DFR_Gypsophila_elegans          KTNLTLWKADLNE.EGSFDKAVDGCSGVFHIATPMDFESKDPE.................
XP_013466134.1_DFR_Medicago_truncatula     KGKLSLWKADLGE.EGSFDEAIKGCTGVFHVATPMDFESKDPE.................
BAF96936.1_DFR_Nicotiana_tabacum           DTNLTLWKADLSV.EGSFDEAIQGCQGVFHVATPMDFESEDPE.................
BAA36183.1_DFR_Oryza_sativa_Japonica_Group KERLTLWKADLGE.EGSFDAAIRGCTGVFHVATPMDFESEDPE.................
BAA12723.1_DFR_Rosa_hybrid_cultivar        ATHLTLWKADLAE.EGSFDEAIKGCTGVFHVATPMDFESKDPE.................
AHL46438.1_DFR_Fragaria_vesca              ATLLTLWKADLDI.EGSFDEAIKGCTGVFHVATPMDFESEDPE.................
AHL46443.1_DFR_Fragaria_x_ananassa         ATRLTLWKADLDV.EGSFDEAIKGCTGVFHVATPMDFESEDPE.................
AHL46445.1_DFR_Fragaria_vesca              ATHLTLWKADLAD.EGSFDEAIKGCTGVFHVATPMDFESKDPE.................
AHM27144.1_DFR_Angelonia_angustifolia      DTNLTLWKADLME.EASFDEAIEGCQGVFHLATPMEFDSKDPE.................
AAT84073.1_DFR_Camellia_sinensis           DTNLTLWKADLNE.EGSFDEAIEGCSGVFHVATPMDFKSKDPE.................
AAS00611.1_DFR_Citrus_sinensis             STHLTLWKADLAE.EGNFDEAIRGCTGVFHLATPMDFESKDPE.................
AAX16491.1_DFR_Crataegus_monogyna          ETHLTLWKADLAD.EGSFDEAIQGCSGVFHVATPMDFESKDPE.................
BAF49325.1_DFR_Delphinium_belladonna       KSKLTLWKADLTE.DGSYDDAIKGCTGVFHVATPMDFESKDPE.................
CAA78930.1_DFR_Gerbera_hybrid_cultivar     QTNLKLWKADLTQ.EGSFDEAIQGCHGVFHLATPMDFESKDPE.................
AAO39816.1_DFR_Malus_domestica             ETHLTLWKADLAD.EGSFDEAIQGCSGVFHVATPMDFESKDPE.................
AAO39817.1_DFR_Malus_domestica             ETHLTLWKADLAD.EGSFDEAIQGCSGVFHVATPMDFESKDPE.................
AAR27015.1_DFR2_Medicago_truncatula        NSKLSLWKADLGE.EGSFDEAIKGCTGVFHVATPMDFESKDPE.................
AEF14420.1_DFR_Onobrychis_viciifolia       KTKLSLWKADLAH.EGSFDEAIKGCTGVFHVATPMDFDSKDPE.................
AAF60298.1_DFR_Petunia_x_hybrida           DTNLTLWKADLTV.EGSFDEAIQGCQGVFHVATPMDFESKDPE.................
AAQ77347.1_DFR_Triticum_aestivum           KERLSIWKADLSE.EGSFDDAIAGCTGVFHVATPMDFDSKDPE.................
AAL89715.1_DFR_Vaccinium_macrocarpon       DTNLTLWKADLNE.EGSFDEAIEGCVGVFHVATPMDFESKDPE.................
AAX12420.1_DFR_Vaccinium_macrocarpon       DTNLTLWKADLNE.EGSFDEAIEGCVGVFHVATPMDFESKDPE.................
CAA33543.1_DFR_Antirrhinum_majus           DTNLTLWKADMTV.EGSFDEAIQGCEGVFHLATSMEFDSVDPE.................
AAB62873.1_DFR_Bromheadia_finlaysoniana    NELLSIWKADLNDIEGSFDEVIRGCVGVFHVATPMNFQSKDPE.................
BAA84940.1_DFR_Camellia_sinensis           DTNLTLWKADLNE.EGSFDEAIEGCSGVFHVATPMDFESKDPE.................
AAC17843.1_DFR_Cymbidium_hybrid_cultivar   NELLSIWKADLNDIDETFDEVTRGSVGLFHVATPMNFQSEDPE.................
CAA91924.1_DFR_Dianthus_caryophyllus       KTNLTLWKADLHE.EGSFDAAVDGCTGVFHIATPMDFESKDPE.................
CAA70345.1_DFR_Forsythia_x_intermedia      DTNLTLWKADMTV.EGSFDEAIQGCEGVFHVATPMDFESKDPE.................
AAC25960.1_DFR_Fragaria_x_ananassa         ATRLTLWKADLDV.EGSFDEAIKGCTGVFHVATPMDFESEDPE.................
BAA12736.1_DFR_Gentiana_triflora           STNLTLLKADLTE.EGSFDEAIHGCHGVFHVATPMDFESKDPK.................
AAD54273.1_DFR1_Glycine_max                KSKLSLWKADLAE.EGSFDEAIKGCTGVFHVATPMDFESKDPE.................
BAA59332.1_DFR_Ipomoea_nil                 EGKLKVWKGVLEE.EGSFDEAIAGCEGVFHVAAAVNFASKDPE.................
BAA36405.1_DFR_Ipomoea_purpurea            EGKLKVWKGVLEE.EGSFDEAIAGCEGVFHVAAAVNFASKDPE.................
AAD49343.1_DFR_Lilium_hybrid_division_VII  DERLTIWKADLSE.DASFDEAINGCTGVYHVATPMDFDSKDPE.................
AAF23884.2_DFR3_Lotus_corniculatus         KTKPTLWKADLAE.EGSFDEAIKGCTGVFHVATPMDFESKNPE.................
BAE19948.1_DFR_Lotus_japonicus             KTNLTIWNADLTE.EGSFDEAIKGCSGVFHVASPMDFNSKDPE.................
BAE19949.1_DFR_Lotus_japonicus             KTKLTLWKADLAE.EGSFDEAIKGCTGVFHVATPMDFESKDPE.................
BAE19950.1_DFR_Lotus_japonicus             KTNLTIWNADLTE.EGSFDEAINGCSGVFHVASPMDFNSKDPE.................
BAE19951.1_DFR_Lotus_japonicus             KTNLTIWNADLTE.EGSFDEAINGCSGVFHLATPMDFNSKDPE.................
BAE19953.1_DFR_Lotus_japonicus             KTKLSLWKADLAE.EGSFDEAIRGCTGVFHVATPMDFESKDPE.................
BAA19658.1_DFR_Perilla_frutescens          DTNLTLWRADMNI.EGSYDEAVQGCECVFHMATPMDFESDDPE.................
BAB20075.1_DFR_Torenia_hybrid_cultivar     DASLRLFKADMNV.EGSFDEAVGSCECVFHMATPMDFESDDPE.................
AAL35830.1_DFR_Triticum_monococcum         KERLSIWKADLSD.QGSFDDAIAGCTGVFHVATPNGLRLPRPRGKYQYPGTLKQSITTEI
QFQ61498.1_DFR_Dryopteris_erythrosora      AERLEIVKADLRV.KGDFDKVAQGCDGVFHLACPTNFLVEDPQ.................
QFQ61499.1_DFR_Dryopteris_erythrosora      DERLQIVRGQLLE.ESGFDDAVNGCDGVFHCACPTEFSFEDPY.................
BAA74700.1_DFR_Ipomoea_purpurea            DTNLTVWKGVMEE.EGSFDEAIAGCEGVFHVATPMDFDSKDPE.................
BAF93896.1_DFR_Iris_x_hollandica           DALLTIWKADLGQ.DGSFDKAVAGCTAVFHVATPMDFESKDPE.................
XP_008797532.1_DFR_Phoenix_dactylifera     SERVTIWKADLDE.EGSFDEAIEGCSGVFHVATPMDFESKDPE.................
AEQ92209.1_DFR_Ipomoea_batatas             DTNLRVWKGVMEE.EGSFDEAIAGCEGVFHVATPMDFDSKDPE.................
BAH98155.1_DFR_Tulipa_gesneriana           DERLTIWKADLND.EGSFDNAINGCTGVFHVATPMDFESKDPE.................
APG32494.1_DFR2_Freesia_hybrid_cultivar    DRLLTIWKADLGE.EGSFDEVVQGCTAVFHVATPMDFESQDPE.................
AAO63026.1_DFR_Allium_cepa                 DTRLSLWEADLLQ.DGSFDHVISGSIAVFHVATPMDFDSIDPE.................
AFP58815.1_DFR_Hyacinthus_orientalis       GSRLTIWKADLND.EGSFDDVIDGCTGVFHVATPMDFESKDPE.................
                                      acc
```

```
                                                            α6                          β5        η1              β6
NP_001268144.1_DFR_Vitis_vinifera                                                                                    TT
                                       100       110       120       130       140

NP_001268144.1_DFR_Vitis_vinifera      ...NEVIKPTIEGMLGIMKSCAAAK.TVRRLVFTSSAGTVNI...QEHQLPVYDESCWSD
TRINITY_DN27402_c0_g1_i3               ...NEVIKPTINGVLDILKACVKAK.TVHRVVFTLSAGAVDV...QEKPKFMYDETCYSD
NP_199094.1_DFR_Arabidopsis_thaliana   ...NEVIKPTVNGMLGIMKACVKAK.TVRRFVFTSSAGTVNV...EEHQKNVYDENDWSD
BAA85261.1_DFR_Arabidopsis_thaliana    ...NEVIKPTVNGMLGIMKACVKAK.TVRRFVFTSSAGTVNV...EEHQKNVYDENDWSD
AEI59122.1_DFR_Medicago_sativa         ...KEVINPTINGLLDIMKACEKAK.TVRRLVFTSSAGTLDV...TEHQKSVIDETCWSD
BAD67185.1_DFR_Spinacia_oleracea       ...NEVIKPTINGMLDIMRSCAKAK..IRRLVFTSSAGTVNV...EEIQKPVYDETCWSD
BAB40789.1_DFR_Lilium_hybrid           ...NEVIKPTINGVIGIMKSCKKAG.TVKRIIFTSSAGTVNV...HEHQMPEYDESSWSD
CAA79154.1_DFR_Solanum_lycopersicum    ...NEVIKPTVRGMLSIIESCAKAN.TVKRLVFTSSAGTLDV...QEDQKLFYDETSWSD
AHZ30596.1_DFR_Prunus_domestica        ...NEVIKPTINGVLDILKACLKAK.TVRRLVFTSSAGTVNI...EENQKSFYDETNWSD
AAD26204.1_DFR_Malus_domestica         ...NEVIKPTINGLLDILKACQKAK.TVRKLVFTSSAGTVNV...EEHQKPVYDESNWSD
AAO39819.1_DFR_Pyrus_communis          ...NEVIKPTINGLLDILKACQKAK.TVRKLVFTSSAGTVNV...EEHQKPVYDESNWSD
AAD56578.1_DFR_Daucus_carota           ...NEVIKPTVQGVLDIITACAKAK.TVKKLIYTSSAGTVNV...REHQLPVYDESNWSD
AKN56970.1_DFR_Gerbera_hybrid_cultivar ...NEIIKPTIEGVLSIIRSCVKAK.TVKKLVFTSSAGTVNG...QEKQLHVYDESHWSD
AOF39984.11_DFR_Brassica_rapa          ...NEVIKPTVNGVLGIMKACDKAK.TVRRIVFTSSAGTVNV...EEHQKNVYDENDWSD
P51106.1_DFR_Hordeum_vulgare           ...NEVIKPTVEGMLSIMRACKEAG.TVKRIVFTSSAGSVNI...EERPRPAYDQDNWSD
BAE19953.1_DFR_Lotus_japonicus         ...NEVIKPTINGLLDILKACEKAK.TVRRLVFTSSAGTVDV...TEHPKPVIDETCWSD
AII26023.1_DFR_Pisum_sativum           ...NEVIKPTINGLLDIMKACEKAK.TVRRLVFTSSAGTVDV...TENPKPVIDETCWSD
NP_001274988.1_DFR_Solanum_tuberosum   ...NEVIKPTVRGMLSIIESCAKAN.TVKRLVFTSSAGTLDV...QEDQKLFYDETSWSD
NP_001152467.2_DFR_Zea_mays            ...NEVIKPTVEGMISIMRACKEAG.TVRRIVFTSSAGTVNL...EERQRPVYDEESWTD
AIZ74402.1_DFR_Anthurium_andraeanum    ...SEMIKPTIEGMLNVLRSCARASSTVRRVVFTSSAGTVSI...HEGRRHLYDETSWSD
AAB62873.1_DFR_Bromheadia_finlaysoniana ...NEVIKPAINGLLGILTSCKKAG.SVKRVIFTSSAGTVNV...EEHQAAVYDENSWSD
CAA91922.1_DFR_Callistephus_chinensis  ...NEIIKPTIEGILSIIRSCAKAK.TVKKLVYTSSAGTVNV...QETQLPVYDESHWSD
AIM58715.1_DFR_Cymbidium_hybrid_cultivar ...NEVIKPTISGLLGILRSCKRVG.TVKRVIFTSSAGTVNV...EEHQATVYDESSWSD
ACZ48698.1_DFR_Fagopyrum_esculentum    ...NEVIKPTINGMLDIMRACLKA..NVRKLVFTSSAGTVNV...EEKQKPVYDETCWSD
ACZ48697.1_DFR_Fagopyrum_tataricum     ...NEVIKPTINGMLDIMKACLKA..NVRKLVFTSSAGTVNV...EEKQKPVYDETCWSD
BAO53730.1_DFR2_Glycine_max            ...NEVIKPTINGLLDIMKACVKAK.TVRRLVFTSSAGTVDV...TEHPNPVIDENCWSD
AAP13055.1_DFR_Gypsophila_elegans      ...NEMIKPTINGMLDIMKSCAKAK..VKRVVFTSSGGTVNV...EETQKAEYDETCWSD
XP_013466134.1_DFR_Medicago_truncatula ...NEMIKPTIKGVLDIMKACLKAK.TVRRFIFTSSAGTLNV...TEDQKPLWDESCWSD
BAF96936.1_DFR_Nicotiana_tabacum       ...NEVIKPTVRGMLSIIESCAKAN.TVKRLVFTSSAGTLDA...QEHQKLFYDETSWSD
BAA36183.1_DFR_Oryza_sativa_Japonica_Group ...NEVVKPTVEGMLSIMRACRDAG.TVKRIVFTSSAGTVNI...EERQRPSYDHDDWSD
BAA12723.1_DFR_Rosa_hybrid_cultivar    ...NEVIKPTINGVLDIMQACLKAK.TVRRLVFTSSAGSVNV...EETQKPVYNESNWSD
AHL46438.1_DFR_Fragaria_vesca          ...NEVIKPTINGMLDIMKACLKAK.TVRRVVFTSSAGAVAI...EEHRKEVYSENNWSD
AHL46443.1_DFR_Fragaria_x_ananassa     ...NEVIKPTINGMLDIMKACLKAK.TVRRVVFTSSAGAVAI...EEHRKEVYSENNWSD
AHL46445.1_DFR_Fragaria_vesca          ...NEVIKPTINGVLDIMKACLKAK.TVRRLVFTASAGSVNV...EETQKPVYNESNWSD
AHM27144.1_DFR_Angelonia_angustifolia  ...NEVIKPTIEGMMSIIRSCAKAK.TVEKVIFTTSAGTLNV...EEHQKPFYDETSWSD
AAT84073.1_DFR_Camellia_sinensis       ...NEVIKPTINGVLSIIRSCTKAK.TVKRLVFTSSAGTVNV...QEHQQPVFDENNWSD
AAS00611.1_DFR_Citrus_sinensis         ...NEVIRPTINGMVSIMRACKNAK.TVRRLVFTSSAGTLDV...EEHRKPVYDETSWSD
AAX16491.1_DFR_Crataegus_monogyna      ...NEVIKPTINGLLDILKACQKAK.TVRKLVFTSSAGTVNV...EEHQKPVYDESNWSD
BAF49325.1_DFR_Delphinium_belladonna   ...NEVIKPTIEGMLGIMKSCVKAK.SVRRLVFTSSAGSVNV...EERQQAEYDENSWTD
CAA78930.1_DFR_Gerbera_hybrid_cultivar ...NEIIKPTIEGVLSIIRSCVKAK.TVKKLVFTSSAGTVNG...QEKQLHVYDESHWSD
AAO39816.1_DFR_Malus_domestica         ...NEVIKPTINGLLDILKACQKAK.TVRKLVFTSSAGTVNV...EEHQKPVYDESNWSD
AAO39817.1_DFR_Malus_domestica         ...NEVIKPTINGLLDILKACQKAK.TVRKLVFTSSAGTVNV...EEHQKPVYDESNWSD
AAR27015.1_DFR2_Medicago_truncatula    ...KEVINPTINGLLDIMKACKKAK.TVRRLVFTSSAGTLDV...TEQQNSVIDETCWSD
AEF14420.1_DFR_Onobrychis_viciifolia   ...NEVIKPTINGLLDIMKACEKAK.TVRRLVFTSSAGTVDV...TEHPKPVIDETCWSD
AAF60298.1_DFR_Petunia_x_hybrida       ...NEVIKPTVRGMLSIIESCAKAN.TVKRLVFTSSAGTLDV...QEQQKLFYDQTSWSD
AAQ77347.1_DFR_Triticum_aestivum       ...NEVIKPTVEGMLSIMRACKEAG.TVKRIVFTSSAGSVNI...EERQRPAYDQDNWSD
AAL89715.1_DFR_Vaccinium_macrocarpon   ...NEVIKPTINGVLSIIKSCTKAK.TVKRLVFTSSAGAVVD...QEHQPLVFDENNWSD
AAX12420.1_DFR_Vaccinium_macrocarpon   ...NEVIKPTINGVLSIIKSCTKAK.TVKRLVFTSSAGTVNV...QEHQQPTFDENNWSD
CAA33543.1_DFR_Antirrhinum_majus       ...NEVIKPTIDGMLNIIKSCVQAK.TVKKFIFTTSGGTVNV...EEHQKPVYDETDSSD
AAB62873.1_DFR_Bromheadia_finlaysoniana ...NEVIKPAINGLLGILTSCKKAG.SVKRVIFTSSAGTVNV...EEHQAAVYDENSWSD
BAA84940.1_DFR_Camellia_sinensis       ...NEVIKPTINGVLSIIRSCTKAK.TVKRLVFTSSAGTVNV...QEHQQPVFDENNWSD
AAC17843.1_DFR_Cymbidium_hybrid_cultivar ...NEVIKPTISGLLGILRSCKRVG.TVKRVIFTSSAGTVNV...EEHQATVYDESSWSD
CAA91924.1_DFR_Dianthus_caryophyllus   ...NEMIKPTINGMLDILKSCVKAK..LRRVVFTSSGGTVNV...EATQKPVYDETCWSA
CAA70345.1_DFR_Forsythia_x_intermedia  ...NEVIKPTIEGFLSLIRSCTKAK.TVKRIVFTSSAGTVNV...EEHQKSVYDETDYSD
AAC25960.1_DFR_Fragaria_x_ananassa     ...NEVIKPTINGMLDIMKACLKAK.TVRRLVFTSSAGAVAI...EEHPKEVYSENNWSD
BAA12736.1_DFR_Gentiana_triflora       ...NEVIKPTIDGFLSIIRSCVKAK.TVKKLVFTSSAGTVDV...QEQQKPVYDENDWSD
AAD54273.1_DFR1_Glycine_max            ...NEVIKPTINGVLDIMKACLKAK.TVRRLIFTSSAGTLNV...IERQKPVFDDTCWSD
BAA59332.1_DFR_Ipomoea_nil             ...NEVIKPAVKGILSIINSCAKAK.TVKKLVFTSSTAAVHI...KETQQLEYDESSWSD
BAA36405.1_DFR_Ipomoea_purpurea        ...NEVIKPAVKGILSIINSCAKAK.TVKKLVFTSSTAAVHI...KETQQLVYDESSWSD
AAD49343.1_DFR_Lilium_hybrid_division_VII ...NEVFQPTINGVLGIMKSCKKAG.TVKRVIFTSSAGTVNV...QENQMPEYDESSWSD
AAF23884.2_DFR3_Lotus_corniculatus     ...NEVIKPTINGVLDIMKACQKAK.TVRRLVFTSSAGTLNV...IEHQKQMFDESCWSD
BAE19948.1_DFR_Lotus_japonicus         ...NEVIKPAINGVLDIMKACLKAK.TVRRLVFTSSAGILSV...SERHKHMLDETCWGD
BAE19949.1_DFR_Lotus_japonicus         ...NEVIKPTINGVLDIMKACQKAK.TVRRLVFTSSAGTLNV...IEHQKQMFDESCWSD
BAE19950.1_DFR_Lotus_japonicus         ...NEVIKPSINGVLDIMKACQKAK.TVRRLVFTSSAGTLNA...VEHQKQMCDESCWSD
BAE19951.1_DFR_Lotus_japonicus         ...NEVIKPTINGVLDIMKACQKAK.TVRRLVFTSSAGTLDA...VEHQKQMFDESCWSD
BAE19953.1_DFR_Lotus_japonicus         ...NEVIKPTINGLLDILKACEKAK.TVRRLVFTSSAGTVDV...TEHPKPVIDETCWSD
BAA19658.1_DFR_Perilla_frutescens      ...NEVIKPTVDGMLSIMRSCTKAK.TVKKLIFTNSAGTLNV...EEHQKPVYNEANWSD
BAB20075.1_DFR_Torenia_hybrid_cultivar ...NEVIKPTVDGMLSIIRSCAKAQ.TVKRLIFTNSAGTLNV...EEHQKPVYDESNWSD
AAL35830.1_DFR_Triticum_monococcum     LLLNEVIKPTVEGMLSIMRACKEAG.TVKRIVFTSSAGSVNI...EERQRPAYDQDNWSD
QFQ61498.1_DFR_Dryopteris_erythrosora  ...KDVIEPAVQGTLNVLNACVKSS.SVKRVIHVSSAAAIRFSG.KEVAEQVFDESCWTD
QFQ61499.1_DFR_Dryopteris_erythrosora  ...KDCIEPAVNGTVNVLKACAKAP.SIKRIVFTSSAAAVRFTQLEDIAGQRFDENCWTD
BAA74700.1_DFR_Ipomoea_purpurea        ...NEVIKPTINGVLNIINSCVKAK.TVKRLVFTSSAGTLNV...QPQQKPVYDETCWSD
BAF93896.1_DFR_Iris_x_hollandica       ...NEVIKPTINGVLSIMRSCKKAG.TVKRVVFTSSAGTVDV...KEHQQTEYDESSWSD
XP_008797532.1_DFR_Phoenix_dactylifera ...NEVIKPTVTGLLSIMKSCKKAG.TVKRVIFTSSAGTVNV...QEHQHSEYEESSWSD
AEQ92209.1_DFR_Ipomoea_batatas         ...NEVIKPAINGVLNIINSCVKAK.TVKRLVFTSSAGTLNV...QPQQKPVYDESCWSD
BAH98155.1_DFR_Tulipa_gesneriana       ...NEVIKPTVNGVLSIMKSCKKAG.TVKRVIFTSSAGTVNV...QEEQMPEYDEDSWSD
APG32494.1_DFR2_Freesia_hybrid_cultivar ...NEVIKPAINGMLSIMRSCKRAG.SVERVIFTSSAGTVNV...EEHQRAEYDENCWSD
AAO63026.1_DFR_Allium_cepa             ...NEVIKPAVNGMLSIMKSCKKAG.TVKRVIFTSSAGTVNV...EEHQKPEYDENSWSD
AFP58815.1_DFR_Hyacinthus_orientalis   ...NEVIKPTVDGMLSILKSCKRAG.TVRRVIYTSSAGTVNV...EERQKPEYDENSWSD
                                  acc
```

```
                                         α7                  α8                         β7      β8          α9
NP_001268144.1_DFR_Vitis_vinifera
                                         150       160 ★     170       180       190       200

NP_001268144.1_DFR_Vitis_vinifera        MEFCRAKKMTAWMYFVSKTLAEQAAWKYAKENNIDFITIIPTLVVGPFIMSSMPPSLITA
TRINITY_DN27402_c0_g1_i3                 LDFILSKKMTGWMYFVAKTIAEKEAFKFAKENNLDFISIIPSLVVGQFLMPSLPPSLITA
NP_199094.1_DFR_Arabidopsis_thaliana     LEFIMSKKMTGWMYFVSKTLAEKAAWDFAEEKGLDFISIIPTLVVGPFITTSMPPSLITA
BAA85261.1_DFR_Arabidopsis_thaliana      LEFIMSKKMTGWMYFVSKSLAEKAAWDFAEEKGLDFISIIPTLVVGPFITTSMPPSLITA
AEI59122.1_DFR_Medicago_sativa           VEFCRRVKMTGWMYFVSKTLAEQEAWKFSKEHNIDFVSIIPPLVVGPFIMPSMPPSLITA
BAD67185.1_DFR_Spinacia_oleracea         MEFCSSKKMTGWMYFVSKTLAEKAAWKFAAENNLDFISIIPPLVVGPFITPTMPPSLITA
BAB40789.1_DFR_Lilium_hybrid             IDFIRRVKMTGWMYFVSKILAEKAAWDFAKENDIQFISIIPTLVVGPFITTTMPPSMLTA
CAA79154.1_DFR_Solanum_lycopersicum      LDFIYAKKMTGWMYFVSKILAEKAAMEEARKNNIDFISIIPPLVVGPFITSTFPPSLITA
AHZ30596.1_DFR_Prunus_domestica          VEFCRSVKMTGWMYFVSKTLAEQAAWKFAKENNIDFITIIPTLVIGPFLMPSMPPSLITG
AAD26204.1_DFR_Malus_domestica           VEFCRSVKMTGWMYFVSKTLAEQAAWKYAKENNIDFITIIPTLVIGPFLMPSMPPSLITG
AAO39819.1_DFR_Pyrus_communis            VEFCRSVKMTGWMYFVSKTLAEQAAWKYAKENNIDFITIIPTLVIGPFLMPSMPPSLITG
AAD56578.1_DFR_Daucus_carota             MDFIYSTKMTAWMYFVSKSLAEKAAWQAAEENNIQFISIIPTLVVGPFISPTFPPSLITA
AKN56970.1_DFR_Gerbera_hybrid_cultivar   LDFIYSKKMTAWMYFVSKTLAEKAAWDAMKGNNISFISIIPTLVVCPFITSTFPPSLVTA
AOF39984.11_DFR_Brassica_rapa            LDFIMSKKMTGWMYFMSKTLAEKAAWDYAKEKGIDFISIIPTLVIGPFITTSMPPSLVTA
P51106.1_DFR_Hordeum_vulgare             IDYCRRVKMTGWMYFVSKALAEKAAMEYASENGLDFISIIPTLVVGPFLSAGMPPSLVTA
BAE19953.1_DFR_Lotus_japonicus           IEFCLRVKMTGWMYFVSKTRAEQEAWKYAKEHNIDFVSVIPPLVVGPFLMPTMPPSLITA
AII26023.1_DFR_Pisum_sativum             VDFCRRVNMTGWMYFVSKTLAEQEAWKYSKEHKIDFVSIIPPLVVGPFLMPSMPPSLITA
NP_001274988.1_DFR_Solanum_tuberosum     LDFIYAKKMTGWMYFVSKILAEKAAMKEAKKNNINFISIIPPLVVGPFITPTFPPSLITA
NP_001152467.2_DFR_Zea_mays              VDFCRRVKMTGWMYFVSKTLAEKAALAYAAEHGLDLVTIIPTLVVGPFISASMPPSLITA
AIZ74402.1_DFR_Anthurium_andraeanum      VDFCRAKKMTGWMYFVSKTLAEKAAWDFAEKNNIDFISIIPTLVNGPFVMPTMPPSMLSA
AAB62873.1_DFR_Bromheadia_finlaysoniana  LHFVTRVKMTGWMYFVSKTLAEKAAWEFVKENAIHFIAIIPTLVVGSFITNEMPPSLITA
CAA91922.1_DFR_Callistephus_chinensis    LDFIYSKKMTAWMYFVSKTLAEKAAMEAAKENNIDFVSIIPPLVVGPFINPTFPPSLITA
AIM58715.1_DFR_Cymbidium_hybrid_cultivar LDFVTRVKMTGWMYFVSKTLAEKAAWEFVSDNDIHFITIIPTLVVGSFLISRMPPSLITA
ACZ48698.1_DFR_Fagopyrum_esculentum      VDFCRRVKMTGWMYFVSKTLAEQAAWKFAEENNMDFISIIPTLVVGPFIMPSFPPSLITA
ACZ48697.1_DFR_Fagopyrum_tataricum       VDFCRRVKMTGWMYFVSKTLAEQAAWKFAEENNMDFISIIPTLVVGPFIMPSFPPSLITA
BAO53730.1_DFR2_Glycine_max              VDFCTRVKMTGWMYFVSKTLAEQEAWKYAKEHNIDFISVIPPLVVGPFLMPTMPPSLITA
AAP13055.1_DFR_Gypsophila_elegans        LDFIRSVKMTGWMYFVSKILAEQAAWKYAAENNLDFISIIPPLVVGPFIMPSMPPSLITA
XP_013466134.1_DFR_Medicago_truncatula   VEFCRRVKMTGWMYFVSKTLAEQEAWKFAKEHNMDFITIIPPLVVGPFLIPTMPPSLITA
BAF96936.1_DFR_Nicotiana_tabacum         LDFIYAKKMTGWMYFVSKILAEKAAMEAAKKKNFDFISIIPPLVVGPFLTPTFPPSLITA
BAA36183.1_DFR_Oryza_sativa_Japonica_Group IDFCRRVKMTGWMYFVSKSLAEKAAMEYAREHGLDLISVIPTLVVGPFISNGMPPSHVTA
BAA12723.1_DFR_Rosa_hybrid_cultivar      VEFCRRVKMTGWMYFASKTLAEQEAWKFAKENNIDFITIIPTLVIGPFLMPSMPPSLITG
AHL46438.1_DFR_Fragaria_vesca            VVFCRKVKMTGWMYFVSKTLAEQAAWKFAKENNIDFITIIPTLVIGPFLAPSMPPSLISG
AHL46443.1_DFR_Fragaria_x_ananassa       VVFCRKVKMTGWMYFVSKTLAEQAAWKFAKENNIDFITIIPTLVIGPFLAPSMPPSLISG
AHL46445.1_DFR_Fragaria_vesca            VEFCRRVKMTGWMYFVSKTLAEQEAWKFAKENNIDFITIIPTLVIGPFLMPAMPPSLITG
AHM27144.1_DFR_Angelonia_angustifolia    LDFIYSKKMTGWMYFVSKYLAEKQAVEAAKLNNIEFISIIPPVVVGPFINPAFPPSLVTA
AAT84073.1_DFR_Camellia_sinensis         LDFINKKKMTGWMYFVSKTLAEKAAWEAAKENNIDFISIIPTLVVGPFIMPTFPPSLITA
AAS00611.1_DFR_Citrus_sinensis           LDFVRSVKMTGWMYFVSKTLAEQAAWKFAEENNIDFISIIPSLVVGPFLTSSMPPSLITA
AAX16491.1_DFR_Crataegus_monogyna        VEFCRSVKMTGWMYFVSKTLAEQAAWKYAKENNIDFITIIPTLVIGPFLMPSMPPSLITG
BAF49325.1_DFR_Delphinium_belladonna     VEFCRTRKMTGWMYFVSKTLAEKAAWEFAQQNNIDFISIIPTLVVGPFLMPSMPPSLITA
CAA78930.1_DFR_Gerbera_hybrid_cultivar   LDFIYSKKMTAWMYFVSKTLAEKAAWDATKGNNISFISIIPTLVVGPFITSTFPPSLVTA
AAO39816.1_DFR_Malus_domestica           VEFCRSVKMTGWMYFVSKTLAEQAAWKYAKENNIDFITIIPTLVIGPFLMPSMPPSLITG
AAO39817.1_DFR_Malus_domestica           VEFCRSVKMTGWMYFVSKTLAEQAAWKYAKENNIDFITIIPTLVIGPFLMPSMPPSLITG
AAR27015.1_DFR2_Medicago_truncatula      VEFCRRVKMTGWMYFVSKTLAEQEAWKFSKEHNIDFVSIIPPLVVGPFIMPSMPPSLITA
AEF14420.1_DFR_Onobrychis_viciifolia     IEFCRRVKMTGWMYFVSKTLAEQEAWKYAKEHNIDFISVIPPLVVGPFLMPTMPPSLITA
AAF60298.1_DFR_Petunia_x_hybrida         LDFIYAKKMTGWMYFASKILAEKAAMEEAKKKNIDFISIIPPLVVGPFITPTFPPSLITA
AAQ77347.1_DFR_Triticum_aestivum         IDFCRRVKMTGWMYFVSKSLAEKAAMEYASENGLDFISIIPTLVVGPFLSAGMPPSLVTA
AAL89715.1_DFR_Vaccinium_macrocarpon     VDFLYDKKMTGWTYFVSKTLAERAAMEAAKEISIDFISIIPTLVVGPFISPTFPPSLITV
AAX12420.1_DFR_Vaccinium_macrocarpon     LDFIYKIKMTGWMYFVSKTLAEKAAWEAAKENNIDFISIIPTLVVGPFIMPTFPPSLITA
CAA33543.1_DFR_Antirrhinum_majus         MDFINSKKMTGWMYFVSKILAEKAGMEAAKENNIDFISIIPPLVVGPFIMPTFPPSLITA
AAB62873.1_DFR_Bromheadia_finlaysoniana  LHFVTRVKMTGWMYFVSKTLAEKAAWEFVKENAIHFIAIIPTLVVGSFITNEMPPSLITA
BAA84940.1_DFR_Camellia_sinensis         LDFINKKKMTGWMYFVSKTLAEKAAWEAAKENNIDFISIIPTLVVGPFIMPTFPPSLITA
AAC17843.1_DFR_Cymbidium_hybrid_cultivar LDFVTRVKMTGWMYFVSKTLAEKAAWEFVSDNDIHFITIIPTLVVGSFLISRMPPSLITA
CAA91924.1_DFR_Dianthus_caryophyllus     LDFIRSVKMTGWMYFVSKILAEQAAWKYAAENNLEFISIIPPLVVGPFIMPSMPPSLITA
CAA70345.1_DFR_Forsythia_x_intermedia    LNFIYSKKMTGWMYFASKILAEKVAWEAAKENSIGFISIIPTLVVGPFIMPTFPPSLITA
AAC25960.1_DFR_Fragaria_x_ananassa       VVFCRKVKMTGWMYFVSKTLAEQAAWKFAKENNIDFITIIPTLVIGPFLAPSMPPSLISG
BAA12736.1_DFR_Gentiana_triflora         LDFINSTKMTGWMYFVSKILAEKAAWEVTKANDIGFISIIPTLVVGPFITTTFPPSLITA
AAD54273.1_DFR1_Glycine_max              VEFCRRVKMTGWMYFVSKTLAEKEAWKFAKEQGLDFITIIPPLVVGPFLMPTMPPSLITA
BAA59332.1_DFR_Ipomoea_nil               LDFIYANKMGGWMYFASKTLAEKEAWKAAKEKQIEFISIIPPLVIGPFLIPTFPLSLVTA
BAA36405.1_DFR_Ipomoea_purpurea          LDFIYANKMGGWMYFASKTLAEKEAWKAAKEKQIEFISIIPPLVIGPFLIPTFPLSLVTA
AAD49343.1_DFR_Lilium_hybrid_division_VII VDFCRRVKMTGWMYFVSKTLAEKAAWEFAKENDIQLISIIPTLVVGPFITSTMPPSMLTA
AAF23884.2_DFR3_Lotus_corniculatus       VEFCRRVKMTGWMYFVSKTLAEQEAWKFAKEHGIDFITIIPPLVVGSFLMPTMPPSLITA
BAE19948.1_DFR_Lotus_japonicus           LEFCKKVKMTGWMYFVSKELAEQEALKFAKENNIDFVSIIPSLVVGPFLMPTMPPSLYTA
BAE19949.1_DFR_Lotus_japonicus           VEFCRRVKMTGWMYFVSKTLAEQEAWKFAKEHGIDFITIIPPLVVGSFLMPTMPPSLITA
BAE19950.1_DFR_Lotus_japonicus           VEFCRRVKMTGWMYFVSKTLAEQEAWKFAQEHDIDFITIIPSLVVGSFLMPTLPPSLTTA
BAE19951.1_DFR_Lotus_japonicus           VEFCRRVKMTGWMYFVSKTLAEQEAWKFAKEHDIDFITTIPSLVVGSFLMPTMPPSLITA
BAE19953.1_DFR_Lotus_japonicus           IEFCLRVKMTGWMYFVSKTRAEQEAWKYAKEHNIDFVSVIPPLVVGPFLMPTMPPSLITA
BAA19658.1_DFR_Perilla_frutescens        LDFIYSKKMTGWMYFVSKILAEKEAMKAAKESNINFISIIPPVVVGPFFMPTFPPSLITA
BAB20075.1_DFR_Torenia_hybrid_cultivar   LDFIYSTKMTGWMYFVSKVLAEKAAIKACKENNIDFISIIPPVVVGPFIIDNWPPSLITA
AAL35830.1_DFR_Triticum_monococcum       IDFCRRVKMTGWMYFVSKALAEKAAMEYASENGLDFISIIPTLVVGPFLSAGMPPSLVTA
QFQ61498.1_DFR_Dryopteris_erythrosora    VDFCIDNKIPGWPYFVGKTLGEKAAVEFAKEHNLDLVVVNPSIVHGPFLLSNIPNSVKDC
QFQ61499.1_DFR_Dryopteris_erythrosora    IEMCERDKPHGWPYFVSKTLSEKKAFELAQEYNLDLVTILPPLVNGPFLIDKIPNSVADA
BAA74700.1_DFR_Ipomoea_purpurea          LDFIYAKKMTGWMYFASKILAEKEAWKATKEKKIDFISIIPPLVVGPFITPTFPPSLITA
BAF93896.1_DFR_Iris_x_hollandica         VDFCRRVKMTGWMYFVSKTLAERATWEFARENGIDFISIIPTLVVGPFITTTMPPSMVTA
XP_008797532.1_DFR_Phoenix_dactylifera   IEFCRRVKMTGWMYFVSKTLAEQAAWEFARENGIHFISIIPTLVVGPFISSSMPPSLITA
AEQ92209.1_DFR_Ipomoea_batatas           LDFIYAKKMTGWMYFASKILAEKEAWKATKEKKIDFISIIPPLVVGPFITPTFPPSLITA
BAH98155.1_DFR_Tulipa_gesneriana         IDFCRRVKMTGWMYFVSKTLAEKAAWEFAKDNDIQLISIIPTLVVGPFITTSMPPSMITA
APG32494.1_DFR2_Freesia_hybrid_cultivar  VEFCRRVKMTGWMYFVSKTLAEKAAWDFALENGIHLITIIPTLVVGPFITTTMPPSMITA
AAO63026.1_DFR_Allium_cepa               IDFCRRVKMTGWMYFVSKSLAEKAAWEFAKANGIDLVTIIPTLVVGAFITTAMPPSMITA
AFP58815.1_DFR_Hyacinthus_orientalis     IEFCRRVKMTGWMYFVSKSLAERAAWDFARENSMDLITIIPTLVVGPFITSTMPPSMITA
                                    acc
```

```
                                           α10     η2   α11      β9            α12               β10       β11        α13
*NP_001268144.1_DFR_Vitis_vinifera*
                                         210       220       230       240       250       260
NP_001268144.1_DFR_Vitis_vinifera        LSPITGNEAHYSIIRQGQFVHLDDLCNAHIYLFENPKAEGRYICSSHDCIILDLAKMLRE
TRINITY_DN27402_c0_g1_i3                 LSPITGTESHYHIIKQGQFVHLDDLCMAHIYLFENPKAQGRYMCSSHEATILELAKLLRE
NP_199094.1_DFR_Arabidopsis_thaliana     LSPITRNEAHYSIIRQGQYVHLDDLCNAHIFLYEQAAAKGRYICSSHDATILTISKFLRP
BAA85261.1_DFR_Arabidopsis_thaliana      LSPITRNEAHYSIIRQGQYVHLDDLCNAHIFLYEQAAAKGRYICSSHDATILTISKFLRP
AEI59122.1_DFR_Medicago_sativa           LSLITGNEAHYSIIKQGQYIHLHDLCLAHIFLFENPKAQGRYICCSHEATIHEVAKLINK
BAD67185.1_DFR_Spinacia_oleracea         LSPITRNEAHYSIIKQGQFVHLDDLCMAHIYLYEHPNAKGRYIASACDATIYDIGKMLRE
BAB40789.1_DFR_Lilium_hybrid             LSLITGNEAHYSILKQIQLVHLDDVCNAHIFLFENPEAIGRYICSSYDTTIYDLARKIKD
CAA79154.1_DFR_Solanum_lycopersicum      LSLIT...AHYGIIKQGQYVHLDDLCEAHIFLYEHPKAEGRFICSSHHAIIYDVAKMVRQ
AHZ30596.1_DFR_Prunus_domestica          LSPLTGNTSHYSIIKRGQFVHLDDLCLSHIYLYEHPKAEGRYICSSHDATIYDIAKLLRE
AAD26204.1_DFR_Malus_domestica           LSPILRNESHYGIIKQGQYVHLDDLCLSHIYLYEHPKAEGRYICSSHDATIHELVKMLRE
AAO39819.1_DFR_Pyrus_communis            LSPILRNESHYGIIKQGQYVHLDDLCLSHIYLYKHPKAEGRYICSSHDATIHELVKMLRE
AAD56578.1_DFR_Daucus_carota             LSPLTGNEAHYSIIKQCQYVHLDDLCESQIFLYEHPTAQGRYICSSHDATIHDVVKLIRE
AKN56970.1_DFR_Gerbera_hybrid_cultivar   LSLITGNEAHYSIIKQGQYVHLDDLCECHIYLYENPKAKGRYICSSHDATIHQLAKIIQD
AOF39984.11_DFR_Brassica_rapa            LSPITRNEAHYSIIRQGQYVHLDDLCNAHIFLYEQAAAKGRYVCSSHDATILTISEFLRQ
P51106.1_DFR_Hordeum_vulgare             LALITGNEAHYSILKQVQLVHLDDLCDAMTFLFEHPEANGRYICSSHDATIHGLARMLQD
BAE19953.1_DFR_Lotus_japonicus           LSLITGNEAHYSIIKQGQYVHLDDLCLAHIFLFENPKAQGRYMCSAYEATIHEVARMINK
AII26023.1_DFR_Pisum_sativum             LSLITGNEAHYTIIKQGQYVHLDDLCLAHIFLFENPKAQGRYICCSHEATIHEVATLIKQ
NP_001274988.1_DFR_Solanum_tuberosum     LSLITGNEAHYGIIKQGQYVHLDDLCEAHIFLYEHPKAEGRFICSSHHAIIYDVAKMVRQ
NP_001152467.2_DFR_Zea_mays              LALITGNAPHYSILKQVQLIHLDDLCDAEIFLFENPAAAGRYVCSSHDVTIHGLAAMLRD
AIZ74402.1_DFR_Anthurium_andraeanum      LALITRNEPHYSILNPVQFVHLDDLCNAHIFLFECPDAKGRYICSSHDVTIAGLAQILRQ
AAB62873.1_DFR_Bromheadia_finlaysoniana  LSLISGNEAHYSILKQAQFVHLDDLCDAHIFVYEHPEANGRYICSSHDSTIYDLANMLKN
CAA91922.1_DFR_Callistephus_chinensis    LSLINGAESHYSIIKQGQYVHLDDLCECHIFLYENPEAKGRYICSKQDATIHQLARMIKQ
AIM58715.1_DFR_Cymbidium_hybrid_cultivar LSLITGNEAHYSILRQAQFVHLDDLCDAHIFLFEHHKANGRYICSSHDSTIYSLAKMLKN
ACZ48698.1_DFR_Fagopyrum_esculentum      LSPITRTEGHYTIIKQCQYVHLDDLCMSHIYLYEKAGSKGRYVCSSDNATIYDLGKMLRN
ACZ48697.1_DFR_Fagopyrum_tataricum       LSPITRTEGHYTIIKQCQYVHLDDLCMSHIYLYEKAGSKGRYVCSSHNATIYDLGKMLRN
BAO53730.1_DFR2_Glycine_max              LSLITGNESHYHIIKQGQFVHLDDLCLGHIFVFENPKAEGRYICCSHEATIHDIAKLLNQ
AAP13055.1_DFR_Gypsophila_elegans        LSPITRTESHYTIIKQGQFVHLDDLCMAHIFLYENPKAQGRYIASACDATIYDIAKMLRE
XP_013466134.1_DFR_Medicago_truncatula   LSPITGNEAHYSIIKQGQFVHLDDLCEAHIFLFEHMEVEGRYLCSACEANIHDIAKLINT
BAF96936.1_DFR_Nicotiana_tabacum         LSLITGNEAHYCIIKQGQYVHLDDLCEAHIFLYEQPKAEGRFICASHHAIIYDVAKMVRE
BAA36183.1_DFR_Oryza_sativa_Japonica_Group LALLTGNEAHYSILKQVQFVHLDDLCDAEIFLFESPEARGRYVCSSHDATIHGLATMLAD
BAA12723.1_DFR_Rosa_hybrid_cultivar      LSPLTGNESHYSIIKQGQFIHLDDLCQSHIYLYEHPKAEGRYICSSHDATIHEIAKLLKG
AHL46438.1_DFR_Fragaria_vesca            LSPLTGNEAHYGIIKQCQYVHLDDLCQSHIFLYEHPKAEGRYICSSHDATIHDIAKLLNE
AHL46443.1_DFR_Fragaria_x_ananassa       LSPLTGNEAHYGIIKQCQYVHLDDLCQSHIFLYEHPKAEGRYICSSHDATIHDIAKLLNE
AHL46445.1_DFR_Fragaria_vesca            LLPLFGNEGHYSIIKQGQFVHLDDLCQSHIYLYEHPKAEGRYICSSHDATIHEIGKLLRE
AHM27144.1_DFR_Angelonia_angustifolia    LSPITGNEAHYTIMKQGQFVHLDDICEAHIFLFEHPKADGRYICSSHDATIYDIARLIRD
AAT84073.1_DFR_Camellia_sinensis         LSPITRNEGHYSIIKQGQFVHLDDLCESHIFLYERPQAEGRYICSSHDATIHDLAKLMRE
AAS00611.1_DFR_Citrus_sinensis           LSPITRNEAHYPIIKQGQFVHLDDLCSAHIFLFEHPNAKGRYICSSHPATILELAKFLRE
AAX16491.1_DFR_Crataegus_monogyna        LSPILRNESHYGIIKQGQYVHLDDLCLSHIYLYKHPKAEGRYICSSHDATIHELVKMLRE
BAF49325.1_DFR_Delphinium_belladonna     LSPITGNQSHYSILKQIQLVHLDDLCNAHIYLFEHPESNGRYICSSHDATITDVANLLRH
CAA78930.1_DFR_Gerbera_hybrid_cultivar   LSLITGNEAHYSIIKQGQYVHLDDLCECHIYLYENPKAKGRYICSSHDATIHQLAKIIKD
AAO39816.1_DFR_Malus_domestica           LSPILRNESHYGIIKQGQYVHLDDLCLSHIYLYEHPKAEGRYICSSHDATIHELVKMLRE
AAO39817.1_DFR_Malus_domestica           LSPILRNESHYGIIKQGQYVHLDDLCLSHIYLYEHPKAEGRYICSSHDATIHELVKMLRE
AAR27015.1_DFR2_Medicago_truncatula      LSLITGYEAHYSIIKQGQYIHLDDLCLAHIFLFENPKAHGRYICCSHEATIHEVAKLINK
AEF14420.1_DFR_Onobrychis_viciifolia     LSLITGNESHYHIIKQGQYVHLDDLCLGHIFLFEHPKPEGRYICCSDEATIHDVAKLINK
AAF60298.1_DFR_Petunia_x_hybrida         LSLITGNEAHYCIIKQGQYVHLDDLCEAHIFLYEHPKADGRFICSSHHAIIYDVAKMVRE
AAQ77347.1_DFR_Triticum_aestivum         LALITGNEAHYSILKQVQLVHLDDLCDAMTFLFEHPEANGRYICSSHDATIHGLARMLRD
AAL89715.1_DFR_Vaccinium_macrocarpon     LSPITGNDPHYSIIKQGQFVHLDDLCESHIYLFEHPEAEGRYICSSHDATIYDLAKIMRR
AAX12420.1_DFR_Vaccinium_macrocarpon     LSPITGNEPHYSIIKQGQFVHLDDLCESHIYLFEHPEAEGRYICSSHDATIYDLAKIMRE
CAA33543.1_DFR_Antirrhinum_majus         LSPITGNEAHYSIIKQCQYVHLDDLCEGHIFLFEYPKAEGRYICSSHDATIYDIAKLITE
AAB62873.1_DFR_Bromheadia_finlaysoniana  LSLISGNEAHYSILKQAQFVHLDDLCDAHIFVYEHPEANGRYICSSHDSTIYDLANMLKN
BAA84940.1_DFR_Camellia_sinensis         LSPITRNEGHYSIIKQGQFVHLDDLCESHIFLYERPQAEGRYICSSHDATIHDLAKLMRE
AAC17843.1_DFR_Cymbidium_hybrid_cultivar LSLITGNEAHYSILRQAQFVHLDDLCDAHIFLFEHHKANGRYICSSHDSTIYSLAKMLKN
CAA91924.1_DFR_Dianthus_caryophyllus     LSPITRTESHYTIIKQGQFVHLDDLCMSHIFLYENPKANGRYIASACAATIYDIAKMLRE
CAA70345.1_DFR_Forsythia_x_intermedia    LSPITGNEAHYSIIDQCQYVHLDDLCEAHIFLYEYPNAEGRYICSSHDATIYDVAKMIRD
AAC25960.1_DFR_Fragaria_x_ananassa       LSPLTGNEAHYGIIKQCQYVHLDDLCQSHIFLYEHAKAEGRYICSSHDATIHDIAKLLNE
BAA12736.1_DFR_Gentiana_triflora         LSLITGNEAHYGIIKQGQFVHLDDLCEAHIFLYEHPEAEGRYICSSHDTTIHDLAKMIRQ
AAD54273.1_DFR1_Glycine_max              LSPITGNEDHYSIIKQGQFVHLDDLCLAHIFLFEEPEVEGRYICSACDATIHDIAKLINQ
BAA59332.1_DFR_Ipomoea_nil               LSPIMGNGLHHNIIKQGKFVHLDDLCEAQIFLYQHPKAGGRFICSSHHATIHDVAKMIRH
BAA36405.1_DFR_Ipomoea_purpurea          LSPIMGNGLHHNIIKQGKFVHLDDLCEAQIFLYQHPEAGGRFICSSHHATIHDVAKMIRH
AAD49343.1_DFR_Lilium_hybrid_division_VII LSLITGNEAHYSILKQIQLVHLDDVCKAHIFLFENPEASGRYICSSYDATIYDLARKIKD
AAF23884.2_DFR3_Lotus_corniculatus       LSPITGNEAHYSIIKQGQYVHLDDLCLAHIFLFEHPESEGRYICSASEATIHDIAKLINS
BAE19948.1_DFR_Lotus_japonicus           LCPITGNEAHYMIMKQSQFVHVDDLCLAHIFLFEHPESEGRYMCSACDANIHDIAKLINT
BAE19949.1_DFR_Lotus_japonicus           LSPITGNEAHYSIIKQGQYVHLDDLCLAHIFLFEHPESEGRYICSASEATIHDIAKLINS
BAE19950.1_DFR_Lotus_japonicus           LSPITGNEAHYSIIKQGQYVHLDDLCLAHIFLFEHPKSEGRYICSASEATIHDIAKLINS
BAE19951.1_DFR_Lotus_japonicus           LSPITGNEAHYSIIKQGQFVHLDDLCLAHIFLFEHPESEGRYILSASEATIHDIAKLINS
BAE19953.1_DFR_Lotus_japonicus           LSLITGNEAHYSIIKQGQYVHLDDLCLAHIFLFENPKAQGRYMCSAYEATIHEVARMINK
BAA19658.1_DFR_Perilla_frutescens        LSPITGNEAHYSIIKQGQFVHVDDLCEAHIFLFEHPAAEGRYICSSHDATIYDIAKMIGE
BAB20075.1_DFR_Torenia_hybrid_cultivar   LSPITGNEAHYSIIKQGQFVHVDDLCEAHIFLSEHPKTEERYICSSHDATIYDIANMIRE
AAL35830.1_DFR_Triticum_monococcum       LALITGNEAHYSILKQVQLVHLDDLCDAMTFLFEHPEANGRYICSSHDATIHGLATMLRD
QFQ61498.1_DFR_Dryopteris_erythrosora    LALITGELFKLPFLARMSYIHTDDLVRAFIFLYENPEASGRYICSAVDASITEVVNIISS
QFQ61499.1_DFR_Dryopteris_erythrosora    ISLVTGDISHHKFIRRISYVHIDDIIDAHIFLYETPAAQGRYNGSCADASITEIAELLKK
BAA74700.1_DFR_Ipomoea_purpurea          LSLITGNQAHYSIIKQGQYVHLDDLCEAHIFLYEHPKAEGRFICSSHHTTIHGLAEMITQ
BAF93896.1_DFR_Iris_x_hollandica         LSFMTGNEAHYHIIKHAQLVHLDDLCAAHIYLLNRPEANGRYICSSHEATIHDLARMVRE
XP_008797532.1_DFR_Phoenix_dactylifera   LSLITGNEAHYSILKQIQFVHLDDLCVAHIFLFEHPEANGRYICSSHDVTIYELARMMKE
AEQ92209.1_DFR_Ipomoea_batatas           LSLITGNQAHYSIIKQGQYVHLDDLCEAHIFLYEHPKAEGRFICSSHHTTIHGLGEMIRQ
BAH98155.1_DFR_Tulipa_gesneriana         LSLITGNDSHYSILKQIQLVHLDDLCIAHIFLFENQEASGRYICSSFDATIWDLARLMKD
APG32494.1_DFR2_Freesia_hybrid_cultivar  LSLITGNEGHYSIIKQAQLVHLDDLCAAHIYLLNHPEAKGRYICSSHDATIYDLARMLRE
AAO63026.1_DFR_Allium_cepa               LSLITGNEAHYSIIKQAQLVHLDDLCEAHILLLNHPKAEGRYICSSHDVTIYDMAKMIRQ
AFP58815.1_DFR_Hyacinthus_orientalis     LSLITGNEAHYSIIKQVQLVHLDDLCEAHVFLLDHPKANGRYICSSHDATIHDLAGMIRR
                                  acc
```

```
                                                                          β12   α14                 α15
NP_001268144.1_DFR_Vitis_vinifera                      TT    TT                   
                                  270         280          290          300             310  ★      320
NP_001268144.1_DFR_Vitis_vinifera                 KYPEY.NIPTEFKGVD.ENLKSVCFSSKKLTDLGF..EFKY.SLEDMFTGAVDTCRA.KG
TRINITY_DN27402_c0_g1_i3                          KFPEN.NIPSKFKDVD.ENLQNIVFSSKKLTDMGF..EFKY.SLEDMFSAAVETCRA.KG
NP_199094.1_DFR_Arabidopsis_thaliana              KYPEY.NVPSTFEGVD.ENLKSIEFSSKKLTDMGF..NFKY.SLEEMFIESIETCRQ.KG
BAA85261.1_DFR_Arabidopsis_thaliana               KYPEY.NVPSTFEGVD.ENLKSIEFSSKKLTEMGF..NFKY.SLEEMFIESIETCRQ.KG
AEI59122.1_DFR_Medicago_sativa                    KYPEF.NVPTKFKDIP.DELEIIKFSSKKITDLGF..KFEY.SLEDMFTGAIETCRE.KG
BAD67185.1_DFR_Spinacia_oleracea                  EYPEY.NIPTKFKDFE.EDMEHVHFSSEKLMGLNF..KFKY.ELKDMYKGAVDTCRT.KG
BAB40789.1_DFR_Lilium_hybrid                      RYPQY.AIPQKFEGID.DQIKPVHFSSKKLIDLGF..KYQY.TFEEMFDEGIRSCIE.KN
CAA79154.1_DFR_Solanum_lycopersicum               KWPEY.YVPTEFKGID.KDLALVSFSSKKLMDIKF..QFKH.TLEDMYKGAIETCRQ.KQ
AHZ30596.1_DFR_Prunus_domestica                   KYPEY.NIPTKFDNIE.ENLTNIHFSSKKLTDQGF..EFKY.NLDDMFVGAVDTCRA.KG
AAD26204.1_DFR_Malus_domestica                    KYPEY.NIPTKFKGID.DNLEPVHFSSKKLREIGF..EFKY.SLEDMFVGAVDACRA.KG
AAO39819.1_DFR_Pyrus_communis                     KYPEY.NIPTKFKGID.DNLEPVHFSSKKLREIGF..EFKY.SLEDMFVGAVDACRA.KG
AAD56578.1_DFR_Daucus_carota                      KWPEY.NVPTEFKGIE.KDLPVISFSSKKLTDMGF..TFKY.NLEDMFKGAIETCRE.KG
AKN56970.1_DFR_Gerbera_hybrid_cultivar            KWPEY.YISTKFPGID.EELPIVSFSSKKLIDTGF..EFKY.NLEDMFKGAIDTCRE.KG
AOF39984.11_DFR_Brassica_rapa                     KYPEY.NVPSTFEGVD.ENLKSIMFSSKKPIDMGF..NFKY.SLEDMLVESIETCRQ.KG
P51106.1_DFR_Hordeum_vulgare                      RFPEY.DIPQKFAGVD.DNLQPIHFSSKKLLDHGF..SFRY.TTEDMFDAAIHTCRD.KG
BAE19953.1_DFR_Lotus_japonicus                    KYPEF.NVPTKFKDIP.DELDIIKFSSKKITDLGF..KFKY.SLEDMYTGAVETCRE.KG
AII26023.1_DFR_Pisum_sativum                      KYPEF.NVPTKFKDIP.DELEIIKFCSKKITDLGF..KFKY.SLEDMFTGAVETCRE.KG
NP_001274988.1_DFR_Solanum_tuberosum              KWPEY.YVPTEFKGID.KDLPIVSFSSKKLMDMGF..QFKY.TLEDMYKGAIETCRQ.KQ
NP_001152467.2_DFR_Zea_mays                       RYPEY.DVPQRFPGIQ.DDLQPVRFSSKKLQDLGF..TFRYKTLEDMFDAAIRTCQE.KG
AIZ74402.1_DFR_Anthurium_andraeanum               RYPEF.DVPTEFGDM..EVFDIISYSSKKLTDLGF..EFKY.SLEDMFDGAIQSCRE.KG
AAB62873.1_DFR_Bromheadia_finlaysoniana           RYATY.AIPQKFKEID.PNIKSVSFSSKKLMDLGF..KYKY.TIEEMFDDAIKTCRD.KN
CAA91922.1_DFR_Callistephus_chinensis             KWPEY.HVPTQFAGID.EELPTVSFSSKKLIDMGF..KFKY.DLEDMFKGAIDSCKE.KG
AIM58715.1_DFR_Cymbidium_hybrid_cultivar          RYATY.DIPLKFKEID.PNIESVSFSSKKLLDLGFKYKYKY.TMEEMFDDAIKTCRD.KN
ACZ48698.1_DFR_Fagopyrum_esculentum               KYPEY.NVPTKFRDFD.ENMEAVSFSSKKLTDEGF..EFKY.SLEDMFVGAVETCRE.KG
ACZ48697.1_DFR_Fagopyrum_tataricum                KYPEY.NVPTKFRDFD.ENMEAVSFSSKKLTDEGF..EFKY.SLEDMFVGAVETCRE.KG
BAO53730.1_DFR2_Glycine_max                       KYPEY.NVLTKFKNIP.DELDIIKFSSKKITDLGF..KFKY.SLEDMFTGAVETCRE.KG
AAP13055.1_DFR_Gypsophila_elegans                 EYPEY.NVPTKFKDYK.EDMDLVHFSSKKLTELGF..EFKY.GLKDMYTGAVETCRA.KG
XP_013466134.1_DFR_Medicago_truncatula            KYPEY.NIPTKFNNIP.DELELVRFSSKKIKDLGF..EFKY.SLEDMYTEAIDTCIE.KG
BAF96936.1_DFR_Nicotiana_tabacum                  KWPEY.YVPTEFKGID.KDLPVVYFSPKKLTDMGF..QFKY.TLEDMYKGAIETCRQ.KQ
BAA36183.1_DFR_Oryza_sativa_Japonica_Group        MFPEY.DVPRSFPGIDADHLQPVHFSSWKLLAHGF..RFRY.TLEDMFEAAVRTCRE.KG
BAA12723.1_DFR_Rosa_hybrid_cultivar               KYPEY.NVPTTFKGIE.ENLPKVHFSSKKLLETGF..EFKY.SLEDMFVGAVDACKE.KG
AHL46438.1_DFR_Fragaria_vesca                     RYPKY.NVPKKFKGIE.ENLTNIHFSSKKLKEMGF..EFKH.SLEDMFTGAVDACRA.KG
AHL46443.1_DFR_Fragaria_x_ananassa                KYPKY.NVPKKFKGIE.ENLTNIHFSSKKLKEMGF..EFKH.SLEDMFTGAVDACRE.KG
AHL46445.1_DFR_Fragaria_vesca                     KYPEY.NVPTKFKGIE.DNLTKVHFSSKKLLETGF..EFKY.SLEDMFVGAVDACKA.KG
AHM27144.1_DFR_Angelonia_angustifolia             KWPEY.NVPMEFKGID.KDIPVVRFSSKKLTSMGF..KFKY.SLEDMYRGAIETCRE.KG
AAT84073.1_DFR_Camellia_sinensis                  KWPEY.NVPTEFKGID.KDLPVVSFSSKKLIGMGF..EFKY.SLEDMFRGAIDTCRE.KG
AAS00611.1_DFR_Citrus_sinensis                    KYPEF.NVPTEFEDVD.ENMKNMLFSSKKLTDLGF..KFKY.SLDDMFTGAVDTCRA.KG
AAX16491.1_DFR_Crataegus_monogyna                 KYPEY.NIPTKFKGID.DNLEPVHFSSKKLREIGF..EFKY.SLEDMFVGAVDACRA.KG
BAF49325.1_DFR_Delphinium_belladonna              KFPEY.NVPTKFMGVD.ENLKAVVFSSKRLKDLGF..NYKY.TMEDMFVGAVTTCRR.NG
CAA78930.1_DFR_Gerbera_hybrid_cultivar            KWPEY.YIPTKFPGID.EELPIVSFSSKKLIDTGF..EFKY.NLEDMFKGAIDTCRE.KG
AAO39816.1_DFR_Malus_domestica                    KYPEY.NIPTKFKGID.DNLEPVHFSSKKLREIGF..EFKY.SLEDMFVGAVDACRA.KG
AAO39817.1_DFR_Malus_domestica                    KYPEY.NIPTKFKGID.DNLEPVHFSSKKLREIGF..EFKY.SLEDMFVGAVDACRA.KG
AAR27015.1_DFR2_Medicago_truncatula               KYPEF.NVPTKFKDIP.DDLEIIKFSSKKITDLGF..IFKY.SLEDMFTGAIETCRE.KG
AEF14420.1_DFR_Onobrychis_viciifolia              IYPEF.NVPTKFKDIP.DELEIIKFSSKKITDLGF..KFKY.TLEDMLTGAVETCRE.KG
AAF60298.1_DFR_Petunia_x_hybrida                  KWPEY.YVPTEFKGID.KDLPVVSFSSKKLTDMGF..QFKY.TLEDMYKGAIDTCRQ.KQ
AAQ77347.1_DFR_Triticum_aestivum                  RFPEY.SIPHKFAGVG.DDLQPIHFSSKKLLDHGF..SFRY.TAEDMFDAAIRTCRE.KG
AAL89715.1_DFR_Vaccinium_macrocarpon              NAPST.MSPTEFKGIP.KDLPNVSFPSKKLIGMGF..QFKY.SLEDMFRGAIDTCQETKG
AAX12420.1_DFR_Vaccinium_macrocarpon              KWPEY.NVPTEFKGIP.KDLPNVSFSSKKLIGMGF..QFKY.SLEDMFRGAIDTCRE.KG
CAA33543.1_DFR_Antirrhinum_majus                  NWPEY.HIPDEFEGID.KDIPVVSFSSKKMIGMGF..IFKY.TLEDMVRGAIDTCRE.KG
AAB62873.1_DFR_Bromheadia_finlaysoniana           RYATY.AIPQKFKEID.PNIKSVSFSSKKLMDLGF..KYKY.TIEEMFDDAIKTCRD.KN
BAA84940.1_DFR_Camellia_sinensis                  KWPEY.NVPTEFKGID.KDLPVVSFSSKKLIGMGF..EFKY.SLEDMFRGAIDTCRE.KG
AAC17843.1_DFR_Cymbidium_hybrid_cultivar          RYATY.DIPLKFKEID.PNIESVSFSSKKLLDLGFKYKYKY.TMEEMFDDAIKTCRD.KN
CAA91924.1_DFR_Dianthus_caryophyllus              EYPEY.NVPTKFKDYK.EDMGQVQFSSKKLTDLGF..EFKY.GLKDMYTAAVESCRA.KG
CAA70345.1_DFR_Forsythia_x_intermedia             KWPKY.DIPTEFEGIG.KDIPVVSFSSKKLIGMGF..TFKY.TLEDMYRGAIETCRE.KG
AAC25960.1_DFR_Fragaria_x_ananassa                KYPKY.NVPKKFKGIE.ENLTNIHFSSKKLKEMGF..EFKH.SLEDMFTGAVDACRE.KG
BAA12736.1_DFR_Gentiana_triflora                  NWPEY.YIPTKLKGID.EDIPVVSFSSNKLIDLGF..QYKY.TLEDMFRGAIDTCKE.KR
AAD54273.1_DFR1_Glycine_max                       KYPEY.KVPTKFKNIP.DQLELVRFSSKKITDLGF..KFKY.SLEDMYTGAIDTCRD.KG
BAA59332.1_DFR_Ipomoea_nil                        NWPEY.YVPSEFKGIE.KELPIVSFSSKKLQEMGF..QFKY.TLEDMYKGAIETLRK.KG
BAA36405.1_DFR_Ipomoea_purpurea                   NWPEY.YVPSEFKGIE.KELPIVSFSSKKLQEMGF..QFKY.TLEDMYKGAIETLRK.KG
AAD49343.1_DFR_Lilium_hybrid_division_VII         RYPKY.AIPQKFEGID.DQIKPVHFSSKKLMDLGF..KYQY.TFEEMFDEGIRSCIE.KK
AAF23884.2_DFR3_Lotus_corniculatus                KYPEY.NIPTKFKNIP.DELELVRFSSKKIKDMGF..EFKY.SLEDMYTGAIDTCKE.KG
BAE19948.1_DFR_Lotus_japonicus                    KYPEY.NVPTKFKNIP.DELELVRFSSKKIKDMGF..QFKY.TLEDMYTGAIDACRE.KG
BAE19949.1_DFR_Lotus_japonicus                    KYPEY.NIPTKFKNIP.DELELVRFSSKKIKDMGF..EFKY.SLEDMYTGAIDTCKE.KG
BAE19950.1_DFR_Lotus_japonicus                    KYPEY.NVPTKFKNIP.DELELVRFSSKKIKDMGF..EFKY.SLEDMYTGAIDTCKE.KG
BAE19951.1_DFR_Lotus_japonicus                    KYPEY.NVPTKI......................F..QMNW.SLSDFHQR.....RS.K.
BAE19953.1_DFR_Lotus_japonicus                    KYPEF.NVPTKFKDIP.DELDIIKFSSKKITDLGF..KFKY.SLEDMYTGAVETCRE.KG
BAA19658.1_DFR_Perilla_frutescens                 NWPEY.HVPTEFEGIH.KDIAVVSFSSKKLVDMGF..SFKY.TLEDMYRGAIESCRE.KG
BAB20075.1_DFR_Torenia_hybrid_cultivar            KWPEY.DVPTEFEGID.KDIPVVRFSSKKLMGMGF..TFKY.TLEDMFREAIETCRD.KG
AAL35830.1_DFR_Triticum_monococcum                RFPEY.SIPHKFPGVD.DDLQPIHFSSKKLLDHGF..SFRY.TAEDMFDAAIRTCRE.KG
QFQ61498.1_DFR_Dryopteris_erythrosora             RYPEF.KIPTDLAYL..GEPVIHYFDSSKLKRLGF..QFNF.TLEDMYHDAIKCCKE.KG
QFQ61499.1_DFR_Dryopteris_erythrosora             LFPKI.KLPDNFDYV..GEILLHYMDNSKLKKLGF..EFKH.GLADMFESAVKCCVE.KG
BAA74700.1_DFR_Ipomoea_purpurea                   NWPEY.YIPSQFKGIE.KELAVVYFSSKKLQDMGF..QFKY.SLEDMYRGAIETLRK.KG
BAF93896.1_DFR_Iris_x_hollandica                  RHPWCGSIPEKFDGIE.KDVRTVHFSSKRLLDLGF..EFKY.TVEEMFDEAIRSCVE.KK
XP_008797532.1_DFR_Phoenix_dactylifera            RYPQY.SIPQKFEGID.DGIGRVCFSSKKLIDLGF..KFRY.SMEDMFDAAIQICRE.KK
AEQ92209.1_DFR_Ipomoea_batatas                    NWPEY.YIPSQFKGIE.KDLPVVYFSSKKLQDMGF..QFKY.SLEDMYRGAIETLRK.KG
BAH98155.1_DFR_Tulipa_gesneriana                  RYPQY.AIPQEFEDID.EKIKPVRFSSKKLMDLGF..NYQY.TIEEMFDEGIRSCTE.KK
APG32494.1_DFR2_Freesia_hybrid_cultivar           RHPWC.SIPQKFKGIN.EGIETVHFSSKKLMDLGF..RFKY.SMEEMFDEAIQTCIE.KK
AAO63026.1_DFR_Allium_cepa                        NYPQY.YIPQQFEGID.KGIQPVRFSSKKLVDLGF..RYKY.SMESMFDEAIKTCVE.RK
AFP58815.1_DFR_Hyacinthus_orientalis              KYHQY.HIPQKIEGMD.DEIQTVRFSSKKLLEFGF..EYKY.TMEEMFDEAIKSCVE.KK
                                             acc
```

*NP_001268144.1_DFR_Vitis_vinifera*

| | |
|---|---|
| | 330 |
| NP_001268144.1_DFR_Vitis_vinifera | L..L......PPS.HE........KP.................................. |
| TRINITY_DN27402_c0_g1_i3 | L..L......PFS............................................... |
| NP_199094.1_DFR_Arabidopsis_thaliana | F..L......PVSLSY........QSISEIK...............TKNENIDV...... |
| BAA85261.1_DFR_Arabidopsis_thaliana | F..L......PVSLSY........QSISEIKVP.............TKNEIIEV...... |
| AEI59122.1_DFR_Medicago_sativa | L..L......PKV............................................... |
| BAD67185.1_DFR_Spinacia_oleracea | L..L......PPS............................................... |
| BAB40789.1_DFR_Lilium_hybrid | L..I......PRQ.TQ........ERYYADDKLNLC........................ |
| CAA79154.1_DFR_Solanum_lycopersicum | L..L......PFS.TR........STAD................................ |
| AHZ30596.1_DFR_Prunus_domestica | L..I......PIS.AE........K................................... |
| AAD26204.1_DFR_Malus_domestica | LIPI......PIP.AE........K................................... |
| AAO39819.1_DFR_Pyrus_communis | L..I......PIP.AE........K................................... |
| AAD56578.1_DFR_Daucus_carota | L..L......PNSTTL........QENDQEKK............................ |
| AKN56970.1_DFR_Gerbera_hybrid_cultivar | M..L......PYS.T............................................. |
| AOF39984.11_DFR_Brassica_rapa | F..L......PVTLPE........HLKSEDKVP.............GSDDNKEI...... |
| P51106.1_DFR_Hordeum_vulgare | L..I......PLG.DV........PAPAAGGKLGA......................... |
| BAE19953.1_DFR_Lotus_japonicus | L..L......PKT.A............................................. |
| AII26023.1_DFR_Pisum_sativum | L..L......PKA.VE............................................ |
| NP_001274988.1_DFR_Solanum_tuberosum | L..L......PFS.TR........SSAD................................ |
| NP_001152467.2_DFR_Zea_mays | L..I......PLA..............TAAGGDGFAS....................... |
| AIZ74402.1_DFR_Anthurium_andraeanum | L..L......PPA.TK........EPSYATEQL........................... |
| AAB62873.1_DFR_Bromheadia_finlaysoniana | L..M......PLN.TE........ELVLAAEKYDEV........................ |
| CAA91922.1_DFR_Callistephus_chinensis | F..L......PYS.TN........E................................... |
| AIM58715.1_DFR_Cymbidium_hybrid_cultivar | L..I......PLH.TE........EMVSANEKFDEV........................ |
| ACZ48698.1_DFR_Fagopyrum_esculentum | L..L......PKT.FE........EI.................................. |
| ACZ48697.1_DFR_Fagopyrum_tataricum | L..L......PKT.FE........EI.................................. |
| BAO53730.1_DFR2_Glycine_max | L..L......PKP.EE........TTVN................................ |
| AAP13055.1_DFR_Gypsophila_elegans | L..L......PLS............................................... |
| XP_013466134.1_DFR_Medicago_truncatula | L..L......PKF.V............................................. |
| BAF96936.1_DFR_Nicotiana_tabacum | L..L......PFS.TQ........STAD................................ |
| BAA36183.1_DFR_Oryza_sativa_Japonica_Group | L..L.....PPLP.PP........PTTAVAGGDGSAG....................... |
| BAA12723.1_DFR_Rosa_hybrid_cultivar | L..L......PPP.TE........R................................... |
| AHL46438.1_DFR_Fragaria_vesca | L..L......PLP.QE........E................................... |
| AHL46443.1_DFR_Fragaria_x_ananassa | L..L......PLP.QE........EE.................................. |
| AHL46445.1_DFR_Fragaria_vesca | L..L......PPP.TE........R................................... |
| AHM27144.1_DFR_Angelonia_angustifolia | L..L......PYS.TH........NQSNGEKKESTLQSLEK.....HSDDQEKVL..... |
| AAT84073.1_DFR_Camellia_sinensis | L..L......PHSFAE........NPVNGNKV............................ |
| AAS00611.1_DFR_Citrus_sinensis | L..L......PLL.CE........N................................... |
| AAX16491.1_DFR_Crataegus_monogyna | L..I......PIP.AE........K................................... |
| BAF49325.1_DFR_Delphinium_belladonna | I..L......PFT............................................... |
| CAA78930.1_DFR_Gerbera_hybrid_cultivar | L..L......PYS.T............................................. |
| AAO39816.1_DFR_Malus_domestica | LIPI......PIP.AE........K................................... |
| AAO39817.1_DFR_Malus_domestica | LIPI......PIP.AE........K................................... |
| AAR27015.1_DFR2_Medicago_truncatula | L..L......PKV............................................... |
| AEF14420.1_DFR_Onobrychis_viciifolia | L..L......PKA.VE............................................ |
| AAF60298.1_DFR_Petunia_x_hybrida | L..L......PFS.PR........SAED................................ |
| AAQ77347.1_DFR_Triticum_aestivum | L..I......PLG.DA........PAPAAAGKLGA......................... |
| AAL89715.1_DFR_Vaccinium_macrocarpon | L..L......PYS.NE........TTANGNGNG........................... |
| AAX12420.1_DFR_Vaccinium_macrocarpon | L..L......PYS.NE........TTANGNGNG........................... |
| CAA33543.1_DFR_Antirrhinum_majus | M..L......PYS.TK........NNKGDEKEP.ILNSLEN....NYNIQDKELF..... |
| AAB62873.1_DFR_Bromheadia_finlaysoniana | L..M......PLN.TE........ELVLAAEKYDEV........................ |
| BAA84940.1_DFR_Camellia_sinensis | L..L......PHSFAE........NPVNGNKV............................ |
| AAC17843.1_DFR_Cymbidium_hybrid_cultivar | L..I......PLH.TE........EMVSANEKFDEV........................ |
| CAA91924.1_DFR_Dianthus_caryophyllus | L..L......PLS............................................... |
| CAA70345.1_DFR_Forsythia_x_intermedia | L..L......PYS.TR........NQANREKKE.LLLNLKK....FHGHDQE........ |
| AAC25960.1_DFR_Fragaria_x_ananassa | L..L......PLP.QE........EE.................................. |
| BAA12736.1_DFR_Gentiana_triflora | M..L......PLSIGH........Q................................... |
| AAD54273.1_DFR1_Glycine_max | L..L......PKP.AE............................................ |
| BAA59332.1_DFR_Ipomoea_nil | L..L......PYS.TK........EPADIEQEQHSG........................ |
| BAA36405.1_DFR_Ipomoea_purpurea | L..L......PYS.TK........EPAGIEQ............................. |
| AAD49343.1_DFR_Lilium_hybrid_division_VII | L..I......PHQ.TQ........ERYYVHDELDLG........................ |
| AAF23884.2_DFR3_Lotus_corniculatus | L..L......PKA.A............................................. |
| BAE19948.1_DFR_Lotus_japonicus | L..L......PKA.A............................................. |
| BAE19949.1_DFR_Lotus_japonicus | L..L......PKA.A............................................. |
| BAE19950.1_DFR_Lotus_japonicus | L..L......PKA.A............................................. |
| BAE19951.1_DFR_Lotus_japonicus | ............................................................ |
| BAE19953.1_DFR_Lotus_japonicus | L..L......PKT.A............................................. |
| BAA19658.1_DFR_Perilla_frutescens | M..L......PF.STQ........IHTNGENKESLSNSQEKHSQINTNGENKDSI..... |
| BAB20075.1_DFR_Torenia_hybrid_cultivar | L..L......PYSSTR........DHIHGEHKIE.......................... |
| AAL35830.1_DFR_Triticum_monococcum | L..I......PLG.DA........PPPAAAGKLGA......................... |
| QFQ61498.1_DFR_Dryopteris_erythrosora | L..L........................................................ |
| QFQ61499.1_DFR_Dryopteris_erythrosora | L..LQYPGEAP................................................. |
| BAA74700.1_DFR_Ipomoea_purpurea | L..L......PYS.TK........EAAAIEEEQETVP....................... |
| BAF93896.1_DFR_Iris_x_hollandica | L..I......P...............LPENGNVDAAA....................... |
| XP_008797532.1_DFR_Phoenix_dactylifera | L..I......PHQ.T..........KEPLCSN............................ |
| AEQ92209.1_DFR_Ipomoea_batatas | L..L......PYS.TK........EPAAIEEEQETVP....................... |
| BAH98155.1_DFR_Tulipa_gesneriana | L..L......PIH.TQ........ELFYIDDKIDLGGRKKNLINEMMRGQSEQLSMYTVE |
| APG32494.1_DFR2_Freesia_hybrid_cultivar | L..I......T...............LPQSGD............................ |
| AAO63026.1_DFR_Allium_cepa | F..I......PLQ.TAVELQLKPYELLEHNNK...............NG........... |
| AFP58815.1_DFR_Hyacinthus_orientalis | L..I......PPQ.AV........EALAENGK............................ |
| *acc* | |

*NP_001268144.1_DFR_Vitis_vinifera*

| | |
|---|---|
| **NP_001268144.1_DFR_Vitis_vinifera** | .....................................................VDGKT.. |
| **TRINITY_DN27402_c0_g1_i3** | .....................................................HQPSVSS |
| **NP_199094.1_DFR_Arabidopsis_thaliana** | ...............................KTGD...GLTDGM.......KPCNKTETG |
| **BAA85261.1_DFR_Arabidopsis_thaliana** | ...............................KTGD...GLTDGM.......KPCNKTETG |
| **AEI59122.1_DFR_Medicago_sativa** | ........................................................EETP |
| **BAD67185.1_DFR_Spinacia_oleracea** | .....................................................YGENENN |
| **BAB40789.1_DFR_Lilium_hybrid** | .........CTK............MTNDKLDLGGMK.........LNSLDEIVRGH.NEQ |
| **CAA79154.1_DFR_Solanum_lycopersicum** | ................................NGKD.............KEAIPISTENY |
| **AHZ30596.1_DFR_Prunus_domestica** | .....................................................HE.AD.. |
| **AAD26204.1_DFR_Malus_domestica** | .....................................................TEAAE.. |
| **AAO39819.1_DFR_Pyrus_communis** | .....................................................TEAAE.. |
| **AAD56578.1_DFR_Daucus_carota** | ...................................................YLFSATDDH |
| **AKN56970.1_DFR_Gerbera_hybrid_cultivar** | ..............................IKNHI.....................NGNH |
| **AOF39984.11_DFR_Brassica_rapa** | ...............................KNGSA..GLTDGM.......VACKKTEPG |
| **P51106.1_DFR_Hordeum_vulgare** | ...................................................LAAGEGQA. |
| **BAE19953.1_DFR_Lotus_japonicus** | ........................................................ETPA |
| **AII26023.1_DFR_Pisum_sativum** | .......................................................SVETE |
| **NP_001274988.1_DFR_Solanum_tuberosum** | ................................NGKD.............KEAIPISTENY |
| **NP_001152467.2_DFR_Zea_mays** | ...............................V...................RAPGETEAT |
| **AIZ74402.1_DFR_Anthurium_andraeanum** | .........................I..............................ATGQ |
| **AAB62873.1_DFR_Bromheadia_finlaysoniana** | .........................................................KEQ |
| **CAA91922.1_DFR_Callistephus_chinensis** | ..............................VKKGL.....................FESS |
| **AIM58715.1_DFR_Cymbidium_hybrid_cultivar** | .........................................................KEQ |
| **ACZ48698.1_DFR_Fagopyrum_esculentum** | ........................................................EKNH |
| **ACZ48697.1_DFR_Fagopyrum_tataricum** | ........................................................EKNH |
| **BAO53730.1_DFR2_Glycine_max** | ................................NV.................LLPKPAETT |
| **AAP13055.1_DFR_Gypsophila_elegans** | .....................................................L...ENH |
| **XP_013466134.1_DFR_Medicago_truncatula** | .........................................................KST |
| **BAF96936.1_DFR_Nicotiana_tabacum** | ................................NGRD.............KETIPLSAENY |
| **BAA36183.1_DFR_Oryza_sativa_Japonica_Group** | ...............................VAGEK.........E...PILGRGTGTA. |
| **BAA12723.1_DFR_Rosa_hybrid_cultivar** | .....................................................VEKQEVD |
| **AHL46438.1_DFR_Fragaria_vesca** | ............................................................ |
| **AHL46443.1_DFR_Fragaria_x_ananassa** | .....................................................TEKRRAG |
| **AHL46445.1_DFR_Fragaria_vesca** | .....................................................VEKQENG |
| **AHM27144.1_DFR_Angelonia_angustifolia** | ............................EKRVNGR.....LDKQFEEQKKELLPIREEKH |
| **AAT84073.1_DFR_Camellia_sinensis** | ............................................................ |
| **AAS00611.1_DFR_Citrus_sinensis** | ............................................................ |
| **AAX16491.1_DFR_Crataegus_monogyna** | .....................................................TEAAE.. |
| **BAF49325.1_DFR_Delphinium_belladonna** | ...................................................NKPKSATKW |
| **CAA78930.1_DFR_Gerbera_hybrid_cultivar** | ..............................IKNHI.....................NGNH |
| **AAO39816.1_DFR_Malus_domestica** | .....................................................TEAAE.. |
| **AAO39817.1_DFR_Malus_domestica** | .....................................................TEAAE.. |
| **AAR27015.1_DFR2_Medicago_truncatula** | ........................................................TETP |
| **AEF14420.1_DFR_Onobrychis_viciifolia** | ........................................................ETTP |
| **AAF60298.1_DFR_Petunia_x_hybrida** | ................................NGHN.............REAIAISAQNY |
| **AAQ77347.1_DFR_Triticum_aestivum** | ...................................................LAAGKGQA. |
| **AAL89715.1_DFR_Vaccinium_macrocarpon** | ................................NG.......................... |
| **AAX12420.1_DFR_Vaccinium_macrocarpon** | ................................NG.......................... |
| **CAA33543.1_DFR_Antirrhinum_majus** | ........................PISEEKHINGQE.NALLSNT...QDKELLPTSEEKR |
| **AAB62873.1_DFR_Bromheadia_finlaysoniana** | .........................................................KEQ |
| **BAA84940.1_DFR_Camellia_sinensis** | ............................................................ |
| **AAC17843.1_DFR_Cymbidium_hybrid_cultivar** | .........................................................KEQ |
| **CAA91924.1_DFR_Dianthus_caryophyllus** | .....................................................L...EHH |
| **CAA70345.1_DFR_Forsythia_x_intermedia** | ............................ELPANG......................KEKH |
| **AAC25960.1_DFR_Fragaria_x_ananassa** | .....................................................TEKRRAG |
| **BAA12736.1_DFR_Gentiana_triflora** | .....................................................KESTDPE |
| **AAD54273.1_DFR1_Glycine_max** | ................................KG.................LFTKPGETP |
| **BAA59332.1_DFR_Ipomoea_nil** | ..........................................................KE |
| **BAA36405.1_DFR_Ipomoea_purpurea** | ...........................................................E |
| **AAD49343.1_DFR_Lilium_hybrid_division_VII** | .........CSK............MTNDKLDLGGSK.........LNSMDEMVRGH.NER |
| **AAF23884.2_DFR3_Lotus_corniculatus** | .........................................................ENP |
| **BAE19948.1_DFR_Lotus_japonicus** | .........................................................ETP |
| **BAE19949.1_DFR_Lotus_japonicus** | .........................................................ENP |
| **BAE19950.1_DFR_Lotus_japonicus** | .........................................................ENP |
| **BAE19951.1_DFR_Lotus_japonicus** | ......................................................KWGSNL |
| **BAE19953.1_DFR_Lotus_japonicus** | ........................................................ETPA |
| **BAA19658.1_DFR_Perilla_frutescens** | .........................LNSQINTNGENKDSILNSH.........DKHSQIH |
| **BAB20075.1_DFR_Torenia_hybrid_cultivar** | ............................................................ |
| **AAL35830.1_DFR_Triticum_monococcum** | ...................................................LAAGEGQA. |
| **QFQ61498.1_DFR_Dryopteris_erythrosora** | ............................................................ |
| **QFQ61499.1_DFR_Dryopteris_erythrosora** | ............................................................ |
| **BAA74700.1_DFR_Ipomoea_purpurea** | .........................LKVEEPT............AIEQKQETVPLKLEEE |
| **BAF93896.1_DFR_Iris_x_hollandica** | ........................GAKDMVHGAEEH.........A...RIAMELEPKKK |
| **XP_008797532.1_DFR_Phoenix_dactylifera** | ...............................KAGEH..............VAMDA...KR |
| **AEQ92209.1_DFR_Ipomoea_batatas** | .........................LKVQEPTKQEATTVPLKPAAIEQKQETVPLKL.EE |
| **BAH98155.1_DFR_Tulipa_gesneriana** | KMFGEQIRSCTESKLLPLQTQELFYINDKIDLGGSK.........MNSIKEMMRGQ.TEQ |
| **APG32494.1_DFR2_Freesia_hybrid_cultivar** | ..........................................................RK |
| **AAO63026.1_DFR_Allium_cepa** | ........................VVTNTIKIVGQM.........VNTKAMITEHEENEP |
| **AFP58815.1_DFR_Hyacinthus_orientalis** | .............................LEVTGEQ.........VDVAKAAMNGE.KEE |

*acc*

*NP_001268144.1_DFR_Vitis_vinifera*

```
NP_001268144.1_DFR_Vitis_vinifera           ............................................................
TRINITY_DN27402_c0_g1_i3                    QNS.........................................................
NP_199094.1_DFR_Arabidopsis_thaliana        ITGERTDAP............................MLAQQMCA...............
BAA85261.1_DFR_Arabidopsis_thaliana         VTGERTDAP............................MLAQQMCA...............
AEI59122.1_DFR_Medicago_sativa              VNDTMKK.....................................................
BAD67185.1_DFR_Spinacia_oleracea            ENPNTIID....................................................
BAB40789.1_DFR_Lilium_hybrid                VSVALQ......................................................
CAA79154.1_DFR_Solanum_lycopersicum         SSGKENAPVANCT.........GKFTNGEI..............................
AHZ30596.1_DFR_Prunus_domestica             DNTVVDVK.................VSG................................
AAD26204.1_DFR_Malus_domestica              ESNLVDVK.................VG.................................
AAO39819.1_DFR_Pyrus_communis               ESNLVDVK.................VGG................................
AAD56578.1_DFR_Daucus_carota                HNGHEKDLFHHSIDKDAI....GKEKRGETES.....LVAA...................
AKN56970.1_DFR_Gerbera_hybrid_cultivar      VNGVHHYIKNNG..........DDQEKGLL.......CCSKEGQ................
AOF39984.11_DFR_Brassica_rapa               MAGEKADSH............................MSAQQICA...............
P51106.1_DFR_Hordeum_vulgare                IGAET.......................................................
BAE19953.1_DFR_Lotus_japonicus              TNGTTQK.....................................................
AII26023.1_DFR_Pisum_sativum                VNDTMKK.....................................................
NP_001274988.1_DFR_Solanum_tuberosum        SSGKENAPVANCT.........GKFTNGEI..............................
NP_001152467.2_DFR_Zea_mays                 IGA.........................................................
AIZ74402.1_DFR_Anthurium_andraeanum         DNGH........................................................
AAB62873.1_DFR_Bromheadia_finlaysoniana     IAVK........................................................
CAA91922.1_DFR_Callistephus_chinensis       INGNVHGQKGNQ..........KIGDEGVK.......LVN....................
AIM58715.1_DFR_Cymbidium_hybrid_cultivar    IAVK........................................................
ACZ48698.1_DFR_Fagopyrum_esculentum         VNGNGH......................................................
ACZ48697.1_DFR_Fagopyrum_tataricum          VNGNGH......................................................
BAO53730.1_DFR2_Glycine_max                 VNDTMRK.....................................................
AAP13055.1_DFR_Gypsophila_elegans           ENVYA.......................................................
XP_013466134.1_DFR_Medicago_truncatula      NK..........................................................
BAF96936.1_DFR_Nicotiana_tabacum            ASGKENSPVANGT.........GKSTNGEI..............................
BAA36183.1_DFR_Oryza_sativa_Japonica_Group  VGAETEALVK..................................................
BAA12723.1_DFR_Rosa_hybrid_cultivar         ESSVVRVK.................VTG................................
AHL46438.1_DFR_Fragaria_vesca               ............................................................
AHL46443.1_DFR_Fragaria_x_ananassa          ............................................................
AHL46445.1_DFR_Fragaria_vesca               DSSVVHVE.................VTG................................
AHM27144.1_DFR_Angelonia_angustifolia       TNGSLEKCSDDQA.KVLLPLPEEKQTNGSLEK...HVMINIKTCE...............
AAT84073.1_DFR_Camellia_sinensis            ............................................................
AAS00611.1_DFR_Citrus_sinensis              ..HVSEVS.................I..................................
AAX16491.1_DFR_Crataegus_monogyna           DSNLVDVK.................VGG................................
BAF49325.1_DFR_Delphinium_belladonna        SN..........................................................
CAA78930.1_DFR_Gerbera_hybrid_cultivar      VNGVHHYIKNND..........DDHEKGLL.......CCSKEGQ................
AAO39816.1_DFR_Malus_domestica              ESNLVDVK.................AG.................................
AAO39817.1_DFR_Malus_domestica              ESNLVDVK.................VG.................................
AAR27015.1_DFR2_Medicago_truncatula         VNDTMKK.....................................................
AEF14420.1_DFR_Onobrychis_viciifolia        VNGTTHK.....................................................
AAF60298.1_DFR_Petunia_x_hybrida            ASGKENAPVANHT.........EMLSNVEV..............................
AAQ77347.1_DFR_Triticum_aestivum            IGAET.......................................................
AAL89715.1_DFR_Vaccinium_macrocarpon        .NGN.....................GNGTI..............................
AAX12420.1_DFR_Vaccinium_macrocarpon        .NGN.....................GNGTI..............................
CAA33543.1_DFR_Antirrhinum_majus            VNGLESALLSKIQDKEVLPTSGVKHAKGQENAL.LPDIANDHTDGRI.............
AAB62873.1_DFR_Bromheadia_finlaysoniana     IAVK........................................................
BAA84940.1_DFR_Camellia_sinensis            ............................................................
AAC17843.1_DFR_Cymbidium_hybrid_cultivar    IAVK........................................................
CAA91924.1_DFR_Dianthus_caryophyllus        LCVFRVTLIFFK................................................
CAA70345.1_DFR_Forsythia_x_intermedia       TNG.........................................................
AAC25960.1_DFR_Fragaria_x_ananassa          ............................................................
BAA12736.1_DFR_Gentiana_triflora            VDEVVKEME............................LIQDSLD................
AAD54273.1_DFR1_Glycine_max                 VNAMHK......................................................
BAA59332.1_DFR_Ipomoea_nil                  PKS.........................................................
BAA36405.1_DFR_Ipomoea_purpurea             PKS.........................................................
AAD49343.1_DFR_Lilium_hybrid_division_VII   VSVALQ......................................................
AAF23884.2_DFR3_Lotus_corniculatus          SNGK........................................................
BAE19948.1_DFR_Lotus_japonicus              SNGIMEK.....................................................
BAE19949.1_DFR_Lotus_japonicus              SNGK........................................................
BAE19950.1_DFR_Lotus_japonicus              SNGITEK.....................................................
BAE19951.1_DFR_Lotus_japonicus              NTA.........................................................
BAE19953.1_DFR_Lotus_japonicus              TNGTTQK.....................................................
BAA19658.1_DFR_Perilla_frutescens           TNGENKESILNSQEK.....HSQIRTNGENKESIFNSLEKHDTDNNQEKELLPIPKEAHA
BAB20075.1_DFR_Torenia_hybrid_cultivar      ............................................................
AAL35830.1_DFR_Triticum_monococcum          IGAET.......................................................
QFQ61498.1_DFR_Dryopteris_erythrosora       ............................................................
QFQ61499.1_DFR_Dryopteris_erythrosora       ............................................................
BAA74700.1_DFR_Ipomoea_purpurea             PIAIEKKQE.......VVP..................LKA....................
BAF93896.1_DFR_Iris_x_hollandica            VK..........................................................
XP_008797532.1_DFR_Phoenix_dactylifera      VN.GLTIQKS..................................................
AEQ92209.1_DFR_Ipomoea_batatas              PTAIEQKQK.......VVP..................LKA....................
BAH98155.1_DFR_Tulipa_gesneriana            LSTAFH......................................................
APG32494.1_DFR2_Freesia_hybrid_cultivar     VENG........................................................
AAO63026.1_DFR_Allium_cepa                  IATH........................................................
AFP58815.1_DFR_Hyacinthus_orientalis        VH....IASR..................................................
```

***acc***

*NP_001268144.1_DFR_Vitis_vinifera*

| | |
|---|---|
| NP_001268144.1_DFR_Vitis_vinifera | . . . . . . . |
| TRINITY_DN27402_c0_g1_i3 | . . . . . . . |
| NP_199094.1_DFR_Arabidopsis_thaliana | . . . . . . . |
| BAA85261.1_DFR_Arabidopsis_thaliana | . . . . . . . |
| AEI59122.1_DFR_Medicago_sativa | . . . . . . . |
| BAD67185.1_DFR_Spinacia_oleracea | . . . . . . . |
| BAB40789.1_DFR_Lilium_hybrid | . . . . . . . |
| CAA79154.1_DFR_Solanum_lycopersicum | . . . . . . . |
| AHZ30596.1_DFR_Prunus_domestica | . . . . . . . |
| AAD26204.1_DFR_Malus_domestica | . . . . . . . |
| AAO39819.1_DFR_Pyrus_communis | . . . . . . . |
| AAD56578.1_DFR_Daucus_carota | . . . . . . . |
| AKN56970.1_DFR_Gerbera_hybrid_cultivar | . . . . . . . |
| AOF39984.11_DFR_Brassica_rapa | . . . . . . . |
| P51106.1_DFR_Hordeum_vulgare | . . . . . . . |
| BAE19953.1_DFR_Lotus_japonicus | . . . . . . . |
| AII26023.1_DFR_Pisum_sativum | . . . . . . . |
| NP_001274988.1_DFR_Solanum_tuberosum | . . . . . . . |
| NP_001152467.2_DFR_Zea_mays | . . . . . . . |
| AIZ74402.1_DFR_Anthurium_andraeanum | . . . . . . . |
| AAB62873.1_DFR_Bromheadia_finlaysoniana | . . . . . . . |
| CAA91922.1_DFR_Callistephus_chinensis | . . . . . . . |
| AIM58715.1_DFR_Cymbidium_hybrid_cultivar | . . . . . . . |
| ACZ48698.1_DFR_Fagopyrum_esculentum | . . . . . . . |
| ACZ48697.1_DFR_Fagopyrum_tataricum | . . . . . . . |
| BAO53730.1_DFR2_Glycine_max | . . . . . . . |
| AAP13055.1_DFR_Gypsophila_elegans | . . . . . . . |
| XP_013466134.1_DFR_Medicago_truncatula | . . . . . . . |
| BAF96936.1_DFR_Nicotiana_tabacum | . . . . . . . |
| BAA36183.1_DFR_Oryza_sativa_Japonica_Group | . . . . . . . |
| BAA12723.1_DFR_Rosa_hybrid_cultivar | . . . . . . . |
| AHL46438.1_DFR_Fragaria_vesca | . . . . . . . |
| AHL46443.1_DFR_Fragaria_x_ananassa | . . . . . . . |
| AHL46445.1_DFR_Fragaria_vesca | . . . . . . . |
| AHM27144.1_DFR_Angelonia_angustifolia | . . . . . . . |
| AAT84073.1_DFR_Camellia_sinensis | . . . . . . . |
| AAS00611.1_DFR_Citrus_sinensis | . . . . . . . |
| AAX16491.1_DFR_Crataegus_monogyna | . . . . . . . |
| BAF49325.1_DFR_Delphinium_belladonna | . . . . . . . |
| CAA78930.1_DFR_Gerbera_hybrid_cultivar | . . . . . . . |
| AAO39816.1_DFR_Malus_domestica | . . . . . . . |
| AAO39817.1_DFR_Malus_domestica | . . . . . . . |
| AAR27015.1_DFR2_Medicago_truncatula | . . . . . . . |
| AEF14420.1_DFR_Onobrychis_viciifolia | . . . . . . . |
| AAF60298.1_DFR_Petunia_x_hybrida | . . . . . . . |
| AAQ77347.1_DFR_Triticum_aestivum | . . . . . . . |
| AAL89715.1_DFR_Vaccinium_macrocarpon | . . . . . . . |
| AAX12420.1_DFR_Vaccinium_macrocarpon | . . . . . . . |
| CAA33543.1_DFR_Antirrhinum_majus | . . . . . . . |
| AAB62873.1_DFR_Bromheadia_finlaysoniana | . . . . . . . |
| BAA84940.1_DFR_Camellia_sinensis | . . . . . . . |
| AAC17843.1_DFR_Cymbidium_hybrid_cultivar | . . . . . . . |
| CAA91924.1_DFR_Dianthus_caryophyllus | . . . . . . . |
| CAA70345.1_DFR_Forsythia_x_intermedia | . . . . . . . |
| AAC25960.1_DFR_Fragaria_x_ananassa | . . . . . . . |
| BAA12736.1_DFR_Gentiana_triflora | . . . . . . . |
| AAD54273.1_DFR1_Glycine_max | . . . . . . . |
| BAA59332.1_DFR_Ipomoea_nil | . . . . . . . |
| BAA36405.1_DFR_Ipomoea_purpurea | . . . . . . . |
| AAD49343.1_DFR_Lilium_hybrid_division_VII | . . . . . . . |
| AAF23884.2_DFR3_Lotus_corniculatus | . . . . . . . |
| BAE19948.1_DFR_Lotus_japonicus | . . . . . . . |
| BAE19949.1_DFR_Lotus_japonicus | . . . . . . . |
| BAE19950.1_DFR_Lotus_japonicus | . . . . . . . |
| BAE19951.1_DFR_Lotus_japonicus | . . . . . . . |
| BAE19953.1_DFR_Lotus_japonicus | . . . . . . . |
| BAA19658.1_DFR_Perilla_frutescens | D R Q E M Q I |
| BAB20075.1_DFR_Torenia_hybrid_cultivar | . . . . . . . |
| AAL35830.1_DFR_Triticum_monococcum | . . . . . . . |
| QFQ61498.1_DFR_Dryopteris_erythrosora | . . . . . . . |
| QFQ61499.1_DFR_Dryopteris_erythrosora | . . . . . . . |
| BAA74700.1_DFR_Ipomoea_purpurea | . . . . . . . |
| BAF93896.1_DFR_Iris_x_hollandica | . . . . . . . |
| XP_008797532.1_DFR_Phoenix_dactylifera | . . . . . . . |
| AEQ92209.1_DFR_Ipomoea_batatas | . . . . . . . |
| BAH98155.1_DFR_Tulipa_gesneriana | . . . . . . . |
| APG32494.1_DFR2_Freesia_hybrid_cultivar | . . . . . . . |
| AAO63026.1_DFR_Allium_cepa | . . . . . . . |
| AFP58815.1_DFR_Hyacinthus_orientalis | . . . . . . . |
| *acc* | |

```
sp|Q4W2K4_LAR_Vitis_vinifera
                                                   1
sp|Q4W2K4_LAR_Vitis_vinifera                  .....MTVSP...........VPS.......................P
TRINITY_DN33042_c3_g1_i3                      .....MTASATL.........FST.......................M
NP_001352050.1_LAR_Glycine_max                .....MVTSPPA.........IPTTT......................
ADD51357.1_LAR_Theobroma_cacao                MKSTNMNGSSPN.........VSE.......................E
CAI56321.1_LAR_Pinus_taeda                    .....MACATDVARQFLPCVQPVPSSMGGETARSINLTCNGLSPPQPQYNAENNHDQDTT
CAI56322.1_LAR_Phaseolus_coccineus            .....MVTS.PT.........IPSHT......................
CAI56320.1_LAR_Hordeum_vulgare_subsp._vulgare .......MAPC..EELQ...EEVA.......................R
CAD79341.1_LAR_Desmodium_uncinatum            ......MTVSGA.........IPSMT......................
AIS92512_LAR_Epimedium_sagittatum             .......MAPPS.........VDSVATF.................SFETCSK
AII26024.1_LAR_Pisum_sativum                  .......MAPTS.........SPPTTL....................AS
AEF14422.1_LAR_Onobrychis_viciifolia          .....MATSPAN.........IPPTL.......................
ABE90657.1_LAR_Medicago_truncatula            .......MAPSS.........S.PTTP....................IS
CAI56326.1_LAR_Vitis_shuttleworthii           .....MTVSP...........VPS.......................L
ABC71327.1_LAR_Lotus_corniculatus             ......MVSTAA.........TPPAT......................
AHA14498.1_LAR_Fagopyrum_tataricum            .....MTVAVTA.........IPE.......................S
AAZ82410.1_LAR_Vitis_vinifera                 .....MTVSP...........VPS.......................P
ADY15310.1_LAR_Prunus_avium                   .....MTVSTCV.........SAA.......................K
BAH89267.1_LAR_Diospyros_kaki                 .....MTVSPSF........AAAA.......................K
AAX12186.1_LAR_Malus_domestica                .....MTVSPSL.........SVA.......................R
AEY62396.1_LAR_Fagopyrum_dibotrys             .....MTVAVTA.........IPE.......................S
AAZ79364.1_LAR_Malus_domestica                .....MTVSSSL.........SVA.......................K
CAI56319.1_LAR_Gossypium_arboreum             .....MTV.............SVA........................
CAI56323.1_LAR_Gossypium_arboreum             MKSTQMNGSYPN.........ES........................E
CAI56324.1_LAR_Gossypium_raimondii            .....MTV.............SVA........................
CAI56325.1_LAR_Gossypium_raimondii            MKSTHMNGSYPN.........ES........................E
CAI56328.1_LAR_Oryza_sativa_Japonica_Group    .......MAPAAQELLL...QEVPQ.....................PRR
CAI26308.1_LAR_Vitis_vinifera                 ..MTVLSVSTPP.........APQAPP....................AA
ABF95070.1_LAR_Oryza_sativa_Japonica_Group    .......MAPAAQELLL...QEVPQ.....................PRR
ADD51358.1_LAR_Theobroma_cacao                MKSTNMNGSSPN.........VSE.......................E
ACI41981.1_LAR_Diospyros_kaki                 .....MTVSPSF........AAAA.......................K
ABC71329.1_LAR_Lotus_corniculatus             .......................M.....................AT
ABH07785.2_LAR_Fragaria_x_ananassa            .....MTVSPSI........ASAA.......................K
ABB77697.1_LAR_Pyrus_communis                 .....MTVSPSL.........SVA.......................I
                                          acc
```

```
                                                   β1          α1                β2           α2        β3
sp|Q4W2K4_LAR_Vitis_vinifera                  10        20        30    ★    40        50        60
sp|Q4W2K4_LAR_Vitis_vinifera                  KG.RVLIAGATGFIGQFVATASLDAHRPTYILARP.GPRS.P.SKAKIFKALEDKGAIIV
TRINITY_DN33042_c3_g1_i3                      GG.SVLIAGATGFIGQYVVQASLDSGRRTYVLVRP.SATACP.SRAKFIKCLEEKGAIIL
NP_001352050.1_LAR_Glycine_max                KD.RVLIIGATGFIGKFVAEASLTSEHPTCLLVRP.GPLV.P.SKDAIVKTFQDKGAIVI
ADD51357.1_LAR_Theobroma_cacao                TG.RTLVVGSGGFMGRFVTEASLDSGRPTYILARS.S.SNSP.SKASTIKFLQDRGATVI
CAI56321.1_LAR_Pinus_taeda                    VATRVLIIGATGFIGRFVAEASVKSGRPTYALVRP.TTLS...SKPKVIQSLVDSGIQVV
CAI56322.1_LAR_Phaseolus_coccineus            KA.RVLIIGATGFIGKFVTEASLLTAHPTYLLLRP.PPLV.P.SKDAIVKTFQEKGAMII
CAI56320.1_LAR_Hordeum_vulgare_subsp._vulgare SG.PALIVGATGYIGRFVAEACLDSGRRTFILVRP.GNAC.P.ARAASVDALLRKGAFVV
CAD79341.1_LAR_Desmodium_uncinatum            KN.RTLVVGGTGFIGQFITKASLGFGYPTFLLVRP.GPVS.P.SKAVIIKTFQDKGAKVI
AIS92512_LAR_Epimedium_sagittatum             AG.RTLIIGATGFIGGFIVDACLASGRPTYILSRS......KSTKVGAHKELQDKGAIVL
AII26024.1_LAR_Pisum_sativum                  KN.RVLIIGATGFMGKFLTEASLSSSHPTYLLIRPGGPLL.S.PKSTTIKTFQDKGAIIV
AEF14422.1_LAR_Onobrychis_viciifolia          KG.RVLIVGATGFMGKFVAEASLSSAHATFLLLRP.GPII.S.SKASIVKAFQDKGARVI
ABE90657.1_LAR_Medicago_truncatula            KG.RVLIVGATGFMGKFVTEASISTAHPTYLLIRP.GPLI.S.SKAATIKTFQEKGAIVI
CAI56326.1_LAR_Vitis_shuttleworthii           KG.RVLIAGATGFIGQFVAAASLDAHRPTYILARP.GPRS.P.SKAKIIKAHEDKGAIIV
ABC71327.1_LAR_Lotus_corniculatus             AG.RILIIGATGFMGQFMTKASLGLGRSTYLLLRP.GSLT.P.SKAAIVKSFQDRGAKVI
AHA14498.1_LAR_Fagopyrum_tataricum            KC.RTLVAGATGFIGRFVTESSLESERPTFILVRP.GPIS.P.SKTKIIKALEDKGAIIV
AAZ82410.1_LAR_Vitis_vinifera                 KG.RVLIAGATGFIGQFVAAASLDAHRPTYILARP.GPRS.P.SKANIFKALEDKGAIIV
ADY15310.1_LAR_Prunus_avium                   NG.RILIVGATGFIGRFVAEASLDAGQPTYVLVRP.GPLD.P.SKADIIKALKDRGAIIL
BAH89267.1_LAR_Diospyros_kaki                 QG.RVLIAGATGFIGQFVAEASLEAGRTTYVLVRS.G....P.SKAKTIKALQEKGAIPI
AAX12186.1_LAR_Malus_domestica                NG.RVLIVGATGFIGRFVAEASLAAGRPTYVLVRP.GPLH.P.SKADTVKSFKHKGAIIL
AEY62396.1_LAR_Fagopyrum_dibotrys             KC.RTLVAGATGFIGRFVTESSLESERPTFILVRP.GPIS.P.SKTKIIKALEDKGAIIV
AAZ79364.1_LAR_Malus_domestica                NG.RVLIAGATGFIGRFVAEASLAAGQPTYVLVRP.GPLH.P.SKADTVKSFKDKGAIIL
CAI56319.1_LAR_Gossypium_arboreum             NG.RVLIVGATGFIGRFVADASLDAGRPTYVLVRP.SSGN.QYSKDKVAKALQDRGAILL
CAI56323.1_LAR_Gossypium_arboreum             AG.QTVVIGSSGFIGRFITEACLDSGRPTYILVRS.S.SNSP.SKASTIKFLQDKGAIVI
CAI56324.1_LAR_Gossypium_raimondii            NG.RVLIVGATGFIGRFVADASLDAGRPTYVLVRP.SSGN.QYSKDKVAKALRDRGAILL
CAI56325.1_LAR_Gossypium_raimondii            TG.QTLVIGSSGFIGRFITEACLDSGRPTYILVRS.S.SNSP.SKASTIKFLQDKGAIVI
CAI56328.1_LAR_Oryza_sativa_Japonica_Group    TG.AALIVGATGYIGRFVAEACLDSGRDTFILVRP.GNAC.P.ARAASVDALRQKGAVVI
CAI26308.1_LAR_Vitis_vinifera                 TGPRTLEVGASGFIGRFVAEASLSSGHPTYVLVRS.SATTSS.SKASTIKSLEDQGAILV
ABF95070.1_LAR_Oryza_sativa_Japonica_Group    TG.AALIVGATGYIGRFVAEACLDSGRDTFILVRP.GNAC.P.ARAASVDALRQKGAVVI
ADD51358.1_LAR_Theobroma_cacao                TG.RTLVVGSGGFMGRFVTEASLDSGRPTYILARS.S.SNSP.SKASTIKFLQDRGATVI
ACI41981.1_LAR_Diospyros_kaki                 QG.RVLIVGATGFIGQFVAEASLEAGRTTYVLVRS.G....P.SKAKTIKALQEKGAIPI
ABC71329.1_LAR_Lotus_corniculatus             KG.RVLIIGATGFMGRFMAEASLAAAHPTYLLVRL..PLI.P.SKATIVKTFQDKGAIVI
ABH07785.2_LAR_Fragaria_x_ananassa            SG.RVLIIGATGFIGKFVAEASLDSGLPTYVLVRP.GPSR.P.SKSDTIKSLKDRGAIIL
ABB77697.1_LAR_Pyrus_communis                 NG.RVLIVGATGFIGRFVAEASLAAGQPTYVLVRP.GPLH.P.SKADTVKSFKHKGAIIL
                                          acc
```

```
sp|Q4W2K4_LAR_Vitis_vinifera                      α3                 β4      η1          α4              β5
                                                    70          80          90         100         110         120
sp|Q4W2K4_LAR_Vitis_vinifera                  Y.GLINEQE...AMEKILKEHEIDIVVSTV.GGESILDQIALVKAMKAVGTIKRFLPSEF
TRINITY_DN33042_c3_g1_i3                      L.GLINDKE...NMEKLLKEHKIDIVISAV.GGENVLDQLVLVEAINAVGTVKRYVPSEF
NP_001352050.1_LAR_Glycine_max                H.GVINNKD...FVEKILKEYEIDIVISAI.GAKSLLDQLILVEAMKSVKTIKRFLPSEF
ADD51357.1_LAR_Theobroma_cacao                Y.GSITDKE...FMEKVLKEHKIEVVISAV.GGGSILDQFNLIEAIRNVDTVKRFLPSEF
CAI56321.1_LAR_Pinus_taeda                    Y.GCLHDHN...SLVKAIR..QVDVVISTV.GGALILDQLKIVDAIKEVGTVKRFLPSEF
CAI56322.1_LAR_Phaseolus_coccineus            H.GVINNKD...FVEKILKEHEIDIVISAI.GAKSLLDQLILVEAMKSLKTIKRFLASEF
CAI56320.1_LAR_Hordeum_vulgare_subsp._vulgare E.GRVDGKDGKRSVETALRAHGIEVVISVM.GGANILDQLGLIKAIQAAGTVKRFLPSEF
CAD79341.1_LAR_Desmodium_uncinatum            Y.GVINDKE...CMEKILKEYEIDVVISLV.GGARLLDQLTLLEAIKSVKTIKRFLPSEF
AIS92512_LAR_Epimedium_sagittatum             Y.GKINDRH...TMENILKDHKIEMVISAV.GGDTLLEQIDLVHAIKAVGSVQRFLPSEF
AII26024.1_LAR_Pisum_sativum                  Y.GVVDNKE...FMEKILKKYEIDIVISAI.GAESLLDQITLVEAMKSIKTVKRFLPSEF
AEF14422.1_LAR_Onobrychis_viciifolia          Y.GVVNNKE...LMEKILKEYEIDIVISAI.GAESLMDQLTLVEAMKSVKSIKRFLPSEF
ABE90657.1_LAR_Medicago_truncatula            Y.GVVNNKE...FVEMILKKYEIDTVISAI.GAESLLDQLTLVEAMKSIKTIKRFLPSEF
CAI56326.1_LAR_Vitis_shuttleworthii           Y.GLINEQE...SMEKILKEHEIDIVVSTV.GGESILDQIALVKAMKAVGTIKRFLPSEF
ABC71327.1_LAR_Lotus_corniculatus             H.GVINDKE...LMVKILKDYEIDVVISLV.GGGNLMDQRTLVDAIKSVKTVKRFLPSEF
AHA14498.1_LAR_Fagopyrum_tataricum            Q.GLINNKE...RMEKILRENEINVVISAV.GGGNILDQIPLVHAIKSVPSVKRFLASEF
AAZ82410.1_LAR_Vitis_vinifera                 Y.GLINEQE...AMEKILKEHEIDIVVSTV.GGESILDQIALVKAMKAVGTIKRFLPSEF
ADY15310.1_LAR_Prunus_avium                   Q.GVISDKA...LMEKLLKEHEIEVVISAV.GGATILDQITLVEAIQAVGTIKRFLPSEF
BAH89267.1_LAR_Diospyros_kaki                 PQGNINDQE...FLEKILKEYEIDVVISAV.GGDSIRDQLTLVHAIKAVGTIKRFLPSEF
AAX12186.1_LAR_Malus_domestica                H.GLISDKT...LMEKILREHEIETVISAV.GGATILDQIALVEAIAAVGTVKRFLPSEF
AEY62396.1_LAR_Fagopyrum_dibotrys             Q.GLINNKE...RMEKILRENEINVVISAV.GGGNILDQIPLVHAIKSVPSVKRFLASEF
AAZ79364.1_LAR_Malus_domestica                H.GLISDKT...LMEKMLREHEIEIVISAV.GGATILDQITLVDAIAAVGTVKRFLPSEF
CAI56319.1_LAR_Gossypium_arboreum             N.GLANDKE...LMVKLLKEHQIEIVISAL.GGATILDQLSLVEAIHSVGTVKRFLPSEF
CAI56323.1_LAR_Gossypium_arboreum             Y.GSITDQE...FMEKVLREYKIEVVISAV.GGESILDQLSLIEAIKNVNTVKRFVPSEF
CAI56324.1_LAR_Gossypium_raimondii            N.GLANDKE...LMVKLLKEHQIEIVISAL.GGATILDQLSLVEAIHSAGTVKRFLPSEF
CAI56325.1_LAR_Gossypium_raimondii            Y.GSITDQE...FMEKVLREYKIEVVISAV.GGESILDQFSLIEAIKNVNTVKRFVPSEF
CAI56328.1_LAR_Oryza_sativa_Japonica_Group    E.GCVGGKEGRKSVEAALRARGVEVVISVM.GGASILDQLGLIEAIRAAGTVKRFLPSEF
CAI26308.1_LAR_Vitis_vinifera                 T.GSIGDKE...VMIEILRKYEIEVVISAV.GGATILDQLTLAEAIKAVGSIKRFLPSEF
ABF95070.1_LAR_Oryza_sativa_Japonica_Group    E.GCVGGKEGRKSVEAALRARGVEVVISVM.GGASILDQLGLIEAIRAAGTVKRFLPSEF
ADD51358.1_LAR_Theobroma_cacao                Y.GSITDKE...FMEKVLKEHKIEVVISAV.GGGSILDQFNLIEAIRNVDTVKRFLPSEF
ACI41981.1_LAR_Diospyros_kaki                 P.GNINDQE...FLEKILKEYEIDVVISAV.GGDSIRDQLTLVHAIKAVGTXKRFLPSEF
ABC71329.1_LAR_Lotus_corniculatus             Q.GVMNDKE...FMQKILKEYQIDVVISTVGGAHGLLDQLTLVEAMKSVNTIKRFLPSEF
ABH07785.2_LAR_Fragaria_x_ananassa            H.GVMSDKP...LMEKLLKEHEIEIVISAV.GGATILDQITLVEAITSVGTVKRFLPSEF
ABB77697.1_LAR_Pyrus_communis                 H.GLISDKT...LMEKILREHEIETVISAV.GGATILDQIALVEAIAAVGTVKRFLPSEF
                                          acc
```

```
sp|Q4W2K4_LAR_Vitis_vinifera                               α5                       β6   β7
                                                  130         140         150         160         170         180
sp|Q4W2K4_LAR_Vitis_vinifera                  GHDVNRADPVEPGLNMYREKRRVRQLVEESGIPFTYICCNSIASWPYYNNIHPSEVLPPT
TRINITY_DN33042_c3_g1_i3                      GHDVDRADPVEPGDKMYMEKRKVRRLIEEYGIPYTYICCNSIASWPYYTNKHPAEVLPPL
NP_001352050.1_LAR_Glycine_max                GHDVDKADPVEPGLTMYKEKRLVRRVVEESGVPFTNICCNSIASWPYHDNCHPSQLPPL
ADD51357.1_LAR_Theobroma_cacao                GHDTDRADPVEPGLTMYEQKRQIRRQIEKSGIPYTYICCNSIAAWPYHDNTHPADVLPPL
CAI56321.1_LAR_Pinus_taeda                    GHDVDRADPVEPALSFYIEKRKVRAVEEAKIPYTYICCNSIAGWPYYYHTHPTELPPK
CAI56322.1_LAR_Phaseolus_coccineus            GHDVDRADPVEPGLTMYKEKQLVRRVVEQSGVPYTNICCNSIASWPYYDNCHPSQLPPL
CAI56320.1_LAR_Hordeum_vulgare_subsp._vulgare GHDVDRARPVGAGLGFYEEKRRVRRAAEAAGVPYTYICCNSIAGWPYFDNMHPSEVRPPL
CAD79341.1_LAR_Desmodium_uncinatum            GHDVDRTDPVEPGLTMYKEKRLVRRAVEEYGIPFTNICCNSIASWPYYDNCHPSQVPPM
AIS92512_LAR_Epimedium_sagittatum             GHDVDRADPVEPGLTMYNEKRKVRRVIEEVGIPYTYICCNSIAAWPYYDNTHPSEVLPPM
AII26024.1_LAR_Pisum_sativum                  GHDVDRADPVEPGLAMYKQKRLVRRVIEESGVPYTYICCNSIASWPYYNNCHPSSLPPL
AEF14422.1_LAR_Onobrychis_viciifolia          GHDVDRADPVEPGLAMYKEKRLVRRVIEQSGVPYTYICCNSIASWPYYDNCHPSQLPPL
ABE90657.1_LAR_Medicago_truncatula            GHDVDRADPVEPGLAMYKQKRLVRRVIEESGVPYTYICCNSIASWPYYDNCHPSQLPPL
CAI56326.1_LAR_Vitis_shuttleworthii           GHDVNRADPVEPGLNMYREKRRVRQLVEESGIPFTYICCNSIASWPYYNNIHPSEVLPPT
ABC71327.1_LAR_Lotus_corniculatus             GHDTDRANPVEPGLTMYKEKRLVRRLIEESGIPYTYICCNSIASWPYHDNCHPSKVPPV
AHA14498.1_LAR_Fagopyrum_tataricum            GHDVDRANPVEPGLTMYLEKRAVRRAIEEAGVPYTHICCNSIASWPYYDNTHPSEVTPPM
AAZ82410.1_LAR_Vitis_vinifera                 GHDVNRADPVEPGLNMYREKRRVRQLVEESGIPFTYICCNSIASWPYYNNIHPSEVLPPT
ADY15310.1_LAR_Prunus_avium                   GHDVDRADPVEPGLTMYLEKRQVRRWVEKSGVPYTYICCNSIASWPYFDNKHPSEVLPPL
BAH89267.1_LAR_Diospyros_kaki                 GHDVDRANPVEPGLAMYIEKRTVRRVVEESGVPYTYICCNSIASWPYYDNTHPSEVLPPL
AAX12186.1_LAR_Malus_domestica                GHDVDRADPVEPGLTMYEEKRKVRRWVEKSGVAYTYICCNSIASWPYFDNTHPSEVLPPL
AEY62396.1_LAR_Fagopyrum_dibotrys             GHDVDRANPVEPGLTMYLEKRAVRRAIEEAGVPYTHICCNSIASWPYYDNTHPSEVTPPM
AAZ79364.1_LAR_Malus_domestica                GHDVDRADPVEPGLTMYLEKRKVRRSVETSGVPYTYICCNSIASWPYFDNTHPSEVLPPL
CAI56319.1_LAR_Gossypium_arboreum             GHDVDRADPVEPGLTMYKEKRQVRRLIEKLEIPYTYICCNSIASWPYHNNTHPSEVIPPL
CAI56323.1_LAR_Gossypium_arboreum             GHDIDRAEPVEPGLTMYEQKRKIRRQIEECGIPYSYICCNSIAAWPYHDNTHPTDVLPPL
CAI56324.1_LAR_Gossypium_raimondii            GHDVDRADPVEPGLTMYKEKRQVRRLIEKLEIPYTYICCNSIASWPYHNNRHPSEVIPPL
CAI56325.1_LAR_Gossypium_raimondii            GHDIDRAEPVEPGLTMYEQKSKIRRQIEECGIPYSYICCNSIAAWPYHDNTHPADVLPPL
CAI56328.1_LAR_Oryza_sativa_Japonica_Group    GHDVDRARPVGAGLRFYEEKRLVRRAAEASGVPYTFICCNSIAGWPYHDSTHPSELPPL
CAI26308.1_LAR_Vitis_vinifera                 GHDIDRAEPVEPGLTMYMEKRKVRRFIEEAAIPYTYICCNSIAAWPYHDNTHPADVLPPL
ABF95070.1_LAR_Oryza_sativa_Japonica_Group    GHDVDRARPVGAGLRFYEEKRLVRRAAEASGVPYTFICCNSIAGWPYHDSTHPSELPPL
ADD51358.1_LAR_Theobroma_cacao                GHDTDRADPVEPGLTMYEQKRQIRRQVEKSGIPYTYICCNSIAAWPYHDNTHPADVLPPL
ACI41981.1_LAR_Diospyros_kaki                 GHDVDRANPVEPGLAMYIEKRRVRRVVEESGVPYTYICCNSIASWPYYDNTHPSEVLPPL
ABC71329.1_LAR_Lotus_corniculatus             GHDVDRADPVEPGLAMYKEKRLVRRVIEESGIPYTYICCNSIASWPYYNNCHPSQLPPL
ABH07785.2_LAR_Fragaria_x_ananassa            GHDVDRADPVEPGLTMYLEKRKVRRAIEKSGVPYTYICCNSIASWPYYDNKHPSEVVPPL
ABB77697.1_LAR_Pyrus_communis                 GHDVDRADPVEPGLTMYVEKRKVRRWVEKSGVPYTYICCNSIASWPYFDNTHPSEVLPPL
                                          acc
```

```
sp|Q4W2K4_LAR_Vitis_vinifera                β8          β9        α6              η2     β10       η3 β11        α7
                                                  1 9 0              2 0 0         ★2 1 0         2 2 0       2 3 0         2 4 0
sp|Q4W2K4_LAR_Vitis_vinifera               DFFQIYGDGNVKAYFVAGTDIGKFTMKTVDDVRTLNKSVHFRPSCNCLNINELASVWEKK
TRINITY_DN33042_c3_g1_i3                   DEFQIYGDGNTKAYFVAGSDIGKFTMKAVDDVRTINKKVHFQPSCNFYNMNELAALWENK
NP_001352050.1_LAR_Glycine_max             DQLQIYGHGNVKAYFVDGIDIGKFTMKVIDDVRTVNKNVHFRPSNNCYSVNELASLWEKK
ADD51357.1_LAR_Theobroma_cacao             DRFKIYGDGTVKAYFVAGTDIGKFTIMSIEDDRTLNKTVHFQPPSNLLNINEMASLWEEK
CAI56321.1_LAR_Pinus_taeda                 EQFEIYGDGSVKAFFVTGDDIGAYTMKAVDDPRTLNKSIHFRPPKNFLNLNELADIWENK
CAI56322.1_LAR_Phaseolus_coccineus         DQLQIYGHGNVKAYFVDGIDIGKFTMKVIDDVKTINKNVHFRPSKNCYSINELASLGEMK
CAI56320.1_LAR_Hordeum_vulgare_subsp._vulgare DRFQIYGDGTVRAFFVAGTDIGKFTVKAAYDARSVNKAVHFRPACNLLSTNEMACLWESK
CAD79341.1_LAR_Desmodium_uncinatum         DQFQIYGDGNTKAYFIDGNDIGKFTMKTIDDIRTLNKNVHFRPSSNCYSINELASLWEKK
AIS92512_LAR_Epimedium_sagittatum          DQFLIYGDGSVKAYFVDGKDIGKFTMKALDDIRTINKSVHFRPPSNCLSINDMATLWEKK
AII26024.1_LAR_Pisum_sativum               DQLHVYGNGNVKAYFVDGFDIGKFTMKIVDDHRTINKNIHFRPSINCYSMNELASLWESK
AEF14422.1_LAR_Onobrychis_viciifolia       DQLHIYGDGRVQAYFIDGYDIGKFTMKVVDDVRTINKNVHFRPPTNCYSMNGLASLWEKK
ABE90657.1_LAR_Medicago_truncatula         DQLHIYGHGNVKAYFVDGYDIGKFTMKVVDDERTINKSVHFRPSTNCYSMNELASLWENK
CAI56326.1_LAR_Vitis_shuttleworthii        DFFQIYGDGNVKAYFVAGTDIGKFTMKTVDDVRTLNKSVHFRPSCNCLNINELASVWEKK
ABC71327.1_LAR_Lotus_corniculatus          DQFLIYGDGSVKAYFVDGNDIGKFTMKAIDDIRTRNKNVHFRPPSNCYSINELASLWEKI
AHA14498.1_LAR_Fagopyrum_tataricum         DRFQIYGDGNVKAYFVAGSDIGKFTMKTLEDPRTLNKMVHFRPPSNYLTINELASLWEKK
AAZ82410.1_LAR_Vitis_vinifera              DFFQIYGDGNVKAYFVAGTDIGKFTMKTVDDVRTLNKSVHFRPSCNCLNINELASVWEKK
ADY15310.1_LAR_Prunus_avium                DQFQIYGDGSVKAYFIDGTDIGKFTIKTVEDIRTINKNVHFRPPSNLYDINGLASLWEKK
BAH89267.1_LAR_Diospyros_kaki              DHFQIYGDGSVKAYFVAGSDIGKFTIKAADDIRTLNKSIHFRPPCNFLNINELASMWEKK
AAX12186.1_LAR_Malus_domestica             DRFQIYGDGTVKAYFVDGTDIGKFTIKTVDDIRTINKNIHFRPPSNLYDINGLASLWEKK
AEY62396.1_LAR_Fagopyrum_dibotrys          DRFQIYGDGNVKAYFVAGSDIGKFTMKTLEDPRTLNKMVHFRPPSNYLTINELASLWEKK
AAZ79364.1_LAR_Malus_domestica             DRFQIYGDGTVKAYFVDGTDIGKFTMKTVDDMRTINKNVHFRPPSNLYDINGLASLWEKK
CAI56319.1_LAR_Gossypium_arboreum          DHFEIYGDGSVKAYFVAGTDIGKFTMKTVDDIRTLNKSVHFRPACNFYNMNELAALWERK
CAI56323.1_LAR_Gossypium_arboreum          DRFQIYGDGTVKAYFVAGSDIGKFTVMSIDDDRTLNKTVHFQPPSNLLNMNEMASLWETK
CAI56324.1_LAR_Gossypium_raimondii         DHFEIYGDGSVKAYFVAGTDIGKFTMKTVDDIRTLNKSVHFRPACNFYNMNELAALWERK
CAI56325.1_LAR_Gossypium_raimondii         DRFQIYGDGTVKAYFVAGSDIGKFTVMSIDDDRTLNKTVHFQPPSNLLNMNEMASLWETK
CAI56328.1_LAR_Oryza_sativa_Japonica_Group DRFQIYGDGDVRAFFVAGSDIGKFTIRAAYDARSINKIVHFRPACNLLSTNEMASLWESK
CAI26308.1_LAR_Vitis_vinifera              DRFHIYGDGSVKAYFVAGTDIGKFTIETINDDRTVNKSLHFRPPSNLVSINELASLWEKK
ABF95070.1_LAR_Oryza_sativa_Japonica_Group DRFQIYGDGDVRAFFVAGSDIGKFTIRAAYDARSINKIVHFRPACNLLSTNEMASLWESK
ADD51358.1_LAR_Theobroma_cacao             DRFKIYGDGTVKAYFVAGTDIGKFTIMSIEDDRTLNKTVHFQPPSNLLNINEMASLWEEK
ACI41981.1_LAR_Diospyros_kaki              DHFQIYGDGSVKAYFVAGSDIGKFTIKAADDIRTLNKSLHFRPPCNFLNINELASMWEKK
ABC71329.1_LAR_Lotus_corniculatus          DQMHIYGDGTVKAYFVDGFDIGKFTMKLVDDARALNKVVHFRPSSNYYSMNELASLWENK
ABH07785.2_LAR_Fragaria_x_ananassa         DQFQIYGDGTVKAYFVDGPDIGKFTMKTVDDIRTMNKNVHFRPSSNLYDINGLASLWEKK
ABB77697.1_LAR_Pyrus_communis              DRFQIYGDGTVKAYFVDGTDIGKFTIKTAKDIRTINKNVHFRPPSNLYDINGLASLWEKK
acc
```

```
sp|Q4W2K4_LAR_Vitis_vinifera                     β12     α8             α9                       β13  α10
                                                  2 5 0     2 6 0     2 7 0     2 8 0     2 9 0   ★ 3 0 0
sp|Q4W2K4_LAR_Vitis_vinifera               IGRTLPRVTVTEDDLLAAAGENIIPQSVVAAFTHDIFIKGCQVNFSIDGPEDVEVTTLYP
TRINITY_DN33042_c3_g1_i3                   IGRTLPRSTISENDLLAAAAENRIPESIVASFTHDIFIRGCQI.FSIEGPDDVDIATLYP
NP_001352050.1_LAR_Glycine_max             IGRTIPRVTISEDDLLAVAAENCIPRSIVASFTHDIFIKGCQVNFNIDGLGDVEISTLYP
ADD51357.1_LAR_Theobroma_cacao             IGRTLPRVTITEEDLLQMAKEMRIPQSVVAALTHDIFINGCQINFSLDKPTDVEVCSLYP
CAI56321.1_LAR_Pinus_taeda                 INRTLPRVSVSAEDLVMIAKANFMPSSIVAALTHDIFINGCQFKFPIQEPHHVEACELYP
CAI56322.1_LAR_Phaseolus_coccineus         IGRTIPRVTISEDDLLAAAAENCIPQSIVASFTHDIFIKGCQVNFSIDGVDDVEISTLYP
CAI56320.1_LAR_Hordeum_vulgare_subsp._vulgare IGRTLPRVTLSKEELLAMAAEDIIPESIVASLTHDIFINGCQTNFGIDGSRDIEISSLYP
CAD79341.1_LAR_Desmodium_uncinatum         IGRTLPRFTVTADKLLAHAAENIIPESIVSSFTHDIFINGCQVNFSIDEHSDVEIDTLYP
AIS92512_LAR_Epimedium_sagittatum          IKKTLPRKTITEEDLLEAANENRIPESIVAAFTHDIFIKGCQINFPIDGIKDLEATSLYP
AII26024.1_LAR_Pisum_sativum               IARKIPRLFVSENDLLAIAAENIIPESVVASLTHDIFINGCQVSYKIDGVHDVEISTLYP
AEF14422.1_LAR_Onobrychis_viciifolia       LGRKIPRVTVSEDDLLGIAAENCIPESIVASITHDIFINGCQVKFHIDGIHDVEIGTLYP
ABE90657.1_LAR_Medicago_truncatula         IARKIPRAIVSEDDLLGIAAENCIPESVVASITHDIFINGCQVNFKIDGIHDVEISTLYP
CAI56326.1_LAR_Vitis_shuttleworthii        IGRTLPRVTVTEDDLLAAAGENIIPQSVVAAFTHDIFIKGCQVNFSIDGPEDVEVTTLYP
ABC71327.1_LAR_Lotus_corniculatus          IGRKIPRAIVSAEDLLAAAAENCIPRSIVAAFTHDIFINGCQINFSIEGPNDIEIGTLYP
AHA14498.1_LAR_Fagopyrum_tataricum         IGKTLPRVTVTEDDLLDLAAENRIPESIVASFTHDIFIKGCQVDFAVDGPNEVEIEKLYP
AAZ82410.1_LAR_Vitis_vinifera              IGRTLPRVTVTEDDLLAAAGENIIPQSVVAAFTHDIFIKGCQVNFSIDGPEDVEVTTLYP
ADY15310.1_LAR_Prunus_avium                IVRTLPRVTITEDDLLTIAAENRIPESIVASFTHDIFIKGCQINFPVEGPHDVEIGTLYP
BAH89267.1_LAR_Diospyros_kaki              IGRSLPRVTVTEQDLLAAAGENIIPQSIVASFTHDIFIKGCQVNFSIDGPNEVELSSLYP
AAX12186.1_LAR_Malus_domestica             IGRTLPRITITENHLLALAAENRIPESIVASFTHDIFIKGCQVNFAVEGPRDVEVGTLYP
AEY62396.1_LAR_Fagopyrum_dibotrys          IGKTLPRVTVTEDDLLDLAAENRIPESIVASFTHDIFIKGCQVDFAVDGPNEVEIEKLYP
AAZ79364.1_LAR_Malus_domestica             IGRTLPRVTVTEDHLLALAADNRIPESIVASFTHDIFIKGCQVNFAVEGPHDVEVGTLYP
CAI56319.1_LAR_Gossypium_arboreum          IRRTLPRVTVTEEDLLSAAAENIIPQSVVASFTHDIFIKGCQINFPIEGPNETEACSLYP
CAI56323.1_LAR_Gossypium_arboreum          IGRVLPRVTITEQDLLQRAQEMRIPQSVVAAITHDIFINGCQINFSLDKTTDVEICSLYP
CAI56324.1_LAR_Gossypium_raimondii         IRRTLPRVTVTEEDLLSAAAENIIPQSVVASFTHDIFIKGCQINFPIEGPNETEACSLYP
CAI56325.1_LAR_Gossypium_raimondii         IGRVLPRVNITEQDLLQRAQEMRIPQSVVAAITHDIFINGCQINFSLDKTTDVEVCSLYP
CAI56328.1_LAR_Oryza_sativa_Japonica_Group IGRTLPRVTLTEEDLIAMAADDIIPESIVASLTHDIFINGCQTHFYIDGPRDIEISSLYP
CAI26308.1_LAR_Vitis_vinifera              IGRTLPRVTVEEDDLLAAAAEMCIPESIVASFTHDIFIKGCQVNFSLDKPTDLEATSLYP
ABF95070.1_LAR_Oryza_sativa_Japonica_Group IGRTLPRVTLTEEDLIAMAADDIIPESIVASLTHDIFINGCQTHFYIDGPRDIEISSLYP
ADD51358.1_LAR_Theobroma_cacao             IGRTLPRVTITEEDLLQMAKEMRIPQSVVAALTHDIFINGCQINFSLDKPTDVEVCSLYP
ACI41981.1_LAR_Diospyros_kaki              IGRSLPRVTVTEQDLLAAAGENIIPQSIVASFTHDIFIKGCQVNFSIDGPNEVELSSLYP
ABC71329.1_LAR_Lotus_corniculatus          VGRKIPRVTISEDDLLALAAENCIPESIVASITHDIFIKGCQVNYNIDGVHDIEIGTLYP
ABH07785.2_LAR_Fragaria_x_ananassa         IGRTLPKVTITENDLLTMAAENRIPESIVASFTHDIFIKGCQTNFPIEGPNDVDIGTLYP
ABB77697.1_LAR_Pyrus_communis              IGRTLPRITITENHLLALAAENRIPESIVASFTHDIFIKGCQVNFAVEGPRDVEVGTLYP
acc
```

```
                                                       α11
sp|Q4W2K4_LAR_Vitis_vinifera
                                                    310       320
sp|Q4W2K4_LAR_Vitis_vinifera                    EDSFRTVEECFGEYIVKMEE..........K.......QPTADSA.............
TRINITY_DN33042_c3_g1_i3                        EETYQTLDECFEEFLVRLNE......K..NE......TTTVAPKP.............
NP_001352050.1_LAR_Glycine_max                  EEAFRSLEDCFDAFAIMIDD......K..........IHKGENK..............
ADD51357.1_LAR_Theobroma_cacao                  DTPFRTINECFEDFAKKIID......N...........AKAVSKP.............
CAI56321.1_LAR_Pinus_taeda                      DIKYTTMEDFFQGYL...........................................
CAI56322.1_LAR_Phaseolus_coccineus              DEEFRSLEDCYEDFAHMIED......N..........IHKGEHK..............
CAI56320.1_LAR_Hordeum_vulgare_subsp._vulgare   DIPFRTIDECFDDYARGLHL........EEE......A.EESKKS.............
CAD79341.1_LAR_Desmodium_uncinatum              DEKFRSLDDCYEDFVPMVHD......K.........IHAGKSGEIKIKDGKPLVQTGT
AIS92512_LAR_Epimedium_sagittatum               ETPFRTLDDCFDDFLTKTVN....KKN.ADE......RTKIVDKPL............
AII26024.1_LAR_Pisum_sativum                    GESFRSMEDCFESFVVMAAD......K..........IRKGENG..............
AEF14422.1_LAR_Onobrychis_viciifolia            GEEFRSLEDCFGDFVHMAVDNNINNNNN.........NHKGENGG.......GVTTTT
ABE90657.1_LAR_Medicago_truncatula              GESFRSLEDCFESFVAMAAD......K..........IHKGENG..............
CAI56326.1_LAR_Vitis_shuttleworthii             EDSFRTVEECFGEYIVKIEE..........K.......QPTADSA.............
ABC71327.1_LAR_Lotus_corniculatus               DEKFRCLEECFKDFVPMTHD.....MN..........VHVGTT...............
AHA14498.1_LAR_Fagopyrum_tataricum              KDKYITIDECFEEFVITSNN......N..KEIIEEVVVTEAFDDE.............
AAZ82410.1_LAR_Vitis_vinifera                   EDSFRTVEECFGEYIVKIEE..........K.......QPTADSA.............
ADY15310.1_LAR_Prunus_avium                     GESFRTLDECFNDFLLKLED......KLELE...........KNK.............
BAH89267.1_LAR_Diospyros_kaki                   DESFRSVDECFDEFAVKMKD......M..HQ.......EGAKDDG.............
AAX12186.1_LAR_Malus_domestica                  GDSFRTLDECFNDFLLKLKD......N..LEP.....VHEE...N.............
AEY62396.1_LAR_Fagopyrum_dibotrys               KDKYITIDECFEEFVITSNN......N..KEIIEEVVVTEAFDDE.............
AAZ79364.1_LAR_Malus_domestica                  GDSFRTLDECFDGFLLKLKD......NLELEL.....LQEEEDQK.............
CAI56319.1_LAR_Gossypium_arboreum               NEPFRTLDDCFNDFLAKMKD......E.NMK.......QSDENTK.............
CAI56323.1_LAR_Gossypium_arboreum               NTSFRTIAECFDDSAKKISD......N...........EKAVSKP.............
CAI56324.1_LAR_Gossypium_raimondii              NEPFRTLDDCFNDFVAKMKD......E.NMK.......QSDENTK.............
CAI56325.1_LAR_Gossypium_raimondii              NTSFRTIAECFDDFAKKISD......N...........EKAVSKP.............
CAI56328.1_LAR_Oryza_sativa_Japonica_Group      DIPFRTIDECFDDYIHVLNL........AEE......AKEEEEKK.............
CAI26308.1_LAR_Vitis_vinifera                   EMQFRTIDECFDEFVEKIMG......G...........QAAAEK..............
ABF95070.1_LAR_Oryza_sativa_Japonica_Group      DIPFRTIDECFDDYIHVLNL........AEE......AKEEEEKK.............
ADD51358.1_LAR_Theobroma_cacao                  DTPFRTINECFEDFAKKIID......N...........AKAVSKP.............
ACI41981.1_LAR_Diospyros_kaki                   DESFRSVDECFDEFAVKMKD......M..HQ.......EGAKDDG.............
ABC71329.1_LAR_Lotus_corniculatus               DKTFRSLEDCFEDFVTMIVE......K..........IHKGENE..............
ABH07785.2_LAR_Fragaria_x_ananassa              EESFRTLDECFNDFLVKVGG......K..LE...........TDK.............
ABB77697.1_LAR_Pyrus_communis                   GDSFRTLDECFNDFLLKLKD......N..LEL.....VQEEKDQK.............
                                           acc

sp|Q4W2K4_LAR_Vitis_vinifera
                                                                    330
sp|Q4W2K4_LAR_Vitis_vinifera                    ...................IANT...................................GP
TRINITY_DN33042_c3_g1_i3                        ...................IGNE...................................KT
NP_001352050.1_LAR_Glycine_max                  ...................IAG...................................TES
ADD51357.1_LAR_Theobroma_cacao                  ...................AASN...................................NA
CAI56321.1_LAR_Pinus_taeda                      ............................................................
CAI56322.1_LAR_Phaseolus_coccineus              ...................ITG...................................TKS
CAI56320.1_LAR_Hordeum_vulgare_subsp._vulgare   .........................................................NAP
CAD79341.1_LAR_Desmodium_uncinatum              IEEINKDIKTLVETQPNEEIKKD..................................MKA
AIS92512_LAR_Epimedium_sagittatum               ..................EVTSK............HSSGAAEHMEVISRHNTGA...AEQ
AII26024.1_LAR_Pisum_sativum                    ...................VAGG..................................TKS
AEF14422.1_LAR_Onobrychis_viciifolia            ..................ATAGT..................................KKT
ABE90657.1_LAR_Medicago_truncatula              ...................VTGG..................................TKA
CAI56326.1_LAR_Vitis_shuttleworthii             ...................IANT...................................GP
ABC71327.1_LAR_Lotus_corniculatus               ..................EINNN..................................RKS
AHA14498.1_LAR_Fagopyrum_tataricum              ...................IGNKKQSNKRNVENEEDASGNKKRSSMNKITSTAAAAKSSH
AAZ82410.1_LAR_Vitis_vinifera                   ...................IANT...................................GP
ADY15310.1_LAR_Prunus_avium                     ...................VSNKT..................................NA
BAH89267.1_LAR_Diospyros_kaki                   ...................IAAQ...................................NH
AAX12186.1_LAR_Malus_domestica                  ...................VSTK...................................NA
AEY62396.1_LAR_Fagopyrum_dibotrys               ...................IGNKKQSNKRNVENEEDASGNKKRSSMNKITSTAAAAKSSH
AAZ79364.1_LAR_Malus_domestica                  ...................VSTE...................................NT
CAI56319.1_LAR_Gossypium_arboreum               ...................QSNEIPP...............................PKP
CAI56323.1_LAR_Gossypium_arboreum               ..................VTASN...................................TD
CAI56324.1_LAR_Gossypium_raimondii              ...................QSNEIPP...............................PKP
CAI56325.1_LAR_Gossypium_raimondii              ..................VTASN...................................TD
CAI56328.1_LAR_Oryza_sativa_Japonica_Group      .........................................................NAP
CAI26308.1_LAR_Vitis_vinifera                   ...................AAGK...................................EG
ABF95070.1_LAR_Oryza_sativa_Japonica_Group      .........................................................NAP
ADD51358.1_LAR_Theobroma_cacao                  ...................AASN...................................NA
ACI41981.1_LAR_Diospyros_kaki                   ...................IAAQ...................................NH
ABC71329.1_LAR_Lotus_corniculatus               ...................VYG...................................TKS
ABH07785.2_LAR_Fragaria_x_ananassa              ...................LAAK...................................NK
ABB77697.1_LAR_Pyrus_communis                   ...................VSTK...................................NA
                                           acc
```

*sp|Q4W2K4_LAR_Vitis_vinifera*

```
                                                            340
                                                              .
sp|Q4W2K4_LAR_Vitis_vinifera                   V.......V.GM......RQVTATCA........................
TRINITY_DN33042_c3_g1_i3                       VRDETAAMV.EP......LIVTATCA........................
NP_001352050.1_LAR_Glycine_max                 V.......V.EA......VPFKASCGE..EPPPKKCSKFVVTNY......
ADD51357.1_LAR_Theobroma_cacao                 I..FVPTAKPGA......LPITAICT........................
CAI56321.1_LAR_Pinus_taeda                     ..................................................
CAI56322.1_LAR_Phaseolus_coccineus             V.......V.EA......VPIMASCGNIYE....................
CAI56320.1_LAR_Hordeum_vulgare_subsp._vulgare  M.......V.EI......LAVYPTCA........................
CAD79341.1_LAR_Desmodium_uncinatum             L.......V.EA......VPISAMG.........................
AIS92512_LAR_Epimedium_sagittatum              I.......I.ESKTDYLHLILSSTDS...............TPYLGYLGS
AII26024.1_LAR_Pisum_sativum                   M.......V.EP......VIITASC.........................
AEF14422.1_LAR_Onobrychis_viciifolia           L.......I.EA......VPITASC.........................
ABE90657.1_LAR_Medicago_truncatula             L.......V.EP......VPITASC.........................
CAI56326.1_LAR_Vitis_shuttleworthii            V.......V.GM......RQVTATCA........................
ABC71327.1_LAR_Lotus_corniculatus              L.......V.EV......APITAMG.........................
AHA14498.1_LAR_Fagopyrum_tataricum             V.......V.EA......LPVPAVC.........................
AAZ82410.1_LAR_Vitis_vinifera                  V.......V.GM......RQVTATCA........................
ADY15310.1_LAR_Prunus_avium                    V.......V.ET......RAVTATCA........................
BAH89267.1_LAR_Diospyros_kaki                  V.......VEKM......LPITAMCA........................
AAX12186.1_LAR_Malus_domestica                 V.......V.ES......RAVTPTCA........................
AEY62396.1_LAR_Fagopyrum_dibotrys              V.......V.EA......LPVPAVC.........................
AAZ79364.1_LAR_Malus_domestica                 V.......V.ES......RTVTATCA........................
CAI56319.1_LAR_Gossypium_arboreum              V.......V.EA......FAITATCA........................
CAI56323.1_LAR_Gossypium_arboreum              I..FVPTAKPEA......LAITAICT........................
CAI56324.1_LAR_Gossypium_raimondii             V.......V.EA......FAITATCA........................
CAI56325.1_LAR_Gossypium_raimondii             I..FVPTAKPEA......LAITAICT........................
CAI56328.1_LAR_Oryza_sativa_Japonica_Group     T.......V.GR......LAIPPTCA........................
CAI26308.1_LAR_Vitis_vinifera                  I..VVPASAPDA......LVITATCA........................
ABF95070.1_LAR_Oryza_sativa_Japonica_Group     T.......V.GR......LAIPPTCA........................
ADD51358.1_LAR_Theobroma_cacao                 I..FVPTAKPGA......LPITAICT........................
ACI41981.1_LAR_Diospyros_kaki                  V.......VEKM......LPITAMCA........................
ABC71329.1_LAR_Lotus_corniculatus              L.......V.EA......VPITASC.........................
ABH07785.2_LAR_Fragaria_x_ananassa             AA....VGV.EP......MAITATCA........................
ABB77697.1_LAR_Pyrus_communis                  V.......V.ES......RAVTPTCA........................
                                            acc
```

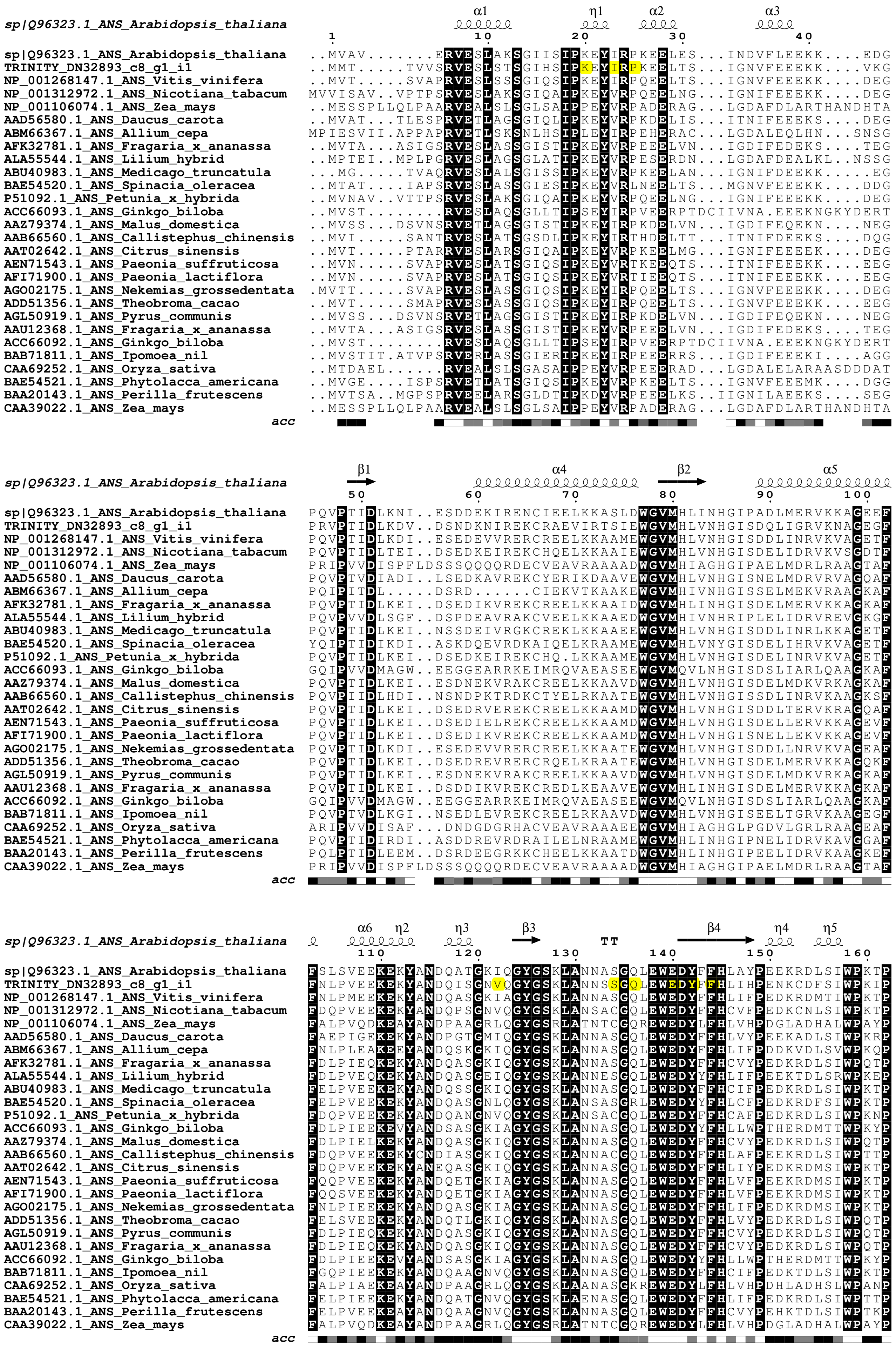

sp|Q96323.1_ANS_Arabidopsis_thaliana
α1
η1
α2
α3
β1
α4
β2
α5
α6
η2
η3
β3
TT
β4
η4
η5
sp|Q96323.1_ANS_Arabidopsis_thaliana
TRINITY_DN32893_c8_g1_i1
NP_001268147.1_ANS_Vitis_vinifera
NP_001312972.1_ANS_Nicotiana_tabacum
NP_001106074.1_ANS_Zea_mays
AAD56580.1_ANS_Daucus_carota
ABM66367.1_ANS_Allium_cepa
AFK32781.1_ANS_Fragaria_x_ananassa
ALA55544.1_ANS_Lilium_hybrid
ABU40983.1_ANS_Medicago_truncatula
BAE54520.1_ANS_Spinacia_oleracea
P51092.1_ANS_Petunia_x_hybrida
ACC66093.1_ANS_Ginkgo_biloba
AAZ79374.1_ANS_Malus_domestica
AAB66560.1_ANS_Callistephus_chinensis
AAT02642.1_ANS_Citrus_sinensis
AEN71543.1_ANS_Paeonia_suffruticosa
AFI71900.1_ANS_Paeonia_lactiflora
AGO02175.1_ANS_Nekemias_grossedentata
ADD51356.1_ANS_Theobroma_cacao
AGL50919.1_ANS_Pyrus_communis
AAU12368.1_ANS_Fragaria_x_ananassa
ACC66092.1_ANS_Ginkgo_biloba
BAB71811.1_ANS_Ipomoea_nil
CAA69252.1_ANS_Oryza_sativa
BAE54521.1_ANS_Phytolacca_americana
BAA20143.1_ANS_Perilla_frutescens
CAA39022.1_ANS_Zea_mays
acc

```
                                              α7                           α8        α9            β5
sp|Q96323.1_ANS_Arabidopsis_thaliana
                                         170          180          190          200          210
sp|Q96323.1_ANS_Arabidopsis_thaliana     SDYIEATSEYAKCLRLLATKVFKALSVGLGL.EP.DRLEKEVGGL.......EELLLQMK
TRINITY_DN32893_c8_g1_i1                 SDYRDATSEYARQLRALATKILSILSLGLGV.EE.GRLEKEVGGV.......EDLLLQMK
NP_001268147.1_ANS_Vitis_vinifera        SDYVPATCEYSVKLRSLATKILSVLSLGLGL.EE.GRLEKEVGGM.......EELLLQKK
NP_001312972.1_ANS_Nicotiana_tabacum     TDYIPATSEYAKQIRNLATKILAVLSIGLRL.EE.GRLEKEVGGM.......EDLLLQMK
NP_001106074.1_ANS_Zea_mays              PDYIAATRDFGRRTRDLASTLLAILSMGLLGTDRGDALEKALTTTTTRTAADDDLLLQLK
AAD56580.1_ANS_Daucus_carota             QDYIPATREYAKELRGLATKLLSALSFGLGL.EE.GRLEKEVGGM.......EELILQMK
ABM66367.1_ANS_Allium_cepa               SDYIEIMQEFGSQLRILASKMLSILSLGLQLPTK.DRLEQELKGP.......EDLLLQLK
AFK32781.1_ANS_Fragaria_x_ananassa       SDCIVATSEYAKELRGLATKMLSILSLGLGL.EE.GRLEKEVGGL.......EELLLQMK
ALA55544.1_ANS_Lilium_hybrid             EDYTEVTKEFAKELRVVVTKMLSMLSQGLGL.ES.GKLEKELGGM.......DDLLMQMK
ABU40983.1_ANS_Medicago_truncatula       ADYTKVTSEYAKELRVLASKIMEVLSLELGL.EG.GRLEKEAGGM.......EELLLQMK
BAE54520.1_ANS_Spinacia_oleracea         ADYIPAVSEYSKELRSLATKILSALSLALGL.EE.GRLEKEVGGI.......EELLLQFK
P51092.1_ANS_Petunia_x_hybrida           TDYTPATSEYAKQIRALATKILTVLSIGLGL.EE.GRLEKEVGGM.......EDLLLQMK
ACC66093.1_ANS_Ginkgo_biloba             SDYIEATDEYGREIRKLVSRILGTLSMELGL.EE.PRMETILGG.........QDLEMQLK
AAZ79374.1_ANS_Malus_domestica           ADYIEATAEYAKQLRELATKVLKVLSLGLGL.DE.GRLEKEVGGL.......EELLLQMK
AAB66560.1_ANS_Callistephus_chinensis    HDYIPATTEYAKQLRALATKILAALSLGLGL.EE.GRLEKEVGGM.......EELLVQLK
AAT02642.1_ANS_Citrus_sinensis           SDYTEATSEYARQLRSLATKILAVLSLGLGL.EE.GRLEKEVGGL.......EELLLQMK
AEN71543.1_ANS_Paeonia_suffruticosa      SDYTEATSEYAKQLRILVTKILSILSIGLGL.EE.GRLEQEVGGM.......EELLLQMK
AFI71900.1_ANS_Paeonia_lactiflora        SDYIEATSEYAKQLRILVTKILSIPSIGLGL.EE.GRLEQEVGGM.......EELLLQMK
AGO02175.1_ANS_Nekemias_grossedentata    SDYIPATCEYSVELRGLATKILSVLSLGLGL.EE.GRLEKEVGGM.......EELLLQKK
ADD51356.1_ANS_Theobroma_cacao           SDYTEVTSEYARQLRVLASKILSALSLCLGL.EE.GRLEKEVGGL.......EELLLQMK
AGL50919.1_ANS_Pyrus_communis            ADYIEATAEYAKQLRELATKVLKVLSLGLGL.DE.GRLEKEVGGL.......EELLLQMK
AAU12368.1_ANS_Fragaria_x_ananassa       SDYIVATSEYAKELRGLATKILSILSLGLGL.EE.GRLEKEVGGL.......EELLLQMK
ACC66092.1_ANS_Ginkgo_biloba             SDYIEATDEYGREIRKLVSRILGTLSMELGL.EE.PRMETILGG.........QDLEMQLK
BAB71811.1_ANS_Ipomoea_nil               SDYIDATKEYAKQLRALATKVLAVLSLGLGL.EE.GRLEKEVGGM.......EELLLQMK
CAA69252.1_ANS_Oryza_sativa              PEYVPVSRDFGGRVRTLASKLLAILSLGLGLPE..ETLERRLRGHEL.AGVDDDLLLQLK
BAE54521.1_ANS_Phytolacca_americana      SDYICAVSEYAKELRALATKILSALSLALGL.EE.PRLEKEVGGL.......EELLLQLK
BAA20143.1_ANS_Perilla_frutescens        PDYIPATSEYAKQLRALATKILSVLSIGLGL.EK.GRLEKEVGGA.......EDLIVQMK
CAA39022.1_ANS_Zea_mays                  PDYIAATRDFGRRTRDLASTLLAILSMGLLGTDRGDALEKALTTTTTRTAADDDLLLQLK
                                    acc
```

```
                                                   TT     β6           β7            β8   TT  β9       TT    β10
sp|Q96323.1_ANS_Arabidopsis_thaliana
                                              220          230          240          250          260          270
sp|Q96323.1_ANS_Arabidopsis_thaliana     INYYPKCPQPELALGVEAHTDVSALTFILHNMVPGLQLFYEGKWVTAKCVPDSIVMHIGD
TRINITY_DN32893_c8_g1_i1                 INYYPKCPQPELALGVEAHTDISSLTFILHNMVPGLQLFYKGKWVTAKCVPNSIIMHIGD
NP_001268147.1_ANS_Vitis_vinifera        INYYPKCPQPELALGVEAHTDVSALTFILHNMVPGLQLFYEGKWVTAKCVPNSIIMHIGD
NP_001312972.1_ANS_Nicotiana_tabacum     INYYPKCPQPELALGVEAHTDVSALTFILHNMVPGLQLFYEGQWVTAKCVPNSIIMHIGD
NP_001106074.1_ANS_Zea_mays              INYYPRCPQPELAVGVEAHTDVSALSFILHNGVPGLQVLHGARWVTARHEPGTIIVHVGD
AAD56580.1_ANS_Daucus_carota             INYYPKCPQPELALGVEAHTDVSALSFILHNTVPGLQLFYGGKWVTAKCVPDSIIVHIGD
ABM66367.1_ANS_Allium_cepa               INYYPKCPQPHLALGVEAHTDVSALSFILHNNVPGLQVLYEGEWVTAKLVPDSLIVHVGD
AFK32781.1_ANS_Fragaria_x_ananassa       INYYPKCPLPELALGVEAHTDISALTFILHNMVPGLQLFYGGKWVTAKCVPNSVVMHIGD
ALA55544.1_ANS_Lilium_hybrid             INYYPKCPQPELALGVEAHTDVSSLTFLLTNMVPGLQLYYGGKWVIAQCVPDSLLVHIGD
ABU40983.1_ANS_Medicago_truncatula       INYYPICPQPELALGVEAHTDVSSLTFLLHNMVPGLQLFYEGKWVTAKCVPDSILMHIGD
BAE54520.1_ANS_Spinacia_oleracea         INYYPKCPQPELALGVEAHTDVSALTFILHNMVPGLQLFYEGKWVTAKCVPNSIIMHIGD
P51092.1_ANS_Petunia_x_hybrida           INYYPKCPQPELALGVEAHTDVSALTFILHNMVPGLQLFYEGQWVTAKCVPNSIIMHIGD
ACC66093.1_ANS_Ginkgo_biloba             INYYPRCPQPELALGVEAHTDISALTFLLHNMVPGFQLFKDGKWVTAKCIPGALIVHIGD
AAZ79374.1_ANS_Malus_domestica           INYYPKCPQPELALGVEAHTDVSALTFILHNMVPGLQLFYEGKWVTAKCVPNSIVMHIGD
AAB66560.1_ANS_Callistephus_chinensis    INYYPKCPQPDVALGVEAHTDVSALTFILHNMVPGLQLFYGGKWVTAKCVPDSIIMHIGD
AAT02642.1_ANS_Citrus_sinensis           INYYPKCPQPELALGVEAHTDVSALTFILHNMVPGLQLFYKDKWVTAKCVPNSIILHIGD
AEN71543.1_ANS_Paeonia_suffruticosa      INYYPKCPQPELALGVEAHTDVSALTFILHNMVPGLQLFYQGKWVTAKCVPNSIIMHIGD
AFI71900.1_ANS_Paeonia_lactiflora        INYYPKCPQPELALGVEAHTDVSALTFILHNMVPGLQLFYQGKWVTAKCVPNSIIMHIGD
AGO02175.1_ANS_Nekemias_grossedentata    INYYPKCPQPELALGVEAHTDVSALTFILHNMVPGLQLFYEGKWVTAKCVPNSIIMHIGD
ADD51356.1_ANS_Theobroma_cacao           INYYPKCPQPELALGVEAHTDVSALTFILHNMVPGLQLFYEGKWITAKCVPNSIIMHIGD
AGL50919.1_ANS_Pyrus_communis            INYYPKCPQPELALGVEAHTDVSALTFILHNMVPGLQLFYEGKWVTAKCVPNSIVMHIGD
AAU12368.1_ANS_Fragaria_x_ananassa       INYYPKCPQPELALGVEAHTDISALTFILHNMVPGLQLFYGGKWVTAKCVPNSIVMHIGD
ACC66092.1_ANS_Ginkgo_biloba             INYYPRCPQPELALGVEAHTDISALTFLLHNMVPGFQLFKDGKWVTAKCIPGALIVHIGD
BAB71811.1_ANS_Ipomoea_nil               INYYPKCPQPELALGVEAHTDVSALTFILHNMVPGLQLFYGGKWVTAKCVPNSIIMHVGD
CAA69252.1_ANS_Oryza_sativa              INYYPRCPRPDLAVGVEAHTDVSALSFILHNGVPGLQVHHAGSWVTARPEPGTIVVHVGD
BAE54521.1_ANS_Phytolacca_americana      INYYPKCPQPELALGVEAHTDVSALTFILHNMVPGLQLFYEGKWVTAKCVPNSIIMHIGD
BAA20143.1_ANS_Perilla_frutescens        INFYPKCPQPELALGWEAHTDVSALTFILHNMVPGLQLFYEDKWVTAKCVPNSIIMHIGD
CAA39022.1_ANS_Zea_mays                  INYYPRCPQPELAVGVEAHTDVSALSFILHNGVPGLQVLHGARWVTARHEPGTIIVHVGD
                                    acc
```

```
                                         α10            β11          β12          TTT  β13   η6
sp|Q96323.1_ANS_Arabidopsis_thaliana
                                                280          290          300          310          320          330
sp|Q96323.1_ANS_Arabidopsis_thaliana     TLEILSNGKYKSILHRGLVNKEKVRISWAVFCEPPKDKIVLKPLPEMVSV..ESPAKFPP
TRINITY_DN32893_c8_g1_i1                 TIEILSNGKYKSILHRGLVNKEKVRISWAVFCEPPKETIILKPLPEVVTE..DEPPLFPP
NP_001268147.1_ANS_Vitis_vinifera        TIEILSNGKYKSILHRGLVNKEKVRISWAVFCEPPKEKIILKPLPETVSE..TEPPLFPP
NP_001312972.1_ANS_Nicotiana_tabacum     TLEILSNGKYKSILHRGVVNKEKIRISWAIFCEPPKEKIILKPLPETITE..AEPPRFPP
NP_001106074.1_ANS_Zea_mays              ALEILSNGRYTSVLHRGLVNREAVRISWVVFCEPPPDSVLLHPLPELVTE..GHPARFTP
AAD56580.1_ANS_Daucus_carota             TLEILSNGKYKSILHRGLVNKEKVRISWAVFCEPPKEKIILKPLPETVTE..AEPPLFPP
ABM66367.1_ANS_Allium_cepa               SLEILSNGIYKSVLHRGLVNKEKVRISWAVFCEPPKDAVVLKPLDEVVTD..DAPARYTP
AFK32781.1_ANS_Fragaria_x_ananassa       TLEILSNGKYKSILHRGLVNKEKVRISWAVFCEPPKEKIILKPLPETVSE..EEPAIFPP
ALA55544.1_ANS_Lilium_hybrid             TLEILSNGRYRSILHRSLVNKERVRISWAVFCEPPKETIVLKPLPELVTE..GAPAKFPP
ABU40983.1_ANS_Medicago_truncatula       TIEILSNGKYKSILHRGLVNKEKVRISWAVFCEPPKEKIILKPLPELVTE..KEPARFPP
BAE54520.1_ANS_Spinacia_oleracea         TIEILSNGKYKSILHRGLVNKEKVRISWAVFCEPPKEKIVLMPLPDLVSG..EEVARYPP
P51092.1_ANS_Petunia_x_hybrida           TIEILSNGKYKSILHRGVVNKEKVRFSWAIFCEPPKEKIILKPLPETVTE..AEPPRFPP
ACC66093.1_ANS_Ginkgo_biloba             QVEILSNGKYKSGLHRGLVNKEKVRISWAVFCDPPKDAMI.GPMKELINE..KNPPLFAA
AAZ79374.1_ANS_Malus_domestica           TLEILSNGKYKSILHRGMVNKEKVRISWAVFCEPPKEKIILKPLPETVSE..EEPAMFPP
AAB66560.1_ANS_Callistephus_chinensis    TIEILSNGKYKSILHRGLVNKEKVRISWAVFCEPPKDKIILKPLPETISE..DEPPLFPP
AAT02642.1_ANS_Citrus_sinensis           TIEILSNGEYKSILHRGLVNKEKVRISWAVFCEPPKDKIILKPLPETVSE..QKPAMFPP
AEN71543.1_ANS_Paeonia_suffruticosa      TIEILSNGKYKSILHRGLVNKDKVRISWAVFCEPPKEKIILKPVLEVVSE..EEPALFPP
AFI71900.1_ANS_Paeonia_lactiflora        TIEILSNGKYKSILHRGLVNKEKVRISWAVFCEPPKEKIILKPLPEVVSE..DEPALFPP
AGO02175.1_ANS_Nekemias_grossedentata    TIEILSNGKYKSILHRGLVNKEKVRISWAVFCEPPKEKIILKPLPETVSE..TEPPLFPP
ADD51356.1_ANS_Theobroma_cacao           TVEILSNGKYKSILHRGLVNKEKVRISWAVFCEPPKEKIILKPLPETVSE..TEPPLFPP
AGL50919.1_ANS_Pyrus_communis            TLEILSNGKYKSILHRGMVNKEKVRISWAVFCEPPKEKIILKPLPETVSE..DEPAMFPP
AAU12368.1_ANS_Fragaria_x_ananassa       TLEILSNGKYKSILHRGLVNKEKVRISWAVFCEPPKEKIILKPLPETVSE..EEPAIFPP
ACC66092.1_ANS_Ginkgo_biloba             QVEILSNGKYKSGLHRGLVNKEKVRISWAVFCDPPKDAMI.GPMKELINE..KNPPLFAA
BAB71811.1_ANS_Ipomoea_nil               TVEILSNGKYKSILHRGVVNREKVRVSWAVFCEPPKDKILLQPLPETVSE..AEPPRFPP
CAA69252.1_ANS_Oryza_sativa              ALEILTNGRYTSVLHRGLVSRDAVRLSWVVFCEPPPESVLLQPVQELLADGAGKPL.FAP
BAE54521.1_ANS_Phytolacca_americana      TIEILSNGKYKSILHRGLVNKEKVRISWAVFCEPPKEKIILKPLTELLTE..EEPARFPP
BAA20143.1_ANS_Perilla_frutescens        TLEILSNGKYKSILHRGLVNKEKVRISWAVFCEPPKEKIVLQPLPETVSE..VEPPRFPP
CAA39022.1_ANS_Zea_mays                  ALEILSNGRYTSVLHRGLVNREAVRISWVVFCEPPPDSVLLHPLPELVTE..GHPARFTP
                                    acc
```

```
                                         β14                       α11
sp|Q96323.1_ANS_Arabidopsis_thaliana     →ℓℓℓℓℓℓℓℓℓℓℓℓ.............................ℓ
                                               340                                      350

sp|Q96323.1_ANS_Arabidopsis_thaliana     RTFAQHIEHKLFGK.............................EQEELVSEKND......
TRINITY_DN32893_c8_g1_i1                 RTFAQHLEHKLFRK.............................SQEALLANK........
NP_001268147.1_ANS_Vitis_vinifera        RTFSQHIQHKLFRK.............................TQEALLSK.........
NP_001312972.1_ANS_Nicotiana_tabacum     RTFAQHMAHKLFKKDDQDAAAEHKVSKKDDPDSAAEHKPFKKDDQDAVAQQKV...LKED
NP_001106074.1_ANS_Zea_mays              RTFKQHLDRKLFKK.............................KQQHKAKAEEE......
AAD56580.1_ANS_Daucus_carota             RTFAQHMAHKLFRK.............................SQEAIDDSKK...VQPQ
ABM66367.1_ANS_Allium_cepa               RTFAQHLERKLFKK.........KVGDL...............DDSDV............
AFK32781.1_ANS_Fragaria_x_ananassa       RTFFEHIQHKLFRQ.............................SQEALVSTKESAALKST
ALA55544.1_ANS_Lilium_hybrid             RTFKQHIQHKLFKK.............................TEEDFTSLK........
ABU40983.1_ANS_Medicago_truncatula       RTFAQHIHHKLFRK.............................DEEEKKDDPK.......
BAE54520.1_ANS_Spinacia_oleracea         RTFAQHVQYKLFRK.............................TQD...P..........
P51092.1_ANS_Petunia_x_hybrida           RTFAQHMAHKLFRKDDKDAAVEHKVFNEDELDTAAEHKVLKKDNQDAVAENKD...IKED
ACC66093.1_ANS_Ginkgo_biloba             KTFKDHTDHKLFKK.............................GQSKKN...........
AAZ79374.1_ANS_Malus_domestica           RTFAEHIQHKLFRK.............................SQGALLPK.........
AAB66560.1_ANS_Callistephus_chinensis    RTFQQHMEHKLFRK.............................NNDAVDPK.........
AAT02642.1_ANS_Citrus_sinensis           RTFQQHIEHKLFRR.............................TQDALLSDEE.......
AEN71543.1_ANS_Paeonia_suffruticosa      RTFAQHIQHKLFRK.............................TQEEFKN..........
AFI71900.1_ANS_Paeonia_lactiflora        RTFAQHIQHKLFRK.............................TQEEFKN..........
AGO02175.1_ANS_Nekemias_grossedentata    RTFAQHIQHKLFRK.............................TQEALLTK.........
ADD51356.1_ANS_Theobroma_cacao           RTFAQHIHHKLFRK.............................TQDGLSN..........
AGL50919.1_ANS_Pyrus_communis            RTFAEHIQHKLFRK.............................SQEALLPK.........
AAU12368.1_ANS_Fragaria_x_ananassa       RTFFEHIQHKLFRQ.............................SQEALVSTKESAALKST
ACC66092.1_ANS_Ginkgo_biloba             KTFKDHTDHKLFKK.............................GQSKKN...........
BAB71811.1_ANS_Ipomoea_nil               RTFAQHIKHKLFRQ............................SDQEAADTPKPD...NDD
CAA69252.1_ANS_Oryza_sativa              RTFKQHVQRKLFKK..........................LKDQQDNNAAA.........
BAE54521.1_ANS_Phytolacca_americana      RTFAQHIQYKLFRK.............................TQDVQAPVSN.......
BAA20143.1_ANS_Perilla_frutescens        RTFAQHLKHKLFRK.............................TDGDLDEKPTY......
CAA39022.1_ANS_Zea_mays                  RTFKQHLDRKLFKK.............................KQQHKAKAEEE......
                                     acc

sp|Q96323.1_ANS_Arabidopsis_thaliana

sp|Q96323.1_ANS_Arabidopsis_thaliana     .......................................................
TRINITY_DN32893_c8_g1_i1                 .......................................................
NP_001268147.1_ANS_Vitis_vinifera        .......................................................
NP_001312972.1_ANS_Nicotiana_tabacum     E..............QNAAAEHKVFKKDNQDAAAEESK.................
NP_001106074.1_ANS_Zea_mays              ............DGGNGDHHRHEPPPQTN..........................
AAD56580.1_ANS_Daucus_carota             EQ.............NNAETD...IPQPEEQQKTEESNPQKIEILKPGEEASSSP
ABM66367.1_ANS_Allium_cepa               .......................................................
AFK32781.1_ANS_Fragaria_x_ananassa       T..............ESAL.......KSTKEAALISTN.................
ALA55544.1_ANS_Lilium_hybrid             .......................................................
ABU40983.1_ANS_Medicago_truncatula       ..........................K............................
BAE54520.1_ANS_Spinacia_oleracea         .......................................................
P51092.1_ANS_Petunia_x_hybrida           EQCGPAEHKDIKEDGQGAAAENKVFKENNQDVAAEESK.................
ACC66093.1_ANS_Ginkgo_biloba             .......................................................
AAZ79374.1_ANS_Malus_domestica           .......................................................
AAB66560.1_ANS_Callistephus_chinensis    .......................................................
AAT02642.1_ANS_Citrus_sinensis           .......................................................
AEN71543.1_ANS_Paeonia_suffruticosa      .......................................................
AFI71900.1_ANS_Paeonia_lactiflora        .......................................................
AGO02175.1_ANS_Nekemias_grossedentata    .......................................................
ADD51356.1_ANS_Theobroma_cacao           .......................................................
AGL50919.1_ANS_Pyrus_communis            .......................................................
AAU12368.1_ANS_Fragaria_x_ananassa       T..............ESAL.......KSTKEAALISTN.................
ACC66092.1_ANS_Ginkgo_biloba             .......................................................
BAB71811.1_ANS_Ipomoea_nil               HH........................QSN..........................
CAA69252.1_ANS_Oryza_sativa              .............ASNGMITK..................................
BAE54521.1_ANS_Phytolacca_americana      .......................................................
BAA20143.1_ANS_Perilla_frutescens        .......................................................
CAA39022.1_ANS_Zea_mays                  ............DGGNGDHHRHEPPPQTN..........................
                                     acc
```

```
BAD89742.1_ANR_Vitis_vinifera                                                          β1
                                                                        1          10         20
BAD89742.1_ANR_Vitis_vinifera                  ...............................MATQHP.....IGKKTACVVGGTGFVASL
TRINITY_DN30161_c9_g1_i2                       ...............................MATQL......NPNKTACVIGGSGFVASL
TRINITY_DN30161_c9_g1_i3                       ...............................MATQL......NPNKTACVIGGSGFVASL
Q9SEV0.2_ANR_AT1G61720_Arabidopsis_thaliana    ..................................MDQTL.THTGSKKACVIGGTGNLASI
AKV89239.1_ANR_Prunus_cerasifera               ...............................MAT.QP.....ISKKTACVIGGTGFVASL
AJK93561.1_ANR_Vicia_faba                      ...............................MASIK.....EIGKKKACVIGGTGFVASM
AII26022.1_ANR_Pisum_sativum                   ...............................MASIK.....EVGKKKACVIGGTGFVASL
AGL81352.1_ANR_Pyrus_communis                  ...............................MATQQP.....ISKKTACVVGGTGFVASL
ACV72641.1_ANR_Gossypium_hirsutum              ...............................MASQL......VGTKRACVVGGSGFVASL
ADD51353.1_ANR_Theobroma_cacao                 ...............................MASQT......VGKKTACVVGGTGYVASL
AAT68773.1_ANR_Camellia_sinensis               ...............................MEAQP......TAPKAACVVGGTGFVAAT
ABM64802.1_ANR_Gossypium_hirsutum              ...............................MASQI......VGTKRACVVGGSGFVASL
AAN77735.1_ANR_Medicago_truncatula             ...............................MASIKQ...IEIEKKKACVIGGTGFVASL
ASU87432.1_ANR_Camellia_sinensis               ...............................MEAQP......TAPKAACVVGGTGFVAAT
AEC10993.1_ANR_Camellia_sinensis               ...............................MEAQP......TAPKAACVVGGTGFVAAT
ADZ58168.1_ANR_Camellia_sinensis               ...........................MAMAMATTTTTTKPMIGAKAACAVGGTGFVAAT
AHJ11240.1_ANR_Camellia_sinensis               ...........................MAMAMATTTTATKPMIGAKAACVVGGTGFVAAT
NP_001267885.1_ANR_Vitis_vinifera              ...............................MATQHP.....IGKKTACVVGGTGFVASL
ABD95362.1_ANR_Fragaria_x_ananassa             ...............................MSTTQP....IISTKSACVIGGTGFVASQ
AEL79861.1_ANR_Malus_domestica                 ...............................MATQQP.....ISKKTACVVGGTGFVASL
BAF56654.1_ANR_Diospyros_kaki                  ...............................MAAAPPP...TLGAKAACVIGGTGFVAAT
ADD51354.1_ANR_Theobroma_cacao                 ...............................MASQT......VGKKTACVVGGTGYVASL
CAD91909.1_ANR_Phaseolus_coccineus             ...............................MATVK.....KIG.KKACVIGGSGFMASC
ABM90632.1_ANR_Lotus_uliginosus                ...............................MASIKP....IYEKKKACVIGGTGFVASL
ABC71336.1_ANR_Lotus_corniculatus              ...............................MASIKP....IYEKKKACVIGGTGFVASL
XP_002317270.2_ANR_Populus_trichocarpa         MHPPMEELNQIKFIFNLARKNSDMIHIFSSSMASQT......K.KNTACVIGGTGFVASL
CAD91910.1_ANR_Gossypium_arboreum              ...............................MASQL......VGTKRACVVGGSGFVASL
AEL79859.1_ANR_Malus_domestica                 ...............................MATQQP.....ISKKTACVVGGTGFVASL
AEL79860.1_ANR_Malus_domestica                 ...............................MATQQP.....ISKKTACVVGGTGFVASL
AEL79861.1_ANR_Malus_domestica                 ...............................MATQQP.....ISKKTACVVGGTGFVASL
ACY30421.1_C.BANa_Brassica_napus               ...............................MTSTDQT....TGTKKVCVIGGTGNLASI
ACY30422.1_C.BANb_Brassica_napus               ...............................MATVDQTA.VTTGTKKACVIGGTGNLASI
ACY30423.1_A.BANa_Brassica_napus               ...............................MTSTDQT....TGTKKVCVIGGTGNLASI
ACY30424.1_A.BANb_Brassica_napus               ...............................MATVDQTA.VTTGTKKACVIGGTGNLASI
ACY30425.1_C.BANa_Brassica_oleracea            ...............................MTSTDQT....TGTKKVCVIGGTGNLASI
ABG76842.1_ANR_Fragaria_x_ananassa             ...............................MATTQP....IISTKSACVIGGTGFVASQ
ABC71337.1_ANR-1_Lotus_corniculatus            ...............................MASIKP....IYEKKKACVIGGTGFVASL
ABC71332.1_ANR1-1_Lotus_corniculatus           ...............................MASIKP....IYEKKKACVIGGTGFVASL
ABC71333.1_ANR1-2_Lotus_corniculatus           ...............................MASIKP....IYEKKKACVIGGTGFVASL
ABC71335.1_ANR1-4_Lotus_corniculatus           ...............................MASIKP....IYEKKKACVIGGTGFVASL
AAZ79363.1_ANR_Malus_domestica                 ...............................MATQQP.....ISKKTACVVGGTGFVASL
AAX12184.1_ANR_Malus_domestica                 ...............................MATQQP.....ISKKTACVVGGTGFVASL
AAZ17408.1_ANR_Malus_domestica                 ...............................MATQQP.....ISKKTACVVGGTGFVASL
EEE86150.1_ANR_Populus_trichocarpa             ...............................MASQL......T.KKTACVIGGTGFVASL
EEE97882.1_ANR_Populus_trichocarpa             ...............................MASQT......K.KNTACVIGGTGFVASL
ABB77695.1_ANR_Pyrus_communis                  ...............................MATQQP.....ISKKTACVVGGTGFVASL
BAD89742.1_ANR_Vitis_vinifera                  ...............................MATQHP.....IGKKTACVVGGTGFVASL
AAZ82409.1_ANR_Vitis_vinifera                  ...............................MATQHP.....IGKKTACVVGGTGFVASL
                                         acc

                                               α1          β2              α2      η1        β3              α3
BAD89742.1_ANR_Vitis_vinifera                        30        40        50        60        70        80
BAD89742.1_ANR_Vitis_vinifera                  LVKLLLQKGYAVNTTVRDPDNQKKVSHLLELQELGDLKIFRADLTDELSFEAPIAGCDFV
TRINITY_DN30161_c9_g1_i2                       LVKLLLQKGYTVHTTVRDPANQKKISHLLPLKELGGLRIFGADLTNEESFNAPIAGCELV
TRINITY_DN30161_c9_g1_i3                       LVKLLLQKGYTVHTTVRDPANQKKISHLLPLKELGGLRIFGADLTNEESFNAPIAGCELV
Q9SEV0.2_ANR_AT1G61720_Arabidopsis_thaliana    LIKHLLQSGYKVNTTVRDPENEKKIAHLRKLQELGDLKIFKADLTDEDSFESSFSGCEYI
AKV89239.1_ANR_Prunus_cerasifera               LVKLLLEKGYAVKTTVRDPDNQKKISHLTALQDLGELEIFAADLTDEGSFDAPIAGCDLV
AJK93561.1_ANR_Vicia_faba                      LIKQLLEKGYAVNTTVRDPDNPKKTSHLVALQSLGELNMFKAELTVEEDFDAPIAGCELV
AII26022.1_ANR_Pisum_sativum                   LIKQLLEKGYAVNTTVRDLDNPKKISHLVALKSLGELNLFKAELTVDGDFDAPIAGCELV
AGL81352.1_ANR_Pyrus_communis                  LVKLLLQKGYAVRTTVRDPDNHKKVSHLTALQELGELEILAGDLTDEGSFDAPIAGCDLV
ACV72641.1_ANR_Gossypium_hirsutum              LVKLLLEKGYAVNTTVRDPDNQKKISHLVTLQELGDLKIFQADLTDEGSFDAPIAGCDLV
ADD51353.1_ANR_Theobroma_cacao                 LVKLLLEKGYAVNTTVRDPDNQKKIPHLVTLQKLGDLKIFRADLTDEGSFDVPIAGCDLV
AAT68773.1_ANR_Camellia_sinensis               LIKLLLEKGYAVNTTVRDPGNQKKTSHLLALKGSGNLKIFRADLTDEQSFDTPVAGCDLV
ABM64802.1_ANR_Gossypium_hirsutum              LVKLLLEKGYAVNTTVRDPDNQKKISHLVTLQELGDLKIFQADLTDEGSFDAPIAGCDLV
AAN77735.1_ANR_Medicago_truncatula             LIKQLLEKGYAVNTTVRDLDSANKTSHLIALQSLGELNLFKAELTIEEDFDAPISGCELV
ASU87432.1_ANR_Camellia_sinensis               LIKLLLEKGYAVNTTVRDPGNQKKTSHLLALKGSGNLKIFRADLTDEQSFDTPVAGCDLV
AEC10993.1_ANR_Camellia_sinensis               LIKLLLEKGYAVNTTVRDPGNQKKTSHLLALKGSGNLKIFRADLTDEQSFDTPVAGCDLV
ADZ58168.1_ANR_Camellia_sinensis               LVKMLLERGYSVNTTVRDPDNKKNISHLVALEGMGNLKIFRADLTDEQSFDAPIAGCDLV
AHJ11240.1_ANR_Camellia_sinensis               LVKMLLERGYSVNTTVRDPDNKKNISHLVALEGMGNLKIFRADLTDEQSFDAPIAGCDLV
NP_001267885.1_ANR_Vitis_vinifera              LVKLLLQKGYAVNTTVRDPDNQKKVSHLLELQELGDLKIFRADLTDELSFEAPIAGCDFV
ABD95362.1_ANR_Fragaria_x_ananassa             LIKLLLEKGYAVRTTVRDPDNLKKISHLTALQELGELTIFRGDLTDEGSFDAAIAGSDLV
AEL79861.1_ANR_Malus_domestica                 LVKLLLQKGYAVRTTVRDPDNHKKVSHLTALQELGELEILAGDLTDEGSFDAPIAGCDLV
BAF56654.1_ANR_Diospyros_kaki                  LIKLLLEKGYAVNTTVRDPDDKKKISHLLTMKSLGSLNIFQAELTDEKSFDAPIAGCDLV
ADD51354.1_ANR_Theobroma_cacao                 LVKLLLEKGYAVNTTVRDPDNQKKIPHLVTLQKLGDLKIFRADLTDEGSLDVPIAGCDLV
CAD91909.1_ANR_Phaseolus_coccineus             LVKQLLHKGYAVNTTVRDPDNAKKISHLLALQSLGDLNIFGADLTGERDFDAPIAGCELV
ABM90632.1_ANR_Lotus_uliginosus                LIKQLLEKGYAVNTTVRDPDNHKKISHLVALQSLGELNIFGGELTVENDFDAPIAGSQLV
ABC71336.1_ANR_Lotus_corniculatus              LIKQLLEKGYAVNTTVRDP....................................GSELV
XP_002317270.2_ANR_Populus_trichocarpa         LIKLLLEKGYAVNTTVRDPDNQKKIAHLIALQNLGDLNIFGADLTNEESFNAPIACCDLV
CAD91910.1_ANR_Gossypium_arboreum              LVKLLLEKGFAVNTTVRDPDNQKKISHLVTLQELGDLKIFQADLTDEGSFDAPIAGCDLV
AEL79859.1_ANR_Malus_domestica                 LVKLLLQKGYAVRTTVRDPDNHKKVSHLTALQELGELEILAGDLTDEGSFDAPIAGCDLV
AEL79860.1_ANR_Malus_domestica                 LVKLLLQKGYAVRTTVRDPDNHKKVSHLTSLQELGELEILAGDLTDEGSFDAPIAGCDLV
AEL79861.1_ANR_Malus_domestica                 LVKLLLQKGYAVRTTVRDPDNHKKVSHLTALQELGELEILAGDLTDEGSFDAPIAGCDLV
ACY30421.1_C.BANa_Brassica_napus               LIDRLLQSGYEVNTTVRDPENVKKMAVLRVLQEQGDLNIFKADLTDEGSFNSPVSGCEYV
ACY30422.1_C.BANb_Brassica_napus               LIKHLLQSGYKVNTTVRDPENEKKMAHLKVLQELGDLKIFKADLTDEGSFTSPISGCEYV
ACY30423.1_A.BANa_Brassica_napus               LIDRLLQSGYEVNTTVRDPENEKKMAVLRVLQEQGDLNIFKADLTDEGSFNSPVSGCEYV
ACY30424.1_A.BANb_Brassica_napus               LIKHLLQSGYKVNTTVRDPENEKKMAHLKVLQELGDLKIFKADLTDEGSFTSPISGCEYV
ACY30425.1_C.BANa_Brassica_oleracea            LIDRLLQSGYEVNTTVRDPENVKKMAVLRVLQEQGDLNIFKADLTDEGSFNSPVSGCEYV
ABG76842.1_ANR_Fragaria_x_ananassa             LIKLLLEKGYAVRTTVRDPDNLKKISHLTALQELGELTIFRGDLTDEGSFDAAIAGSDLV
ABC71337.1_ANR-1_Lotus_corniculatus            LIKQLLEKGYAVNTTVRDPDNHKKISHLLVLQSLGELNIFGGELTVENDFDTPIEGSELV
ABC71332.1_ANR1-1_Lotus_corniculatus           LIKQLLEKGYAVNTTVRDPDNHKKISHLLALQSLGELNIFGGELTVENDFDTPIAGSELV
ABC71333.1_ANR1-2_Lotus_corniculatus           LIKQLLEKGYAVNTTVRDPDNHKKISHLLALQSWGELNIFEGELTVENDFDTPIAGSELV
ABC71335.1_ANR1-4_Lotus_corniculatus           LIKQLLEKGYAVNTTVRDPDNHKKISHLLALQSLGELNIFGGELTVENDFDTPIEGSELV
AAZ79363.1_ANR_Malus_domestica                 LVKLLLQKGYAVRTTVRDPDDRKKVSHLTALQELGELEIFAGDLTDEGSFDAPIAGCDLV
AAX12184.1_ANR_Malus_domestica                 LVKLLLQKGYAVRTTVRDPDNHKKVSHLTALQELGELEIFAGDLTDEGSFDAPIAGCDLV
AAZ17408.1_ANR_Malus_domestica                 LVKLLLQKGYAVRTTVRDPDNHKKVSHLTALQELGELEILAGDLTDEGSFDAPIAGCDLV
EEE86150.1_ANR_Populus_trichocarpa             LVKLLLEKGYAVNTTVRDPDNQKKVAHLIALQNLGDLNIFGADLTDEESFNAPIAGCELV
EEE97882.1_ANR_Populus_trichocarpa             LIKLLLEKGYAVNTTVRDPDNQKKIAHLIALQNLGDLNIFGADLTNEESFNAPIACCDLV
ABB77695.1_ANR_Pyrus_communis                  LVKLLLQKGYAVRTTVRDPDNHKKVSHLTALQELGELEILAGDLTDEGSFDAPIAGCDLV
BAD89742.1_ANR_Vitis_vinifera                  LVKLLLQKGYAVNTTVRDPDNQKKVSHLLELQELGDLKIFRADLTDELSFEAPIAGCDFV
AAZ82409.1_ANR_Vitis_vinifera                  LVKLLLQKGYAVNTTVRDPDNQKKVSHLLELQELGDLKIFRADLTDELSFEAPIAGCDFV
                                         acc
```

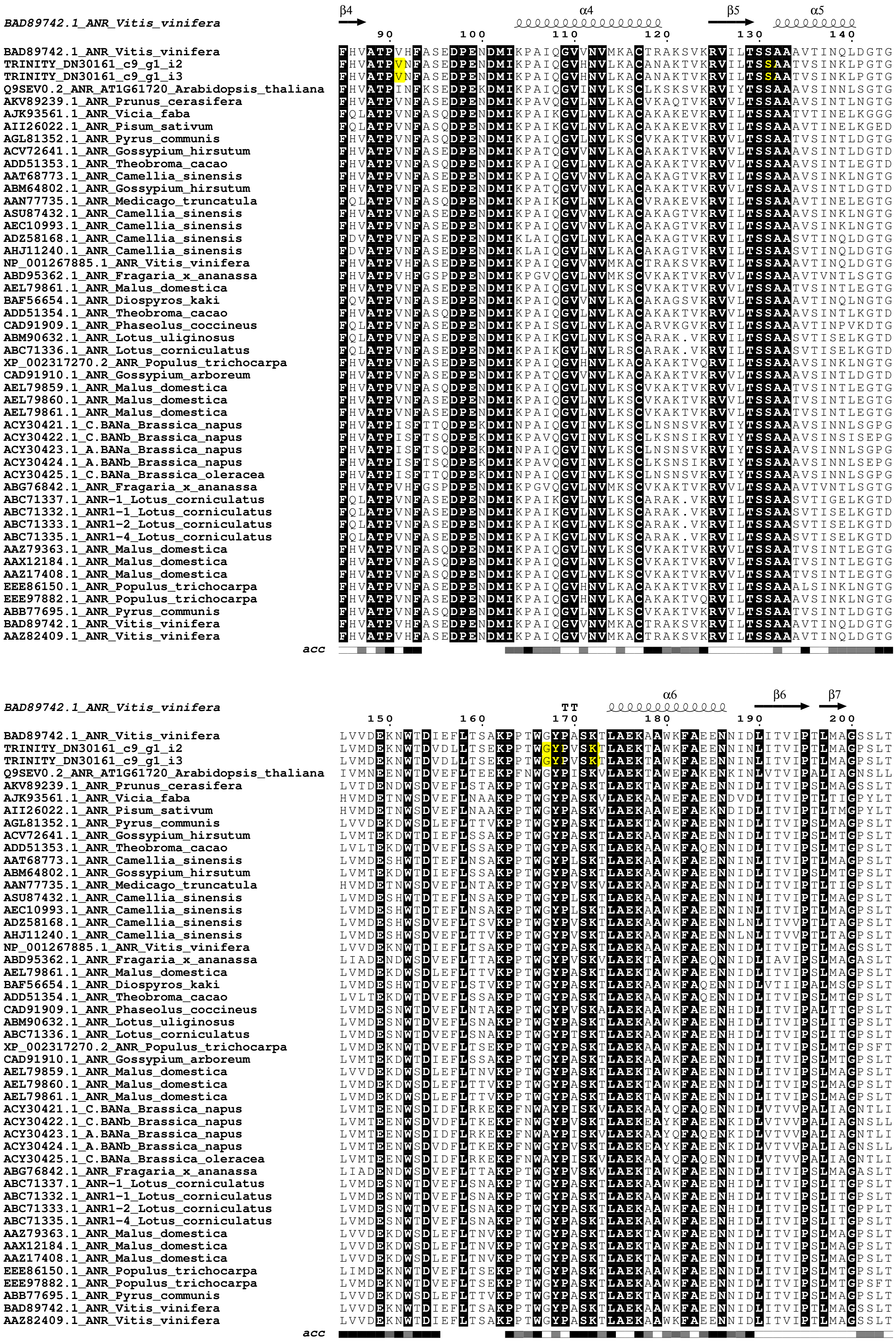

BAD89742.1_ANR_Vitis_vinifera
β4
α4
β5
α5
90 100 110 120 130 140
BAD89742.1_ANR_Vitis_vinifera FHVATPVHFASEDPENDMIKPAIQGVVNVMKACTRAKSVKRVILTSSAAAVTINQLDGTG
TRINITY_DN30161_c9_g1_i2 FHVATPVNFASEDPENDMIKPAIQGVHNVLKACANAKTVKRVILTSSAATVSINKLPGTG
TRINITY_DN30161_c9_g1_i3 FHVATPVNFASEDPENDMIKPAIQGVHNVLKACANAKTVKRVILTSSAATVSINKLPGTG
Q9SEV0.2_ANR_AT1G61720_Arabidopsis_thaliana FHVATPINFKSEDPEKDMIKPAIQGVINVLKSCLKSKSVKRVIYTSSAAAVSINNLSGTG
AKV89239.1_ANR_Prunus_cerasifera FHVATPVNFASEDPENDMIKPAVQGVLNVLKACVKAQTVKRVVLTSSAAAVSINTLNGTG
AJK93561.1_ANR_Vicia_faba FQLATPVNFASQDPENDMIKPAIKGVLNVLKACAKAKEVKRVILTSSAATVTINELKGGG
AII26022.1_ANR_Pisum_sativum FQLATPVNFASQDPENDMIKPAIKGVLNVLKACAKAKEVKRVILTSSAAAVTINELKGED
AGL81352.1_ANR_Pyrus_communis FHVATPVNFASQDPENDMIKPAIQGVLNVLKSCVKAKTVKRVVLTSSAATVSINTLDGTG
ACV72641.1_ANR_Gossypium_hirsutum FHVATPVNFASEDPENDMIKPATQGVVNVLKACAKAKTVKRVVLTSSAAAVSINTLDGTD
ADD51353.1_ANR_Theobroma_cacao FHVATPVNFASQDPENDMIKPAIQGVLNVLKACAKAKTVKRVVLTSSAAAVSINTLEGTD
AAT68773.1_ANR_Camellia_sinensis FHVATPVNFASEDPENDMIKPAIQGVVNVLKACAKAGTVKRVILTSSAAAVSINKLNGTG
ABM64802.1_ANR_Gossypium_hirsutum FHVATPVNFASEDPENDMIKPATQGVVNVLKACAKAKTVKRVVLTSSAAAVSINTLDGTD
AAN77735.1_ANR_Medicago_truncatula FQLATPVNFASQDPENDMIKPAIKGVLNVLKACVRAKEVKRVILTSSAAAVTINELEGTG
ASU87432.1_ANR_Camellia_sinensis FHVATPVNFASEDPENDMIKPAIQGVVNVLKACAKAGTVKRVILTSSAAAVSINKLNGTG
AEC10993.1_ANR_Camellia_sinensis FHVATPVNFASEDPENDMIKPAIQGVVNVLKACAKAGTVKRVILTSSAAAVSINKLNGTG
ADZ58168.1_ANR_Camellia_sinensis FDVATPVNFASEDPENDMIKLAIQGVLNVLKACAKAGTVKRVILTSSAASVTINQLDGTG
AHJ11240.1_ANR_Camellia_sinensis FDVATPVNFASEDPENDMIKLAIQGVLNVLKACAKAGTVKRVILTSSAASVTINQLDGTG
NP_001267885.1_ANR_Vitis_vinifera FHVATPVHFASEDPENDMIKPAIQGVVNVMKACTRAKSVKRVILTSSAAAVTINQLDGTG
ABD95362.1_ANR_Fragaria_x_ananassa FHVATPVHFGSPDPENDMIKPGVQGVLNVMKSCVKAKTVKRVVLTSSAAAVTVNTLSGTG
AEL79861.1_ANR_Malus_domestica FHVATPVNFASEDPENDMIKPAIQGVLNVLKSCVKAKTVKRVVLTSSAATVSINTLEGTG
BAF56654.1_ANR_Diospyros_kaki FQVATPVNFASEDPENDMIKPAIQGVVNVLKACAKAGSVKRVVLTSSAAAVTINQLNGTG
ADD51354.1_ANR_Theobroma_cacao FHVATPVNFASQDPENDMIKPAIQGVLNVLKACAKAKTVKRVVLTSSAAAVSINTLKGTD
CAD91909.1_ANR_Phaseolus_coccineus FQLATPVNFASEDPENDMIKPAISGVLNVLKACARVKGVKRVILTSSAAAVTINPVKDTG
ABM90632.1_ANR_Lotus_uliginosus FQLATPVNFASEDPENDMIKPAIKGVLNVLKSCARAK.VKRVILTSSAASVTISELKGTD
ABC71336.1_ANR_Lotus_corniculatus FQLATPVNFASEDPENDMIKPAIKGVLNVLRSCARAK.VKRVILTSSAASVTISELKGTD
XP_002317270.2_ANR_Populus_trichocarpa FHVATPVNFASEDPENDMIKPAIQGVHNVLKACAKAKTVQRVILTSSAAAVSINKLNGTG
CAD91910.1_ANR_Gossypium_arboreum FHVATPVNFASEDPENDMIKPATQGVVNVLKACAKAKTVKRVVLTSSAAAVSINTLDGTD
AEL79859.1_ANR_Malus_domestica FHVATPVNFASQDPENDMIKPAIQGVLNVLKSCVKAKTVKRVVLTSSAATVSINTLEGTG
AEL79860.1_ANR_Malus_domestica FHVATPVNFASEDPENDMIKPAIQGVLNVLKSCVKAKTVKRVVLTSSAATVSINTLEGTG
AEL79861.1_ANR_Malus_domestica FHVATPVNFASEDPENDMIKPAIQGVLNVLKSCVKAKTVKRVVLTSSAATVSINTLEGTG
ACY30421.1_C.BANa_Brassica_napus FHVATPISFTTQDPEKDMINPAIQGVINVLKSCLNSNSVKRVIYTSSAAAVSINNLSGPG
ACY30422.1_C.BANb_Brassica_napus FHVATPISFTSQDPEKDMIKPAVQGVINVLKSCLKSNSIKRVIYTSSAAAVSINSISEPG
ACY30423.1_A.BANa_Brassica_napus FHVATPISFTSQDPEKDMINPAIQGVINVLKSCLNSNSVKRVIYTSSAAAVSINNLSGPG
ACY30424.1_A.BANb_Brassica_napus FHVATPISFTSQDPEKDMIKPAVQGVINVLKSCLKSNSIKRVIYTSSAAAVSINNLSEPG
ACY30425.1_C.BANa_Brassica_oleracea FHVATPISFTTQDPEKDMINPAIQGVINVLKSCLNSNSVKRVIYTSSAAAVSINNLSGPG
ABG76842.1_ANR_Fragaria_x_ananassa FHVATPVHFGSPDPENDMIKPGVQGVLNVMKSCVKAKTVKRVVLTSSAAAVTVNTLSGTG
ABC71337.1_ANR-1_Lotus_corniculatus FQLATPVNFASEDPENDMIKPAIKGVLNVLKSCARAK.VKRVILTSSAASVTIGELKGTD
ABC71332.1_ANR1-1_Lotus_corniculatus FQLATPVNFASEDPENDMIKPAIKGVLNVLKSCARAK.VKRVILTSSAASVTISELKGTD
ABC71333.1_ANR1-2_Lotus_corniculatus FQLATPVNFASEDPENDMIKPAIKGVLNVLKSCARAK.VKRVILTSSAASVTISELKGTD
ABC71335.1_ANR1-4_Lotus_corniculatus FQLATPVNFASEDPENDMIKPAIKGVLNVLKSCARAK.VKRVILTSSAASVTISELKGTD
AAZ79363.1_ANR_Malus_domestica FHVATPVNFASQDPENDMIKPAIQGVLNVLKSCVKAKTVKRVVLTSSAATVSINTLEGTG
AAX12184.1_ANR_Malus_domestica FHVATPVNFASQDPENDMIKPAIQGVLNVLKSCVKAKTVKRVVLTSSAATVSINTLEGTG
AAZ17408.1_ANR_Malus_domestica FHVATPVNFASQDPENDMIKPAIQGVLNVLKSCVKAKTVKRVVLTSSAATVSINTLEGTG
EEE86150.1_ANR_Populus_trichocarpa FHVATPVNFASEDPENDMIKPAIQGVHNVLKACAKAKTVKRVILTSSAAALSINKLNGTG
EEE97882.1_ANR_Populus_trichocarpa FHVATPVNFASEDPENDMIKPAIQGVHNVLKACAKAKTVQRVILTSSAAAVSINKLNGTG
ABB77695.1_ANR_Pyrus_communis FHVATPVNFASQDPENDMIKPAIQGVLNVLKSCVKAKTVKRVVLTSSAATVSINTLDGTG
BAD89742.1_ANR_Vitis_vinifera FHVATPVHFASEDPENDMIKPAIQGVVNVMKACTRAKSVKRVILTSSAAAVTINQLDGTG
AAZ82409.1_ANR_Vitis_vinifera FHVATPVHFASEDPENDMIKPAIQGVVNVMKACTRAKSVKRVILTSSAAAVTINQLDGTG
acc
BAD89742.1_ANR_Vitis_vinifera
TT
α6
β6
β7
150 160 170 180 190 200
BAD89742.1_ANR_Vitis_vinifera LVVDEKNWTDIEFLTSAKPPTWGYPASKTLAEKAAWKFAEENNIDLITVIPTLMAGSSLT
TRINITY_DN30161_c9_g1_i2 LVMDEKNWTDVDLLTSEKPPTWGYPVSKTLAEKTAWKFAEENNINLITVIPTLMAGPSLT
TRINITY_DN30161_c9_g1_i3 LVMDEKNWTDVDLLTSEKPPTWGYPVSKTLAEKTAWKFAEENNINLITVIPTLMAGPSLT
Q9SEV0.2_ANR_AT1G61720_Arabidopsis_thaliana IVMNEENWTDVEFLTEEKPFNWGYPISKVLAEKTAWEFAKENKINLVTVIPALIAGNSLL
AKV89239.1_ANR_Prunus_cerasifera LVTDENDWSDVEFLSTAKPPTWGYPASKTLAEKTAWKFAEENNIDLITVIPSLMAGSSLT
AJK93561.1_ANR_Vicia_faba HVMDETNWSDVEFLNAAKPPTWGYPASKVLAEKAAWEFAEENDVDLITVIPTLTIGPYLT
AII26022.1_ANR_Pisum_sativum HVMDETNWSDVEFLNAAKPPTWGYPASKVLAEKAAWEFAEKNDIDLITVIPTLTMGPYLT
AGL81352.1_ANR_Pyrus_communis LVVDEKDWSDLEFLTTVKPPTWGYPASKTLAEKTAWKFAEENNIDLITVIPSLMAGPSLT
ACV72641.1_ANR_Gossypium_hirsutum LVMTEKDWTDIEFLSSAKPPTWGYPASKTLAEKAAWKFAEENNIDLITVIPSLMTGPSLT
ADD51353.1_ANR_Theobroma_cacao LVLTEKDWTDVEFLSSAKPPTWGYPASKTLAEKAAWKFAQENNIDLITVIPSLMTGPSLT
AAT68773.1_ANR_Camellia_sinensis LVMDESHWTDTEFLNSAKPPTWGYPLSKTLAEKAAWKFAEENNINLITVIPTLMAGPSLT
ABM64802.1_ANR_Gossypium_hirsutum LVMTEKDWTDIEFLSSAKPPTWGYPASKTLAEKAAWKFAEENNIDLITVIPSLMTGPSLT
AAN77735.1_ANR_Medicago_truncatula HVMDETNWSDVEFLNTAKPPTWGYPVSKVLAEKAAWKFAEENNIDLITVIPTLTIGPSLT
ASU87432.1_ANR_Camellia_sinensis LVMDESHWTDTEFLNSAKPPTWGYPLSKTLAEKAAWKFAEENNINLITVIPTLMAGPSLT
AEC10993.1_ANR_Camellia_sinensis LVMDESHWTDTEFLNSAKPPTWGYPLSKTLAEKAAWKFAEENNINLITVIPTLMAGPSLT
ADZ58168.1_ANR_Camellia_sinensis LVMDESHWSDVEFLTSVKPPTWGYPVSKTLAEKAAWKFAEENNLNLITVVPTLTAGPSLT
AHJ11240.1_ANR_Camellia_sinensis LVMDESHWSDVEFLTTVKPPTWGYPVSKTLAEKAAWKFAEENNLNLITVVPTLTAGPSLT
NP_001267885.1_ANR_Vitis_vinifera LVVDEKNWTDIEFLTSAKPPTWGYPASKTLAEKAAWKFAEENNIDLITVIPTLMAGSSLT
ABD95362.1_ANR_Fragaria_x_ananassa LIADENDWSDVEFLTTAKPPTWGYPVSKVLAEKTAWKFAEQNNIDLIAVIPSLMAGASLT
AEL79861.1_ANR_Malus_domestica LVMDEKDWSDLEFLTTVKPPTWGYPASKTLAEKTAWKFAEENNIDLITVIPSLMAGPSLT
BAF56654.1_ANR_Diospyros_kaki LVMDESHWTDVEFLSTVKPPTWGYPASKTLAEKAAWKFAQENNIDLVTIIPALMSGPSLT
ADD51354.1_ANR_Theobroma_cacao LVLTEKDWTDVEFLSSAKPPTWGYPASKTLAEKAAWKFAQENNIDLITVIPSLMTGPSLT
CAD91909.1_ANR_Phaseolus_coccineus LVMDESNWTDVEFLNTAKPPTWGYPVSKALAEKAAWKFAEENHIDLITVIPALTTGPSLT
ABM90632.1_ANR_Lotus_uliginosus LVMDESNWTDVEFLSNAKPPTWGYPASKTLAEKAAWKFAEENHIDLITVIPSLITGPSLT
ABC71336.1_ANR_Lotus_corniculatus LVMDESNWTDVEFLSNAKPPTWGYPTSKTLAEKAAWKFAEENHIDLITVIPSLITGPSLT
XP_002317270.2_ANR_Populus_trichocarpa LVMDEKNWTDVEFLTSEKPPTWGYPASKTLAEKAAWKFAEENNIDLITVIPSLMTGPSFT
CAD91910.1_ANR_Gossypium_arboreum LVMTEKDWTDIEFLSSAKPPTWGYPASKTLAEKAAWKFAEENNIDLITVIPSLMTGPSLT
AEL79859.1_ANR_Malus_domestica LVVDEKDWSDLEFLTNVKPPTWGYPASKTLAEKTAWKFAEENNIDLITVIPSLMAGPSLT
AEL79860.1_ANR_Malus_domestica LVMDEKDWSDLEFLTTVKPPTWGYPASKTLAEKTAWKFAEENNIDLITVIPSLMAGPSLT
AEL79861.1_ANR_Malus_domestica LVMDEKDWSDLEFLTTVKPPTWGYPASKTLAEKTAWKFAEENNIDLITVIPSLMAGPSLT
ACY30421.1_C.BANa_Brassica_napus LVMTEENWSDIDFLRKEKPFNWAYPISKVLAEKAAYQFAQENNIDLVTVVPALIAGNTLI
ACY30422.1_C.BANb_Brassica_napus LVMTEENWSDVDFLTKEKPFNWGYPVSKTLAEKEAYKFAEENKIDLVTVVPALIAGNSLL
ACY30423.1_A.BANa_Brassica_napus LVMTEENWSDIDFLRKEKPFNWAYPISKVLAEKAAYQFAQENKIDLVTVVPALIAGNTLI
ACY30424.1_A.BANb_Brassica_napus LVMTEENWSDVDFLTKEKPFNWGYPVSKTLAEKEAYKFAEENKIDLVTVVPALIAGNSLL
ACY30425.1_C.BANa_Brassica_oleracea LVMTEENWSDIDFLRKEKPFNWAYPISKVLAEKAAYQFAQENNIDLVTVVPALIAGNTLI
ABG76842.1_ANR_Fragaria_x_ananassa LIADENDWSDVEFLTTAKPPTWGYPVSKVLAEKTAWKFAEENNIDLITVIPSLMAGASLT
ABC71337.1_ANR-1_Lotus_corniculatus LVMDESNWTDVEFLSNAKPPTWGYPASKTLAEKAAWKFAEENHIDLITVIPSLITGPSLT
ABC71332.1_ANR1-1_Lotus_corniculatus LVMDESNWTDVEFLSNAKPPTWGYPASKTLAEKAAWKFAEENHIDLITVIPSLITGPSLT
ABC71333.1_ANR1-2_Lotus_corniculatus LVMDESNWTDVEFLSNAKPPTWGYPASKTLAEKAAWKFAEENHIDLITVIPSLITGPPLT
ABC71335.1_ANR1-4_Lotus_corniculatus LVMDESNWTDVEFLSNAKPPTWGYPASKTLAEKAAWKFAEENHIDLITVIPSLITGPSLT
AAZ79363.1_ANR_Malus_domestica LVVDEKDWSDLEFLTNVKPPTWGYPASKTLAEKTAWKFAEENNIDLITVIPSLMAGPSLT
AAX12184.1_ANR_Malus_domestica LVVDEKDWSDLEFLTNVKPPTWGYPASKTLAEKTAWKFAEENNIDLITVIPSLMAGPSLT
AAZ17408.1_ANR_Malus_domestica LVVDEKDWSDLEFLTNVKPPTWGYPASKTLAEKTAWKFAEENNIDLITVIPSLMAGPSLT
EEE86150.1_ANR_Populus_trichocarpa LIMDEKNWTDVEFLTSEKPPTWGYPASKTLAEKAAWKFAEENNIDLITVIPSLMTGPSLT
EEE97882.1_ANR_Populus_trichocarpa LVMDEKNWTDVEFLTSEKPPTWGYPASKTLAEKAAWKFAEENNIDLITVIPSLMTGPSFT
ABB77695.1_ANR_Pyrus_communis LVVDEKDWSDLEFLTTVKPPTWGYPASKTLAEKTAWKFAEENNIDLITVIPSLMAGPSLT
BAD89742.1_ANR_Vitis_vinifera LVVDEKNWTDIEFLTSAKPPTWGYPASKTLAEKAAWKFAEENNIDLITVIPTLMAGSSLT
AAZ82409.1_ANR_Vitis_vinifera LVVDEKNWTDIEFLTSAKPPTWGYPASKTLAEKAAWKFAEENNIDLITVIPTLMAGSSLT
acc

```
                                              α7          α8          β8    α9
BAD89742.1_ANR_Vitis_vinifera
                                              210       220       230       240       250       260
BAD89742.1_ANR_Vitis_vinifera                 SD.VPSSIGLAMSLITGNEFLIN.GMKGMQMLSGSVSIAHVEDVCRAHIFVAEKESASGR
TRINITY_DN30161_c9_g1_i2                      SD.IPSSIHLATSLITGNEFLIN.GLKGMQILSGSISITHVEDVCRAHVFLGEKESASGR
TRINITY_DN30161_c9_g1_i3                      SD.IPSSIHLATSLITGTH...........LSASID.......................
Q9SEV0.2_ANR_AT1G61720_Arabidopsis_thaliana   SD.PPSSLSLSMSFITGKEMHVT.GLKEMQKLSGSISFVHVDDLARAHLFLAEKETASGR
AKV89239.1_ANR_Prunus_cerasifera              PD.VPSSIGLAMSLITGNDFLINHALKGMQLLSGSISITHVEDVCRAHIFLAEKESASGR
AJK93561.1_ANR_Vicia_faba                     QD.MPSSVGLAMSLITGNDFLVN.ALKGMQFLSGSISITHVEDICRAHIFVAEKESASGR
AII26022.1_ANR_Pisum_sativum                  QD.MPSSVGLAMSLITGNDFLVN.ALKGMQFLSGSISITHVEDICRAHIFVAEKESASGR
AGL81352.1_ANR_Pyrus_communis                 PD.VPSSIGLAMALITGDDFLINMALKGMQMLSGSISITHVEDVCRAHIFLAEKESASGR
ACV72641.1_ANR_Gossypium_hirsutum             PI.VPSSIGLATSLISGNEFLIN.ALKGMQMLSGSISITHVEDECRAHVFLAEKESASGR
ADD51353.1_ANR_Theobroma_cacao                PD.VPSSIGLATSLLSGNEFLVN.ALKGMQMLSGSISITHVEDVCRAHVFLAEKESGSGR
AAT68773.1_ANR_Camellia_sinensis              AD.VPSSIGLAMSLITGNEFLIN.GLKGMQMLSGSISISHVEDVCRAHVFVAEKESASGR
ABM64802.1_ANR_Gossypium_hirsutum             PI.VPSSIGLATSLISGNEFLIN.ALKGMQMLSGSISITHVEDVCRAHVFLAEKESASGR
AAN77735.1_ANR_Medicago_truncatula            QD.IPSSVAMGMSLLTGNDFLIN.ALKGMQFLSGSISITHVEDICRAHIFVAEKESTSGR
ASU87432.1_ANR_Camellia_sinensis              AD.VPSSIGLAMSLITGNEFLIN.GLKGMQMLSGSISISHVEDVCRAHVFVAEKESASGR
AEC10993.1_ANR_Camellia_sinensis              AD.VPSSIGLAMSLITGNEFFIN.GLKGMQMLSGSISISHVEDVCRAHVFVAEKESAFGR
ADZ58168.1_ANR_Camellia_sinensis              SE.VPNSIELAMSLITGNEFLID.GLKGMQILSGSISITHVEDVCGAHIFVAEKESASGR
AHJ11240.1_ANR_Camellia_sinensis              SE.VPNSIELAMSLMTGNEFLID.GLKGMQILSGSISITHVEDVCGAHIFVAEKESASGR
NP_001267885.1_ANR_Vitis_vinifera             SD.VPSSIGLAMSLITGNEFLIN.GMKGMQMLSGSVSIAHVEDVCQAHIFVAEKESASGR
ABD95362.1_ANR_Fragaria_x_ananassa            PD.IPSSIGLATSLITGNEFLIN.GLKGMQMLSGSISITHVEDVCRAHIFLAEKESASGR
AEL79861.1_ANR_Malus_domestica                PD.VPSSIGLAMSLITGNDFLINMALKGMQMLSGSISISHVEDVCRAHIFLAEKESASGR
BAF56654.1_ANR_Diospyros_kaki                 VD.PPSSMGLAMSLITGNEFLIN.ALKGMQMLSGSISIAHVEDVCRAHIFVAEKKSASGR
ADD51354.1_ANR_Theobroma_cacao                PD.VPSSIGLATSLLSGNEFLVN.ALKGMQMLSGSISITHVEDVCRAHVFLAEKESASGR
CAD91909.1_ANR_Phaseolus_coccineus            PD.IPSSVGLATSLITGNDFLIN.ALKGMQFLSGSISITHVEDICRAHIYVAENESSSGR
ABM90632.1_ANR_Lotus_uliginosus               PD.IPSSVGLATSLITGNDFLIN.AMKGMQLLSGSISITHVEDVCRAHIFVAEKESASGR
ABC71336.1_ANR_Lotus_corniculatus             PD.IPSSVGLATSLITGNDFLIN.AMKGMQLLSGSISITHVEDVCRAHIFVAEKQSASGR
XP_002317270.2_ANR_Populus_trichocarpa        PH.IPDSINLAMSLITGNKFLIN.GLKGMQMLSGSISITHVEDVCRAHIFLAEKESASGR
CAD91910.1_ANR_Gossypium_arboreum             PI.VPSSIGLATSLISGNEFLIN.ALKGMQMLSGSISITHVEDVCRAHVFLAEKESASGR
AEL79859.1_ANR_Malus_domestica                PD.VPSSIGLAMALITGDDFLINMALKGMQMLSGSISIAHVEDVCRAHIFLAEKESASGR
AEL79860.1_ANR_Malus_domestica                PD.VPSSIGLAMSLITGNDFLINMALKGMQMLSGSISISHVEDVCRAHIFLAEKESASGR
AEL79861.1_ANR_Malus_domestica                PD.VPSSIGLAMSLITGNDFLINMALKGMQMLSGSISISHVEDVCRAHIFLAEKESASGR
ACY30421.1_C.BANa_Brassica_napus              DDPPPSSLSLSMSLITRKEMHLN.ALKEMQKLSGSISFIHVDDLACAHLFLAEKETASGR
ACY30422.1_C.BANb_Brassica_napus              SD.PPSSLSLSMSLITGKEMHLS.GLKEMQKLSGSISFIHVDDLARAHMFLAEKETASGR
ACY30423.1_A.BANa_Brassica_napus              DDPPPSSLSLSMSLITRKEMHLN.ALKEMQKLSGSISFIHVDDLACAHLFLAEKETASGR
ACY30424.1_A.BANb_Brassica_napus              SD.PPSSLSLSMSLITGKEMHLS.GLKEMQKLSGSISFIHVDDLARAHMFLAEKETASGR
ACY30425.1_C.BANa_Brassica_oleracea           DDPPPSSLSLSMSLITRKEMHLN.ALKEMQKLSGSISFIHVDDLACAHLFLAEKETASGR
ABG76842.1_ANR_Fragaria_x_ananassa            PD.IPSSIGLATSLITGNEFLIN.GLKGMQMLSGSISITHVEDVCRAHIFLAEKESASGR
ABC71337.1_ANR-1_Lotus_corniculatus           PD.IPSSVGLATSLITGNDFLIN.AMKGMQLLSGSISITHVEDVCRAHIFVAEKQSASGR
ABC71332.1_ANR1-1_Lotus_corniculatus          PD.IPSSVGLATSLITGNDFLIN.AMKGMQLLSGSISITHVEDVCRAHIFVAEKQSASGR
ABC71333.1_ANR1-2_Lotus_corniculatus          PD.IPSSVGLATSLITGNDFLIN.AMKGMQLLSGSISITHVEDVCRAHIFVAEKQSASGR
ABC71335.1_ANR1-4_Lotus_corniculatus          PD.IPSSVGLATSLITGNDFLIN.AMKGMQLLSGSISITHVEDVCRAHIFVAEKQSASGR
AAZ79363.1_ANR_Malus_domestica                PD.VPSSIGLAMALITGDDFLINMALKGMQMLSGSISIAHVEDVCRAHIFLAEKESASGR
AAX12184.1_ANR_Malus_domestica                PD.VPSSIGLAMALITGDDFLINMALKGMQMLSGSISIAHVEDVCRAHIFLAEKESASGR
AAZ17408.1_ANR_Malus_domestica                PD.VPSSIGLAMALITGDDFLINMALKGMQMLSGSISIAHVEDVCRAHIFLAEKESASGR
EEE86150.1_ANR_Populus_trichocarpa            LD.IPSSVHLSMSLITGNEFLKN.ALKGMQMLSGSISITHVEDVCRAHIFLAEKESASGR
EEE97882.1_ANR_Populus_trichocarpa            PH.IPDSINLAMSLITGNKFLIN.GLKGMQMLSGSISITHVEDVCRAHIFLAEKESASGR
ABB77695.1_ANR_Pyrus_communis                 PD.VPSSIGLAMALITGDDFLINMALKGMQMLSGSISITHVEDVCRAHIFLAEKESASGR
BAD89742.1_ANR_Vitis_vinifera                 SD.VPSSIGLAMSLITGNEFLIN.GMKGMQMLSGSVSIAHVEDVCRAHIFVAEKESASGR
AAZ82409.1_ANR_Vitis_vinifera                 SD.VPSSIGLAMSLITGNEFLIN.GMKGMQMLSGSVSIAHVEDVCQAHIFVAEKESASGR
                                        acc

                                              β9  β10   α10                    TT                α11
BAD89742.1_ANR_Vitis_vinifera
                                                   270       280       290       300       310       320
BAD89742.1_ANR_Vitis_vinifera                 YICCAANTSVPELAKFLSKRYPQYKVPTDFGDFPPKSKLIISSEKLVKEGFSFKYGIEEI
TRINITY_DN30161_c9_g1_i2                      YICCGVNTSVPELAKFLNQRYPNYKVPIEFGDFPSKAKLVLSSEKLIGEGFSFKYGIEEI
TRINITY_DN30161_c9_g1_i3                      ............................................................
Q9SEV0.2_ANR_AT1G61720_Arabidopsis_thaliana   YICCAYNTSVPEIADFLIQRYPKYNVLSEFEEGLSIPKLTLSSQKLINEGFRFEYGINEM
AKV89239.1_ANR_Prunus_cerasifera              YICCAVNTSVPELAKFLNERYPEYKVPTEFGDFPSKAKLILSSEKLIKEGFDFKYDIEQI
AJK93561.1_ANR_Vicia_faba                     YICCAHNTSVPELAKFLSKRYPQYKVPTEFEDCPSKAKLIISSEKLIKEGFSFKYGIVET
AII26022.1_ANR_Pisum_sativum                  YICCAQNTCVPELAKFFSKRYPQYKVPTKFEDCPSKAKLIIFFEKLIKEGFGFKYGIGET
AGL81352.1_ANR_Pyrus_communis                 YICCAANTGVPELAKFLNKRYPQYKVPTEFGDFPSKAKLIISSEKLIKEGFDFKYGIEEI
ACV72641.1_ANR_Gossypium_hirsutum             YICSAVNTSVPELAKFLNKRYPDFKVPTDFGDFPSKPKLIISSEKLISEGFSFKYGIEEI
ADD51353.1_ANR_Theobroma_cacao                YICCAVNSSVPELAKFLNQRYPEFKVPTDFGDFPSKAKLIISSDKLINEGFSFKFGIEEI
AAT68773.1_ANR_Camellia_sinensis              YICCAVSTSVPELAKFLNKRYPEYNVPTDFGDFPSKAKLILSSEKLTKEGFSFKYGIEEI
ABM64802.1_ANR_Gossypium_hirsutum             YICSAVNTSVPELAKFLNKRYPDFKVPTDFGDFPSKPKLIISSEKLISEGFSFKYGIEEI
AAN77735.1_ANR_Medicago_truncatula            YICCAHNTSVPELAKFLSKRYPQYKVPTEFDDFPSKAKLIISSGKLIKEGFSFKHSIAET
ASU87432.1_ANR_Camellia_sinensis              YICCAVSTSVPELAKFLNKRYPEYNVPTDFGDFPSKAKLILSSEKLTKEGFSFKYGIEEI
AEC10993.1_ANR_Camellia_sinensis              YICCAVNTSVPELAKFLNKRYPEYNVPTDFGDFPSKAKLILSFEKLTKEGFSFKYGIEEI
ADZ58168.1_ANR_Camellia_sinensis              YICCGVNSSVPELARFLNKRYPQYNVPTDFGDLPSKAKLIISSEKLIKEGFSFKYGIEEI
AHJ11240.1_ANR_Camellia_sinensis              YICCSVNSSVPELARFLNKRYPQYNVPTDFGDLPSKAKLIISSEKLIKEGFSFKYGIEEI
NP_001267885.1_ANR_Vitis_vinifera             YICCAANTSVPELAKFLSKRYPQYKVPTDFGDFPPKSKLIISSEKLVKEGFSFKYGIEEI
ABD95362.1_ANR_Fragaria_x_ananassa            YICCAENSSVPEVAKFLSKRYPEYKVPTEFGDFPSKAKTILPSEKLKKEGFTFKFGIEDI
AEL79861.1_ANR_Malus_domestica                YICCAANTSVPELARFLNKRYPQYKVPTEFGDFPSKAKLIISSEKLIQEGFDFKYGIEEI
BAF56654.1_ANR_Diospyros_kaki                 YICCAVNTSVPELAKFLSKKYPQYNVPTDFGDFPSKAKLVLSSDKLTKEGFSFQYGMEDI
ADD51354.1_ANR_Theobroma_cacao                YICCAVNSSVPELAKFLNQRYPEFKVPTDFGDFPSKAKLIISSDKLINEGFSFKFGIEEI
CAD91909.1_ANR_Phaseolus_coccineus            YIVSAHSTSVPELAKFLSKRYPQYKVPTEFDDCPSKAKLTISSEKLVKEGFSFKYGIEEI
ABM90632.1_ANR_Lotus_uliginosus               YICCAHNTSVPELAKFLNKRYPQYKVSTEFNDFPAKAKLIISPEKLIKEGFSFKYGVEEI
ABC71336.1_ANR_Lotus_corniculatus             YICCAHNTSVPELAKFLNKRYPQYKVSTEFNDFPAKAKLIISPEKLIKEGFSFKYGVEEI
XP_002317270.2_ANR_Populus_trichocarpa        YICCGVNTSVVELAKFLNKRYPQYQVPTDCGDFPSEAKLIITSEKLSSEGFSFKYGIEET
CAD91910.1_ANR_Gossypium_arboreum             YICSAVNTSVPELAKFLNKRYPDFKVPTDFGDFPSKPKLIISSEKLISEGFSFKYGIEEI
AEL79859.1_ANR_Malus_domestica                YICCAANTGVPELAKFLNKRYPQYKVPTEFGDFPSEAKLIISSEKLIKEGFDFKYGIEEI
AEL79860.1_ANR_Malus_domestica                YICCAANTSVPELARFLNKRYPQYKVPTEFGDFPSKAKLIISSEKLIKEGFDFKYGIEEI
AEL79861.1_ANR_Malus_domestica                YICCAANTSVPELARFLNKRYPQYKVPTEFGDFPSKAKLIISSEKLIQEGFDFKYGIEEI
ACY30421.1_C.BANa_Brassica_napus              YICCSYNTNIPELADFLRKRYPRYNVLSEFEECLATAKLTLSSEKLINEGFEFEHDIDEM
ACY30422.1_C.BANb_Brassica_napus              YICCYYNTNVPEIADFLRRRYPMYNVLSEFEEGLSSAKLTLSSEKLIKEGFRFEYGISEM
ACY30423.1_A.BANa_Brassica_napus              YICCSYNTNIPELADFLRKRYPRYNVLSEFEECLSTAKLTLSSEKLINEGFQFEHDVEEM
ACY30424.1_A.BANb_Brassica_napus              YICCYYNTNVPEIADFLRRRYTKYNVLSEFEECLSSAKLTLSSEKLIKEGFRFEYGISEM
ACY30425.1_C.BANa_Brassica_oleracea           YICCSYNTSIPELADFLRKRYPRYNVLSEFEECLAAAKLTLSSEKLINEGFEFEHDIDEM
ABG76842.1_ANR_Fragaria_x_ananassa            YICCAENSSVPEVAKFLSKRYPEYKVPTEFGDFPSKAKTILSSEKLKKEGFTFKYGIEDI
ABC71337.1_ANR-1_Lotus_corniculatus           YICCAHNTSVPELAKFLNKRYPQYKVSTEFNDFPAKAKLIISPEKLIKEGFSFKYGVEEI
ABC71332.1_ANR1-1_Lotus_corniculatus          YICCAHNTSVPELAKFLYKRYPQYKVSTEFNDFPAKAKLIISPEKLIKEGFSFKYGVEEI
ABC71333.1_ANR1-2_Lotus_corniculatus          YICCAHNTSVPELAKFLNKRYPQYKVSTEFNDFPAKAKLIISPEKLIKEGFSFKYGVEEI
ABC71335.1_ANR1-4_Lotus_corniculatus          YICCAHNTSVPELAKFLNKRYPQYKVSTEFNDFPAKAKLIISPEKLIKEGFSFKYGVEEI
AAZ79363.1_ANR_Malus_domestica                YICCAANTGVPELAKFLNKRYPQYKVPTEFGDFPSEAKLIISSEKLIKEGFDFKYGIEEI
AAX12184.1_ANR_Malus_domestica                YICCAANTGVPELAKFLNKRYPQYKVPTEFGDFPSEAKLIISSEKLIKEGFDFKYGIEEI
AAZ17408.1_ANR_Malus_domestica                YICCAANTGVPELAKFLNKRYPQYKVPTEFGDFPSEAKLIISSEKLIKEGFDFKYGIEEI
EEE86150.1_ANR_Populus_trichocarpa            YICCAVNTSVVELAEFLNKRYPQYQVPTDFGDFPSKAKLAITSEKLISEGFSFKYGIEEV
EEE97882.1_ANR_Populus_trichocarpa            YICCGVNTSVVELAKFLNKRYPQYQVPTDCGDFPSEAKLIITSEKLSSEGFSFKYGIEEI
ABB77695.1_ANR_Pyrus_communis                 YICCAANAGVPELAKFLNKRYPQYKVPTEFGDFPSKAKLIISSEKLIKEGFDFKYGIEEI
BAD89742.1_ANR_Vitis_vinifera                 YICCAANTSVPELAKFLSKRYPQYKVPTDFGDFPPKSKLIISSEKLVKEGFSFKYGIEEI
AAZ82409.1_ANR_Vitis_vinifera                 YICCAANTSVPELAKFLSKRYPQYKVPTDFGDFPPKSKLIISSEKLVKEGFSFKYGIEEI
                                        acc
```

| | |
|---|---|
| *BAD89742.1_ANR_Vitis_vinifera* | α12 |
| | 330 |
| **BAD89742.1_ANR_Vitis_vinifera** | YDESVEYFKAKGLLQN. |
| **TRINITY_DN30161_c9_g1_i2** | YDQTVEYLKGKGILK.. |
| **TRINITY_DN30161_c9_g1_i3** | ................. |
| **Q9SEV0.2_ANR_AT1G61720_Arabidopsis_thaliana** | YDQMIEYFESKGLIKAK |
| **AKV89239.1_ANR_Prunus_cerasifera** | YDQAVDYFKAKGLLQN. |
| **AJK93561.1_ANR_Vicia_faba** | FDHTVEYLKTKGILKE. |
| **AII26022.1_ANR_Pisum_sativum** | YAQTIEYLKIKGVLKK. |
| **AGL81352.1_ANR_Pyrus_communis** | YDQTVEYFKAKGLLQN. |
| **ACV72641.1_ANR_Gossypium_hirsutum** | YDQTVEYLKSKGLLK.. |
| **ADD51353.1_ANR_Theobroma_cacao** | YDQTVEYMNAKGLLK.. |
| **AAT68773.1_ANR_Camellia_sinensis** | YDQSVEYFKAKGILKN. |
| **ABM64802.1_ANR_Gossypium_hirsutum** | YDQTVEYLKSKGLLK.. |
| **AAN77735.1_ANR_Medicago_truncatula** | FDQTVEYLKTQGIK... |
| **ASU87432.1_ANR_Camellia_sinensis** | YDQSVEYFKAKGILKN. |
| **AEC10993.1_ANR_Camellia_sinensis** | YDQSGEYFKVKGILKN. |
| **ADZ58168.1_ANR_Camellia_sinensis** | FDHSVAYLKTKGLLQN. |
| **AHJ11240.1_ANR_Camellia_sinensis** | FDHSVAYLKTKGLLQN. |
| **NP_001267885.1_ANR_Vitis_vinifera** | YDESVEYFKAKGLLQN. |
| **ABD95362.1_ANR_Fragaria_x_ananassa** | YDQTVEYLKLKGVLQN. |
| **AEL79861.1_ANR_Malus_domestica** | YDQTVEYFKAKGLLQN. |
| **BAF56654.1_ANR_Diospyros_kaki** | YDQSVEYFKAKGILKN. |
| **ADD51354.1_ANR_Theobroma_cacao** | YDQTVEYMNAKGLLK.. |
| **CAD91909.1_ANR_Phaseolus_coccineus** | YDQTVEYLKNKGTLKN. |
| **ABM90632.1_ANR_Lotus_uliginosus** | FDQTLEYLKTKGALKN. |
| **ABC71336.1_ANR_Lotus_corniculatus** | FDQTLEYLKTKGALKN. |
| **XP_002317270.2_ANR_Populus_trichocarpa** | YDQTVEYFKANGLLN.. |
| **CAD91910.1_ANR_Gossypium_arboreum** | YDQTVEYLKSKGLLK.. |
| **AEL79859.1_ANR_Malus_domestica** | YDQTVEYFKAKGLLQK. |
| **AEL79860.1_ANR_Malus_domestica** | YDQTVEYFKAKGLLQN. |
| **AEL79861.1_ANR_Malus_domestica** | YDQTVEYFKAKGLLQN. |
| **ACY30421.1_C.BANa_Brassica_napus** | YDQMVEYFKTNRWA... |
| **ACY30422.1_C.BANb_Brassica_napus** | YDEMTKYFESKGLIKP. |
| **ACY30423.1_A.BANa_Brassica_napus** | YDQMVEHFKTNRWA... |
| **ACY30424.1_A.BANb_Brassica_napus** | YDEMTEYFESKGLIKP. |
| **ACY30425.1_C.BANa_Brassica_oleracea** | YDQMVEYFKTNRWA... |
| **ABG76842.1_ANR_Fragaria_x_ananassa** | YDQTVEYLKLKGVLQN. |
| **ABC71337.1_ANR-1_Lotus_corniculatus** | FDQTLEYLKTKGALKN. |
| **ABC71332.1_ANR1-1_Lotus_corniculatus** | FDQTLEYLKTKGALKN. |
| **ABC71333.1_ANR1-2_Lotus_corniculatus** | FDQTLEYLKTKGALKN. |
| **ABC71335.1_ANR1-4_Lotus_corniculatus** | FDQTLEYLKTKGALKN. |
| **AAZ79363.1_ANR_Malus_domestica** | YDQTVEYFKAKGLLQN. |
| **AAX12184.1_ANR_Malus_domestica** | YDQTVEYFKAKGLLQN. |
| **AAZ17408.1_ANR_Malus_domestica** | YDQTVEYFKAKGLLQK. |
| **EEE86150.1_ANR_Populus_trichocarpa** | YDQTVEYFKAKGLLN.. |
| **EEE97882.1_ANR_Populus_trichocarpa** | YDQTVEYFKANGLLN.. |
| **ABB77695.1_ANR_Pyrus_communis** | YDQTVEYFKAKGLLQN. |
| **BAD89742.1_ANR_Vitis_vinifera** | YDESVEYFKAKGLLQN. |
| **AAZ82409.1_ANR_Vitis_vinifera** | YDESVEYFKAKGLLQN. |
| ***acc*** | |
